# The disadvantage of derivation: conserved systematic flaws in primary data have repeatedly biased the phylogenetic inference of Temnospondyli (Tetrapoda, Amphibia)

**DOI:** 10.1101/2022.06.22.496729

**Authors:** Bryan M. Gee

**Affiliations:** Burke Museum and Department of Biology, University of Washington, Seattle, WA, USA

## Abstract

Phylogenetic analyses and their resultant tree topologies underlie paleobiological studies. Regardless of the type of study, the relationships of focal taxa are foundational, whether implemented in a qualitative or a quantitative framework. This reliance places a premium on the continued refinement of both phylogenetic methods and inference. Temnospondyls are a diverse clade of non-amniote (‘amphibian’) tetrapods whose phylogenetic relationships have been extensively explored due to their speciose nature, widespread occurrence in Paleozoic and Mesozoic paleoenvironments, and putative relationship to extant amphibians. Despite being studied by a diversity of workers, there is only one dataset that is widely employed to test the broad-scale relationships of Temnospondyli, that of Schoch (2013). This dataset has been reused in several high-profile studies testing the question of lissamphibian origins, and the original resultant topology has been widely adopted by taxonomic specialists and non-specialists alike. However, close examination of this matrix reveals discernible patterns of problematic codes related to non-homology, dependency, and unsubstantiated data (e.g., codes for postcranial characters for taxa with no known postcrania). These patterns, in conjunction with their prevalence, warrant a thorough survey of the entire matrix and subsequent reanalysis of its various forms to test whether previously published findings regarding the relationships of temnospondyls and the origins of lissamphibians are substantiated. A thorough reassessment of this matrix and several of its high-profile derivates revealed that the phylogeny of temnospondyls is more poorly known than depicted by the literature and that certain hypotheses of lissamphibian origins within Temnospondyli lack phylogenetic support.

## INTRODUCTION

Temnospondyls are a speciose clade of non-amniote tetrapods with a cosmopolitan geographic distribution and a temporal range of more than 200 million years (late Carboniferous to Early Cretaceous). This clade is notable for (1) longstanding hypothesized ancestry to modern amphibian (lissamphibian) clades (e.g., e.g., Moodie, 1916; Watson, 1919, 1926, 1940; Parsons and Williams, 1963; Bolt, 1969, 1977, 1991; Milner, 1988, 1993; Trueb and Cloutier, 1991; Gardner, 2001; Ruta et al., 2003; Schoch and Carroll, 2003; Carroll, 2004, 2007; Schoch and Milner, 2004; Zhang et al., 2005; Ruta and Coates, 2007; Anderson, 2008; Anderson et al., 2008; Sigurdsen and Bolt, 2009; Sigurdsen and Green, 2011; Maddin and Anderson, 2012; Maddin et al., 2012; Pardo et al., 2017; Daza et al., 2020; Schoch et al., 2020b); (2) a wide range of morphological (and presumed ecological) diversity, which exceeds that observed in lissamphibians (e.g., Hammer, 1979; Sengupta and Ghosh, 1993; Laurin and Soler-Gijón, 2001; Stayton and Ruta, 2006; Witzmann et al., 2009; Mukherjee et al., 2010, 2019; Sanchez et al., 2010a, 2010b; Fortuny et al., 2011b, 2016; Angielczyk and Ruta, 2012; Konietzko-Meier and Klein, 2013; Sanchez and Schoch, 2013; McHugh, 2014, 2015; Schoch, 2009, 2010, 2014; Marcé-Nogué et al., 2015; Lautenschlager et al., 2016; Teschner et al., 2017, 2020; Penrice and Deeming, 2018; Witzmann and Ruta, 2018; Pérez-Ben et al., 2019, 2020; Carter et al., 2021; Fig. 1); and (3) frequent occurrence in non-marine paleoenvironments in the late Paleozoic and early Mesozoic (e.g., Panchen, 1977; Cosgriff, 1984; Milner, 1987, 1994; Shubin and Sues, 1991; Sennikov, 1996; Lucas, 1998, 2010; Warren et al., 2001; Piñeiro et al., 2007a; Kear et al., 2016; Bandyopadhyay and Ray, 2020; Nonsrirach et al., 2020; Romano et al., 2020; Fig. 2). Temnospondyls are therefore of great import to paleontologists and neontologists seeking to address questions ranging from lissamphibian origins and the interrelationships of early tetrapods to Phanerozoic terrestrial ecosystem dynamics and the role of mass extinctions in tetrapod evolution (e.g., Sennikov, 1996; Shishkin, 2007; Ruta and Benton, 2008; Sidor et al., 2013; Brocklehurst et al., 2017, 2018; Button et al., 2017; Shishkin and Novikov, 2017; Dunne et al,., 2018; Pardo et al., 2019; Allen et al., 2020; Romano et al., 2020; Mancuso et al., 2021; Vigletti et al., 2021; Liu et al., 2022).

**Figure 1.**
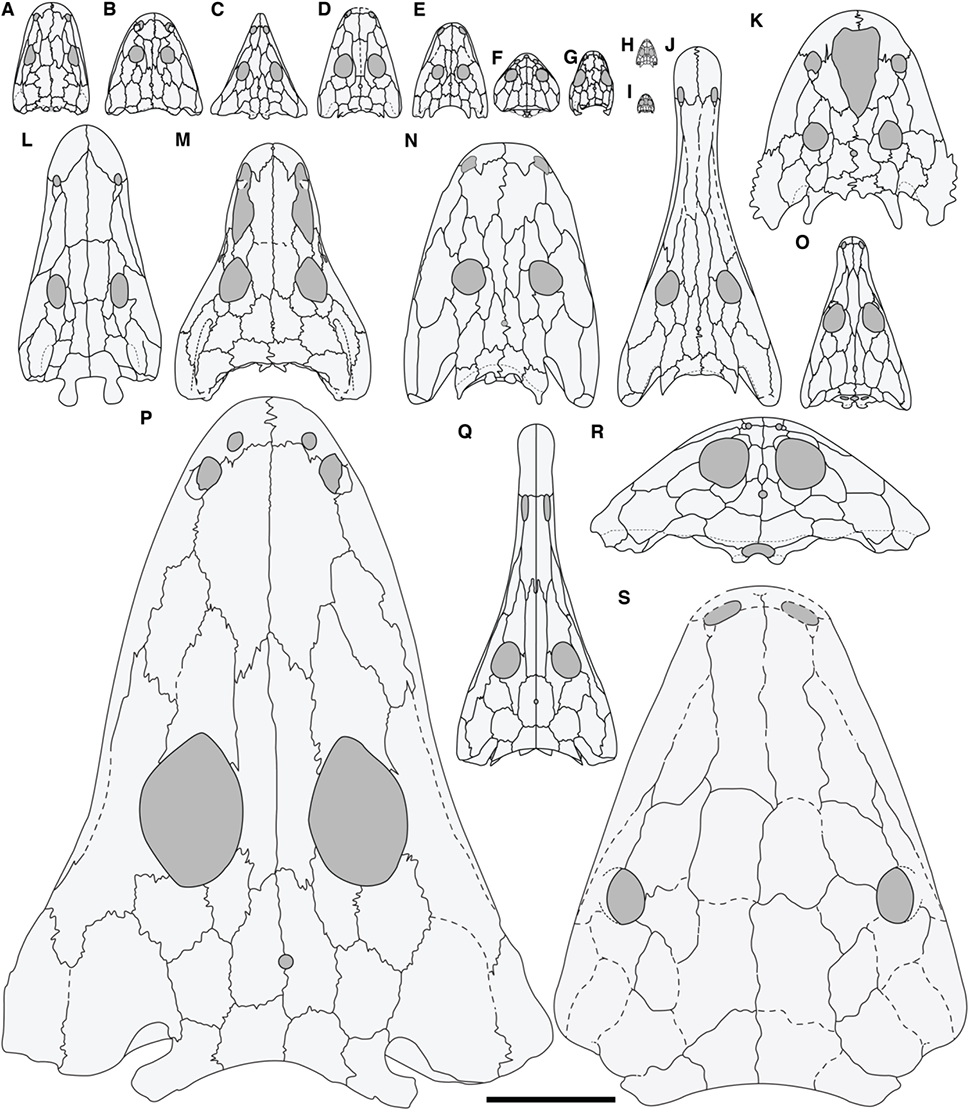
Representative temnospondyl skulls shown in dorsal view illustrate the cranial diversity and size range present in this clade. **A**, the ‘dendrerpetid’ *Dendrerpeton helogenes* (after Holmes et al., 1998); **B**, the trimerorhachid dvinosaur *Trimerorhachis insignis* (after Milner and Schoch, 2013); **C**, the rhytidosteid *Laidleria gracilis* (after Warren, 1998a); **D**, the rhinesuchid *Broomistega putterilli* (after Shishkin and Rubidge, 2000); **E**, the lydekkerinid *Lydekkerina huxleyi* (after Schoch and Milner, 2000); **F**, the dvinosaur *Acroplous vorax* (after Englehorn et al., 2008); **G**, the amphibamiform *Micropholis stowi* (after Schoch and Rubidge, 2005); **H**, the lapillopsid *Lapillopsis nana* (after Yates, 1999); **I**, the amphibamiform *Doleserpeton annectens* (after Sigurdsen and Bolt, 2010); **J**, the stereospondylomorph *Archegosaurus decheni* (after Witzmann, 2005); **K**, the zatracheid *Zatrachys serratus* (after Schoch and Milner, 2013); **L**, the edopoid *Cochleosaurus bohemicus* (after Sequeira, 2003); **M**, the trematopid dissorophoid *Acheloma cumminsi* (after Polley and Reisz, 2011); **N**, the stereospondylomorph *Sclerocephalus haeuseri* (after Schoch and Witzmann, 2009b); **O**, the trematosaur *Trematolestes hagdorni* (after Schoch, 2006); **P**, the capitosaur *Mastodonsaurus giganteus* (after Schoch, 1999); **Q**, the rhinesuchid *Australerpeton cosgriffi* (after Eltink et al., 2016); **R**, the plagiosaurid *Gerrothorax pulcherrimus* (after Schoch and Witzmann, 2012); **S**, the indeterminate temnospondyl *Saharastega moradiensis* (after Damiani et al., 2006). Silhouettes are scaled relative to each other and are based on the original scaling in the source literature (i.e. they are not scaled to the maximum known or inferred size of the given taxon); scale bar equal to 10 cm.

**Figure 2.**
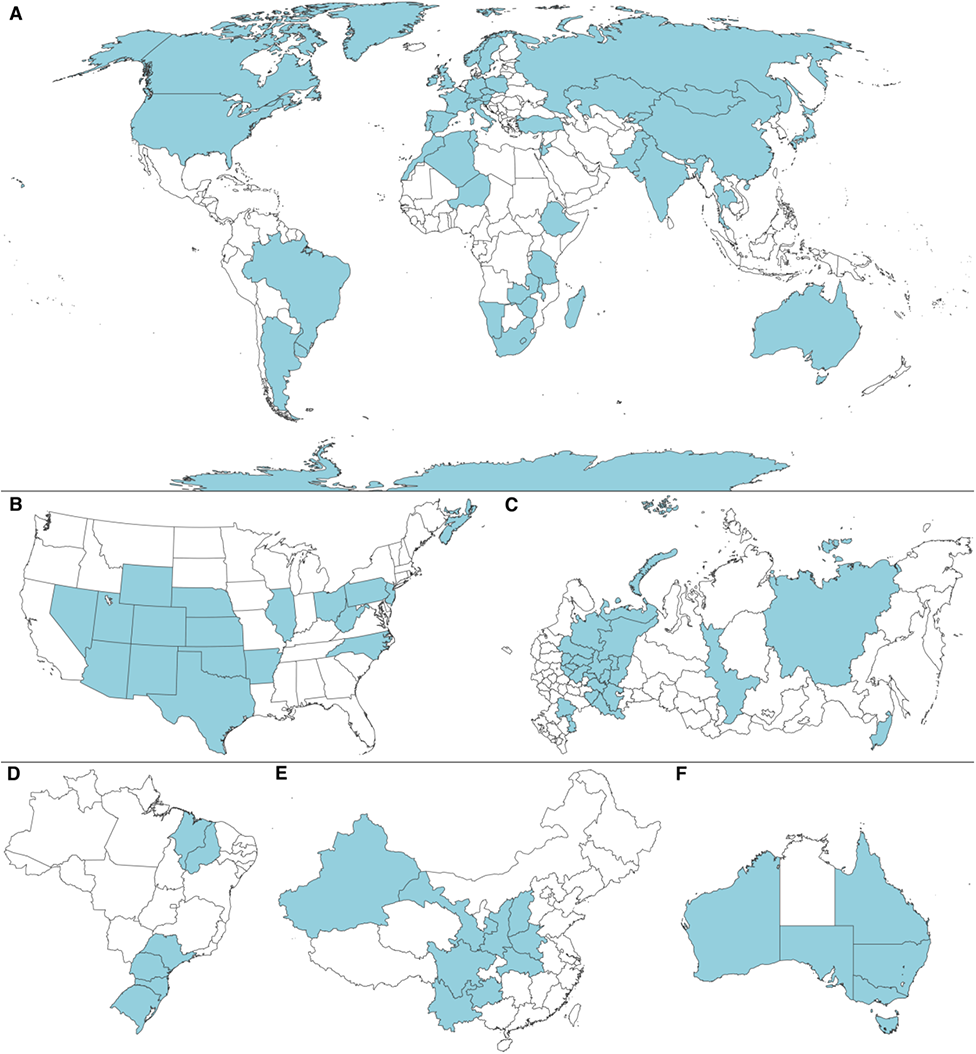
Distribution of temnospondyl body fossils. **A**, distribution on a global map; **B**, distribution within the continental United States and the Canadian Maritimes; **C**, distribution within Russia; **D**, distribution within Brazil; **E**, distribution within China; **F**, distribution within Australia. Specific focal regions represent both the largest countries by surface area and some of the most productive regions for temnospondyl body fossils. Occurrence data was sourced directly from the literature and is cited in Appendix 4. Map depictions do not represent opinions on jurisdictional claims and were sourced from FreeVectorMaps.com.

The phylogeny of temnospondyls and its various subclades has been frequently examined using computer-assisted methods (e.g., Eltink et al., 2017, 2019; Fernández-Coll et al., 2017; Marsicano et al., 2017, 2021; Marzola et al., 2017; Pacheco et al., 2017; Pardo et al., 2017; Chakravorti and Sengupta, 2018; Gee et al., 2018, 2021; Liu, 2018; Schoch, 2018a, 2018b, 2018c, 2019, 2021a, 2021b; Schoch and Witzmann, 2018; Atkins et al., 2019; Buffa et al., 2019; Schoch and Voigt, 2019; Dilkes, 2020; Gee, 2020b, 2021; Maisch, 2020; Schoch et al., 2020a; Werneburg et al., 2020; Schoch and Milner, 2021; Schoch and Sobral, 2021; Arbez et al., 2022; Schoch & Sues, 2022; Werneburg et al., 2022). The large body of work underscores the import of phylogenetic analyses as the means of inferring evolutionary narratives and informing taxonomic frameworks. However, the large number of studies and diversity of working groups belies the small number of independent matrices currently used to test temnospondyl relationships. The vast majority of studies published within the past decade used pre-existing matrices that were largely replicated with little to no modification beyond the addition or deletion of taxa (Fig. 3).

**Figure 3.**
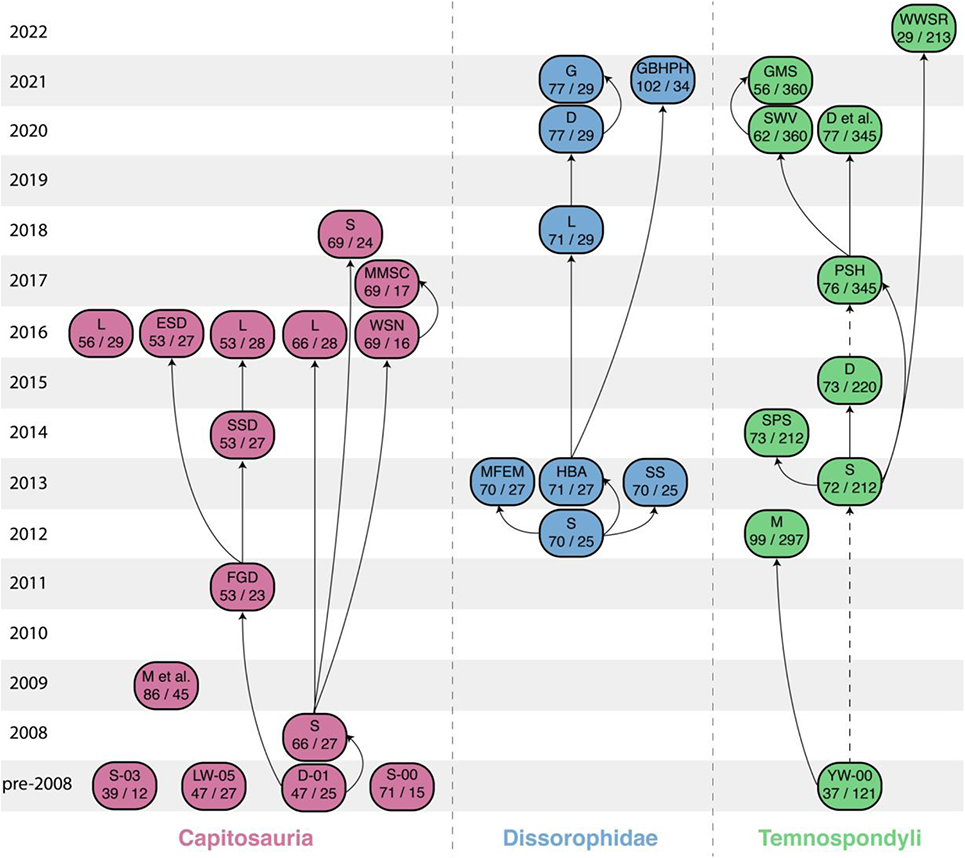
Genealogy of phylogenetic matrices used to analyze select temnospondyl clades and Temnospondyli, showing conservation of matrices across studies by different workers. Temnospondyl-wide analyses are represented by green boxes; Capitosauria (red) and Dissorophidae (blue) were selected because these are relatively well-studied from a phylogenetic perspective. Initials represent those of the authors’ surnames; studies prior to 2008 are appended with the year of publication. Depicted studies are those that focus on the intrarelationships of the focal clade and exclude studies testing the position of nominal taxa in broad-scale analyses (e.g., Schoch et al., 2007, for *Sclerothorax*). Supertrees are not included (e.g., Ruta et al., 2007); conference abstracts are also omitted (e.g., So et al., 2018; Kligman et al., 2021). Arrows depict express derivation; dashed arrows depict inferred derivation. Abbreviations for capitosaur analyses: D, Dilkes (2015, 2020); D-01, Damiani (2001); D et al., Daza et al. (2020); ESD, Eltink et al. (2017); FGD, Fortuny et al. (2011a); G, Gee (2021); GMS, Gee et al. (2021); GBHPH, Gee et al. (2021); HBA, Holmes et al. (2013); L, (Liu, 2016, 2018); LW-05 (Liu and Wang, 2005); M, McHugh (2012); M et al. (Maganuco et al., 2009); MFEM, Maddin et al. (2013); MMSC, Marzola et al. (2017); PSH, Pardo et al. (2017); S-00, Schoch (2000); S-03, Steyer (2003); S, Schoch (2008b, 2012, 2013 2018c); SS (Schoch and Sues, 2013); SPS, Strapasson et al. (2014); SSD (Sidor et al., 2014); SWV, Schoch et al. (2020); WSN Witzmann et al. (2016); WWSR, Werneburg et al. (2022); YW-00, Yates and Warren (2000).

The underlying non-independence between analyses is exemplified by analyses of phylogenetic relationships of Temnospondyli in its entirety (Fig. 3). There is only one recently published matrix that samples the entire clade—Schoch (2013)—others are either unpublished dissertation chapters (e.g., Pawley, 2006; McHugh, 2012) or are skewed toward a subset of temnospondyls (e.g., Ruta & Bolt, 2006; Maganuco et al., 2009, 2014). The topology recovered by Schoch (2013) has been extensively employed as a backbone for phylogenetic comparative method studies (e.g., Anderson et al., 2013; Organ et al., 2016; MacIver et al., 2017; Ruta et al., 2018; Tarailo, 2018; Witzmann and Ruta, 2018; Pérez-Ben et al., 2019; Dickson et al., 2021). It has also been frequently employed to map character distributions (e.g., Witzmann, 2013, 2016; Konietzko-Meier et al., 2014, 2020; McHugh, 2015; Witzmann and Voigt, 2015; Danto et al., 2016, 2017, 2019; Gee et al., 2017; Witzmann and Werneburg, 2017; Schoch, 2019). Finally, the matrix itself has been expanded in several stepwise studies to address lissamphibian origins, under a presumption of origin(s) within Temnospondyli (Pardo et al., 2017; So et al., 2018, 2021; Daza et al., 2020; Schoch et al., 2020b; Kligman et al., 2021). In both these derivates, which were adopted by other workers, and other derivates that were not adopted by a subsequent study (e.g., Strapasson et al., 2014; Dilkes, 2015; Gee et al., 2021; Werneburg et al., 2022), the taxon and character sampling have been variably modified, but virtually all of the original codes of Schoch (2013), and subsequently those introduced for characters added by Pardo et al. (2017) and Schoch et al. (2020b), have not been modified. There is therefore not only a connectivity in the source of temnospondyl-wide matrices but also a broad consistency in the primary dataset (individual codes).

The non-independence of propagated datasets can be double-edged. When workers continue to propagate a matrix without substantive modification beyond revisions associated with publication of novel data, this results in an implicit perception that the integrity of the primary data is relatively high and that workers are converging on a consensus of relationships. Ideally, this consistency would be reflected in a consistent topology, which is largely true in derivates of Schoch (2013) that tend to differ only in the relationships of lissamphibians. However, if the integrity of the primary data is found to be compromised, this means that there has been systemic propagation of errors, and that all subsequent derivate matrices or studies that utilized any resultant topologies may have been compromised. As part of ongoing studies (Kligman et al., 2021; Gee et al., in prep) that intended to narrowly expand one of the most recent derivates (Schoch et al., 2020b) to address the relationships of select taxa, a large number of questionable codes were identified in those studies’ focal taxa (e.g., *Lapillopsis*). These included codes that directly contradicted recent descriptive work; codes for skeletal regions not known for a given taxon; and codes for characters that should be treated as inapplicable for a given taxon. These observations necessitated a more thorough systematic review of the entire matrix, which identified broader patterns, both within characters and within taxa, that necessitated thousands of coding modifications (Kligman et al., 2021). Given the scope of the modifications, I felt that a dedicated study was needed to thoroughly document and justify coding revisions; to reanalyze previous matrices to determine whether their original findings were compromised by errors in the primary data; and to discuss the implications of the new findings with respect to temnospondyl phylogeny and lissamphibian origins.

## MATERIALS and METHODS

### Source Matrices and History

This study reanalyzes previously published matrices that trace their roots back to the temnospondyl matrix of Schoch (2013), as visualized in Figure 3 and as detailed in Table 1. Per the publisher’s website, Schoch’s manuscript was accepted in November 2011 but was only published online in January 2013, meaning that the content was finalized by the end of 2011 and is now more than a decade old. This gap between acceptance and publication explains why revisions of sampled taxa that were published in 2012 or 2013 (e.g., *Branchiosaurus salamandroides*; Werneburg, 2012; *Gerrothorax pulcherrimus*; Schoch and Witzmann, 2012; *Trimerorhachis insignis*; Milner and Schoch, 2013) were not cited by Schoch (2013).

**Table 1.** Summary details of the family of matrices deriving from Schoch (2013). In the original analysis of Schoch (2013); one multistate character was explicitly stated to be ordered, another was explicitly stated to be unordered, and the remaining 19 were not clarified. Pardo et al. (2017) was stated to be a direct derivate from Schoch (2013), but it incorporated specific changes made by Dilkes (2015), without attribution, including several made from Dilkes’ personal observations; it therefore is classified as a derivate of that study as well. In the same vein, while Pardo et al. made no express mention of coding modifications or lack thereof, comparison with previous derivates led to the identification of many modifications (see Appendix 3.2). For the in-group taxa, the number before the slash indicates the number of temnospondyl OTUs, and the number after the slash indicates the number of lissamphibian OTUs

| Study | In-group | Characters | Ordering | Coding modifications ? | Program | Derived from |
| --- | --- | --- | --- | --- | --- | --- |
| Schoch (2013) | 70 (61)* / 0 | 212 | Not stated | — | PAUP* 3.1; TNT 1.0 | — |
| Strapasson et al. (2014) | 71 / 0 | 212 | Some | No | TNT 1.1 | Schoch (2013) |
| Dilkes (2015) | 71 / 0 | 220 | Some | Yes | PAUP* 4.0a10 | Schoch (2013) |
| Pardo et al. (2017) | 63 / 11 | 345 | No | Not stated | PAUP* 4.0b159; MrBayes 3.1.2 | Schoch (2013) / Dilkes (2015) |
| Daza et al. (2020) | 63 / 12 | 345 | Not stated | No | TNT 1.5 | Pardo et al. (2017) |
| Schoch et al. (2020) | 44 / 13 | 360 | Not stated | Yes | TNT 1.0 | Pardo et al. (2017) |
| Gee et al. (2021) | 51 / 0 | 360 | No | Yes | PAUP* 4.0b169 | Schoch et al. (2020) |
| Werneburg et al. (2022) | 27 / 0 | 213 | Some | Yes | PAUP* 3.1 | Schoch (2013) |

Other relevant aspects of this matrix’s history are as follows. Firstly, Schoch (2013) analyzed the matrix in two different programs (PAUP*, TNT), each with a full (72 operational taxonomic units [OTUs], 70 temnospondyls) and a reduced (63 OTUs, 62 temnospondyls) taxon sample. He also mentioned ‘variant analyses’ with 66 OTUs (p. 693 therein), but it is unclear how many of these analyses were performed or which taxa were excluded. Only the results of one analysis were figured and described in detail (i.e. number and length of MPTs, cladogram topology, support metrics): the reduced taxon sample when analyzed in PAUP*. For reasons unstated, Schoch opted to “prefer” this topology over the equivalent TNT analysis, which recovered a different number and length of MPTs and a different strict consensus. The preference for the reduced taxon sample is more apparent (wildcards included in the full sample leading to loss of resolution).

Because the PAUP* topology of the reduced taxon sample is the only one figured and discussed in detail, this topology is the one widely employed by other workers, and it remains widely employed as a backbone. However, some workers have cast doubt on whether one of the reduced taxon sample analyses did not recover any MPTs (i.e. only suboptimal trees were found; Marjanović and Laurin, 2019). As the PAUP* analysis recovered MPTs of a longer length than the equivalent TNT analysis (633 steps vs. 628 steps), it can be conjectured that the PAUP* analysis did not find any MPTs, which would be problematic as this topology is the one frequently employed by other workers.

This study re-analyzes two versions of this matrix: the original of Schoch (2013) and the second-order derivate of Pardo et al. (2017). The original is re-analyzed because it serves as the foundation for many other studies and is the most widely used topology for studies comparative method studies that require a phylogenetic backbone. Disparity in taxon sampling and the addition of lissamphibians in some derivates has resulted in more instability within Temnospondyli in derivates, which is likely one reason that the original reported topology remains the most widely used topology for various purposes. The derivate of Pardo et al. (2017) is re-analyzed because this is the only derivate to make substantial taxon and character additions, with 13 new OTUs and 133 new characters. Other derivates have made comparatively few modifications (Strapasson et al., 2014; Dilkes, 2015; Daza et al., 2020; Schoch et al., 2020b; Gee et al., 2021). The expanded taxon sample of Pardo et al. includes lissamphibians, and their analysis produced a novel hypothesis of a diphyletic origin of Lissamphibia from within Temnospondyli. This hypothesis was tested using both Bayesian and parsimony methods, the only time that likelihood methods were employed on this family of matrices. This hypothesis has generated substantial controversy due to the high-profile nature of the topic, and to date, has not been supported by independent studies (e.g., Marjanović & Laurin, 2019; Daza et al., 2020; Santos et al., 2020; Schoch et al., 2020b; Kligman et al., 2021).

### Objectives and Motivations

I consider it important to establish the motivations for this study, as I recognize that the reanalysis of previous studies can be a sensitive topic, and I want to avoid any misinterpretations or unsubstantiated assumptions about this work. This study is not motivated by any personal or professional animus towards the author(s) of any previous study that have employed this matrix. This study is not a prelude to a broader series of papers targeting matrices of either a specific worker or a specific taxonomic group. While some workers may be inclined to extend the results of this study beyond the confines of this specific family of matrices to other matrices by the same author(s), I do not endorse this, either implicitly or explicitly. The implications of this study should be confined to the datasets analyzed here. Finally, this study is not motivated by a desire to ‘cherry-pick’ (proving that at least one possible error exists) or to ‘prove’ the truism that phylogenetic matrices inherently become outdated as new data emerge. The magnitude of necessary revisions will further bolster this final point.

This study represents, in part, a collation of work done as part of two ongoing projects (Gee et al., in prep; Kligman et al., 2021). Those studies identified scoring discrepancies in their respective focal taxa (e.g., the coding of *Doleserpeton* as lacking symphyseal dentition, contra Sigurdsen and Bolt, 2010; and the absence of coding for any lacrimal characters in, despite documentation of this element by Yates, 1999). In isolation, these issues could have been considered to fall within a reasonable range of inadvertent error, but I observed that many of these problematic codes were part of a broader pattern. For example, *Doleserpeton, Lapillopsis*, and 38 other taxa that were coded by Schoch (2013) as lacking osteoderms (original character state 209–0) were coded as having a ‘simple set of osteoderms arranged in one layer if present’ (original 210–0) and ‘narrow median osteoderms’ (original 211–0). Osteoderms cannot be arranged in one layer or be narrow if they are absent. Such patterns were repeatedly identified across the matrix, and it became apparent that vetting and modification of codes could not be restricted to focal taxa, as this would introduce bias to the dataset.

It may be questioned why a separate study is warranted when many of the scoring modifications are implemented in other studies (Gee et al., in prep; Kligman et al., 2021). I have three primary motivations for a separate study. The first is the treatment of previous phylogenetic studies when newer results are available (either independently or as derivates). By convention, a newer study does not inherently invalidate an older one if there are discrepancies – due to differences such as character and taxon sampling or to methodological nuances, both studies can have equally accurate results insofar as they faithfully represent the analysis of a given dataset. For example, Daza et al. (2020) used a minimally modified derivate of Pardo et al.’s (2017) matrix (adding only one OTU) and did not recover the diphyly hypothesis, but the former utilized implied-weights parsimony versus Pardo et al.’s equal-weights parsimony. The former’s novel hypothesis is not inherently invalidated merely by being post-dated. However, if a matrix does not accurately reflect the primary data (within a reasonable margin of human error), then its results are inherently flawed, regardless of whether this arises from high error rate, new data, or any combination in between, and the results of such an analysis can be invalidated on those grounds. Such a matrix and its resultant topologies cannot be propagated in good faith once it is known that there are systemic issues or shortcomings, and at minimum, the encapsulating study cannot stand as an equally weighted line of evidence in any debate.

The second motivation is that because this matrix has been repeatedly propagated with minimal modification by previous workers, any errors in the original are almost certainly present in the derivates, and whether these errors have also affected derivates must be assessed. It can therefore be reasonably hypothesized that implementing necessary coding changes would exert a measurable influence on the topology based on their sheer number, and as a result, that previous results are flawed. However, this cannot be assumed *a priori.* Because this family of matrices and their previously published topologies are likely to be propagated and utilized in the debate over lissamphibian origins, as well as in studies for which a topology of Temnospondyli is needed, a reanalysis of at least one derivate is warranted.

The final motivation is that phylogenetic revision, including assessments of the relationships across Temnospondyli, was not the focus of the motivating studies, and there is therefore little discussion of the implications of widespread unsubstantiated character codings (which is then relegated to supplemental information). Methodological research has identified shortcomings of extensive use of supplementary materials, such as inconsistent formatting and access; higher probability of not being thoroughly reviewed during peer review; lower readership compared to the primary article; separation from the main text; and undercounting of citations (e.g., Evangelou et al., 2005; Maunsell, 2010; Borowski, 2011; Kenyon and Sprague, 2014, 2016; Pop and Salzberg, 2015; Moore and Beckerman, 2016; Shutler and Murray, 2016). Of these, I am most concerned about the prospect of reduced readership, which, when paired with the lack of dedicated discussion to the implications of such widespread recoding, greatly limits the visibility and utility of these revisions. Therefore, I felt that a dedicated study that allows for a full discussion of these issues, their redress and implications, and future directions for temnospondyl phylogenetics would be warranted.

### Assessment of Previous Codes

As previously mentioned, reexamination of previous codes was first conducted as part of two other studies, Kligman et al. (2021), which used the matrix of Schoch et al. (2020b); and Gee et al. (in prep), which combined the taxon sample of Pardo et al. (2017) with the character sample of Schoch et al. (2020b). Between these two, the majority of previously sampled taxa have been reexamined for all previously coded characters. However, nine of the taxa originally coded by Schoch (2013) were not carried forward into any Pardo et al.’s derivate and had to be reexamined for the 212 characters used by Schoch as part of this study: the tupilakosaurid dvinosaurs *Thabanchuia oomie* and *Tupilakosaurus wetlugensis*; the micromelerpetid dissorophoid *Branchierpeton amblystomum*; the branchiosaurid dissorophoid *Branchiosaurus salamandroides*; and the capitosaurs *Watsonisuchus gunganji*, *Wetlugasaurus angustifrons, Eryosuchus garjainovi*, and *Eocyclotosaurus wellesi*.

The basic principles used to assess codes are outlined here, with additional comments and detailed examples given in Appendix 2.1. Examination of codes utilized the highest evidentiary standard for changing codes – the literature had to expressly contradict the original code to warrant being changed. This included previously coded cells where a different condition is documented in the literature, previously coded cells where the literature does not indicate that the requisite skeletal region is known for that given taxon, and previously uncoded cells where the literature documents the requisite region to a sufficient degree to code. Although I personally prefer to avoid the use of reconstructions when coding matrices (Gee, 2020b, 2021), it is apparent from previous codes that reconstructions were regularly utilized by previous workers, and I followed that approach here for consistency. The only instance in which I did not utilize reconstructions is when it was entirely conjectural. For example, taxa represented only by one specimen, either in totality or for a given skeletal region (e.g., *Gerobatrachus*, *Siderops*), must be reconstructed in part based strictly on conjecture, even if it is informed by putative close relatives. In particular, older reconstructions could not have accounted for the possibility that some subjective decisions might prove influential in future phylogenetic analyses.

### Analytical Parameters

All analyses were run on a personal computer (MacBook Pro, mid-2015 model; 16 GB of RAM). This is not a dedicated analytical computer and therefore computation times may have been increased when running multiple processes in addition to the analyses presented here. The most recent versions of PAUP* (4.0a169) and TNT (1.5) were employed here (Goloboff and Catalano, 2016; Swofford, 2001). I mirrored the original parameters wherever they were specified.

One outstanding question is character ordering. Schoch (2013) made no clear statement about the treatment of the 21 multistate characters. Only one (character 67, tabular [ventral crest]) was stated to be ordered in his character list; one (character 158, presacral count) was stated to be unordered; and there was no comment on the remaining 19. The one stated to be unordered is perplexing because the character states form an unambiguous morphocline as a count-based character. Strapasson et al. (2014) and Werneburg et al. (2022), both first-order derivates, only ordered character 67, indicating they inferred all characters to be unordered unless expressly stated as such. This was followed here, as the default treatment in PAUP* and TNT is unordered. Pardo et al. (2017) expressly stated that all characters were unordered.

If the objective of the reanalyses was solely to recover the topology that the original workers ‘should’ have recovered if unsubstantiated cells were correctly coded, then characters should be entirely unordered for most of the reanalyses except perhaps that of Schoch (2013). However, if a secondary objective is to infer a more accurate topology, then character ordering must be considered because it has biological relevance. While no ordering is the algorithmic default, it is not the biological default and thus not neutral. An unordered multistate character implies a scenario in which transformation between states is equally likely, whereas some character states are configured in a way that they form a logical morphocline (e.g., presacral vertebral count). Empirical and simulated studies have shown that reasonable character ordering produces a more accurate solution (e.g., Fröbisch and Schoch, 2009a; Grand et al., 2013; Rineau et al., 2015, 2018). Therefore, analyses were run with no ordering and select ordering, but because this is not fundamentally a methods paper, I did not explore other parameters such as implied weighting or the use of multiple operational outgroups.

The means of computing support metrics was also sometimes lacking in detail for previous analyses. For example, Schoch (2013) and Pardo et al. (2017) gave no details as to the number of replicates or the type of search employed during bootstrapping. Given the size of the matrix and the lengthy time to perform heuristic searches in PAUP*, performing any statistically meaningful number of bootstrap replicates (here considered to be ≥ 1,000 replicates) would be prohibitively costly and infeasible without a dedicated instrument. Therefore, I performed 100,000 “fast” stepwise-addition replicates in PAUP*. This option was chosen because the computation time of the original searches in PAUP* typically exceeded six hours, and in one instance, exceeded 24 hours; bootstrapping with the same parameters as the topological search (e.g., 10,000 random addition sequence replicates) or even with simple addition sequence was simply not feasible. TNT has faster processing time, and it is feasible to perform 20,000 replicates using the ‘traditional’ search option. Bootstrap values reported from TNT are absolute frequencies to be most comparable with those reported by PAUP*.

Bremer decay index was calculated in both by rerunning the heuristic search, retaining trees of progressively longer length. In PAUP*, strict consensus trees of each stepwise iteration have to be manually compared, whereas in TNT, the ‘*bsupport*’ command will compile this index from the series of stepwise iterations. In TNT, I progressively searched for trees of one step longer with an increased holding capacity of 100,000 trees with each search (e.g., the search for trees of a length of five steps longer had a capacity of 500,000 trees). In PAUP*, some of the initial searches for MPTs were rather costly with respect to time (> 24 hours), even though they tended to find at least one MPT within the first 500 replicates. Preliminary Bremer searches also indicated that all suboptimal trees were recovered within the first 500 replicates. Because searches for progressively longer trees take a progressively longer time, it was not considered reasonable to run the analysis with 10,000 replicates for an additional five steps given the limitations of my personal computer (certain reanalyses would have required over a week to complete one Bremer sequence). Therefore, for the calculation of this index in PAUP*, the original search was rerun with 1,000 random addition sequence replicates, holding one tree per step, and with all other parameters unchanged. In all instances, these limited searches still recovered the same sets of MPTs, so these parameters were then used to search for the progressively longer trees. The holding capacity was increased by 2,000,000 trees with each step, reflecting that PAUP* tends to recover more trees than the equivalent search in TNT due to the default branch collapsing rule and that for the matrices analyzed here, the necessary RAM exceeded that available between 10 and 12 million trees. For this reason, certain reanalyses were not able to calculate out to a Bremer support value of five.

### Details of Reanalyses

#### Reanalysis of Schoch (2013)

Schoch coded 72 taxa (70 temnospondyls) for 212 characters. As previously mentioned, the only figured and discussed topology is from his PAUP* analysis with only 63 taxa (62 temnospondyls); other analyses were briefly discussed with respect to wildcard taxa. The restricted dataset was analyzed with heuristic searches in both PAUP* and TNT, although methodological details were lacking (e.g., which type of heuristic search was used in PAUP*). Per Schoch (p. 679 therein):

> “The data were analysed with two different software packages, TNT 1.0 and PAUP 3.1/MacClade 3.0. The same datasets were run in both programs, resulting in generally very similar tree topologies. The search was always conducted in the heuristic mode, employing branch swapping (TBR) options. In TNT, new technology search options were also used in addition to traditional search options. All analyses were run in the ACCTRAN mode.”

Because most pertinent details were not specified, I ran a check to see whether I could recover the original results (1,080 MPTs with length 638 steps for the restricted taxon sample) using the original matrix with the specified parameters and all other settings left as defaults. I first employed a heuristic search in PAUP* with 10,000 random addition sequence replicates, holding 10 trees per step and with the seed taxon = 0 (*Proterogyrinus scheelei*), with tree-bisection- and-reconnection (TBR) with a reconnection limit = 8. Although most of these were not specified by Schoch, they are standard based on other studies (e.g., they mirror those of Pardo et al., 2017). All other parameters were defaults of the current version of PAUP* (4.0a build 169 for Macintosh), including: branches collapsed if maximum length = 0 (‘rule 3’); multiple states interpreted as polymorphism; and treatment of “gap” states as missing data. The maximum number of trees was set to continually increase such that it did not truncate the analysis.

##### *Proterogyrinus scheelei* was the operational outgroup

The reanalysis recovered substantially more MPTs that are ten steps shorter than the original results (9,608 MPTs with length 628 steps; CI = 0.371; RI = 0.803; Figs. 4–5). MPTs were distributed across six disparately sized tree islands. The result was the same regardless of whether character 67 of the matrix was ordered (see previous comments on uncertainty regarding the original character ordering). The differing number of MPTs indicates that the original tree length reported by Schoch (2013) was not a typographic error in the MPT length. The treatment of multistate cells (which affects MPT length but not count) is irrelevant because there were no multistate cells in the original matrix. Reanalysis in TNT also recovered many more MPTs of a shorter length than originally reported (1,456 MPTs with length of 628 steps; CI = 0.371; RI = 0.803; Supp. Figs. 1–2); this MPT length is five steps shorter than the MPTs reported from TNT by Schoch but is the same as that recovered by my PAUP* renalysis. These results were also unaffected by the ordering of character 67. The smaller number of MPTs in TNT is due to the different default branch-collapsing rules; when PAUP* is set to collapse branches with a minimum length = 0 (‘rule 1’ and default in TNT), it also recovered 1,456 MPTs. Similarly, when TNT was set to collapse branches with a maximum length = 0 (‘rule 3’ and default in PAUP*), it recovered 9,608 MPTs (1,308 of which were only recovered after the second round of branch swapping). This demonstrates that the discrepancy in MPT length between these programs as reported by Schoch (2013) were not the result of different default settings.

**Figure 4.**
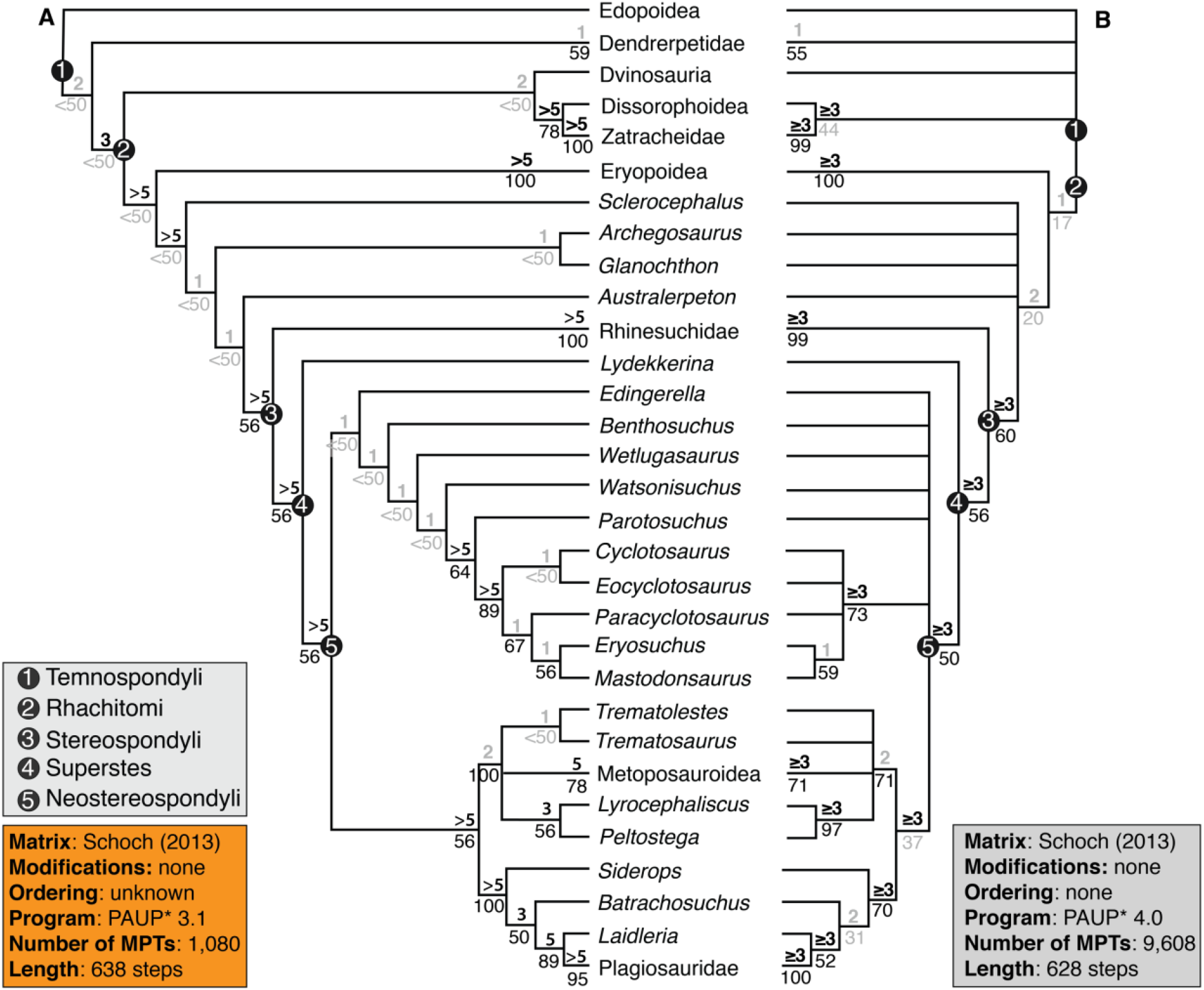
Comparison of the strict consensus topology from Schoch’s (2013) analysis using PAUP* 3.1(1,080 MPTs, length = 638 steps) with the strict consensus topology recovered from reanalysis of Schoch’s original matrix using PAUP* 4.0a169 (9,608 MPTs, length = 628 steps) with all multistate characters unordered and the restricted taxon sample. **A**, original topology figured and described by Schoch (2013:figs. 3–5); **B**, newly recovered topology from the reanalysis. The latter consensus is the same regardless of whether character 67 is ordered or unordered. Values above lines represent Bremer decay index; values below lines represent bootstrap support. Values in gray represent those below the thresholds for strong support (bootstrap ≥ 50%; Bremer index ≥ 3). Collapsed nodes representing only two species (Dendrerpetidae [*Balanerpeton*, *Dendrerpeton*], Eryopoidea [*Eryops*, *Onchiodon*], Metoposauroidea [*Callistomordax*, *Metoposaurus*], Plagiosauridae [*Gerrothorax*, *Plagiosuchus*], Rhinesuchidae [*Rhineceps*, *Uranocentrodon*], Zatracheidae [*Acanthostomatops*, *Zatrachys*]) have their support values listed at the tips and are condensed for visual clarity. Relationships of more speciose nodes (Dissorophoidea, Dvinosauria, Edopoidea) are presented in Figure 5.

**Figure 5.**
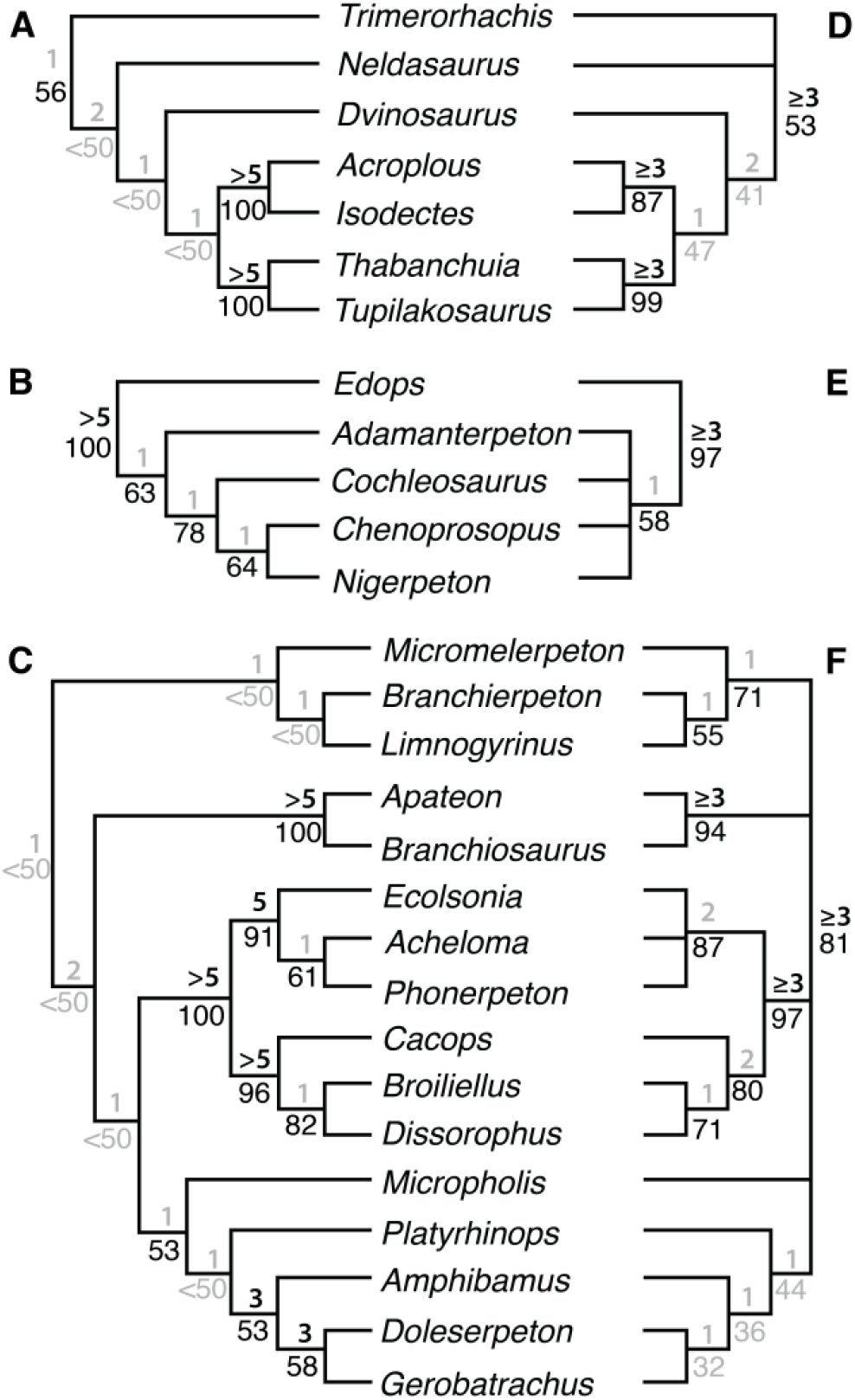
Specific relationships of Dissorophoidea, Dvinosauria, and Edopoidea recovered from reanalysis of the original matrix of Schoch (2013) in PAUP* 4.0a169 (9,608 MPTs, length = 628 steps) with the restricted taxon sample. **A**, original topology of Dvinosauria as described but not figured by Schoch (2013:682); **B**, original topology of Edopoidea; **C**, original topology of Dissorophoidea; **D**, newly recovered topology of Dvinosauria; **E**, newly recovered topology of Edopoidea; **F**, newly recovered topology of Dissorophoidea. The consensus of Schoch (2013) is the same regardless of whether character 67 is ordered or unordered. Values in gray represent those below the thresholds for strong support (bootstrap ≥ 50%; Bremer index ≥ 3).

Further demonstrating the discrepancy is not one related to a typographic error in MPT number or length, the strict consensus topologies also differ. The topologies differed between programs in Schoch (2013) but are identical in my reanalysis. The new topology still recovered many of the major clades (e.g., Edopoidea, Dvinosauria, Dissorophoidea, Stereospondyli), and no taxon was recovered in a drastically different position, but there is substantial loss of resolution, including a major basal polytomy of ‘Dendrepretidae,’ Dvinosauria, Edopoidea, and Dissorophoidea + Zatracheidae (Figs. 4–5; Supp. Figs. 1–8). The use of a majority-rule consensus tree (not shown) neither recovers the same topology as the strict consensus reported by Schoch (2013) nor a similar degree of resolution. This is another line of evidence that the original study recovered legitimately different sets of putative MPTs than I did here. Bootstrap support is lower for nearly all nodes in the reanalysis, although there is a wide range in the difference between overlapping nodes, and a few nodes increased in bootstrap frequency. Given the uncertainty over Schoch’s bootstrapping methods, it is plausible that if the original heuristic search failed to find any MPTs, then bootstrapping with the same heuristic search parameters would also be compromised.

**Figure 6.**
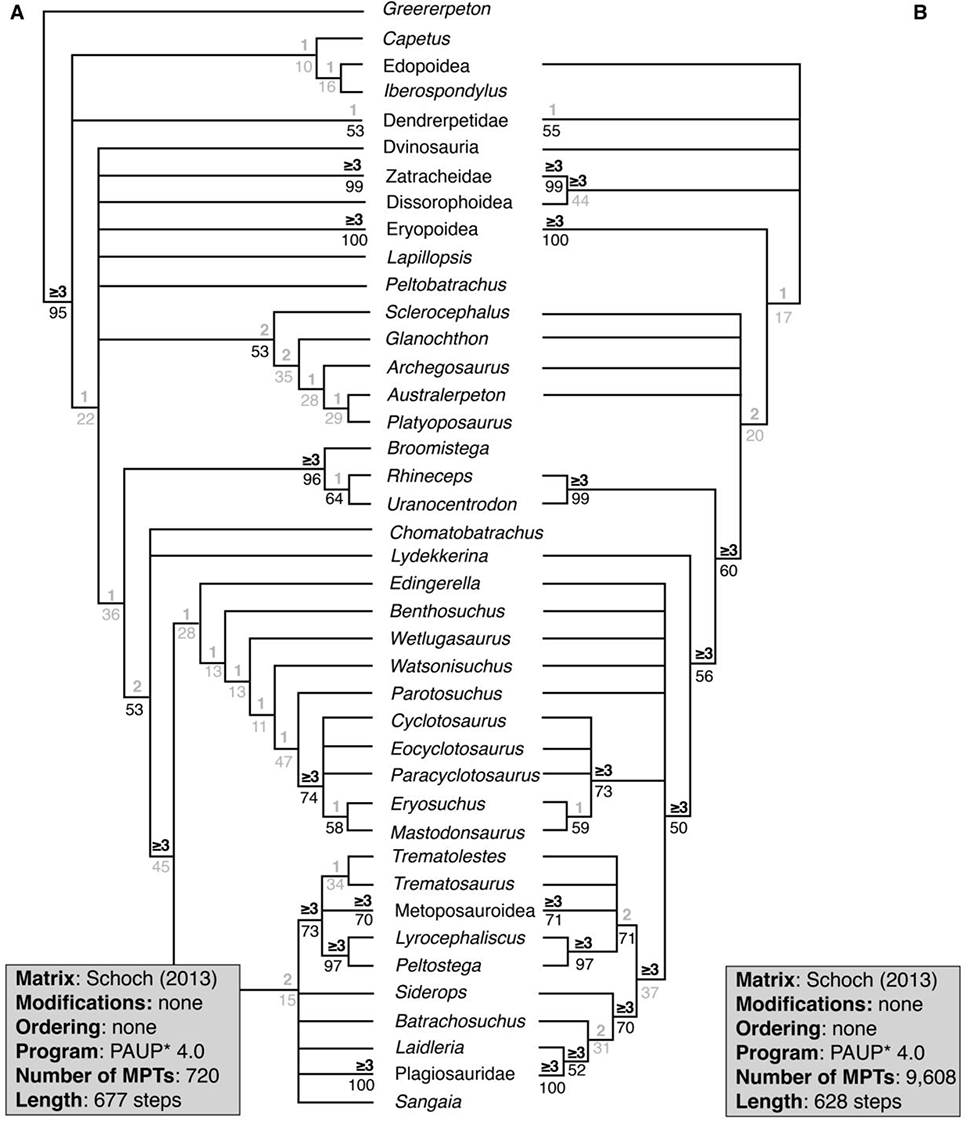
Comparison of the strict consensus topologies recovered from reanalysis of the original Schoch (2013) matrix using PAUP* 4.0a169 (720 MPTs, length = 677 steps), with differing taxon samples and only character 67 ordered. **A**, topology of the full taxon sample; **B**, topology of the restricted taxon sample (reproduced from Fig. 4B). Values above lines represent Bremer decay index; values below lines represent bootstrap support. Values in gray represent those below the thresholds for strong support (bootstrap ≥ 50%; Bremer index ≥ 3). Collapsed nodes representing only two species (Dendrerpetidae [*Balanerpeton*, *Dendrerpeton*], Eryopoidea [*Eryops*, *Onchiodon*], Metoposauroidea [*Callistomordax*, *Metoposaurus*], Plagiosauridae [*Gerrothorax*, *Plagiosuchus*], Rhinesuchidae [*Rhineceps*, *Uranocentrodon*], Zatracheidae [*Acanthostomatops*, *Zatrachys*]) have their support values listed at the tips and are condensed for visual clarity. Relationships of more speciose nodes (Dissorophoidea, Dvinosauria, Edopoidea) are presented in Figure 7.

**Figure 7.**
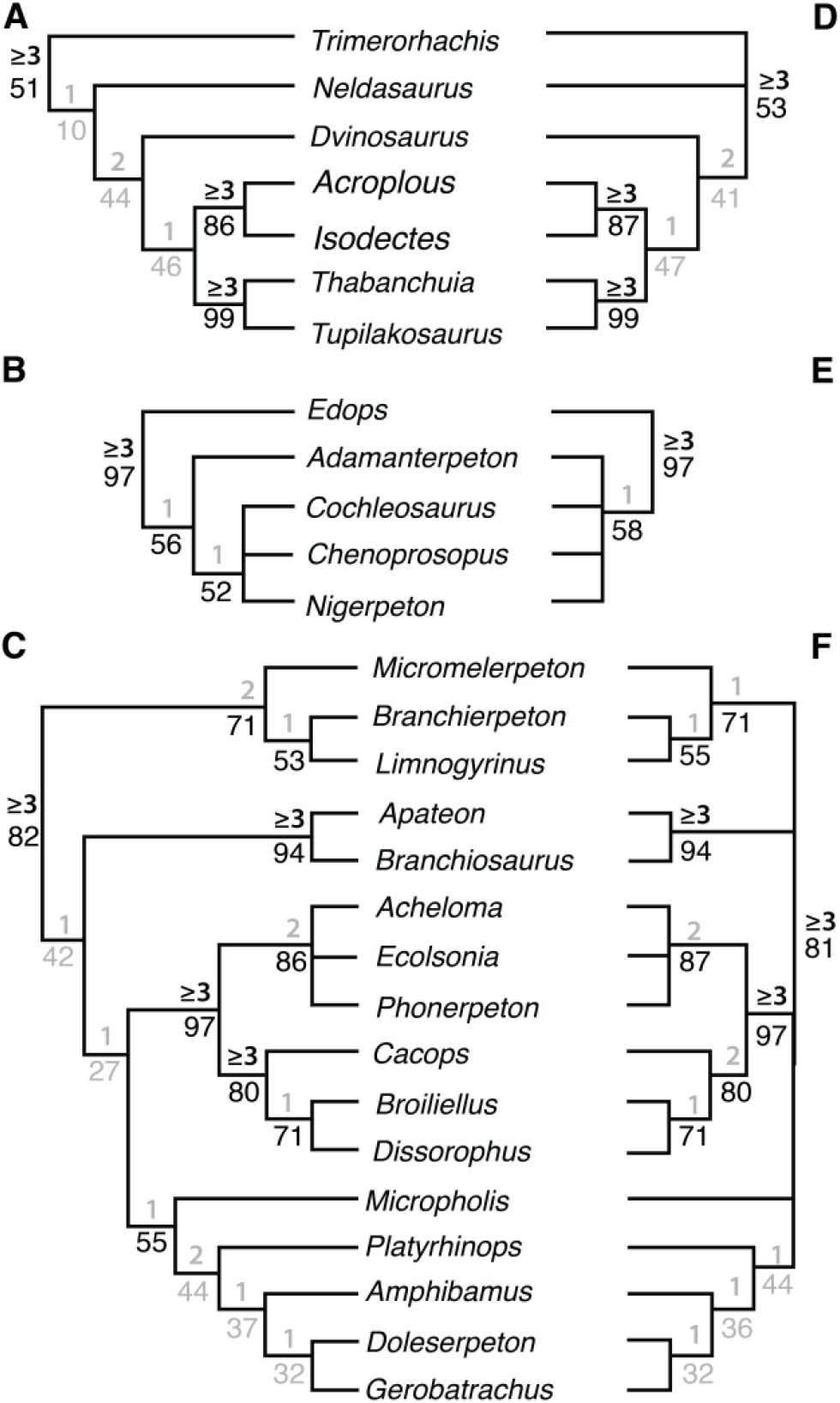
Specific relationships of Dissorophoidea, Dvinosauria, and Edopoidea recovered from reanalysis of the original matrix of Schoch (2013) in PAUP* 4.0a169 (720 MPTs, length = 677 steps), with differing taxon samples and only character 67 ordered. **A**, newly recovered topology of Dvinosauria from reanalysis from the full taxon analysis; **B**, newly recovered topology of Edopoidea from reanalysis from the full taxon analysis; **C**, newly recovered topology of Dissorophoidea from reanalysis from the full taxon analysis; **D**, newly recovered topology of Dvinosauria from the restricted taxon analysis; **E**, newly recovered topology of Edopoidea from the restricted taxon analysis; **F**, newly recovered topology of Dissorophoidea from the restricted taxon analysis. Parts D-F are reproduced from Figure 5D-F. Values in gray represent those below the thresholds for strong support (bootstrap ≥ 50%; Bremer index ≥ 3).

**Figure 8.**
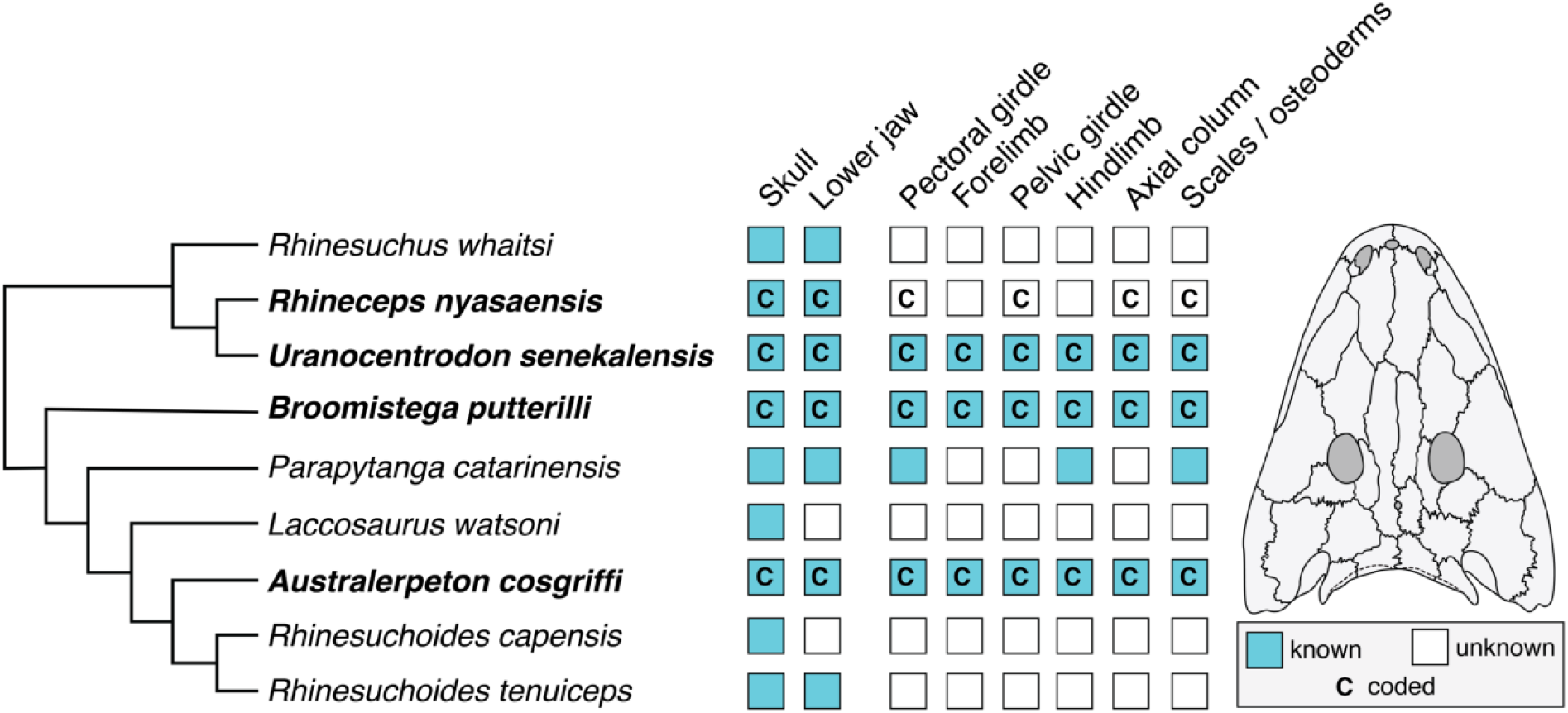
Comparison of skeletal representation and character coding among rhinesuchids sampled in this family of matrices. Topology follows that of Eltink et al. (2019) and includes *Australerpeton cosgriffi* (as with Marsicano et al., 2017, but contra Schoch, 2013, and derivates, including reanalyses of this study). A silhouette of *Rhineceps nyasaensis* (after Schoch and Milner, 2000) is provided as a representative of a sampled rhinesuchid.

Two testable hypotheses are considered here with respect to the broader discrepancies. The first is that character 67 was not the only multistate character to be ordered. When all 21 multistate characters were ordered, PAUP* recovered 1,332 MPTs with a length of 639 steps (distributed among three tree islands; CI = 0.365; RI = 0.805; Supp. Figs. 3–4), and TNT recovered 252 MPTs with the same length. These are longer than the purported MPTs reported by Schoch for either PAUP* or TNT and indicate that it is hypothetically possible that a certain, presently unknown combination of ordered and unordered multistate characters could reproduce the original results. However, without any documentation from the original study and no evidence otherwise, this is mere conjecture. It is hypothetically possible that more than one combination could recover MPTs with the original reported length, and it is also possible that no combination of character ordering can reproduce the original; ordering characters does not always increase the length of MPTs (e.g., treatment of character 67 did not affect results) or will not do so in a proportionate fashion. For example, characters 67, 75, 110, 143, 145, 158, 163, 170, 182, 187, and 191 can be reasonably inferred to occur along a morphocline. Reanalysis in PAUP* with these characters ordered recovered 3,360 MPTs with a length of 629 steps (distributed among two tree islands; CI = 0.370; RI = 0.805; Supp. Fig. 5). Even though these account for more than half of all multistate characters, the resultant MPTs were only one step longer than when none were ordered. This test therefore recovers no support that a different unspecified combination of ordered characters explains the discrepancy.

The second hypothesis, and the one favored here, is that the original search was incomplete and only located a local optimum, at least within PAUP*. Schoch (2013) did not specify any details with regards to heuristic searches in either TNT or PAUP*, such as whether ‘random’ addition sequences were employed (‘simple’ is the default in PAUP*) or, if random addition was employed, how many replicates were run. Further details regarding the TNT analyses were also lacking; the ‘traditional’ and New Technology Search options are different types of heuristic searches, and the latter has many more options than the former. When ‘simple’ addition was used (default in PAUP*), it only recovered 2,720 MPTs belonging to a single island (< 30% of all MPTs). When 10 random addition sequence replicates were used, holding one tree per step (defaults for this type of addition sequence in PAUP*), the search sometimes recovered all MPTs and sometimes recovered only a subset. Reanalysis with default parameters in TNT (10 random addition sequence replicates holding 10 trees per step) recovered only 70 MPTs (< 5% of all MPTs) after the initial search and only 1,072 MPTs (< 75% of all MPTs) after a second round of branch-swapping. The need for a second round of TBR in TNT is not universally known such that some other previous studies have only reported partial set of MPTs from the first round of branch swapping (see Langer et al., 2017; Marjanović and Laurin, 2019). Even with 10,000 random addition sequence replicates and the restricted taxon sample, TNT still failed to recover all MPTs in the first round of searching (1,427 of 1,456 MPTs).

These reanalyses demonstrate that it is not only possible but probable that incomplete searches of the original matrix (either too few replicates or too few rounds of TBR) failed to recover all MPTs. Given that this was documented in the most recent versions of each program (PAUP* 4.0 and TNT 1.5), the same possibility likely existed in the older versions utilized by Schoch (2013; PAUP* 3.1 and TNT 1.0). Arguably, those searches might have been even more incomplete and failed to identify any MPTs due to greater computational limitations. The difference in MPT length between equivalent PAUP* and TNT analyses (638 steps and 633 steps, respectively) in Schoch’s original analysis is one of the most conspicuous lines of evidence for a failure of that analysis to recover any MPTs. It is here corroborated by the recovery of MPTs of the same length in both programs upon thorough reanalysis (628 steps). It is thus likely that the original PAUP* search of Schoch (2013) failed to find any MPTs. Holding all else equal, TNT has more efficient search algorithms such that it has a better chance of finding either a better (or global) optimum than PAUP* or a larger subset of MPTs. Whether Schoch’s original TNT analysis found any MPTs is also doubtful, as those trees are five steps longer than the MPTs found in my reanalysis. Therefore, the original TNT analysis may only have found a better local optimum than PAUP*. This hypothesis is supported by the results from PAUP* showing that the MPTs were distributed among six tree islands (regardless of branch collapsing rule); incomplete searches often result from an inability to locate multiple islands given the structure of heuristic algorithms. An incomplete search could explain why Strapasson et al. (2014), a first-order derivate of Schoch (2013), recovered MPTs of the same length (633 steps) as those reported by Schoch for his TNT analysis despite the addition of the highly fragmentary *Parapytanga catarinensis* (coded for 33 of 212 characters); it would be unexpected for OTU addition with no other modifications to produce the same shortest tree length (e.g., compare Fortuny et al., 2011a, and Sidor et al., 2014).

Finally, support for an incomplete search in the original study can be gleaned from another temnospondyl study. Liu (2016) reanalyzed the unmodified capitosaur matrix of Schoch (2008) using a newer version of PAUP* and recovered MPTs of a distinctly shorter length than reported by the original study. Like Schoch (2013), Schoch (2008) did not specify what kind of heuristic search was run, and the latter also used PAUP* 3.1. The reanalysis of the original 2008 matrix by Liu (2016) used PAUP* 4.0 beta 1.0 and the parameters of Sidor et al. (2014, which followed that of Fortuny et al., 2011a:561) with 10,000 replicates, holding one tree per step, reasonable parameters for a small matrix. As with Schoch (2013), Schoch’s (2008) original search was likely incomplete and did not identify any MPTs.

To summate, reanalysis of the original Schoch (2013) matrix recovered MPTs of a length of 628 steps in both PAUP* 4.0 and TNT 1.5 (Figs. 4–5; Supp. Figs. 1–5). These trees are five steps shorter than those reported from TNT 1.0 by Schoch and are ten steps shorter than those reported from PAUP* 3.1 by Schoch. While it is impossible to ‘know’ the discrepancies without a reanalysis using every possible combination of methodological parameters (a prohibitively comprehensive undertaking), the present evidence suggests that neither of the original searches found the global optimum, either due to incomplete searches (e.g., low number of replicates) or to computational limitations of older versions of the programs. Because the reanalysis here demonstrated that PAUP* and TNT recover MPTs of the same length using the same parameters and the same matrix, the observation that the original MPTs recovered by PAUP* were longer than those recovered by TNT is considered as unequivocal evidence that Schoch’s original PAUP* search failed to find any MPTs, which would explain the differences between the strict consensus topologies recovered by each program. Because Schoch “preferred” the PAUP* topology, it is the only one figured and described in detail, and consequently, it is the one that has been propagated by numerous authors who did not reanalyze the underpinning matrix. In other words, it does not matter whether the TNT analysis found all, some, or none of the MPTs because its resultant topology has never been figured or used by any other worker. Given the findings here, the original PAUP* topology reported by Schoch should be abandoned.

I also reanalyzed the original matrix with all taxa sampled, but it cannot be compared to the original results because those were not described (e.g., number of MPTs, tree length), provided (i.e. MPTs), or figured in sufficient detail by Schoch (2013). Three of the nine taxa (*Broomistega*, *Platyoposaurus*, *Sangaia*) that Schoch excluded were not commented on in a fashion that allows for confident reconstruction of the topology from description alone. Since some clades were not recovered as monophyletic when wildcards were included (e.g., the lydekkerinids *Chomatobatrachus* and *Lydekkerina* form a grade), it cannot be assumed that uncertain taxa formed clades with nominal relatives (e.g., *Broomistega* with rhinesuchids). Reanalysis in PAUP* with only character 67 ordered recovered 720 MPTs with a length of 677 steps (distributed across four tree islands; CI = 0.344; RI = 0.797; Figs. 6–7). The resultant topology experienced further loss of resolution from the reduced taxon reanalysis (Figs. 4–5), although resolution was also restored in some areas (e.g., the basal polytomy). Certain results contradict those described by Schoch (e.g., a large polytomy with branches for *Lapillopsis nana*, *Peltobatrachus pustulatus*, Dvinosauria, Dissorophoidea, Zatracheidae, Eryopoidea, a clade of *Australerpeton* and all non-stereospondyl stereospondylomorphs, and Stereospondyli). Certain parts of the tree are equally resolved but with a different topology (e.g., Dissorophoidea is now divided into large- and small-bodied clades rather than Micromelerpetidae and Branchiosauridae forming progressively diverging clades at the base).

#### Reanalysis of Pardo et al. (2017)

Pardo et al. (2017) coded 76 OTUs (63 temnospondyls) for 345 characters. There is no express mention of any scoring changes to cells previously coded by Schoch (2013), although some were inadvertently identified as part of this study, but I have verified the conjecture of Marjanović & Laurin (2019) that Pardo et al. (2017) incorporated many of the changes made by Dilkes (2015), some of which were strictly based on personal observation. I also identified several dozen additional changes not made by Dilkes (2015). Both are detailed in Appendix 3. This dataset was analyzed with a heuristic search in PAUP* and a MCMC search in MrBayes. Per Pardo et al. (2017:E594):

> “76 taxa and 345 morphological characters were analyzed using Bayesian posterior probability in MrBayes v. 3.1.2 and using parsimony in PAUP* 4.0a151. The parsimony analysis was performed using the tree bisection–reconnection branch-swapping algorithm of PAUP*. All characters had equal weight, and none were ordered. For the Bayesian analysis, we used the default Mk model in MrBayes with variable character rates and running a Markov chain Monte Carlo for 10 million generations, sampling the posterior distribution every 100 generations. For the parsimony analysis, a heuristic search (random addition sequence with 10,000 replicates) was performed […]”

These settings were followed here. All other parsimony settings in PAUP* were left as with the reanalysis of Schoch (2013), such as holding 10 trees per step rather than the default of one, except for treatment of multistate cells, which should be set to ‘variable’ (differentiates partial uncertainty from polymorphism). This reanalysis confirmed the original results (882 MPTs with a length of 1,514 steps). If multistate taxa were treated as ‘polymorphism’ (the default), the same MPTs were merely a longer length (1,532 steps). The Bayesian analysis employed the script for their analysis that was provided in the supplemental information. Reanalysis using MrBayes v. 3.2 (Ronquist et al., 2012) confirmed the original results. It should be noted that if the average standard deviation of split frequencies (ASDSF, reported in real-time by MrBayes) is used to assess convergence, with a threshold of 0.01 (Lakner et al., 2008), only slightly more than 1.5 million generations were needed to reach convergence.

#### Institutional abbreviations

**MCN**, Museu de Ciências Naturais da Fundação Zoobotânica do RS, Porto Alegre, Brazil; **SAM-PK**, Iziko South African Museum, Cape Town, South Africa; **UMCZ**, University Museum of Zoology, Cambridge, England; **UMVT**, Museu de História da Vida e da Terra da Universidade do Vale do Rios dos Sinos, São Leopoldo, Brazil; **URGS**, Universidad Rio Grande do Sul, Vale do Rio dos Sinos, Brazil.

## RESULTS

### Summary of Coding Modifications

While it is not possible to include the full list of coding modifications and their discussion in the main text (see Appendix 2.4–2.5), they are considered to be ‘Results’ such that general trends, with select examples, are provided here to illustrate the shortcomings of previous versions of this matrix and their redress here. Three primary patterns were identified here and are termed as follows: (1) duplicated coding; (2) block coding; and (3) dependent/redundant coding. Character numbering of this section refers to that of Schoch (2013), Pardo et al. (2017), and derivates of the latter.

#### Duplicated coding

This pattern occurred when a taxon was coded for a character pertaining to a skeletal region that is unknown in that taxon and for which the particular code(s) is the same (‘duplicated’) as a certain subset of taxa that would be considered to be closely related, either by historical placement or by previous phylogenetic studies. Two examples are provided below.

Four rhinesuchids are sampled in this family of matrices: *Australerpeton cosgriffi* (which was not recovered as a rhinesuchid by Schoch, 2013, and derivates), *Broomistega putterilli*, *Rhineceps nyasaensis*, and *Uranocentrodon senekalensis*. Of these, *R. nyasaensis* is the most poorly known, with a holotype comprising two fragments of the same lower jaw (SAM-PK-7866) and a referred specimen comprising a nearly complete skull and fragments of both lower jaws (UMCZ T.259; Watson, 1962:figs. 6–8). Postcrania are entirely unknown, as is apparent from Schoch and Milner’s (2000) synoptic review of stereospondyls and Marsicano et al.’s (2017) phylogenetic revision of Rhinesuchidae. However, Schoch (2013) coded 32 of the 54 postcranial characters (58%) for this OTU. This is in stark contrast to Eltink et al. (2016, 2019) who did not code any of the 55 postcranial characters (only some overlap with Schoch, 2013) for this taxon in spite of personal observation of this taxon (note that Schoch was a co-author on the 2019 study). There is no other species of *Rhineceps*, nor another species previously assigned to the genus and now placed elsewhere, so the OTU cannot be a genus-level composite OTU. Postcranial codes for *R. nyasaensis* span the axial column, the pectoral and pelvic girdles, and dermal ossifications. Most rhinesuchids are similarly unknown for the postcranial skeleton, but coincidentally, the three taxa with sufficient postcrania to code these regions are the other three sampled rhinesuchids (Schoch and Milner, 2000; Shishkin and Rubidge, 2000; Marsicano et al., 2017; Fig. 8). All 32 unsubstantiated postcranial codes of *R. nyasaensis* were identical to the codes of *U. senekalensis*. These codes were also identical to *B. putterilli* when the latter was coded (i.e. they differ only in distribution of missing data; note, however, that most of the postcranial codes of *B. putterilli* were added by Pardo et al., 2017, without comment or documentation). Some differences in character states are noted from the scorings of the Brazilian *A. cosgriffi*, which is unsurprising given the correspondent cranial disparity and geographic separation from the other three taxa (all from southern Africa).

The second example is the rhytidosteid *Sangaia lavina*. Rhytidosteids are a morphological disparate group (Dias-da-Silva and Marsicano, 2011) compared to other temnospondyl families and have consistently been recovered as paraphyletic or polyphyletic (e.g., certain analyses of Dias-da-Silva and Marsicano, 2011; McHugh, 2012; Schoch, 2013, and derivates; Maganuco et al., 2014). *Sangaia lavina* was described on the basis of a partial skull with most of the roof and occiput intact but lacking almost all of the palate (UMVT 4302; Dias-da-Silva et al., 2006:figs. 2–3) and a paratype consisting of the basal plate of the parasphenoid and pterygoid (UMVT 4303; Dias-da-Silva et al., 2006:figs. 4–5). Dias-da-Silva and Marsicano’s (2011) revision of Rhytidosteidae listed two additional specimens, PV 0497 T and MCN PV 2606, which were stated to be “skull fragments.” PV 0497 T was described and figured as an indeterminate temnospondyl by Dias-da-Silva (1998:103–104, fig. 2D), and MCN PV 2606 was described and figured as an indeterminate rhytidosteid by Dias-da-Silva et al. (2005:169, fig. 6). Both only preserve skeletal regions found in the holotype. Schoch (2013:table 1) listed only material reposited at the URGS (which he did not personally examine) and Dias-da-Silva et al. (2006) as the sources.

Based on the literature, no information is available about the: premaxillae, vomers, palatines, ectopterygoids, lower jaws, or postcranial skeleton. Except for the dorsal surface of the premaxillae, these regions were not reconstructed, and *Sangaia lavina* was mostly left uncoded for palatal characters by Dias-da-Silva and Marsicano (2011), who originally described the taxon (Dias-da-Silva et al., 2006); those that were coded pertained to the posterior regions that are preserved. Schoch (2013) coded 23 characters that require the premaxillae and/or anterior palate to be known, even though there is no indication that these are preserved in any specimen. Another two such characters were coded by Pardo et al. (2017), who also gave no indication of personal examination, and other six such characters were coded by Schoch et al. (2020b). There were also nine postcranial characters coded between Schoch (2013) and Schoch et al. (2020b). Many of the unsubstantiated codes for cranial characters were for successive characters, such as those for the dorsal exposure of the premaxilla (10–14), for the choana (99–101), and for anterior palatal depressions or openings (91–95). Most of these are not presence/absence characters and require characterization of a specific condition (e.g., the shape of the choana, original characters 99 and 101). For these characters, the codes of *S. lavina* were the same as the condition that was most common in other stereospondyls (e.g., an oval choana).

These examples illustrate issues with codes that seem to have been assumed on the basis of inferred relatedness, an inherently circular approach to phylogenetics. Because of the stringent evidentiary standard required to change codes, it is possible that other codes like these exist for other taxa but that they did not meet the evidentiary standard needed to overturn the original code. An example is the ilium of the trematopid *Phonerpeton pricei*, an element which has never been described or figured (Olson, 1941; Dilkes, 1990). The taxon was coded for the derived condition for all five iliac characters by Schoch (2013). These derived conditions also occur in all other sampled olsoniforms (some represent synapomorphies sensu Schoch and Milner, 2014) and could have been assumed by inferred relatedness. Because Dilkes (1990:224) mentioned “fragments of pectoral and pelvic girdles” for one specimen, I did not change these codes, but there is no clear indication that the ilium is known to any appreciable degree.

#### Block coding

This pattern differs from duplicated coding in that it presents at the character scale rather than at the taxon scale. In this pattern, it appears as if a subset of taxa with a specific condition were coded for one state and (nearly) all other taxa were coded for the opposing state, regardless of whether that condition is verified or can even be assessed. This pattern presented most often in characters with an unusually low frequency of missing data compared to other characters for the same skeletal region that should have similar frequencies of missing data. As with duplicated coding, block coding was more readily identifiable among characters for less well represented skeletal regions. Like duplicated coding, codes may have been assumed by inferred relatedness. Three examples are given below.

Characters 198–202 of Schoch (2013) and derivates refer to the ilium, more specifically to either the entire shaft or to the distal end of the shaft. Because the postcrania of some sampled taxa is entirely unknown (e.g., *Zatrachys, Sangaia*), with many others for which the ilium is not among the known postcrania (e.g., *Nigerpeton*), a relatively high frequency of missing data is expected. The number of taxa coded as unknown for each character by Schoch (2013) is as follows: 27 (198, ilium, shaft); 30 (199, ilium, dorsal end); 27 (200, ilium, height); 19 (201, ilium, orientation); and 2 (202, ilium, tip). Character 202 was conspicuous because nearly every OTU was coded. This character refers to whether the dorsal end of the ilium is double- or single-headed and thus requires that the iliac blade and its terminus are preserved. Presumably, any taxon for which this can be determined should also be able to be coded for character 199, which refers to whether the dorsal end (‘tip’) of the ilium is ‘continuous’ or ‘much broadened,’ yet character 199 was left uncoded for 28 taxa that were coded for character 202. Almost every taxon was coded for 202–1 (single-headed); only *Proterogyrinus, Trimerorhachis*, and *Dvinosaurus* were coded for 202–0. For 15 taxa, character 202 was the only one of the five iliac characters that was coded: *Capetus, Iberospondylus, Zatrachys, Lapillopsis, Rhineceps*, *Broomistega, Chomatobatrachus*, *Sangaia*, *Laidleria*, *Batrachosuchus*, *Edingerella*, *Watsonisuchus*, *Trematosaurus*, *Eocyclotosaurus*, and *Cyclotosaurus*. Several of these have no documentation of any postcrania (e.g., *Zatrachys*, *Watsonisuchus*), including two mentioned above as examples of duplicated coding (*Rhineceps*, *Sangaia*). Based on the number of taxa with unsubstantiated codes, it is interpreted here that this character was coded for the few taxa with a distinct apomorphy and was then coded for the opposing state for almost every other taxon on the assumption that the character is strongly dichotomous. It seems highly improbable that unpublished distal ends of the iliac shaft are known for all 15 taxa and have yet to be published. Furthermore, Schoch (2013:table 1) noted a lack of personal observation for several of these taxa. The only two uncoded taxa, the rhytidosteid *Peltostega* and the trematosaur *Lyrocephaliscus*, share no apparent attribute that would explain their coding compared to other taxa (*Peltostega* has no known postcrania at all, while anterior postcrania are known from *Lyrocephaliscus*). It may be noteworthy that these were recovered as sister taxa in Schoch’s analysis and derivates despite being from nominally different families.

Character 28 of Schoch (2013) referred to the presence or absence of palpebral ossicles. Because these occur in the orbit, it could be argued that any taxon with a preserved orbit could be coded; indeed, 66 of the 72 taxa sampled by Schoch were coded. In this instance, it is the pattern of the six uncoded taxa that was conspicuous, as all six were dissorophoids: non-branchiosaurid amphibamiform *Gerobatrachus*, the dissorophids *Cacops* and *Dissorophus*, and all three trematopids, *Acheloma, Ecolsonia*, and *Phonerpeton*. All other dissorophoids except for the two branchiosaurids were coded for 28–1; no other temnospondyl was coded for 28–1. These omissions cannot be considered a reflection of low sample size. Of the six, only *Gerobatrachus* is known from only one cranial specimen (e.g., Williston, 1910; Olson, 1941; DeMar, 1968; Berman et al., 1985; Dilkes, 1990; Anderson et al., 2020), while many taxa coded for 28–0 are also known from only one specimen (e.g., *Batrachosuchus*, *Siderops*), or from only one orbit (e.g., *Sangaia*). For most taxa, the scleral ring is not actually known despite being a specifier in the character states. One interpretation for this coding pattern is that it was assumed that the six uncoded dissorophoids did have palpebral ossifications that were lost post-mortem and that coding them for ‘ossifications absent’ would be misleading since many dissorophoids do have palpebral ossifications (e.g., Berman et al., 2010; Holmes et al., 2013; Maddin et al., 2013; Schoch and Sues, 2013). This would require a deliberate decision to not code them that was not applied to other taxa of similar or lesser preservational quality and completeness. Furthermore, it is circular logic to assume that all non-branchiosaurid dissorophoids had palpebral ossifications, as these are unknown in many such taxa with well-preserved cranial material (e.g., *Acheloma*), or to assume that palpebral ossifications could not occur in another clade (e.g., they occur in some ‘microsaurs’; Maddin et al., 2011; Huttenlocker et al., 2013; Pardo et al., 2015; Gee et al., 2019). *Balanerpeton* has non-sclerotic ring ossifications that are sometimes considered to be palpebral ossifications (compare Milner and Sequeira, 1993, with Sigurdsen and Bolt, 2010).

Finally, character 197 of Schoch (2013) referred to manual digit count and was coded for 41 of the 72 taxa sampled by Schoch (2013). Temnospondyls have long been assumed to share a condition of four manual digits (197–1; e.g., Warren and Snell, 1991; Dilkes, 2015; Konietzko-Meier et al., 2020), and this is a common rationale for associating tetradactyl handprints with temnospondyls (e.g., Stimson et al., 2012, 2016; Marsicano et al., 2014; Bird et al., 2020; Cisneros et al., 2020; Mujal and Schoch, 2020). Predictably, every previously coded temnospondyl other than *Metoposaurus* was coded for a tetradactyl manus; *Metoposaurus* was only correctly coded as pentadactyl because of personal communication to Schoch (Konietzko-Meier et al., 2020:1153). However, even semi-articulated manuses are extremely rare for temnospondyls. Dilkes (2015), who surveyed the carpus and tarsus of certain taxa, added eight characters related to these elements to the matrix of Schoch (2013) and carried over all 72 taxa, but only 25 temnospondyls were coded for at least one of the new characters. Only two (*Balanerpeton*, *Eryops*) were coded for all eight. Finally, while many temnospondyls did not ossify their carpals (196–0), there were several taxa coded as unknown for the ossification of the carpals but for which four manual digits were coded (e.g., *Edops*). No manus of *Edops* is complete enough as to be certain of four manual digits, yet so uncertain as to whether at least one carpal was ossified (there is no mention of the manus in the last description by Romer and Witter, 1942; in the synopsis of non-stereospondylomorphs by Schoch and Milner, 2014; or in the review of the carpus in Temnospondyl by Dilkes, 2015). This pattern suggests that there was a blanket working assumption that most sampled temnospondyls had a tetradactyl manus, even if no such material is known.

It appears that certain unstated assumptions were made in coding characters like these that could be perceived as strongly dichotomous among temnospondyls. These assumptions may not be unreasonable, but phylogenetic analyses are tests of, not tests based on, assumptions. There is at least one example that introduced definitive errors. Original character 57 referred to a ventral exposure of the jugal on the palate (the insula jugalis) that is present posterior to the ectopterygoid and that frames the subtemporal vacuity anteriorly in most stereospondylomorphs. In some studies, it was expressly identified as a synapomorphy of Stereospondylomorpha (e.g., Yates and Warren, 2000) or the more inclusive Eryopiformes (e.g., Eltink et al., 2019). This character was coded for 65 of the 72 taxa by Schoch. All coded stereospondyls were coded as 57–1 (exposure present). There is, however, no unequivocal evidence for this exposure in at least three stereospondyls coded as such (Fig. 9): *Rhineceps* (Watson, 1962:233, 235, fig. 7); *Edingerella* (Steyer, 2003:548, fig. 2; Maganuco et al., 2009:12–17, fig. 10); and *Trematolestes* (Schoch, 2006:34, fig. 4B). Watson’s description of *Rhineceps* may be considered more skeptically due to datedness, as this exposure was reconstructed by Schoch and Milner (2000:fig. 54) and was coded as present by Marsicano et al. (2017; character 73 therein), but the equivalent character was left uncoded by Eltink et al. (2019; character 55 therein). *Edingerella* and *Trematolestes* were more recently (re)described. The condition of *Trematolestes* is unclear from specimen illustrations, but Schoch (2006:34) stated that the “lateral and posteriormost portions of the palate are preserved.” Because some codes for this OTU could only be substantiated from the reconstruction (Schoch, 2006:fig. 4B), the lack of a described or reconstructed ventral exposure of the jugal was incorporated here. *Edingerella* definitively lacks this exposure based on previous studies. Furthermore, Holmes (1984:453) identified this exposure in the outgroup, *Proterogyrinus*, in which the “processus alaris” separates the ectopterygoid from the subtemporal vacuity (as in temnospondyls), and Milner and Sequeira (1998:272) identified this exposure in the edopoid *Adamanterpeton*. A small exposure is also found in the dvinosaur *Dvinosaurus* (e.g., Bystrow, 1938:fig. 1; Schoch and Milner, 2014:fig. 21B). The character thus has a patchier distribution within Stereospondyli and is more common among non-stereospondylomorphs than was originally coded. Collectively, this demonstrates how assumptions of a stark dichotomy introduced errors to the matrix.

**Figure 9.**
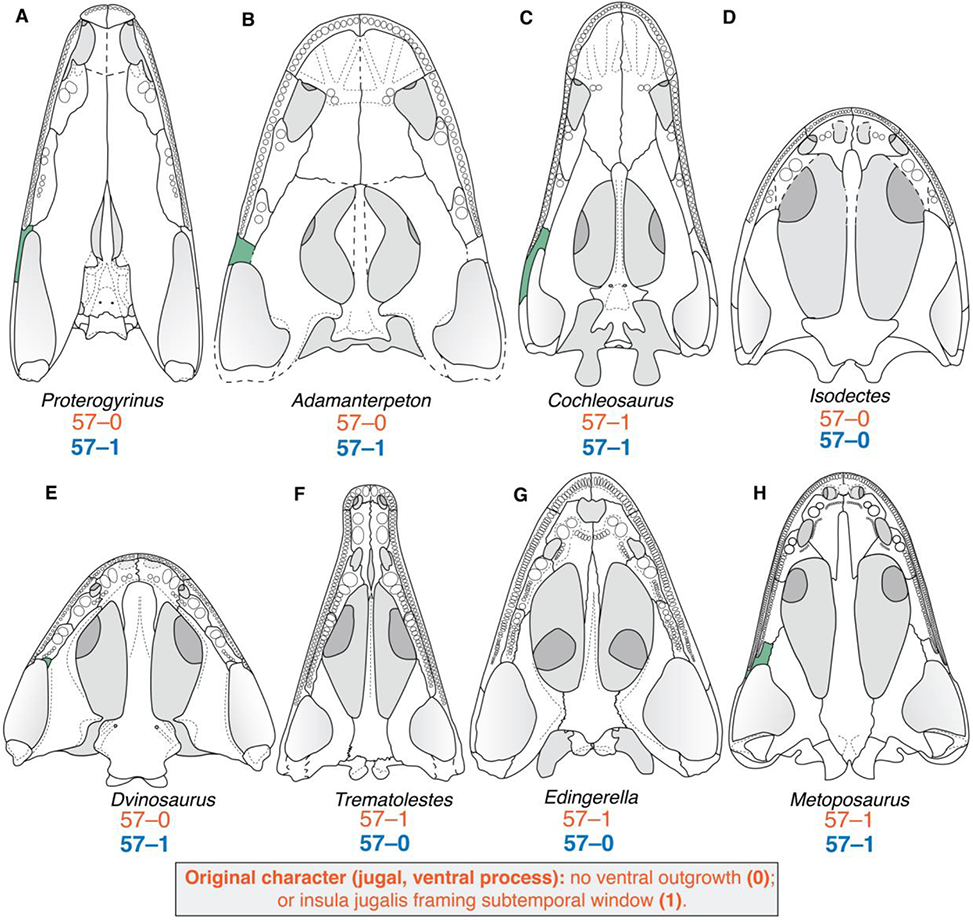
Representative temnospondyl skulls in ventral view, shown to compare taxa apparently lacking a ventral palatal exposure of the jugal with select taxa having this exposure. **A**, the anthracosaur *Proterogyrinus scheelei*, which served as the outgroup in this study (after Holmes, 1984); **B**, the edopoid *Adamanterpeton ohioensis* (after Milner and Sequeira, 1998); **C**, the edopoid *Cochleosaurus bohemicus* (after Sequeira, 2003); **D**, the ‘dendrerpetid’ *Balanerpeton woodi* (after Milner and Sequeira, 1993); **E**, the dvinosaur *Dvinosaurus primus* (after Schoch and Milner, 2014); **F**, the dvinosaur *Isodectes obtusus* (after Sequeira, 1998); **G**, the trematosaur *Trematolestes hagdorni* (after Schoch, 2006); **H**, the capitosaur *Edingerella madgascariensis* (after Maganuco et al., 2009); **I**, the metoposaurid *Metoposaurus krasiejowensis* (after Sulej, 2007). Orange text represent the original character and codes from Schoch (2013); blue text represents the revised codes. Silhouettes not to scale.

**Figure 10.**
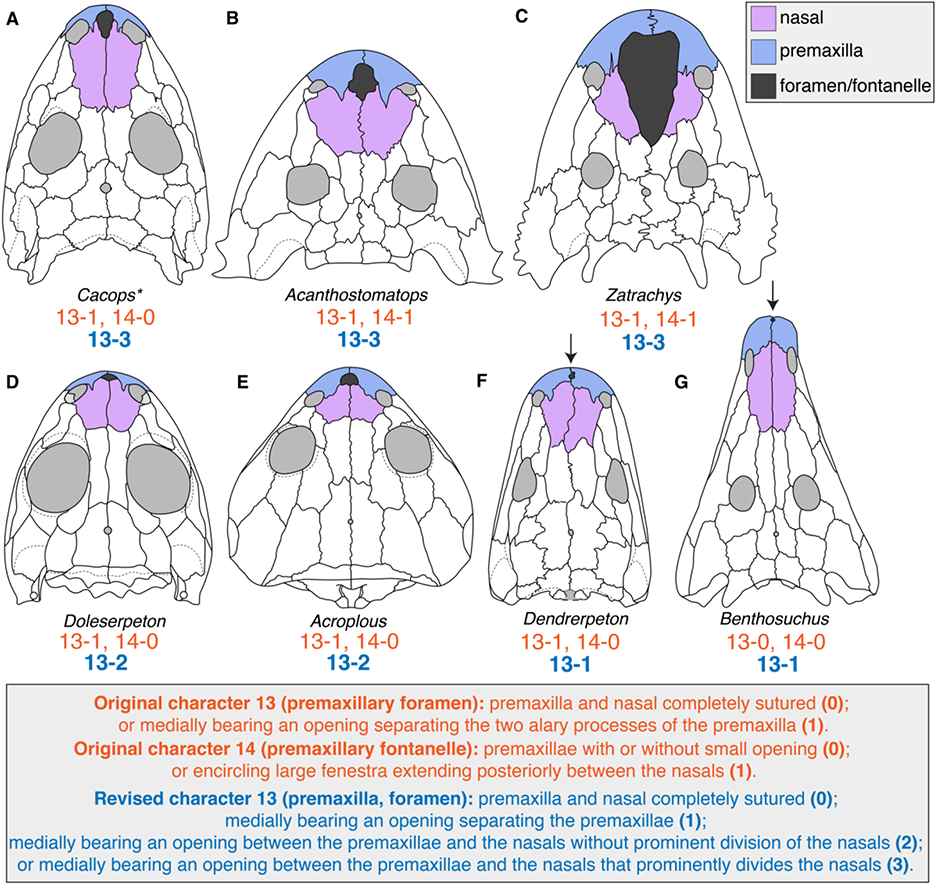
Representative temnospondyl skulls with median rostral openings shown in dorsal view to illustrate the partial redundancy in the original coding of characters 13 (premaxillary foramen) and 14 (premaxillary fontanelle). **A**, the dissorophid *Cacops morrisi* (after Reisz et al., 2009); **B**, the zatracheid *Acanthostomatops vorax* (after Schoch and Milner, 2014); **C**, the zatracheid *Zatrachys serratus* (after Schoch and Milner, 2014); **D**, the stereospondylomorph *Archegosaurus decheni* (after Witzmann, 2005); **E**, the amphibamiform *Doleserpeton annectens* (after Sigurdsen and Bolt, 2010); **F**, the dvinosaur *Acroplous vorax* (after Englehorn et al., 2008); **G**, the ‘dendrepetid’ *Dendrerpeton helogenes* (after Holmes et al., 1998). Arrows are used to indicate the small foramen in *Dendrerpeton* and *Benthosuchus.* In contrast to other comparative cranial diagrams, both sides of the skull are colored here to demonstrate the degree to which the rostral opening divides the premaxillae and the nasals. Orange text represents the original characters and codes from Schoch (2013); blue text represents the revised character and codes. The asterisk (*) next to *Cacops* is because the OTU was restricted to *Cacops aspidephorus* (in which the presence / size of a fontanelle is unclear; Anderson et al., 2020) rather than as a composite including *C. morrisi* (in which a fontanelle is entirely defined; Reisz et al., 2009), the revised code reflects the proper code if the latter were utilized as the sole OTU. Silhouettes not to scale.

#### Character dependency/redundancy

The third pattern is dependent/redundant coding. Character dependencies and their treatment has been exhaustively covered in the literature (e.g., Maddison, 1993; Lee and Bryant, 1999; Strong and Lipscomb, 1999; Sereno, 2007; Vogt, 2018, and references therein); the underlying point is that dependent characters are problematic. For the sake of this study, characters can be defined as ‘primary’ or ‘secondary.’ A primary character refers to the presence or absence of a given feature (e.g., otic notch), and a secondary character refers to different aspects of that feature (e.g., otic notch shape), which requires that said feature is present. This division is equivalent to the ‘neomorphic’ (presence/absence) and ‘transformational’ (other attributes) binary of Sereno (2007). If the primary character is coded as ‘absent,’ any secondary character cannot be coded. However, I identified a large number of characters where their treatment failed to account for different forms of dependency, collectively accounting for hundreds of problematic codes. Two examples are given below.

In full dependency, a primary character must be ‘present’ in order for any secondary character to be coded. This is the ‘ontological dependency’ of Vogt (2018). One example is the trio of osteoderm characters (original characters 209–211). Character 209 expresses absence (209–0) versus two forms of presence (209–1 [restricted to midline], 209–2) [forming carapacial covering]). Character 210 expresses whether osteoderms lie in a single series (210–0) or in a double series with overlap of one series by another (210–1). Character 211 expresses relative width of osteoderms, either narrow (211–0) or wide (211–1). Few temnospondyls have osteoderms (e.g., DeMar, 1966; Panchen, 1959; Hellrung, 2003; Dilkes and Brown, 2007; Witzmann and Soler-Gijón, 2010), and only eight taxa sampled across all versions of the matrix were coded as 209–1 or 209–2. Therefore, only eight taxa should have been coded for characters 210 and 211. However, of the 72 taxa sampled by Schoch (2013), 70 were coded for both characters 210 and 211, specifically for a ‘single series’ of ‘narrow’ osteoderms (210–0, 211–0). This includes nearly four dozen taxa expressly coded as lacking osteoderms (209–0) and more than a dozen taxa for which character 209 was originally uncoded (e.g., *Adamanterpeton*, *Acroplous*, *Zatrachys*, *Benthosuchus*). It is inherently impossible for osteoderms to be a ‘single series’ or ‘narrow’ when they either do not exist or their presence/absence cannot be assessed (as in taxa with no known postcrania, like *Zatrachys*). This mistreatment of the character is differentiated from block coding because none of the secondary characters’ states (210 and 211) are applicable to a taxon lacking the primary feature (osteoderms, 209–1 or 209–2). Coding taxa without osteoderms for a single series of narrow osteoderms falsely homologized taxa with no osteoderms (e.g., *Doleserpeton*) with taxa that have a single series of narrow osteoderms (e.g., *Cacops*). Examples of other features with a primary and at least one secondary character include the lateral line grooves (original 33–36), the otic notch (original 51–52), the tabular horn (original 65–66), and pleurocentra (original 168–170); each of these primary features is absent in at least one of the taxa sampled by Schoch (2013). Note that a full dependency exists even when a primary character is not sampled. For example, characters 210 and 211 still cannot be coded for taxa without osteoderms in an analysis without character 209 (i.e. omitting character 209 would not rectify the problem). This point is particularly important with the expanded character and taxon sample of Pardo et al. (2017), which includes lissamphibians, whose crania are greatly reduced compared to early tetrapods, as many characters were expressly designed for temnospondyls or other extinct tetrapods. There were several hundred erroneous codes for a secondary character that requires the presence of a feature that is absent in many sampled taxa.

In partial dependency, a certain state of one character always correlates with a certain state of another. In most instances, this is the ‘semantic dependency’ of Vogt (2018). Original characters 13 (premaxillary foramen) and 14 (premaxillary fontanelle) were binary characters that referred to openings in the snout at the midline (Fig. 10). State 14–1 (fontanelle present) was restricted to the zatracheids *Acanthostomatops* and *Zatrachys*, but it likely represents merely an enlarged premaxillary foramen because the position (between at least the premaxilla and nasal) is the same as taxa coded for 13–1 (premaxillary foramen present) and 14–0 (fontanelle absent) like many dissorophoids. The zatracheids were coded for both 13–1 and 14–1, but there is only one median opening, so the presence of a fontanelle required the presence of a foramen because a fontanelle was merely a greatly enlarged foramen (14–1 predicts 13–1, but the inverse is not true). Coding the zatracheids as having both a premaxillary foramen and a premaxillary fontanelle without accounting for the partial dependency resulted in upweighting of the median rostral opening. Zatracheids are only united by one shared feature in this regard – a proportionately large rostral opening – but as originally coded, they were united by two shared states, an apomorphic large rostral opening and a small rostral opening shared with a variety of other taxa. By the same token, any taxon without any medial rostral opening shared two states, foramen absent and fontanelle absent, with similar taxa (13–0 predicts 14–0, but the inverse is not true). In this case, zatracheids should either have been coded as inapplicable for character 13, or, as I did here, both characters should be combined into a multistate character. A third approach (not taken here), following Sereno (2007) would be to restrict one character to the presence/absence of a rostral opening (the neomorphic character) and to restrict the other to the relative size (the transformational character). Although the coding following Sereno’s philosophy would be most similar to that based on the original construction by Schoch, the construction itself was problematic as detailed above. In analyses intending to be equally weighted, failure to account for dependencies will bias certain anatomical features and artificially increase support for certain nodes. Certainly, an argument can be made for weighting of certain features, but in the osteoderm example, the absence of osteoderms was unintentionally overweighted, whereas it seems more appropriate to upweight the presence of osteoderms as a rare, derived, neomorphic feature that was independently acquired in a handful of clades. Assigning this weight a priori then becomes another parameter in need of specification and justification.

#### Patterns across matrices

Problematic codes were not evenly distributed within the matrices. The overwhelming majority of these codes were for cells originally coded by Schoch (2013). This observation is particular important because this is the original matrix, which means that these codes were propagated in all subsequent versions unless otherwise stated. Of the three classes of problematic codes identified above, none was confidently identified in taxa and/or characters added by Pardo et al. (2017). The only systemic issue identified in the latter matrix was asymmetric treatment of a character (e.g., a tentacle was coded as absent for most temnospondyls [321–0] but was left as unknown in non-metoposaurid trematosaurs and early diverging stereospondyls). In these instances, there was some pattern of omissions, but no clear pattern for why certain taxa were not coded, in contrast to the palpebral ossifications example given above. These occurrences were rare, and there were no systemic issues identified within a given taxon except for the brachyopid OTU, for which Pardo et al. (2017) coded many lower jaw characters.

These observations indicate that the majority of patterns relate to different coding practices by different workers, which is also evidenced in codes for the fifteen new characters added by Schoch et al. (2020b), most of which are simple presence/absence characters. There is very little missing data for these characters, seven of which were coded for all 62 sampled taxa. These seven characters include the presence/absence of the interclavicle and cleithrum, even though postcrania are entirely unknown for many of the sampled taxa. Most of these characters, appear to have been subjected to block coding because they are almost all strongly dichotomous (e.g., a given state only occurred in lissamphibians). Corrections and modifications to these codes were addressed by Kligman et al. (2021) and are not detailed here.

### Reanalysis of Schoch (2013)

The original matrix comprised 72 taxa (70 temnospondyls) and 212 characters, for a total of 15,264 cells. The modified matrix comprises the same 72 taxa and 207 characters, not all of which are identical to those in the original matrix, for a total of 14,094 cells. Seventeen multistate characters can be reasonably inferred to occur along a morphocline (39, 67, 74–75, 103, 107, 110, 137–138, 143, 145, 158, 163, 170, 182, 187, 191; all numbering follows the original scheme). Of the 207 characters, none is constant and six are parsimony-uninformative. The original matrix contained zero multistate cells (polymorphisms or partial uncertainty); 1,731 cells coded as unknown (11.3%); and 28 cells coded as inapplicable (0.720%). The revised matrix contains 95 polymorphisms and 39 cells coded as partially uncertainty (134 multistate cells; 0.950%); 2,284 cells coded as unknown (16.2%); and 491 cells coded as inapplicable (3.48%).

#### Without character ordering

The analysis of the modified version of the matrix in PAUP* 4.0a169 with 63 taxa and no characters ordered recovered 1,044 MPTs with a length of 964 steps (CI = 0.355; RI = 0.715; CPU time = 00:36:13; Fig. 11). MPTs were distributed across two tree islands of dissimilar size (1–981, 982–1044). The equivalent analysis in TNT recovered 900 MPTs with a length of 874 steps (CI = 0.288; RI = 0.715; CPU time = 00:02:21; Supp. Fig. 6). Twenty-six MPTs were only recovered after the second round of branch swapping. The strict consensus is identical between the analyses, is largely resolved, and departs from the original results of Schoch (2013) and my reanalysis of the original matrix here (Figs. 4–5). Firstly, the earliest diverging clade is Dvinosauria, followed by a polytomy of *Balanerpeton*, *Dendrerpeton*, and all other temnospondyls; in contrast, Schoch reported Edopoidea as the earliest diverging clade, a ‘Dendrerpetidae’ clade, and Dvinosauria as the sister group to Dissorophoidea + Zatracheidae. The relationships of the highly nested short-faced taxa (Brachyopoidea, Plagiosauridae, *Laidleria*) is now completely unresolved other than a monophyletic Plagiosauridae, whereas previously, *Laidleria* and Plagiosauridae nested within Brachyopoidea as sister taxa. Finally, the capitosaur *Edingerella* is recovered as the sister taxon to the clade encompassing brachyopoids, plagiosaurids, and trematosaurs; and the capitosaur *Wetlugasaurus* is recovered as the sister taxon to the clade containing all other higher stereospondyls (brachyopoids, capitosaurs, plagiosaurids, trematosaurs). Compared to the strict consensus recovered from the reanalysis of Schoch’s original matrix (Figs. 4, 5), the differences are almost exclusively related to the resolution of previously unresolved nodes. The loss of resolution in the more inclusive Brachyopoidea is the only large-scale change. Minor loss of resolution has also occurred within Dissorophoidea.

**Figure 11.**
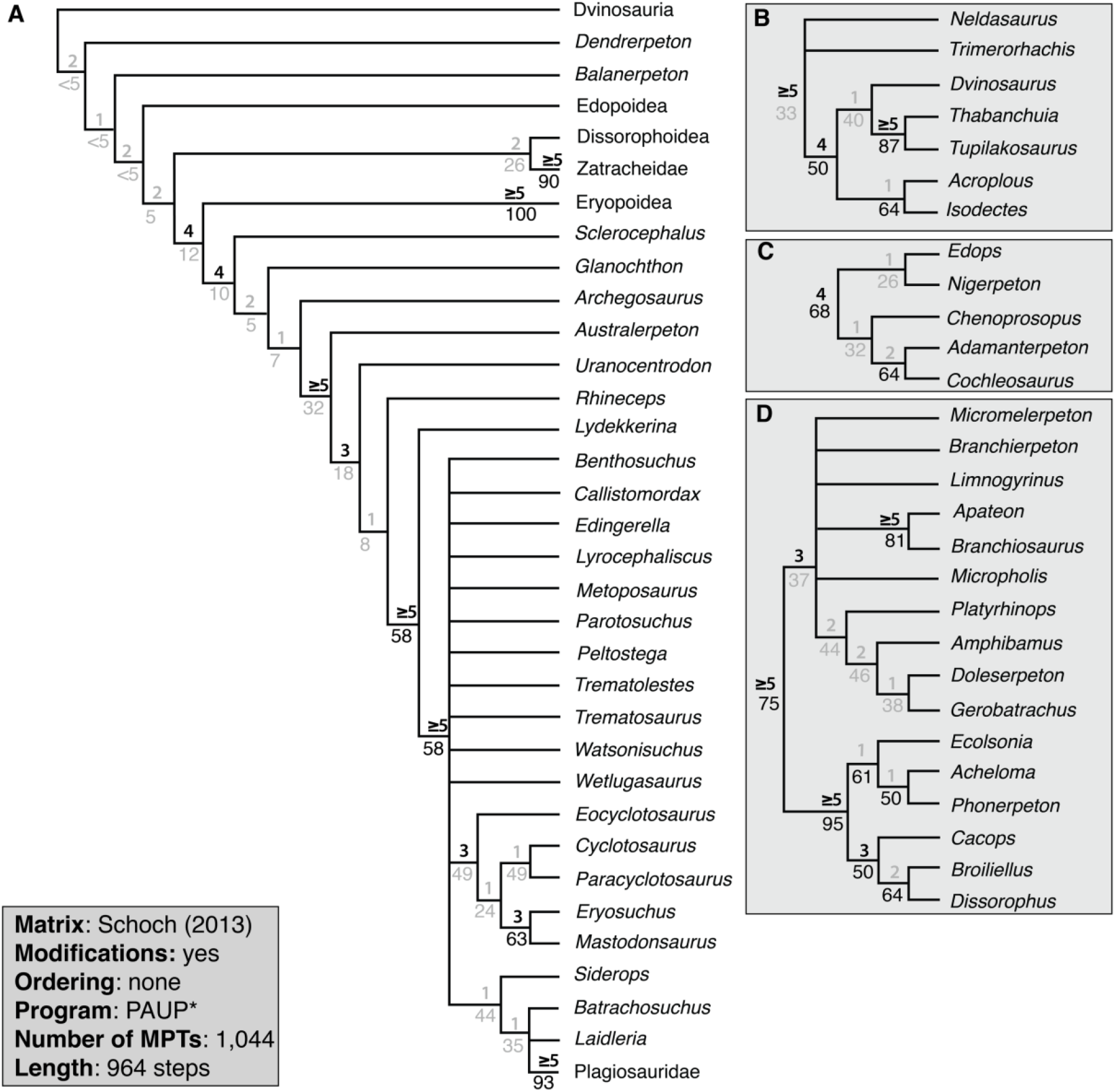
Strict consensus topology recovered from analysis of the modified Schoch (2013) matrix using PAUP*, with the restricted taxon sample and all multistate characters unordered. **A**, topology of Temnospondyli, with select clades collapsed for visual clarity; **B**, topology of Dvinosauria; **C**, topology of Edopoidea; **D**, topology of Dissorophoidea. Values above lines represent Bremer decay index; values below lines represent bootstrap support. Values in gray represent those below the thresholds for strong support (bootstrap ≥ 50%; Bremer index ≥ 3). Collapsed nodes representing only two species (Dendrerpetidae [*Balanerpeton*, *Dendrerpeton*], Eryopoidea [*Eryops*, *Onchiodon*], Metoposauroidea [*Callistomordax*, *Metoposaurus*], Plagiosauridae [*Gerrothorax*, *Plagiosuchus*], Rhinesuchidae [*Rhineceps*, *Uranocentrodon*], Zatracheidae [*Acanthostomatops*, *Zatrachys*]) have their support values listed at the tips and are condensed for visual clarity.

Statistical support is generally low across the tree. There were no nodes recovered in the 50% majority-rule bootstrap tree (values here reported from PAUP*) that were not recovered in the strict consensus other than a Rhinesuchidae that includes all three nominal members (*Australerpeton*, *Rhineceps*, *Uranocentrodon*; 54%); *Trimerorhachis* + *Neldasaurus* (56%); and *Acheloma* + *Phonerpeton* (long-snouted trematopids, 58%).

The results of the original analysis with the full taxon sample were not figured or reported in detail (e.g., number and length of MPTs) by Schoch (2013). The analysis of the modified matrix with all 72 taxa and no characters ordered recovered 45 MPTs with a length of 1,063 steps (distributed across one tree island; CI = 0.332; RI = 0.705; CPU time = 00:08:53; Fig. 12). The equivalent analysis in TNT with no characters ordered recovered 45 MPTs with a length of 966 steps (CI = 0.27; RI = 0.705; CPU time = 00:02:31; Supp. Fig. 7). All MPTs were recovered on the first round of branch swapping. The strict consensus is identical between these analyses and is largely unchanged from the analysis with the restricted taxon sample (Fig. 11). Notable departures include a large basal polytomy of Edopoidea, Dendrerpetidae, the wildcard *Capetus*, and a clade of all non-dvinosaur temnospondyls; and the partial collapse of the higher stereospondyls (Brachyopoidea, Capitosauria, Plagiosauridae, Trematosauria). However, other parts of the tree are more resolved, such as Brachyopoidea (inclusive of *Laidleria* and plagiosaurids) and the recovery of Dendrerpetidae. Most of the wildcards sampled in this analysis were recovered adjacent to the clade they have historically been allied with or placed in; examples include *Greererpeton* outside of Temnospondyli; *Capetus* in a basal polytomy with ‘Dendrerpetidae,’ Edopoidea, and all other non-dvinosaurs; and *Platyoposaurus* as the latest diverging non-stereospondyl stereospondylomorph. The two more notable depatures are *Peltobatrachus*, in a polytomy with *Iberospondylus*, Dissorophoidea, Zatracheidae, and Eryopiformes; and *Lapillopsis*, which nests securely within Stereospondyli in the clade including all members of Superstes other than nominal lydekkerinids. *Peltobatrachus* has historically been considered to be a stereospondyl (e.g., Panchen, 1959) and has sometimes been recovered as such (e.g., Eltink et al., 2019). *Lapillopsis* has also historically been considered to be a stereospondyl (Yates, 1999), but previous derivates of this matrix recovered it as the sister taxon to Dissorophoidea, reflecting homoplasy between small-bodied terrestrial taxa that originally led to hypothesized relatedness (Warren and Hutchinson, 1990).

**Figure 12.**
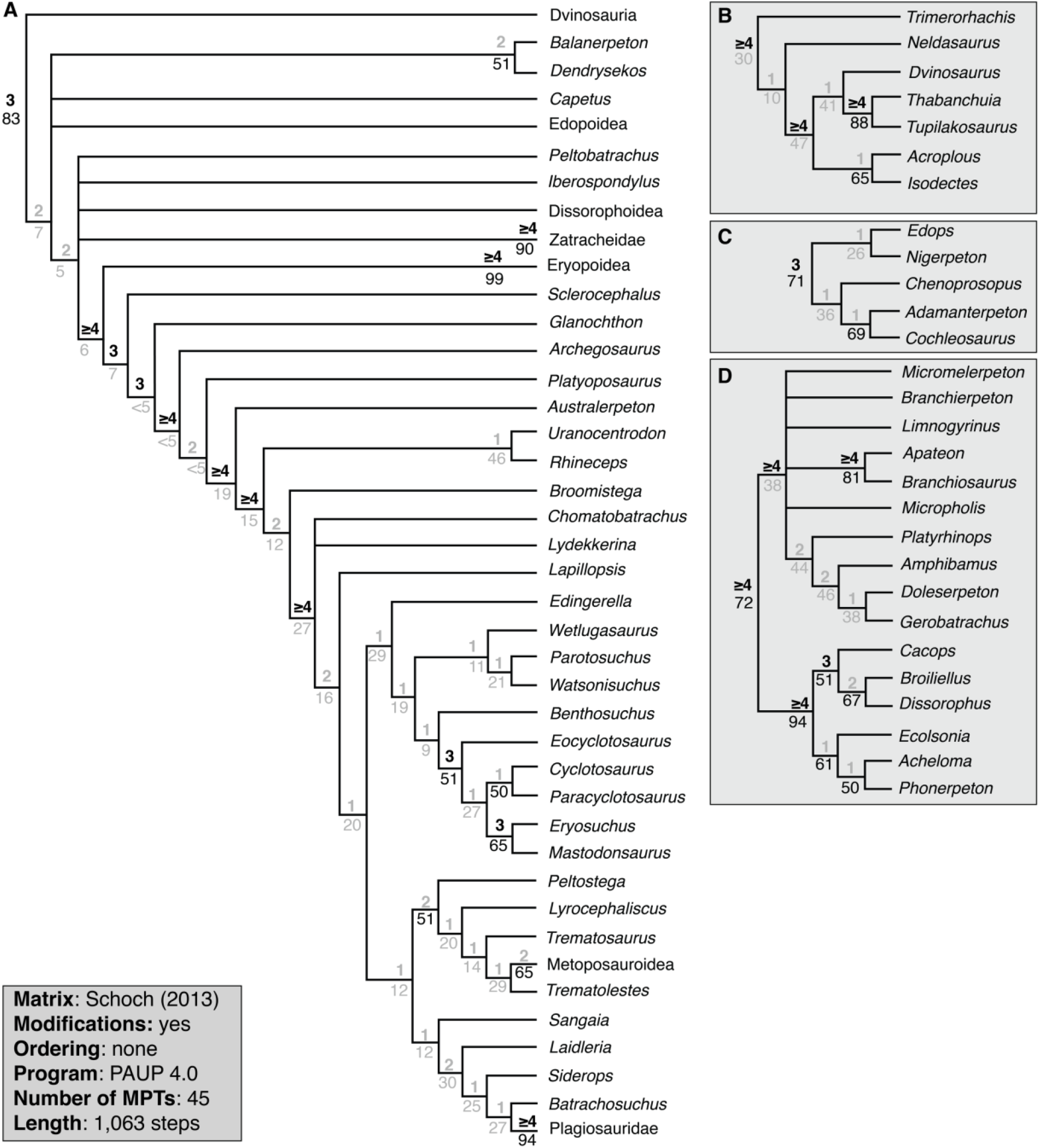
Strict consensus topology recovered from analysis of the modified Schoch (2013) matrix using PAUP*, with the full taxon sample and all multistate characters unordered. **A**, topology of Temnospondyli, with select clades collapsed for visual clarity; **B**, topology of Dvinosauria; **C**, topology of Edopoidea; **D**, topology of Dissorophoidea. Values above lines represent Bremer decay index; values below lines represent bootstrap support. Values in gray represent those below the thresholds for strong support (bootstrap ≥ 50%; Bremer index ≥ 3). Collapsed nodes representing only two species (Dendrerpetidae [*Balanerpeton*, *Dendrerpeton*], Eryopoidea [*Eryops*, *Onchiodon*], Metoposauroidea [*Callistomordax*, *Metoposaurus*], Plagiosauridae [*Gerrothorax*, *Plagiosuchus*], Rhinesuchidae [*Rhineceps*, *Uranocentrodon*], Zatracheidae [*Acanthostomatops*, *Zatrachys*]) have their support values listed at the tips and are condensed for visual clarity.

#### With select character ordering

The reanalysis in PAUP* with the restricted taxon sample and certain multistate characters ordered recovered 513 MPTs with a length of 973 steps (distributed across one tree island; CI = 0.353; RI = 0.716; CPU time = 00:39:46; Fig. 13). The equivalent reanalysis in TNT recovered 405 MPTs with a length of 882 steps (CI =0.286; RI = 0.716; CPU time = 00:02:04; Supp. Fig. 8). Two MPTs were only recovered with an additional round of branch swapping. The strict consensus is nearly identical to that recovered without any character ordering (Fig. 12; Supp. Fig. 7). Two newly formed polytomies were identified. The first is the partial collapse of Capitosauria, with *Benthosuchus*, *Watsonisuchus*, and *Parotosuchus* now recovered as single branches of a polytomy with a branch for the clade of all remaining higher stereospondyls. *Edingerella* shifted to this polytomy from its previous position as the sister taxon to the clade of brachyopoids, plagiosaurids, and trematosaurs. *Wetlugasaurus* remains the earliest diverging member of Neostereospondyli, and the remaining nominal capitosaurs form a clade as they did in the reanalysis without character ordering. The second new polytomy is a trichotomy, with branches for Metoposauridae, a clade of all remaining nominal trematosaurs (and *Peltostega*), and the clade of brachyopoids, plagiosaurids, and *Laidleria*. *Trematosaurus* and *Trematolestes* are no longer recovered as exclusive sister taxa. Statistical support is mixed, with strong Bremer support (> 2) for many nodes, including most of the historically recovered major nodes (e.g., Stereospondyli), but bootstrap support is generally weak (< 50%) for most nodes except for very distinctive clades with relatively few sampled taxa (e.g., Plagiosauridae with two sampled taxa and 93% bootstrap support). Even fewer nodes have strong support based on both metrics.

**Figure 13.**
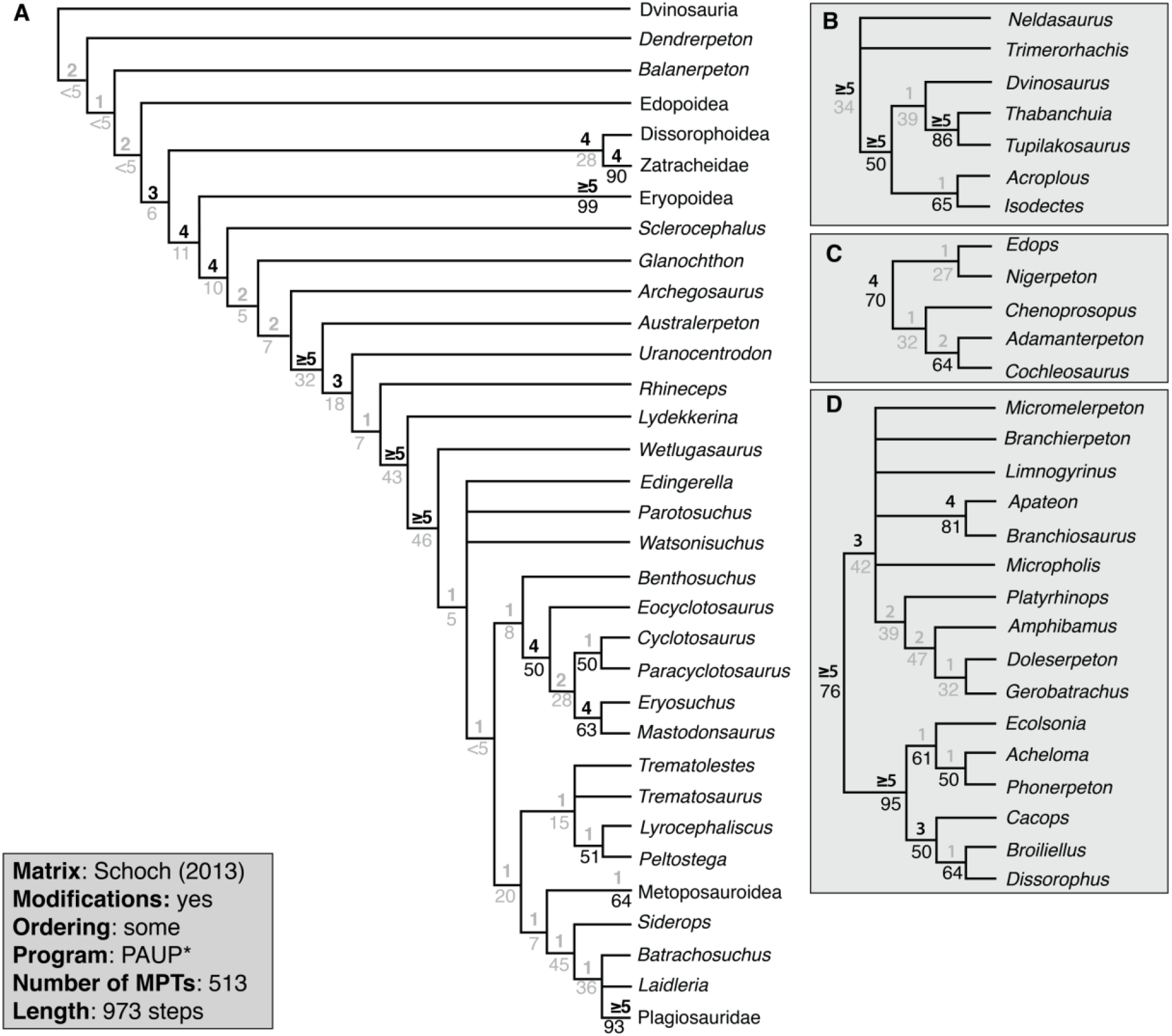
Strict consensus topology recovered from analysis of the modified Schoch (2013) matrix using PAUP*, with the restricted taxon sample and certain multistate characters ordered. **A**, topology of Temnospondyli, with select clades collapsed for visual clarity; **B**, topology of Dvinosauria; **C**, topology of Edopoidea; **D**, topology of Dissorophoidea. Values above lines represent Bremer decay index; values below lines represent bootstrap support. Values in gray represent those below the thresholds for strong support (bootstrap ≥ 50%; Bremer index ≥ 3). Collapsed nodes representing only two species (Dendrerpetidae [*Balanerpeton*, *Dendrerpeton*], Eryopoidea [*Eryops*, *Onchiodon*], Metoposauroidea [*Callistomordax*, *Metoposaurus*], Plagiosauridae [*Gerrothorax*, *Plagiosuchus*], Rhinesuchidae [*Rhineceps*, *Uranocentrodon*], Zatracheidae [*Acanthostomatops*, *Zatrachys*]) have their support values listed at the tips and are condensed for visual clarity.

With the full taxon sample and certain multistate characters ordered, PAUP* recovered 255 MPTs with a length of 1,073 steps (CI = 0.330; RI = 0.706; CPU time = 00:08:52; Fig. 14). MPTs were distributed across two tree islands (1–45, 46–255). The equivalent TNT analysis recovered 255 MPTs with a length of 976 steps (CI = 0.263; RI = 0.706; CPU time = 00:02:33; Supp. Fig. 9). One MPT was only recovered with the second round of branch swapping. The strict consensus is much less resolved than the reduced taxon sample analysis, with noticeable collapse of nodes near the base of Temnospondyli and within higher stereospondyls. All of the newly recovered polytomies involve taxa that were excluded from the reduced taxon sample. The same pattern of mixed statistical support is identified.

**Figure 14.**
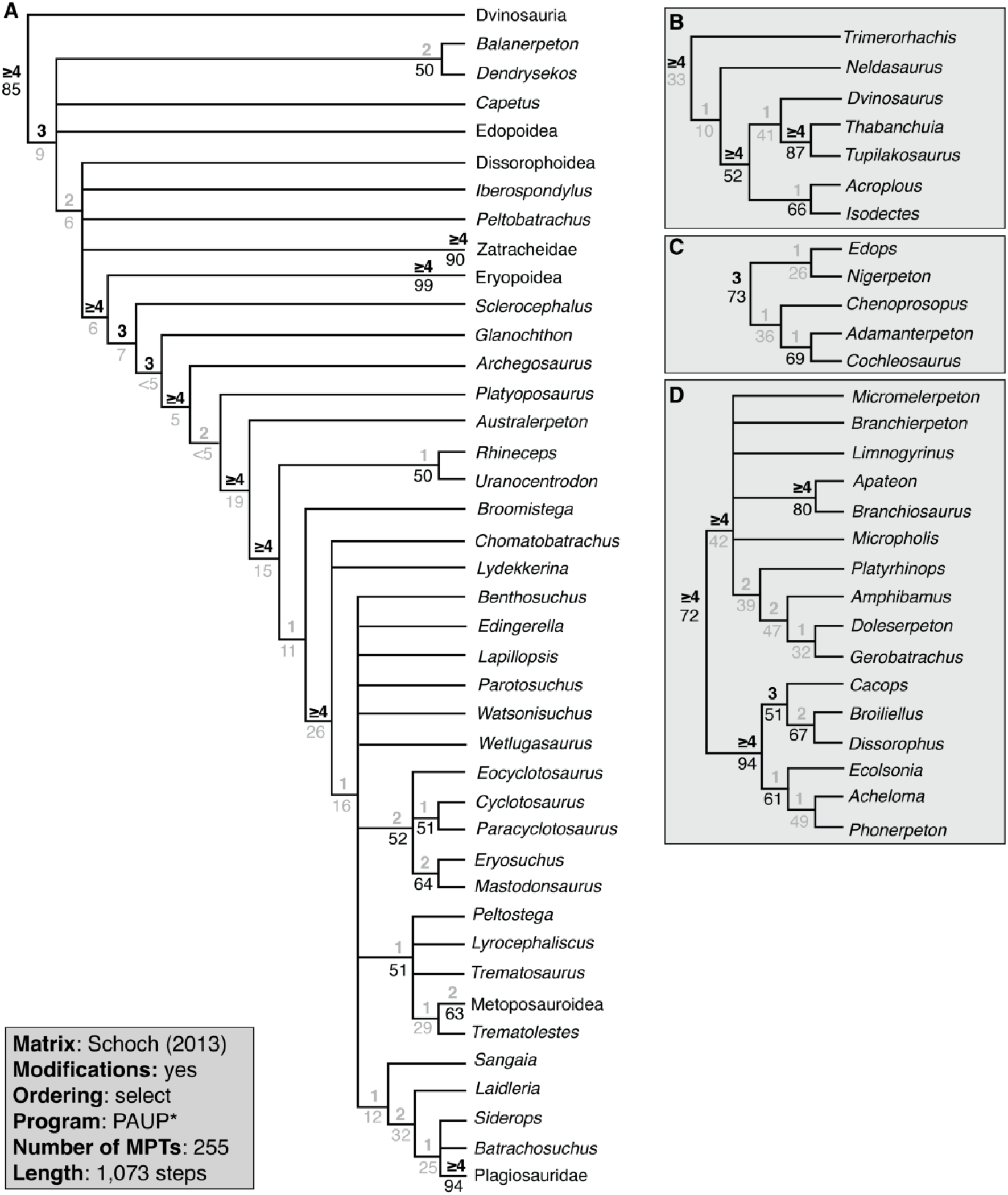
Strict consensus topology recovered from analysis of the modified Schoch (2013) matrix using PAUP*, with the full taxon sample and certain multistate characters ordered. **A**, topology of Temnospondyli, with select clades collapsed for visual clarity; **B**, topology of Dvinosauria; **C**, topology of Edopoidea; **D**, topology of Dissorophoidea. Values above lines represent Bremer decay index; values below lines represent bootstrap support. Values in gray represent those below the thresholds for strong support (bootstrap ≥ 50%; Bremer index ≥ 3). Collapsed nodes representing only two species (Dendrerpetidae [*Balanerpeton*, *Dendrerpeton*], Eryopoidea [*Eryops*, *Onchiodon*], Metoposauroidea [*Callistomordax*, *Metoposaurus*], Plagiosauridae [*Gerrothorax*, *Plagiosuchus*], Rhinesuchidae [*Rhineceps*, *Uranocentrodon*], Zatracheidae [*Acanthostomatops*, *Zatrachys*]) have their support values listed at the tips and are condensed for visual clarity.

### Reanalysis of Pardo et al. (2017)

The original matrix comprised 76 taxa (65 temnospondyls) and 345 characters for a total of 26,620 cells. The modified matrix comprises the same 76 taxa and 334 characters, not all of which are identical to those in the original matrix, for a total of 25,384 cells. Forty-one multistate characters can be reasonably inferred to occur along a morphocline (39, 67, 74–75, 103, 107, 110, 137–138, 143, 145, 158, 163, 170, 182, 187, 191, 213, 214, 226, 229, 242–243, 254, 262, 264, 266, 269, 271, 273, 279, 290, 292, 294, 298, 300–302, 304, 311, 341; all numbering follows the original scheme). Of the 334 characters, eight are constant and 18 are parsimony-uninformative. The original matrix contained 18 multistate cells (all partial uncertainty; 0.067%); 6,556 cells coded as unknown (24.6%); and 559 cells coded as inapplicable (2.01%). The revised matrix contains 119 polymorphisms and 104 cells coded as partially uncertainty (223 total multistate cells; 0.878%); 5,909 cells coded as unknown (23.2%); and 1,507 cells coded as inapplicable (5.93%).

#### Without character ordering

The original analysis in PAUP* recovered 882 MPTs with a length of 1,514 steps. The reanalysis with no characters ordered recovered 26 MPTs with a length of 1,882 steps (CI = 0.292; RI = 0.715; CPU time = 01:21:11 Fig. 15). The strict consensus of these MPTs is much more resolved than that of the original study (not reported). The new strict consensus does not differ substantially from the majority-rule consensus that Pardo et al. (2017) did report, but there is one major caveat: a monophyletic origin of Lissamphibia from within amphibamiform dissorophoids is now recovered. This result is recovered in spite of restoring several original codes for focal taxa that were changed by Schoch et al. (2020) – examples include restoring ‘lacrimal absent’ (21–1) in *Chinlestegophis* and *Rileymillerus* – and the uncoding of many cells for highly nested amphibamiforms like *Doleserpeton* and *Gerobatrachus* (e.g., treating the presence/absence of the ectopterygoid as unknown in *Doleserpeton*, rather than as absent, a feature that would be shared with lissamphibians and that was previously coded for this taxon). One other notable shift regarding Lissamphibia is that Karauridae is no longer the sister group to Anura + Caudata but instead the sister group of Caudata, which places it within Batrachia as it has historically been considered (e.g., Skutschas and Martin, 2011). Other departures from the original reported topology include the nesting of *Lapillopsis* within Stereospondyli, rather than as the sister taxon to Dissorophoidea; and the recovery of the trematosaur *Benthosuchus* as more closely related to trematosaurs than to capitosaurs (the inverse was true in previous analyses).

**Figure 15.**
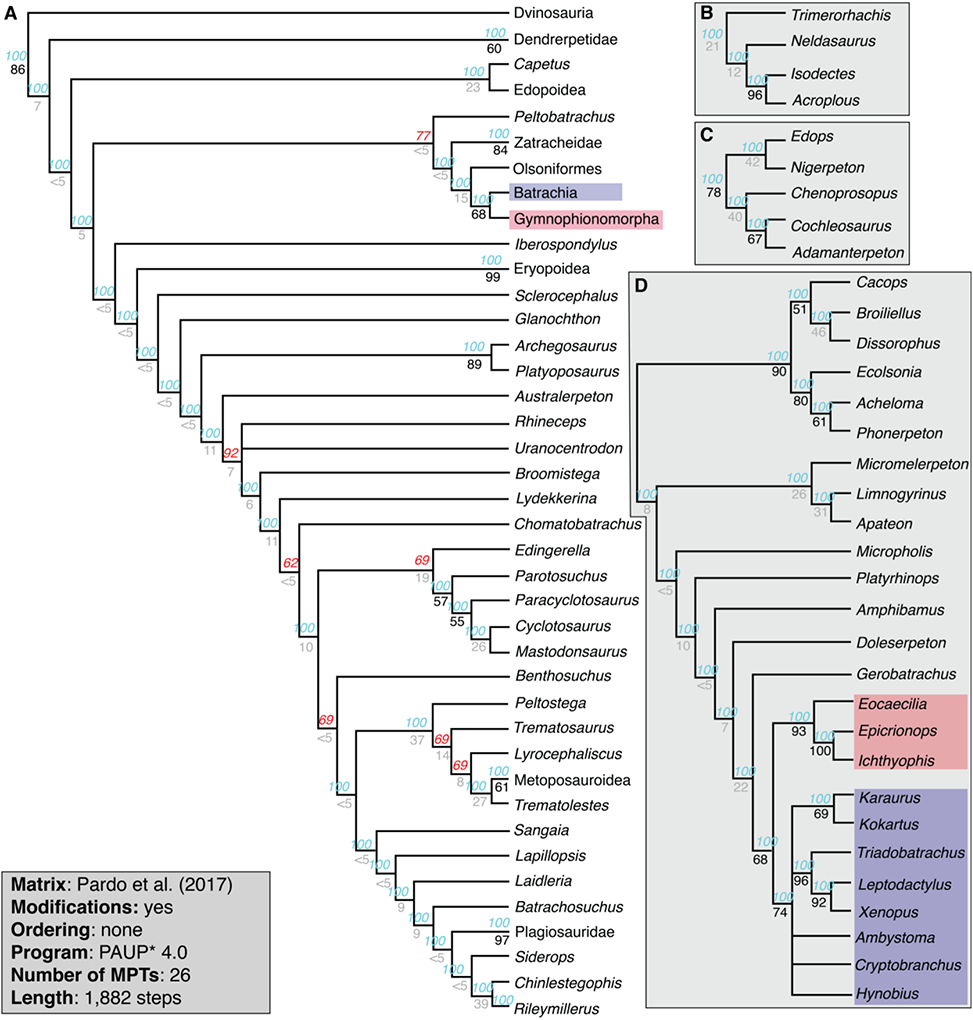
50%-majority-rule consensus topology recovered from analysis of the modified Pardo et al. (2017) matrix using PAUP*, with all multistate characters unordered. **A**, topology of Temnospondyli, with select clades collapsed for visual clarity; **B**, topology of Dvinosauria; **C**, topology of Edopoidea; **D**, topology of Dissorophoidea. Note that the majority-rule consensus is only given because this was the original consensus reported by Pardo et al. (2017); I do not endorse the use of this consensus for inferring evolutionary relationships. Values above lines represent nodal frequency, with nodes not recovered in all MPTs listed in red text; values below lines represent bootstrap support. Values in gray represent those below the thresholds for strong support (bootstrap ≥ 50%;). Collapsed nodes representing only two species (Dendrerpetidae [*Balanerpeton*, *Dendrerpeton*], Eryopoidea [*Eryops*, *Onchiodon*], Metoposauroidea [*Callistomordax*, *Metoposaurus*], Plagiosauridae [*Gerrothorax*, *Plagiosuchus*], Rhinesuchidae [*Rhineceps*, *Uranocentrodon*], Zatracheidae [*Acanthostomatops*, *Zatrachys*]) have their support values listed at the tips and are condensed for visual clarity. Colored boxes follow those of Pardo et al. for Batrachia and Gymnophionomorpha.

The equivalent Bayesian analysis (no character ordering) converged within 10 million generations (CPU time = 15:38:19), as with the original analysis. The 50%-majority-rule consensus is similar to the parsimony strict consensus, but with minor loss of resolution near the base of Temnospondyli and within higher stereospondyls. As with the parsimony analysis, a monophyletic Lissamphibia is recovered within Dissorophoidea. However, the intrarelationships of Lissamphibia differ markedly, most prominently in the paraphyly of Caudata relative to Gymnophionomorpha. Paraphyly of Dissorophidae with respect to Trematopidae is another peculiar aspect of the topology. Most nodes have strong posterior support (> 70%).

#### With select character ordering

With certain multistate characters ordered, the analysis of the modified matrix recovered 140 MPTs with a length of 1,958 steps (CI = 0.283; RI = 0.651; CPU time = 03:27:22; Fig. 17). The strict consensus is now largely unresolved, although examination of the majority-rule consensus (depicted here to be equivalent with that of Pardo et al., 2017, but not endorsed as a means of inferring relatedness) reveals that most of the nodes that were recovered in the analysis without character ordering occur in > 90% of MPTs recovered by this analysis. A monophyletic Lissamphibia is recovered in the strict consensus, but it is unplaced with respect to any particular temnospondyl clade (it nests within Dissorophoidea in 91% of MPTs).

The equivalent Bayesian analysis converged within 10 million generations (CPU time = 15:56:08). It should be noted that MrBayes cannot handle ordered characters with more than six states, so character 213 (skull length, nine states) had to be left unordered, but the 40 other characters ordered in the parsimony analysis were ordered. The 50%-majority-rule consensus tree is broadly similar to the equivalent Bayesian analysis without any character ordering, being similar in recovering polytomies near the base of Temnospondyli and within higher stereospondyls. Like that Bayesian analysis but unlike the corresponding parsimony analysis with select ordering, a monophyletic Lissamphibia nests within Dissorophoidea. Certain peculiarities like the paraphyly of Caudata are shared with the other Bayesian analysis, but others have been resolved along the lines of previous studies (e.g., the monophyly of Dissorophidae and Trematopidae is now recovered).

## DISCUSSION

### Interpretation of Unsubstantiated Codes

This study thoroughly documented extensive errors and other unsubstantiated codes in the most widely used phylogenetic matrix used to analyze the relationships of temnospondyls. The majority of these problematic codes can be traced back to the original matrix of Schoch (2013) and therefore have been propagated in every derivate. Consequently, their misleading influence extends to every subsequent study. Especially in large matrices, random typographic errors are to be expected, although this does not necessarily scale with size (e.g., discussion by Simões et al., 2017, 2018, and Laing et al., 2017). The point of this study was not to demonstrate that at least one error exists in the matrix of Schoch (2013) or its derivates. Instead, the purpose was to document clear patterns of substantial number of unequivocal errors and other unsubstantiated codes that were identified using a stringent evidentiary standard and whose correction leads to noticeably different topological results. As I argue here, these codes largely cannot be attributed to the most common sources of discrepancies: typographic errors and unpublished data. I am less interested in definitively identifying the source of the large number of errors and unsubstantiated codes than I am in the fact that it was demonstrated here that they have compromised the inference of temnospondyl relationships and related discussions about lissamphibian origins. However, I also consider it important to address possible criticisms of this study by addressing different hypotheses for the findings presented here.

A large number of unequivocal errors are philosophical in nature and relate to the ‘full dependency’ of characters. Examples include taxa coded for several secondary characters related to the lateral line grooves (original characters 34–36) and to osteoderms (original 210–211) when these taxa lack these primary features (original 33–1 and 209–1 / 209–2). Other examples include characters related to the lacrimal, to the ‘adult condition,’ to the otic notch, to anterior palatal openings, to the basioccipital, to squamation, and to intraspecific ontogeny. Notably, in many of these instances (e.g., the lateral line and osteoderm examples), taxa were uniformly erroneously coded for state 0. Although state 0 need not be the plesiomorphic condition, it appears that characters were originally constructed as such; the operational outgroup, *Proterogyrinus,* was coded for 209 of the original 212 characters, of which 208 were for state 0. Because these errors were widespread and largely uniform within a given character, they are almost certainly not typographic, nor can they be related to personal observation given the nature of the dependency. Instead, it is possible that Schoch (2013) adopted a practice in which taxa that should be treated as inapplicable were uniformly coded for the plesiomorphic state in order to more strongly differentiate the subset of taxa with an apomorphic condition. However, such an approach is neither a common practice in contemporary phylogenetics nor philosophically sound.

The more debatable coding modifications are to individual taxa for which underlying patterns are more sporadic or non-existent. It should first be acknowledged that unpublished data were almost certainly included in the original matrix of Schoch (2013), even if there was no express statement to this end. Several sampled taxa were subsequently redescribed by Schoch and colleagues (Schoch and Witzmann, 2012, 2018; Milner and Schoch, 2013; Schoch, 2018a, 2018b, 2019), and it is a reasonable hypothesis that the data in these publications was utilized by Schoch (2013). Four of the six (*Gerrothorax*, *Parotosuchus*, *Trematosaurus*, *Trimerorhachis*) exhibit relatively low rates of both overall coding changes and specific errors; most changes were related to a newly coded polymorphism from a single-state original code. The other two (*Limnogyrinus*, *Neldasaurus*) have a large number of coding changes but a relatively low error rate, a contrast that can be explained by the observation that most coding changes made here were to previously uncoded cells. *Neldasaurus* was peculiarly uncoded for a large number of postcranial characters that could be assessed from Chase’s (1965) original description. *Limnogyrinus* is thus the only OTU where a substantial number of coding changes could be linked to a recent revision (Schoch and Witzmann, 2018). Previously, the taxon was not well-characterized (and some specimens were misattributed; e.g., Milner and Sequeira, 2003), so the character coding patterns of this OTU suggest that the data published by Schoch and Witzmann (2018) were not fully known at the time of Schoch (2013).

Despite the probable incorporation of some degree of unpublished data for these taxa, subsequently published studies did not substantially revise or expand the osteology reported in previously published literature (e.g., Case, 1935; Schoch and Milner, 2000; Hellrung, 2003; Pawley, 2007; Jenkins et al., 2008) with the exception of certain reconstructions (e.g., the posterior skull margin of *Neldasaurus*) and *Limnogyrinus*. In other words, there is no evidence that character codings present in post-2013 studies that came from unpublished observations exerted a substantial influence on the coding of these taxa or that unpublished data was widely integrated and relatively influential. There is therefore no grounds for assuming that unpublished data explains taxa with higher error rates. Of the derivates of this matrix, Dilkes (2015) was the only one to expressly incorporate personal observations into coding modifications, and no other study provided a list of personally examined taxa. This underscores the point that personal observations can be more confirmatory than contradictory or revelatory. It cannot be assumed that most unsubstantiated codes are validated by unpublished data.

The argument that unpublished data are not responsible for most of the unsubstantiated codes can be further assessed by examining codes for the 10 taxa that Schoch (2013:table 1) did not personally examine as part of his study: *Australerpeton*, *Chomatobatrachus*, *Ecolsonia*, *Gerobatrachus*, *Laidleria*, *Lapillopsis*, *Nigerpeton*, *Sangaia*, *Siderops*, and *Watsonisuchus*. Five of these, *Australerpeton*, *Gerobatrachus, Lapillopsis*, *Sangaia*, and *Siderops*, were characterized by both many coding changes and a high error rate. Two of these (*Gerobatrachus*, *Siderops*) are only known by their holotypes, and another (*Sangaia*) is only known from cranial material. An example of unsubstantiated codes is those for the five characters related to ornamentation (3–7) for *Gerobatrachus*; as the holotype is only exposed in ventral view (Anderson et al., 2008), there is no basis for coding these. *Gerobatrachus* was coded the same as other amphibamiforms for all five characters. Many of these discrepancies are independent of perceptions of the validity of reconstructions for phylogenetic coding (e.g., the anterior palate of *Sangaia* has never been documented or reconstructed, yet this OTU was coded for many such characters contra Dias et al., 2006, and Dias-da-Silva and Marsicano, 2011). *Lapillopsis* is noteworthy because its high error rate is a combination of coding practices of both Schoch (2013) and Pardo et al. (2017). This OTU was substantially modified by Pardo et al., without documentation or justification, and often in perplexing fashion (e.g., treating the presence/absence of the lacrimal as unknown, contra Yates, 1999, and all subsequent workers). Many of the cells that I changed represent reversions to the original codes of Schoch (2013). Collectively, examination of these ten taxa reveals high error rates, especially for characters that could have been block coded.

A second counterpoint is the nature of the unsubstantiated codes across the entire taxon sample. Compared to codes treated as mischaracterizations (coded for one state when it should be coded for another), a majority of the coding changes implemented here relate to originally coded cells that cannot be substantiated from the literature. The inability to corroborate them refers not only to the inability to determine whether a specific condition is present (e.g., a tetradactyl manus versus a pentadactyl manus) but also to the complete lack of documentation that establishes that the requisite skeletal region is known at all (e.g., the forelimb). As noted above in the discussion of dependencies, many unsubstantiated cells were coded identically to a putative close relative (e.g., all dubious postcranial codes of *Rhineceps* were identical to those of *Uranocentrodon*), which raises questions about their reproducibility.

Finally, the magnitude of unpublished data needed to justify unsubstantiated codes for skeletal regions for which there is no documentation is very high across the entire taxon sample. This is most evident in assessing taxa known from very few specimens (e.g., *Gerobatrachus*, *Sangaia*) but can also be observed in taxa with disproportionate representation of one skeletal region (e.g., *Zatrachys* is known from many skulls, but its postcrania are unknown; Langston, 1953; Schoch, 1999; Schoch and Milner, 2014). The following are examples of presently unknown skeletal regions of various taxa that would have to both exist and to have been personally observed prior to Schoch (2013) to justify the original codes: (1) the stapes of *Proterogyrinus*; (2) branchial denticles in *Greererpeton*; (3) the ilium and scalation of *Nigerpeton* and *Zatrachys*; (4) articulated presacral and caudal sequences of *Acroplous* (sufficient to estimate total count); (5) articulated presacral and caudal sequences, the humerus, and the hindlimb of *Thabanchuia*; (6) the interclavicle, ilium, and squamation of *Chomatobatrachus*; (7) a second skull of *Siderops kehli* for which cranial sutures are preserved in their entirety; (8) the lower jaws, ribs, pectoral girdle, and scalation of either *Batrachosuchus browni* or *Bathignathus watsoni*; (9) the anterior palate, intercentra, pectoral girdle, manus, ilium, squamation, and ribs of *Sangaia*; and (10) the pectoral girdle, ilium, and squamation of *Watsonisuchus*. These examples include taxa not personally examined by Schoch (*Siderops*, *Sangaia*); span Temnospondyli and the outgroups; and encompass taxa with both dated descriptions (e.g., Watson, 1956, for the brachyopid OTU) and recent descriptions (e.g., Englehorn et al., 2008, for *Acroplous*). Greater confidence in assessing the role of unpublished data in unsubstantiated codes can be conferred when vetting non-stereospondylomorphs (and brachyopids and plagiosaurids), as the review by Schoch and Milner (2014) provided lists of known material for these taxa. This included references to explicitly unpublished material (e.g., *Tambachia trogallas*, p. 66 therein; many specimens of *Micromelerpeton credneri*, p. 44 therein). As it post-dates Schoch (2013), it can be more confidently concluded, for example, that *Bathignathus watsoni* and *Batrachosuchus browni* are only known from partial skulls (Schoch and Milner, 2014:98), which is consistent with earlier studies (e.g., Warren and Marsicano, 2000:465), and therefore that any non-cranial codes introduced for the brachyopid OTU by Schoch (2013) and probably those introduced by Pardo et al. (2017) are erroneous.

To summate, unpublished data and typographic errors are unlikely to be the primary source of errors in Schoch (2013) given the magnitude of these unsubstantiated codes and the patterns observed. At least some of the character-level patterns may be attributed to an unstated decision to code taxa for the inferred plesiomorphic state when they should be coded as inapplicable. I also want to reiterate that I applied a very stringent standard in recoding – codes had to expressly contradict the literature with respect to a condition or to whether an element is even known in order to be modified. Finally, because Schoch’s (2013) manuscript was accepted in 2011, more than a decade has transpired. If there in fact remains substantial amounts of unpublished data that corroborate many of the unsubstantiated codes of that study, the lack of timely publication presents a barrier to accessibility and reproducibility and would be problematic. Unpublished data are irreproducible in peer review and should not be afforded the same burden of proof to overturn as codes that are substantiated by the literature, which is reproducible. For a variety of reasons, other workers may not be able to reproduce one worker’s personal observations even if attempting to do so (e.g., through specimen damage or loss). Therefore, workers who are skeptical of some of my coding modifications must consider whether it is known that specimens that expressly corroborate these codes exist or whether this is merely a conjectural defense. Because of the nature of the fossil record, practically anything is ‘possible’ regarding personal observations, even for well-known taxa (e.g., newly documented polymorphism, newly documented ontogenetic stages). For this reason, it should also be considered that there are many more questionable codes that were not identified or overturned merely because of the high evidentiary standard applied here.

### What is the Best Supported Phylogeny of Temnospondyli?

Even prior to this study, it was unclear whether a defensible consensus on temnospondyl relationships had truly emerged. Other broad-scale analyses have recovered discordant results in the relationships of higher-level taxa (e.g., Milner, 1990; Yates and Warren, 2000; Pawley, 2006; Ruta et al., 2007; McHugh, 2012; Fig. 20), such as discrepancies in which clade diverged first (most often considered to be ‘Dendrerpetidae,’ Dvinosauria, or Edopoidea) and disagreement over the position of eryopoids (i.e. the Euskelia hypothesis of Yates and Warren, 2000, and McHugh, 2012, versus the Eryopiformes hypothesis of Schoch, 2013). Simply because Schoch’s (2013) topology and results have been almost universally adopted does not mean that the relationships of temnospondyls are robustly known. The accuracy of inference is not determined by its alignment with previous topologies and concepts or its relative usage compared to other studies. All studies can be accurate within the confines of their unique datasets insofar as the results are the most parsimonious or the most likely yet be discordant with other studies.

There are further discrepancies when comparing the original topology of Schoch (2013) with those of derivates (e.g., Dilkes, 2015; Pardo et al., 2017). Some of these likely relate to the inclusion of putative wildcards like *Capetus* and *Lapillopsis*, to the broader taxon sample, or to the different operational outgroups. Most significant among these is the inadvertent discovery that Schoch’s (2013) original PAUP* analysis did not recover any MPTs, and therefore, the topology that has been widely propagated is a consensus of suboptimal trees that includes unsubstantiated relationships. Whether studies that used this topology for quantitative studies would be affected by the unsupported resolution is beyond the scope of this study, but this is an inherent concern for methods that are highly sensitive to topology.

Finally, there is the issue that finer-scale relationships in broad-scale analyses are frequently discordant with large-scale analyses. Examples with respect to Schoch (2013) and derivates include the lack of a Branchiosauridae nested within the historical ‘Amphibamidae’ (contra Fröbisch and Schoch, 2009; Schoch, 2018b); the paraphyly of Amphibamidae (sensu Schoch, 2018, and derivates thereof), Lydekkerinidae (contra Dias-da-Silva and Hewison, 2013), Sclerocephalidae (contra Eltink et al., 2019; Schoch, 2021a), and Rhinesuchidae (contra Eltink et al., 2016, 2019; Marsicano et al., 2017); the polyphyly of Rhytidosteidae (contra Dias-da-Silva and Marsicano, 2011); the nesting of plagiosaurids within Brachyopoidea (contra Warren and Marsicano, 2000); the position of early diverging capitosaurs and trematosaurs like *Edingerella* and *Benthosuchus* (contra Schoch, 2018c, 2019, among others); and the placement of ‘wildcards’ like *Lapillopsis* and *Peltobatrachus*. Some of these discrepancies reflect longstanding disagreements over the placement or monophyly of various clades. Others have been proposed phenetically but never substantiated by phylogenetic analyses, such as *Peltobatrachus* as a plagiosaurid (Panchen, 1959); and brachyopids or metoposaurids as dvinosaurs (e.g., Watson, 1956; Shishkin, 1973; Coldiron, 1978; Hunt, 1993; Milner in Schoch and Milner, 2014).

### Lissamphibian Origins

The debate over lissamphibian origins remains controversial (e.g., Pardo et al., 2017; Schoch, 2018; Marjanović and Laurin, 2019; Daza et al., 2020; Santos et al., 2020; Schoch et al., 2020; Kligman et al., 2021; So et al., 2021). One of the primary challenges is that analyses are often skewed towards either lepospondyls or temnospondyls, the two historical candidates. A proper survey of the issue requires extensive expertise in both groups, which are both among the most speciose of Paleozoic clades. This conundrum has been compounded by the hypothesis of a diphyletic origin within Temnospondyli (Pardo et al., 2017; So et al., 2021), as historical debates focused specifically on the Permo-Carboniferous temnospondyls and neither addressed nor fully sampled the predominantly Mesozoic stereospondyls in the few analyses with more equal sampling of lepospondyls and non-stereospondyl temnospondyls (e.g., Marjanović and Laurin, 2019).

Here I demonstrated that the original results reported by Pardo et al. (2017) were compromised by the propagation of a large number of errors from Schoch (2013). Consequently, there is no strong evidence for their novel diphyly hypothesis, which has not been supported by subsequent independent studies (e.g., Daza et al., 2020; Santos et al., 2020; Schoch et al., 2020b, but see So et al., 2018, 2021, of which Pardo and Huttenlocker of Pardo et al. are involved). Without character ordering (as in their original study), the analysis of the modified matrix recovered a monophyletic origin from within dissorophoids (Figs. 15–16), not their novel hypothesis. With that said, I did not reanalyze the slightly modified derivate matrices of Pardo et al. that were utilized by Daza et al. (2020) or Schoch et al. (2021), so the support for the traditional temnospondyl hypothesis that those studies recovered may also be more equivocal than is presently depicted. Regarding the diphyly hypothesis, like the largely disregarded polyphyly hypothesis (e.g., Anderson et al., 2008), this novel hypothesis should also be treated with a greater degree of skepticism compared to the classic lepospondyl and temnospondyl hypotheses unless novel independent data that support it are produced.

**Figure 16.**
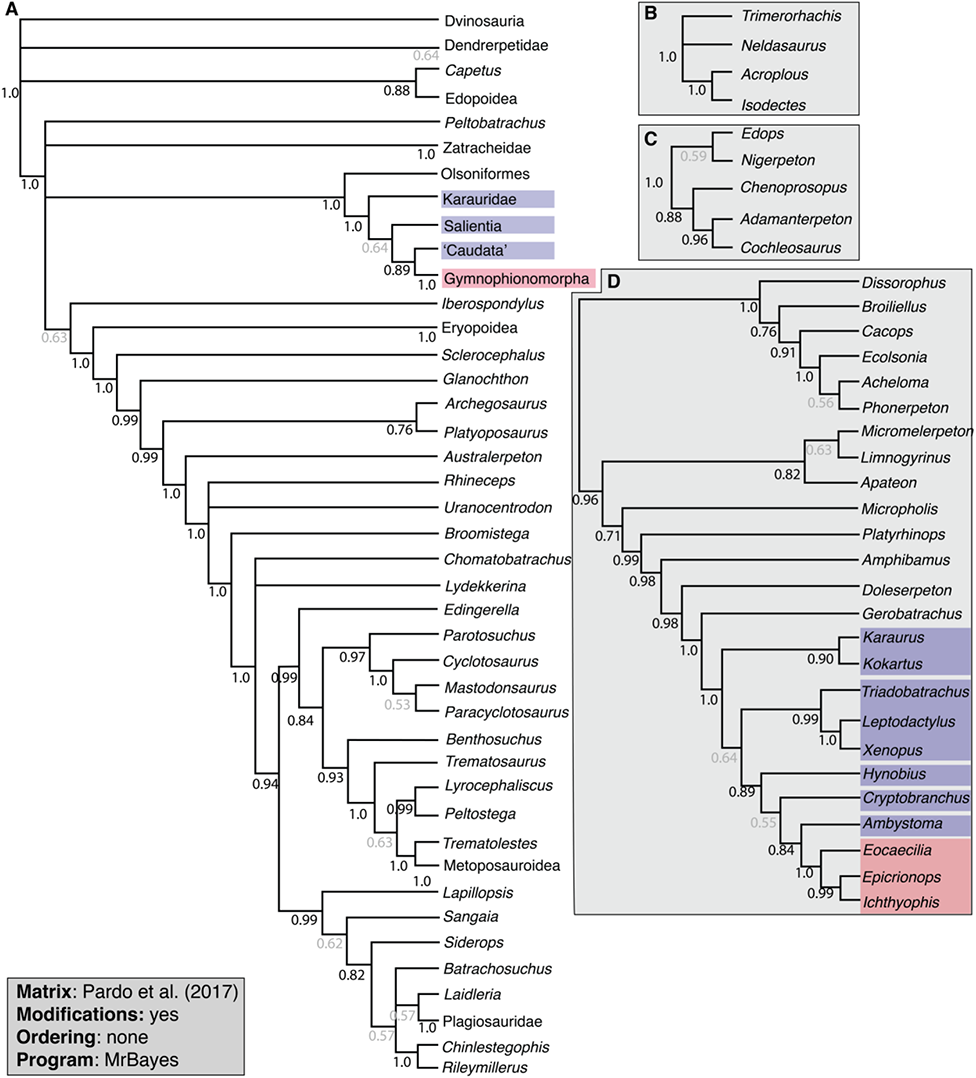
50%-majority-rule topology recovered from analysis of the modified Pardo et al. (2017) matrix using MrBayes, with all multistate characters unordered. **A**, topology of Temnospondyli, with select clades collapsed for visual clarity; **B**, topology of Dvinosauria; **C**, topology of Edopoidea; **D**, topology of Dissorophoidea. Values below lines represent posterior probabilities, with values in gray represent those below the thresholds for strong support ≥ 70%). Collapsed nodes representing only two species (Dendrerpetidae [*Balanerpeton*, *Dendrerpeton*], Eryopoidea [*Eryops*, *Onchiodon*], Metoposauroidea [*Callistomordax*, *Metoposaurus*], Plagiosauridae [*Gerrothorax*, *Plagiosuchus*], Rhinesuchidae [*Rhineceps*, *Uranocentrodon*], Zatracheidae [*Acanthostomatops*, *Zatrachys*]) have their support values listed at the tips and are condensed for visual clarity. Colored boxes follow those of Pardo et al. for Batrachia and Gymnophionomorpha.

**Figure 17.**
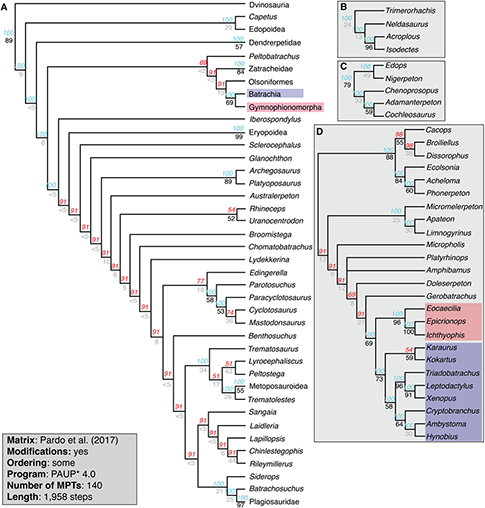
50%-majority-rule consensus topology recovered from analysis of the modified Pardo et al. (2017) matrix using PAUP*, with certain multistate characters ordered. **A**, topology of Temnospondyli, with select clades collapsed for visual clarity; **B**, topology of Dvinosauria; **C**, topology of Edopoidea; **D**, topology of Dissorophoidea. Note that the use of a majority-rule consensus is only because this was the original consensus reported by Pardo et al. (2017); I explicitly do not endorse the use of this consensus for inferring evolutionary relationships. Values above lines represent nodal frequency, with nodes not recovered in all MPTs listed in red text; values below lines represent bootstrap support. Values in gray represent those below the thresholds for strong support (bootstrap ≥ 50%; Bremer index ≥ 3). Collapsed nodes representing only two species (Dendrerpetidae [*Balanerpeton*, *Dendrerpeton*], Eryopoidea [*Eryops*, *Onchiodon*], Metoposauroidea [*Callistomordax*, *Metoposaurus*], Plagiosauridae [*Gerrothorax*, *Plagiosuchus*], Rhinesuchidae [*Rhineceps*, *Uranocentrodon*], Zatracheidae [*Acanthostomatops*, *Zatrachys*]) have their support values listed at the tips and are condensed for visual clarity. Colored boxes follow those of Pardo et al. for Batrachia and Gymnophionomorpha.

**Figure 18.**
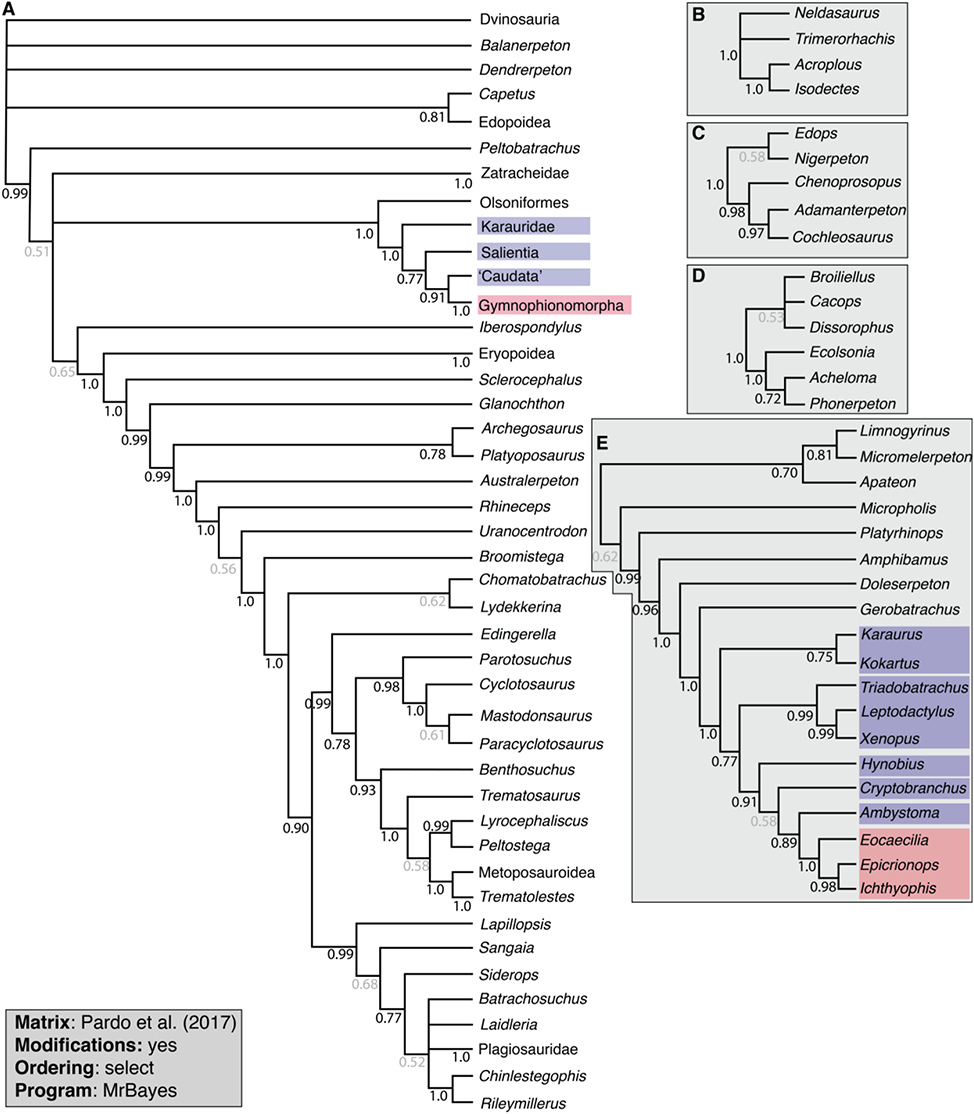
50%-majority-rule topology recovered from analysis of the modified Pardo et al. (2017) matrix using MrBayes, with certain multistate characters ordered. **A**, topology of Temnospondyli, with select clades collapsed for visual clarity; **B**, topology of Dvinosauria; **C**, topology of Edopoidea; **D**, topology of Olsoniformes; E, relationships of ‘Amphibamiformes,’ which here includes Micromelerpetidae and Lissamphiba. Dissorophoidea is divided into two subparts in this figure and not in others due to space constraints within the figure, but Olsoniformes is the sister taxon to Amphibamiformes (inclusive of Lissamphiba) as with all other analyses of this matrix. Values below lines represent posterior probabilities, with values in gray represent those below the thresholds for strong support ≥ 70%). Collapsed nodes representing only two species (Dendrerpetidae [*Balanerpeton*, *Dendrerpeton*], Eryopoidea [*Eryops*, *Onchiodon*], Metoposauroidea [*Callistomordax*, *Metoposaurus*], Plagiosauridae [*Gerrothorax*, *Plagiosuchus*], Rhinesuchidae [*Rhineceps*, *Uranocentrodon*], Zatracheidae [*Acanthostomatops*, *Zatrachys*]) have their support values listed at the tips and are condensed for visual clarity. Colored boxes follow those of Pardo et al. for Batrachia and Gymnophionomorpha.

**Figure 19.**
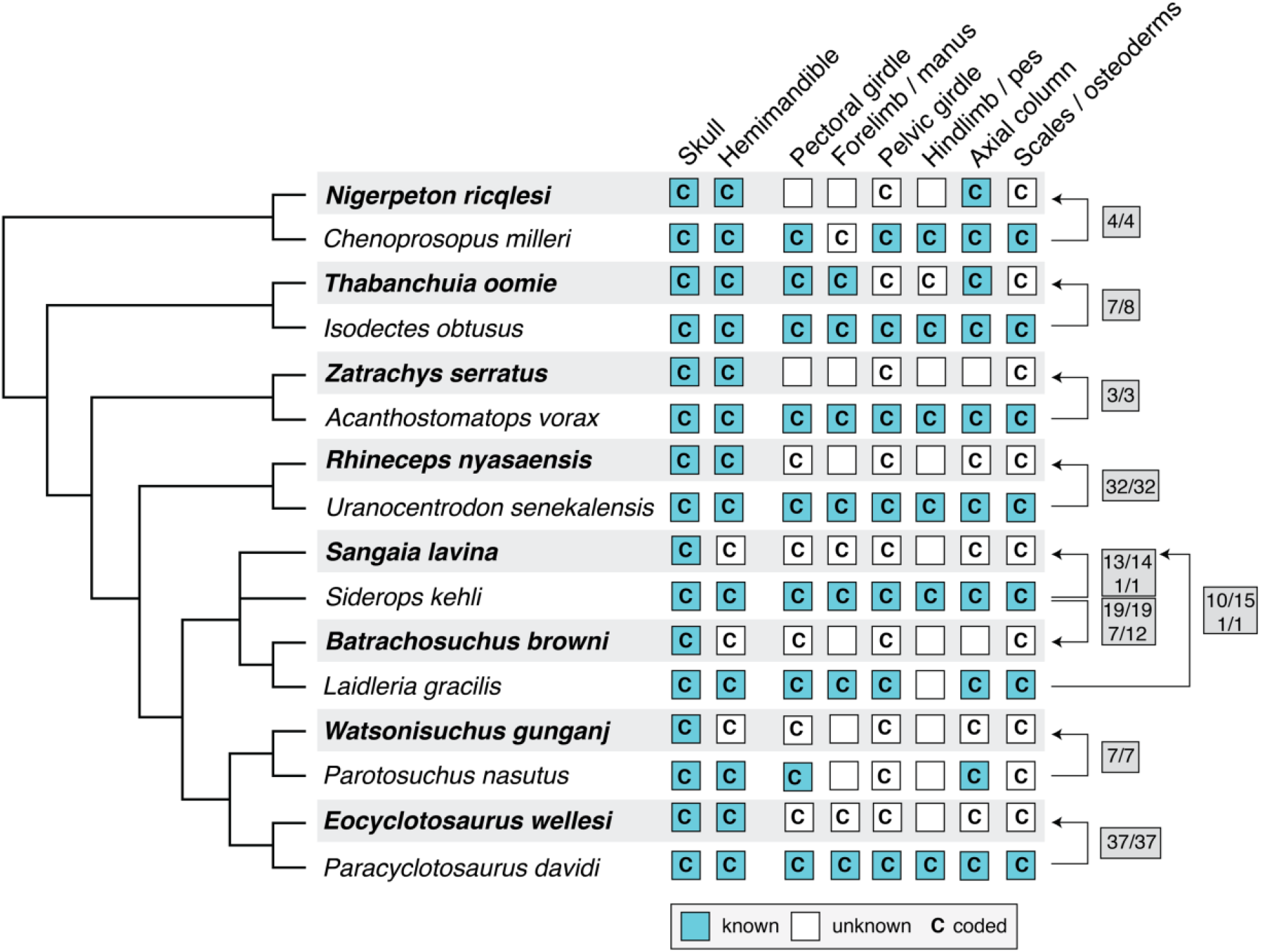
Comparison of patterns of problematic codes among Temnospondyli. Taxa listed in bold text are those with patterns of unsubstantiated codes that appear to have been duplicated from a putative close relative for which the code is substantiated; this relationship is indicated by the arrows on the right. The number before the slash (/) represents the number of problematic codes where the code is the same as the possible ‘source taxon,’ and the number after represents the total number of problematic codes for that taxon (e.g., there were eight problematic codes for *Thabanchuia oomie*, of which seven were coded the same as in *Isodectes obtusus*). For *Sangaia lavina* and *Batrachosuchus browni*, the top set of numbers is for characters originally coded by Schoch (2013), and the bottom set of numbers is for characters originally coded by Pardo et al. (2017). Characterization of a skeletal region as ‘known’ is based on the literature and includes even cursory mentions of such material (i.e. it does not refer to a more stringent standard of whether such material has been described in detail).

**Figure 20.**
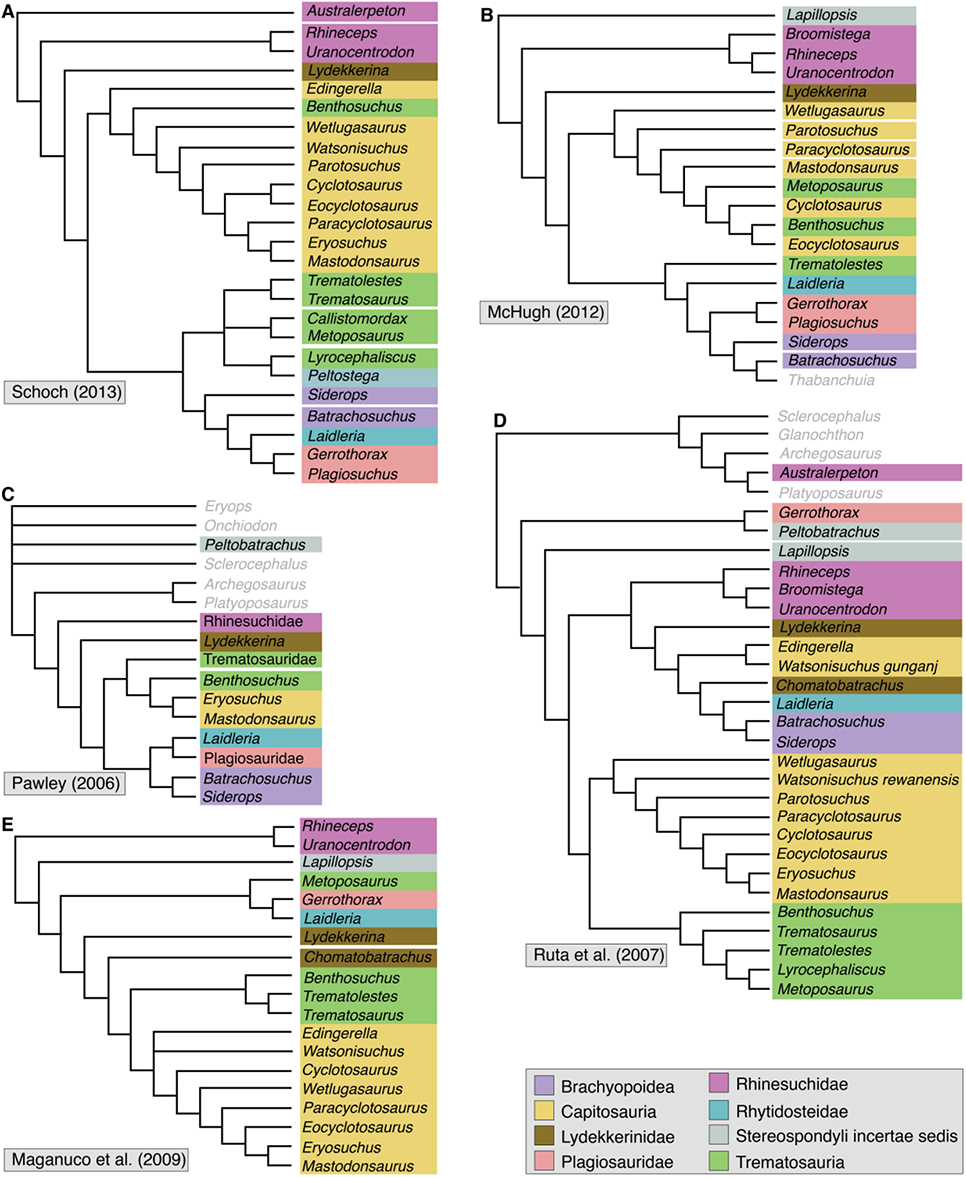
Comparison of the phylogenetic relationships and composition of Stereospondyli in recent analyses. **A**, original strict consensus of Schoch (2013); **B**, strict consensus of McHugh (2012); **C**, strict consensus of Pawley (2006); **D**, supertree of Ruta et al. (2007); **E**, strict consensus of Maganuco et al. (2009). Colored boxes delineate different clades and the present nominal placement of taxa. Taxa listed in gray text are not nominal stereospondyls. Phylogenies of studies other than Schoch (2013) have been collapsed to only depict relationships of taxa sampled by Schoch for more direct comparison.

### Future Directions

This analysis focused very narrowly on one aspect of phylogenetic analyses, namely the codes, and in a few instances, character construction. It did not address other aspects of the primary data, mainly taxon and character sampling, let alone analytical approaches other than character ordering. Substantial addition of new characters or modification to the taxon sample, as well as substantial character modifications that are not considered essential, begins to rise to the level of creating an entirely new matrix, as it requires either introduction of substantial new codes by a different worker than the original author or substantial recoding, again by a different worker. This section highlights some possible avenues for future work to address both temnospondyl relationships and the origin of lissamphibians.

#### Taxon sampling

Despite the finding that the original results of Schoch (2013) were flawed in presenting strict consensus of suboptimal trees, the relationships of temnospondyls have been generally consistent across previous derivates. This should not be taken as evidence of a consensus but rather of constraints imposed by the relative paucity of modifications made subsequent to the original study. While certain relationships align with historical concepts (e.g., a monophyletic Dissorophoidea), others are almost certainly spurious, or at least in conflict with most historical concepts (e.g., *Lapillopsis* clustering with dissorophoids rather than with stereospondyls; Yates, 1999; Yates and Sengupta, 2002; Eltink et al., 2019). Not all nine of the taxa identified as “problematic” by Schoch (2013) are clearly wildcards, either upon reanalysis of the original matrix or upon reanalysis of subsequent derivates, and it is not clear how “problematic” taxa were originally identified (e.g., with an Adams consensus or the IterPCR method of TNT; Goloboff and Szumik, 2015). A taxon cannot be “problematic” merely because it deviates from longstanding taxonomic frameworks. *Lapillopsis* can be shown to have been recovered in a spurious position on the basis of a large number of miscodings, while the colosteid *Greererpeton*, the stereospondylomorph *Platyoposaurus*, the rhinesuchid *Broomistega*, and the lydekkerinid *Chomatobatrachus* were all recovered in approximately their predicted positions (adjacent to other nominal members of their traditional clades). *Capetus* tended to cluster with edopoids, a longstanding association (e.g., Sequeira & Milner, 1993), and *Iberospondylus* tended to cluster with eryopiforms, either as their sister taxon or in a polytomy with Eryopiformes. *Peltobatrachus* and *Sangaia* remain more poorly situated, the former due to its combination of peculiar anatomy and spatiotemporal occurrence, and the latter due to the uncertainty over rhytidosteid monophyly.

The existing taxon sample is skewed towards overrepresentation of capitosaurs, dissorophoids, and dvinosaurs, for which finer-scale analyses are available that more densely sample and infer the intrarelationships of these clades (e.g., Schoch, 2008b, 2012, 2018b, 2018c, 2019; Schoch and Voigt, 2019; Gee, 2021; Marsicano et al., 2021). Schoch’s (2013) original taxon sample includes five edopoids, seven dvinosaurs, 16 dissorophoids, and nine capitosaurs, in contrast to only two brachyopoids, two eryopoids, and five trematosaurs. This representation is not proportional to the number of presently valid taxa in each clade; there are only eight species of edopoids, yet five were sampled, for example. Some of the taxa from overrepresented clades could be omitted if the overarching objective of an analysis of Temnospondyli is only elucidating the coarse-scale (family level and higher) relationships. Schoch et al. (2020) did just this, reducing their sample by 18 temnospondyls, most of which were from proportionately oversampled clades (e.g., only two edopoids and five capitosaurs were retained). The overall relationships among higher clades remained unchanged. Conversely, the taxon sample should be expanded for brachyopids (only *Batrachosuchus*), chigutisaurids (only *Siderops*), and nominal rhytidosteids (*Laidleria, Peltostega, Sangaia*) to address outstanding questions related to whether rhytidosteids form a clade (Dias-da-Silva and Marsicano, 2011; Maganuco et al., 2014) and whether plagiosaurids nest within Brachyopoidea or are the sister group to this clade.

Finally, there is the possibility that known tetrapods not presently considered to be temnospondyls might in fact be temnospondyls and have not been recognized as such because they have only been tested in relatively narrow analyses. This ambiguity can sometimes result from outdated definitions (e.g., Schoch, 2013; Marjanović and Laurin, 2019) but also owes to the paucity of temnospondyl records from the Viséan (e.g., Milner and Sequeira, 1993; Werneburg et al., 2019; Bird et al., 2020) and the disputed affinities of certain Carboniferous taxa (e.g., the “St. Louis tetrapod” of Clack et al., 2012; *Caerorhachis*, compare Holmes and Carroll, 1977, Ruta et al., 2001, and Pawley, 2006; *Doragnathus*, compare Smithson, 1980, with Clack and Milner, 2015; certain ‘lepospondyl’ nectrideans, compare Carroll et al., 1998, with Pardo, 2014), which collectively creates ambiguity regarding the earliest stages of temnospondyl evolution.

#### Character sampling

Neither the original matrix of Schoch (2013) nor any derivate captures all of the characters used in fine-scale analyses at the family or superfamily level. In some instances, such characters would be parsimony-uninformative because of limited taxon sampling, but there remain avenues to augment the existing character sample in order to test some of the aforementioned hypotheses. For example, a twisted quadrate ramus of the pterygoid is a rhytidosteid synapomorphy (Dias-da-Silva and Marsicano, 2011), but this condition is not captured in the existing matrix and could, in part, explain the lack of monophyly among the three sampled nominal rhytidosteids. The neurocranium remains a poorly sampled region, in large part because this is a historically difficult area to access and thus to code. It is hoped that additional tomographic work in temnospondyls (e.g., Robinson et al., 2005; Čerňanský et al., 2016; Arbez et al., 2017, 2022; Gee et al., 2019a; Gee, 2020a; Marsicano et al., 2021) will produce further clarity in the way that it has for ‘microsaurian’ lepospondyls (Maddin et al., 2011; Huttenlocker et al., 2013; Pardo et al., 2015; Szostakiwskyj et al., 2015; Pardo and Anderson, 2016; Gee et al., 2019b, 2020; MacDougall et al., 2021). There are also avenues for expanding certain characters that only capture generalized trends. For example, character 4 (ornamentation) only differentiates taxa with grooves (most temnospondyls) from those with pustules (e.g., *Micropholis*, *Gerrothorax*), but there are more specific ornamentation patterns that are considered to have diagnostic value (e.g., the ‘spider-web’ pattern of rhytidosteids).

#### Character construction

A final consideration within the confines of the existing character sample is the way in which these characters are constructed. This topic has been extensively covered in broad contexts in the literature (e.g., Wilkinson, 1995; Rieppel and Kearney, 2006; Sereno, 2007; Simões et al., 2017, Vogt, 2017, 2018, and references therein). I emphasize that in this study, following my approach of minimizing changes to the existing matrix, characters were not modified unless they were unequivocally problematic. As a result, there remains substantive room to refine existing characters to ensure independence or to strengthen the argument for homology, and I felt that these would be better suited for a study that actively sought to expand and to refine all characters, not merely those that are overtly problematic. Examples of characters that were modified here include some numerical characters for which the original formulation did not capture all possible conditions of sampled taxa (e.g., original character 158 for presacral count did not have any state that captured taxa with 22, 26, 27, or 28 positions). Another example is the use of excessive specifiers in character states that create conflict in whether to code a character literally or only ‘in a general sense.’ An example of the latter is given below.

Character 20 refers to the lateral margin of the nasal, and the original two states were: “(0) straight; or (1) stepped, with lateral excursion anterior to lacrimal.” State 1 could be restricted to ‘stepped,’ but the original construction meant that this character was conditional on the presence of a lacrimal as the reference point. Among taxa with a lacrimal, this may differentiate some taxa from those with a nasal that only steps anterior to the prefrontal. The original coding of this character is asymmetric, however (Fig. 21). Taxa with a stepped lateral margin that did not extend anterior to the lacrimal were coded for 20–0 (straight), such as *Lydekkerina* (Jeannot et al., 2006). Furthermore, taxa without lacrimals were also coded for this character, sometimes for 20–1 (stepped; e.g., the anurans *Xenopus* and *Leptodactylus*). Any taxon without a lacrimal that is coded for 20–1 is inherently erroneous; the nasal cannot ‘step’ in front of something that does not exist. Furthermore, because this is a binary character, there is inherently a partial dependency for any taxon without a lacrimal because only one state is viable. However, this does not mean that the margin is straight (20–0), and coding a taxon without a lacrimal for this state by process of elimination creates a false homology with a taxon with a straight lateral margin and a lacrimal. Here, the reference to the lacrimal was removed, and the character was recoded on the basis of whether the suture is longitudinally straight, obliquely straight, or stepped.

**Figure 21.**
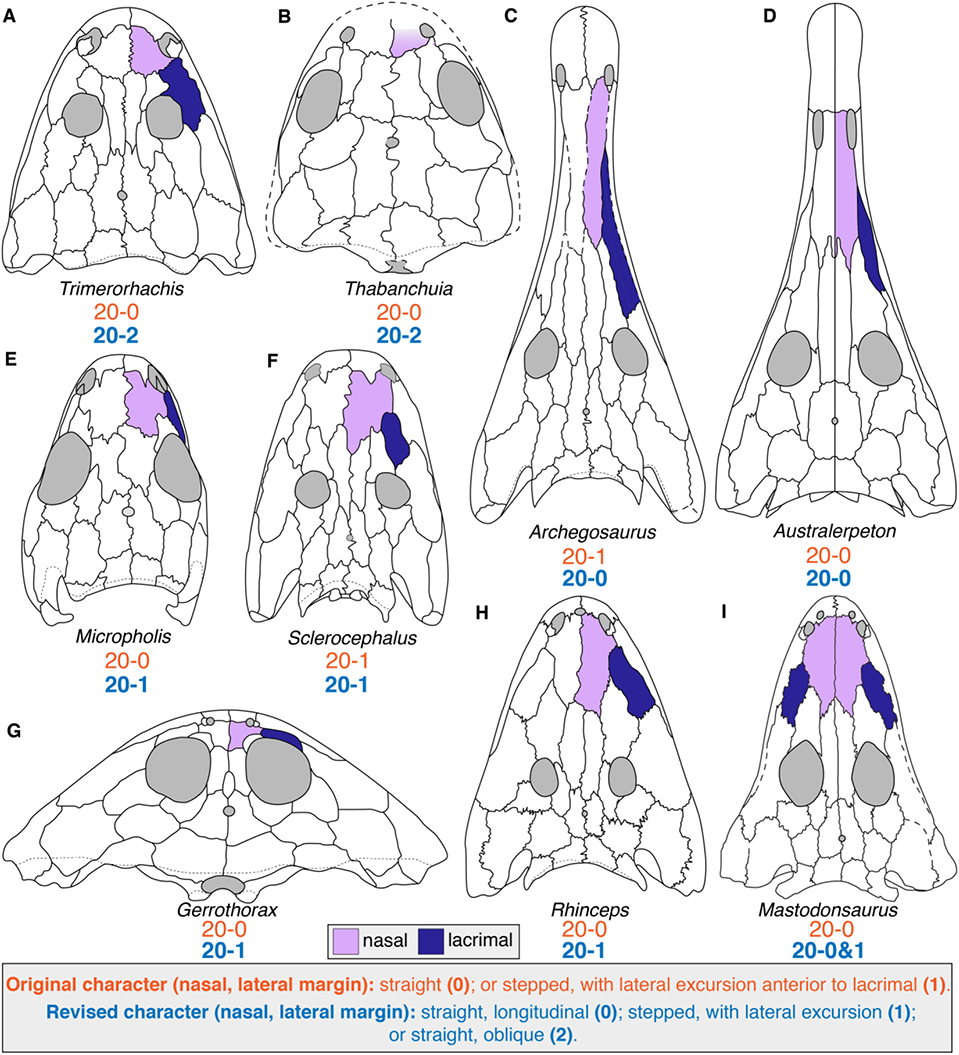
Comparison of nasal morphology among temnospondyls to demonstrate previous coding inconsistencies in the treatment of character 20 (nasal, lateral margin). **A**, the dvinosaur *Trimerorhachis insignis* (after Milner & Schoch, 2013); **B**, the dvinosaur *Thabanchuia oomie* (after Warren, 1998b); **C**, the stereospondylomorph *Archegosaurus decheni* (after Witzmann, 2005); **D**, the rhinesuchid *Australerpeton cosgriffi* (after Eltink et al., 2016); **E**, the long-headed morph of the amphibamiform *Micropholis stowi* (after Schoch & Rubidge, 2005); **F**, the stereospondylomorph *Sclerocephalus haeuseri* (after Schoch & Witzmann, 2009a); **G**, the plagiosaurid *Gerrothorax pulcherrimus* (after Schoch & Witzmann, 2012); **H**, the rhinesuchid *Rhineceps nyasaensis* (after Schoch & Milner, 2000); **I**, the capitosaur *Mastodonsaurus giganteus* (after Schoch, 1999). Note that *T. oomie* lacks a lacrimal. Orange text represent the original character and codes from Schoch (2013); blue text represent the revised character and codes. Silhouettes not to scale.

Other examples of features that are cited as reference points but that are not always present include the tabular horn, the otic notch, and the intertemporal. Inclusion of these features as specifiers creates uncertainty in whether these characters were coded literally or “in the spirit of” (ignoring accessory specifiers when they are absent). Such uncertainty creates potential for asymmetry as different working groups modify a matrix and perhaps approach coding differently. Excessive specifiers became a particular challenge with the addition of lissamphibians, which have substantial reduction and/or consolidation of discrete cranial and mandibular elements (e.g., the caecilian maxillopalatine) such that they lack distinct homologues of many temnospondyl elements or features.

Another area in which addition of lissamphibians and non-temnospondyls has created prominent conflict is in compound characters. Character 145 expressed the presence or absence of dentition on the anterior and middle coronoids (compared to character 146, which referred only to dentition on the posterior coronoid). Almost all temnospondyls have three coronoids (only one occurs in *Plagiosuchus*). While the plesiomorphic condition is three dentigerous coronoids, many stereospondyls only have dentition on the posteriormost coronoid (character 144), such that grouping the other two coronoids into one character did not create substantial conflict because nearly all taxa without teeth on the anterior coronoid also lack teeth on the middle coronoid (an exception is *Callistomordax*; Schoch, 2008a). However, in caecilians, studies have identified only one tooth-bearing element that originates lingual to the dentary in the symphysial region before co-ossifying with the dentary to form the pseudodentary; this has most commonly been homologized with the anteriormost coronoid since a tooth-bearing anterior coronoid is plesiomorphic in tetrapods. As a result, character 145 should not be coded for caecilians because there is no middle coronoid to assess, and it would be preferable to now split this character into separate ones for each coronoid (e.g., Ruta and Bolt, 2006, characters 179, 184, 187). The same issue of homology also arose for the lepospondyls added by Schoch et al. (2020b); *Brachydectes* has only one coronoid (Pardo and Anderson, 2016), while *Rhynchonkos* has two (Szostakiwskyj et al., 2015).

Finally, binning continuous data (e.g., skull length-to-width ratio) into character discrete states has long been a challenging aspect of phenotypic datasets (e.g., Archie, 1985; Felsenstein, 1988; Goldman, 1988; Chappill, 1989; Stevens, 1991; Thiele, 1993). This issue can hardly be settled here, and the direct use of continuous data should also be explored (e.g., Felsenstein, 1973; Rae, 1998; Goloboff et al., 2006; Brocklehurst et al., 2016; Romano et al., 2017; Parins-Fukuchi, 2018). The following discussion focuses on specific temnospondyl characters within the current paradigm, which favors discretization of such data into distinct states based on semi-arbitrary divisions. There are dozens of characters that invoke size (e.g., character 9: preorbital region less than twice the length of the posterior skull table or more than twice as long). If such characters continued to be used, many will have to be modified to construct more equivalent states or to ensure that all possible conditions are captured by the states. For example, character 133 refers to the size of the quadrate trochlea and had two states: (0) medial bulge only slightly larger than lateral one; or (1) being at least two times longer and twice as wide. State 0 is more ambiguous, and in this absolute binary, it must be interpreted as ‘less than twice as long and less than twice as wide’ by virtue of state 1, but this was not the original formulation and is not equivalent to the prescribed ‘only slightly larger’ definition. A related issue is the capture of ‘intermediate’ conditions. For example, character 16 refers to the internarial distance and has two states: (0) narrower than interorbital distance; or (1) wider. This character does not capture the hypothetical condition where these two distances are equal. Some other binary characters (e.g., 29, interorbital distance relative to orbit) capture the intermediate state with the use of ‘greater/less than or equal to’ language. However, which state the ‘equal to’ condition is grouped with is probably arbitrary. The primary shortcoming to splitting it into a separate state is that it is much narrower than ‘greater than’ and ‘less than,’ and as a result, risks rendering the character parsimony-uninformative if only one sampled OTU has this condition. The use of morphometric data to guide these thresholds represents one approach in addition to direct incorporation of continuous data (not feasible in all programs).

#### Backbone construction

The findings of this study pose a major challenge to workers seeking to use a phylogenetic backbone in quantitative studies. Any one of the topologies recovered here is preferable to the original topology of Schoch (2013), now demonstrated to represent a consensus of suboptimal trees. However, the new results are probably best considered only as a temporary solution because of the conservative approach taken to modifications. There are assuredly other unsubstantiated codes that can be overturned with a lower evidentiary standard (or re-examination of the relevant material), as well as avenues improve the dataset (e.g., expanded character sampling) and the analytical approach (e.g., implied-weights parsimony, well-parametrized Bayesian).

As I previously discussed (Gee, 2021), workers should avoid the supertree of Ruta et al. (2007) because of its datedness and discordance with computer-assisted non-supertree analyses across taxonomic scales. The alternative approach is the collation of topologies that were recovered from computer-assisted non-supertree analyses to form a non-computed (informal) supertree. This has frequently been employed when there is dense sampling of a relatively exclusive clade (family-level or below) and thus where fine-scale relationships (which are unlikely to resolve as well in a broader analysis) are important (e.g., Angielczyk and Ruta, 2012; Witzmann and Ruta, 2018; Pardo et al., 2019). An example for temnospondyls is given in Figure 22. Each node is not explained in detail, but the frequent use of polytomies reflects a greater lack of consensus in certain areas of the tree, such as the relationships at the base of Temnospondyli; the sister taxon of Stereospondylomorpha; the monophyly of Rhytidosteidae; and the composition of Brachyopoidea (Milner, 1990; Warren, 1998a; Warren & Marsicano, 2000; Yates and Warren, 2000; Pawley, 2006; Ruta and Bolt, 2006; Dias-da-Silva and Marsicano, 2011; McHugh, 2012; Schoch, 2013; Maganuco et al., 2014; Marjanović and Laurin, 2019). Many wildcards (e.g., *Peltobatrachus*) also remain poorly resolved and thus were placed in relatively basal positions here.

**Figure 22.**
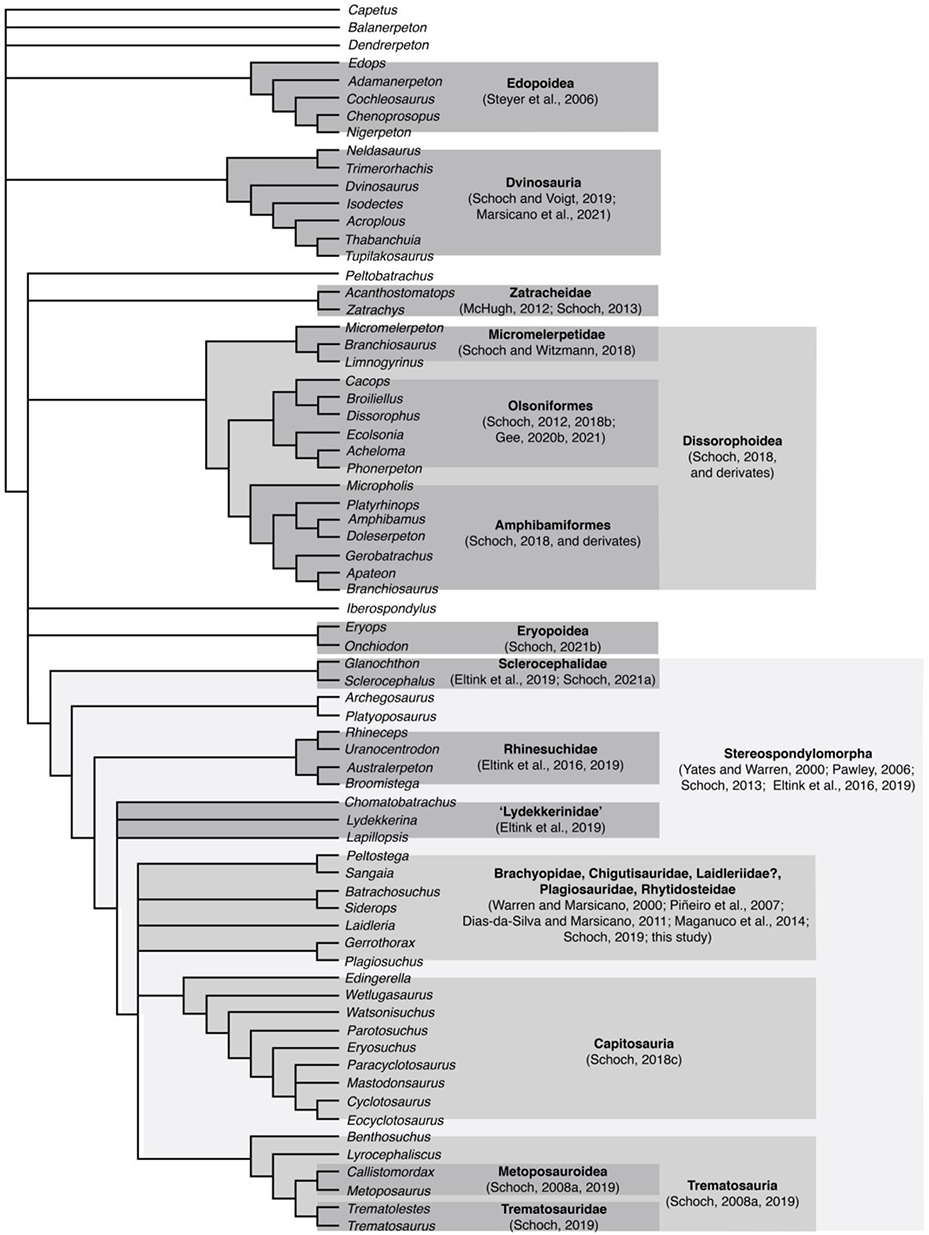
Example of a non-computed supertree compiled by collating topologies from recent computer-assisted phylogenetic analyses of different temnospondyl subclades.

## CONCLUSIONS

1. The matrix of Schoch (2013) contains a large number of unsubstantiated codes that have been propagated into subsequent derivates, including a number of high-profile publications addressing the longstanding question of lissamphibian origins (Pardo et al., 2017; Daza et al., 2020; Schoch et al., 2020b). These codes were identified using a high evidentiary standard – the literature had to expressly contradict the existing code or expressly indicate that a coded skeletal region is not in fact known.

a. It is possible that some of the discrepancies relate to personal observations. However, given that there have been minimal scoring changes made to this matrix’s lineage, it can be inferred that this is a not a major source of discrepancies, and given the time elapsed since the original matrix was accepted in its final form (November 2011), any such personal observations have not been entered into the official scientific record for over a decade and thus are inaccessible to other workers. Such personal observations would have to include large amounts of novel data, in particular for taxa for which certain skeletal regions are entirely unknown (e.g., postcrania of *Zatrachys serratus*), another line of evidence that unpublished data are not a substantial source of discrepancies.
b. Many unsubstantiated codes are unequivocal errors related to non-homology and dependency. For example, taxa cannot be coded for a character differentiating ‘narrow’ and ‘wide’ osteoderms when they are coded as lacking osteoderms at the same time. This falsely homologizes taxa that do have narrow osteoderms with taxa lacking osteoderms but that were coded as having narrow osteoderms. As a second example, taxa should not be coded for characters where one state requires the presence of the ectopterygoid when they are coded as lacking an ectopterygoid. Absence of secondary features like an ectopterygoid entering the interpterygoid vacuity or ectopterygoid dentition are inherently linked to the primary absence of this element. Scoring them as such also produces unequivocal non-homology.
c. Although the source of errors may be speculated on, the most important point is that these errors exist and that their influence has yielded results that are not reproducible or supported. Future workers should carefully scrutinize this matrix for further areas that require redress.
2. Reanalysis of the original Schoch (2013) matrix without modifications recovered substantially more MPTs of a distinctly shorter length than originally reported and that produce a different consensus topology. The original topology, which has been widely adopted, is notably more resolved than what is actually the presently most parsimonious solution. Therefore, even workers who are skeptical of some or all of the scoring revisions or even the merit of this study should be aware that this original topology does not accurately reflect the results of the utilized parsimony criteria.
3. Reanalysis of the Schoch (2013) and Pardo et al. (2017) matrices with their respective modifications recovered different topologies than the original analyses in all iterations. This includes newly recovered positions of putative wildcard taxa that are more congruent with their phenetic and/or historical placement (e.g., *Lapillopsis nana* nesting at the base of Stereospondyli rather than clustering with dissorophoids). The diphyly hypothesis of lissamphibian origins of Pardo et al. (2017) is not supported upon reanalysis and should be downweighted in discussions of lissamphibian origins unless new data to support it are produced.
4. The phylogeny of temnospondyls is poorly resolved. This provides an impetus for renewed efforts to construct novel characters across a broad sample of temnospondyl species. Additionally, temnospondyl workers should explore alternative methods, including implied weights parsimony and Bayesian inference, which are less susceptible to homoplasy than traditional parsimony.
5. Workers who are not seeking to test the relationships of temnospondyls but who require a topology for a quantitative or qualitative backbone should expressly state their justifications for utilizing a given topology or set of topologies. Pardo et al. (2019) have suggested that the topologies of focused phylogenetic studies at more exclusive levels be collated into non-computed supertrees based on a generalized concept of the relationships of higher taxa within the focal clade, with areas of instability intentionally left as polytomies. I believe that this method is the best approach for the immediate future until further testing, preferably of independent matrices, can be conducted.

## Supporting information

Phylogenetic datasets

## ACKNOWLEDGMENTS

I thank the following people for many valuable discussions on various temnospondyls and matters related to systematics: J. Anderson, C. Beightol, A. Huttenlocker, H. Maddin, A. Kufner, A. Mann, D. Marjanović, A. Marsh, A. Milner, J. Pardo, W. Parker, C. Sidor, and C. So. Thanks to C. Sidor for looking over an earlier draft of this manuscript. TNT is freely provided by the Willi Hennig Society. This study was supported by NSF ANT-1947094 (to C. Sidor).

