## Supplementary material for "The disadvantage of derivation: conserved systematic flaws in primary data have repeatedly biased the phylogenetic inference of Temnospondyli (Tetrapoda, Amphibia)": Phylogenetic datasets: README.rtf

Author: Bryan Gee
Currently reposited at: bioRxiv

CONTENTS

* 1. Introduction
* 2. Matrices
* 3. Analytical methods
	* 3.1. PAUP* settings
	* 3.2. TNT settings
	* 3.3. MrBayes settings
* 4. Most parsimonious trees (MPTs)

———

1. Introduction

This README file contains information on how phylogenetic matrices were constructed and analyzed for this study as well as how to locate the relevant MPTs that were output from the parsimony analyses. Compared to the main text, this file goes into slightly greater detail about parameters, being explicit about which defaults were used (as the default may change over the development of a program or can be manually set to a different setting by a user).

2. Matrices

Matrices were sourced from the original literature (a Word document of text strings for Schoch, 2013; the text of a NEXUS script for Pardo et al., 2017). Because neither was provided as a direct NEXUS file, both are provided as NEXUS files here. Matrices were edited in Mesquite version 3.6 (build 917) (https://www.mesquiteproject.org/). Two base matrices are provided: a modified version of Schoch’s (2013) matrix and a modified version of Pardo et al.’s (2017) matrix. Two different forms of each matrix are given. One contains only the characters that were analyzed, and thus the numbering scheme follows the revised scheme given in the appendices. The other contains characters that were omitted such that the numbering scheme follows the original numbering scheme; this one is most useful for comparing directly to the appendices when referencing specific character changes. Note that these were the versions used to conduct the analysis. These characters have ‘DELETE’ appended in front of their original character name, have their original states omitted, and have been set to be excluded and will be automatically set as such when reading in the NEXUS file (or a derivative from the file). The filenames are appended with the number of characters in each (207 vs. 212 for Schoch; 334 vs. 345 for Pardo et al.). 
	Character ordering, as specified in the main text and the appendices, is also set in the NEXUS files. Character partitions were created strictly for visual organizational purposes and are neither based on a previous study nor intended for specific implementation in this study; excluded characters were not assigned to any partition. Annotations indicate the original code and the new code (e.g., ‘0 -> 1’ indicates the original code was ‘0’ and the new code is ‘1.’ The reader is referred to the appendices for details on these changes.

TNT files (.tnt) were generated by exporting the NEXUS file from Mesquite. I did not convert gaps to missing (default) and used the default option of ‘Windows (CR+LF)’ for the End of line character.

The NEXUS file specifically for MrBayes was also exported from Mesquite. I simplified names as required for MrBayes (default).

3. Analytical methods

These analyses were run on a Mid 2015 MaBook Pro with a 2.5 Ghz Intel Core i7, 16 GB of RAM, and macOS Mojave version 10.14.6.

—The version of PAUP* that was utilized is version 4.0a (build 169) for Macintosh (http://phylosolutions.com/paup-test/); the incompatibility of the GUI version of PAUP* with MacOS 10.15+ is the primary reason why version 10.14 was maintained. The version of TNT that was utilized is version 1.5 (http://www.lillo.org.ar/phylogeny/tnt/), as made freely available by the Willi Hennig Society. The GUI version of the program, only available for Windows, was run through Wine (https://www.winehq.org/). The version of MrBayes that was utilized is version 3.2.7a (http://nbisweden.github.io/MrBayes/download.html).

3.1. PAUP* settings
—Maxtrees were set at 1,000,000 and to automatically increase by 1,000,000.
—The default Rooting option of ‘Outgroup’ and ‘Root tree at internal node with basal polygamy’ we’re applied.
—Parsimony searches utilized the following defaults: collapsing branches when maximum length is zero (Rule 3); ACCTRAN character optimization; allowing assignment of states not observed in terminal taxa to internal nodes (“short-cuts”) and only those states that canoe identified as partial shortcuts by the “3+1” test; use of minimum-possible single-character lengths as the “minimum” values for calculating CI, RI, and RC; and treatment of “gap” states as “missing data.” The one default that was not employed here relates to treatment of taxa coded as having multiple states (default is polymorphism); I treated these as variable (respecting “()” versus “{}”).
—Heuristic searches utilized the following defaults: using stepwise addition to get starting trees for branch-swapping; swapping only on the best trees when multiple starting trees exist; 10,000 random addition sequence replicates; TBR with a reconnection limit = 8, swapping only on the best trees. The one default that was not employed here is holding 10 trees per step rather than the default of 1.

3.2. TNT settings
—Max trees were set at 99,999 with all other memory default parameters employed (e.g., 500 Mb of general RAM)
—Tree collapsing was done when minimum length = 0 (Rule 1)
—When consensing, taxa missing from some trees were excluded, and branches unsupported under the selected collapsing criterion were temporarily collapsed
—Parsimony searches utilized the following defaults: starting on Wagner trees for the initial search, with random seed = 1; TBR; 10,000 replications, saving 10 trees per replication. The second round of branch-swapping was then run using trees from RAM as the starting trees.
—Bootstrapping was performed using standard bootstrap resampling, outputting absolute frequencies, with 10,000 replicates and a traditional search, using groups from the strict consensus to map the output. 
—Bremer support was calculated by iteratively searching for trees one step longer using the line ‘hold X; sub Y; bbreak = TBR;” where ‘X’ is the total number of trees to hold and ‘Y’ is the number of steps beyond the shortest trees to search for (e.g., Y = 2 means the search holds all MPTs and all suboptimal trees of 1 or 2 steps longer). The command ’bsupport’ then collates the results of these iterative searches to output Bremer support on the strict consensus tree.  

3.3. MrBayes settings
—The MrBayes analysis largely followed the settings that were automatically exported from Mesquite (e.g., ‘set autoclose=yes’; ‘lset nst=1’; rates=gamma). The MCMC analysis was run for the default of 10,000,000 generations with four chains, using the default burn-in of 0.25 and the default sample frequency of 1,000 generations. The only default that was modified was the print frequency. MrBayes is set to terminate the analysis after completing the specified number of generations unless the Average Standard Deviation of Split Frequencies (ASDSF) exceeds 0.1, in which case it prompts the user whether to continue the analysis; the ASDSF dropped well below 0.1 for all analyses here and were thus terminated after the 10 million generations. The analysis was then summarized using the ‘sumt’ command.

4. Most parsimonious trees (MPTs)

The most parsimonious trees (MPTs) from the PAUP* and TNT analyses are provided here in nested folders within a single ZIP file. They are named using the following format:

Author name_modifications_taxa_ordering_number of MPTs x MPT length.

—Author name is either ‘Schoch2013’ (for Schoch, 2013) or ‘Pardo2017’ (for Pardo et al., 2017).
—Modifications are either ‘original’ (only applicable for the Schoch, 2013, reanalyses) or ‘modified’
—Taxa is only applicable to the reanalyses of Schoch (2013) and is either ‘63taxa’ (restricted taxon sample) or ’72taxa’ (full taxon sample).
—Ordering is ‘noordering’ (no multistate characters ordered), 'selectordering' (certain multistate characters ordered), ‘allordered’ (all multistate characters ordered; only applicable for the Schoch, 2013, reanalyses), or ’67ordered’ (only original character 67 ordered; only applicable for the Schoch, 2013, reanalyses).

For example, “Pardo_modified_noordering_26x1882.nex” refers to the reanalysis of the modified matrix of Pardo et al. (2017), with all multistate characters unordered and which recovered 26 MPTs of a length of 1,882 steps.

[END of document]
