## Supplemental appendices for "The disadvantage of derivation: conserved systematic flaws in primary data have repeatedly biased the phylogenetic inference of Temnospondyli (Tetrapoda, Amphibia)"

#### Appendix 1. Overview of matrices.

The revised matrices analyzed here are, in the strictest sense, revised versions of those employed by Schoch (2013) and Pardo et al. (2017a), but they represent the collation of work conducted as part of Kligman et al. (2021; on a putative Triassic stem caecilian) and Gee et al. (in prep; on a novel lapillopsid-like taxon). These studies revised the most recently published derivate in this family of matrices (Schoch et al., 2020). As a result, revisions are 'top-down' – that is to say, they began with the most recent versions of this family of matrices, even when producing revised versions of older matrices. Therefore, coding modifications made to intermediate derivates that were subsequently propagated (e.g., those of Dilkes, 2015, that were incorporated into Pardo et al., 2017) were accepted unless expressly stated otherwise (see Appendix 3 for more).

The Schoch (2013) matrix is the original in this family of matrices and represents the collation of smaller-scale analyses of various temnospondyl clades (e.g., Schoch & Rubidge,

2005; Schoch, 2008a, 2008b). It sampled 72 OTUs, 70 of which were temnospondyls (although eight were excluded from the final analysis for which results are discussed and figured in detail) for 212 characters.

The matrix of Pardo et al. (2017a) is the more derived of the two reanalyzed here and represents a second-order derivate; although not stated as such in that study, the majority of coding modifications (but not character additions) made by Dilkes (2015), a first-order derivate, were incorporated. This was noted by Marjanović & Laurin (2019:116) and independently verified here. Furthermore, there was coding modifications novel to Pardo et al. (2017a) that were also not documented or identified by previous workers (see Appendix 3.2). The most notable modifications made by Pardo et al. were the addition of 133 new characters and 11 OTUs, including extant and extinct lissamphibians, for a total of 76 OTUs sampled for 345 characters. Pardo et al. excluded nine temnospondyls that were originally sampled by Schoch (*Dvinosaurus primus, Thabanchuia oomie, Tupilakosaurus wetlugensis, Branchierpeton amblystomum, Branchiosaurus salamandroides, Watsonisuchus gunganj*, *Wetlugasaurus angustifrons*, *Eryosuchus garjainovi*, *Eocyclotosaurus wellesi*), but they also retained several of the "wildcard" taxa excluded by Schoch in most of the latter's analyses (e.g., *Lapillopsis nana*, *Peltobatrachus pustulatus*) and added *Chinlestegophis jenkinsi* and *Rileymillerus cosgriffi* to produce a total of 65 temnospondyls.

### Appendix 2. Coding modifications

#### Appendix 2.1. Recoding philosophy and methods.

Many codes in previous matrices cannot be replicated from the literature alone (i.e., they are either erroneous or are predicated entirely on personal observations). While such discrepancies may be most apparent for taxa with dated descriptions, such as *Cyclotosaurus robustus*, *Metoposaurus diagnosticus*, and *Uranocentrodon senekalensis*, questionable codes are hardly exclusive to such taxa. **In determining whether to adjust an existing code, I opted for a conservative approach and applied the highest evidentiary standard ('beyond a reasonable doubt') in assessing codes. Codes were adjusted only if the literature expressly contradicted the previous code, either by documenting a different condition or by indicating that a given skeletal region is unknown in a given taxon.** If a feature or condition was not clearly documented in the literature but was previously coded, I kept that coding provided that the literature established that the general region is known for the taxon. An example is given below.

The trematopid *Phonerpeton pricei* was coded for five characters related to the ilium (198–202). The original (and only) descriptions of this taxon by Olson (1941:153–155; as "*Acheloma pricei*") and Dilkes (1990:234–237) make no mention of the ilium, nor do they provide a description or a figure. This taxon was coded identically to other olsoniform dissorophoids in which the ilium is sufficiently known to code these five characters, and it could be reasonably inferred (though it should not be coded solely on this basis) that *Phonerpeton pricei* had the same apomorphic conditions by virtue of its olsoniform affinities. However, because Dilkes (1990:224) listed one specimen as consisting of "fragments of pectoral and pelvic girdles," I did not change these codes because I are unable to conclude 'beyond a reasonable doubt' from the literature that the ilium is wholly unknown, even if there is also no express confirmation of any part of the ilium being known. This is not the most conservative approach from the perspective of *coding* (which would be to code it as unknown for these characters), but it is the most conservative approach from the perspective of *recoding* (which gives the benefit of the doubt to previous workers if the literature does not unequivocally contradict the codes). I opted for this approach to make the reanalyses as comparable as possible to previous iterations of this matrix, but there are a substantial number of unmodified codes that exist in this 'gray area' of being neither confidently substantiated nor invalidated by the literature, which will need to be addressed going forward. It is important to be transparent that codes for certain characters were more likely to be modified (e.g., those with counts or presence/absence binaries) compared to those that are more ambiguous or undoubtedly subjective (e.g., 'long' vs. 'short'). By the same token, codes taxa represented by only a few, or only one, specimens were more likely to be confidently recoded.

The same burden of evidence was applied when considering whether to code previously uncoded cells. For example, the 'dendrerpetid' *Dendrerpeton helogenes* was coded as unknown for dentigerous palatal ossicles (character 139), but Godfrey et al. (1987:800, fig. 1D) identified numerous small plates between the mandibles. They called these "gular" osteoderms in their figure 1D (character 149) but compared them to similar features in the holotype of the amphibamid *Platyrhinops lyelli* that were identified as by Carroll (1964:231) as "denticulate skin covering" and as dentigerous ossicles by Clack & Milner (2010:289), and the position is more in line with palatal plates rather than with gular osteoderms. On one hand, Godfrey et al. (1987) did not identify ornamentation on these plates that would provide positive evidence that they are osteoderms, but they also did not identify denticles that would provide positive evidence of a dentigerous palatal feature. Consequently, I did not code either character 139 or 149 for this taxon, but other workers may consider a lower evidentiary standard (e.g., 'more likely than not') to be sufficient to code either character. This approach applies also to entire characters that may require a high fidelity of preservational quality. For example, a narial flange is considered to be a dissorophoid synapomorphy and is coded as absent in all other sampled temnospondyls, even though most taxa do not appear to be preserved or prepared in a fashion that would actually permit assessment of the ventral surface of the skull roof medial to the naris (e.g., in dorsoventrally compressed specimens from coal deposits).

I treated characters literally. For example, a number of characters are compound in nature, presumably because they frequently co-occur. An example is character 129 (palatine, ectopterygoid [width]). This refers to the width of both palatal elements, which is greatly reduced in many amphibamiforms (129–1) and which has been qualitatively cited as a shared feature supporting lissamphibian affinities (e.g., Schoch, 2019b). However, the construction of the character clearly specifies both the palatine *and* the ectopterygoid, not one or the other. Therefore, any taxon without a palatine or an ectopterygoid (or both) cannot be coded for this character. Other examples include character 50 (postfrontal and postorbital either shorter than the supratemporal and the parietal or longer) and character 148 (anterior and middle coronoid dentition present or absent). Issues with compound characters have primarily arisen as a result of the addition of lissamphibians by Pardo et al. (2017a), as these taxa have far fewer cranial elements than temnospondyls. This approach to interpretation of characters was superseded only by philosophical problems (e.g., non-homology, see 'Terminology' [Section 2.2] below).

In line with my conservative recoding approach, I also did not modify characters unless I felt that doing so was essential for avoiding redundancies or overt dependencies, or if the existing character clearly failed to capture the full range of variation. For example, character 158 (presacral count) originally had three states: more than 28; 23–25; and fewer than 21. This character did not capture taxa with presacral counts of 21, 22, 26, 27, or 28 positions (which are documented in some sampled taxa, e.g., *Callistomordax kugleri*, *Paracyclotosaurus davidi*;

Watson, 1958; Schoch, 2008a), and the addition of a fourth character state by Schoch et al. (2020) to specifically differentiate the further reduction observed in batrachians does not resolve this. This character was thus modified here to fully encompass all possible presacral counts. However, I did not attempt to modify characters with ambiguous qualitative distinctions (e.g., character 307, femur 'long' vs. 'short') in order to minimize the number of necessary recodings. There is therefore substantial room to refine existing characters following the recommendations of other workers for standardizing character construction (e.g., Sereno, 2007).

#### Appendix 2.2. Operational terminology.

For brevity in the sections below and to remain consistent and transparent with respect to all of the changes I made to the matrix, I outline how specific terms are employed with respect to the coding changes that I made. The following terms are made with respect to the **original state coded.** Character numbering in this section refers to the original numbering of Schoch (2013), Pardo et al. (2017a), and Schoch et al. (2020) regardless of the omission of several characters of previous studies (14, 47, 48, 61, 78, 81, 100, 115, 253, 259, 334); the new numbering scheme is detailed along with the revised character list in Appendix 6.

- An **unsubstantiated code** is classified as one where the original coding was for a specific state (i.e., not unknown or inapplicable) and was subsequently modified; the following terms define the four primary types of unsubstantiated codes.
    - A **mischaracterized code** is when the taxon was coded for one state but should be coded for another character state. An example is the original coding of the zatracheids *Acanthostomatops vorax* and *Zatrachys serratus* as having a prefrontal and postfrontal separated by the frontal (42–1). In fact, these taxa have a wide prefrontal-postfrontal contact (42–0, the plesiomorphic condition; e.g., Langston, 1953; Schoch, 1997; Witzmann & Schoch, 2006; Urban & Berman, 2007), and this character in particular appears to have frequently been mischaracterized.
    - An **overgeneralized code** is when the taxon was coded for one state but should be coded for two or more as part of a polymorphism. In the absence of any previously specified protocol regarding the treatment of non-ontogenetic intraspecific variation, polymorphisms were coded across all scales within a species (between populations, within populations, within individuals) except in cases of explicit ontogenetic transformations, and no consideration was given to the relative frequency of either a polymorphism or of a specific state.
    - An **untestable code** is when the taxon cannot be coded for the character because the characterized feature is unknown in that taxon. An example is the original coding of the rhytidosteid *Sangaia lavina* as having four manual digits (197–1). However, the original description by Dias-da-Silva et al. (2006:382) only noted two specimens, one of which is a left half of the skull and the other of which is a palatal fragment (i.e. no postcrania were represented). Two specimens previously published as Rhytidosteidae indet. (Dias-da-Silva, 1998; Dias-da-Silva et al., 2005) and subsequently listed as referred material of *S. lavina* (Dias-da-Silva & Marsicano, 2011) also do not consist of any postcranial material, and no new material has been published since. Therefore, *S. lavina* cannot be coded for any postcranial features, let alone one that requires a high quality of preservation to

assess like digit count. Untestable states were recoded as unknown (?) unless they were related to a dependency (see next point) and are termed as much because there is no way for the code to be tested or reproduced from the literature.

- An **inapplicable code** is when the taxon should not be coded for a character because it is dependent on the presence of a feature that is known to be absent. Dependencies are extensively discussed in the literature (e.g., Maddison's 1993 example of two characters: presence/absence of a tail and tail color; see also Strong & Lipscombe, 1993; Brazeau, 2011; Simões et al., 2017). There are two types of dependencies, although individual characters are not identified as such.
  - The first is a violation of two basic tenets of character sampling: independence and homology. For example, if the ectopterygoid is absent (346–1), it is technically true that ectopterygoid fangs are absent (88–1); that the palatine ramus is only formed by the pterygoid (123–0) rather than by both the pterygoid and the ectopterygoid; and that a lateral exposure of the ectopterygoid is absent (132–0). This holds regardless of whether there is an express character for the presence/absence of the ectopterygoid. Some of these binary characters are specifically stated in 'presence/absence' language, while others merely express it in a different fashion. For example, character 124, with states 'pterygoid contacts only the palatine' versus 'pterygoid-contacts palatine and ectopterygoid,' could alternatively be expressed as 'pterygoid-ectopterygoid contact absent' versus 'pterygoid-ectopterygoid contact present.' They all correspond to the same primary character (ectopterygoid), and their secondary absences are all intrinsically linked to the same primary condition (absence of the ectopterygoid). These characters are therefore non-independent and coding secondary conditions like the absence of ectopterygoid fangs will overweight this absence.
    - In the example here, *Doleserpeton* has traditionally been coded as lacking an ectopterygoid (although this is equivocal at best based on the literature, see below), a feature shared with lissamphibians that is not found in any other temnospondyl. Considering that there are 10 secondary characters related to the ectopterygoid, it should be considered that this absence has overweighted this inferred homology in the debate over lissamphibian origins under the presumption that it lies within Temnospondyli. While the 'intent' of employing these secondary characters is not expressly stated, presumably it is intended as a means of differentiating between the subset of taxa that all have an ectopterygoid. Therefore, coding taxa without an ectopterygoid for these secondary presence/absence characters will obfuscate differentiation of taxa because there is a false equivalency (unequivocal non-homology) between absence due to absence of ectopterygoid (a consequence) and absence in the presence of ectopterygoid (a condition).
  - The second type of dependency is when a dependent secondary character cannot be coded under any interpretation because it does not express presence/absence but rather characterizes a feature assumed to be present.

Presence/absence may be explicitly defined in another character but does not need to be for this dependency to exist. An example in this matrix is character 211 (osteoderms, width), which has two states: 'narrow median osteoderms' or 'transversely extended plates.' Any taxon without any osteoderms (209–0), as is the condition of most temnospondyls, cannot be coded for osteoderm width, as osteoderms do not exist. However, most taxa without osteoderms were originally coded for 211–0 ('narrow').

- An **omission** occurs when the original coding was listed as unknown (?), but now available published literature allows for coding of the character. An example is the original coding of the trematopids *Acheloma cumminsi* and *Phonerpeton pricei* as being unknown for lacrimal-orbit contact (220). However, the lacrimal enters the orbit in these taxa (220–1), as in most dissorophoids and is well-documented in the historical literature (e.g., Olson, 1941; Dilkes & Reisz, 1987; Dilkes, 1990, 1993; Polley & Reisz, 2011). In the specific context of this study, I only coded these characters if the literature is sufficient to confidently do so; there may therefore be some missing data that other workers would be able to rectify through personal observation or access to relatively inaccessible publications (e.g., obscure journals, print-only materials). Omissions are not considered errors by default, but as is discussed below and in the main text, there are non-random patterns to omissions within certain characters that hint at unstated coding philosophies whose basis is questionable.

The following terms are made with respect to the changes implemented by us to the original code:

- A **correction** is a coding change made based on the identification of an unsubstantiated code. All types of unsubstantiated codes are considered to be in need of correction.
- A **recoding** is a coding change made based on a modification to a character, such as the addition of a new character state, or a coding change made based on a modification to a taxon, such as the restriction of the former composite OTU of *Cacops aspidephorus* + *Cacops morrisi* to strictly *Cacops aspidephorus*.
- A **change to inapplicable** refers specifically a change from '?' to '-'. This has no significance from an algorithmic perspective, as inapplicable cells are treated as missing data, but as with Pardo et al. (2017a), I prefer to explicitly express inapplicable cells as such. My motivation here is for future use of this matrix; when inapplicable cells are left as unknown, the presence of this code cannot differentiate between a cell that was accidentally missed (an omission), a cell that was intentionally left as '?' because the feature is unknown (missing data), and a cell that was intentionally left as '?' because the character is inapplicable to the taxon ('coded').

#### Appendix 2.3. Literature sources

The following table outlines the literature that I used in this study for various taxa. Note that it lists only taxon-specific literature to avoid redundancy, but synoptic work such as Jupp & Warren's (1986) review of stereospondyl lower jaws, Warren & Snell's (1991) review of stereospondyl postcrania, Warren & Davey's (1992) review of temnospondyl and early tetrapod dentition, Witzmann's (2007) review of the scalation, Witzmann's (2013) review of the hyobranchial apparatus, and the taxonomic reviews by Schoch & Milner (2000, 2014) were also utilized and are cited in relevant parts of this appendix.

**Table 1. List of sampled taxa and associated literature used to examine and recode the matrices of Schoch (2013) and Pardo et al. (2017a).** Taxa are listed in the same order as they were in previous analyses. References that are bolded were also expressly utilized by previous workers. Taxa with an asterisk (*) represent replacements of a congeneric taxon from the original analyses; further details are found in Appendix 2.5.1. Although I have personally examined material of some of these taxa, such observations were not made in the specific context of this study, and therefore express denotation of these taxa is omitted.

| Taxon | Literature |
|---|---|
| ***Proterogyrinus scheelei*** | Holmes (1984) |
| ***Greererpeton burkemorani*** | Smithson (1982); **Godfrey (1989a, 1989b)**; Bolt & Lombard (2001) |
| ***Edops craigi*** | **Romer & Witter (1942)**; Schoch (2017) |
| ***Adamanterpeton ohioensis*** | **Milner & Sequeira (1998)** |
| ***Cochleosaurus bohemicus*** | **Sequeira (2004, 2009)** |
| ***Nigerpeton ricqlesi*** | Sidor et al. (2005); **Steyer et al. (2006);** Sidor (2013) |
| ***Chenoprosopus milleri*** | **Langston (1953)** |
| ***Dendrerpeton helogenes*** | Godfrey et al. (1987); **Holmes et al. (1998)** |
| ***Balanerpeton woodi*** | **Milner & Sequeira (1993)** |
| ***Trimerorhachis insignis*** | **Case (1935); Pawley (2007);** Milner & Schoch (2013) |
| ***Neldasaurus wrightae*** | **Chase (1965);** Schoch (2018a) |
| ***Dvinosaurus primus*** | **Bystrow (1938)** |
| ***Acroplous vorax*** | Coldiron (1978); **Englehorn et al. (2008)** |
| ***Isodectes obtusus*** | Watson (1956); **Sequeira (1998)** |
| ***Thabanchuia oomie*** | **Warren (1998a)** |
| ***Tupilakosaurus wetlugensis*** | Shishkin (1961) |
| ***Capetus palustris*** | **Sequeira & Milner (1998)** |
| ***Iberospondylus schultzei*** | **Laurin & Soler-Gijón (2001, 2006)** |
| ***Acanthostomatops vorax*** | **Witzmann & Schoch (2006)** |
| ***Zatrachys serratus*** | **Langston (1953);** Schoch (1997) |
| ***Micromelerpeton credneri*** | **Boy (1995)** |
| ***Limnogyrinus elegans*** | Werneburg (2012); Schoch & Witzmann (2018) |
| ***Branchierpeton amblystomum*** | **Werneburg (1991)** |
| ***Branchiosaurus salamandroides*** | Werneburg (2012) |
| ***Apateon pedestris*** | Schoch (2004); Schoch & Milner (2008) |
| ***Micropholis stowi*** | **Schoch & Rubidge (2005)** |
| ***Platyrhinops lyelli*** | **Clack & Milner (2010)** |
| ***Amphibamus grandiceps*** | **Milner (1982)** |
| ***Doleserpeton annectens*** | **Sigurdsen & Bolt (2010)** |
| ***Gerobatrachus hottoni*** | **Anderson et al. (2008)** |
| ***Dissorophus multicinctus*** | **DeMar (1968);** Dilkes & Brown (2007); Dilkes (2009, 2020) |
| ***Cacops aspidephorus*** | **Williston (1910);** Anderson (2005); Dilkes & Brown (2007); Dilkes (2009); Anderson et al. (2020) |
| ***Broiliellus texensis*** | Williston (1914); DeMar (1966) |
| ***Acheloma cumminsi*** | Williston (1909); Olson (1941); Dilkes & Reisz (1987); **Polley & Reisz (2011);** Gee et al. (2019) |
| ***Phonerpeton pricei*** | Olson (1941); **Dilkes (1990,** 1993) |
| ***Ecolsonia cutlerensis*** | Vaughn (1969); **Berman et al. (1985)** |
| ***Onchiodon labyrinthicus*** | **Boy (1990)** |
| ***Eryops megacephalus*** | **Sawin (1941); Pawley & Warren (2006)** |
| ***Sclerocephalus haeuseri*** | **Schoch & Witzmann (2009a)** |
| ***Glanochthon latirostris*** | **Schoch & Witzmann (2009b)** |

| | |
|---|---|
| Archegosaurus decheni | **Witzmann (2005)** |
| Australerpeton cosgriffi | Barberena (1998); **Dias & Schultz (2003);** Eltink & Langer (2014); Eltink et al. (2016); Azevedo et al. (2017) |
| Platyoposaurus stuckenbergi | Schoch & Milner (2000) |
| Lapillopsis nana | **Yates (1999)** |
| Peltobatrachus pustulatus | **Panchen (1959)** |
| Rhineceps nyasaensis | **Watson (1962)** |
| Uranocentrodon senekalensis | **Van Hoepen (1915);** Broom (1930); Watson (1962); Marsicano et al. (2017) |
| Broomistega putterilli | **Shishkin & Rubidge (2000)**; Fernandez et al. (2013) |
| Lydekkerina huxleyi | **Pawley & Warren (2005);** Jeannot et al. (2006); Hewison (2007) |
| Chomatobatrachus halei | **Cosgriff (1974);** Warren et al. (2006) |
| Sangaia lavina | Dias-da-Silva (1998); **Dias-da-Silva et al.** (2005, **2006**); Dias-da-Silva & Marsicano (2011) |
| Peltostega erici | Cosgriff & Zawiskie (1979); **Janvier (1983);** Schoch & Milner (2000) |
| Laidleria gracilis | **Kitching (1957); Warren (1998)** |
| Siderops kehli | **Warren & Hutchinson (1983)** |
| Batrachosuchus browni | **Watson (1956);** Welles & Estes (1969); Chernin (1977); Warren & Marsicano (2000) |
| Plagiosuchus pustuliferus | Hellrung (2003); **Damiani et al. (2009)** |
| Gerrothorax pulcherrimus | **Hellrung (2003);** Jenkins et al. (2008); Schoch & Witzmann (2012) |
| Edingerella madagascariensis | Lehman (1961); Warren & Hutchinson (1988); Steyer (2003); **Maganuco et al. (2009)** |
| Watsonisuchus gunganj | **Warren (1980)** |
| Benthosuchus sushkini | **Bystrow & Efremov (1940);** Damiani (2001) |
| Trematosaurus brauni | Schoch & Milner (2000); Schoch (2019) |
| Trematolestes hagdorni | **Schoch (2006)** |
| Lyrocephaliscus euri | Mazin & Janvier (1983); Schoch & Milner (2000) |
| Callistomordax kugleri | **Schoch (2008a)** |
| *Metoposaurus krasiejowensis | Sulej (2002, 2007); Konietzko-Meier & Wawro (2007); Konietzko-Meier et al. (2020) |
| Wetlugasaurus angustifrons | Schoch & Milner (2000) |
| Parotosuchus nasutus | Schoch & Milner (2000); Schoch (2018c) |
| Eryosuchus garjainovi | Schoch & Milner (2000) |
| Paracyclotosaurus davidi | **Watson (1958)** |
| *Eocyclotosaurus appetolatus | Rinehart et al. (2015); Rinehart & Lucas (2016) |
| *Cyclotosaurus intermedius | **Sulej & Majer (2005)** |
| Mastodonsaurus giganteus | Schoch (1999) |
| Rileymillerus cosgriffi | Bolt & Chatterjee (2000); Martz et al. (2012) |
| Chinlestegophis jenkinsi | Pardo et al. (2017a) |
| Eocaecilia micropodia | Jenkins et al. (2007) |
| Kokartus honorarius | Skutschas & Martin (2011) |
| Karaurus sharovi | Shishkin (2000); Carroll (2007) |
| Triadobatrachus massinoti | Ascarrunz et al. (2016) |
| Leptodactylus mystacinus | Ponssa (2008) |
| Xenopus tropicalis | Goodman et al. (2021) |
| Cryptobranchus allegheniensis | Reese (1906); Carroll & Holmes (1980); MorphoSource (scan via David Blackburn / Florida Museum of Natural History) |
| Ambystoma opacum | Cope (1889); Stokely & Holle (1954); Tihen (1958) |
| Hynobius spp. | Xiong et al. (2007); Vassilieva et al. (2015); Hara & Nishikawa (2021) |
| Ichthyophis bannanicus | DigiMorph (scan via D. Cannatella / AmphibiaTree) |
| Epicrionops bicolor | DigiMorph (scan via D. Cannatella / AmphibiaTree) |

#### Appendix 2.4. Character-level modifications.

##### Appendix. 2.4.1. Unspecified character states.

A number of multistate characters occur in the matrix where the numerically highest state was not listed in the character list of previous studies. All of these characters are characters from the original Schoch (2013) matrix, but most of the ambiguous character states first appeared in the matrix of Pardo et al. (2017a). Pardo et al. (2017a)'s typeset character list does not reflect these added states, and there is no comment in either the main text or supplemental information about character changes, but these added states can mostly be gleaned from their NEXUS script in their supplemental information.

**Character numbering in this section refers to the original numbering of Schoch (2013), Pardo et al. (2017a), and Schoch et al. (2020**) regardless of the omission of several characters of previous studies (14, 47, 48, 61, 78, 81, 100, 115, 253, 259, 334); the new numbering scheme is detailed in Appendix 6. Note that characters discussed in this section are written without modification from those listed by previous studies, with any modifications to these characters detailed in the revised character list. Taxa are listed only by the genus for readability, as no genus has more than one species sampled in this matrix. Lists of taxa are ordered following their order in the matrix.

- **3. Ornament (elements). Reticulate ridges of various sizes (0), or isolated pustules (1).**
    - State 2 was defined in the script of Pardo et al. (2017a) as ' absent'; this was originally coded for most lissamphibians other than karaurids and *Epicrionops*. With that said, the condition of *Eocaecilia* (densely distributed series of isolated, shallow pits; Jenkins et al., 2007) was not differentiated from the essentially smooth skull roof of batrachians (i.e. it was more of a catch-all state for ornamentation not predominated by ridges or pustules). This is also problematic because *Proterogyrinus* was coded for 3–0 (ridges), but *Proterogyrinus* has ornamentation that is relatively dissimilar to temnospondyls and *Greererpeton*, with minute pits and grooves (Holmes, 1984), rather than ridges or pustules. Therefore, state 2 is restricted to 'dense covering of minute pits' like those found in *Proterogyrinus* and *Eocaecilia*, and a new state 3 is defined as 'sparsely distributed pits or absent.' Batrachians and gymnophionans were recoded for 3–3. *Eocaecilia* was left as 3–2, and *Proterogyrinus* was recoded to 3–2.
- **9. Preorbital region (length). Less than twice the length of posterior skull table (0), or more (1).**
    - State 2 was originally defined as 'equal [in length]' by Pardo et al. (2017a), but this condition would be covered by state 0; state 2 is redefined herein as 'twice as long.'
- **26. Orbit location. Medial, framed by wide jugals laterally (0), or lateral emplacement, framed by very slender jugals (1).**
    - State 2 was coded for *Platyrhinops*, *Amphibamus*, *Doleserpeton*, and *Gerobatrachus* among temnospondyls; for *Ambystoma, Hynobius*, *Cryptobranchus*, *Leptodactylus*, and *Xenopus* among extant lissamphibians; and for *Eocaecilia*, *Karaurus*, and *Triadobatrachus* among extinct lissamphibians. This was one character for which the uncertain state is not listed in either the character list or NEXUS script and was apparently intended to differentiate

lissamphibians and putative close dissorophoid relatives with greatly enlarged orbits from other in-group taxa. Personal communication with J. Pardo for another study (Gee et al., in prep) clarified that state 2 refers to enlarged orbits that are framed by both slender jugals and slender frontals. *Eocaecilia* was therefore recoded to state 1 because it has wide frontals and consequently small, more laterally situated orbits (Fig. 1H).

- The original restriction of state 2 to highly nested amphibamiforms is not surprising, but it is unclear why other taxa with proportionately large orbits that can be said to have both slender jugals and slender frontals were not recoded to state 2, including the early diverging amphibamiform *Micropholis*, micromelerpetids, the highly nested branchiosaurid amphibamiforms, and *Lapillopsis* (e.g., Schoch & Milner, 2014:figs. 25–27; Fig. 1E, 1F). The orbits of *Platyrhinops* in particular are proportionately much smaller than in other amphibamiforms coded for state 2, which might relate to its large size; the interorbital region is wider than the orbits, and therefore the frontals are not slender, and the orbits are more distinctly laterally situated (Clack & Milner, 2010; Fig. 1D). In other words, if *Platyrhinops* is coded as state 2, all of the aforementioned taxa should be as well. I recoded *Platyrhinops* to state 1 and recoded *Apateon, Branchiosaurus*, and *Lapillopsis* to state 2.

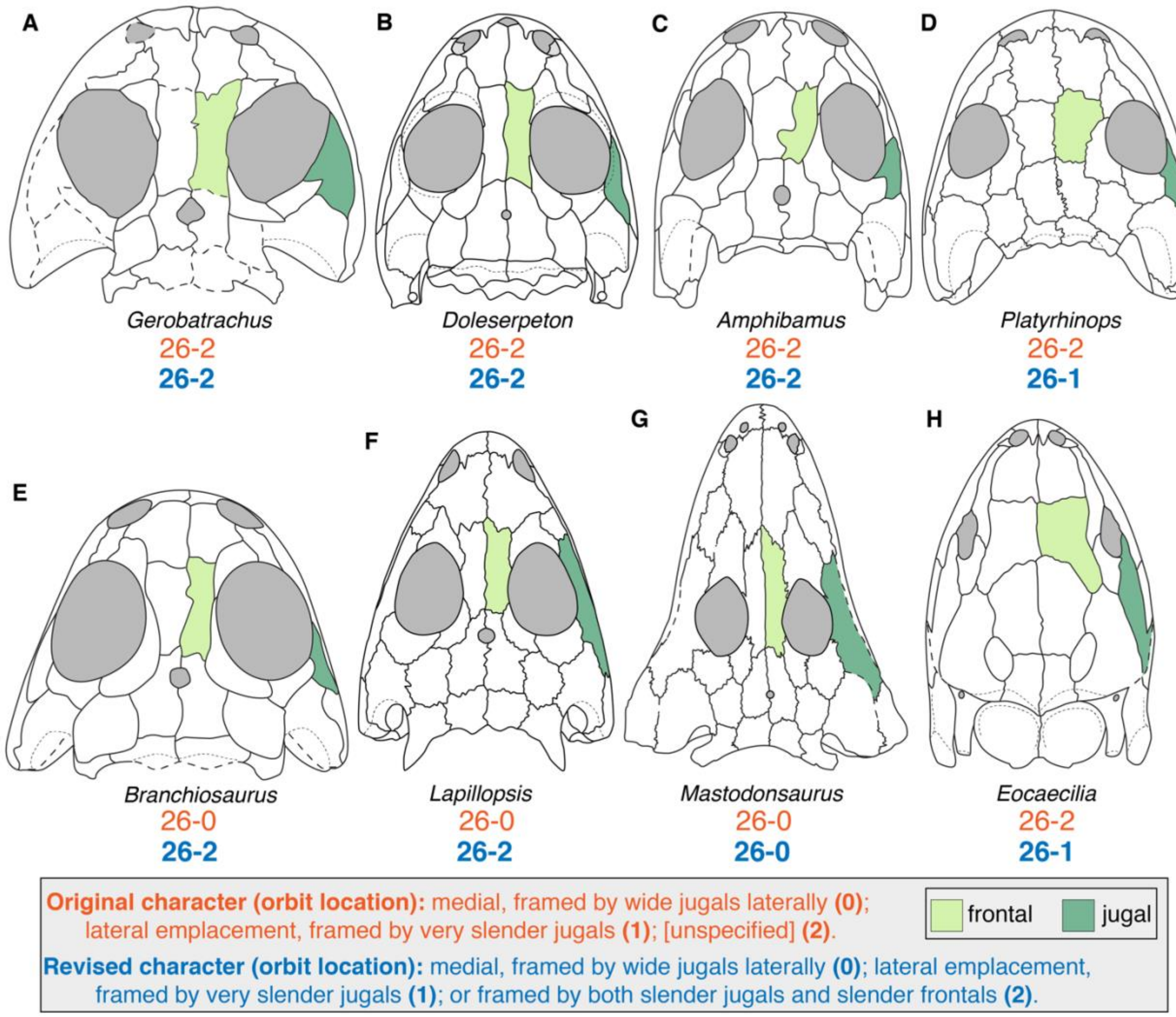

**Figure 1. Comparison of orbit location among temnospondyls and lissamphibians. A**, the amphibamiform *Gerobatrachus hottoni* (after Schoch & Milner, 2014); **B**, the amphibamiform *Doleserpeton annectens* (after Sigurdsen & Bolt, 2010); **C**, the amphibamiform *Amphibamus grandiceps* (after Schoch & Milner, 2014); **D**, the amphibamiform *Platyrhinops lyelli* (after Clack & Milner, 2010); **E**, the branchiosaurid *Branchiosaurus salamandroides* (after Schoch & Milner, 2014); **F**, the stereospondyl *Lapillopsis nana* (after Yates, 2000); **G**, the capitosaur *Mastodonsaurus giganteus* (after Schoch, 1999); **H**, the gymnophionomorph *Eocaecilia micropodia* (after Jenkins et al., 2006). Orange text represents the original character and codes from Pardo et al. (2017; some carried over from Schoch, 2013); blue text represents the revised character and codes. Silhouettes not to scale.

- **54. Supratemporal (lateral margin). Rectangular with straight sagittal lateral margin (1), or posterolaterally constricted by expanded otic notch (1).**
  - State 2 was defined in the script of Pardo et al. (2017a) as ‘small and oval in shape.’ However, this state would be better placed with character 53 (referring to proportions of the supratemporal) rather than with one referring only to the lateral margin (the inclusion of ‘rectangular’ in state 0 is superfluous and was removed here to avoid overgeneralization of the anatomy across the taxon sample). It is clearly a compound state, prescribing both size (without defining how size was determined) and shape. This state was simply omitted. *Chinlestegophis* and *Eocaecilia* were recoded to 54–0.

- **112. Cultriform process (width). Base not wider than rest, clearly set off from basal plate (0), or merging continuously into plate (1).**
    - State 2 was ‘flaring anteriorly.’
- **158. Presacral count. More than 28 (0), or 23-25 vertebrae (1), or less than 23 (2).**
    - State 3 was introduced by Schoch et al. (2020) and is only considered here because it could be useful for differentiating taxa with low presacral counts. Schoch et al. did not provide a revised character list or a script with character names and definitions, and their list of coding changes does not include any mention of character alterations, but there is a clear pattern for this new state based on how taxa were coded. State 3 was coded for *Amphibamus* (18? positions) and *Gerobatrachus* (17 positions) among temnospondyls; for *Ambystoma* (14 positions), *Hynobius* (15–18 positions), *Cryptobranchus* (9 positions), *Leptodactylus* (8 positions), and *Xenopus* (8 positions) among extant lissamphibians; and for *Karaurus* (12–13 positions), *Triadobatrachus* (15 positions), and *Triassurus* (16 positions) among extinct lissamphibians (number in parentheses represents presacral count). *Amphibamus* represents the upper bound for taxa coded for state 3, with a presumed 18 presacral positions (e.g., Clack & Milner, 2010; Schoch & Milner, 2014, contra Schoch et al., 2020, in which they state it has 24). If state 3 is ‘<19 presacral positions,’ this would also align with why *Apateon* (19–22 positions) was coded for state 2. Therefore, 158–3 is formally defined as ‘< 19 positions.’
- **201. Ilium (orientation). Main axis of shaft inclined posterodorsally (0), or vertical (1).**
    - State 2 was defined in the script of Pardo et al. (2017a) as ‘anteriorly.’

##### Appendix. 2.4.2. Modification to existing characters.

The following comments pertain to characters in their entirety, including but not limited to, issues with character construction, character dependencies, and character redundancies. **Character numbering in this section refers to the original numbering of Schoch (2013), Pardo et al. (2017a), and Schoch et al. (2020) regardless of the omission of several characters of previous studies (14, 47, 48, 61, 78, 81, 100, 115, 253, 259, 334); the new numbering scheme is detailed along with the revised character list.** Taxa are listed only by the genus for readability, as no genus has more than one species sampled in this matrix. Lists of taxa are ordered following their order in the matrix. For brevity, references are only provided when deemed necessary (e.g., newly coding a polymorphism), and the reader is directed to the list of utilized references for otherwise uncited taxa. Unless otherwise stated, coding modifications were made across all versions of the matrix.

- **2 (skull, height); 134 (palate structure); 245 (skull, occipital profile):** these characters require an undistorted skull with the occiput preserved and exposed in order to be coded. Taxa only represented by flattened or disarticulated specimens should not be coded for either, even if they are reconstructed, as a given element can vary in the degree of curvature or orientation (e.g., quadratojugal, pterygoid) such that width or lateral extent are not a reliable proxy.
    - Codes were recoded from 2–0 to unknown for *Proterogyrinus, Cochleosaurus, Adamanterpeton, Balanerpeton, Acanthostomatops, Capetus, Iberospondylus, Micromelerpeton, Limnogyrinus, Branchierpeton, Branchiosaurus, Apateon,*

*Platyrhinops, Doleserpeton, Amphibamus, Gerobatrachus, Onchiodon, Sclerocephalus, Glanochthon, Trematolestes, Callistomordax, Triadobatrachus, Karaurus, Kokartus*, and *Eocaecilia*.

  - Codes were recoded from 134–0 to unknown for *Proterogyrinus, Cochleosaurus, Adamanterpeton, Balanerpeton, Acanthostomatops, Capetus, Iberospondylus, Micromelerpeton, Limnogyrinus, Branchierpeton, Branchiosaurus, Apateon, Platyrhinops, Doleserpeton, Amphibamus, Gerobatrachus, Onchiodon, Sclerocephalus, Glanochthon, Trematolestes, Callistomordax, Triadobatrachus,* and *Kokartus*.
  - Codes were recoded from 245–0 to unknown for *Cochleosaurus, Balanerpeton, Acanthostomatops, Capetus, Iberospondylus, Apateon, Platyrhinops, Doleserpeton, Amphibamus, Gerobatrachus, Onchiodon, Sclerocephalus, Glanochthon, Trematolestes, Callistomordax, Triadobatrachus, Kokartus,* and *Eocaecilia*. Codes were recoded from 245–2 to unknown for *Proterogyrinus*. This character was originally parsimony-uninformative and remains as such following modifications.

- **4 (ornament, snout):** a third state (numerous circular to subcircular pits) is added to capture the condition found in gymnophionomorphs.
  - *Ichthyophis*, *Epicrionops*, and *Eocaecilia* were recoded to the new 4–2.
- **10 (premaxilla, alary process):** originally, the absence of an alary process (10–0) was only coded for the outgroups, edopoids, and the dvinosaur *Acroplous.*
  - **10. Premaxilla (alary process).** Absent (0), or present, forming a posterior hook-like indentation (1). [*added by Schoch, 2013*]

Schoch (2013) listed an alary process as an unambiguous synapomorphy for all post-edopoid temnospondyls (his Eutemnospondyli), but there is substantial variation in premaxillary morphology among this subset such that it is unclear what feature is being coded (Fig. 2). Edopoids have a long premaxilla that results in a long prenarial region, but this is captured by other characters, and the obliquely oriented premaxilla-nasal suture is shared with many other taxa coded as having an alary process (10–1; e.g., *Lydekkerina*, *Laidleria*, *Mastodonsaurus*; Warren, 1998a; Schoch, 1999; Jeannot et al., 2006; Fig. 2A). There are also taxa with essentially horizontal sutures that were coded for 10–1 (e.g., *Apateon*, *Gerrothorax*, *Trematolestes*; Hellrung, 2003; Schoch, 2006; Schoch & Milner, 2008; Schoch & Witzmann, 2012; Fig. 2C, 2G).

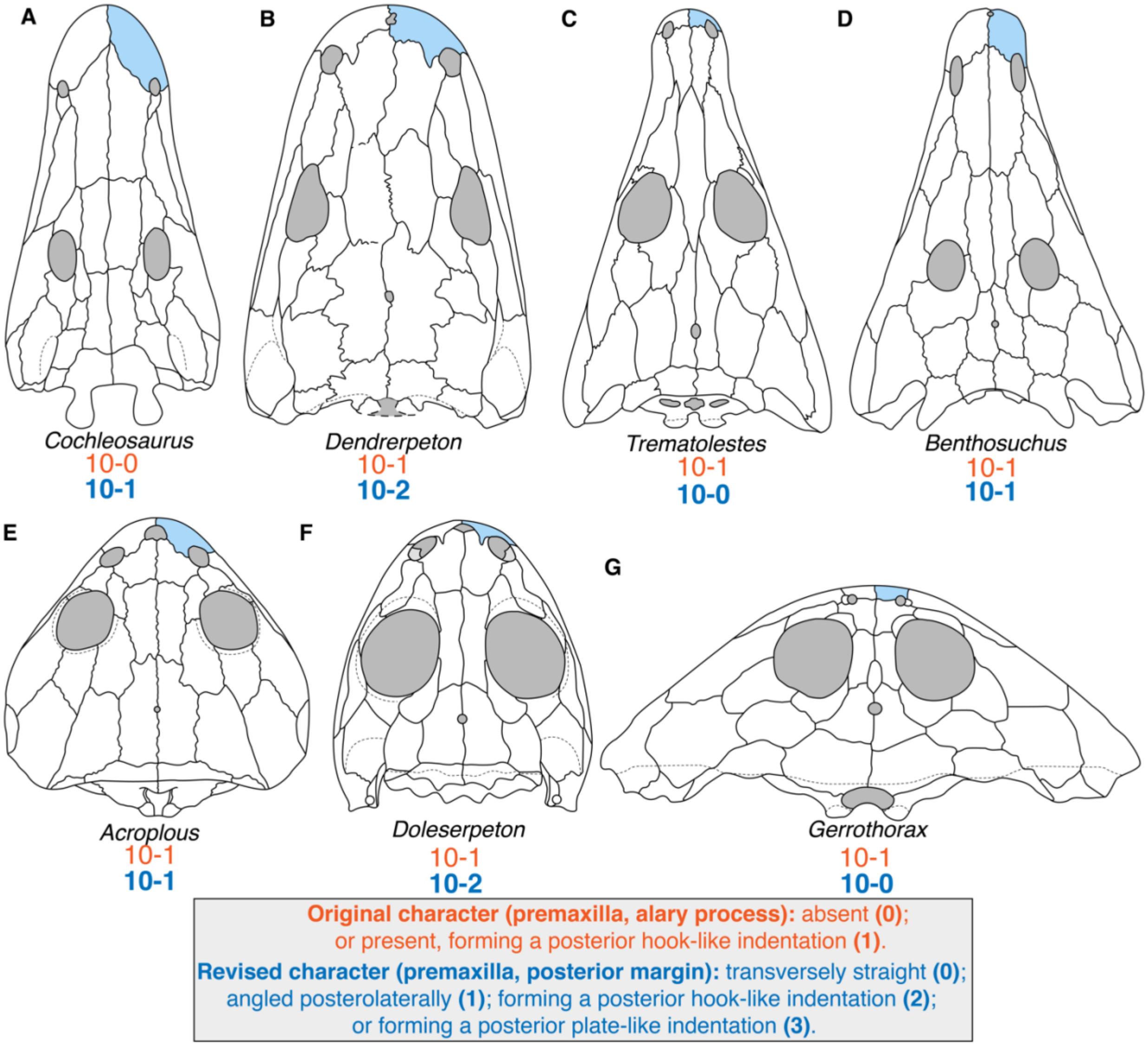

**Figure 2. Comparison of the posterior margin of the premaxilla among temnospondyls.** **A**, the edopoid *Cochleosaurus bohemicus* (after Sequeira, 2004); **B**, the 'dendrerpetid' *Dendrerpeton acadianum* (after Holmes et al., 1998); **C**, the trematosaur *Trematolestes hagdorni* (after Schoch, 2006); **D**, the trematosaur *Benthosuchus sushkini* (after Schoch, 2008); **E**, the dvinosaur *Acroplous vorax* (after Englehorn et al., 2008); **F**, the amphibamiform *Doleserpeton annectens* (after Sigurdsen & Bolt, 2010); **G**, the plagiosaurid *Gerrothorax pulcherrimus* (after Schoch & Witzmann, 2012). Orange text represents the original character and codes from Schoch (2013); blue text represents the revised character and codes. Silhouettes not to scale.

Other workers appear to use a differing, and arguably more consistent, definition of an 'alary process.' For example, per Ruta & Bolt (2006:appendix 2): "the alary process is a subtriangular bony lappet that arises from the posterolateral margin of the dorsal ramus of the premaxilla, and extends posteriorly overlapping the nasal along or near the margin of the external nostril." This definition specifies a discrete process, which is why it may be treated as absent in edopoids because the massive premaxillae of this clade do not have a distinct process on the dorsal surface; this distinction is similar to character 5 of Yates & Warren (2000). On this basis, taxa with an unequivocal alary process (10–1) would include 'dendrerpetids' (e.g., Milner & Sequeira, 1993; Holmes et al., 1998; Fig. 2B), some dvinosaurs (e.g., *Isodectes*; Sequeira, 1998), and many dissorophoids (e.g., *Micromelerpeton, Amphibamus, Cacops, Ecolsonia*; Berman et al., 1985; Boy, 1995; Reisz et al., 2009; Schoch & Milner, 2014; Fig. 2G). In these taxa, there is a discrete posterior process with a partially to fully defined lateral margin. However, under the same more explicit definition, many non-edopoids coded as having an alary process by Schoch (2013) do not in fact have such a process. To remove ambiguity, a second state is added here to differentiate between taxa with a fully distinct process (the new 10–2) and those with a slanted posterior margin (the new 10–1).

Schoch et al. (2020) added another character referring to the alary process (350 therein).

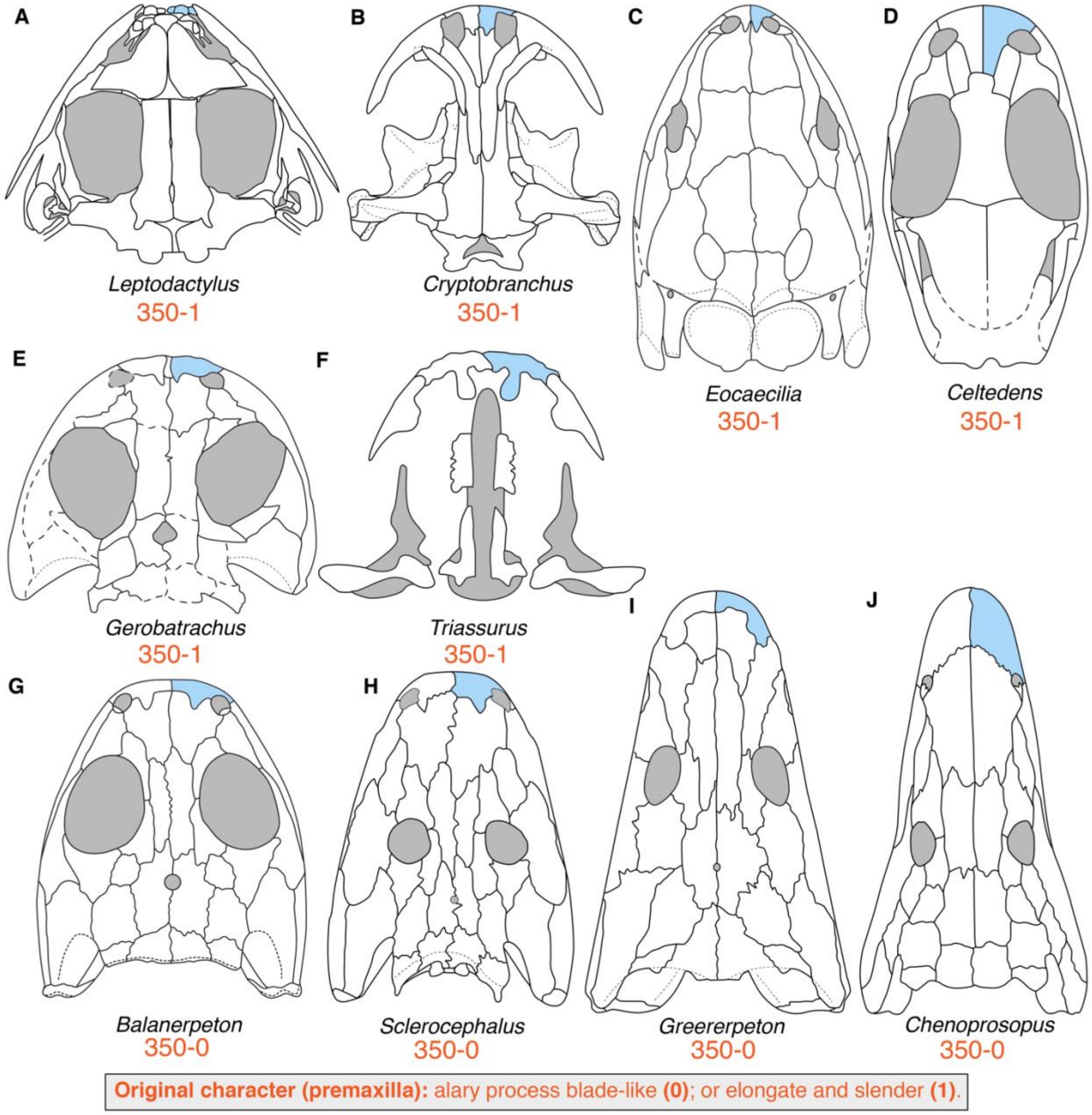

**Figure 3. Comparison of alary process morphologies among temnospondyls and lissamphibians. A**, the anuran *Leptodactylus mystacinus* (after Ponssa, 2008); **B**, the caudate *Cryptobranchus allegheniensis* (after Carroll &

Holmes, 1980); **C**, the gymnophionomorph *Eocaecilia jenkinsi* (after Jenkins et al., 2007); **D**, the albanerpetontid *Celtedens ibericus* (after McGowan, 2002); **E**, the amphibamiform *Gerobatrachus hottoni* (after Schoch & Milner, 2014); **F**, the urodele *Triassurus sixtelae* (after Schoch et al., 2020); **G**, the 'dendrerpetid' *Balanerpeton woodi* (after Milner & Sequeira, 1993); **H**, the stereospondylomorph *Sclerocephalus haeuseri* (after Schoch & Witzmann, 2009a); **I**, the colosteid *Greererpeton burkemorani* (after Smithson, 1982); **J**, the edopoid *Chenoprosopus milleri* (after Schoch & Milner, 2014). Note that *C. milleri* is not considered to have an alary process and should not have been coded for this character. Orange text represents the original character and codes from Schoch et al. (2020). Silhouettes not to scale.

- **350. Premaxilla.** Alary process blade-like (0), or elongate and slender (1). [*added by Schoch et al., 2020*]

State 1 was only coded for all lissamphibians and *Gerobatrachus* (Fig. 3A-F), but I was not able to find a common condition among these taxa (e.g., Schoch et al., 2020:fig. 3; Fig. 3) – the premaxilla is not even known in *Triadobatrachus* (Ascarrunz et al., 2016) – and because of the above uncertainty over what represents an alary process. At least *Kokartus* and *Celtedens* have a broader, more plate-like process compared to the slender, triangular process found in some temnospondyls and in *Eocaecilia* (Fig. 3C-D, 3G-H). *Leptodactylus* has a dorsal projection of the premaxilla that is termed the alary process, but it does not have a posterior process on the skull roof, resulting in a rectangular dorsal profile (Fig. 3A).

While the original coding of this character cannot be reproduced, there is value in further differentiating a broad, plate-like process from a slender process. This is appended to character 10 as the new state 3, and taxa were recoded as part of the recoding for the entire character.

- Codes were recoded from 10–0 to the new 10–2 for *Greererpeton*.
- Codes were recoded from 10–0 to the revised 10–1 for *Edops*, *Cochleosaurus, Nigerpeton*, *Chenoprosopus*, and *Acroplous*.
- Codes were changed from 10–1 to the new 10–2 for *Dendrerpeton, Balanerpeton, Trimerorhachis, Neldasaurus, Isodectes, Capetus, Iberospondylus, Acanthostomatops*, *Micromelerpeton*, *Limnogyrinus*, *Micropholis*, *Platyrhinops, Amphibamus*, *Doleserpeton, Dissorophus*, *Cacops*, *Broiliellus*, *Acheloma, Phonerpeton*, *Ecolsonia, Onchiodon*, *Eryops*, *Sclerocephalus, Archegosaurus, Lapillopsis*, *Lyrocephaliscus*, *Hynobius*, *Karaurus*, *Cryptobranchus*, and *Eocaecilia*.
- Codes were changed from 10–1 to the new 10–3 for *Celtedens* and *Kokartus*.
- Codes were recoded from 10–1 to 10–0 for *Dvinosaurus*, *Branchierpeton, Branchiosaurus*, *Apateon*, *Australerpeton*, *Platyoposaurus*, *Rhineceps, Uranocentrodon*, *Broomistega*, *Lydekkerina*, *Chomatobatrachus*, *Gerrothorax, Edingerella*, *Trematosaurus*, *Trematolestes*, *Callistomordax*, *Wetlugasaurus, Parotosuchus*, *Paracyclotosaurus*, and *Leptodactylus*.
- The following taxa were newly coded for a polymorphism: *Adamanterpeton* (0 → 0&1; Milner & Sequeira, 1993:figs. 6, 9); *Zatrachys* (1 → 1&2; Schoch, 1997:figs. 1–2); *Metoposaurus* (1 → 0&1&2; Sulej, 2007:figs. 4–15).
- The following taxa were newly coded for partial uncertainty: *Gerobatrachus* (1 → 2/3) and *Ambystoma* (1 → 2/3).
- The following taxa were newly coded for 10–2: *Ichthyophis* and *Epicrionops*.

    - *Thabanchuia*, *Tupilakosaurus*, *Peltobatrachus*, *Sangaia*, *Peltostega*, *Siderops*, *Batrachosuchus*, *Plagiosuchus*, *Watsonisuchus*, were all originally coded for 10–1 but were uncoded because the premaxilla-nasal suture is conjectural.
- **13 (premaxillary foramen); 14 (premaxillary fontanelle):** character 13 referred to a small opening at the tip of the snout typically called the 'internarial fenestra', 'internarial pit,' or 'internarial fontanelle' in temnospondyls (e.g., Dilkes & Reisz, 1987:3; Clack & Milner, 2010:279; Milner, 2018:217; Schoch, 2019b:144). Character 14 appeared to refer to the enlarged opening found in zatracheids that is convergent with a large fontanelle found in many lissamphibians that is larger than the 'foramen' of temnospondyls but smaller than the 'fontanelle' of zatracheids.
    - **13. Premaxillary foramen.** Premaxilla and nasal completely sutured (0), or medially bearing an opening separating the two alary processes of the premaxilla (1). [*added by Schoch, 2013*]
    - **14. Premaxillary fontanelle.** Premaxillae with or without small opening (0), or encircling large fenestra extending posteriorly between the nasals (1). [*added by Schoch, 2013*]

All taxa coded for 14–1 (fontanelle present) were coded for 13–1 (fenestra present), but in none of these taxa are there two separate openings framed between the premaxilla and nasal (the internasal opening of *Kokartus* is considered to be a separate condition not homologous to 14–1 because it is entirely within the nasals; Skutschas & Martin, 2011). This is therefore a partial dependency (there must be a premaxillary foramen whenever there is a premaxillary fontanelle; or conversely, there cannot be a fontanelle if there is no foramen; Fig. 4). There is also a question of whether *Acanthostomatops* should be coded the same as *Zatrachys*. While a large fontanelle diagnoses the clade (e.g., Schoch & Milner, 2014), the opening in *Acanthostomatops* is substantially smaller than in either *Zatrachys* or in *Dasyceps* (the latter not sampled; Paton, 1975). The incompletely defined fontanelle in the recently described *Chemnitzion richteri* was inferred to be similar to that of *Acanthostomatops* (Werneburg et al., 2022). It barely divides the nasals posteriorly in *Acanthostomatops* (it fully divides them in *Zatrachys* and divides them for about half the length in *Dasyceps*), and it can therefore be argued that *Acanthostomatops* has a condition closer to non-zatracheids like *Cacops* in this respect. That *Acanthostomatops* has a slightly larger fontanelle than *Cacops* is a combination of premaxillary elongation (for which there is already a character) and overall skull proportions (more equant in width and length).

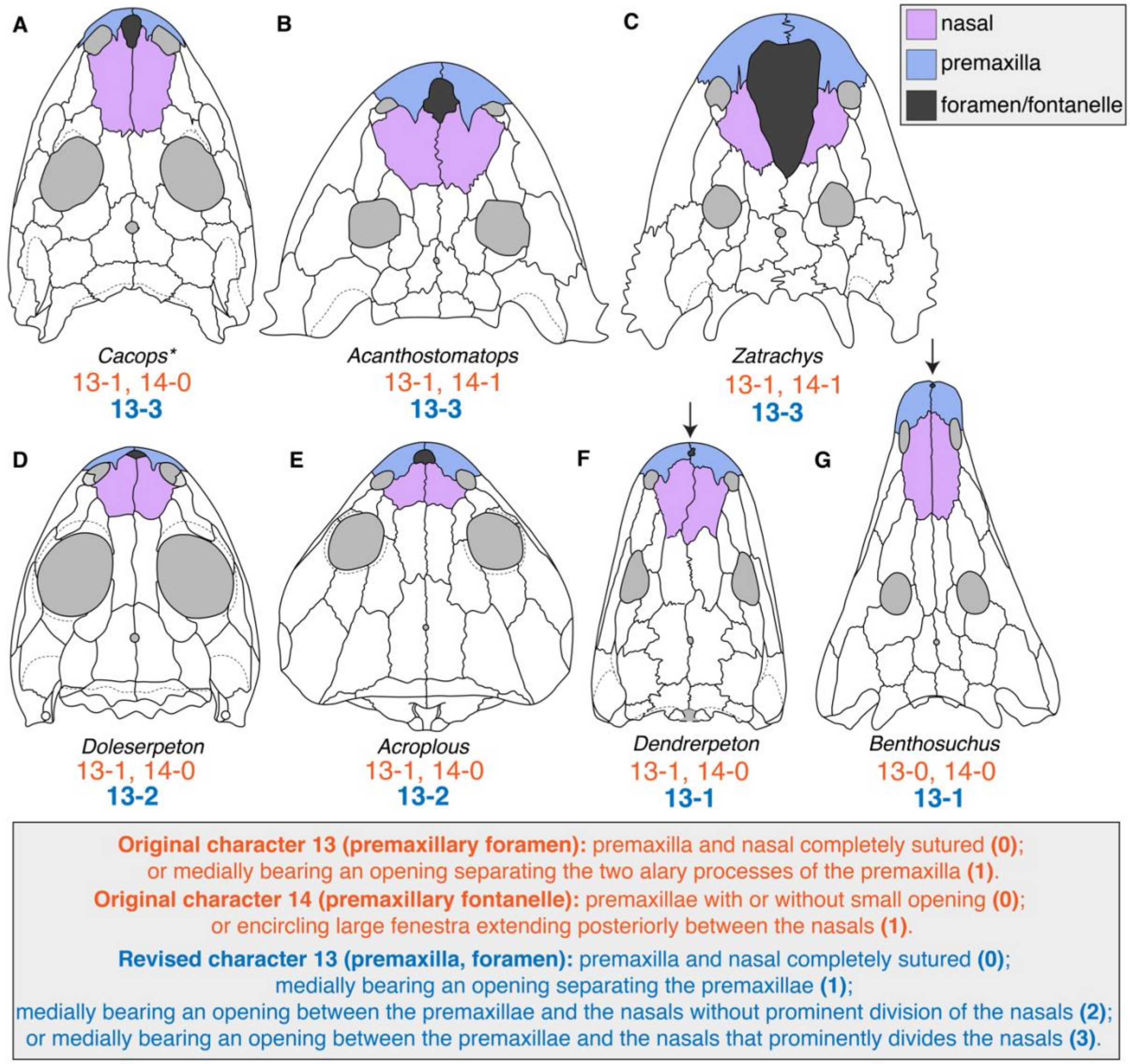

**Figure 4. Comparison of taxa with median rostral openings to illustrate the partial redundancy in the original coding of characters 13 (premaxillary foramen) and 14 (premaxillary fontanelle). A**, the dissorophid *Cacops morrisi* (after Reisz et al., 2009); **B**, the zatracheid *Acanthostomatops vorax* (after Schoch and Milner, 2014); **C**, the zatracheid *Zatrachys serratus* (after Schoch and Milner, 2014); **D**, the stereospondylomorph *Archegosaurus decheni* (after Witzmann, 2005); **E**, the amphibamiform *Doleserpeton annectens* (after Sigurdsen and Bolt, 2010); **F**, the dvinosaur *Acroplous vorax* (after Englehorn et al., 2008); **G**, the ‘dendrerpetid’ *Dendrerpeton helogenes* (after Holmes et al., 1998). Arrows are used to indicate the small foramen in *Dendrerpeton* and *Benthosuchus.* In contrast to other comparative cranial diagrams, both sides of the skull are colored here to demonstrate the degree to which the rostral opening divides the premaxillae and the nasals. The asterisk (*) next to *Cacops* is because the OTU was restricted to *Cacops aspidephorus* (in which the presence / size of a fontanelle is unclear; Anderson et al., 2020) rather than as a composite including *C. morrisi* (in which a fontanelle is entirely defined; Reisz et al., 2009), the revised code reflects the proper code if the latter were utilized as the sole OTU. Orange text represents the original character and codes from Schoch (2013); blue text represents the revised character and codes. Silhouettes not to scale.

- Characters 13 and 14 were first combined into a three-state character for the median rostral opening to minimize the number of inapplicable codes that would need to be introduced if they remained separate. A fourth state was added to further differentiate conditions. The new 13–1 is defined as an opening entirely framed within the premaxillae (e.g., *Acheloma*, *Benthosuchus*). The new 13–2 (equivalent to the old 13–1 and 14–0) is defined as an opening that is framed by both the premaxillae and the nasals but that does not prominently divide the nasals (e.g., *Acroplous*, *Phonerpeton*). The new 13–3 (equivalent to the old 13–1 and 14–1) is defined as an opening that prominently divides the nasals (e.g., *Acanthostomatops*, *Zatrachys*). Because I consider the condition of *Acanthostomatops* to be closer to that of *Cacops* (viz. *C. morrisi*; Reisz et al., 2009; Fig. 4A-B) than to *Zatrachys* (Fig. 4C), an additional state for 'greatly enlarged fontanelle' is not created, as it would render the character as parsimony-uninformative by virtue of occurring only in *Zatrachys* (a comparably sized opening occurs in no other taxon sampled in this family of matrices). Incidentally, because *Cacops* is restricted here to *C. aspidephorus*, in which the presence of any fenestra is equivocal (Anderson et al., 2020; in contrast to *C. morrisi* and to the original coding, Fig. 4A), only zatracheids were coded for the new 13–3 among temnospondyls. An example of a non-zatracheid other than *C. morrisi* that is not sampled here and that would be coded for 13–3 is the amphibamiform *Tersomius texensis* (Maddin et al., 2013). This new multistate character is not ordered.

- Codes were recoded from 13–1 to 13–2 for *Acroplous*, *Apateon*, *Doleserpeton*, *Phonerpeton*, *Rhineceps*, *Kokartus*, and *Hynobius*.
- Codes were recoded from 13–1 to 13–3 for *Acanthostomatops*, *Zatrachys*, and *Leptodactylus*.
- Codes were recoded from 13–1 to 13–1/2 for *Dendrerpeton* (the interpretation of Arbez et al., 2022, conflicts with that of Holmes et al., 1998).
- *Cochleosaurus* was corrected from 13–0 to 13–0&2. Sequeira (2004:25) noted the presence of a small foramen in some specimens (it is not reconstructed in that paper or subsequent reconstructions).
- *Micromelerpeton* was corrected from 13–0 to 13–2 because this taxon has an oval internarial fenestra (Boy, 1995; Schoch & Milner, 2014).
- *Benthosuchus* was corrected from 13–0 to 13–1 because this taxon has a small oval fenestra entirely within the premaxillae (e.g., Bystrow & Efremov, 1940; Schoch & Milner, 2000; Damiani, 2001).
- *Xenopus* was corrected from 13–1 to 13–0 because the genus has a fused nasal and there is no opening within it (e.g., Henderson, 2002:24; Goodman et al., 2021:fig. 2A). *Ambystoma* and *Karaurus* were also corrected from 13–1 to 13–0 because they have no apparent opening between the unfused nasals or premaxillae in the most mature specimens (e.g., Parker, 1887:pl. 27.6; Skutschas & Martin, 2011:825, 828; Schoch et al., 2020:fig. 3).
- *Thabanchuia*, *Tupilakosaurus*, *Branchiosaurus*, *Peltobatrachus*, *Sangaia*, *Peltostega*, *Siderops*, *Batrachosuchus*, and *Watsonisuchus* were originally coded for 13–0 (and 14–0) but were uncoded because published specimens are insufficient to code this character. *Cacops* was uncoded as part of the restriction of the OTU to *C. aspidephorus* (Anderson et al., 2020).

- **20 (nasal, lateral margin):** the original character is shown below:

- **20. Nasal (lateral margin).** Straight (0), stepped, with lateral excursion anterior to lacrimal (1). [*added by Schoch, 2013*]

The original definition is problematic because it specified the lacrimal as a reference point and therefore should not have been coded for any state for taxa lacking a lacrimal (but was; e.g., *Siderops, Batrachosuchus*). Furthermore, there is a wide variety of nasal morphologies that cannot be reconciled with the original codes (Fig. 5).

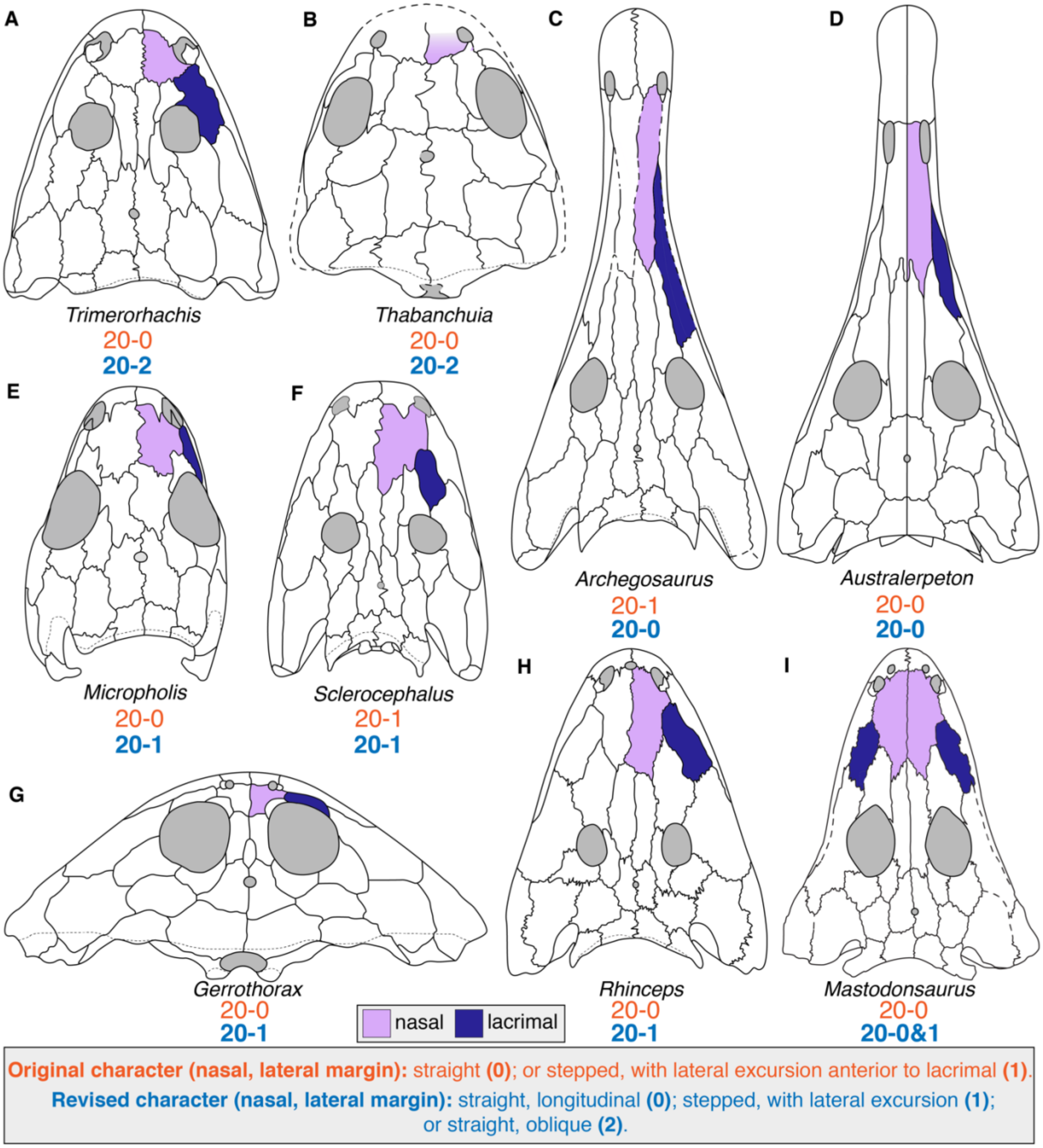

**Figure 5. Comparison of nasal morphology among temnospondyls to demonstrate previous coding inconsistencies in the treatment of character 20 (nasal, lateral margin). A**, the dvinosaur *Trimerorhachis insignis* (after Milner & Schoch, 2013); **B**, the dvinosaur *Thabanchuia oomie* (after Warren, 1998b); **C**, the stereospondylomorph *Archegosaurus decheni* (after Witzmann, 2005); **D**, the rhinesuchid *Australerpeton cosgriffi*

(after Eltink et al., 2016); **E**, the long-headed morph of the amphibamiform *Micropholis stowi* (after Schoch & Rubidge, 2005); **F**, the stereospondylomorph *Sclerocephalus haeuseri* (after Schoch & Witzmann, 2009a); **G**, the plagiosaurid *Gerrothorax pulcherrimus* (after Schoch & Witzmann, 2012); **H**, the rhinesuchid *Rhineceps nyasaensis* (after Schoch & Milner, 2000); **I**, the capitosaur *Mastodonsaurus giganteus* (after Schoch, 1999). Note that *T. oomie* lacks a lacrimal. Orange text represents the original character and codes from Schoch (2013); blue text represents the revised character and codes. Silhouettes not to scale.

Schoch (2013) only coded *Sclerocephalus*, *Glanochthon*, and *Archegosaurus* for 20–1 (Fig. 5C, 5F). Pardo et al. (2017a) changed *Lapillopsis* to 20–1 (without comment or documentation) and coded several lissamphibians for this state. Firstly, the condition of *Archegosaurus* (e.g., Witzmann, 2005:figs. 2–5; Fig. 5C) is much closer to that of many stereospondyls in which the nasal has a slightly angled contact with the lacrimal and then extends straight from the anterior terminus of the lacrimal (e.g., *Trematosaurus*, *Cyclotosaurus*; Schoch & Milner, 2000:fig. 101; Schoch, 2019a:figs. 2, 4–5); even Holmes' (1984:fig. 3) reconstruction of *Proterogyrinus* depicts this slight 'step.' Other taxa coded for 20–0 have a small lateral expansion just posterior to the naris that separates the lacrimal from the opening (e.g., *Rhineceps*, *Wetlugasaurus*; Watson, 1962:fig. 6; Schoch & Milner, 2000:fig. 73; Damiani, 2001:fig. 31A; Marsicano et al., 2017; Fig. 5H) or a small lateral expansion that abuts the lacrimal but that does not expand anterior to it (e.g., *Micromelerpeton*, *Micropholis*; Boy, 1995:figs. 1–2; Schoch & Rubidge, 2005:figs. 1–3; Fig. 5E); these would warrant coding as 20–1 because the lateral margin is more 'stepped' than it is 'straight'. There are also taxa in which the nasal does not contact the lacrimal at any point but does expand laterally (e.g., in *Callistomordax*, the expansion is anterior to the prefrontal; Schoch, 2008:figs. 1–3). Finally, there are taxa with straight but sharply angled lateral margins (e.g., *Trimerorhachis*, *Tupilakosaurus*; Shishkin, 1961:fig. 1; Milner & Schoch, 2013:figs. 1–2; Fig. 5A-B); these typically do not expand anterior to the lacrimal (if present), but neither are they necessarily homologous with taxa with longitudinally straight margins (e.g., some individuals of *Cochleosaurus* and *Platyrhinops*; Sequeira, 2004:figs. 1, 3–4, 6; Clack & Milner, 2010:figs. 1–2, 5, 9).

- I removed the mention of the lacrimal because the lacrimal-naris relationship is accounted for in character 219, and as noted above, some taxa lack lacrimals and would have to be treated as inapplicable if left unchanged, and others have a stepped lateral margin that does not expand anterior to the lacrimal. Therefore, taxa were recoded strictly on the basis of the lateral suture morphology. A third state is added to differentiate between sutures that are continually straight and longitudinally oriented (original and new 20–0) and sutures that are straight and obliquely oriented (the new 20–2). Taxa with slightly laterally convex sutures are considered to be 20–0.
- Codes were recoded from 20–0 to the revised 20–1 for *Proterogyrinus*, *Adamanterpeton*, *Nigerpeton*, *Balanerpeton*, *Neldasaurus*, *Isodectes*, *Tupilakosaurus*, *Limnogyrinus*, *Micromelerpeton*, *Branchierpeton*, *Branchiosaurus*, *Apateon*, *Micropholis*, *Amphibamus*, *Rhineceps*, *Broomistega*, *Chomatobatrachus*, *Sangaia*, *Plagiosuchus*, *Gerrothorax*, *Edingerella*, *Benthosuchus*, *Callistomordax*, *Metoposaurus*, *Wetlugasaurus*, *Eocyclotosaurus*, *Chinlestegophis*, *Ichthyophis*, and *Epicrionops*.
- Codes were recoded from 20–1 to the new 20–2 for *Trimerorhachis*, *Dvinosaurus*, *Acroplous*, *Thabanchuia*, *Xenopus*, *Karaurus*, and *Kokartus*.

- Codes were recoded from 20–0 to the new 20–2 for *Acanthostomatops*.
- Codes were recoded from 20–1 to 20–0 for *Archegosaurus*.
- *Cryptobranchus* was newly coded for 20–0.
- *Peltobatrachus*, *Siderops*, and *Rileymillerus* were originally coded for 20–0 but were uncoded here because this suture is not sufficiently preserved to recode (Panchen, 1959; Warren & Hutchinson, 1983; Bolt & Chatterjee, 2000). *Leptodactylus* was uncoded from 20–1 because it has a straight anteromedially angled suture; this is not accounted for in the existing states and should not be homologized with the new 20–2 for an anterolaterally angled suture.
- The revision of this character led to many new polymorphisms: *Greererpeton* (20-–0 → 20–0&1; Smithson, 1982; Godfrey, 1989); *Edops* (20–0 → 20–0&1; Romer & Witter, 1942); *Cochleosaurus* (20–0 → 20–0&1; Sequeira, 2004); *Capetus* (20–0 → 20–0&1; Sequeira & Milner, 1998); *Zatrachys* (20–0 → 20–1&2; Schoch, 1997); *Micromelerpeton* (20–0 → 20–1&2; Boy, 1995); *Platyrhinops* (20–0 → 20–0&1; Clack & Milner, 2010); *Gerobatrachus* (20–0 → 20–1/2; Anderson et al., 2008); *Broiliellus* (20–0 → 20–0&1; Schoch, 2012); *Ecolsonia* (20–0 → 20–0&1; Berman et al., 1985); *Onchiodon* (20–0 → 20–0&1; Boy, 1990); *Eryops* (20–0 → 20–0&1; Sawin, 1941); *Uranocentrodon* (20–0 → 20–0&1; Marsicano et al., 2017); *Lydekkerina* (20–0 → 20–0&1; Jeannot et al., 2006); *Peltostega* (20–0 → 20–0/1; Schoch & Milner, 2000; Janvier, 1983); *Batrachosuchus* (20–0 → 20–0&1; Schoch & Milner, 2014); *Trematosaurus* (20–0 → 20–0&1; Schoch, 2019a); *Eryosuchus* (20–0 → 20–0&1; Schoch & Milner, 2000); *Mastodonsaurus* (20–0 → 20–0&1; Schoch, 1999).

- **22 (lacrimal, lateral suture); 23 (lacrimal, position); 24 (lacrimal, posterior extension); 43 (prefrontal, process); 59 (jugal-lacrimal); 219 (lacrimal-naris); 220 (lacrimal-orbit); 221 (lacrimal, orbital processes):** any taxon coded for 21–1 (lacrimal absent) cannot be coded for these characters.
  - The most recent redescription of *Laidleria* (Warren, 1998a:171), stated that the holotype is not well-preserved posterior to the nares, and “for this reason it is not possible to determine whether septomaxillae or lacrimals are present.” However, the most recent revision of Rhytidosteidae (Dias-da-Silva & Marsicano, 2011), which involved personal examination of this taxon, did treat the lacrimal as absent (state 12–1 therein). This treatment is followed here.
  - Pardo et al. (2017a) originally coded *Rileymillerus* as lacking a lacrimal (following Bolt & Chatterjee, 2000), but this was changed to unknown by Schoch et al. (2020) without detailed rationale. The original code is restored here.
  - Character 22: Codes were corrected from 22–0 (lateral suture parallel to medial suture) for *Thabanchuia, Tupilakosaurus, Laidleria, Siderops,* and *Batrachosuchus.*
  - Character 24: Codes were corrected from 24–0 (restricted to the antorbital region) for *Thabanchuia, Tupilakosaurus, Laidleria, Siderops,* and *Batrachosuchus.*
  - Character 43: Codes were corrected from 43–0 (prefrontal-lacrimal suture simple) for *Thabanchuia, Tupilakosaurus, Laidleria, Siderops, Batrachosuchus,* and *Ichthyophis*.
  - Character 59: Codes were corrected from 59–0 (jugal-lacrimal separated by orbit or palate bones) for *Chinlestegophis*.

    - For all eight characters, any taxon originally coded as unknown was updated to inapplicable for consistency.
- **25 (orbit and naris):** state 1 (separated only by narrow gap of bone) was only coded for the dvinosaurs *Isodectes* and *Acroplous* (among temnospondyls; Fig. 6A, 6C). However, it is applicable to all trematopids (both those sampled here and those excluded; Fig. 6E, 6G), in which a posterior expansion of the naris incises into the lacrimal (18–1), creating a narrow separation of orbit and naris that is proportionately narrower than in those dvinosaurs (e.g., Berman et al., 1985, 1987, 2010, 2011; Dilkes, 1990; Sumida et al., 1998; Polley & Reisz, 2011; Milner, 2018; Gee & Reisz, 2020). The three sampled trematopids were previously coded for 25–0 (well separated).

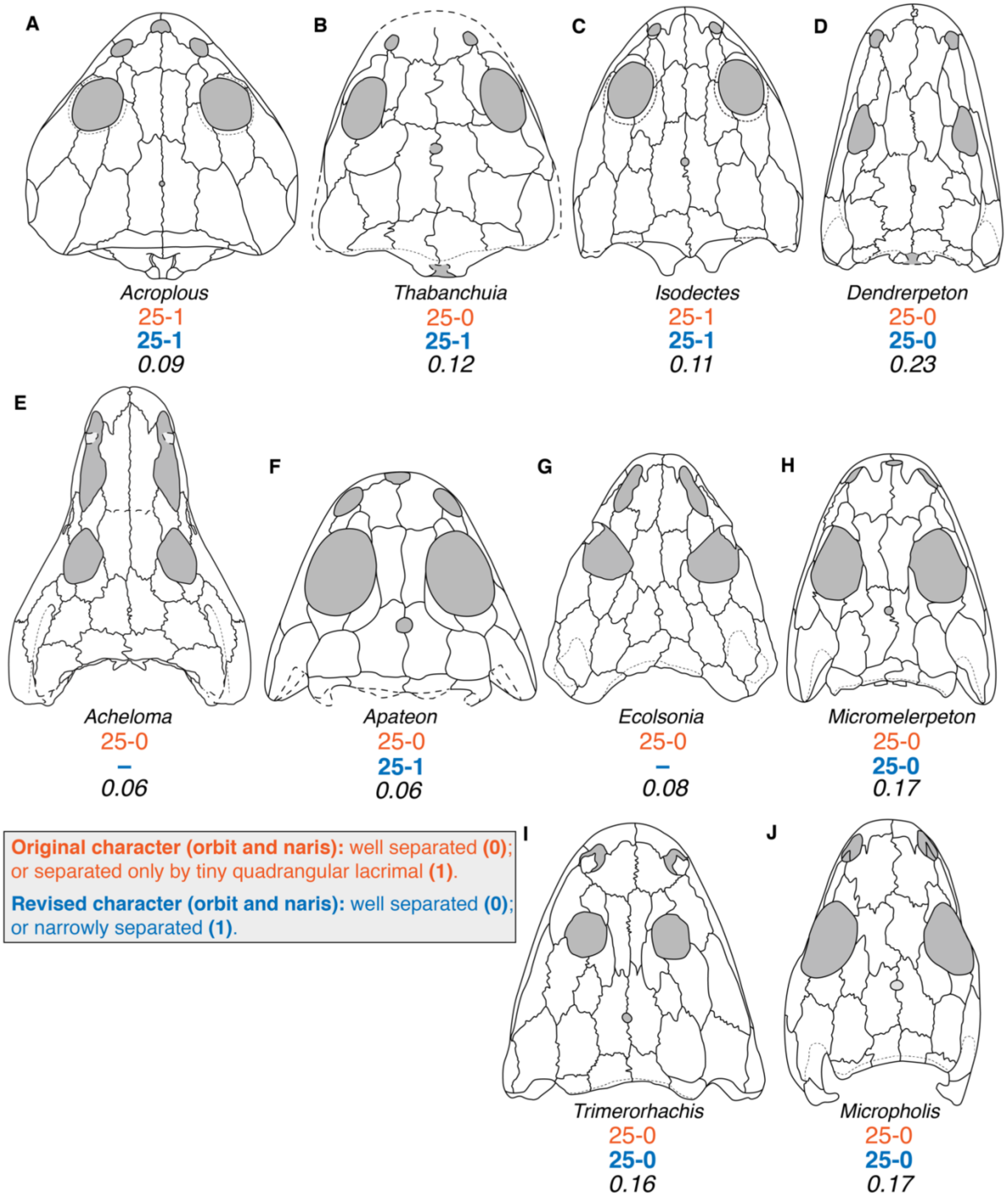

**Figure 6. Comparison of the orbit-naris separation among temnospondyls. A**, the dvinosaur *Acroplous vorax* (after Englehorn et al., 2008); **B**, the dvinosaur *Thabanchuia oomie* (after Warren, 1998b); **C**, the dvinosaur *Isodectes obtusus* (after Sequeira, 1998); **D**, the 'dendrerpetid' *Dendrerpeton acadianum* (after Holmes et al., 1998); **E**, the trematopid *Acheloma cumminsi* (after Polley & Reisz, 2011); **F**, the branchiosaurid *Apateon pedestris* (after Schoch & Milner, 2014); **G**, the trematopid *Ecolsonia cutlerensis* (after Berman et al., 1985); **H**, the micromelerpetid *Micromelerpeton credneri* (after Schoch & Milner, 2014); **I**, the dvinosaur *Trimerorhachis insignis* (after Milner & Schoch, 2013); **J**, the long-headed morph of the amphibamiform *Micropholis stowi* (after Schoch & Rubidge, 2005). Measurements represent the ratio of the length between the naris and orbit and the midline skull

length. The former is measured as the shortest straight line between the two openings, and the latter is measured from the tip of the premaxilla to the occipital margin of the postparietal. Based on the depicted examples, a ratio of 0.15 or lower might be considered as a threshold for 'narrow' separation, but this is not explicitly defined here in the character as it is semi-arbitrary. Orange text represents the original character and codes from Schoch (2013); blue text represents the revised character and codes. Silhouettes not to scale.

    - *Acheloma*, *Phonerpeton*, and *Ecolsonia* were corrected to inapplicable from 25–0 to avoid redundancy because the elongate naris that typifies this clade is the result of posterior expansion towards the orbit, thereby producing a narrow separation.
    - State 1 is also applicable to some branchiosaurids (e.g., *Apateon pedestris*; Schoch & Milner, 2008:fig. 1; Schoch & Milner, 2014:fig. 26; Fig. 6F) and plagiosaurids (e.g., *Plagiosuchus*; Damiani et al., 2009:fig. 7A). *Apateon*, *Gerrothorax*, and *Plagiosuchus* were corrected to 25–1 from 25–0. *Thabanchuia* was also corrected from 25–0 to 25–1 (Warren, 1998b:fig. 5; Schoch & Milner, 2014:fig. 20).

- **25 (orbit and naris); 35 (infraorbital sulcus); 44 (prefrontal-jugal); 45 (prefrontal-maxilla):** modified along the lines of character 20 to remove reference specifically to the lacrimal. No codes were changed.
- **28 (palpebral ossifications):** this is a complicated character to code because these ossifications, as well as the scleral ring, may be predisposed to be dislodged post-mortem. As noted in the main text, the previous codes are conspicuous because nearly all temnospondyls, including some known from singletons and those without any known scleral rings, were coded (for 28–0), but six dissorophoids were left as unknown, while most other dissorophoids were coded for 28–1. It could be reasonably inferred that they were present in these six taxa treated as unknown (i.e. coding them as 'absent' would produce hypothetically false conflicting information). While this is a reasonable approach, it must be applied uniformly, and it cannot be presumed that palpebral ossifications were absent in all other temnospondyl clades (they occur in some 'microsaurs' and the 'dendrerpetid' *Balanerpeton*, for example, and are not a dissorophoid autapomorphy). There are two approaches that can be taken: either to code the character as preserved, without considering the possibility of post-mortem loss; or to only code the character when sclerotic ring plates are at least partially preserved, as the absence of these plates would indicate that preservation was also insufficient for palpebral ossifications to be preserved, if present. The latter approach is taken here for two reasons: (1) it is more conservative and accounts for taphonomy; and (2) both states technically require scleral ring ossicles to be present to be coded.
    - The following taxa were uncoded from 28–0 (no palpebral ossifications) because the literature does not establish the presence of sclerotic ring plates in these taxa: *Proterogyrinus, Greererpeton, Adamanterpeton, Nigerpeton, Chenoprosopus, Trimerorhachis, Neldasaurus, Acroplous, Isodectes, Tupilakosaurus, Iberospondylus, Zatrachys, Eryops, Glanochthon, Australerpeton, Lapillopsis, Peltobatrachus, Rhineceps, Broomistega, Lydekkerina, Chomatobatrachus, Sangaia, Peltostega, Laidleria, Siderops, Batrachosuchus, Plagiosuchus, Gerrothorax, Watsonisuchus, Edingerella, Benthosuchus, Trematosaurus, Trematolestes, Lyrocephaliscus, Callistomordax, Metoposaurus, Parotosuchus, Paracyclotosaurus, Eocyclotosaurus,* and *Mastodonsaurus*.

- **29 (pineal foramen); 33 (lateral line sulci); 154 (ceratobranchials); 155 (basibranchial); 156 (hypobranchial elements); 188 (scapula); 191 (humerus, adult shaft); 356 (palatine):** all of these characters specifically invoke the condition of adults. This means any taxon represented only by juvenile individuals cannot be coded for these characters.
    - Codes were corrected to unknown for characters 29, 154, 156, 188, and 191, for *Gerobatrachus*, which Anderson et al. (2008:515) interpreted as represented by a holotype skeleton (the only specimen) that "belonged to a juvenile individual."
    - Codes were corrected to unknown for characters 29, 154, 155 156, 188, and 191 for *Branchiosaurus* and *Branchierpeton*, and for all of these characters except for 155 for *Limnogyrinus*; Schoch (2013:681), among others, remarked that these taxa are only known from immature specimens.
    - Codes were corrected to unknown for characters 154, 155, and 156 for *Micromelerpeton*; the hyobranchial apparatus is only known from juveniles (Boy, 1995:438–439; Witzmann, 2013:151).
- **34 (sulci, skull roof); 35 (infraorbital sulcus); 36 (supraorbital sulcus):** any taxon coded for 33–1 (lateral line sulci absent) cannot be coded for any of these characters. Codes for characters 34–36 are strange; for example, nearly all taxa coded for 33–1 were left uncoded for character 34, except for *Micropholis stowi* (34–0). This may be a missed correction from a previous matrix, as Pardo et al. (2017a), following Dilkes (2015), corrected taxa to inapplicable for this character if they were coded for 33–1.
    - Character 34: Codes were corrected from 34–0 (sulci present throughout skull roof if present) for *Micropholis*.
    - Character 35: Codes were corrected from 35–0 (infraorbital sulcus with simple curve) for *Edops, Adamanterpeton, Cochleosaurus, Chenoprosopus, Dendrerpeton, Balanerpeton, Capetus, Acanthostomatops, Zatrachys, Apateon, Micropholis, Platyrhinops, Amphibamus, Doleserpeton, Dissorophus, Cacops, Broiliellus, Acheloma, Phonerpeton, Ecolsonia, Onchiodon*, and *Eryops*.
    - Character 36: Codes were corrected from 36–0 (supraorbital sulcus passing entirely through nasal) for *Edops, Adamanterpeton, Cochleosaurus, Chenoprosopus, Dendrerpeton, Balanerpeton, Capetus, Acanthostomatops, Zatrachys, Apateon, Micropholis, Platyrhinops, Amphibamus, Doleserpeton, Dissorophus, Cacops, Broiliellus, Acheloma, Phonerpeton, Ecolsonia, Onchiodon*, and *Eryops*.
    - *Gerobatrachus hottoni* was originally coded for 36–0 but uncoded for 33, 34, and 35. This was corrected to '?' for character 36 (see above comments on coding of character 33 based on the 'adult' condition; note also that the single specimen is only exposed ventrally; Anderson et al., 2008).
    - *Branchiosaurus* was originally coded for 33–1, but because this prescribes the adult condition (see previous point), it was uncoded. However, it was coded for 34–0, 35–0, and 36–0, and regardless of the change to character 33, none of these should be coded for this taxon; all were changed to inapplicable.
    - For all three characters, all taxa originally coded as unknown were updated to inapplicable for consistency.
- **36 (supraorbital sulcus):** this character cannot be modified like characters 25 and 35 to omit the reference to the lacrimal in one state because there is a difference between some

taxa in which the groove passes onto the nasal and prefrontal but not the lacrimal (e.g., *Callistomordax, Cyclotosaurus, Parotosuchus, Uranocentrodon, Wetlugasaurus*; Schoch & Milner 2000; Sulej & Majer, 2005; Schoch, 2008b, 2018c; Fig. 7B, 7C, 7F, 7H) and others in which it passes onto these elements and the lacrimal (e.g., *Benthosuchus, Lyrocephaliscus, Mastodonsaurus, Metoposaurus, Trematosaurus*; Schoch, 1999; Schoch & Milner, 2000; Damiani, 2001; Sulej, 2007; Schoch, 2019a; Fig. 7A, 7D, 7E, 7G).

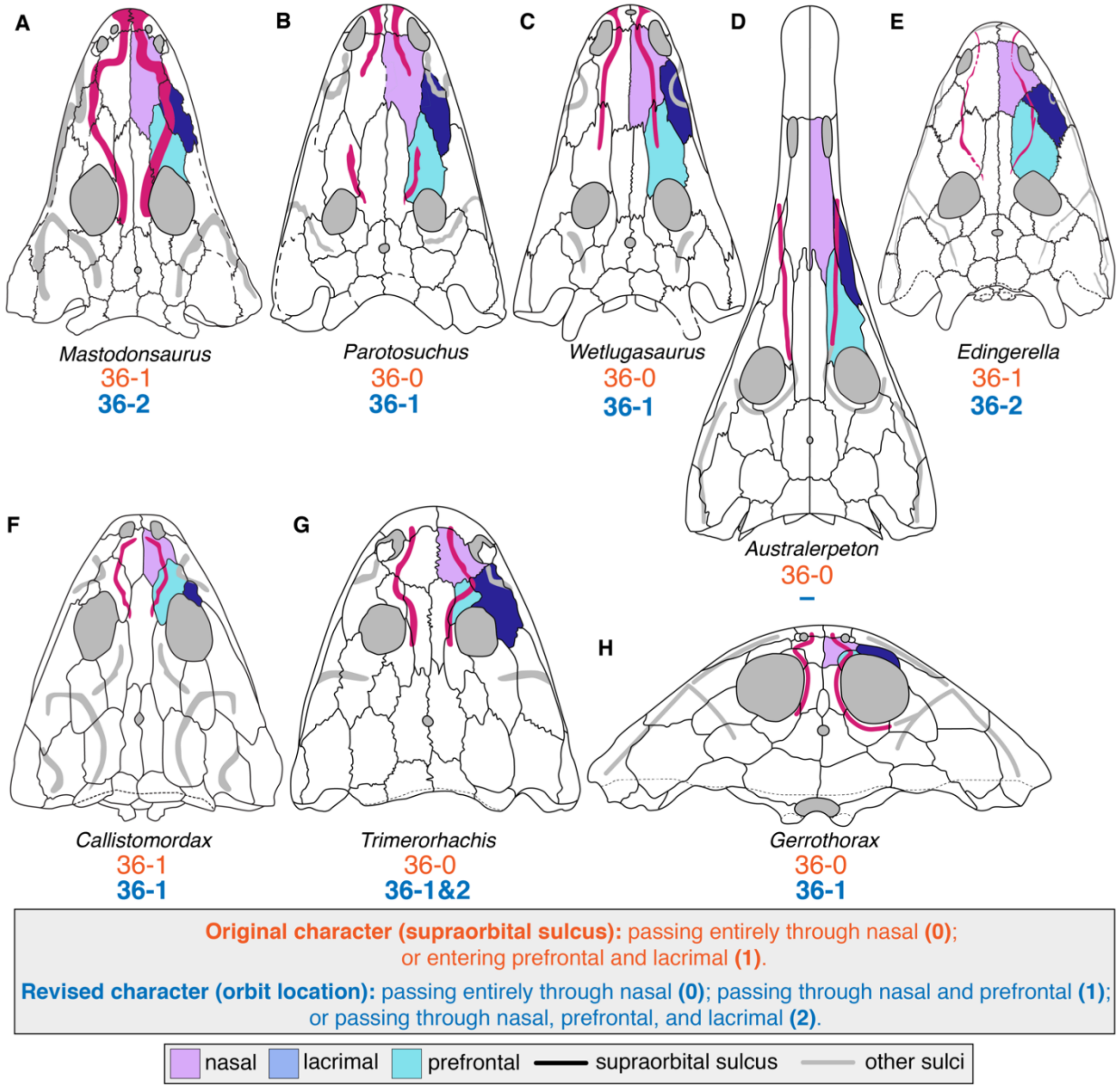

**Figure 7. Comparison of the supraorbital sulcus among temnospondyls. A**, the capitosaur *Mastodonsaurus giganteus* (after Schoch, 1999); **B**, the capitosaur *Parotosuchus nasutus* (after Schoch, 2018); **C**, the capitosaur *Wetlugasaurus angustifrons* (after Schoch, 2008); **D**, the rhinesuchid *Australerpeton cosgriffi* (after Eltink et al., 2016); **E**, the capitosaur *Edingerella madagascariensis* (after Maganuco et al., 2009); **F**, the metoposauroid *Callistomordax kugleri* (after Schoch, 2008); **G**, the dvinosaur *Trimerorhachis insignis* (after Milner & Schoch, 2013); **H**, the plagiosaurid *Gerrothorax pulcherrimus* (after Schoch & Witzmann, 2012). Orange text represents the original character and codes from Schoch (2013); blue text represents the revised character and codes. Silhouettes not to scale.

There are many existing inconsistencies in coding – for example, *Callistomordax* and *Trimerorhachis* both have a lacrimal and a supraorbital groove that passes from the nasal onto the prefrontal (but not the lacrimal; Schoch, 2008a; Milner & Schoch, 2013), but the former was coded for 36–1 and the latter for state 0 (every non-stereospondylomorph was coded for 36–0). Some of these inconsistencies stem from the construction of the states – state 0 was evidently treated as a 'catch-all' for any taxon without any lateral line grooves, with a supraorbital groove restricted to the nasal (e.g., *Nigerpeton*; Steyer et al., 2006), or with a supraorbital groove on the nasal and other elements but not the lacrimal.

- **36. Supraorbital sulcus.** Passing entirely through nasal (0), entering prefrontal and lacrimal (1). [*added by Schoch, 2013*]

Additionally, the condition in which it contacts the nasal and prefrontal but not the lacrimal is not captured. The character is modified to add a state for this. This character cannot be coded for taxa with this sulcus but without either lacrimals or prefrontals (e.g., *Laidleria*, *Peltostega*; Warren, 1998a; Schoch & Milner, 2000), as it would establish a partial dependency with other characters referring to the presence/absence of these elements. There are a handful of taxa with lacrimals, prefrontals, and a supraorbital sulcus that are not captured by any state or the original or revised character and that should also not be coded – an example is *Australerpeton*, in which the groove passes from the maxilla through the lacrimal and onto the prefrontal (no nasal contact; Eltink et al., 2016). These taxa were treated as inapplicable here (Fig. 7D), but future workers may wish to add these as states for a Bayesian analysis in which the number of taxa coded for each state is irrelevant.

- Codes were corrected from 36–0 (supraorbital sulcus passing entirely through nasal) for *Thabanchuia*, *Tupilakosaurus*, *Laidleria*, *Siderops*, *Batrachosuchus*, and *Plagiosuchus*, which either lack lacrimals (21–1) or prefrontals (314–1).
- Codes were recoded from 36–0 to the new 36–1 for *Greererpeton*, *Dvinosaurus Micromelerpeton*, *Archegosaurus*, *Uranocentrodon*, *Lydekkerina*, *Gerrothorax*, *Wetlugasaurus*, *Parotosuchus*, and *Cyclotosaurus*.
- *Neldasaurus* was newly coded for the new 36–1.
- Codes were recoded from 36–0 to the new 36–2 for *Rhineceps*, *Chomatobatrachus*, and *Paracyclotosaurus*.
- Codes were recoded from 36–1 to the new 36–2 for *Sangaia*, *Edingerella*, *Benthosuchus*, *Trematosaurus*, *Lyrocephaliscus*, *Metoposaurus*, *Eocyclotosaurus*, and *Mastodonsaurus*.
- *Trimerorhachis* was recoded from 36–1 to 36–1&2 (Milner & Schoch, 2013:fig. 2).
- *Glanochthon* was uncoded as there is no explicit description or clear figuring of this specific sulcus (Schoch & Witzmann, 2009b:126, fig. 2C); *Broomistega* was uncoded as the groove appears to be restricted to the lacrimal and prefrontal, which is not captured in the original or modified characters (Shishkin & Rubidge, 2000:659, fig. 3D); *Watsonisuchus* was uncoded because the lateral line grooves are very poorly preserved, and this groove is too incomplete to assess (Warren, 1980:26, 29).

- **39 (prefrontal, frontal):** this character did not encompass the state where the prefrontal terminates posterior to the anterior extent of the frontal (this rarely occurs and only in taxa added by Pardo et al., 2017a). A second state was added for this.

- **39. Prefrontal, frontal.** Prefrontal substantially longer than frontal (0), or ending at one level with frontal (1). [*added by Schoch, 2013*]

The language for 39–0 was changed to more clearly indicate that this refers to the relative anterior extent of these elements, not to the elements' relative length; this ambiguity may have caused some confusion based on examination of erroneous codes added by Pardo et al. (2017a); because this is not certain, these corrections are listed in the taxon-specific section.

  - Codes were updated to 39–2 for *Rileymillerus* and *Cryptobranchus*.
- **36 (supraorbital sulcus); 39 (prefrontal, frontal); 40 (prefrontal, anterior end); 41 (prefrontal, lateral suture); 42 (prefrontal-postfrontal); 43 (prefrontal, process); 44 (prefrontal-jugal); 45 (prefrontal-maxilla); 226 (prefrontal contributes to external naris)**: these characters cannot be coded for taxa coded as 314–1 (prefrontals absent).
  - Codes were corrected from 36–0 (see above); 39–0 (prefrontal substantially longer than frontal); 40–1 (prefrontal anterior end wide and blunt); 41–0 (prefrontal lateral suture straight or concave); 42–1 (prefrontal-postfrontal separated by frontal); 43–0 (prefrontal-lacrimal suture simple); 44–1 (prefrontal-jugal in contact); and 45–0 (prefrontal-maxilla separated) for *Plagiosuchus*; the code for character 226 was changed from unknown to inapplicable for consistency for this taxon.
- **41 (prefrontal, lateral suture):** the first issue with this character is that one state refers to the lateral margin of the prefrontal and the other to the medial expansion (or lack thereof) of the jugal. This creates issues for taxa without a prefrontal-jugal contact, as hypothetically both conditions could exist within a single taxon without this contact and yet be distinct from either standalone condition in a taxon with this contact.
  - **41. Prefrontal (lateral suture).** Straight or concave (0), or with marked medial projection of jugal near the orbit (1). [*added by Schoch, 2013*]

As originally coded, 41–1 (medially expanded jugal) was only coded for *Uranocentrodon*, *Rhineceps*, and *Broomistega* (this may have contributed to the exclusion of *Australerpeton* from Rhinesuchidae in the original analysis). *Uranocentrodon* is the clearest example of this condition among the three. In this taxon, the lateral margin of the prefrontal is angled posterolaterally along the contact with the lacrimal, extends straight posteriorly for much of the contact with the jugal, and turns sharply medially at the end, producing the medial expansion of the jugal (e.g., Schoch & Milner, 2000:fig. 55; Marsicano et al., 2017:fig. 4A-B; Fig. 8C). The condition in *Rhineceps* is somewhat different; from the posterior end of the lacrimal, the prefrontal simply extends in a straight suture angled posteromedially (Watson, 1962:fig. 6; Schoch & Milner, 2000:fig. 54; Marsicano et al., 2017:fig. 7; Fig. 8B). In *Broomistega*, this relationship can only be coded from juveniles and is closer to *Rhineceps*, although the suture is less angled (Shishkin & Rubidge, 2000:fig. 3; Fig. 8A). The smallest individual has a small step, but the medial turn of the suture is separated from the orbit to form a short posteromedially angled suture that then turns into a longitudinally straight suture terminating at the orbit.

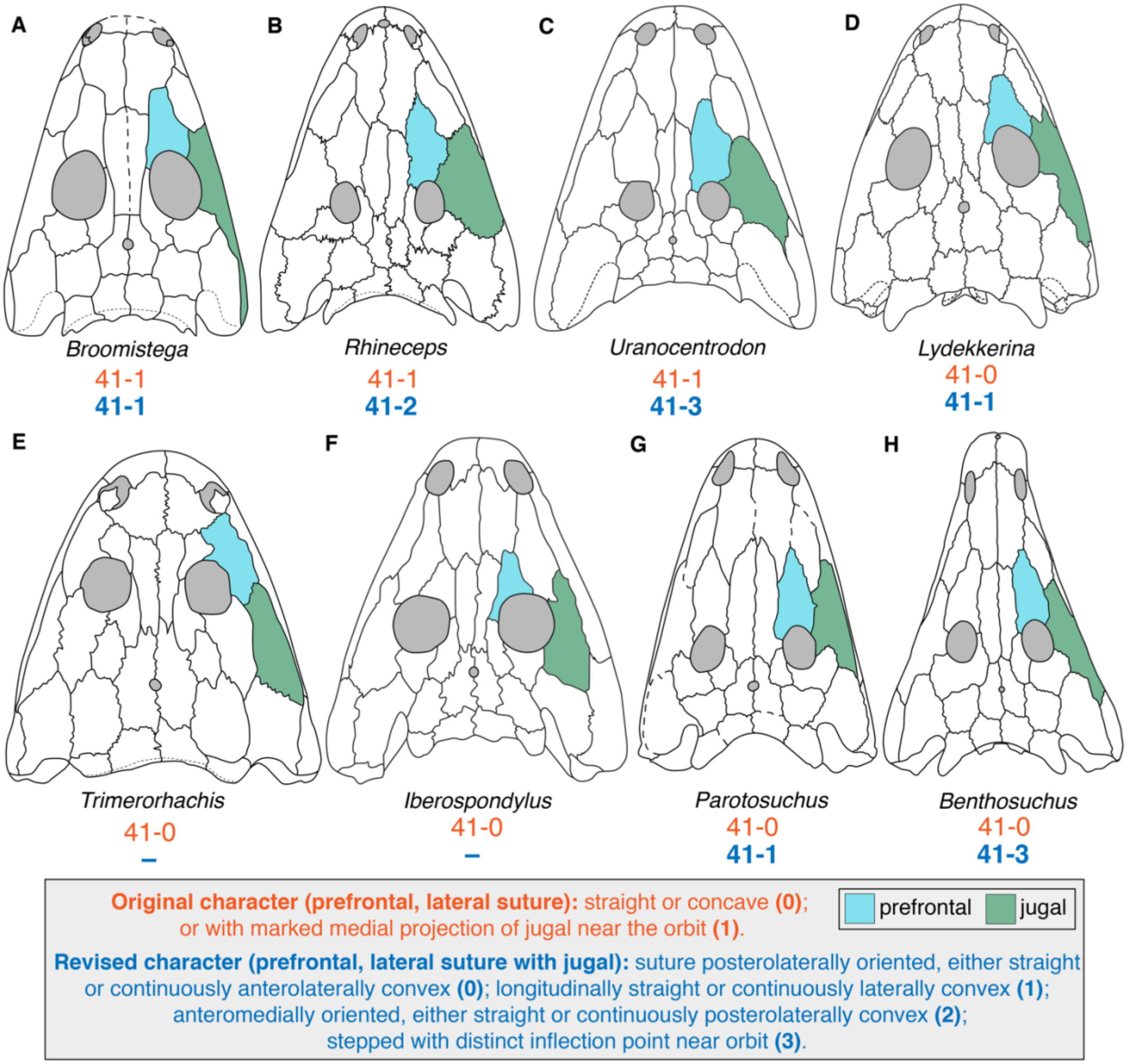

**Figure 8. Comparison of prefrontal-jugal suture morphologies among temnospondyls. A**, the rhineuschid *Broomistega putterilli* (after Shishkin & Rubidge, 2000); **B**, the rhinesuchid *Rhineceps nyasaensis* (after Schoch & Milner, 2000); **C**, the rhinesuchid *Uranocentrodon senekalensis* (after Schoch & Milner, 2000); **D**, the lydekkerinid *Lydekkerina huxleyi* (after Hewison, 2007); **E**, the dvinosaur *Trimerorhachis insignis* (after Milner & Schoch, 2013); **F**, the eryopiform *Iberospondylus schultzei* (after Schoch & Milner, 2014); **G**, the capitosaur *Parotosuchus nasutus* (after Schoch, 2018); **H**, the trematosaur *Benthosuchus sushkini* (after Schoch, 2008). Orange text represents the original character and codes from Schoch (2013); blue text represents the revised character and codes. Silhouettes not to scale.

Based on the original codes, the character was specifically about whether the jugal impinges upon the prefrontal's lateral margin. Therefore, it conceivably should not have been coded for any taxon without a jugal-prefrontal contact (44–0), as the jugal cannot impinge on an element that it does not contact (but was), or this specifier should be removed and the character merely coded on the basis of the prefrontal's lateral margin. Some taxa have markedly convex margins in the absence of a prefrontal-jugal contact (e.g., *Trimerorhachis*; Milner & Schoch, 2013:fig. 1A; *Neldasaurus*; Schoch, 2018a:fig.

3A; Fig. 8E). A convex margin in the presence of a prefrontal-jugal contact also occurs in taxa like *Lydekkerina* (e.g., Jeannot et al., 2006:fig. 2A; Fig. 8D).

Then there is the issue that the character was not properly coded for non-rhinesuchids. The condition of *Broomistega* is not unlike that of some individuals of *Cochleosaurus* (Sequeira, 2004:figs. 1, 6) and *Australerpeton* (e.g., Eltink et al., 2016:fig. 5). The same condition as in *Uranocentrodon* also occurs in *Iberospondylus* (Laurin & Soler-Gijón, 2001:fig. 1, 2006:fig. 2; Fig. 8F), *Eryops* (e.g., Schoch & Milner, 2014:fig. 41G), *Benthosuchus* (e.g., Schoch & Milner, 2000:fig. 74; Fig. 8H), *Watsonisuchus* (Warren, 1980:figs. 2, 4), and *Paracyclotosaurus* (e.g., Watson, 1958:fig. 5). To a lesser degree, it is also found in *Nigerpeton* (Steyer et al., 2006:figs. 2–3), *Edingerella* (e.g., Maganuco et al., 2009:fig. 6), *Wetlugasaurus* (e.g., Schoch & Milner, 2000:fig. 73), and *Eocyclotosaurus* (e.g., Schoch, 2000:fig. 1; Rinehart et al., 2015:fig. 2); the posterolateral step forms an obtuse angle rather than a right angle on account of the suture being angled rather than mostly transverse. In general, the condition like that of *Uranocentrodon* is common among capitosaurs (e.g., Schoch & Milner, 2000:figs. 92–104).

In an attempt to maintain the "spirit" of the original character, it is first restricted to taxa with a prefrontal-jugal contact and the specifier of the jugal medially expanding is maintained. The character is then specified to specifically the prefrontal-jugal suture and therefore excludes the lateral suture with the lacrimal (if present, or with a substituting element); this mitigates issues related to the complexity of the lateral suture when assessed in its entirety. There are four main types of suture that can be discerned within these confines: (1) posterolaterally angled margin, either straight or continuously anterolaterally convex (e.g., *Trematosaurus*; Schoch, 2019a:figs. 2, 4–6); (2) longitudinally straight or continuously laterally convex (e.g., *Sclerocephalus*, *Glanochthon*; Schoch & Witzmann, 2009a:figs. 4–5; 2009b:fig. 2) ; (3) straight, anteromedially angled margin (e.g., *Rhineceps*); and (4) longitudinally straight with distinct inflection point posteriorly near the orbit (e.g., *Uranocentrodon*, *Benthosuchus*, *Watsonisuchus*). These are defined as the new states 0, 1, 2, and 3, respectively, to form a new unordered multistate character, and all taxa were recoded.

- Codes were recoded from 41–0 to inapplicable for the following taxa that lack a prefrontal-jugal contact: *Proterogyrinus, Greererpeton, Dendrerpeton, Balanerpeton, Trimerorhachis, Neldasaurus, Dvinosaurus, Acroplous, Isodectes, Thabanchuia, Tupilakosaurus, Capetus, Iberospondylus, Acanthostomatops, Zatrachys, Micromelerpeton, Limnogyrinus, Branchierpeton, Branchiosaurus, Apateon, Micropholis, Platyrhinops, Amphibamus, Doleserpeton, Gerobatrachus, Dissorophus, Cacops, Broiliellus, Acheloma, Phonerpeton, Ecolsonia* and *Callistomordax.*
- *Karaurus, Ambystoma, Hynobius, Cryptobranchus, Ichthyophis,* and *Eocaecilia* were left as unknown for character 44 because they lack jugals; they were updated to inapplicable for this character and all changed from 41–0 to inapplicable.
- *Gerrothorax* was originally coded for 44–1 but should be coded for 44–0; the lacrimal widely separates these elements (Schoch & Witzmann, 2012). It was then changed from 41–0 to inapplicable.
- *Peltobatrachus* was properly coded as unknown for character 44, so it was changed from 41–0 to unknown.

    - *Chinlestegophis* and *Rileymillerus* were oddly coded as unknown for character 44 (the elements are well-separated by a LEP), so they were coded for 44–0 and changed from 41–0 to inapplicable.
- **42 (prefrontal-postfrontal):** a large number of taxa originally coded for this character by Schoch (2013) were coded for the wrong state.
    - **42. Prefrontal-postfrontal.** In contact (0), or separated by frontal (1). [*added by Schoch, 2013*]

Given that more than two dozen taxa were erroneously coded, this does not seem to be a random typographic error, but some taxa were correctly coded (i.e. this is not an instance in which coded states have merely been inverted from the listed character).

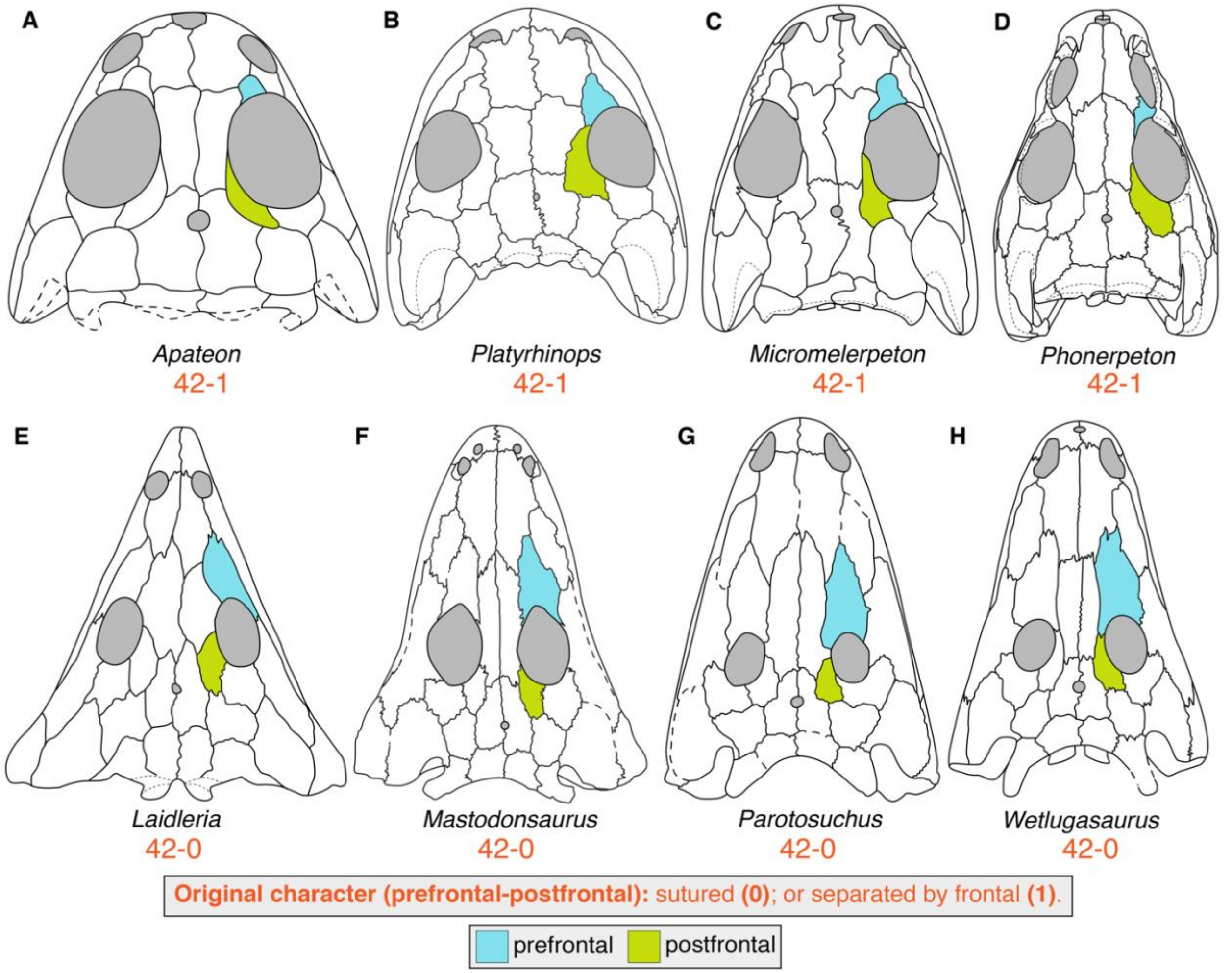

**Figure 9. Comparison of prefrontal-postfrontal sutural relationships among temnospondyls. A**, the branchiosaurid *Apateon pedestris* (after Schoch & Milner, 2014); **B**, the amphibamiform *Platyrhinops lyelli* (after Clack & Milner, 2010); **C**, the micromelerpetid *Micromelerpeton credneri* (after Schoch & Milner, 2014); **D**, the trematopid *Phonerpeton pricei* (after Dilkes, 1990); **E**, the rhytidosteid *Laidleria gracilis* (after Warren, 1998a); **F**, the capitosaur *Mastodonsaurus giganteus* (after Schoch, 1999); **G**, the capitosaur *Parotosuchus nasutus* (after Schoch, 2018c); **H**, the capitosaur *Wetlugasaurus angustifrons* (after Schoch, 2008b). Orange text represents the original character and codes from Schoch (2013). Silhouettes not to scale.

State 1 ('separated by frontal') occurs in the majority of dissorophoids (but not in early-diverging Carboniferous taxa) and most capitosaurs (Fig. 9A-D, 9F-G). All dissorophoids were originally coded for 42–1, even though Carboniferous taxa like *Limnogyrinus*,

*Branchiosaurus*, and *Platyrhinops* have a prefrontal-postfrontal contact (e.g., Clack & Milner, 2010; Schoch & Milner, 2014; Schoch & Witzmann, 2018; Fig. 9B). Conversely, all capitosaurs were originally coded for state 0 ('in contact'), even though the majority of capitosaurs, both those sampled and in general, lack this contact (e.g., Schoch, 1999, 2008b, 2018c; Schoch & Milner, 2000; Damiani, 2001; Fig. 9F-G), a feature separating them from most other stereospondyls (but not from *Laidleria*; Fig. 9E).

- o Codes were corrected from 42–0 to 42–1 for *Laidleria*, *Edingerella*, *Watsonisuchus*, *Parotosuchus*, *Eryosuchus*, *Paracyclotosaurus*, *Cyclotosaurus*, and *Mastodonsaurus*.
- o Codes were corrected from 42–1 to 42–0 for *Greererpeton*, *Trimerorhachis*, *Dvinosaurus*, *Acroplous*, *Isodectes*, *Thabanchuia*, *Tupilakosaurus*, *Acanthostomatops*, *Zatrachys*, *Limnogyrinus*, *Branchiosaurus*, *Platyrhinops*, *Amphibamus*, *Onchiodon*, *Eryops*, *Siderops*, and *Batrachosuchus*.

- **42 (prefrontal-postfrontal); 50 (postorbital, postfrontal); 61 (intertemporal and postorbital); 218 (postfrontal shape)**: these characters cannot be coded for taxa coded for 315–1 (postfrontals absent).
    - o Codes were corrected from 42–1 (see above); 50–0 (postorbital and postfrontal shorter than supratemporal and parietal); and 61–0 (postorbital lateral to postfrontal) for *Plagiosuchus*; the code for character 218 was changed from unknown to inapplicable for consistency for this taxon.
- **47 (postorbital, shape); 48 (postorbital, end); 49 (postorbital); 50 (postorbital, postfrontal); 61 (intertemporal and postorbital); 231 (shape of postorbital); 232 (parietal-postorbital contact):** these characters cannot be coded for any taxon coded as 230–1 (postorbital absent).
    - o Codes were corrected from 47–0 (postorbital long triangular, wedged deeply between squamosal and supratemporal); 48–0 (posterior end acutely triangular); 49–0 (not wider than orbit); 50–0 (see above); 61–0 (see above) for *Plagiosuchus*; the codes for characters 231 and 232 were changed from unknown to inapplicable for consistency for this taxon.
    - o *Epicrionops* and *Ichthyophis* were corrected from 47–1 and 50–0 because there is no postorbital in these taxa (correctly coded as 230–1; character 48 was originally uncoded for these taxa). *Eocaecilia* was corrected from 50–0 on the same basis; it was not originally coded for character 47.
    - o Codes for these seven characters were updated from unknown to inapplicable for *Triadobatrachus*, *Leptodactylus*, *Xenopus*, *Karaurus*, *Ambystoma*, *Hynobius*, *Cryptobranchus*, *Eocaecilia*, *Epicrionops*, and *Ichthyophis* for consistency.
- **47 (postorbital, shape); 231 (shape of postorbital):** these binary characters, refer to the same attribute of the postorbital (character definitions shown below).
    - o **47. Postorbital (shape)**. Long triangular, wedged deeply between squamosal and supratemporal (0), or short (1). [*added by Schoch, 2013*]
    - o **231. Shape of postorbital**. Irregular trapezoid (0), triangular, apex caudal (1). [*added by Pardo et al., 2017a*]

Character 47 is problematic because 47–1 ('short') is not a shape but rather refers to some aspects of proportions (inferred here to be the sagittal length relative to the transverse width).

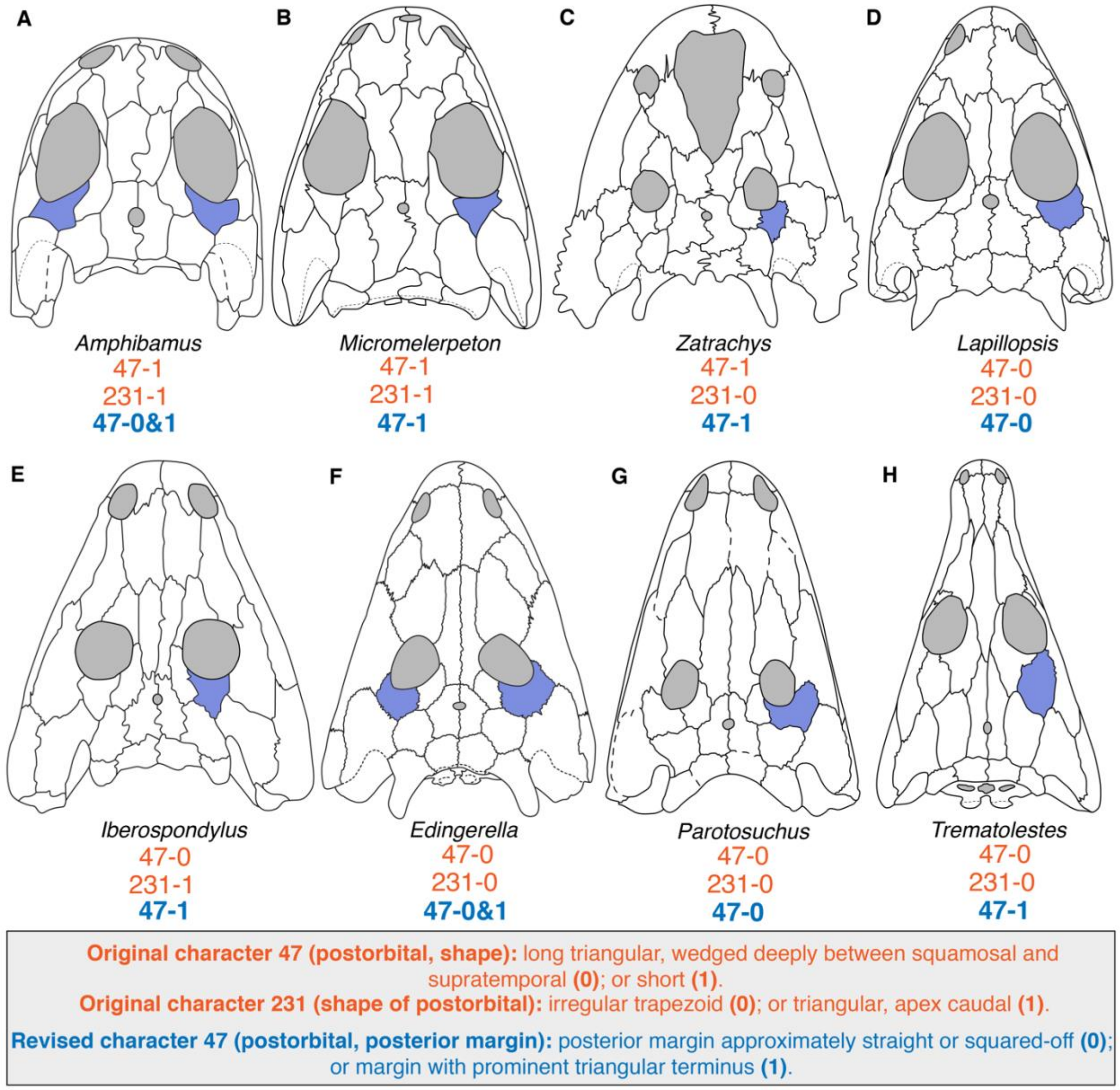

**Figure 10. Comparison of the posterior margin of the postorbital among temnospondyls. A**, the amphibamiform *Amphibamus grandiceps* (after Schoch & Milner, 2014); **B**, the micromelerpetid *Micromelerpeton credneri* (after Schoch & Milner, 2014); **C**, the zatracheid *Zatrachys serratus* (after Schoch & Milner, 2014); **D**, the stereospondyl *Lapillopsis nana* (after Yates, 2000); **E**, the eryopiform *Iberospondylus schultzei* (after Schoch & Milner, 2014); **F**, the capitosaur *Edingerella madagascariensis* (after Maganuco et al., 2009); **G**, the capitosaur *Parotosuchus nasutus* (after Schoch, 2018); **H**, the trematosaur *Trematolestes hagdorni* (after Schoch, 2006). Both postorbitals are colored in *A. grandiceps* and *E. madagascariensis* to depict the polymorphism. Orange text represents the original character and codes from Schoch (2013) and Pardo et al. (2017a); blue text represents the revised character and codes. Silhouettes not to scale.

State 1 occurred only in all zatracheids, all dissorophoids, and *Lapillopsis* in the original matrix (Fig. 10A-D; this may be one reason why *Lapillopsis* was spuriously recovered close to Dissorophoidea + Zatracheidae in previous analyses). Its approximate equivalent state (231–0) was coded in the vast majority of taxa sampled by Pardo et al. (2017a); 231–1 was coded for ‘dendrerpetids,’ *Acanthostomatops* (but not *Zatrachys*), *Capetus*, micromelerpetids, branchiosaurids, and amphibamids. On balance, the coding of character 231 seems more accurate when comparing taxa with conflicting codes. For example, *Micromelerpeton credneri* has a distinctly triangular postorbital that markedly divides the supratemporal and the squamosal and therefore should have been coded for 47–0 (231–1 is more accurate; e.g., Schoch & Milner, 2014:fg. 27A; Fig. 10B). Character 47 is therefore replaced by character 231, which is then modified to clarify the language (the differentiation in shape is essentially related only to the posterior margin). All taxa were then recoded. Character 231's position was not replaced or modified (omitted); the relative length of the postorbital is already partially captured in character 50, so no substitute character is proposed here.

- Codes were recoded from 47–0 to the revised 47–1 for *Proterogyrinus, Greererpeton, Edops, Cochleosaurus, Nigerpeton, Chenoprosopus, Dendrerpeton*, *Balanerpeton, Trimerorhachis, Neldasaurus, Acroplous, Isodectes, Capetus*, *Iberospondylus, Onchiodon, Eryops, Sclerocephalus, Glanochthon, Archegosaurus*, *Platyoposaurus, Peltobatrachus, Chomatobatrachus, Laidleria, Benthosuchus, Trematosaurus, Trematolestes, Lyrocephaliscus, Callistomordax, Metoposaurus*, *Eryosuchus, Eocyclotosaurus, Cyclotosaurus,* and *Mastodonsaurus*.
- Codes were recoded from 47–1 to the revised 47–0 for *Platyrhinops, Cacops, Broiliellus, Acheloma, Phonerpeton,* and *Lapillopsis*.
- *Amphibamus* was recoded from 47–1 to 47–0&1 (Schoch & Milner, 2014:fig. 30A); *Tupilakosaurus, Lydekkerina,* and *Edingerella* were recoded from 47–0 to 47–0&1 (Shishkin, 1961:fig. 1; Steyer, 2003:figs. 1–2, 6; Jeannot et al., 2006:figs. 2, 6–7; Maganuco et al., 2009:fig. 6; Schoch & Milner, 2014:fig. 20I).

- **47 (postorbital, shape); 48 (postorbital end):**
    - **47. Postorbital (shape)**. Long triangular, wedged deeply between squamosal and supratemporal (0), or short (1). [*added by Schoch, 2013*]
    - **48. Postorbital (end)**. Posterior end acutely triangular (0), or with finger like projection (1). [*added by Schoch, 2013*]

State 48–1 was coded for all edopoids and for no other temnospondyls. This condition is exemplified by *Cochleosaurus* (e.g., Sequeira, 1998:figs. 2, 6; Fig. 11C) and *Adamanterpeton* (Milner & Sequeira, 1998:figs. 6, 9; Fig. 11D). It is (presumably) differentiated from 48–0 (acutely triangular) by the abrupt narrowing posteriorly. *Chenoprosopus* and *Nigerpeton* do not have this finger-like process, despite being coded as such (Langston, 1953:fig. 5; Steyer et al., 2006:figs. 2–3; Figs. 11A-B). The condition may be variable in smaller individuals of *Chenoprosopus* (compare Hook, 1993:fig. 1, with Reisz et al., 2005:fig. 2), but the largest individuals described by Langston (see also Schoch & Milner, 2014:fig. 13E) do not have the abrupt tapering found in other edopoids. The condition of *Dendrerpeton* (Holmes et al., 1998:fig. 4A; Fig. 11E; originally coded for 48–0) is closer to that of *Cochleosaurus* or *Adamanterpeton,* while the condition of *Chenoprosopus* and *Nigerpeton* is closer to that of most other non-stereospondyls.

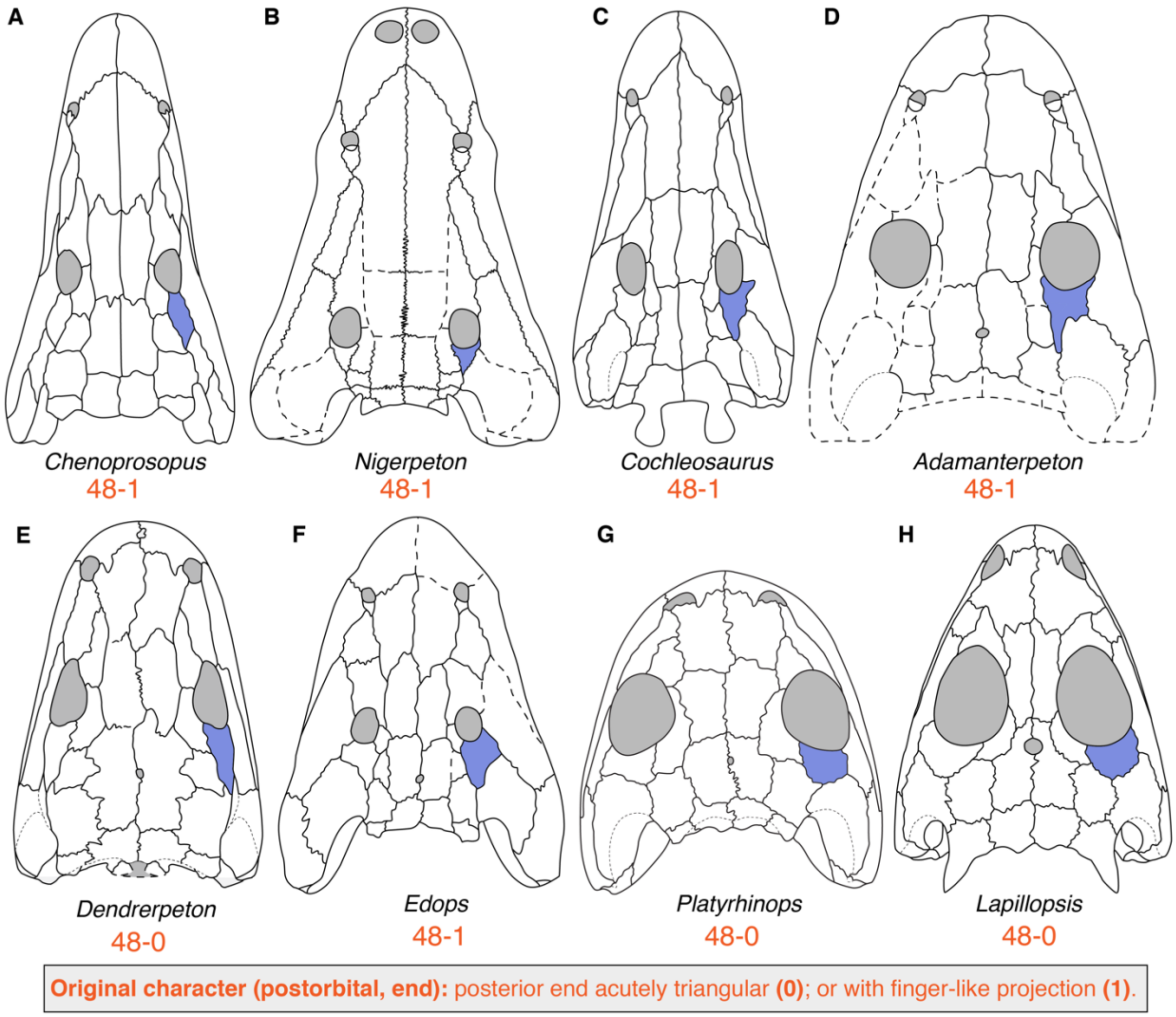

**Figure 11. Comparison of the posterior process of the postorbital among temnospondyls. A**, the edopoid *Chenoprosopus milleri* (after Schoch & Milner, 2014); **B**, the edopoid *Nigerpeton ricqlesi* (after Steyer et al., 2006); **C**, the edopoid *Cochleosaurus bohemicus* (after Sequeira, 2004); **D**, the edopoid *Adamanterpeton ohioensis* (after Milner & Sequeira, 1998); **E**, the ‘dendrerpetid’ *Dendrerpeton acadianum* (after Holmes et al., 1998); **F**, the edopoid *Edops craigi* (after Schoch & Milner, 2014); **G**, the amphibamiform *Platyrhinops lyelli* (after Clack & Milner, 2010); **H**, the stereospondyl *Lapillopsis nana* (after Yates, 2000). Orange text represents the original character and codes from Schoch (2013). Silhouettes not to scale.

    - Any taxon coded for 48–0 will be coded for 47–1 (the inverse is not true). To avoid this dependency, and because character 48 only refers to a more specific aspect of the same general feature (the posterior margin of the postorbital), these characters are merged. Character 48 is not modified or replaced. Any taxon coded for 48–1 would automatically be coded for the new 47–2.
        - *Chenoprosopus* and *Nigerpeton* would have been recoded to the new 47–2, but because they lack the same anatomy in other edopoids, they were left as 47–1.
- **50 (postorbital, postfrontal); 53 (supratemporal); 54 (supratemporal, width); 56 (semilunar flange); 217 (supratemporal exposure on occiput)**: these characters cannot be coded for any taxon coded as 216–1 (supratemporal absent).

    - Character 50: Codes were corrected from 50–0 for *Ichthyophis* and *Epicrionops*.
    - Character 56: Codes were corrected from 56–0 (supratemporal without ventral projection into otic notch) for *Leptodactylus* and *Xenopus.*
    - Codes for these five characters were updated from unknown to inapplicable for the above taxa (when originally coded as unknown) as well as for *Triadobatrachus, Ichthyophis*, and *Epicrionops* for consistency.
- **52 (otic notch, position); 55 (supratympanic flange); 56 (semilunar flange); 229 (large otic notch approaching orbit):** any taxon coded for 51–1 (otic notch absent, straight transverse posterior skull margin) cannot be coded for these characters (as was properly applied originally for character 52: otic notch position).
    - Character 55: Codes were corrected from 55–0 (squamosal continuously ornamented around margin of otic notch) for *Greererpeton, Dvinosaurus, Acroplous, Isodectes, Thabanachuia, Tupilakosaurus, Sangaia, Laidleria, Batrachosuchus, Plagiosuchus*, and *Gerrothorax*.
    - Character 56: Codes were corrected from 56–0 (supratemporal without ventral projection into otic notch) for *Greererpeton, Dvinosaurus, Acroplous, Isodectes, Thabanachuia, Tupilakosaurus, Sangaia, Laidleria, Batrachosuchus, Plagiosuchus*, and *Gerrothorax*.
    - Character 229: Codes were corrected from 229–0 (large otic notch approaching orbit absent) for *Greererpeton, Acroplous, Isodectes, Sangaia, Laidleria, Batrachosuchus, Plagiosuchus, Gerrothorax, Rileymillerus, Ambystoma, Hynobius, Cryptobranchus, Ichthyophis*, and *Epicrionops*. Note that 229–0 is not redundant with 51–1; 229–0 was coded for many taxa with otic notches, including most temnospondyls.
    - Codes for these four characters were updated from unknown to inapplicable for the above taxa (when originally coded as unknown) as well as for *Karaurus* and *Kokartus* for consistency.
- **53 (supratemporal, proportions):**
    - **53. Supratemporal (proportions).** Longer than wide (0), or quadrangular, giving a foreshortened posterior skull table (1). [*added by Schoch, 2013*]

Originally, the majority of temnospondyls were coded for 53–0 (Fig. 12). However, a modest number of taxa have subquadratic supratemporals but were not coded for 53–1.

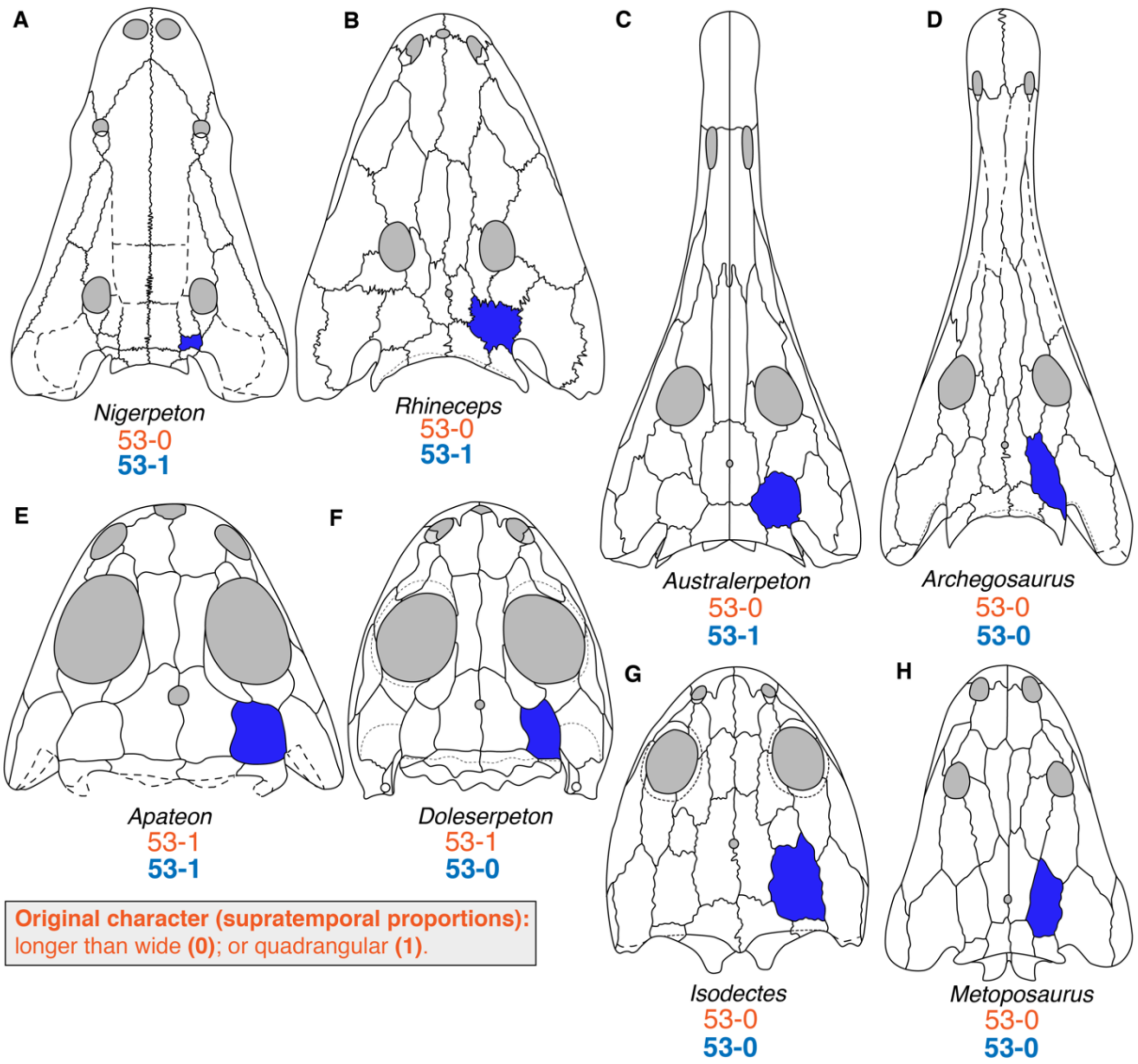

**Figure 12. Comparison of supratemporal proportions among temnospondyls.** **A**, the edopoid *Nigerpeton ricqlesi* (after Steyer et al., 2006); **B**, the rhinesuchid *Rhineceps nyasaensis* (after Schoch & Milner, 2000); **C**, the rhinesuchid *Australerpeton cosgriffi* (after Eltink et al., 2016); **D**, the stereospondylomorph *Archegosaurus decheni* (after Witzmann, 2005); **E**, the branchiosaurid *Apateon pedestris* (after Schoch & Milner, 2014); **F**, the amphibamid *Doleserpeton annectens* (after Sigurdsen & Bolt, 2010); **G**, the dvinosaur *Isodectes obtusus* (after Sequeira, 1998); **H**, the metoposaurid *Metoposaurus krasiejowensis* (after Sulej, 2007). Orange text represents the original character and codes from Schoch (2013). Silhouettes not to scale.

This discrepancy may be a consequence of the compound nature of 53–1, which also specified a shortened posterior skull table (even though this is accounted for in characters 9 and 46). This specifier is therefore removed to ensure that the character is only coded based on the proportions of the supratemporal.

- Codes were corrected from 53–0 to 53–1 for *Nigerpeton*, *Chenoprosopus* (this is an ontogenetic transformation; compare Langston, 1953:figs. 5–7, with Hook, 1993:fig. 1, and Reisz et al., 2005:fig. 2), *Dvinosaurus*, *Thabanchuia*, *Tupilakosaurus*, *Platyrhinops*, *Australerpeton*, *Lapillopsis* (this is an ontogenetic transformation; Yates, 1999:fig. 2), *Rhineceps*, *Parotosuchus*, and *Cyclotosaurus*.

    - Codes were corrected from 53–1 to 53–0 for *Doleserpeton*, *Dissorophus*, *Broiliellus*, and *Eocaecilia*.
    - *Laidleria* and *Batrachosuchus* were recoded from 53–0 to 53–0&1 (Warren, 1998a:figs. 2, 4; Schoch & Milner, 2014:fig. 46E).
- **54 (supratemporal, width):**
    - **54. Supratemporal (width).** Rectangular, with straight sagittal lateral margin (0), or posterolaterally constricted by expanded otic notch (1). [*added by Schoch, 2013*]

This is another character where the states are not clearly opposing or mutually exclusive, creating ambiguity in the coding. State 1 was originally only coded for *Platyrhinops*, *Amphibamus*, and *Karaurus* (Fig. 13A-B). However, a number of other taxa also have a posterolateral incision in the supratemporal (e.g., the edopoids *Edops*, *Adamanterpeton*, *Chenoprosopus*; Romer & Witter, 1942:fig. 1; Langston, 1953:figs. 5–7; Milner & Sequeira, 1998:figs. 6, 8; Schoch & Milner, 2014:fig. 13; Figs. 13C-D).

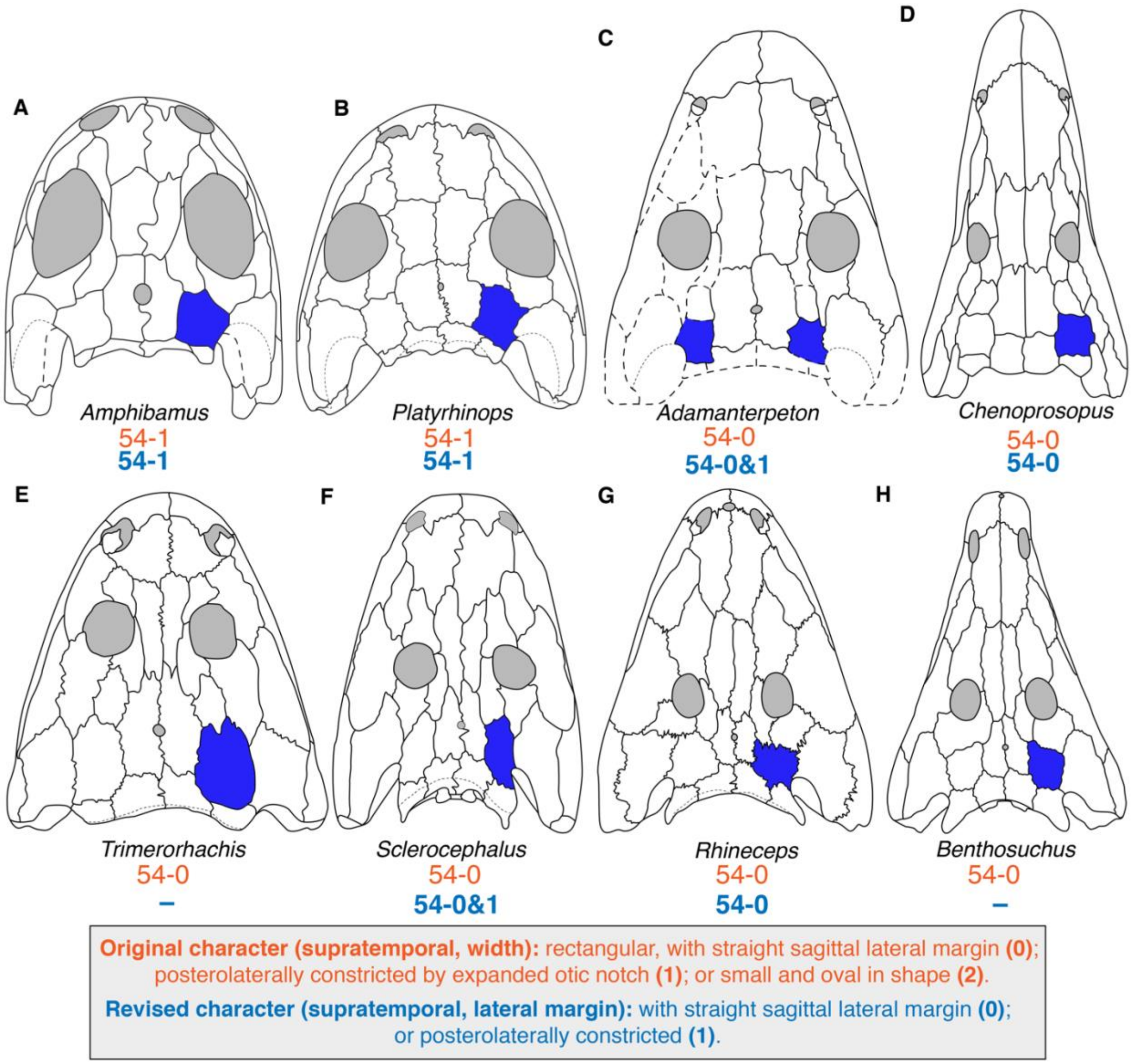

**Figure 13. Comparison of the posterolateral margin of the supratemporal among temnospondyls. A**, the amphibamiform *Amphibamus grandiceps* (after Schoch & Milner, 2014); **B**, the amphibamiform *Platyrhinops lyelli* (after Clack & Milner, 2010); **C**, the edopoid *Adamanterpeton ohioensis* (after Milner & Sequeira, 1998); **D**, the edopoid *Chenoprosopus milleri* (after Schoch & Milner, 2014); **E**, the dvinosaur *Trimerorhachis insignis* (after Milner & Schoch, 2013); **F**, the stereospondylomorph *Sclerocephalus haeuseri* (after Schoch & Witzmann, 2009a); **G**, the rhinesuchid *Rhineceps nyasaensis* (after Schoch & Milner, 2000); **H**, the trematosaur *Benthosuchus sushkini* (after Schoch, 2008). Both supratemporals are colored in *A. ohioensis* to depict the polymorphism. Orange text represents the original character and codes from Schoch (2013); blue text represents the revised character and codes. Silhouettes not to scale.

Additionally, the inclusion of ‘by an expanded otic notch’ in this state creates issues because the character should then not be coded for any taxon without a notch (but was). Even if this specifier is removed, 54–0 would be guaranteed by 62–1 (squamosal-tabular sutured), as the supratemporal might narrow slightly in width posterolaterally, but there is no taxon with a marked medial expansion of the squamosal to produce a ‘constricted’ lateral margin in the fashion found in taxa originally coded for 54–1. Indeed, every taxon coded for 62–1 was coded for 54–0 (but future workers might consider the condition of

*Parotosuchus*, with a squamosal-tabular contact, to meet the criterion for a constricted supratemporal and to thus dispel this dependency; Schoch, 2018c:fig. 3B). Neither the presence of an otic notch (51–0) nor a squamosal-tabular separation (62–0) guarantees a constricted lateral margin.

  - The specifier of the otic notch was removed since it is functionally assured even without its inclusion and to avoid issues in taxa without an otic notch.
  - Any taxon coded for 62–1 was recoded to inapplicable: *Trimerorhachis, Neldasaurus, Dvinosaurus, Isodectes, Thabanchuia, Tupilakosaurus, Acanthostomatops, Zatrachys, Lydekkerina, Chomatobatrachus, Sangaia, Peltostega, Laidleria, Batrachosuchus, Plagiosuchus, Gerrothorax, Edingerella, Watsonisuchus, Benthosuchus, Trematosaurus, Trematolestes, Lyrocephaliscus, Callistomordax, Metoposaurus, Wetlugasaurus, Parotosuchus, Eryosuchus, Paracyclotosaurus, Eocyclotosaurus, Cyclotosaurus, Mastodonsaurus, Chinlestegophis*, and *Eocaecilia*. For *Chinlestegophis* and *Eocaecilia*, this dependency is a second reason why the addition of a state 2 for 'small and oval' by Pardo et al. (2017a) was overturned.
  - *Greererpeton* was originally coded for 62–0 but lacks an otic notch and has a broad squamosal-tabular contact (Smithson, 1982:44; Godfrey, 1989). It was corrected to 62–1 and accordingly recoded to inapplicable for character 54.
  - *Edops, Nigerpeton, Chenoprosopus, Dendrerpeton, Balanerpeton, Australerpeton,* and *Rhineceps* were corrected from 54–0 to 54–1.
  - *Adamanterpeton* and *Sclerocephalus* were recoded to 54–0&1 (Milner & Sequeira, 1998:figs. 6, 9; Schoch & Witzmann, 2009a:figs. 4–5).

- **61 (intertemporal and postorbital); 232 (parietal-postorbital):** the language for character 61 was somewhat convoluted, but it expressed the same feature as character 232 – postorbital-parietal contact.
  - **61. Intertemporal and postorbital.** Postorbital lateral to postfrontal (0), or expanding medially to replace intertemporal, contacting parietal (1). [*added by Schoch, 2013*]
  - **232. Parietal-postorbital contact.** Absent (0), present (1). [*added by Pardo et al., 2017a*]

  This contact occurs in *Greererpeton* (originally coded as 61–0 and 232–1); *Dvinosaurus, Thabanchuia,* and *Tupilakosaurus* (originally coded as such, 61–1 and 232–1); *Chomatobatrachus* (originally coded as 61–0 and 232–0&1); and is polymorphic in *Metoposaurus krasiejowensis* (Sulej, 2007:figs. 4–15). The discrepancy for *Greererpeton* relates to the compound nature of 61–1, which also invoked an absent intertemporal; in this taxon, the contact occurs in the presence of the intertemporal, which is itself polymorphic. The discrepancy for *Chomatobatrachus* relates to discrepancies between Cosgriff (1974:fig. 31), who figured a broad contact on one side of the holotype and a point contact on the other, and Warren et al. (2006:fig. 4), who indicated more uncertainty in the general sutures and who depicted a wide separation on both sides. Character 61 is maintained in its position but is replaced by the wording of character 232 for clarity, and character 232 was not modified or replaced by a new character.
  - *Greererpeton* was recoded to 'contact present' (the old and new 61–1); *Chomatobatrachus* is maintained as 'contact absent' (the old and new 61–0); and the newly substituted *M. krasiejowensis* is coded as polymorphic. All other codes

were unchanged as the character states are directly equivalent (i.e. both 61–1 and 232–1 were for parietal-postorbital contact).

- **62 (squamosal-tabular); 63 (tabular and squamosal); 65 (tabular, horn); 66 (tabular, extension); 67 (tabular, ventral crest); 71 (posterior skull rim); 234 (parietal-tabular):** these characters cannot be coded for taxa coded for 239–1 (tabular absent; exclusive to lissamphibians).
    - Character 63: Codes were corrected from 63–0 (tabular and squamosal forming either squamosal embayment or straight posterior margin) for *Karaurus*, *Kokartus*, *Ambystoma*, *Hynobius*, and *Cryptobranchus*.
    - Character 66: Codes were corrected from 66–0 (tabular horn pointing posteriorly if present) for *Greererpeton, Acroplous, Isodectes, Thabanchuia, Tupilakosaurus, Batrachosuchus, Plagiosuchus,* and *Gerrothorax*.
    - Character 71: Codes were corrected from 71–0 (quadrate trochlea posterior to tabular horns) for *Triadobatrachus.* Codes were corrected from 71–1 (quadrate trochlea at one level or anterior to tabular horns) for *Leptodactylus, Xenopus, Karaurus, Kokartus, Ambystoma, Hynobius, Cryptobranchus,* and *Eocaecilia.*
    - Character 234: Codes were corrected from 234–1 for *Eocaecilia*.
    - Codes for these seven characters were updated from unknown to inapplicable for the above taxa (when originally coded as unknown) as well as for *Ichthyophis* and *Epicrionops* for consistency.
- **65 (tabular, horn); 66 (tabular, extension)**: character 66 cannot be coded for taxa coded for 65–1 (tabular horn absent).
    - *Limnogyrinus* was recoded to 65–1 because Schoch & Witzmann (2018:298, figs. 1–2) figured and described a small tabular with a straight posterior and lateral margin and thus lacking a horn. It was then recoded as inapplicable for character 66.
    - *Chinlestegophis* was recoded to 65–1 because Pardo et al. (2017a:Appendix) expressly questioned whether this was homologous with the tabular horn of other temnospondyls given its occipital position, as with brachyopoids (they cited *Batrachosuchus*, which was coded as lacking a tabular horn [65–1] in their matrix). It was then recoded as inapplicable for character 66.
    - *Rileymillerus* was updated from unknown to inapplicable for character 66.
    - The treatment of rhytidosteids for these characters is complicated. Some workers consider there to be a tabular horn when the tabular extends posterolaterally (e.g., in *Sangaia*; Dias-da-Silva et al., 2006) such that the posterior skull margin is not straight or posteriorly convex (in contrast to brachyopoids and plagiosaurids, which definitively lack tabular horns). In this interpretation, the tabular horn is present and merely not ‘separate’; instead, it is jointly formed by the squamosal, precluding an otic notch. *Laidleria* also has this condition, but Warren (1998b:171) did not consider it to have a tabular horn. Here, both *Sangaia* and *Laidleria* were considered to have tabular horns and were recoded to 65–0. *Sangaia* was recoded to 66–0/1 (the posterolateral portion is oriented at nearly 45 degrees), and *Laidleria* was recoded to 66–1 (it points mostly laterally). *Peltostega* has a ‘separated’ tabular horn like non-rhytidosteids and did not require modification.

- **69 (quadratojugal-maxilla):** codes for this binary character appear to be reversed because taxa without this contact ('separation by jugal') were largely coded for 69–0 ('in contact'), such as the majority of stereospondyls, while taxa with this contact were largely coded for 69–1 ('contact absent'), such as *Greererpeton* and dissorophoids. This pattern only appeared among taxa originally coded by Schoch (2013); taxa subsequently added have no pattern of errors. However, some taxa were correctly coded by Schoch (e.g., *Gerrothorax*, *Trematolestes*), indicating this was not a mere inversion of coded states from the listed character, and therefore, all taxa were reexamined rather than automatically reversing codes for all taxa originally coded by Schoch.
    - Codes were reversed (corrected) for *Proterogyrinus, Greererpeton, Edops, Adamanterpeton, Cochleosaurus, Nigerpeton, Dendrerpeton, Balanerpeton, Trimerorhachis, Neldasaurus, Acroplous, Isodectes, Capetus, Iberospondylus, Acanthostomatops, Zatrachys, Micromelerpeton, Limnogyrinus, Branchierpeton, Branchiosaurus, Apateon, Micropholis, Platyrhinops, Amphibamus, Doleserpeton, Gerobatrachus, Dissorophus, Cacops, Broiliellus, Acheloma, Phonerpeton, Ecolsonia, Onchiodon, Eryops, Sclerocephalus, Glanochthon, Archegosaurus, Australerpeton, Platyoposaurus, Lapillopsis, Rhineceps, Uranocentrodon, Broomistega, Lydekkerina, Chomatobatrachus, Plagiosuchus, Watsonisuchus, Lyrocephaliscus, Metoposaurus, Parotosuchus, Paracyclotosaurus,* and *Cyclotosaurus.*
    - Codes were corrected to unknown for *Siderops* (lateral and palatal surfaces of the quadratojugal are unknown; Warren & Hutchinson, 1983:6), and *Batrachosuchus* (no maxillary sutures are discernible or preserved; Chernin, 1977:102).
- **71 (posterior skull rim):** this character originally specified two reference points: the tabular horns and the quadrate condyles. However, not all taxa have tabular horns (65–1), yet such taxa were coded for this character (e.g., *Batrachosuchus*, *Plagiosuchus*). The original codes were correct if the 'tabular horn' reference point was changed to the 'posteriormost extent of the tabular,' provided that the tabular is present (i.e. excluding lissamphibians); in taxa with horns, the horn is the posteriormost extent, and taxa without horns could then be properly coded. This change was implemented and did not require any coding modifications.
- **72 (occipital flange); 73 (postparietal-exoccipital); 236 (postparietal-squamosal); 237 (postparietal length):** these characters cannot be coded for any taxon coded for 235–2 (postparietals absent).
    - Codes for these four characters were updated from unknown to inapplicable for *Triadobatrachus, Leptodactylus, Xenopus, Karaurus, Ambystoma, Hynobius, Cryptobranchus, Ichthyophis,* and *Epicrionops*.
- **73 (postparietal-exoccipital); 108 (parasphenoid-exoccipital); 118 (pterygoid-exoccipital); 136 (exoccipital condyles); 137 (exoccipital condyles); 247 (jugular foramen):** these characters should not be coded when the exoccipitals are unossified, as often occurs in small neotenic dissorophoids (e.g., Boy, 1972; Werneburg, 1991, 2012; Schoch & Milner, 2008). The absence of the exoccipitals is thus phylogenetically relevant, but there is no character or state that refers to the lack of ossification (any character requiring them to be ossified still should not have been coded).

    A state for 'exoccipitals unossified' could have been placed with one of several characters related to the contact of the exoccipitals with another element; here, it is placed with character 73 simply because this is the first such character. ‘Exoccipitals unossified’ is appended as the new state 2 to create an unordered new multistate character, and state 0 is modified to refer to ‘contact absent when exoccipitals are ossified.’ The treatment when postparietals are absent (see previous point) is unaffected by this modification.

  - *Acanthostomatops*, *Limnogyrinus*, *Branchierpeton*, and *Branchiosaurus* were recoded from 73–1 (postparietal and exoccipital sutured) to 73–2.
  - All four were changed from 108–0 (parasphenoid suturing with exoccipitals) and 118 (no pterygoid-exoccipital contact). *Acanthostomatops* was also changed from 136–1 (trilobed condyle with basioccipital forming ventral part of facet) and 137–0 (exoccipital condyles short and broad base, projecting only with posterior half behind occipital rim); the other taxa were updated from unknown to inapplicable.

- **74 (postfenestral window):** as originally designed, this binary character seemed to be intended to differentiate a few temnospondyls with very small, circular posttemporal fenestrae (e.g., *Laidleria, Gerrothorax*; Warren, 1998a; Schoch & Witzmann, 2012; Witzmann et al., 2012) from all other sampled taxa.
  - **74. Postfenestral window.** Large opening, having at least double the width of the foramen magnum (0), or reduced to a tiny foramen (1). [*added by Schoch, 2013*]

  However, 74–0 specifies fenestra that are more than twice as wide as the foramen magnum; even if the greatest width of the fenestra is compared to the narrowest width of the foramen, this condition is not found in many temnospondyls originally coded for 74–0 (e.g., *Trimerorhachis*, *Micropholis*, *Callistomordax*; Schoch & Rubidge, 2005:fig. 4E; Schoch, 2008a:fig. 1C; Milner & Schoch, 2013:fig. 6E; Fig. 14C-D).

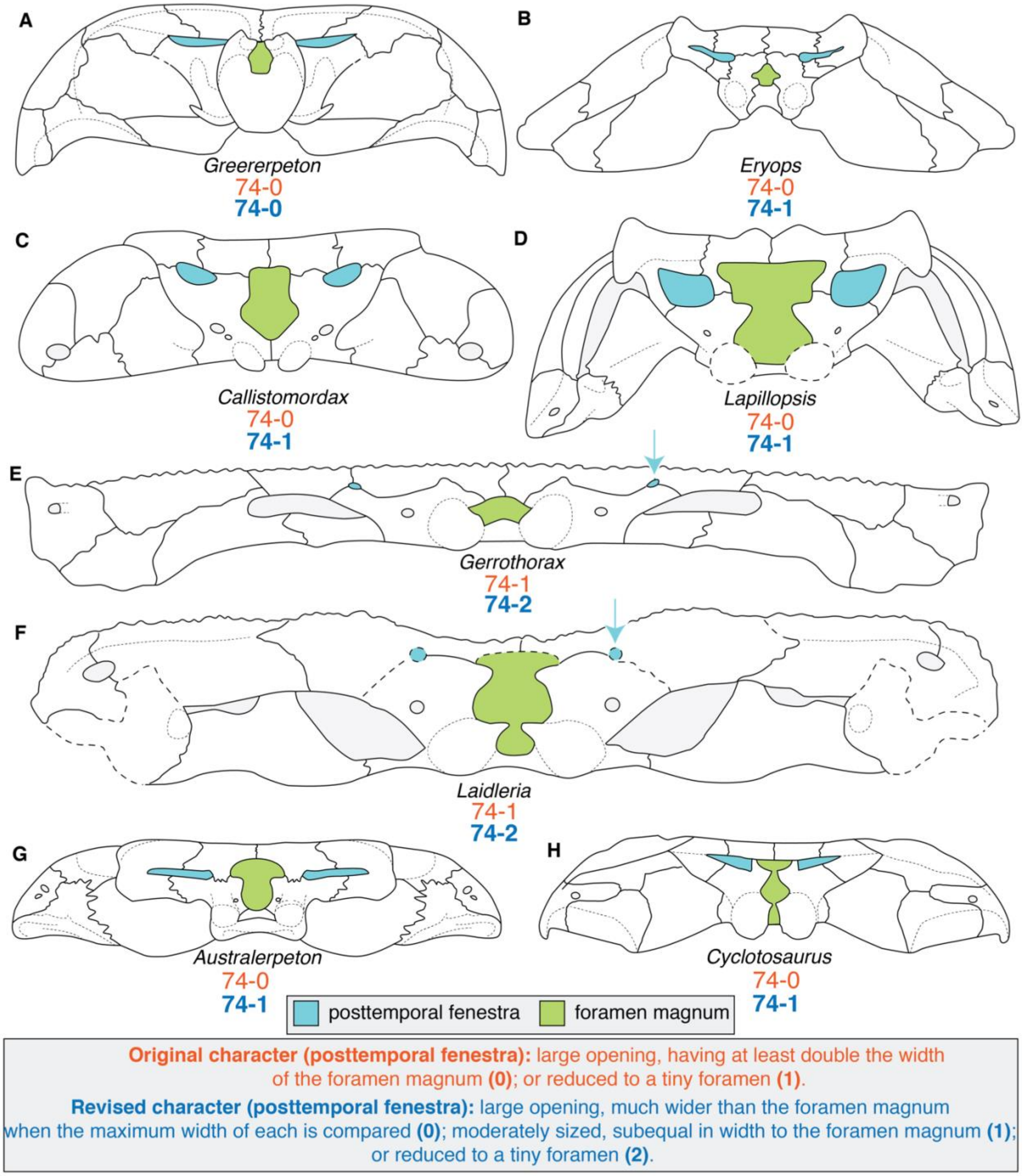

**Figure 14. Comparison of the relative width of the posttemporal fenestrae and foramina magna in temnospondyls and *Greererpeton*. A**, the colosteid *Greererpeton burkemorani* (after Bolt & Lombard, 2001); **B**, the eryopoid *Eryops megacephalus* (after Schoch & Milner, 2014); **C**, the metoposauroid *Callistomordax kugleri* (after Schoch, 2008); **D**, the stereospondyl *Lapillopsis nana* (after Yates, 1999); **E**, the plagiosaurid *Gerrothorax pulcherrimus* (after Schoch & Witzmann, 2012); **F**, the rhytidosteid *Laideria gracilis* (after Warren, 1998a); **G**, the rhinesuchid *Australerpeton cosgriffi* (after Eltink et al., 2016); **H**, the capitosaur *Cyclotosaurus intermedius* (after Sulej & Majer, 2005). Arrows are given to highlight the particularly small posttemporal fenestrae of *G. pulcherrimus* and *L. gracilis*. Orange text represents the original character and codes from Schoch (2013); blue text represents the revised character and codes. Silhouettes not to scale.

In most instances, the widths are subequal at best, or the fenestra is smaller (narrower, see *Callistomordax*; Fig. 14C). Much of the ambiguity stems from the variability in the shape of the foramen magnum, which can be keyhole-shaped in taxa without ossification of the synotic tectum (e.g., most stereospondylomorphs; Yates, 1999:fig. 11; Hewison, 2007:fig. 33; Eltink et al., 2016:fig. 7; Schoch, 2018c:fig. 7D) and which can thus be quite wide dorsally but very constricted at the mid-height and of moderate width ventrally. However, reconstructions of the occiput of *Greererpeton* genuinely depict much wider fenestrae (e.g., Smithson, 1982:fig. 12; Fig. 14A), so state 0 as originally defined does exist within the matrix. To remedy this, the character is modified to specify that the width comparison is specifically based on the greatest width of each opening. State 0 is modified to remove the 'twice as large' specifier and is instead changed to 'much wider'. One state was then added to bridge the gap with original state 1 (now 74–2); this represents fenestrae of a more subequal size (see *Callistomordax* and *Metoposaurus* for examples). All taxa were recoded. The character is also renamed, as these openings are more commonly called the 'posttemporal fenestrae,' to differentiate them from the paraquadrate foramina (more ventrolaterally situated) and any possible occipital foramina for cranial nerves or vasculature.

- Codes were recoded from 74–0 to the revised 74–1 for *Edops, Trimerorhachis, Neldasaurus, Dvinosaurus, Isodectes, Tupilakosaurus, Micropholis, Doleserpeton, Dissorophus, Cacops, Acheloma, Phonerpeton, Eryops, Sclerocephalus*, *Archegosaurus*, *Australerpeton, Platyoposaurus*, *Peltobatrachus, Rhineceps, Uranocentrodon, Broomistega, Lydekkerina, Chomatobatrachus*, *Sangaia*, *Peltostega, Edingerella, Benthosuchus, Trematosaurus, Trematolestes, Lyrocephaliscus, Callistomordax*, *Wetlugasaurus*, *Parotosuchus*, *Eryosuchus*, *Paracyclotosaurus*, *Cyclotosaurus*, *Mastodonsaurus, Rileymillerus*, and *Chinlestegophis*.
- Codes were recoded from 74–0 to the new 74–2 for *Batrachosuchus*, *Metoposaurus*, and *Eocyclotosaurus*.
- Codes were recoded from 74–1 to the new 74–2 for *Laidleria* and *Gerrothorax*.
- *Nigerpeton* was originally coded for 74–0, but Steyer et al. (2006:23, fig. 3) described a small, circular opening, and this is corrected to 74–2.
- *Watsonisuchus* was originally coded for 74–0 but the holotypes of *W. gunganj* and '*W. rewanensis*' have differing conditions that overlap the revised states 0 and 1 (Warren, 1980:fig. 5), and this OTU was recoded as polymorphic.
- *Plagiosuchus* was originally coded for 74–1, but Damiani et al. (2009:359) stated that the opening was "large." Their only figure of the occiput (fig. 1C therein) is not of sufficient quality to refine the code beyond partial uncertainty for state 0 and the revised state 1.
- *Proterogyrinus* was uncoded because it has no openings that are homologous with the posttemporal fenestrae (the 'posttemporal fossae' are typically absent in embolomeres).
- *Adamanterpeton, Cochleosaurus*, *Chenoprosopus, Dendrerpeton, Balanerpeton, Acroplous, Thabanchuia, Capetus, Iberospondylus, Acanthostomatops, Micromelerpeton, Limnogyrinus*, *Apateon, Platyrhinops, Amphibamus, Broiliellus, Ecolsonia, Onchiodon, Glanochthon,* and *Plagiosuchus* were uncoded from 74–0 because the literature does not clearly document the presence of a

well-exposed, undistorted occiput (e.g., Milner & Sequeira, 1993:figs. 2–4; 1998:figs. 2–6), and in many instances, expressly states that these exposures or some constituent elements of the occiput are not preserved or were not ossified (e.g., Witzmann & Schoch, 2006:373; Schoch & Witzmann, 2018:298). This should not be interpreted to mean that such specimens do not exist or that the occiput could not be reconstructed (but was not for any of these taxa), but because of the modification to the character, it was not possible to recode these taxa, so they are conservatively left as unknown. *Siderops* was reconstructed but is almost entirely conjectural based on the preservation of the single specimen and is therefore also uncoded. *Triadobatrachus* was uncoded from 74–1 because Ascarrunz et al. (2016:213) stated that the foramen magnum was "completely indiscernible."

- **78 (dentition, upper jaw):** The two states of this character were not clearly differentiated.
  - **78. Dentition (upper jaw).** Conical to slightly curved inwards (0), or caniniform (1). [*added by Schoch, 2013*]

  State 1 was coded only for the trematopids *Acheloma*, *Phonerpeton*, and *Ecolsonia*. Caniniform dentition in the form of a region of proportionately enlarged teeth on the anterior part of the maxilla is common to trematopids and differentiates them from their sister group (dissorophids), but similar regions of enlarged teeth also occur in taxa like eryopoids and *Sclerocephalus* (e.g., Sawin, 1941; Boy, 1990; Schoch & Witzmann, 2009a). This accounts for the morphology captured in 37–1 (maxilla anterior margin laterally convex due to enlarged teeth). This state was not originally coded for trematopids (as it was for eryopoids and sclerocephalids), but the condition does occur in trematopids (e.g., Berman et al., 1985; Dilkes & Reisz, 1987; Dilkes, 1990, 1993; Polley & Reisz, 2011). Based on the redundancy that correctly coding trematopids as 37–1 would create, it might be presumed that 'caniniform' refers to a morphology, not to size (noting also that marginal tooth homogeneity is captured in character 77), as 78–0 is 'conical to slightly curved inwards.' This does not clearly invoke a contrasting morphology like a compressed tooth with a keel or with carinae (which occurs in some stereospondyls, like *Metoposaurus* and *Siderops*; Warren & Hutchinson, 1983; Milner & Schoch, 2004), and in any event, trematopids do have conical teeth that curve slightly inwards and that lack carinae (e.g., Berman et al., 1985:12; Dilkes, 1990:226; Berman et al., 2010:306; Gee et al., 2019:fig. 3). This character is therefore omitted.
- **81 (dentition, vomer); 251 (denticles on vomer):** these characters refer to the same feature.
  - **81. Dentition (vomer).** Tooth patches present at least in small specimens (0), or dentition entirely restricted to vomerine fangs (1). [*added by Schoch, 2013*]
  - **251. Denticles on vomers.** Present (0), absent (1). [*added by Pardo et al., 2017*]

  It is interpreted that "tooth" in character 81 refers only to a shagreen of denticles, as this character was coded as 81–1 (dentition entirely restricted to vomerine fangs) in *Limnogyrinus*, which has teeth larger than denticles, but smaller than the marginal teeth, on the vomer (e.g., Schoch & Witzmann, 2018), and in most stereospondyls, which frequently have transvomerine and parachoanal dentition that is of a similar relative size. Character 251 is maintained as is, and character 81 is omitted. In order to differentiate whether the transvomerine dentition is a single row (as in many stereospondyls) or a transversely oriented patch (as in *Limnogyrinus*, among other unsampled non-stereospondyls, e.g., *Nyranerpeton amilneri*; Werneburg, 2012), character 85 is modified. Both characters are recoded in their entirety to account for dependencies. The original characters are shown below:

- Specifically for the reanalysis of Schoch (2013), the original character 81 was restored since character 251 was not in the original version of this matrix; codes were subsequently checked. The new character 81 was not added as a new character to this specific version of the matrix since the reanalysis intended to make as few changes as possible, and broader character addition was not implemented.
    - Because character 251 was not sampled in Schoch (2013), character 81 was restored for analysis of this particular matrix, with codes vetted in full. The modification to character 85, which captures the patch of transvomerine teeth in *Apateon* and *Limnogyrinus*, is maintained.
        - *Greererpeton* was newly coded for 81–1 from unknown (Smithson, 1982). *Iberospondylus*, *Broiliellus*, *Ecolsonia*, and *Broomistega* were newly coded for 81–0 from unknown (Berman et al., 1985; Shishkin & Rubidge, 2000; Laurin & Soler-Gijón, 2006).
        - *Nigerpeton* was corrected from 81–0 to 81–1; Steyer et al. (2006:24) stated that there were no accessory teeth or denticles. *Branchierpeton* was corrected from 81–1 to 81–0 based on Werneburg (1991). *Lydekkerina* was corrected from 81–1 to 81–0; Jeannot et al. (2006:830) noted the presence of vomerine shagreens, making note of overpreparation in some areas. This is an autapomorphy among lydekkerinids. *Cyclotosaurus* was changed from 81–1 to 81–0 based on the substitution of *C. intermedius* for *C. robustus*; the presence of palatal denticles is autapomorphic for the former within *Cyclotosaurus*.
        - *Neldasaurus* was also uncoded from 81–1; Schoch (2018a) stated that the vomer was probably covered by small teeth that were removed during preparation. *Isodectes* was uncoded from 81–1 because Sequeira (1998:252) remarked on how preparation made it difficult to discern the distribution of denticles on the palate. Her subsequent comment that "residual pits indicating the position of the denticle field appear to be confined to the anterior pterygoid" is thus probably a statement that denticles were at least present on the pterygoid, not an inference that they were only present on the pterygoid. *Branchiosaurus* was uncoded from 81–1 because the most recent revision by Werneburg (2012) did not confidently identify the vomer. *Sangaia* was uncoded from 81–1 because the vomers are not known (Dias-da-Silva et al., 2006). *Peltostega* was uncoded from 81–1 because Cosgriff & Zawiskie (1979:17) remarked that this taxon was (at the time) the only rhytidosteid lacking palatal denticles but noted a few on the pterygoid and suggested that there may have been a more extensive covering that

was weathered away. The only specimen with the vomers preserved is the fragmentary snout figured by Janvier (1983:fig. 3B), and there are no comments or illustrations that definitively characterize the condition.

- **87 (parasphenoid, shagreen); 253 (denticles on parasphenoid):** characters 87 and 253 are clearly redundant and both express binary 'presence/absence' conditions.
    - **87. Parasphenoid (shagreen)**. Tooth patches present (0), or entirely absent (1). [*added by Schoch, 2013*]
    - **253. Denticles on parasphenoid.** Present (0), absent (1). [*added by Pardo et al., 2017a*]

  Character 87 is retained simply because it occurs earlier in the matrix. However, coded states differ for several taxa:
    - *Trimerorhachis* was most recently described by Milner & Schoch (2013:107), who stated that "the presence of a denticle field on the ventral side of the basal plate of the parasphenoid is not clear; if present, it must have been small and faintly developed." The conservative way to reconcile the differences (87–0, 253–1) is to code it as unknown, but with the addition of a third state (see below for character 352), this can be coded as partial uncertainty for 87–0/1.
    - *Dissorophus multicinctus* has not been redescribed since DeMar (1968:1217) who stated that only vomerine palatal teeth could be confidently identified (indicating preservational conditions may not have been sufficient for preservation of denticles). Schoch (2012:fig. 3D) has reconstructed this taxon without parasphenoidal dentition, as with most dissorophids, but no primary data have been newly provided to substantiate this (Milner's 2003 redescription of a junior synonym, "*Longiscitula houghae*" does not address this), and various attributes of this reconstruction has been disputed by Dilkes (2015, 2020). The conservative way to reconcile the differences (87–0, 253–1) is to code it as unknown.
    - *Cacops aspidephorus* was most recently described by Anderson et al. (2020:7) who stated that a small surface area between the basal tubera and the basicranial articulation is covered by a shagreen of denticles, in contrast to other species of *Cacops* and other dissorophids. This code is updated to 87–0 from 87–1.
    - *Sclerocephalus haeuseri* was most recently described by Schoch & Witzmann (2009a:148), who made no comment on dentition of the parasphenoid but who did remark that there are denticles on the palatal bones. Schoch (2003) noted a decrease in the presence of denticles on this element throughout early ontogeny, which was shown by Boy (1988:figs. 3–4). I feel that this is sufficient to correct the code to 87–1 from 87–0 and is considered a mischaracterization.
    - *Uranocentrodon senekalensis* has not been appreciably redescribed since Broom (1930:fig. 2), who depicted denticles on the parasphenoid. This is in agreement with both earlier work (e.g., Haughton, 1925:228) and the coding by Marsicano et al. (2017), the most recent revision of Rhinesuchidae. This is considered sufficient to maintain the code as 87–0.
    - *Benthosuchus sushkini* has not been appreciably redescribed since Bystrow & Efremov (1940:figs. 67–69), in which it is repeatedly figured with a denticle field on the parasphenoid. This follows how previous workers have coded the taxon (e.g., Damiani, 2001; Schoch, 2008b), although it is not always figured as such.

Novikov (2012b:402–403), in describing a new species of *Benthosuchus*, lists parasphenoid denticles as being present. This is considered sufficient to maintain the code as 87–0.

  - *Trematosaurus brauni* was most recently described by Schoch (2019:51) who compared the topography of the basal plate to other stereospondyls with fine pitting and denticles; this taxon has also been previously reconstructed as having denticles (e.g., Schoch & Milner, 2000:fig. 77). This is considered sufficient to maintain the code as 87–0.
  - *Lyrocephaliscus euri* was most recently described by Mazin & Janvier (1983:16, 19), who stated that denticles are found on the parasphenoid, as also depicted by Schoch & Milner (2000:fig. 83) and described by Säve-Söderbergh (1935:23). This is considered sufficient to maintain the code as 87–0.
  - The following taxa were coded for character 87 but not for character 253 and were left as originally coded for character 87: *Micromelerpeton*, *Platyrhinops*, *Broiliellus*, *Acheloma*, *Phonerpeton*, *Eryops*, *Sangaia*, and *Kokartus*.
  - The following taxa were coded for character 253 but not for character 87, and the condition of the former (denticles absent) was newly applied to the latter: *Karaurus*.

  The following taxa also had their codes changed but did not have an original discrepancy in codes:

  - *Nigerpeton* was corrected from 87–0 to 87–1; Steyer et al. (2006:22–23) made no comments on parasphenoidal dentition and noted that dentigerous fragments on the cultriform process were likely the ossicles normally situated within the interpterygoid vacuities (character 139).
  - *Capetus* was uncoded from 87–0 because Sequeira & Milner (1993:670) stated that there are patches of denticulate bone in the interpterygoid region but also expressly stated that it is unknown whether these are from the basal plate of the parasphenoid or some of the loose plates that would have covered the interpterygoid vacuities.
  - *Iberospondylus* was newly coded for 87–0 (Laurin & Soler-Gijón, 2006:287–288).
  - *Edingerella* was corrected from 87–0 to 87–1; Steyer (2003) and Maganuco et al. (2009) both described the surface as "granular," but the latter expressly noted the absence of denticles. The same feature appears work by in Lehman (1961:fig. 9), but all three of these studies referred to it as an "area aspera," which is not equivalent to a denticle field.
  - *Gerrothorax* was corrected from 87–0 to 87–0&1; this is a well-documented polymorphism (Schoch & Witzmann, 2012:377–378, fig. 4).
  - *Callistomordax* was corrected from 87–0 to 87–1; Schoch (2008a:90) stated that the area aspera is heavily ornamented but edentulous.

- **87 (parasphenoid, shagreen); 115 (cultriform process, dentition):** character 115 (cultriform process, dentition) was intended to capture a further variation on parasphenoid dentition (expanded onto the cultriform process), which overlaps with character 352 that was added by Schoch et al. (2020) and not technically sampled here. The new multistate character is not rearranged since the three states do not necessarily form a morphocline.

- **115. Cultriform process (dentition)**. Main shaft edentulous except for base (0), or with elongate tooth patch (1). [*added by Schoch, 2013*]
- **352. (no title).** Denticle field on basal plate present or not (0), or greatly expanded anteriorly along cultriform process (1). [*added by Schoch et al., 2020*]

States 115–1 and 352–1 are 'synonymized' and subsequently appended to character 87 as a new state 2, and the position of character 115 is not replaced. This approach is preferable to the use of binary characters where one state is clearly a convoluted compound state, as with 352–1 (capturing both an absence and a presence in the same state), and mitigates issues with the original coding of character 115 in which taxa without any parasphenoid denticles that should have been coded as inapplicable had been coded for 115–0.

- All taxa coded for 115–1 and/or 352–1 were recoded to the new 87–2: *Acanthostomatops, Zatrachys, Micropholis, Amphibamus, Platyrhinops, Peltobatrachus, Eocaecilia*. Both zatracheids were originally coded for 115–1 and 352–0, but the former is clearly the correct treatment of their condition (e.g., Schoch, 1997:fig. 3; Witzmann & Schoch, 2006:fig. 4).
- Codes were recoded from 87–0 to the new 87–2 for *Chenoprosopus, Dvinosaurus, Limnogyrinus, Branchierpeton, Broomistega,* and *Wetlugasaurus*(e.g., Langston, 1953; Werneburg, 1991; Shishkin, 1973; Schoch & Milner, 2000, 2014; Shishkin & Rubidge, 2000; Witzmann & Schoch, 2006; Schoch & Witzmann, 2018).
- *Micromelerpeton* was recoded as polymorphic for states 0 and 2 (Boy, 1995:fig. 3), and *Gerrothorax* was recoded as polymorphic for states 0 and 1 (Schoch & Witzmann, 2012:381–382, fig. 4). *Adamanterpeton* and *Watsonisuchus* were coded as partial uncertainty for states 0 and 2 because denticles occur on the basal plate, but the posterior cultriform process is not sufficiently preserved to ascertain whether denticles extended onto it (Warren, 1980:28; Milner & Sequeira, 1998:276, 278).

- **88 (ectopterygoid, fangs); 123 (pterygoid, ectopterygoid); 124 (pterygoid-palatine-ectopterygoid); 125 (palatine, ectopterygoid); 128 (palatine, ectopterygoid, ontogeny); 129 (palatine, ectopterygoid, width); 130 (palatine, ectopterygoid, continuous tooth row); 131 (ectopterygoid length); 132 (laterally exposed ectopterygoid, LEE); 133 (ectopterygoid, Y-shaped):** any taxon coded for state 346–1 (ectopterygoid absent; exclusive to lissamphibians) cannot be coded for these characters.
  - Character 88: Codes were corrected from 88–1 (ectopterygoid fangs absent) for *Triadobatrachus*, and *Eocaecilia*.
  - Character 123: Codes were corrected from 123–0 (palatine ramus exclusively formed by pterygoid) for *Triadobatrachus, Leptodactylus, Xenopus, Ambystoma, Hynobius, Cryptobranchus, Ichthyophis, Epicrionops*, and *Eocaecilia.*
  - Character 124: Codes were corrected from 124–0 for (pterygoid contacting both ectopterygoid and palatine) for *Leptodactylus, Ichthyophis,* and *Epicrionops*. Codes were corrected from 124–1 (pterygoid only contacting ectopterygoid) for *Triadobatrachus*.
  - Character 125: Codes were corrected from 125–0 (palatine and ectopterygoid with simple, transverse suture) for *Triadobatrachus*.

    - Character 128: Codes were corrected from 128–0 (palatine and ectopterygoid maintain their width throughout ontogeny) for *Ichthyophis* and *Epicrionops*.
    - Character 129: Codes were corrected from 129–0 (palatine and ectopterygoid much wider than maxilla) for *Ichthyophis, Epicrionops*, and *Eocaecilia*. Codes were corrected from 129–1 (palatine and ectopterygoid as narrow as maxilla) *Triadobatrachus* and *Leptodactylus*.
    - Character 130: Codes were corrected from 130–0 (palatine and ectopterygoid without continuous tooth row) for *Triadobatrachus*, *Leptodactylus*, and *Eocaecilia*. Codes were corrected from 130–1 (palatine and ectopterygoid with continuous tooth row) for *Ichthyophis* and *Epicrionops*.
    - Character 131: Codes were corrected from 131–1 (ectopterygoid markedly shorter than palatine) for *Triadobatrachus*.
    - Character 132: Codes were corrected from 132–0 (lateral exposure of ectopterygoid absent) for *Triadobatrachus*.
    - Character 133: Codes were corrected from 133–0 (ectopterygoid with continuous maxillary suture) for *Triadobatrachus*.
    - Codes for these 10 characters were updated from unknown to inapplicable for the above taxa (when originally coded as unknown) as well as for *Karaurus, Kokartus,* and *Cryptobranchus* for consistency.
- **92 (anterior palatal opening(s)):** this character requires at least one palatal opening to be present between the vomer and premaxilla; the identical name to character 91 and the 'if present' specifier for state 92–0 evinces that they are linked. Taxa coded for state 91–0 (vomer and premaxilla with continuous suture) cannot be coded for this character.
    - **91. Anterior palatal opening(s)**. Vomer and premaxilla with continuous suture (0), or perforated to accommodate symphyseal fangs (1). [*added by Schoch, 2013*]
    - **92. Anterior palatal opening(s).** Unpaired if present (0), or paired (1). [*added by Schoch, 2013*]
    - Codes were corrected to inapplicable from 92–0 (opening unpaired if present) for *Proterogyrinus, Edops, Adamanterpeton, Cochleosaurus, Chenoprosopus, Dendrerpeton, Balanerpeton, Capetus, Iberospondylus, Acanthostomatops, Zatrachys, Branchierpeton, Apateon, Micropholis, Platyrhinops, Amphibamus, Doleserpeton, Gerobatrachus, Dissorophus, Cacops, Broiliellus, Acheloma, Phonerpeton, Ecolsonia, Onchiodon, Eryops, Sclerocephalus, Glanochthon, Archegosaurus, Platyoposaurus, Plagiosuchus, Gerrothorax, Paracyclotosaurus, Leptodactylus, Ambystoma,* and *Hynobius,*
    - *Xenopus* and *Kokartus* were originally coded as unknown for character 91, so they were recoded to unknown from 92–0.
    - *Micromelerpeton* was originally coded as 91–0, but this condition is intraspecifically variable (Boy, 1995:fig. 3).
    - *Cryptobranchus, Ichthyophis, Epicrionops,* and *Eocaecilia* were updated to inapplicable from unknown for character 92.
- **95 (anterior palatal depression):** this character requires an anterior palatal depression to be present. Taxa that lack such a depression cannot be coded for this character.
    - Codes were corrected from 95–0 (posterior rim round if present) to inapplicable for *Dendrerpeton* and *Balanerpeton*. More taxa may lack such a depression, but

this is often difficult to discern from illustrations or older photographs in the literature.

- **99 (choana, lateral):** the original coding of this character for taxa coded by Schoch (2013) appears to be largely inverted from the listed character. Most taxa were coded for 99–0 (‘choana anterolaterally expanded with triangular outline’), but most taxa lack this expansion. Additionally, the few taxa originally coded for 99–1 (‘choana oval’) have markedly disparate morphology. *Lyrocephaliscus* genuinely has a simple oval choana (e.g., Schoch & Milner, 2000:fig. 83), like most other temnospondyls and *Greererpeton*, whereas *Dendrerpeton* has the marked lateral expansion that is captured in 99–0 (e.g., Holmes et al., 1998:fig. 4B), but both *Lyrocephaliscus* and *Dendrerpeton* were coded for 99–1. This indicates that there is not merely a reversal of states, and codes were systematically checked.
    - Codes were reversed (corrected) from 99–0 to 99–1 for *Proterogyrinus, Greererpeton, Balanerpeton, Trimerorhachis, Neldasaurus, Dvinosaurus, Acroplous, Isodectes, Thabanchuia, Tupilakosaurus, Capetus, Iberospondylus, Micromelerpeton, Limnogyrinus, Branchierpeton, Branchiosaurus, Apateon, Micropholis, Platyrhinops, Amphibamus, Doleserpeton, Gerobatrachus, Dissorophus, Cacops, Broiliellus, Acheloma, Phonerpeton, Ecolsonia, Onchiodon, Eryops, Sclerocephalus, Glanochthon, Archegosaurus, Australerpeton, Platyoposaurus, Lapillopsis, Rhineceps, Uranocentrodon, Broomistega, Lydekkerina, Chomatobatrachus, Laidleria, Siderops, Batrachosuchus, Plagiosuchus, Gerrothorax, Edingerella, Watsonisuchus, Benthosuchus, Trematosaurus, Trematolestes, Callistomordax, Metoposaurus, Wetlugasaurus, Parotosuchus, Eryosuchus, Paracyclotosaurus, Eocyclotosaurus, Cyclotosaurus, Mastodonsaurus*, and *Rileymillerus*.
    - Codes were reversed (corrected) from 99–1 to 99–0 for *Cochleosaurus* and *Dendrerpeton*.
- **99 (choana, lateral); 100 (choana, medial); 101 (choana, width):** these three characters all capture aspects of the shape of the choana.
    - **99. Choana (lateral).** Anterolaterally expanded with triangular outline (0), or oval (1).
    - **100. Choana (medial).** Medial margin straight or gently convex (0), or anteromedially expanded giving choana a reniform outline (1).
    - **101. Choana (width).** Elongated oval to slit-like (0), or wide round (1).

  There are essentially four different choanal morphologies among temnospondyls (Fig. 15). The most common (though not necessarily plesiomorphic) condition is a symmetrical oval opening (Fig. 15A), which also occurs in *Greererpeton* (Smithson, 1982:fig. 11; the condition is uncertain in the operational outgroup, *Proterogyrinus*, because of the absence of a completely known vomer; Holmes, 1984). A similar condition is a symmetrical circular opening, as found in taxa like *Edops* (Romer & Witter, 1942:fig. 3), *Onchiodon* (Boy, 1990:fig. 3B), and *Plagiosuchus* (Damiani et al., 2009:figs. 3, 7B; Fig. 15B). These two morphologies are differentiated by character 101. Character 99 then refers to one apomorphic condition, found in taxa like *Dendrerpeton* (Holmes et al., 1998:fig. 4; Fig. 15C), *Acanthostomatops* (Witzmann & Schoch, 2006:fig. 4), and *Zatrachys* (Schoch, 1997:figs. 3–4), in which there is an anterolateral expansion in the anterior half of the choana. Character 100 refers to a different apomorphic condition, which is expansion anteromedially, as in *Amphibamus* (Schoch & Milner, 2014:fig. 30B) and *Acheloma* (Polley & Reisz, 2011:figs. 1, 7; Fig. 15D). For either apomorphic condition captured by characters 99 and 100, the profile may either be teardrop-shaped (‘triangular’), expanding anteriorly, or more like an upside-down ‘L,’ with a sharper angle to one lateral margin (*Dendrerpeton* is perhaps the best example of the latter).

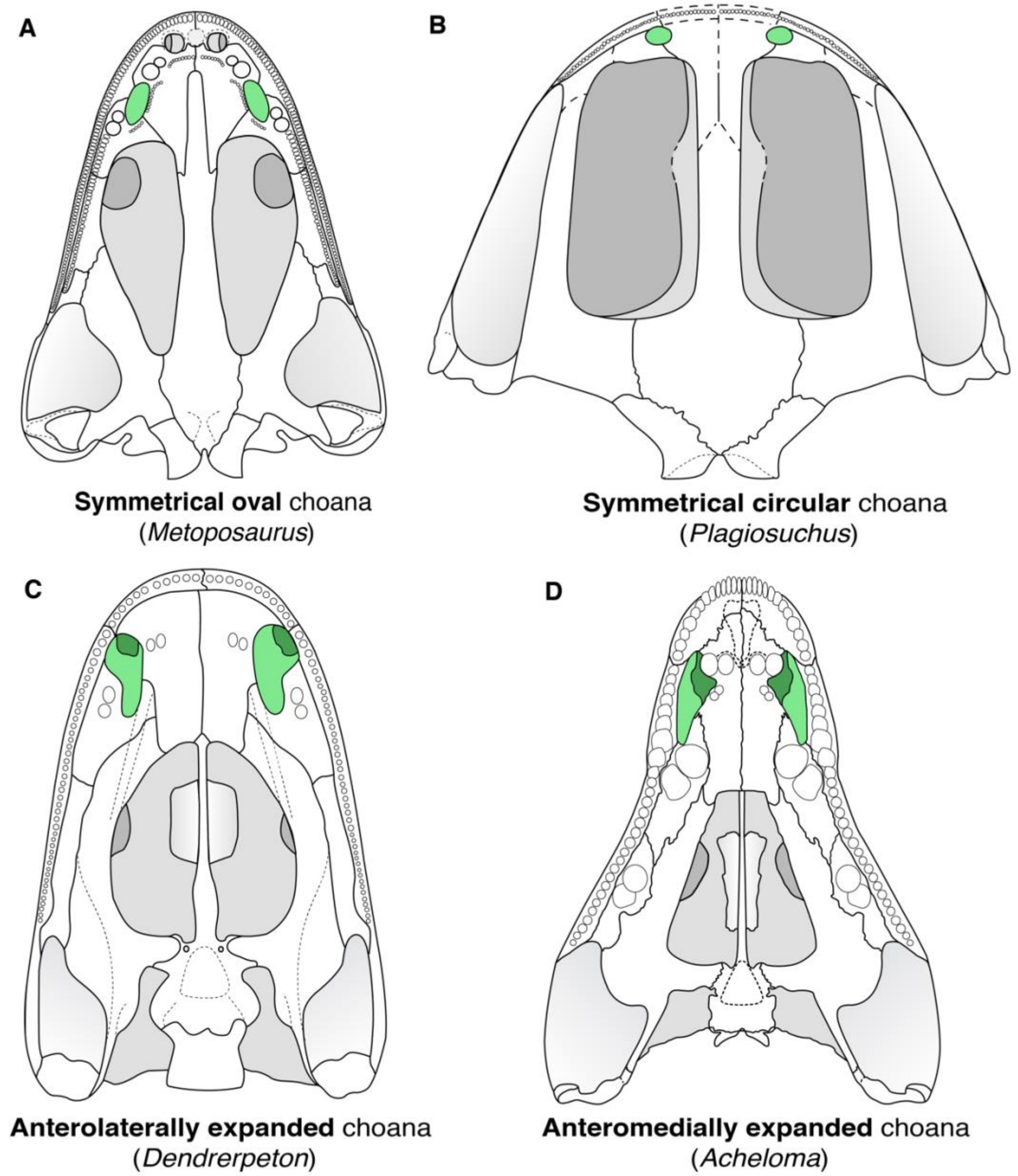

**Figure 15. Comparison of the choanal morphologies among temnospondyls. A**, the metoposaurid *Metoposaurus krasiejowensis* (after Sulej, 2007); **B**, the stereospondyl *Lapillopsis nana* (after Yates, 1999); **C**, the ‘dendrerpetid’ *Dendrerpeton acadianum* (after Holmes et al., 1998); **D**, the trematopid *Acheloma cumminsi* (after Polley & Reisz, 2011). Coronoid denticles (where present) are omitted for simplicity. Silhouettes not to scale.

As the characters are defined, any taxon coded for 99–0 (triangular outline) or 100–1 (reniform outline) should not be coded for character 101 because such a taxon could not have either an oval, slit-like opening or a round opening. However, there is no apparent recognition of this incompatibility in the previous codes, and it is hard to reconcile them. For example, *Micropholis*, *Amphibamus*, and *Doleserpeton* were all coded for 100–1, but the latter two were coded for 101–1, whereas *Micropholis* was coded for 101–0. The choana of *Doleserpeton* is the narrowest (‘slit-like’) of the three, and while that of *Amphibamus* is perhaps the shortest, it is still longer than wide and hardly ‘round.’ Therefore, even if character 101 were restricted to proportions, rather than to both proportions and shape (which are not always correlated), none of these three should be coded for 101–1. There is no taxon with a choana in the shape of an equilateral triangle, and thus any taxon with a choana as wide as it is long would have a circular (‘wide round’) choana.

Finally, the addition of lissamphibians has greatly complicated the treatment of these three characters. Among temnospondyls, no taxon is coded for both 99–0 and 100–1; these should be mutually exclusive based on the prescription of shape, as the opening cannot be both ‘triangular’ and ‘reniform.’ However, *Leptodactylus*, *Ambystoma*, *Hynobius*, and *Cryptobranchus* were all coded for this combination by Pardo et al. (2017a). None of these taxa has a condition similar to either the ‘triangular’ choana of a taxon like *Dendrerpeton* or the ‘reniform’ choana of *Micropholis*. Equally peculiar is that *Triadobatrachus* and *Xenopus* were not coded for character 99 but were coded for character 100 (for state 1). Undoubtedly the problem arises when attempting to fit the unique morphologies of batrachians with greatly reduced palates into characters meant for early tetrapods. The choana is much less defined, especially laterally, in many batrachians due to the loss of the palatine (Fig. 16).

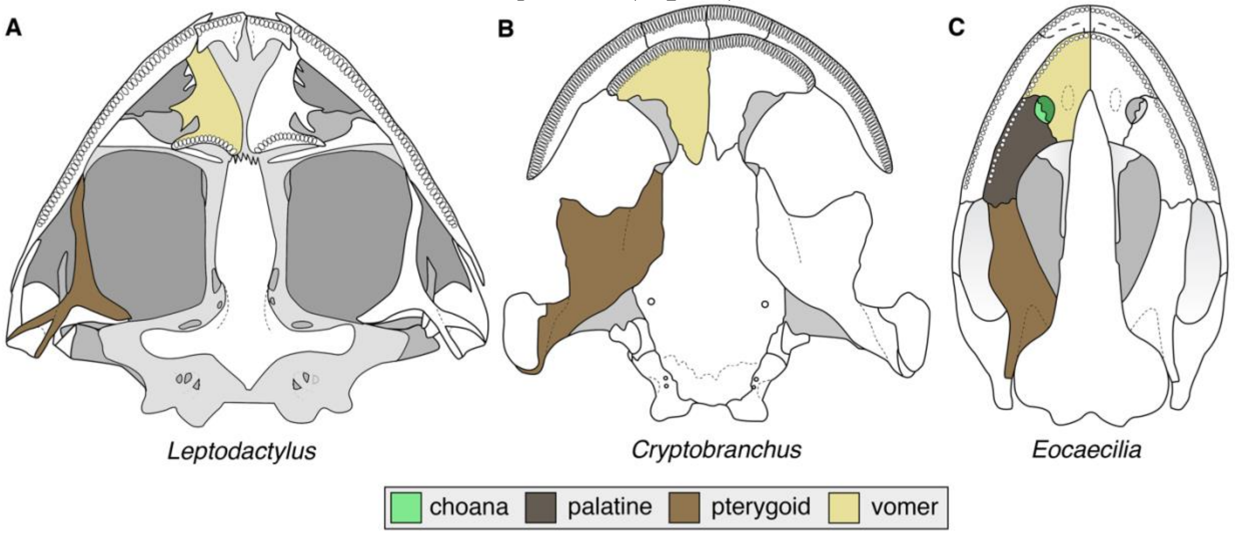

**Figure 16. Comparison of palatal anatomy among lissamphibians to demonstrate the poor definition of the choana in batrachians due to reduction of the palatal elements. A**, the anuran *Leptodactylus mystacinus* (after Ponssa, 2008:fig. 2); **B**, the caudate *Cryptobranchus alleghenienis* (after Carroll & Holmes, 1980:fig. 5); **C**, the gymnophionomorph *Eocaecilia micropodia* (after Jenkins et al., 2007:fig. 1B). Silhouettes not to scale.

Finally, differing orientations of the choana may have produced conflicting codes – many batrachians have a transversely oriented oval choana. If this is thought of as merely the result of a rotation from a condition in which an oval choana is more sagittally oriented, as part of a broader reconfiguration of the overall skull shape, then it should not be considered as either anterolaterally or anteromedially expanded. Conversely, if it is thought of as a transverse expansion of a circular choana, it could be treated as having been expanded in both directions.

- Character 101 is modified to refer explicitly to shape. States 99–1 and 100–0 are appended to this character, with the caveat that the language is modified to refer to expansion towards the aboral (‘lateral’) or oral (‘medial’) direction; this mitigates the issue in which the choana may have the same overall shape but have been sharply rotated into a transverse orientation. These are appended to the end of character 101, and the new multistate character is unordered. For batrachians, because the choana is not defined posteriorly/laterally/orally, it is not possible to code it for this redefined character because recognition of expansion inherently requires a reference point (i.e. these should not have been coded for original characters 99 or 100). All taxa were recoded.
- Character 99 is then repurposed to capture the primary orientation of the choana (more sagittal vs. more transverse), thus negating the above corrections made to the original character 99 coding (which is still detailed in case future workers wish to utilize the original character). Because orientation is determined by the long axis, any taxon with a perfectly equilateral (circular) choana should not be coded for this revised character. All taxa were recoded.
- Character 100 is not replaced or modified. The laterally/posteriorly open choana of batrachians is accounted for with any character referring to the absence of the palatine.

- **102 (vomer-pterygoid); 124 (pterygoid-palatine-ectopterygoid); 125 (palatine, ectopterygoid); 126 (palatine, vomer); 127 (LEP); 128 (palatine, ectopterygoid, ontogeny); 129 (palatine, ectopterygoid, width); 130 (palatine, ectopterygoid, continuous tooth row); 131 (ectopterygoid length); 248 (anterior palatine); 252 (denticles on palatines):** any taxon coded for 248–2 (palatine absent; exclusive to caudates) cannot be coded for these characters.
    - State 248–2 overlaps with state 356–1 in the character added by Schoch et al. (2020), which refers simply to the presence or absence of the palatine. Whereas Pardo et al. (2017a) only coded *Ambystoma*, *Hynobius*, and *Cryptobranchus* for 248–2, Schoch et al. (2020) coded these taxa and both karaurids for 356–1 (the equivalent ‘absent’ state). Karaurids were left uncoded for character 248 by Pardo et al. The treatment of their condition in this study follows that of Schoch et al.; the absence of the palatine can be easily discerned in both dorsal and palatal views in lissamphibians given the extensive modification of the palate.
    - Character 102: this character could be modified such that state 1 (‘separated by palatine’) omits any reference to the palatine. However, this would falsely homologize the condition of some caudates, in which reduction of elements in the suborbital region results in a wide separation along the lateral margin, with the common condition in temnospondyls, in which the pterygoid and vomer have lateral extents but are separated. All caudates were originally left as unknown for this character, so it is left as is, and no coding modifications were required.
    - Character 127: Codes were corrected from 127–0 (LEP absent) for *Ambystoma*.
    - Character 252: Codes were corrected from 252–0 (denticles on palatine present) for *Karaurus*. Codes were corrected from 252–1 (denticles on palatine absent) for *Ambystoma*, *Hynobius*, and *Cryptobranchus*.

- Codes for these 10 characters were updated from unknown to inapplicable (when originally coded as unknown) for the above taxa.

- **248 (anterior palatine)**: the original coding of character 248 by Pardo et al. (2017a), which was subsequently changed for many taxa by Schoch et al. (2020) is difficult to understand.
  - **248. Anterior palatine.** Short anteromedial process articulating with vomer at choana (0), long anteromedial process, more medial than lateral (1), or palatine absent (2). [*added by Pardo et al., 2017a*]

  States 0 and 1 differentiated taxa on the basis of the anteromedial process of the palatine. Pardo et al. coded 248–1 for certain stereospondyls (*Lydekkerina*, *Siderops, Batrachosuchus*, *Plagiosuchus*, *Gerrothorax, Edingerella*, *Lyrocephaliscus*), though not for many others (e.g., *Benthosuchus*, *Trematolestes*, *Rileymillerus*), as well as in *Triadobatrachus*, *Leptodactylus,* and *Xenopus* (the last one as partial uncertainty). Schoch et al. stated that codes were changed for some taxa because “a caecilian-type anteromedial process of the palatine is only present in apodans, not stereospondyls.”

  All three gymnophionomorphs were originally coded for 248–0, as was *Rileymillerus.* Schoch et al. changed all stereospondyls previously coded for 248–1 to 248–0 but also changed all three gymnophionomorphs to 248–1. This removed a possible homology with highly nested stereospondyls. The confusion is undoubtedly related to the ambiguity of the states; 248–1 probably means ‘anteromedial process longer than anterolateral process,’ rather than that the anteromedial process is more medial than lateral (by virtue of being the anteromedial process, it is more medial than lateral). If so, 248–1 (longer anteromedial process) would be exemplified by *Apateon* (e.g., Schoch & Milner, 2014:fig. 27G; Fig. 17B), originally coded as such by Pardo et al. and retained by Schoch et al. However, stereospondyls changed from 248–1 to 248–0 by Schoch et al. would also exemplify this condition (e.g., *Lydekkerina*; Jeannot et al., 2006:fig. 2; *Gerrothorax*; Schoch & Witzmann, 2012:fig. 3; Fig. 17C). Other taxa originally coded for 248–1 are more equivocal (e.g., *Laidleria*; Warren, 1998a:figs. 2, 4; *Edingerella*; Maganuco et al., 2009:fig. 10; Fig. 17D) in that they are more similar to taxa originally coded for 248–0. Some taxa originally coded for 248–0 definitively have a longer anteromedial process (e.g., *Balanerpeton*; Milner & Sequeira, 1993:figs. 1, 3, 5B; *Amphibamus*; Schoch & Milner, 2014:fig. 30B; *Acheloma*; Polley & Reisz, 2011:fig. 1; *Mastodonsaurus*; Schoch, 1999:figs. 8, 13, 15).

  Alternatively, the character could be referring to the orientation of the palatine-vomer suture; in most stereospondyls, the suture is angled posteromedially towards the midline (as in gymnophionomorphs but also in *Greererpeton* and as reconstructed in *Proterogyrinus*), whereas in most non-stereospondyls, it is oriented posterolaterally or sagittally. However, this interpretation would be complicated by taxa with prominently interdigitated sutures (e.g., *Balanerpeton*, *Platyrhinops*; Milner & Sequeira, 1993:fig. 5B; Clack & Milner, 2010:fig. 9b), horizontal sutures (e.g., *Isodectes*; Sequeira, 1998:fig. 9B), or sinuous or curved sutures (e.g., *Dendrerpeton*, *Acheloma*; Holmes et al., 1998:fig. 4B; Polley & Reisz, 2011:fig. 1). However, this interpretation of the character would not align with the length specifiers employed in the original character.
  - The language of the character is modified to differentiate the relative length of the two processes that frame the choana. This removed the ambiguity over how to measure each process when the oral (‘inner’) process diverges from the choanal

margin (as in *Balanerpeton* or *Isodectes*). There is sufficient variation among temnospondyls that states can be created for the intermediate state (processes of equal length) without producing a parsimony-uninformative character. This intermediate state becomes the revised state 1, the revised state 0 refers to a longer process along the aboral ('outer' or typically 'lateral') margin, the revised state 2 refers to a longer process along the oral ('inner' or typically 'medial') margin, and the new state 3 refers to an absent palatine. The use of 'oral' and 'aboral' is preferred to 'medial' and 'lateral' because in some taxa, it seems more likely that an oval choana has rotated based on a broader change to skull shape in which palatal elements are also rotated, rather than shortening anteroposteriorly and widening mediolaterally (e.g., *Gerrothorax*).

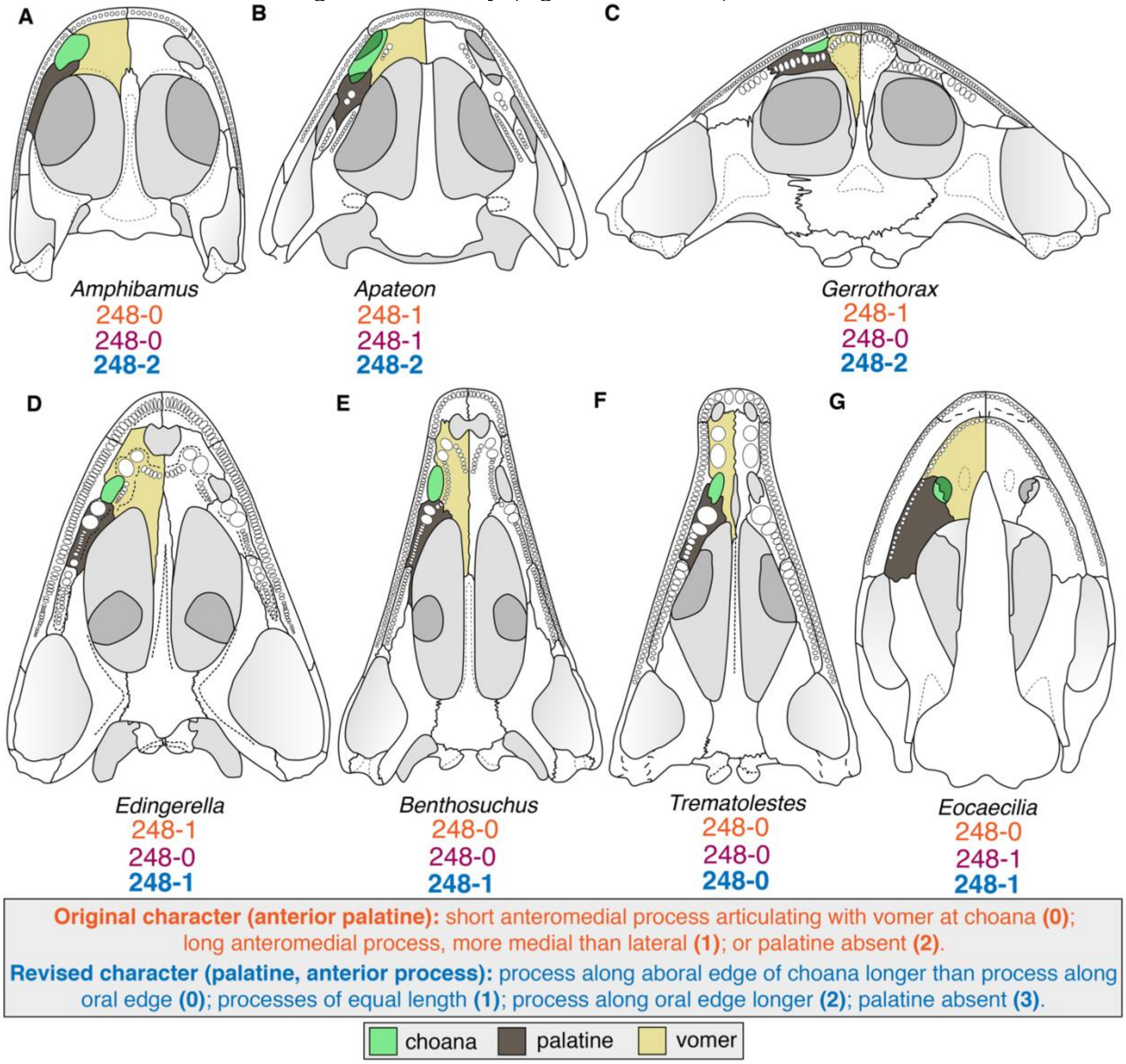

**Figure 17. Comparison of circum-choanal elements in temnospondyls. A**, the amphibamid *Amphibamus grandiceps* (after Schoch & Milner, 2014); **B**, the branchiosaurid *Apateon pedestris* (after Schoch & Milner, 2014); **C**, the plagiosaurid *Gerrothorax pulcherrimus* (after Schoch & Witzmann, 2012); **D**, the capitosaur *Edingerella madagascariensis* (after Maganuco et al., 2009); **E**, the trematosaur *Benthosuchus sushkini* (after Schoch, 2008b); **F**,

the trematosaur *Trematolestes hagdorni* (after Schoch, 2006); **G**, the gymnophionomorph *Eocaecilia micropodia* (after Jenkins et al., 2007). Orange text represents the original character and codes from Schoch (2013); red text represents the original codes from Pardo et al. (2017a); blue text represents the revised character and codes. Silhouettes not to scale.

  - Codes were recoded from 248–0 to 248–2 for *Proterogyrinus*, *Trimerorhachis, Amphibamus, Dissorophus, Acheloma*, *Eryops*, *Lydekkerina*, *Plagiosuchus, Gerrothorax*, *Lyrocephaliscus*, *Callistomordax*, and *Parotosuchus.*
  - Codes were recoded from 248–0 to 248–1 for *Greererpeton*, *Adamanterpeton, Cochleosaurus, Isodectes, Dendrerpeton*, *Capetus*, *Iberospondylus*, *Zatrachys, Micropholis, Doleserpeton, Phonerpeton*, *Onchiodon*, *Sclerocephalus, Glanochthon, Archegosaurus*, *Platyoposaurus*, *Rhineceps*, *Broomistega, Batrachosuchus, Edingerella*, *Benthosuchus*, *Paracyclotosaurus*, *Rileymillerus,*
  - Codes were recoded from 248–1 to 248–2 for *Apateon*, *Gerobatrachus, Lydekkerina, Plagiosuchus*, *Gerrothorax*, *Triadobatrachus*, and *Leptodactylus*.
  - The three gymnophionomorphs were treated as 248–1 on the basis that the palatine (or palatine portion of the maxillopalatine) practically encircles the entire choana and therefore the processes are of subequal length (Fig. 17G).
  - *Laidleria* was recoded from 248–1 to 248–0.
  - *Cacops*, *Broiliellus, Ecolsonia*, *Siderops*, and *Paracyclotosaurus* were recoded from 248–0 to partial uncertainty for 0/1/2; in these taxa, the palatine is identifiable in some form, but it is too incompletely known or defined to code a single state. *Limnogyrinus* and *Chinlestegophis* were recoded from unknown.
  - *Trematosaurus* was newly coded as 248–0.
  - *Micromelerpeton*, *Chomatobatrachus*, and *Cyclotosaurus* were newly coded as 248–1.
- **103 (vomer, anterior part); 254 (cultriform process-vomer):** character 103 was originally treated as differentiating a rare apomorphic condition (103–1) and was only coded in the five edopoids among temnospondyls (Fig. 18A-C), but the condition of a vomer with an anterior region as long or longer than the posterior region occurs in a number of others coded for 103–0 (e.g., some dvinosaurs, some dissorophoids, zatracheids, some long-snouted stereospondylomorphs; Fig. 18D, 18G), as well as in many lissamphibians (as coded for the latter by Pardo et al., 2017a).
  - **103. Vomer (anterior part).** Anterior portion shorter than behind anterior level of choana (0), or as long or longer (1).

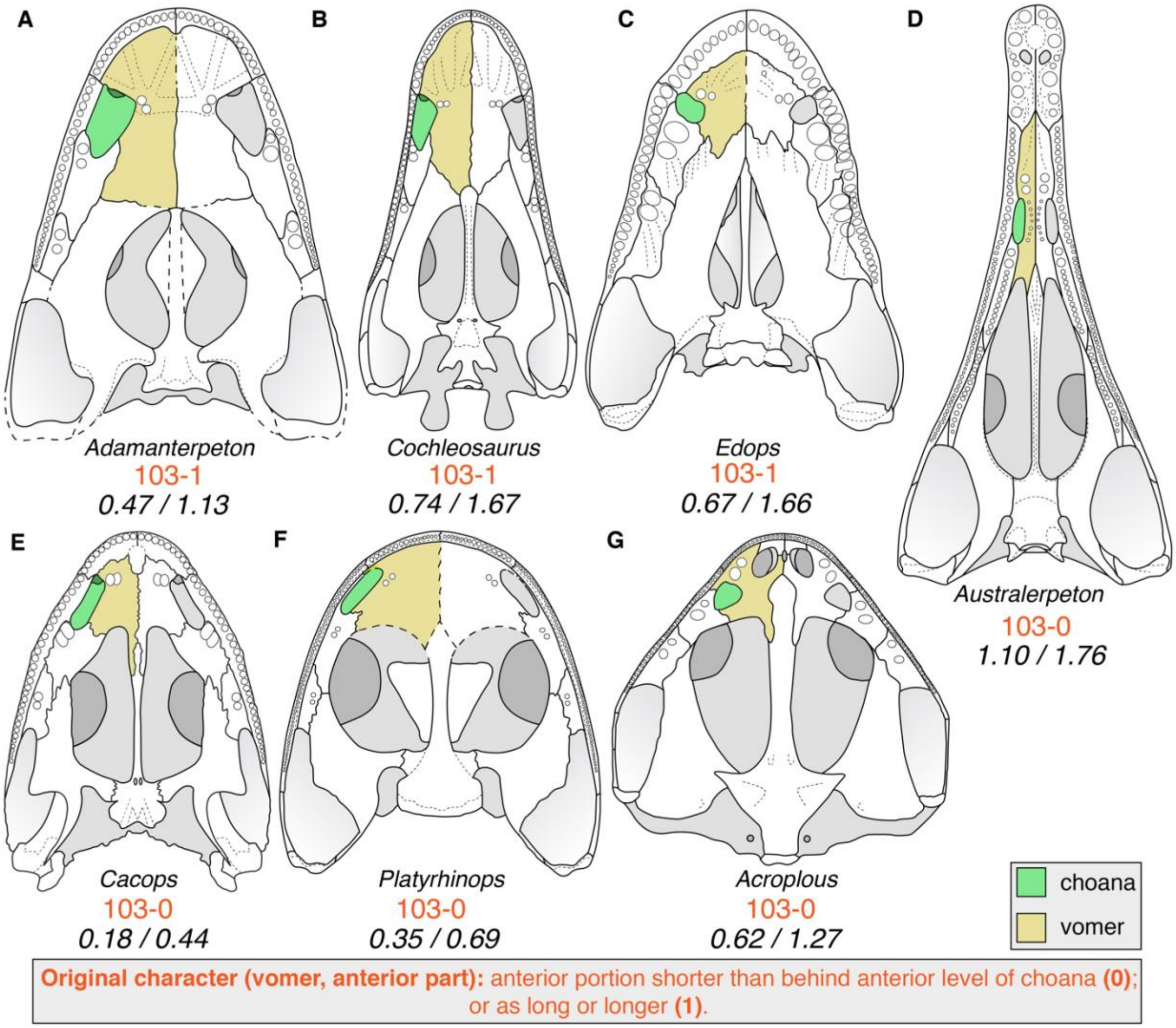

**Figure 18. Comparison of the relative length of the vomer anterior and posterior to the choana among temnospondyls. A**, the edopoid *Adamanterpeton ohioensis* (after Milner & Sequeira, 1998); **B**, the edopoid *Cochleosaurus bohemicus* (after Sequeira, 2004); **C**, the edopoid *Edops craigi* (after Romer & Witter, 1942); **D**, the rhinesuchid *Australerpeton cosgriffi* (after Eltink et al., 2016); **E**, the dissorophid *Cacops morrisi* (after Reisz et al., 2009); **F**, the amphibamiform *Platyrhinops lyelli* (after Clack & Milner, 2010); **G**, the dvinosaur *Acroplous vorax* (after Englehorn et al., 2008). Coronoid denticles (where present) are omitted for simplicity. Measurements represent the ratio of pre-choanal length to post-choanal length. Both are measured from the anteriormost and posteriormost extent of the choana, respectively, to the anteriormost and posteriormost extent of the vomer, respectively, inclusive of any lengthy posterior process along the medial margin of the interpterygoid vacuity (as in *C. morrisi*). Orange text represents the original character and codes from Schoch (2013). Silhouettes not to scale.

It appears that coding largely correlates with whether there is an extensive posteromedial process of the vomer along the lateral edge of the cultriform process (which produces a longer post-choanal region of the vomers), but this begins to overlap with character 254 (cultriform process-vomer). There is also the issue that as defined, the choana itself is factored into the comparison, which would overlap with the set of choanal characters (originally 99–101). Finally, because the choana is included in the ‘posterior region,’ it is not even clear that most edopoids can be coded for 103–1; their coding would only be validated if the ‘posterior region’ was restricted to the region of the vomer posterior to the level of the posterior choanal margin.

There are three conditions that produce a ‘broad’ cultriform process contact with the vomer (254–1; Fig. 19): (1) posteromedial processes of the vomers that extend along the lateral edge of a cultriform process that terminates at or posterior to the anterior margin of the interpterygoid vacuities (e.g., *Phonerpeton, Lapillopsis*; Fig. 19A-B); (2) a cultriform process extending well past the level of the anterior margin of the interpterygoid vacuities that partially subdivides the vomers along the midline (e.g., *Cochleosaurus*; Fig. 19E-F); and (3) both (e.g., *Dvinosaurus, Acroplous*, many stereospondyls; Fig. 19C-D). Note that any taxon with posteromedial processes of the vomers will have partially subdivided vomers, which is why the position relative to the level of the interpterygoid vacuities is important. It is also possible to have posteromedial processes of the vomer that meet along the midline to truncate the cultriform process well posterior to the anterior level of the vacuities (e.g., *Benthosuchus*, *Trematosaurus*). These conditions differ appreciably and should not be treated as homologous.

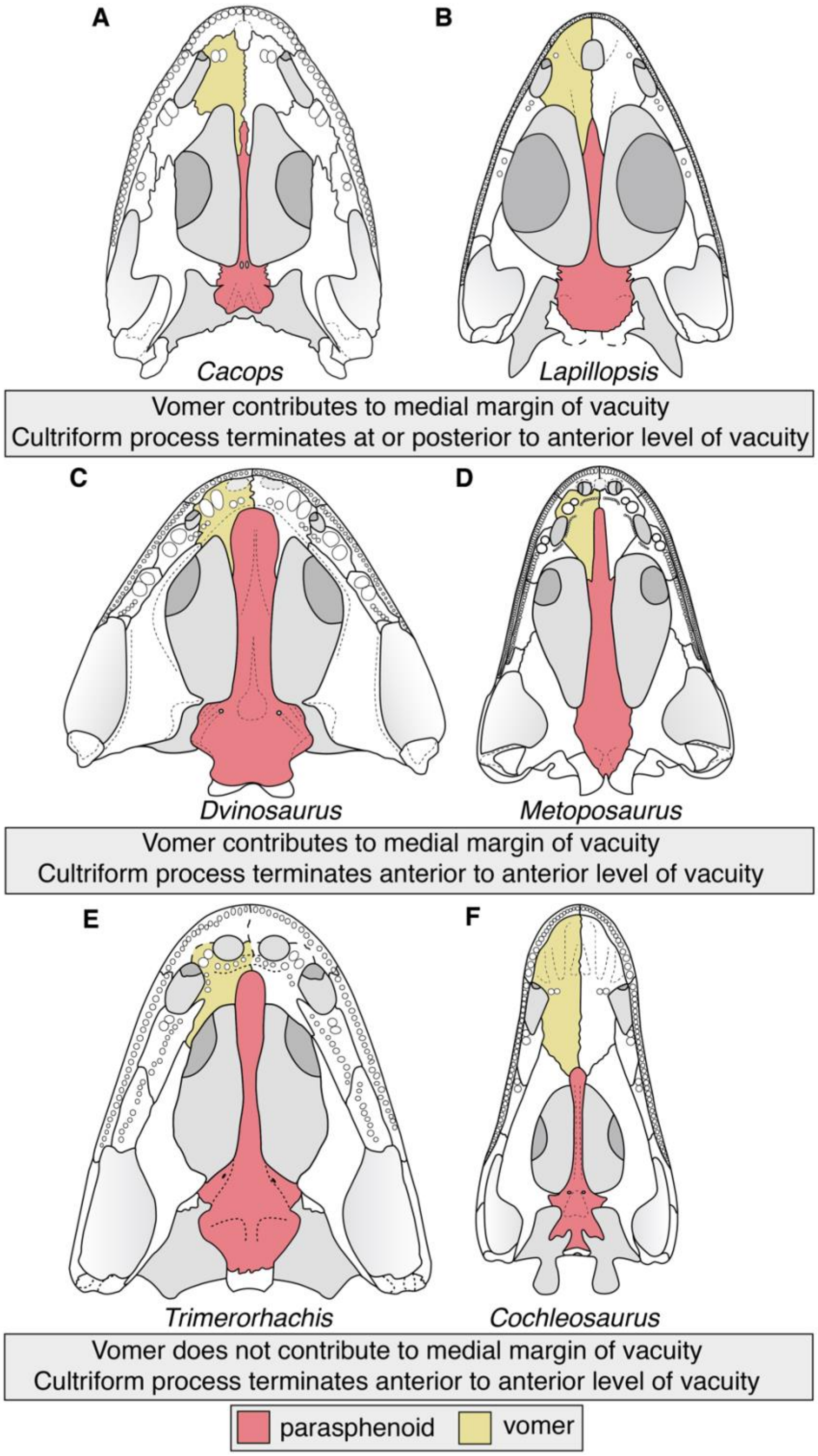

**Figure 19. Comparison of the vomer-cultriform process relationship among temnospondyls. A**, the dissorophid *Cacops morrisi* (after Reisz et al., 2009); **B**, the stereospondyl *Lapillopsis nana* (after Yates, 1999); **C**, the dvinosaur *Dvinosaurus primus* (after Schoch & Milner, 2014); **D**, the metoposaurid *Metoposaurus krasiejowensis* (after Sulej, 2007); **E**, the dvinosaur *Trimerorhachis insignis* (after Milner & Schoch, 2013); **F**, the edopoid *Cochleosaurus bohemicus* (after Sequeira, 2004). Coronoid denticles (where present) are omitted for simplicity. Silhouettes not to scale.

There are then two conditions that produce a vomer where the pre-choanal region is at least as long if not longer than the post-choanal region (regardless of whether this is defined as from the anterior or posterior choanal margin): (1) relatively elongated vomer (e.g., some edopoids; concurrent with the elongation of the roofing elements); or (2) the absence of posteriorly extensive posteromedial processes regardless of whether they are subdivided or not by the cultriform process (e.g., *Doleserpeton*; the posterolateral process rarely extends substantially further posterior than the choana; see comments on character 104 below). These conditions are demonstrably not homologous and should not be treated as such either.

- In my opinion, the original treatment of character 103 was merely meant to capture the marked preorbital elongation of the edopoids, which is already reflected in character 9 (preorbital length) and character 11 (premaxilla, prenarial region). It is not surprising that such modification of the skull roof also resulted in pre-choanal elongation on the palate, and these characters should not be considered independent. Therefore, it is completely revised to refer to the presence and relative length of the posteromedial processes of the vomer, and all taxa were recoded.
- Character 254, if left unmodified, would overlap with the redefined character 103. Therefore, it is modified to refer to the anterior extent of the cultriform process, which removes the ambiguity in defining 'broad' versus 'narrow.' All taxa were recoded for this character.

- **104 (vomer, extension):** there is some ambiguity in what feature this character refers to because the name and character states are somewhat discordant.
  - **104. Vomer (extension).** Vomer contacts pterygoid lateral to choana (0), or extending posteriorly (1). [*added by Schoch, 2013*]

Although the name suggests that it relates to the posterior extent of the vomer, state 104–0 specifies a contact with the pterygoid, implying that it is specifically about the longitudinal position of the vomer-pterygoid suture relative to the choana, and therefore, any taxon coded for state 102–1 (vomer-pterygoid separated by palatine) should not have been coded for this character (but was; e.g., many dissorophoids).

It is also inferred that 104–0 should be 'vomer contacts pterygoid medial to choana,' as the pterygoid never extends onto or along the lateral side of the choana in the sampled taxa. Additionally, the original coding for character 104 is difficult to understand. State 104–1 (extending posteriorly) was originally coded for taxa like *Trimerorhachis* in which the pterygoid terminates well posterior to the choana, and the vomer-pterygoid contact is formed by a posteriorly extensive process of the vomer that is both medial and posterior to the choana (e.g., Milner & Schoch, 2013:fig. 1B; Fig. 20A). However, other taxa with the same condition of a pterygoid-vomer contact well posterior to the level of the posterior choanal margin (e.g., *Adamanterpeton*, *Acheloma, Acanthostomatops*; Milner & Sequeira, 1998:fig. 10; Witzmann & Schoch, 2006:fig. 4; Polley & Reisz, 2011:fig. 1) were coded for 104–0 (vomer contacts pterygoid medial to choana). This ambiguity is further complicated by taxa in which the posterolateral portion of the vomer extends to a level posterior to the choana (not necessarily directly behind the choana), but the pterygoid is anteriorly extensive and reaches to the level of the posterior choanal margin; the vomer-pterygoid contact is therefore not a point but a long obliquely oriented overlapping suture, and its anteroposterior extent is not a level but a range (e.g., *Dendrerpeton*, *Eryops*, *Sclerocephalus*; Sawin, 1941:pl. 2; Holmes et al., 1998:fig. 4B; Schoch & Witzmann, 2009a:fig. 4B; Fig. 20E-F).

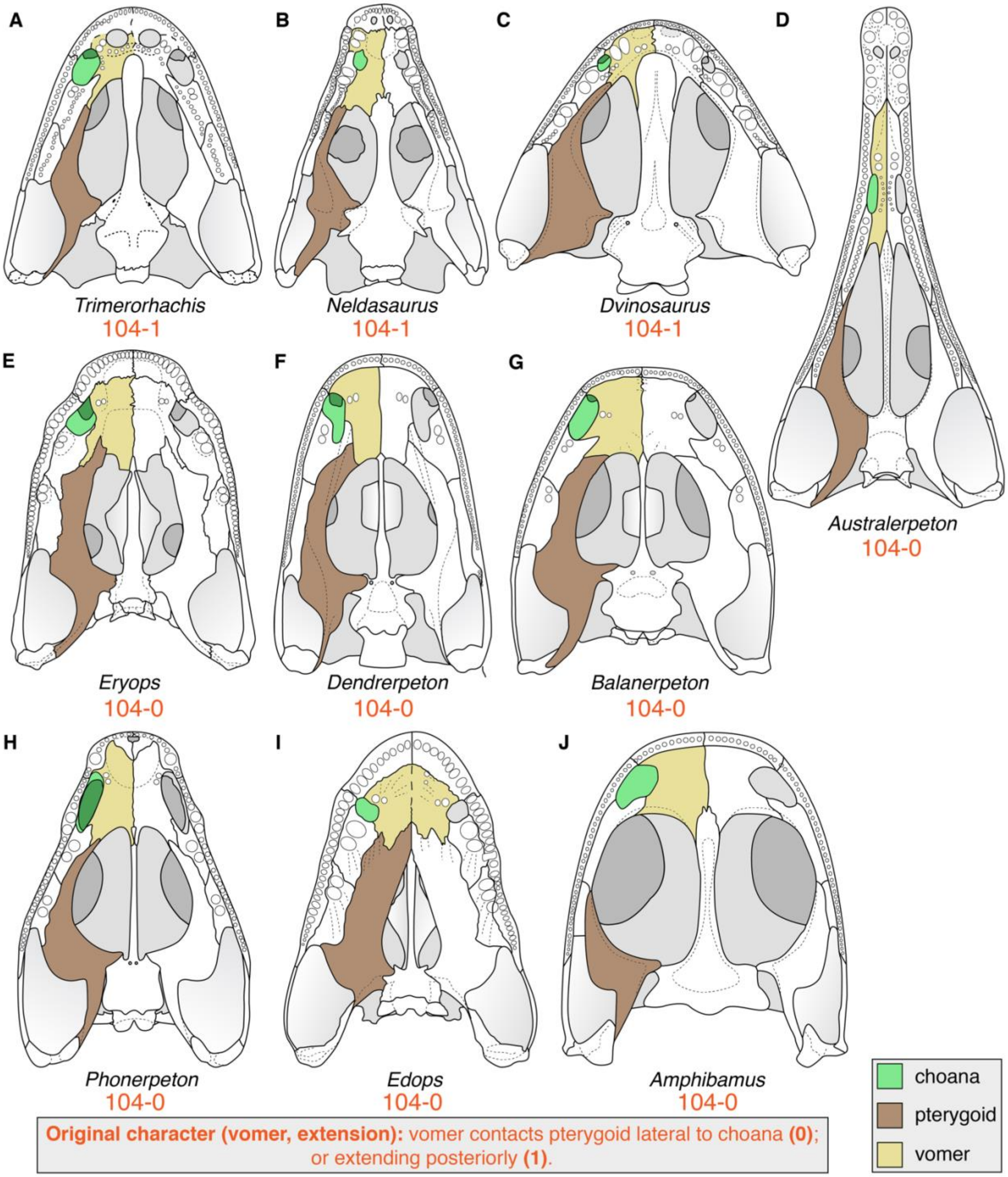

**Figure 20. Comparison of the posterior extent and sutural relationships of the vomer among temnospondyls.** **A**, the dvinosaur *Trimerorhachis insignis* (after Milner & Schoch, 2013); **B**, the dvinosaur *Neldasaurus wrightae* (after Schoch, 2018a); **C**, the dvinosaur *Dvinosaurus primus* (after Schoch & Milner, 2014); **D**, the rhinesuchid *Australerpeton cosgriffi* (after Eltink et al., 2016); **E**, the eryopoid *Eryops megacephalus* (after Sawin, 1941); **F**, the ‘dendrerpetid’ *Dendrerpeton helogenes* (after Holmes et al., 1998); **G**, the ‘dendrerpetid’ *Balanerpeton woodi* (after

Milner & Sequeira, 1993); **H**, the trematopid *Phonerpeton pricei* (after Dilkes, 1990); **I**, the edopoid *Edops craigi* (after Romer & Witter, 1942); **J**, the amphibamid *Amphibamus grandiceps* (after Schoch & Milner, 2014). Orange text represents the original character and codes from Schoch (2013). Silhouettes not to scale.

Across all previous derivates of this matrix, only *Trimerorhachis*, *Neldasaurus*, and *Dvinosaurus* were coded for 104–1 (Fig. 20A-C). There is no apparent condition shared by only these three and not by any other sampled taxon with respect to the vomer-pterygoid contact. One possible way to modify the character would be to refer to the posteriormost extent of the posterolateral end of the vomer, independent of its relationship with the pterygoid. In temnospondyls (but not *Greererpeton*, for example), the vomer extends posterior to the level of the posterior choanal margin, regardless of whether it meets the pterygoid or not, and a modification to this end would serve only to separate the outgroup from the ingroup (but would allow any taxon with a preserved vomer to be coded). Alternatively, the character could be changed to refer to some aspect of vomer-palatine contact; in some taxa, the anteriormost extent of the suture is directly posterior to the choana (e.g., *Platyoposaurus*), whereas in others, it at least extends partially along the medial margin of the choana (e.g., dissorophoids). This is the approach taken here, and in spite of the modifications made to character 248, there is no partial dependency with any of the states of that character.

- Codes were recoded from 104–0 to 104–1 for *Edops*, *Nigerpeton*, *Balanerpeton*, *Acroplous*, *Isodectes*, *Thabanchuia*, *Acanthostomatops*, *Onchiodon*, *Archegosaurus*, *Australerpeton*, *Platyoposaurus*, *Lapillopsis*, *Broomistega*, *Laidleria*, *Plagiosuchus*, *Benthosuchus*, *Trematosaurus*, *Eocyclotosaurus*, and *Cyclotosaurus*.
- Codes were recoded from 104–1 to 104–0 for *Trimerorhachis* and *Dvinosaurus*.
- The following taxa were all recoded from 104–0 to 104–0&1: *Sclerocephalus* (Schoch & Witzmann, 2009a:fig. 4B); *Rhineceps* (Watson, 1962:fig. 7); *Batrachosuchus* (Chernin, 1977:fig. 5b); *Gerrothorax* (Schoch & Witzmann, 2012:fig. 3A); *Edingerella* (Maganuco et al., 2009:fig. 10).
- The three gymnophionomorphs were newly coded for 104–0.
- *Proterogyrinus* was uncoded from 104–0 because the vomer is nearly entirely unknown and thus its sutural relationships are unknown (Holmes, 1984:452). *Limnogyrinus* was uncoded from 104–0 because the sutures are not well-resolved in this area (Schoch & Witzmann, 2018:fig. 3). *Branchiosaurus* was uncoded from 104–0 because the vomer is poorly known (Werneburg, 2012:12). *Iberospondylus* was uncoded from 104–0 because the sutures are not well-resolved in this area (Laurin & Soler-Gijón, 2001:fig. 2; 2006:figs. 1, 5). *Peltobatrachus* was uncoded from 104–0 because the sutures are not well-resolved in this area (Panchen, 1959:figs. 1b, 4). *Uranocentrodon* was uncoded from 104–0 because there are no recent figures of the palate with defined sutures in this region. *Peltostega* was uncoded from 104–0 because the vomer is nearly entirely unknown and thus its sutural relationships are unknown (e.g., Cosgriff & Zawiskie, 1979:fig. 10d; Janvier, 1983). *Paracyclotosaurus* was uncoded from 104–0 because this suture is not fully resolved (Watson, 1958:fig. 1).
- *Rileymillerus* and *Eocaecilia* were newly coded for 104–0.

- **106 (basicranium, contact), 107 (basicranium, suture):** character 107 requires a sutural contact between the parasphenoid and the pterygoid to be present and cannot be coded for any taxa coded for 106–0 (joint between parasphenoid and pterygoid) but was for many such taxa. Based on the original codes, it is inferred that a simple overlapping contact between the parasphenoid and the pterygoid is treated as 106–0 (e.g., *Micropholis*).

  - Codes were corrected from 107–0 (suture [if present] much shorter than basal plate, reaching at best 40% its length) for *Proterogyrinus, Greererpeton, Dendrerpeton, Balanerpeton, Trimerorhachis, Neldasaurus, Acroplous, Isodectes, Dvinosaurus, Capetus, Iberospondylus, Micromelerpeton, Limnogyrinus, Branchierpeton, Branchiosaurus, Apateon, Micropholis, Platyrhinops, Amphibamus, Doleserpeton, Gerobatrachus, Sclerocephalus, Glanochthon, Archegosaurus* and *Platyoposaurus.*
  - All lissamphibians were updated from unknown to inapplicable for character 107 because they were coded for 106–0.

- **107 (basicranium, suture):** the two states fail to capture a wide range of morphologies; 107–0 was defined for a sutural contact less than 40% the length of the basal plate, while 107–1 was defined for a contact nearly as long as the basal plate (this can be reasonably approximated to be around 90% of the length when translated to a quantitative threshold). Many early diverging stereospondyls (brachyopoids, lydekkerinids, rhinesuchids; e.g., Watson, 1962:fig. 7; Chernin, 1977:fig. 5b; Jeannot et al., 2006:fig. 7; Fig. 21C, 21H) have sutures that are slightly more than half as long as the basal plate and therefore fall into neither of these states; these were usually originally coded for 107–0. State 0 is more accurately exemplified by taxa like edopoids and olsoniforms in which the basicranial articulation is formed by a slender bar-like process with a relatively short longitudinal contact that is typically much less than 40% of the length (e.g., Dilkes, 1990:fig. 1B; Sequeira, 2004:fig. 7; Polley & Reisz, 2011:fig. 1; Fig. 21B). Meanwhile, 107–1 is best exemplified by taxa like capitosaurs and trematosaurs in which the contact is more than 80% of the length (e.g., Schoch, 1999:fig. 13; 2019:fig. 7; Schoch & Milner, 2000:figs. 74, 83, 97, 101; Rinehart et al., 2015:fig. 3; Fig. 21E-G). An intermediary state is added for taxa in which the contact is about half the length and no more than three-quarters of the length; it is added between existing states to create an ordered multistate character.

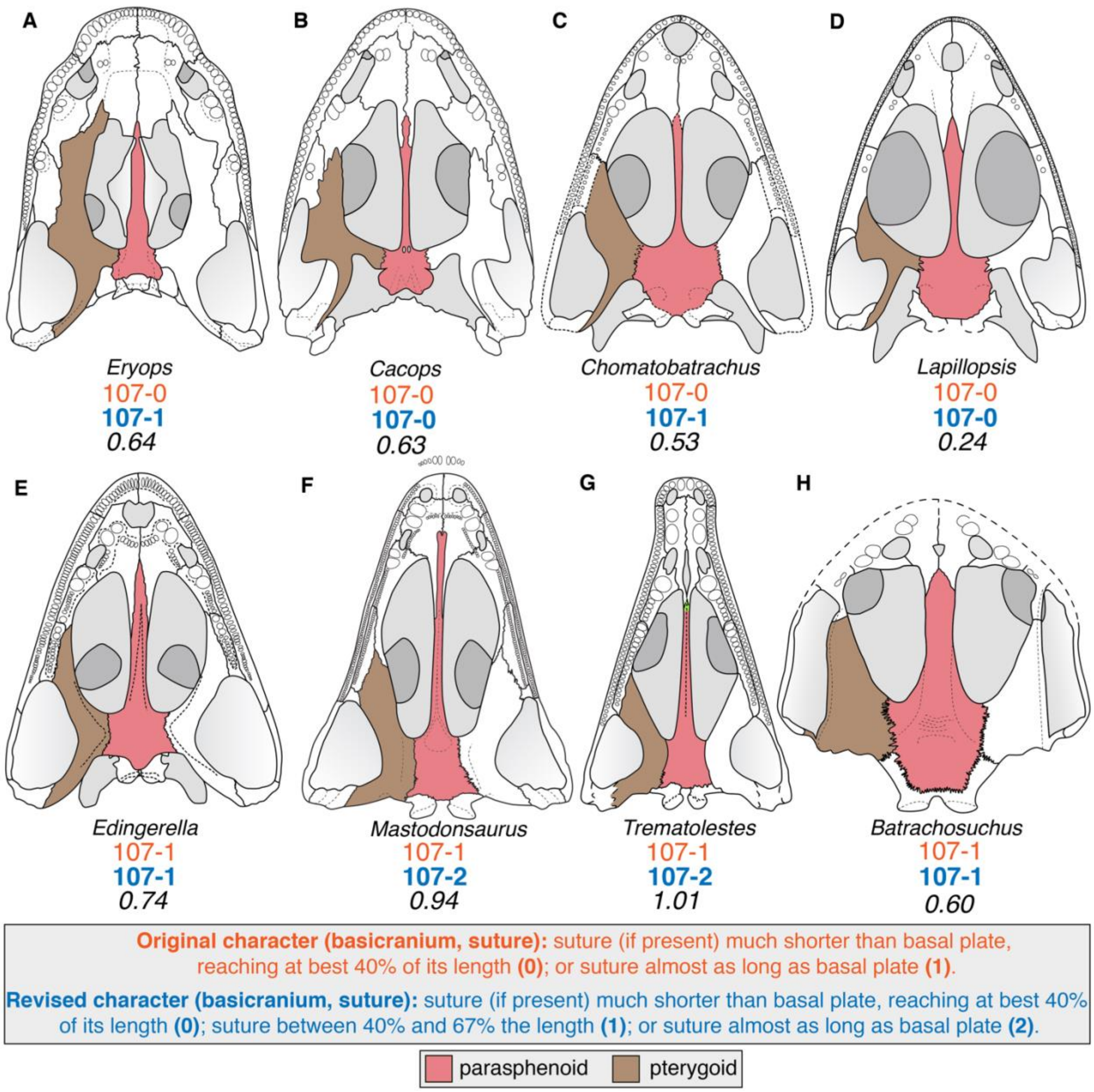

**Figure 21. Comparison of the relative length of the basicranial suture among temnospondyls. A**, the eryopoid *Eryops megacephalus* (after Sawin, 1945); **B**, the dissorophid *Cacops morrisi* (after Reisz et al., 2009); **C**, the lydekkerinid *Chomatobatrachus halei* (after Warren et al., 2006); **D**, the stereospondyl *Lapillopsis nana* (after Yates, 1999); **E**, the capitosaur *Edingerella madagascariensis* (after Maganuco et al., 2009); **F**, the capitosaur *Mastodonsaurus giganteus* (after Schoch, 1999); **G**, the trematosaur *Trematolestes hagdorni* (after Schoch, 2006); **H**, the brachyopid *Batrachosuchus browni* (after Chernin, 1977). Coronoid denticles (where present) are omitted for simplicity. Measurements represent the ratio of basicranial suture length to basal plate length. The suture is measured from end-to-end as a straight line (i.e. ignoring any curvature or complex interdigitation), and basal plate length is measured from the posteriormost extent of the interpterygoid vacuity to the posteriormost extent of the parasphenoid. Orange text represents the original character and codes from Schoch (2013); blue text represents the revised character and codes. Silhouettes not to scale.

- *Acanthostomatops* should not have been coded for the original 107–0 and was recoded to the new 107–1. It has a long and most transversely oriented overlapping contact, and because of the anteroposteriorly short basal plate, the

contact is actually quite long. It differs in this respect from *Zatrachys*, which has the more typical bar-like process found in olsoniforms.
    - *Onchiodon* and *Eryops* both have bar-like processes but also short basal plates such that they are also recoded from 107–0 to the new 107–1.
    - *Australerpeton, Uranocentrodon, Rhineceps, Broomistega, Lydekkerina,* and *Chomatobatrachus* were recoded from 107–0 to the new 107–1.
    - *Tupilakosaurus* was recoded from 107–0 to the revised 107–2.
    - *Gerrothorax, Edingerella, Watsonisuchus, Benthosuchus, Trematosaurus, Trematolestes, Lyrocephaliscus, Callistomordax, Wetlugasaurus, Parotosuchus, Eryosuchus, Eocyclotosaurus, Cyclotosaurus, Mastodonsaurus,* and *Chinlestegophis* were recoded from 107–1 to the revised 107–2.
    - *Laidleria* was recoded from 107–1 to 107–1/2 because of asymmetry of the skull (Warren, 1998a:fig. 4B).
    - *Thabanchuia* was uncoded from 107–0 because most of the suture is reconstructed, and the relationships with the pterygoid and the basioccipital posteriorly are not fully defined (Warren, 1998b:fig. 5). As reconstructed, the suture is well over 50% of the length of the basal plate (107–1/2), but the conservative approach is to treat this as unknown because the condition varies widely among dvinosaurs.
    - *Paracyclotosaurus* was uncoded because the sutures are conjectural (Watson, 1958:fig. 1; Damiani, 2001:fig. 21).
    - *Rileymillerus* was uncoded from 107–1 because Bolt & Chatterjee (2000:676) stated that crushing of this region results in the inability to identify any sutures between the parasphenoid, pterygoid, and exoccipital. Because the exoccipitals can have variable anterior extent (truncating the parasphenoid-pterygoid suture; e.g., *Batrachosuchus browni*; Chernin, 1977) and may contact each other medially (truncating the basal plate), this feature cannot be reliably coded simply on the basis of a broad basicranial articulation.
- **109 (basicranium, carotids):** the carotid foramina are commonly found on the ventral surface of the parasphenoid in early tetrapods, although they can be difficult to identify due to their small size. State 0 (‘ventrally near base of cultriform process’) is common to many non-stereospondyls (e.g., *Balanerpeton*; Milner & Sequeira, 1993). State 1 (‘at posterolateral corner of bone’) is found in taxa like *Sclerocephalus* (Schoch & Milner, 2009). However, many temnospondyls have been described without any trace of carotid foramina, which in stereospondyls, are typically exposed only dorsally, which is not a common exposure (or commonly figured) in stereospondyls (e.g., Sulej, 2007; Eltink et al., 2016; Arbez et al., 2017). Almost all taxa were coded for this character, with a clear division at Eryopiformes; all stereospondylomorphs were coded for 109–1 (eryopids should have been coded for this state as well). All other taxa were coded for 109–0 (except for *Thabanchuia* and *Gerobatrachus*, which were uncoded). This is suggestive of “block coding,” especially because the literature establishes that there are no foramina or grooves anywhere on the ventral surface for some taxa, and others were coded for the wrong state.
    - The following taxa were recoded from 109–0 to unknown because the literature expressly states that there is no ventral expression in adults: *Greererpeton* (Smithson, 1982:54; Godfrey, 1989a:142, fig. 5a); *Adamanterpeton* (Milner &

Sequeira, 1998:10); *Neldasaurus* (Chase, 1965:178; Schoch, 2018a:7); *Isodectes* (Sequeira, 1998:251); and *Micropholis* (Schoch & Rubidge, 2005:508). Given the starkly dichotomous distribution of codes, it is possible that there are other unsubstantiated codes for other taxa for which there is no express description either way and for which figures do not depict foramina (e.g., *Capetus*).

- The following taxa were recoded from 109–1 to unknown because the literature expressly states that no foramina were identified: *Plagiosuchus* (Damiani et al., 2009:359). *Rileymillerus* was recoded from 109–1 to unknown because Bolt & Chatterjee (2000) described in detail how damaged and crushed the basal plate of the parasphenoid was; the extensive fracturing precludes confident determination.
- The following taxa were corrected from 109–1 to 109–0: *Onchiodon* (Boy, 1990:fig. 3); *Eryops* (Sawin, 1941:435–436, 441, pls. 8–10); *Glanochthon* (Schoch & Witzmann, 2009b:fig. 4D); *Archegosaurus* (Witzmann, 2005:147–148, figs. 17, 19); and *Peltobatrachus* (Panchen, 1959:215, fig. 2).

- **112 (cultriform process width); 343 (parasphenoid, cultriform process):** these characters refer to the same feature but with drastically different character states:
    - **112. Cultriform process (width):** Base not wider than rest, clearly set off from basal plate (0), merging continuously into plate (1), flaring anteriorly (2). [*added by Schoch, 2013*]
    - **343. Parasphenoid (cultriform process):** Narrow, tapering anteriorly (0), spatulate and parallel-sided (1). [*added by Pardo et al., 2017a*]

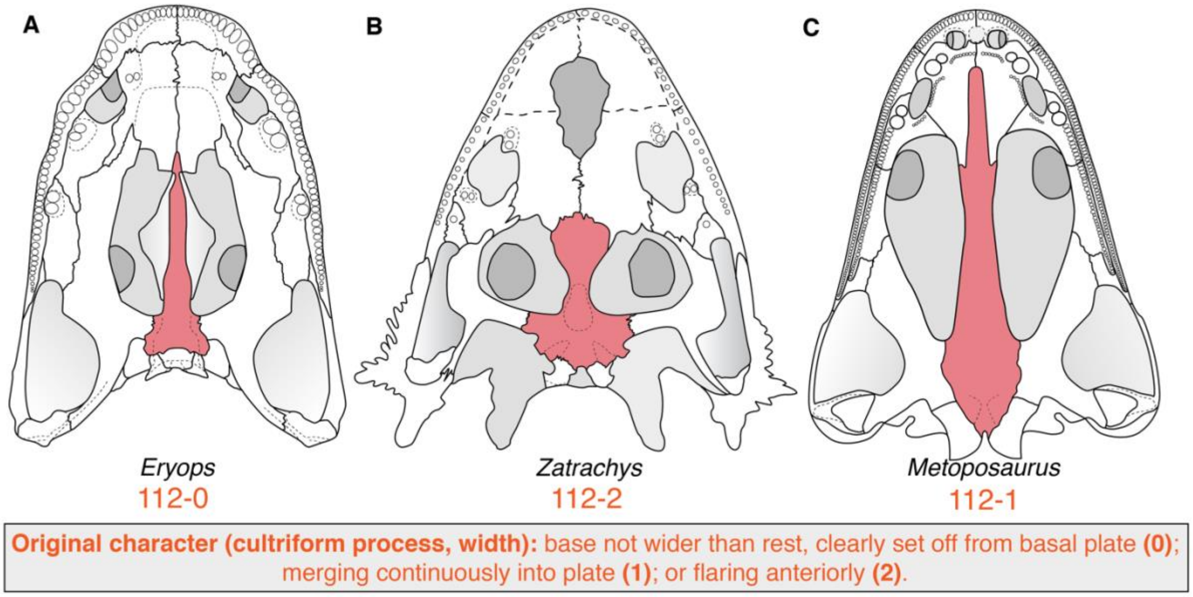

**Figure 22. Comparison of the basal width of the cultriform process among temnospondyls. A**, the eryopoid *Eryops megacephalus* (after Sawin, 1945); **B**, the zatracheid *Zatrachys serratus* (after Schoch & Milner, 2014); **C**, the metoposaurid *Metoposaurus krasiejowensis* (after Sulej, 2007). Coronoid denticles (where present) are omitted for simplicity. Orange text represents the original character and codes from Schoch (2013) and Pardo et al. (2017a). Silhouettes not to scale.

States 112–0 and 112–1 refer to the posterior portion where the cultriform process merges with the basal plate and whether this is smoothly confluent (as in stereospondyls) or more abrupt (as in non-stereospondyls; Fig. 22A). Conversely, 112–2 refers to the anterior portion and is exemplified by the condition seen in somewhat aberrant taxa like *Zatrachys* (e.g., Schoch, 1997:figs. 3–4; Figs. 22B, 23B), although some taxa have a more gradual expansion anteriorly (e.g., *Amphibamus*, *Lapillopsis*; Schoch & Milner (2014:fig. 30B; Yates, 1999:fig. 5B; Fig. 23E). The condition of these taxa is undoubtedly distinct from taxa without an anterior expansion, but 112–2 would be better placed with character 343, which refers to changes to the width throughout the process, rather than with character 112.

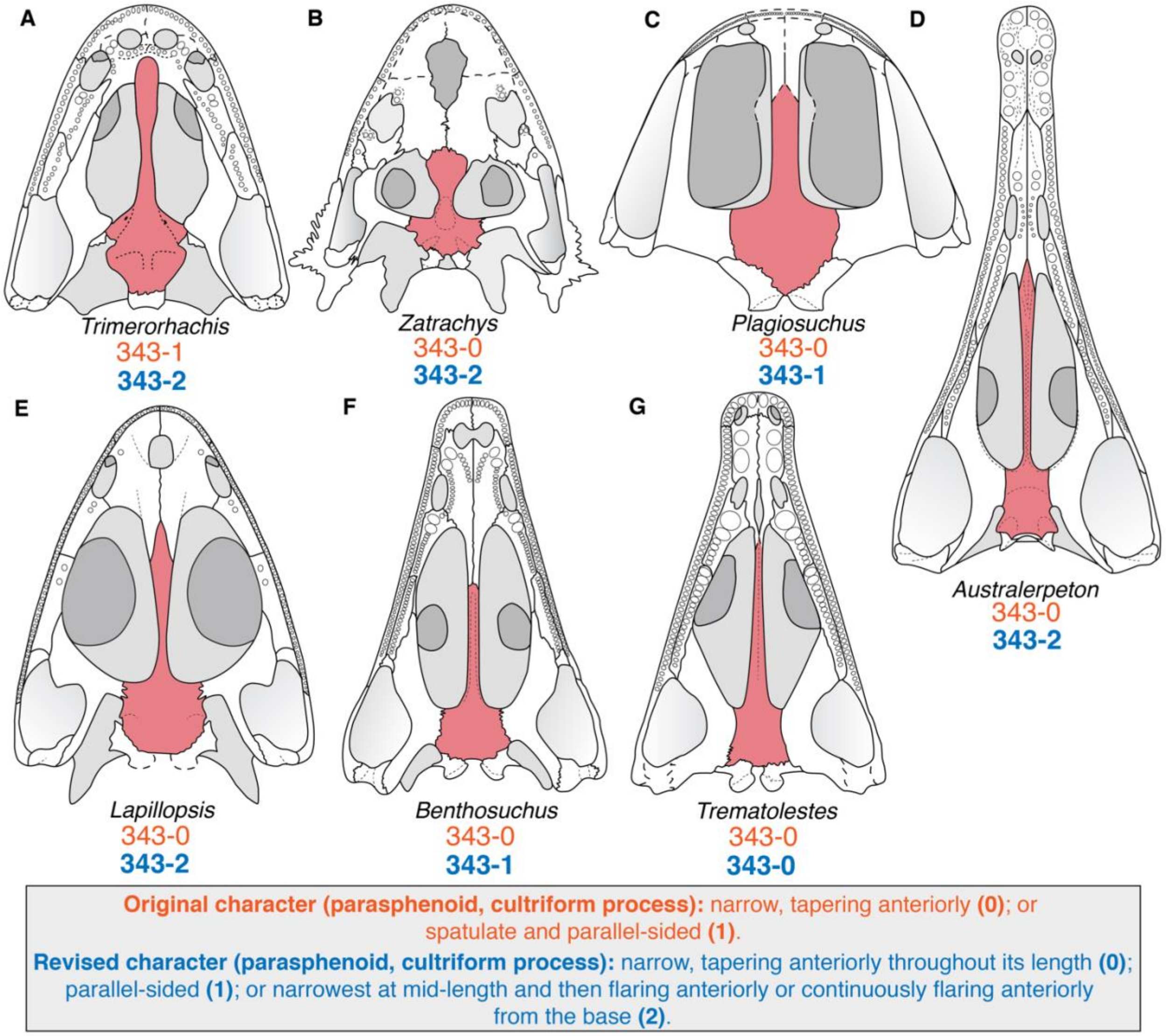

**Figure 23. Comparison of cultriform process width among temnospondyls. Comparison of cultriform process width among temnospondyls. A**, the dvinosaur *Trimerorhachis insignis* (after Milner & Schoch, 2013); **B**, the zatracheid *Zatrachys serratus* (after Schoch & Milner, 2014); **C**, the plagiosaurid *Plagiosuchus pustuliferus* (after Damiani et al., 2009); **D**, the rhinesuchid *Australerpeton cosgriffi* (after Eltink et al., 2016); **E**, the stereospondyl *Lapillopsis nana* (after Yates, 1999); **F**, the trematosaur *Benthosuchus sushkini* (after Schoch, 2008); **G**, the trematosaur *Trematolestes hagdorni* (after Schoch, 2006). Orange text represents the original character and codes from Pardo et al. (2017a); blue text represents the revised character and codes. Coronoid denticles (where present) are omitted for simplicity. Silhouettes not to scale.

I make this change here to make character 112 encapsulate mutually exclusive states referring only to the posterior region and to make character 343 more inclusive, as taxa with an anterior expansion should not have been previously coded for either state (but were). The new 343–2 is slightly reworded for clarity, and 343–1 removes reference to 'spatulate,' as this "flatness" is already captured in character 113 (cultriform process, structure), and 343–0 makes no mention of morphology, only changes in width. The restriction of character 112 to two states referring to the posterior part of the process is also applied to the reanalysis of Schoch (2013) despite the absence of character 343. Future workers may consider further differentiating taxa with a continuous expansion anteriorly (e.g., *Lapillopsis*) from those with a constriction at the mid-length (e.g., *Zatrachys*).

  - Codes were changed from 112–2 to 112–0 for *Balanerpeton, Acroplous, Zatrachys*, *Hynobius,* and *Cryptobranchus*.
  - Codes were changed from 112–2 to 112–1 for *Platyrhinops, Kokartus,* and *Ambystoma.*
  - *Gerobatrachus* (no basal plate is preserved, and the cultriform process is largely absent) and *Sangaia* (no cultriform process preserved) were uncoded from 112–2 and 112–1, respectively.
  - Codes were changed from 343–0 to the new 343–2 for *Edops*, *Acanthostomatops, Zatrachys*, *Platyrhinops*, *Amphibamus*, *Sclerocephalus*, *Glanochthon*, *Archegosaurus*, *Australerpeton, Lapillopsis, Lydekkerina, Edingerella, Rileymillerus,* and *Triadobatrachus*.
  - Codes were changed from 343–0 to the revised 343–1 for *Capetus*, *Micropholis*, *Cacops*, *Platyoposaurus, Broomistega, Plagiosuchus, Benthosuchus, Lyrocephaliscus*, and *Callistomordax*.
  - Codes were changed from 343–1 to the new 343–2 for *Trimerorhachis, Neldasaurus*, *Isodectes, Rhineceps, Laidleria, Karaurus,* and *Kokartus.*
  - Codes were changed from 343–1 to 343–0 for *Epicrionops.*
  - *Micromelerpeton* was newly coded for 343–1&2 (Boy, 1995:fig. 3); *Limnogyrinus* was newly coded for 343–0 (Schoch & Witzmann, 2018); *Cacops* was newly coded for 343–1 (Williston, 1910); *Broiliellus* was newly coded for 343–2 (Williston, 1914); *Chomatobatrachus* was newly coded for 343–2 (Warren et al., 2006); *Trematosaurus* was newly coded for 343–0 (Schoch, 2019a); *Metoposaurus* was newly coded for 343–2 (Sulej, 2007); *Cyclotosaurus* was newly coded for 343–0 (Sulej & Majer, 2005).
  - *Adamanterpeton* was uncoded from 343–0 (most of the process is unknown; Milner & Sequeira, 1998:276); *Nigerpeton* was uncoded from 343–0 (the anterior terminus is unknown; Steyer et al., 2006:22–23); *Gerobatrachus* was uncoded from 343–0 (following character 112); *Paracyclotosaurus* was uncoded from 343–0 (the posterior part of the process is not defined; Watson, 1958:fig. 1).
- **116 (parasphenoid, posterolateral process); 334 (parasphenoid, margins):** both characters appear to refer to the lateral margins of the basal plate of the parasphenoid.
  - **116. Parasphenoid (posterolateral process).** Posterolateral margin straight (0), or with lateral wing (1). [*added by Schoch, 2013*]
  - **334. Parasphenoid (margins)**. Absence of lateral constriction posterior to basicranial articulations (0), weak constriction posterior to basicranial articulations present (1), strong constriction posterior to basicranial articulations, reaching to the carotid foramina (2). [*added by Pardo et al., 2017a*]

The posterolateral wing (116–1) is an inconsistently defined and identified feature; Jeannot et al. (2006:829), who termed these ‘palatal lappets,’ considered these “wings” to really be lateral extremities of the crista muscularis, which frames a posteroventrally facing pocket and which produces an anteriorly situated notch in the lateral surface of the parasphenoid just behind the basicranial articulation, as in *Lydekkerina* (Fig. 24E). Eltink et al. (2016:849), who termed these ‘lateral flanges,’ indicated the presence of the corresponding notches in taxa other than lydekkerinids, rhinesuchids, and *Peltobatrachus*, the only taxa for which 116–1 was coded in this matrix (e.g., these flanges also occur in *Lapillopsis*; Fig. 24C). Meanwhile, *Chomatobatrachus*, coded for 116–1, does have cristae and pockets, but the lateral extent of the crista does not produce a discontinuous lateral margin; the basal plate tapers continuously posteriorly (Warren et al., 2006:fig. 4; Fig. 24F). A similar condition is shown in *Trimerorhachis*, which has the crista muscularis but not the deep pockets found in stereospondyls (Milner & Schoch, 2013:fig. 1D; Fig. 24B) but which was coded for 116–0 (posterolateral margins straight).

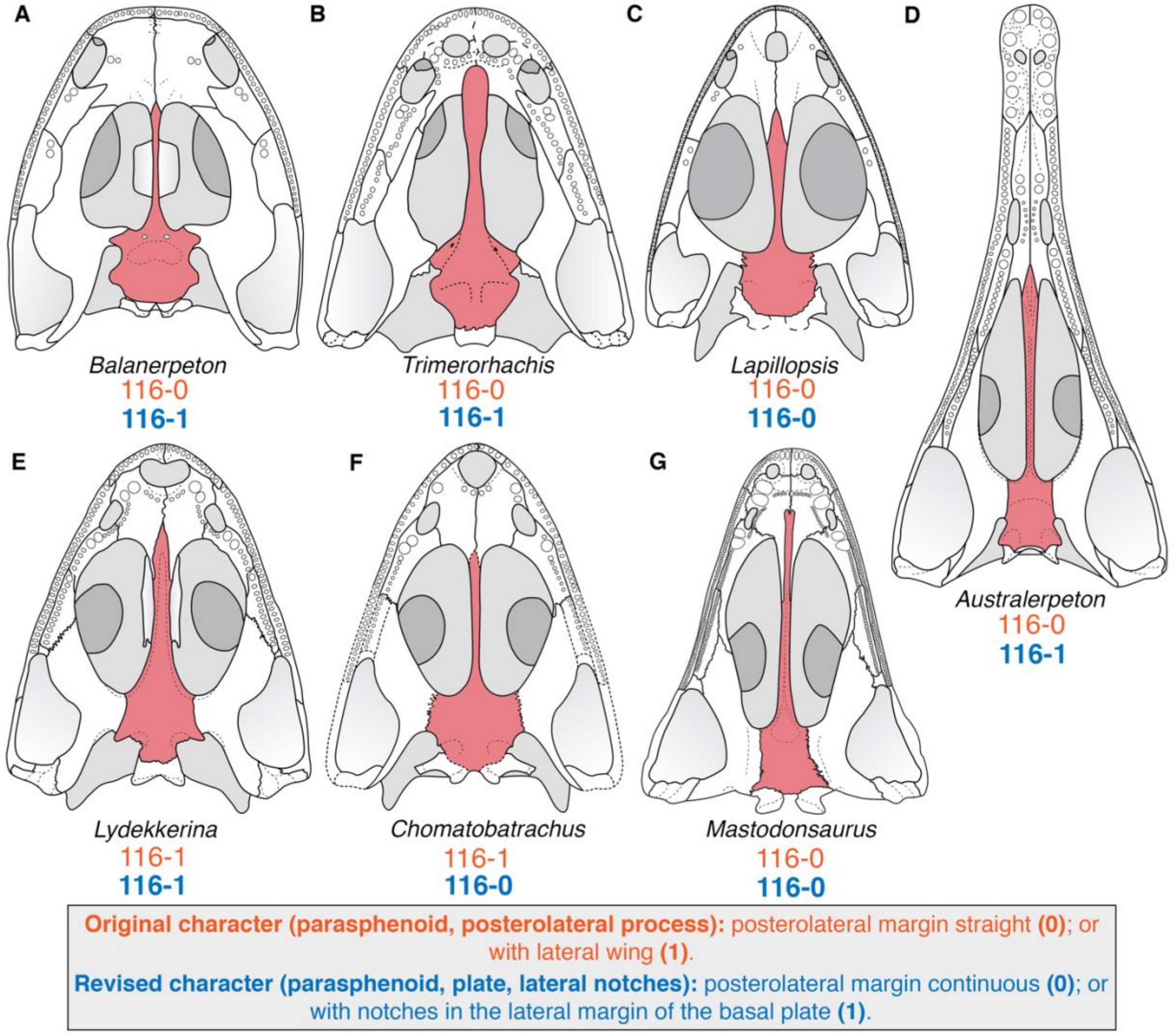

**Figure 24. Comparison of the posterolateral margin of the parasphenoid among temnospondyls. A**, the ‘dendrerpetid’ *Balanerpeton woodi* (after Milner & Sequeira, 1993); **B**, the dvinosaur *Trimerorhachis insignis* (after Milner & Schoch, 2013); **C**, the stereospondyl *Lapillopsis nana* (after Yates, 1999); **D**, the rhinesuchid *Australerpeton cosgriffi* (after Eltink et al., 2016); **E**, the lydekkerinid *Lydekkerina huxleyi* (after Hewison, 2007); **F**,

the lydekkerinid *Chomatobatrachus halei* (after Warren et al., 2006); **G**, the capitosaur *Mastodonsaurus giganteus* (after Schoch, 1999). Coronoid denticles (where present) are omitted for simplicity. Orange text represents the original character and codes from Schoch (2013); blue text represent the revised character and codes. Silhouettes not to scale.

A different condition is found in *Dendrerpeton* (Holmes et al., 1998:fig. 4b), *Balanerpeton* (Milner & Sequeira, 1993:fig. 5B; Fig. 24E); *Platyrhinops* (Clack & Milner, 2010:fig. 9b); *Micropholis* (Schoch & Rubidge, 2005:fig. 2); *Dissorophus* (Schoch, 2012:fig. 3D); *Cacops* (Williston, 1910:pl. 11); *Neldasaurus* (Schoch, 2018a:fig. 3D); *Cochleosaurus* (Sequeira, 2004:fig. 7); and *Zatrachys* (Schoch, 1997:fig. 3) in which there are no cristae or pockets, but the parasphenoid does expand posterolaterally, forming the same notch posterior to the basicranial articulation. If anything, it is these taxa that have a 'lateral wing,' rather than taxa where the crista projects slightly. However, these taxa were also coded for 116–0. If the character really refers to whether the cristae project laterally off the edge of the plate, then it would (1) need to be coded for additional taxa like *Australerpeton* and *Lapillopsis*; and (2) should be rewritten as this kind of character (along the lines of at least one of characters 90–93 of Marsicano et al., 2017, or characters 105–108, 122 of Eltink et al., 2019). As is, character 116 is not only confusing, but it is also not fully independent of character 334. Any taxon coded for 334–1 or 334–2 (weak or strong lateral constriction posterior to basicranial articulations) should be coded for 116–0.

- This exact 'lateral wing' character was sampled by Eltink et al. (2019, as character 110). It was coded in most sampled taxa, including all rhinesuchids and in various non-stereospondyls, including *Trimerorhachis*, *Eryops*, *Archegosaurus*, and *Platyoposaurus stuckenbergi*, but not in *Peltobatrachus* or *Chomatobatrachus* (contra Schoch, 2013). *Lapillopsis* was left as unknown, also contra Schoch (2013).
    - To ameliorate the ambiguity, character 116 is adjusted to refer to laterally facing notches in the lateral surface just posterior to the basicranial articulation. Notches must be framed on both ends; the continually tapering basal plates found in some stereospondyls are not considered to have notches. There are established characters to expand on the morphology of the crista muscularis and the parasphenoidal pockets that are sampled in other matrices but that are not sampled here following the conservative approach to modifications. All taxa were recoded, as there are frequently notches in non-stereospondyls as well.
- In the original coding of character 334 (lateral constriction of parasphenoid) by Pardo et al. (2017a), only one taxon was coded for 334–1 '(weak constriction'; *Acroplous*) and one for 334–2 ('strong constriction'; *Leptodactylus*). Despite being coded for 334–0 ('no constriction'), most temnospondyls exhibit at least minor constriction of the parasphenoid posterior to the basicranial articulation (Fig. 25), either being bordered by medially expanding exoccipitals (e.g., *Callistomordax*; Schoch, 2008a:fig. 1B) or with gradually tapering margins in the absence of this contact laterally (e.g., *Sclerocephalus*; Schoch & Witzmann, 2009a:fig. 4B).

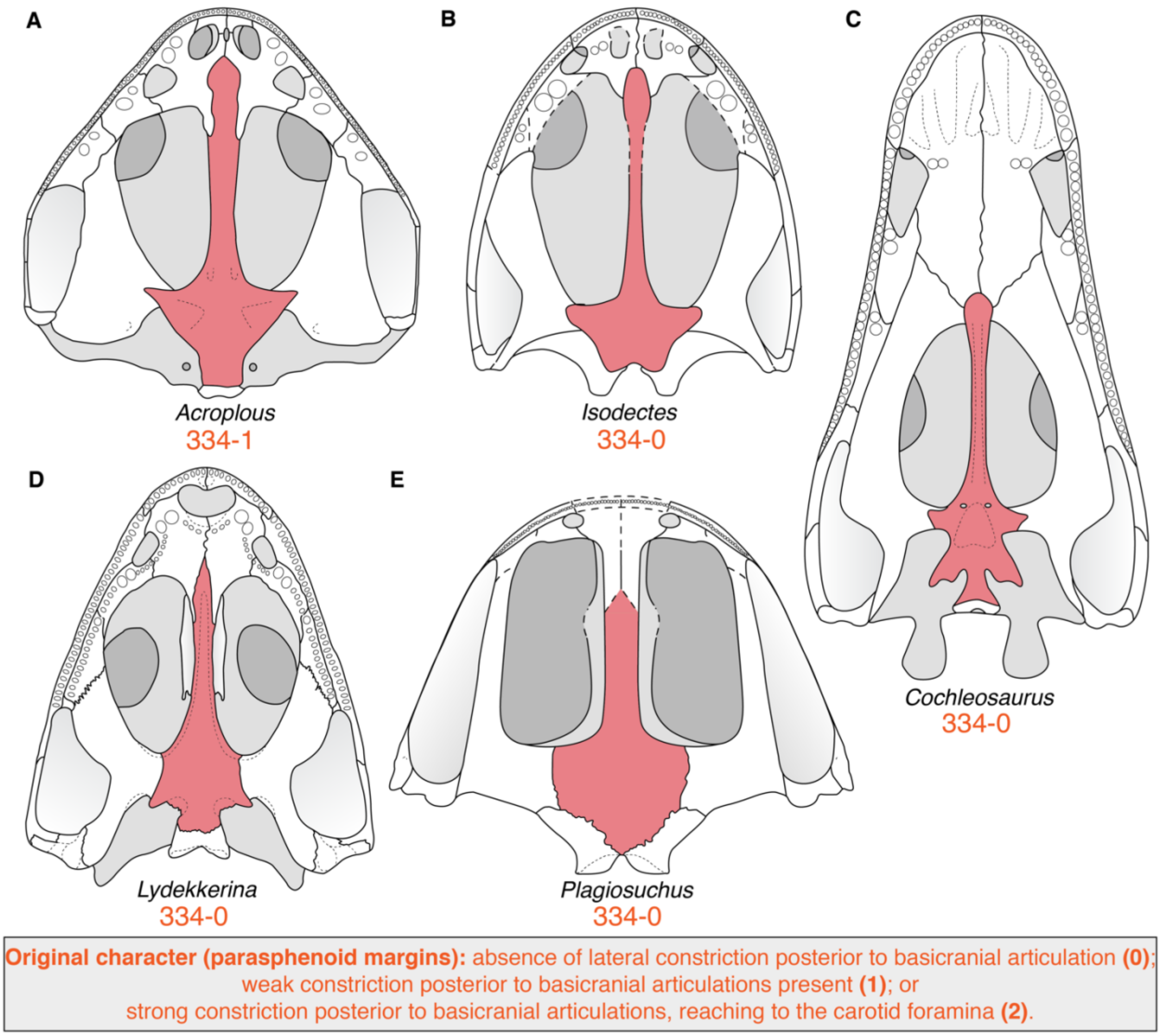

**Figure 25. Comparison of the posterior width of the parasphenoid among temnospondyls. A**, the dvinosaur *Acroplous vorax* (after Englehorn et al., 2008); **B**, the dvinosaur *Isodectes obtusus* (after Sequeira, 1998); **C**, the edopoid *Cochleosaurus bohemicus* (after Sequeira, 2004); **D**, the lydekkerinid *Lydekkerina huxleyi* (after Hewison, 2007); **E**, the plagiosaurid *Plagiosuchus pustuliferus* (after Damiani et al., 2009). Orange text represents the original character and codes from Pardo et al. (2017a). Silhouettes not to scale.

Any taxon with a lateral notch associated with a 'lateral wing' of character 116 (either in the original or in the revised formulations) inherently has lateral constriction, even if there is more posterior lateral expansion to form a notch. *Isodectes* has straight, obliquely oriented lateral margins (Sequeira, 1998:fig. 9B; Fig. 25B) and is more similar to *Acroplous* than to other temnospondyls but was coded for 334–0. This character is further complicated by any taxon with a short posterior process of the parasphenoid that divides the exoccipitals along the midline (e.g., *Chenoprosopus*; Langston, 1953:fig. 8). It is thus apparent that 334–0 captured a wide range of morphologies that are not clearly homologous due to a lack of precision. Finally, there is the issue that using the carotid foramina as a specifier in state 2 is problematic because the position of the foramina (if they are exposed ventrally at all) is highly variable; they may be as far anterior as the base of the cultriform process or as posterior as the posterolateral corner of the basal plate (original character 109).

        - Character 334 could be redefined in a number of ways, but it is apparent that the contour of the lateral margins is too complex in some taxa and too variable among temnospondyls to be properly discretized into a single character that is fully distinct from either the original or the revised character 116. Because it was previously parsimony-uninformative, the character is omitted here.

- **119 (pterygoid, basioccipital); 136 (occipital condyle); 138 (basioccipital, length):** any taxon without an ossified basioccipital cannot be coded for these characters.
    - Codes were corrected from 119–0 (no pterygoid-basioccipital contact) for *Acanthostomatops*, *Limnogyrinus, Branchierpeton, Branchiosaurus, Apateon, Micropholis, Platyrhinops, Amphibamus, Doleserpeton, Lapillopsis, Broomistega, Sangaia*, *Peltostega, Laidleria, Siderops, Batrachosuchus, Gerrothorax, Watsonisuchus, Benthosuchus, Callistomordax, Metoposaurus, Parotosuchus, Eryosuchus, Eocyclotosaurus, Triadobatrachus, Leptodactylus, Xenopus, Karaurus, Kokartus, Ambystoma, Hynobius*, and *Cryptobranchus.* Codes were corrected from 119–1 (pterygoid-basioccipital sutured lateral to parasphenoid) for *Rileymillerus.*
    - Similar to the absence of ossified exoccipitals, the absence of an ossified basioccipital is phylogenetically informative. A third state was added to character 138 for the absence of a basioccipital, as this is phylogenetically informative for differentiating paedomorphic taxa, and this state fits better with this character than with character 119. Provided that the occiput is largely complete and undistorted, taxa without described or figured basioccipitals were coded for this state. Codes were recoded from 138–1 (basioccipital foreshortened to a narrow posterior rim of the palatal bone) to the new 138–2 for *Acanthostomatops*, *Limnogyrinus, Branchierpeton, Branchiosaurus, Apateon, Micropholis, Platyrhinops, Amphibamus, Doleserpeton, Lapillopsis, Broomistega, Sangaia, Peltostega, Laidleria, Siderops, Batrachosuchus, Gerrothorax, Watsonisuchus, Benthosuchus*, *Callistomordax*, *Metoposaurus*, *Parotosuchus*, *Eryosuchus, Eocyclotosaurus*, *Rileymillerus*, and *Triadobatrachus*. Codes were updated from unknown to the new 138–2 for *Leptodactylus, Xenopus, Karaurus, Kokartus, Ambystoma*, *Hynobius,* and *Cryptobranchus*.
    - Character 136 prescribes two conditions for the occipital condyle, one with a major basioccipital contribution to form a trilobed condyle and one with a reduced contribution. Regardless of whether an unossified basioccipital (the new 138–2) is considered as 'reduced' (136–1) or not, any taxon coded for 138–2 should not be coded for character 136, either because of a partial dependency (if 'absent' is considered to fall under 'reduced') or because of an inapplicability (if 'absent' is not considered to fall under 'reduced'). To avoid having to recode many taxa for this character, the mention of the basioccipital was omitted from
    - this character.
- **122 (pterygoid width):** the original coding of character 122 by Schoch (2013) is difficult to interpret despite being strongly dichotomous.

- **122. Pterygoid (width).** Palatine and quadrate regions forming slender rami (0), or broad shelves (1). [*added by Schoch, 2013*]

Broad rami (122–1) were coded for the majority of post-lydekkerinid stereospondyls. Within this, there is noticeable variation, particularly within capitosaurs (Fig. 26F-H); *Edingerella*, *Watsonisuchus*, *Wetlugasaurus*, *Parotosuchus,* and *Eryosuchus* were coded as having slender rami (122–0).

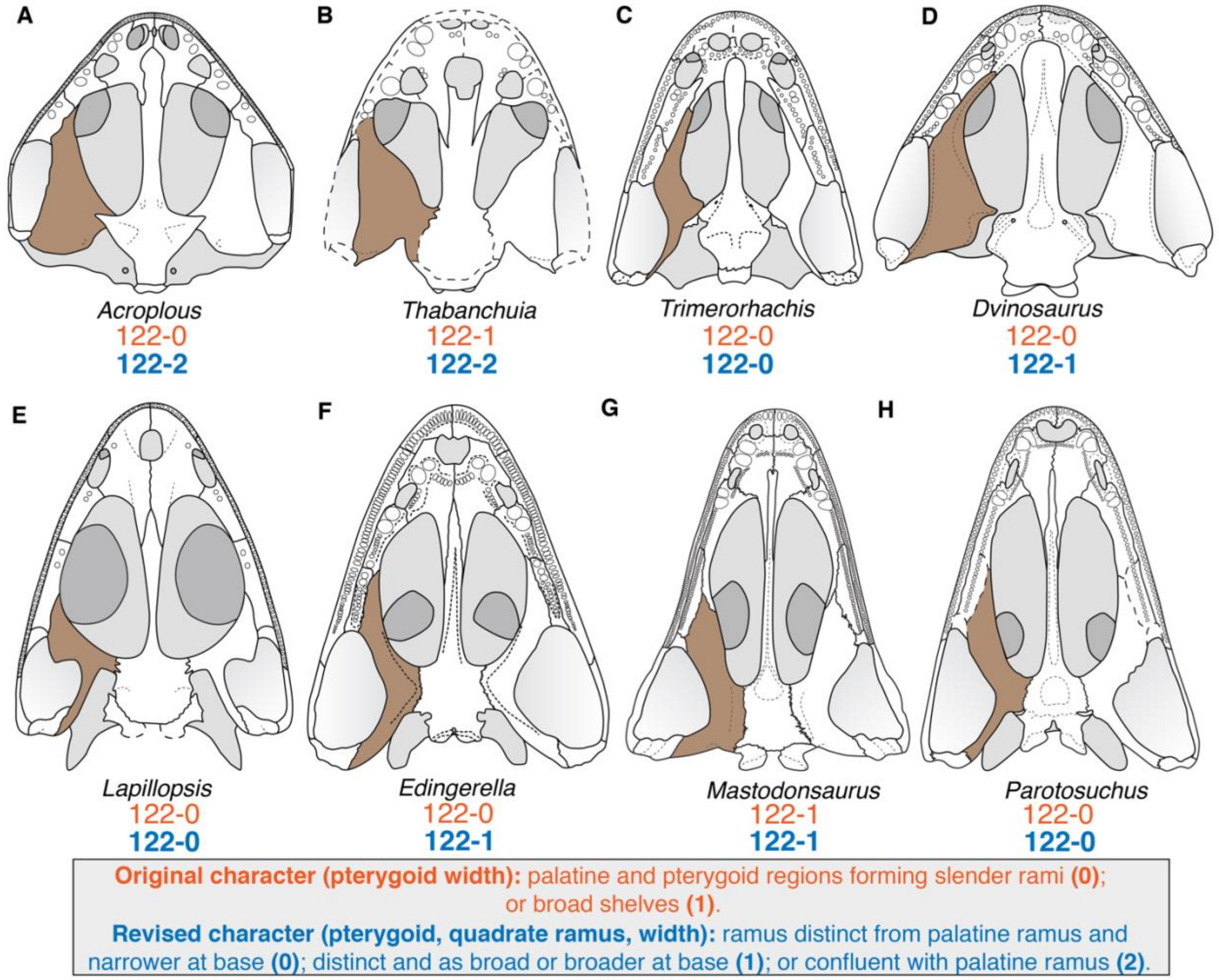

**Figure 26. Comparison of the relative width of the palatine and quadrate rami among temnospondyls. A**, the dvinosaur *Acroplous vorax* (after Englehorn et al., 2008); **B**, the dvinosaur *Thabanchuia oomie* (after Warren, 1998b); **C**, the dvinosaur *Trimerorhachis insignis* (after Milner & Schoch, 2013); **D**, the dvinosaur *Dvinosaurus primus* (after Schoch & Milner, 2014); **E**, the stereospondyl *Lapillopsis nana* (after Yates, 1999); **F**, the capitosaur *Edingerella madagascariensis* (after Maganuco et al., 2009); **G**, the capitosaur *Mastodonsaurus giganteus* (after Schoch, 1999); **H**, the capitosaur *Parotosuchus nasutus* (after Schoch, 2018). Coronoid denticles (where present) are omitted for simplicity. Orange text represents the original character and codes from Schoch (2013); blue text represent the revised character and codes. Silhouettes not to scale.

Differentiation of states appears largely related to the quadrate ramus, which can be mostly vertically oriented and therefore slender in ventral view in many non-stereospondyls (e.g., *Balanerpeton*, *Dendrerpeton*; Milner & Sequeira, 1993: figs. 1, 5B; Holmes et al., 1998:fig. 4B), rather than mostly horizontal and thus wide in ventral view. The palatine ramus is always more or less horizontally oriented unless it has a ventrally angled transverse flange, and it is always as wide or wider than the quadrate ramus, especially in taxa with a transverse flange of the pterygoid. However, this does not clarify how width is being assessed – is it the relative width of the two rami (more disparate in non-stereospondyls) or relative to some unspecified palatal landmark? Comparative figures do not clarify why some capitosaurs were coded for 122–0 and others were coded for 122–1 (e.g., comparison of *Benthosuchus* and *Mastodonsaurus* by Schoch, 2008b:fig. 5). All capitosaurs other than *Wetlugasaurus* and perhaps *Parotosuchus* definitively have wider quadrate rami than in the plesiomorphic condition (e.g., Smithson, 1982:figs. 6, 11; Holmes, 1984:fig. 3). Also peculiarly coded are many dvinosaurs, which have a broad yet deeply vaulted quadrate ramus that is essentially indistinguishable from the palatine ramus along their lateral edge (e.g., Shishkin, 1961:fig. 2; Warren, 1998b:fig. 5B; Englehorn et al., 2008:fig. 7B). State 1 was only coded in the tupilakosaurids *Thabanchuia* and *Tupilakosaurus*, but the condition is essentially the same in *Acroplous* (Fig. 26A-B). In *Dvinosaurus* and *Isodectes*, the ramus is confluent with the palatine ramus anterolaterally and therefore quite wide in this region, but it tapers sharply posterolaterally (Fig. 26D). Finally, *Proterogyrinus*, on account of having very narrow and small interpterygoid vacuities, was coded for 122–0. It could only be conceivably coded for state 0 if the coding was exclusively based on the distal end of the quadrate ramus.

  - The character is linguistically restricted to the quadrate ramus to avoid a compound character that could warrant many taxa being coded as inapplicable. Both the original and the revised versions are somewhat redundant with character 353 of Schoch et al. (2020), which, based upon how it was originally coded, relates to the relative length of the quadrate ramus (but which linguistically invokes both length and orientation in ventral view). Therefore, character 122 is expressly restricted to the width to further differentiate them, and this is assessed specifically by comparison with the width of the palatal ramus using the base of each process as the reference point. This relative width comparison allows for a more reproducible treatment while minimizing modifications to existing codes. Finally, the use of the 'base' of each ramus makes it difficult to code taxa like *Acroplous* and *Siderops* in which the quadrate and palatine rami are essentially undifferentiated because the lateral margin is nearly straight. To address this condition, a third state was added in which the quadrate and palatine rami are essentially merged along the lateral margin.
  - Codes were recoded from 122–0 to the revised 122–1: *Edingerella, Watsonisuchus*, *Benthosuchus, Eryosuchus, Rileymillerus*, *Karaurus,* and *Kokartus*.
  - Codes were recoded from 122–0 to the new 122–2: *Dvinosaurus* and *Acroplous*.
  - Codes were recoded from 122–1 to the new 122–2: *Thabanchuia*, *Tupilakosaurus*, *Siderops*, *Batrachosuchus*, *Plagiosuchus*, *Ichthyophis*, and *Epicrionops*.
- **124 (pterygoid-palatine-ectopterygoid); 125 (palatine, ectopterygoid):** any taxon coded for 125–1 ('palatine with posterolateral process excluding the ectopterygoid from interpterygoid vacuity and contacting pterygoid') is assured of state 124–0 ('pterygoid contacting both ectopterygoid and palatine') unless either the palatine or the ectopterygoid is absent, in which case these characters cannot be coded.
  - To remove this dependency, the language of character 125 was altered to remove any mention of the palatine contacting the pterygoid in state 1; a posteromedial

process (resulting in a suture that is not straight) can be present yet not contact the pterygoid, as in *Adamanterpeton* (Milner & Sequeira, 1998:fig. 10) or *Lydekkerina* (Hewison, 2007:fig. 32). This avoided having to change any existing codes for 124–0 to inapplicable. I also clarified state 0 as referring to a straight suture; many taxa have sutures without a posteromedial process that are not transverse but set at sharply oblique angles.

- **128 (palatine, ectopterygoid (ontogeny)); 180 (interclavicle, ontogeny); 190 (humerus, ontogeny):** these characters express ontogeny. Therefore, any taxon for which no ontogenetic trajectory is available (either in general or for the specific region/element) should not be coded. This inherently applies to any taxon only represented by one specimen (e.g., *Batrachosuchus/Bathignathus*, *Gerobatrachus*, *Siderops*) but also to any taxon only represented by specimens of the same size class (e.g., *Thabanchuia*) and to any taxon where the specified region is only known from specimens of a similar size (e.g., *Broiliellus* for the interclavicle).
    - Character 128: Codes were corrected for *Edops*, *Adamanterpeton, Nigerpeton, Thabanchuia*, *Tupilakosaurus, Capetus, Iberospondylus, Gerobatrachus, Broiliellus, Peltobatrachus, Sangaia, Laidleria, Siderops, Batrachosuchus, Watsonisuchus,* and *Paracyclotosaurus*.
    - Character 180: Codes were corrected for *Acroplous, Thabanchuia*, *Capetus, Broiliellus, Rhineceps, Laidleria, Siderops*, and *Paracyclotosaurus*.
    - Character 190: Codes were corrected for *Ecolsonia* and *Trematolestes.*
- **137 (exoccipital condyles):** this character refers to the exposure of the exoccipitals in dorsal view.
    - **137. Exoccipital condyles.** Short and broad base, projecting only with their posterior half behind the rim of the skull table (0), or almost the complete element posterior to level of occipital flange (1). [*added by Schoch, 2013*]

The wording of this character is somewhat confusing, as it is really the entire exoccipital in dorsal view, rather than specifically the condyles; when the exoccipital is dorsally exposed, it is related to the sloping of the occiput, not to the length of the condyles. The other problem is that 137–0, which refers to the less visible condition, does not account for when the exoccipitals are entirely obscured in dorsal view; this is common in early diverging Paleozoic taxa (e.g., edopoids, eryopoids, 'dendrerpetids,' many dissorophoids; e.g., Sequeira, 2004:fig. 7; Steyer et al., 2006:figs. 2–3; Schoch, 2012:fig. 2). This condition is not homologous with that observed in other taxa in which the condyles but little else of the exoccipital (i.e. the paraoccipital process or the ascending column) is exposed dorsally, as in many stereospondylomorphs (e.g., Shishkin & Rubidge, 2000:fig. 2; Maganuco et al., 2009:fig. 6; Eltink et al., 2016:fig. 5). Certain taxa like brachyopoids and some dvinosaurs then have a markedly projecting/sloping occiput that exposes practically the entire exoccipital in dorsal view (e.g., Englehorn et al., 2008:fig. 7; Damiani et al., 2009:fig. 7; Schoch & Witzmann, 2012:fig. 1; Pardo et al., 2017a:fig. 1J). It is also complicated in lissamphibians because the exoccipital typically co-ossifies with other elements to form a large, dorsally exposed os basale.

To remedy this, state 0 is restricted to taxa with the exoccipitals entirely obscured in dorsal view (any taxon without ossified exoccipitals should be treated as inapplicable). State 1 refers to a limited exposure of primarily the condylar region, and the new state 2 is the former state 1 and refers to a broad exposure. Importantly, any taxon with an os basale should not be coded for this character because the exoccipital portion is undefined. Additionally, there is a separate character (335) that refers to the dorsal exposure of the os basale.

  - Codes were recoded from 137–0 to 137–2 for *Proterogyrinus, Dvinosaurus, Acroplous, Isodectes, Thabanchuia, Tupilakosaurus, Chomatobatrachus, Laidleria, Cyclotosaurus,* and *Mastodonsaurus*.
  - Code s were recoded from 137–0 to the revised 137–1 for *Greererpeton, Trimerorhachis, Doleserpeton, Acheloma, Phonerpeton, Eryops, Sclerocephalus, Australerpeton, Peltobatrachus, Broomistega, Lydekkerina, Sangaia, Edingerella, Benthosuchus, Watsonisuchus, Trematolestes, Wetlugasaurus, Parotosuchus, Paracyclotosaurus, Karaurus,* and *Kokartus*.
  - Codes recoded from 137–0 to 137–1/2 for *Platyoposaurus, Eryosuchus,* and *Eocyclotosaurus* were.
  - *Trematosaurus* was recoded from 137–0 to 137–1&2 (Schoch, 2019a:53, fig. 6A).
  - Codes were recoded from 137–1 to the new 137–2 for *Batrachosuchus, Plagiosuchus, Gerrothorax, Callistomordax, Metoposaurus, Chinlestegophis,* and *Rileymillerus*.
  - Codes were recoded from 137–1 to 137–1/2 for *Peltostega* and *Lyrocephaliscus*.
  - *Dendrerpeton* was uncoded from 137–0 because none of the specimens attributed to *D. helogenes* (sensu Schoch & Milner, 2014, and Arbez et al., 2022) have the exoccipitals. *Gerobatrachus* was uncoded from 137–0 because the exoccipitals are unknown in the holotype. *Iberospondylus* was uncoded because the exoccipitals and basioccipital are dislodged in known specimens. *Siderops* was uncoded from 137–1 because the posterior skull margin is unknown.
  - *Triadobatrachus*, *Leptodactylus*, *Xenopus*, *Hynobius*, *Ichthyophis*, and *Epicrionops* were recoded from 137–0 to inapplicable on the basis of having a co-ossified exoccipital. *Ambystoma*, *Cryptobranchus*, and *Eocaecilia* were recoded from 137–1 to inapplicable on the same basis.
  - *Karaurus* and *Kokartus* were newly coded for 137–1.

- **141 (postglenoid area, types):** the classification of postglenoid areas (PGAs) was originally developed by Warren & Black (1985) and Jupp & Warren (1986), specifically for Triassic stereospondyls. Existing codes are maintained for non-stereospondyl temnospondyls, but future workers may want to consider whether this scheme can really be applied to the entire clade, notwithstanding the general consensus that this binary is oversimplified and that features of both types can occur within a single individual, let alone a single taxon (e.g., Schoch, 1999:80; Sulej, 2007:60, and see alternative scheme of Maryanska & Shishkin, 1996). The extensive consolidation of the gymnophionomorph jaw into the pseudoangular means that this classification scheme is particularly inappropriate for such taxa since it relies on the differentiation of posterior lower jaw elements.
  - Codes were uncoded from 141–1 (type 2 PGA) for *Ichthyophis*, *Epicrionops*, and *Eocaecilia*.
  - *Proterogyrinus* and *Greererpeton* were uncoded from 141–0 as the outgroups since their lower jaws differ markedly from temnospondyls (e.g., Holmes, 1984; Ahlberg & Clack, 1998; Bolt & Lombard, 2001).

- **144 (preglenoid process):** all taxa coded for 262–2 (surangular absent) cannot be coded for this character because it specifies the labial side of the surangular in both states.
    - **144. Preglenoid process.** Labial side of surangular with straight dorsal margin anterior to glenoid (0), or forming dorsal projection well above the level of the glenoid articulation (1). [*added by Schoch, 2013*]
    - Codes were corrected from 144–0 (labial side of surangular with straight dorsal margin anterior to glenoid) for *Triadobatrachus*. Codes were corrected from 144–1 (labial side of surangular forming dorsal projection well above the level of the glenoid articulation) for *Ambystoma*, *Hynobius*, and *Cryptobranchus*. Codes were updated from unknown to inapplicable for *Leptodactylus* and *Xenopus*.

Originally, only *Trematosaurus*, *Trematolestes*, *Callistomordax*, and *Metoposaurus* were coded for 144–1 (Fig. 27A-D). However, the condition among them varies widely, as shown below.

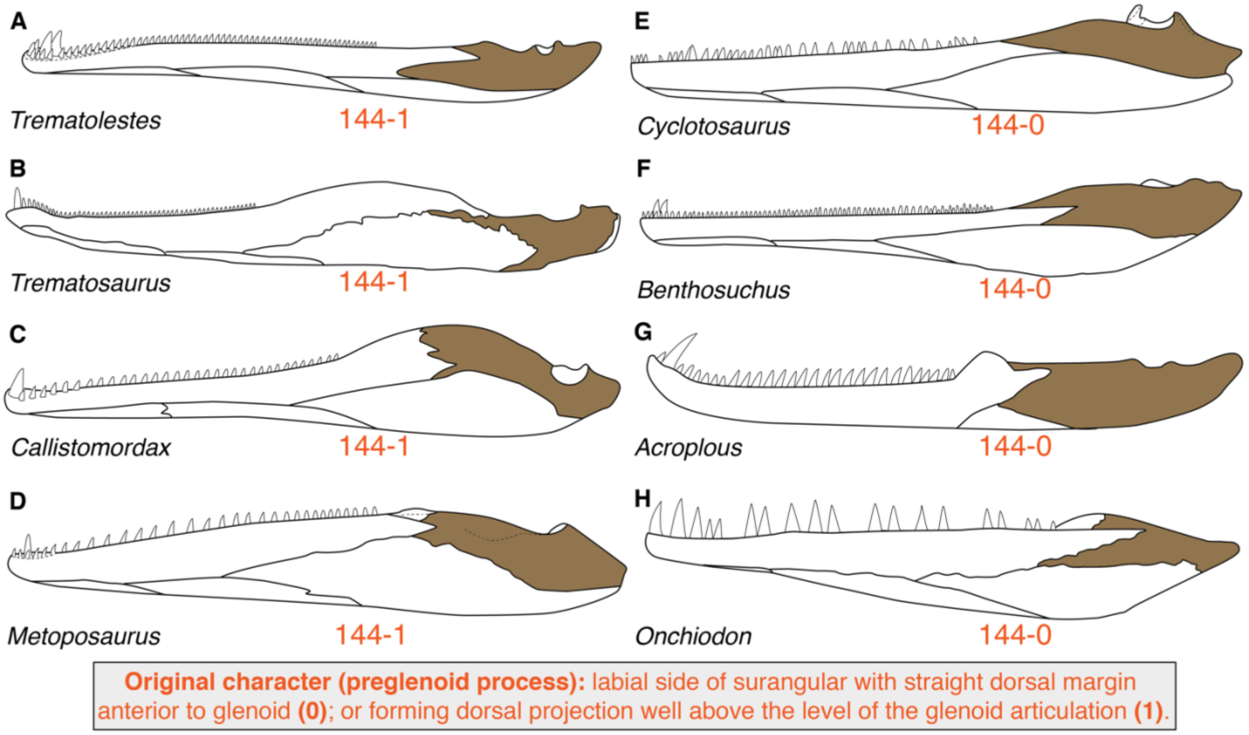

**Figure 27. Comparison of the dorsal margin of the surangular anterior to the glenoid in temnospondyls. A**, the trematosaur *Trematolestes hagdorni* (after Schoch, 2006); **B**, the trematosaur *Trematosaurus brauni* (after Schoch, 2019a); **C**, the metoposauroid *Callistomordax kugleri* (after Schoch, 2008a); **D**, the metoposaurid *Metoposaurus krasiejowensis* (after Sulej, 2007); **E**, the capitosaur *Cyclotosaurus intermedius* (after Sulej & Majer, 2005); **F**, the trematosaur *Benthosuchus sushkini* (after Damiani, 2001); **G**, the dvinosaur *Acroplous vorax* (after Englehorn et al., 2008); **H**, the eryopoid *Onchiodon labyrinthicus* (after Boy, 1990). Orange text represents the original character and codes from Schoch (2013). Silhouettes not to scale.

This is further complicated by the nature of the states, with 144–0 seemingly specifying whether the dorsal margin is straight (versus concave or convex) and 144–1 specifying a position relative to the glenoid. It is important to note that in either case, this character refers only to the surangular portion of the coronoid process.

In *Trematosaurus*, the surangular essentially appears flat dorsally and does not appear to contribute to the coronoid process (Schoch, 2019a:fig. 6E; Fig. 27B). This differs markedly from *Callistomordax* in which there is a very tall coronoid process to which the surangular has a major contribution and thus a dorsally convex margin (Schoch, 2008a:fig. 1E-F; Fig. 27C). *Trematolestes*, in which the mandible is the most poorly known, was reconstructed by Schoch (2006:fig. 4D-E; Fig. 27A) as having a short coronoid process entirely formed by the surangular. *Metoposaurus* has a short coronoid process primarily formed by the surangular, and it is very slightly dorsally convex (Sulej, 2007:fig. 3; Fig. 27D). Based on the character states, it is unclear what makes this collection of conditions distinct from all other taxa; dorsally convex surangulars are also found in taxa like *Benthosuchus* (e.g., Schoch & Milner, 2000:fig. 71; Fig. 27F), *Eocyclotosaurus* (e.g., Rinehart et al., 2015:fig. 6), and perhaps *Rileymillerus* (Bolt & Chatterjee, 2000:fig. 4.1), all coded for 144–0.

A character related to the relative height of the glenoid was added by Pardo et al. (2017a, character 269). Therefore, I treated character 144 as on referring to whether the dorsal margin of the surangular was essentially straight or whether it was convex.

  - Codes were recoded from 144–0 to 144–1 for *Greererpeton*, *Dvinosaurus*, *Benthosuchus*, *Paracyclotosaurus*, *Cyclotosaurus*, and *Eocyclotosaurus*.
  - *Thabanchuia* was uncoded from 144–0 because there is no express description of the surangular (due to articulation with the skull), and the reconstruction depicted a dorsal convexity (Warren, 1998b:fig. 6). *Rileymillerus* was uncoded from 144–0 because Bolt & Chatterjee (2000:677) stated that the dorsal aspect of the surangular was poorly visible; their figure 4.1 depicted a dorsal convexity (144–1).

- **146 (symphyseal teeth):** a third state was added to differentiate between a parasphymphyseal row set an angle relative to the dentary tooth row (often following the lingual margin of the symphysial region) and one set parallel to the dentary tooth row. This is a change made as part of the revisions by Kligman et al. (2021) and serves to further differentiate the gymnophionomorph condition from certain temnospondyls. Note that symphyseal tusks are not considered as part of this character.
  - *Ichthyophis*, *Epicrionops*, and *Eocaecilia* were changed from 146–1 to the new 146–2.
  - *Doleserpeton* was originally coded as lacking adsymphyseal teeth, but there is clearly a second, lingually situated row on the dentary (Sigurdsen & Bolt, 2010:1366, fig. 6A) that is parallel to the labial row. This taxon is corrected to the new 146–2. *Gerrothorax* was also originally coded as lacking such teeth but Schoch & Witzmann (2012:378) described a short row of 2–5 teeth posterior to the marginal row and on the symphysis that were a continuation of the coronoid series (this does not appear in their figure 5A). These teeth were also described by Hellrung (2003:39) and are similarly absent from the reconstruction (figure 15 therein).
  - *Siderops* was corrected from 146–0 to the revised 146–1 (Warren & Hutchinson, 1983:25, fig. 15); *Callistomordax* was corrected from 146–0 to partial uncertainty for 146–1/2 (Schoch, 2008a:91).
  - *Metoposaurus* was updated from 146–1 to 146–0&1 based on the taxonomic substitution of *M. krasiejowensis* (Konietzko-Meier & Wawro, 2007).
  - *Chinlestegophis* was recoded from 146–1 to partial uncertainty for states 1 and 2 (the original description does not make clear their alignment).

- *Edops* was newly coded for 146–0; there are only paired symphyseal tusks (Romer & Witter, 1942).

- **147 (posterior coronoid teeth), 148 (anterior, middle coronoid teeth); 272 (coronoid teeth):** the original coding for characters 147 and 148 is difficult to interpret.
  - **147. Posterior coronoid teeth.** Present (0), or absent (1). [*added by Schoch, 2013*]
  - **148. Anterior, middle coronoid teeth.** Present (0), or absent (1). [*added by Schoch, 2013*]

  *Proterogyrinus*, which has denticles on the posterior and anterior coronoids (middle coronoid is unknown; Holmes, 1984), was coded for state 0 for both characters. Conversely, *Greererpeton* was coded for state 1 for both characters even though it has denticles across the coronoids and a pair of 'fangs' on the anterior coronoid (the question of whether the middle and posterior coronoid suture has been obliterated is also an issue; Bolt & Lombard, 2001:1032). Most non-stereospondyls were coded the same as *Proterogyrinus* and have denticles but no other form of dentition on all three coronoids. Therefore, whether these two characters are restricted to 'true teeth' or are considered to encompass any form of dentition, previous codes cannot be reconciled. Additionally, the grouping of the anterior and middle coronoids into one character reflects the observation that the most common conditions in temnospondyls are either dentition on all three coronoids or only on the posterior coronoid, but there are exceptions among the taxa sample (e.g., *Siderops*, *Callistomordax*). Character 272 also refers to coronoid teeth but appears to have been coded only on 'true teeth' of a subequal size to the marginal dentition; the coding of character 273 (coronoid teeth, size) also reflects this.

  Within the context of Schoch (2013), classifying both characters as encompassing any form of dentition would require the fewest coding modifications. In the expanded character sample of Pardo et al. (2017a), these three characters could be collectively redefined to each refer to dentition of an individual coronoid, if not further for each combination of a coronoid and different forms of dentition (e.g., denticles on anterior coronoid) (e.g., Ruta & Bolt, 2006; McHugh, 2012; Marjanović & Laurin, 2019). However, adopting this approach becomes more complicated when considering the addition of taxa without three coronoids, as the homology becomes uncertain. In the context of these matrices, there is one temnospondyl (*Plagiosuchus*) that has fewer than three coronoids, while *Greererpeton* has two or three (see above comment; Smithson, 1982; Bolt & Lombard, 2001). The three gymnophionomorphs lack a distinct coronoid in adults, but a distinct dentulous ossification is present during development that then fuses with the dentary to form the pseudodentary and the double marginal tooth row. Based on neontological data, this has often been inferred to be specifically the anteriormost coronoid based on its formation near the symphysis and the distribution of teeth in early tetrapods (e.g., Müller et al., 2005; Theska et al., 2019). Pardo et al. (2017a), who introduced character 271 for coronoid count, did not code gymnophionomorphs for this character, but they did code coronoid teeth as present (272–1), presumably to capture their hypothesized transition from the tooth row on all three coronoids in *Chinlestegophis*.

  The issue of coronoid homology becomes more vexing when considering the lepospondyls added by Schoch et al. (2020); *Rhynchonkos* has two coronoids (Szostakiwskyj et al., 2015), *Brachydectes* has one (Pardo & Anderson, 2016), and

*Batropetes* has zero (Glienke, 2013, 2015). Without a clear means of inferring homology in taxa with fewer than three coronoids, modifying characters to refer to individual coronoids is problematic. One alternative is to not code any taxon with fewer than three coronoids for any character invoking a specific coronoid (as with Marjanović & Laurin, 2019). This is undoubtedly the most conservative means of *coding* but not necessarily of *recoding*. Such a conservative approach is unlikely to be favored by all workers either. Therefore, the solution implemented here is as follows.

- Character 272 was treated as referring only to 'true teeth,' exclusive of denticles and 'tusks.' It was then modified to differentiate taxa with only teeth on the posteriormost coronoid from taxa with teeth on all present coronoids; this removed the practical issue of coronoid homology, although it homologizes hypothetical taxa with the same condition but with different numbers of coronoids (e.g., *Plagiosuchus* with only one coronoid, which is edentulous, and *Metoposaurus* with three coronoids, all of which are edentulous). The creation of a state for teeth restricted to the posteriormost coronoid differentiates many capitosaurs and trematosaurs from other stereospondyls. This new state is added as the new state 1 because it could be conceivably treated as an ordered character.
    - The only taxa for which this redefinition creates problems is *Callistomordax*, with three coronoids but with teeth only on the middle coronoid; *Lydekkerina*, with three coronoids but with teeth only on the anterior coronoid; and *Siderops*, with three coronoids but with teeth only on the middle and posterior coronoid. These taxa were treated as inapplicable. Holmes (1984:35) identified at least one tooth of subequal size to the marginal dentition on the anterior coronoid of *Proterogyrinus*, but Ahlberg & Clack (1998:35) have suggested this was a dislodged or misidentified marginal tooth. This taxon is changed to unknown here; if Holmes' interpretation was valid, it would have to be treated as inapplicable.
    - Codes were recoded from 272–0 to 272–1 for *Benthosuchus* and *Mastodonsaurus*.
    - Codes were recoded from 271–0 to 272–2 for *Trimerorhachis, Doleserpeton, Ecolsonia, Onchiodon, Sclerocephalus, Archegosaurus, Platyoposaurus,* and *Chomatobatrachus*. The condition of *Ecolsonia* (a "shagreen of teeth"; Berman et al., 1985:17) is interpreted as a semantic difference of referring a denticle field.
    - Codes were recoded from  272–1 to 272–2 for *Edops, Adamanterpeton, Cochleosaurus, Balanerpeton, Acroplous, Iberospondylus, Australerpeton, Rhineceps, Broomistega,* and *Plagiosuchus*.
    - *Chenoprosopus, Dendrerpeton, Isodectes, Acanthostomatops, Zatrachys, Micromelerpeton, Platyrhinops, Amphibamus, Dissorophus, Cacops, Phonerpeton, Eryops, Glanochthon, Metoposaurus, Paracyclotosaurus,* and *Cyclotosaurus* were newly coded for 272–2.
    - *Ichthyophis, Epicrionops*, and *Eocaecilia* were changed from 272–0 to 272–0/1 to account for the single embryological coronoid.
    - *Isodectes* and *Rileymillerus* were changed from 272–0 to 272–0/1 because the dentition of only the posterior coronoid is confidently known.

        - *Lapillopsis* was changed from 272–0 to unknown because teeth were only confidently identified on the middle coronoid. Yates (1999) reasonably inferred that teeth were present on all three coronoids (subsequently verified in the other lapillopsids *Manubrantlia* and *Rotaurisaurus*) on the basis that no temnospondyl only has teeth on the middle coronoid, but this condition has subsequently been identified in *Callistomordax* (Schoch, 2008a), so the condition is best treated as unknown.
        - *Kokartus* was newly coded for 272–0/1, also related to the apparent retention of only one coronoid.
    - Characters 147 and 148 were then repurposed as characters for coronoid denticles and tusks, respectively. Most taxa with denticles have denticles on all three coronoids. However, some taxa lack denticles on the anterior coronoid (*Trimerorhachis*) or on both the anterior and middle coronoid (*Acroplous, Onchiodon*). These are differentiated from taxa with denticles on all three coronoids but are not further differentiated to maintain the parsimony-informative nature of the character. Coronoid tusks (new 148–1) only occur in two taxa, *Greererpeton* and *Thabanchuia*, and therefore a third state to differentiate them (restricted to anterior coronoid in the former) was not added.
        - Codes were changed from 147–1 to 147–2 for *Callistomordax, Metoposaurus, Ichthyophis, Epicrionops*, and *Eocaecilia*.
        - Codes were changed from 147–0 to 147–2 for *Australerpeton, Plagiosuchus, Gerrothorax, Edingerella, Benthosuchus, Wetlugasaurus, Eryosuchus, Cyclotosaurus, Eocyclotosaurus, Mastodonsaurus, Chinlestegophis, Rileymillerus*, and *Kokartus*.
        - *Lapillopsis, Paracyclotosaurus*, and *Rileymillerus* were newly coded for 147–2.
        - *Edops, Adamanterpeton, Dendrerpeton*, and *Zatrachys* were newly coded for 147–0.
        - *Neldasaurus* was newly coded for 147–0/1 (Chase, 1965).
        - *Acanthostomatops, Glanochthon*, and *Platyoposaurus* were changed from 147–0 to 147–0/1 (Boy, 1988, 1993; Schoch & Milner, 2000).
        - Codes were changed from 147–1 to 147–0 for *Greererpeton*.
        - Codes were changed from 147–0 to 147–1 for *Trimerorhachis* and *Acroplous*.
        - *Doleserpeton* was changed from 147–1 to inapplicable because the presence of an anterior coronoid is uncertain, the probable middle coronoid (the anteriormost preserved) has denticles, and the probable posterior coronoid (the posteriormost preserved) has no dentition.
        - *Cochleosaurus* was uncoded from 147–0 because the anteriormost coronoid is unknown (Sequeira, 2004:30). *Apateon* and *Micropholis* were uncoded from 147–1 and 148–1 on the basis that a confident determination of absence cannot be made given disarticulation and obscuring of the lower jaw elements and/or articulation with the skull. *Batrachosuchus* was uncoded from 147–0 because no lower jaw is known for this taxon. *Wetlugasaurus* and *Eryosuchus* were uncoded for characters because there is no recent literature describing or figuring the dentition of

the lower jaw. Both have been coded as having coronoid teeth (e.g., Damiani, 2001, character 45–1 therein, which was adopted and maintained for these two by Schoch, 2008b). However, as one counterpoint, Schoch & Milner (2000:fig. 90) depicted several capitosauroid mandibles in lingual view and did not show any dentition for *Eryosuchus*, but they did for *Benthosuchus* and *Mastodonsaurus* (indicating this was not an illustrative preference). They can be coded for 147–2 because character 45 of Damiani and Schoch differentiates teeth from denticles as the two states. *Trematosaurus* was uncoded from 147–0 and 148–0 because teeth are not commonly preserved in the material, and the coronoids are so badly preserved that their sutures cannot be discerned (Schoch, 2019a:51, 53). *Parotosuchus* was uncoded from 147–1 and 148–1 because Schoch (2018:119) only mentions a well-preserved anterior coronoid.

- Codes were changed from 148–1 to 148–0 for *Trimerorhachis*, *Acroplous*, *Lydekkerina, Plagiosuchus, Edingerella*, *Metoposaurus, Eocyclotosaurus, Eryosuchus, Cyclotosaurus,* and *Mastodonsaurus.*
- Codes were changed from 148–0 to 148–1 for *Thabanchuia*.
- *Edops*, *Adamanterpeton*, *Dendrerpeton, Acanthostomatops*, *Zatrachys,*
- *Acheloma*, *Lapillopsis, Broomistega*, *Paracyclotosaurus*, and *Kokartus* were newly coded for 148–0.
- *Ambystoma* and *Hynobius* were recoded from 148–1 to inapplicable because they have no coronoids (271–3).

- For the reanalysis of Schoch (2013), the original character 147 and 148 were restored (as character 272 is not sampled) and were revised to differentiate among the distribution of teeth and denticles (both absent, teeth only, denticles only, both present). The presence of coronoid tusks/fangs is not accounted for here given the rare occurrence. Character 148 was restricted to only the middle coronoid. The rationale here is that it was likely lumped with the anterior coronoid because there are some stereospondyls (mostly capitosaurs) in which dentition is restricted to the posteriormost coronoid (e.g., *Benthosuchus*, *Mastodonsaurus*). In other words, it was assumed that the other two coronoids would always share a condition. This is not the case among sampled taxa; *Trimerorhachis*, *Siderops*, and *Callistomordax* have dentulous middle coronoids but edentulous anterior coronoids. There is no sampled taxon with three coronoids that definitively only has dentition on the anterior coronoid.
  - *Edops*, *Adamanterpeton*, *Dendrerpeton*, *Neldasaurus*, *Zatrachys*, *Acheloma*, and *Broomistega* were newly coded as 147–0.
  - Codes were recoded from 147–1 to 147–3 for *Micropholis, Doleserpeton, Callistomordax,* and *Metoposaurus.*
  - Codes were recoded from 147–1 to 147–2 for *Greererpeton*.
  - Codes were recoded from 147–0 to 147–3 for *Australerpeton, Plagiosuchus*, and *Cyclotosaurus*.
  - *Paracyclotosaurus* was newly coded for 147–3.
  - Codes were recoded from 147–0 to 147–1 for *Gerrothorax, Edingerella, Benthosuchus, Wetlugasaurus, Eryosuchus, Eocyclotosaurus,* and *Mastodonsaurus*.

- Codes were recoded from 147–0 to 147–1/2 for *Siderops*. The partial uncertainty reflects the tenuous nature of the encapsulating statement by Warren & Hutchinson (1983:25); it can be inferred that if replacement teeth were only sometimes preserved, that the patch of denticles on the middle coronoid might not represent the full extent of a denticle field.
- Codes were recoded from 148–1 to 148–2 for *Greererpeton* and *Siderops*.
- Codes were recoded from 148–1 to 148–0 for *Trimerorhachis,* and *Lydekkerina*.
- *Edops*, *Adamanterpeton, Dendrerpeton*, *Acanthostomatops*, *Zatrachys*, and *Broomistega* were newly coded as 148–0.
- *Isodectes* and *Acanthostomatops* were uncoded from 148–0 because there is no clear description of the middle coronoid dentition (Boy, 1988; Sequeira, 1998). *Micropholis* was uncoded from 148–1 because the anterior and middle coronoids have not been positively identified (Schoch & Rubidge, 2005). *Platyoposaurus* was uncoded because the condition of the middle coronoid is unclear based on the literature available to me; Schoch & Milner (2000:fig. 39) only depicted denticles on the posterior coronoid of *Platyoposaurus*, which would be discordant with other early diverging stereospondylomorphs.
- *Wetlugasaurus* and *Eryosuchus* were recoded to 148–1/3 because the literature available to me did not clearly describe the middle coronoid dentition. Schoch & Milner (2000:fig. 90) depicted *Eryosuchus* as edentulous, but this taxon has been repeatedly coded as having coronoid teeth (e.g., Damiani, 2001, character 45; and derivates thereof). While most capitosaurs only have dentition on the posterior coronoid, some (e.g., *Edingerella, Watsonisuchus*; Warren & Hutchison, 1988; Steyer, 2003) have a more extensive tooth row. This uncertainty is reflected in their differing original codes; *Wetlugasaurus* was coded as having anterior and middle coronoid dentition, whereas *Eryosuchus* was coded as lacking it.
- *Lapillopsis* was newly coded for 148–1.
- Codes were recoded from 148–0 to 148–3 for *Australerpeton* and *Benthosuchus*.
- Codes were recoded from 148–0 to 148–1 for *Gerrothorax* and *Callistomordax*.
- Codes were recoded from 148–1 to 148–3 for *Metoposaurus.*
- Codes were recoded from 148–1 to 148–1/3 for *Eocyclotosaurus.* This follows my reinterpretation of an elongate ‘splenial’ with teeth just anterior to the adductor fossa in *E. appetolatus*; if this element is a coronoid, it probably represents the plesiomorphic three coronoids without discernible sutures and thus it cannot be determined whether the teeth are restricted to the posteriormost coronoid.
- The following taxa were newly coded as 148–1: *Paracyclotosaurus.*
- *Plagiosuchus* was recoded from 148–1 to inapplicable because there are no anterior or middle coronoids in this taxon.
- *Tupilakosaurus* was uncoded for both characters (state 0) because the coronoids are unknown (e.g., Warren, 1998b:153). *Apateon* and

    - *Branchiosaurus* were uncoded for both characters (state 1) because the disarticulation of specimens obscures the coronoids in most specimens, and definitively proving that dentition was absent. *Batrachosuchus* was uncoded for both characters (147–0, 148–0) because no lower jaw is known for either *Batrachosuchus browni* or *Bathignathus watsoni* (Watson, 1956; Welles & Estes, 1969; Warren & Marsicano, 2000; Schoch & Milner, 2014). *Parotosuchus* and *Trematosaurus* were uncoded for both characters (147–0, 148–0) because the medial surface is very poorly known such that sutures to confidently identify all three coronoids are not even known, and there is no description of any dentition on the coronoids (Schoch, 2018c:119, fig. 7G; 2019:51, 53, fig. 6G-H).

- **272 (coronoid teeth); 273 (coronoid teeth, size):** any taxon coded for 272–1 (as clarified above) cannot be coded for character 273.
    - *Doleserpeton, Ecolsonia, Onchiodon, Sclerocephalus*, *Archegosaurus*, *Platyoposaurus*, and *Chomatobatrachus* were originally coded as 272–0 but lack teeth of a subequal size to the marginal dentition on the coronoids; these were recoded to 272–1 and thus changed to inapplicable for character 273.
    - *Chenoprosopus*, *Dendrerpeton, Isodectes, Acanthostomatops*, *Zatrachys, Micromelerpeton, Platyrhinops, Amphibamus, Dissorophus, Cacops, Eryops, Glanochthon, Metoposaurus*, *Paracyclotosaurus,* and *Cyclotosaurus* were originally uncoded and were newly coded for 272–2 here. As a result, they were then updated from unknown to inapplicable for character 273.
- **149 (mandibular osteoderms):** state 1 refers to “a mosaic of ventral osteoderms between mandible and dermal pectoral girdle.” On this basis, taxa without both a known mandible and pectoral girdle in articulation should not be coded for this taxon because the taphonomic loss of one or both could also have resulted in the loss of such osteoderms.
    - Codes were corrected from 149–0 for *Cochleosaurus*, *Nigerpeton*, *Sangaia*, and *Batrachosuchus*.
- **150 (stapes, quadrate process):** this appears to be another character in which codes were reversed for most taxa; *Sclerocephalus* and *Archegosaurus*, which have a well-figured stapes with a quadrate process (Witzmann, 2005:fig. 30; Schoch & Witzmann, 2009a:fig. 6F-G) were coded as lacking this process (150–0), whereas taxa like *Edops* and the majority of stereospondyls that actually lack this process (Schoch, 2017) were coded as having one (150–1). However, some taxa were originally coded correctly (e.g., *Glanochthon* lacks a process and was coded for 150–0 by Schoch, 2013; Schoch & Witzmann, 2009b).
    - Codes were reversed (corrected) from 150–1 to 150–0 for *Edops*, *Dendrerpeton, Balanerpeton, Trimerorhachis, Acanthostomatops, Micromelerpeton, Limnogyrinus, Branchierpeton, Branchiosaurus, Apateon, Micropholis, Platyrhinops, Amphibamus, Doleserpeton, Dissorophus, Cacops, Broiliellus, Acheloma, Phonerpeton, Onchiodon, Eryops, Rhineceps, Uranocentrodon, Lydekkerina, Siderops, Edingerella, Watsonisuchus, Benthosuchus, Trematosaurus, Trematolestes, Lyrocephaliscus, Metoposaurus, Parotosuchus, Paracyclotosaurus, Eocyclotosaurus, Mastodonsaurus*, *Xenopus*, *Ichthyophis, Epicrionops*, and *Eocaecilia*.

    - Codes were reversed (corrected) from 150–0 to 150–1 for *Dvinosaurus, Sclerocephalus*, and *Archegosaurus.*
    - *Proterogyrinus* was uncoded as there is no evidence for the stapes (Holmes, 1984; Clack et al., 2016); *Batrachosuchus* was uncoded as there is no mention of the stapes or other parts of the otic capsule (Chernin, 1977).
    - Codes were uncoded from 150–1 for *Isodectes, Trematolestes, Wetlugasaurus,* and *Eryosuchus* because the literature available to me did not clearly describe or figure the stapes' morphology.
    - *Lapillopsis* was newly coded for 150–0.
- **152 (stapes, shape):** mention of the presence of a quadrate process for state 0 was removed, as most taxa coded for state 0 do not have this process (150–0). This did not require any coding modifications.
- **153 (stapes, curvature):** as indicated by the name of this character, the feature being highlighted is curvature of the stapes. State 1, however, prescribes at least three different attributes: curvature (no curvature), length ("abbreviated"), and the presence of a vertically aligned and laterally situated otic notch.
    - **153. Stapes (curvature).** Stapes with pronounced dorsodistal curvature (0), or abbreviated without such curvature, directed laterally towards vertically aligned otic notch (1). [*added by Schoch, 2013*]

  This is problematic because state 0 prescribed only curvature, and there are long, non-curved stapes (e.g., *Mastodonsaurus*) and long, curved stapes not directed towards a vertically aligned otic notch (e.g., *Trimerorhachis*). There is also the broader issue of coding taxa without an otic notch. The character is therefore restricted to merely curvature.
    - Codes were recoded from 153–0 to 153–1 for *Edops, Dvinosaurus, Acroplous, Iberospondylus, Acanthostomatops, Lydekkerina, Siderops, Edingerella, Watsonisuchus, Benthosuchus, Metoposaurus, Parotosuchus, Eocyclotosaurus,* and *Mastodonsaurus.*
    - Codes were recoded from 153–0 to unknown for *Proterogyrinus* and *Batrachosuchus* (see comments for character 150) and for *Trematosaurus* (Schoch, 2019a:53). Codes were recoded from 153–0 to unknown for *Isodectes, Trematolestes, Wetlugasaurus,* and *Eryosuchus* because the literature available to me did not clearly describe or figure the stapes' morphology. To avoid introduce spurious codings, I did not merely invert them. *Cyclotosaurus* was uncoded as a result of the taxonomic substitution of *C. intermedius*.
    - Codes were recoded from 153–1 to 153–0 for *Apateon*.
    - *Zatrachys* and *Ecolsonia* were newly coded for 153–0.
    - *Ambystoma, Hynobius, Cryptobranchus, Ichthyophis, Epicrionops,* and *Eocaecilia* were newly coded for 153–1.
- **158 (presacral count):** the three original character states did not encompass the full range of possible presacral counts; any taxon with 22, 26, 27, or 28 presacral vertebrae was not captured here.
    - **158 (presacral count).** More than 28 (0), or 23-25 vertebrae (1), or less than 21 (2). [*added by Schoch, 2013*]

  The addition of a state 3 for less than 19 positions (inferred, state not actually specified) by Schoch et al. (2020) does not resolve this. Considering that these states are arbitrarily differentiated, there is no compelling reason to merely leave taxa with these non-captured counts as uncoded (e.g., *Trematolestes* with 22 presacral vertebrae). I therefore modified state 1 to be '23–28' and state 2 to be '19–22.' Schoch et al.'s addition of state 3 is maintained.

- *Acroplous* was uncoded from 158–0 because all specimens with vertebrae consist only of isolated or fragmentary elements; there is not even a sufficiently complete partial column that has permitted proper estimation of presacral count previously (Hotton, 1959; Coldiron, 1978).
- *Thabanchuia* was uncoded from 158–0 because the most positions preserved in any specimen is seven (Warren, 1998b).
- *Acanthostomatops* was recoded from 158–1 to 158–2. Witzmann & Schoch (2006:374) stated that "has a rather short trunk composed of 21–22 presacral vertebrae." This range would not fit within a single character state in the original character (and is thus an erroneous code), but it fits within state 2 of the reformulated version.
- *Branchierpeton* was recoded from 158–0 to 158–1. Werneburg (1991) stated that the count varied between 25 and 28 positions, a range restricted to the revised state 1 and that should have been treated as partial uncertainty in the original formulation.
- *Platyrhinops* is largely known from incomplete specimens, but the most complete one has 25–26 presacral positions (Carroll, 1964:234; Clack & Milner, 2009:289). Therefore, this taxon was recoded to 158–1, rather than 158–2.
- *Iberospondylus* has 22 presacral positions (including the atlas and axis; Laurin & Soler-Gijón, 2006:290) and was recoded from 158–1 to 158–2. As it has 23 caudal positions, the original code may have been a misreading.
- *Australerpeton* was recoded from 158–1 to unknown. Dias & Schultz (2003:32) reported 18 positions in a sub-adult individual and 20 positions in an adult individual, but it is apparent from their figures (figs. 2–5 therein) that these are not complete vertebral columns, and the counts (which do not differentiate presacral from total preserved count) merely indicate the preservation. Eltink & Langer (2014:fig. 11) reconstructed 26 presacral positions, which was not applicable to the original state 1, but because a similar presacral count character was left uncoded for this taxon by Eltink et al. (2019, character 167 therein), it is uncoded here.
- *Siderops* has 23 presacral positions (Warren & Hutchinson, 1983:26), which means it was recoded to 158–1, rather than 158–2.
- *Trematolestes* was recoded from 158–1 to 158–2. Regardless of the reformulation, it was originally an erroneous code; Schoch (2006a:36) stated that approximately 22 presacral vertebrae are present exclusive of the atlas and axis (24 total presacral).
- *Callistomordax* was recoded from 158–0 to 158–1. Schoch (2008a:92) stated that the presacral count is 26–28, which is not applicable to the original or the new (unchanged) state 0.
- *Paracyclotosaurus* was recoded from 158–0 to 158–1. Watson (1958:246) identified position 28 as the sacral position, which means there are between 25

    - and 27 presacral positions depending on whether the atlas and axis are counted; either is not applicable to state 0.
    - *Mastodonsaurus* has long had an uncertain presacral count; historical workers' estimates ranged from 27–30, which does not fit within a single character state in either the original or revised formulation of this character. Schoch (1999a:93) stated that at least 27 were present and suggested a range of 27–29 (which still does not fall within one character state); he described two specimens, one with 28 positions and one with 30 positions. It was recoded from 158–0 to 158–0/1, which represents a correction.
- **171 (ribs, length); 296 (ribs anterior to sacrum):** both of these characters relate to rib length, although it is formalized somewhat differently between them (original definitions shown below).
    - **171. Ribs (length).** Moderately elongate thoracic ribs curved distoventrally (0), or such ribs foreshortened without distal curvature (1). [*added by Schoch, 2013*]
    - **296. Ribs anterior to sacrum.** Short (0), long (1). [*added by Pardo et al., 2017a*]

  Character 171 is named as if it addresses length, but it referenced both length and curvature, assuming that they are always correlated, which creates a problematic compound character, independent of character 296. Character 296 only addresses length. Codes for these characters are not the same, either before or after other coding changes that were made to both; this reflects the inherent subjectivity in differentiating 'long' versus 'short' and that these characters were coded by different workers. Other versions of this character (e.g., character 88 in Schoch, 2018b) have invoked a reference point: the length of three vertebral positions. However, presacral rib length is typically variable, being broadest in the anterior half of the trunk in most taxa and tapering sharply towards the sacrum. This is only further complicated by the fact that while most taxa are coded, complete presacral columns or even partial columns that can be precisely positioned are not known for most.

  Character 171 is here restricted to curvature, and character 296 is retained for relative length. Both characters were recoded for all taxa (see further comments below on a pattern of discrepancies within character 296).
    - Codes for character 296 seem to have been inverted; *Gerobatrachus*, for example, has relatively short ribs like in lissamphibians (Anderson et al., 2008) but was coded for 'long' ribs (296–1), whereas lissamphibians were coded for 'short' ribs (296–0). Most temnospondyls have much longer ribs than in lissamphibians, but they were coded for 'short' ribs. In order to maintain consistency, the character was recoded for all taxa.
- **173 (ribs, uncinate blades); 174 (ribs, uncinate spines):** the differing names between these characters creates confusion because it is unclear if character 174 should only be coded if a taxon has uncinate spines (173–0).
    - **173. Ribs (uncinate blades).** If present, small and spine-like (0), or extensive and blade-like (1). [*added by Schoch, 2013*]
    - **174. Ribs (uncinate spines).** Short (0), or elongated, as long as shaft (0). [*added by Schoch, 2013*]

  Most taxa were coded for both 173–0 and 174–0, but eryopoids and non-stereospondyl stereospondylomorphs, *Iberospondylus, Australerpeton*, *Peltobatrachus, Paracyclotosaurus, Eryosuchus,* and *Mastodonsaurus* were coded for 173–1 and 174–0 while *Trematolestes* and *Callistomordax* were coded for 173–0 and 174–1. The combination of 173–1 and 174–0 is the one in conflict because of how character 174 is named; if a taxon has 'extensive and blade-like' processes, presumably it should not be coded for 'short' spines. To be clear, some taxa do have multiple processes or expansions termed 'uncinate processes' in some of the ribs (e.g., *Eryops*; Moulton, 1974:fig. 16; *Mastodonsaurus*; Schoch, 1999:fig. 34), but this is relatively uncommon, so it is interpreted that these do not refer to different processes (e.g., *Sclerocephalus* and *Australerpeton*, coded for 173–1 and 174–0, have only one process; Dias & Schultz, 2003:fig. 10; Schoch & Witzmann, 2009a:fig. 7). Some taxa also lack uncinate processes, but were coded for state 0 for both characters (e.g., *Limnogyrinus*; e.g., Werneburg, 2012:fig. 24), which likely reflects the apparent tendency of Schoch (2013) to code inapplicable cells for the inferred plesiomorphic state.

A third state is added to character 173 for 'uncinate processes absent.' This is appended as the new state 2 to create an unordered multistate character. Character 174 is also modified to specify that the longest uncinate process is longer than the respective rib for state 1, as there is typically variation along the trunk in taxa originally coded for 174–1.

- Codes were recoded from 173–0 to 173–2 and from 174–0 to inapplicable for *Proterogyrinus* (Holmes, 1984:figs. 1, 23), *Cochleosaurus* (Sequeira, 2009:143, fig. 5), *Dendrerpeton* (Holmes et al., 1998:fig. 1), *Balanerpeton* (Milner & Sequeira, 1993:341), *Laidleria* (Kitching, 1958:fig. 17), *Trimerorhachis* (Pawley, 2007:876, fig. 4), *Neldasaurus* (Chase, 1965:fig. 10), *Dvinosaurus* (e.g., Bystrow, 1938:fig. 26), *Acroplous* (Coldiron, 1978:fig. 1), *Isodectes* (e.g., Milner, 1982:fig. 1), *Thabanchuia* (Warren, 1998b:fig. 7), *Acanthostomatops* (Witzmann & Schoch, 2006:376, figs. 6–9), *Micromelerpeton* (e.g., Fröbisch et al., 2014:fig. 1), *Limnogyrinus* (e.g., Werneburg, 2012:fig. 24), *Branchierpeton* (e.g., Credner, 1881:fig. 2), *Branchiosaurus* (e.g., Werneburg, 2012:fig. 4), *Apateon* (e.g., Sanchez et al., 2010:appendix A), *Platyrhinops* (Carroll, 1964:fig. 23; Clack & Milner, 2010:fig. 1), *Amphibamus* (e.g., Gregory, 1950:fig. 4), *Doleserpeton* (Sigurdsen & Bolt, 2010:figs. 1, 7), *Gerobatrachus* (Anderson et al., 2008:fig. 1), *Acheloma* (Williston, 1909:fig. 6), *Gerrothorax* (Nilsson, 1946:fig. 4),
- The name of character 174 suggests that it should only apply to taxa with 'spine-like' uncinate processes. Therefore, the following taxa coded as 173–1 were changed from 174–0 to inapplicable: *Iberospondylus*, *Onchiodon*, *Eryops*, *Sclerocephalus*, *Glanochthon*, *Archegosaurus*, *Australerpeton*, *Platyoposaurus*, *Peltobatrachus*, *Paracyclotosaurus*, and *Mastodonsaurus*. *Sclerocephalus* should either be recoded to 173–0 or as polymorphic; the anterior ribs' uncinate processes are narrow and spine-like, and the more posterior ones are not necessarily 'blade-like,' only less offset from the rib shaft (e.g., Schoch & Witzmann, 2009:fig. 7).
- *Greererpeton* was originally coded for 173–0 and 174–0, but Godfrey (1989b:13, 15, fig. 9) described uncinate flanges that are for all intents and purposes 'extensive and blade-like.' It was changed to 173–1 and to inapplicable for character 174.
- *Edops* was originally coded as unknown for character 173 and as 174–0, but Romer & Witter (1942:953) stated that only rib heads were known, and no new

postcrania of this taxon has since been described. Character 174 was recoded to unknown. The same was applied to *Edingerella*; Lehman (1961:32) was unsure of their presence, and while the holotype figured by Steyer (2003:fig. 1A) appears to lack them, one anterior rib may have an uncinate process. *Benthosuchus* and *Eocyclotosaurus* were originally coded for 173–? and 174–0; they were changed to 173–0 (e.g., Bystrow & Efremov, 1940:fig. 40; Rinehart et al., 2016:fig. 44).

    - *Broiliellus* was recoded to unknown from state 0 for both characters; the ribs are unknown among the known material (e.g., DeMar, 1968:fig. 4). *Plagiosuchus* was changed to unknown from state 0 for both characters because the postcrania have not been described in any fashion since von Huene (1922), whose figured specimen does not depict ribs.
    - *Siderops* was recoded from 173–0 to 173–0&2; Warren & Hutchinson (1983:31, fig. 19) only indicated uncinate processes on a few anterior ribs, and Warren & Snell (1991:51) stated that abdominal ribs in this taxon lack uncinate processes.
    - All lissamphibians added by Pardo et al. (2017a) were not coded for either character, presumably because most lissamphibians lack uncinate processes. All taxa with preserved ribs were updated to 173–2 and to inapplicable for character 174.
- **179 (interclavicle, length); 180 (interclavicle, ontogeny); 181 (interclavicle, central ornamented area); 182 (interclavicle, proportions); 183 (interclavicle, posterior margin); 184 (interclavicle, anterior margin); 185 (interclavicle, anterior stylus); 186 (interclavicle, posterior stylus); 187 (interclavicle, clavicles):** any taxon coded for 347–1 (interclavicle absent) in the matrix of Schoch et al. (2020) cannot be coded for these characters. Note that this still applies even to the matrices reanalyzed here in which character 347 was not sampled.
    - Character 179: Codes were corrected from 179–0 (interclavicle shorter than posterior skull) for *Triadobatrachus* and *Leptodactylus*.
    - Codes for these nine characters were updated from unknown to inapplicable (when originally coded as unknown) for the above taxa as well as for *Xenopus, Karaurus, Kokartus, Ambystoma, Hynobius, Cryptobranchus, Ichthyophis, Epicrionops,* and *Eocaecilia.*
- **183 (interclavicle, posterior margin); 186 (interclavicle, posterior stylus):** this character cannot be coded if a stylus is absent due to a transversely straight posterior margin (183–1). This holds true even if 186–0 (posterior end rounded or blunt) is intended to be the same as 183–1 (posterior margin transversely straight / without posterior process).
    - Codes were corrected from 186–0 (posterior end rounded or blunt) for *Plagiosuchus* and *Gerrothorax*.
- **197 (manual digit count):** the taphonomic bar for complete preservation of the manus and confident determination of only four digits (since a pentadactyl manus is known from at least two temnospondyls, contrary to the historical paradigm of a tetradactyl manus; Watson, 1958; Konietzko-Meier et al., 2020) should be very high. Despite this, many temnospondyls, including nearly all non-stereospondyls, were coded for 197–1 (tetradactyl manus); this includes several taxa for which the presence/absence of carpals was treated as unknown (character 196). The paucity of even partially complete manuses

is indicated by Dilkes (2015), both in his qualitative comparisons and in his coding of new phylogenetic characters related to the manus.

- Codes were corrected from 197–1 on the basis that the literature establishes that there are no known manual elements or that the manus is too incomplete to estimate digit count: *Edops*, *Thabanchuia*, *Cacops* (this holds regardless of the construction of this OTU), *Broiliellus, Siderops, Australerpeton,* and *Sangaia*. Many other taxa have a more tentative tetradactyl interpretation that may have been biased by the historical paradigm (e.g., *Cochleosaurus*; Sequeira, 2009:147), but these codes were not changed here.

- **203 (pubis):** state 0 was prescribed for an unossified pubis, with state 1 prescribedfor an ossified pubis, but the codes were largely reversed for taxa originally coded by Schoch (2013): taxa described as having an ossified pubis (e.g., *Cacops, Eryops*) were coded for 203–0, and taxa described as lacking an ossified pubis (e.g., *Acanthostomatops*, *Glanochthon*) were coded for 203–1. However, a few taxa coded were coded correctly (e.g., *Mastodonsaurus*), so all codes were vetted.
    - Codes were reversed (corrected) from 203–0 to 203–1 for *Proterogyrinus, Greererpeton, Cochleosaurus, Dendrerpeton, Balanerpeton, Micropholis, Amphibamus, Doleserpeton, Gerobatrachus, Dissorophus, Cacops, Broiliellus, Acheloma, Phonerpeton, Onchiodon, Eryops, Peltobatrachus, Uranocentrodon, Broomistega, Lydekkerina,* and *Gerrothorax*.
    - Codes were reversed (corrected) from 203–1 to 203–0 for *Trimerorhachis, Dvinosaurus, Acroplous, Isodectes, Acanthostomatops, Micromelerpeton, Limnogyrinus, Branchierpeton, Branchiosaurus, Apateon, Glanochthon, Archegosaurus, Platyoposaurus, Siderops, Plagiosuchus, Benthosuchus, Trematolestes, Callistomordax, Metoposaurus, Eocyclotosaurus,* and *Triadobatrachus*.
    - *Sclerocephalus* was corrected from 203–0 to 203–0&1 (Schoch & Witzmann, 2009a:157).
    - *Neldasaurus, Ichthyophis,* and *Epicrionops* were newly coded for 203–0.
    - *Platyrhinops* and *Australerpeton* were newly coded for 203–1.
- **207 (gastral squamation); 208 (squamation):** character 208 requires the presence of dermal scales, and taxa without these ossifications cannot be coded. Because these ossifications could hypothetically be lost from even complete skeletons post-mortem, the codes are changed to unknown rather than to inapplicable when they were changed. It is also somewhat unclear whether it is supposed to only refer to gastral squamation (i.e. if such squamation is present, what form does it take) or whether it is a broader character referring to squamation on any part of the body.
    - **207. Gastral squamation.** Ossified ventral dermal scales 0), or scales absent (1). [*added by Schoch, 2013*]
    - **208. Squamation.** Dermal scales oval to spindle-shaped (0) or broad-oval with “microsaur-type” ornament (1). [*added by Schoch, 2013*]

Twenty-eight taxa that were left as unknown for character 207 were coded for 208–0: *Adamanterpeton, Nigerpeton, Chenoprosopus, Neldasaurus, Acroplous, Tupilakosaurus, Iberospondylus, Zatrachys, Doleserpeton, Gerobatrachus, Dissorophus, Cacops, Broiliellus, Acheloma, Phonerpeton, Lapillopsis, Peltobatrachus, Rhineceps, Chomatobatrachus, Batrachosuchus, Watsonisuchus, Benthosuchus, Trematosaurus,*

*Wetlugasaurus, Eryosuchus, Eocyclotosaurus, Cyclotosaurus,* and *Mastodonsaurus*. Several of these taxa have no known postcrania (*Zatrachys, Rhineceps, Batrachosuchus*); this in particular is suggestive of block coding for character 208. The remaining taxa are a mixed bag. A few (e.g., *Neldasaurus*, *Acroplous, Doleserpeton*) are described with scales without a clear position, but most others have never been described with scales of any form or position.

Three taxa were coded for 207–1 (gastral squamation absent) but for 208–0 (dermal scales oval to spindle-shaped): *Acanthostomatops, Platyrhinops,* and *Edingerella*. All three appear to be in error. *Acanthostomatops* definitively has scales, and both ventral and dorsal scales have been noted (Witzmann & Schoch, 2006:381). Hook & Baird (1984) described outlines of gastralia in a large specimen of *Platyrhinops* that were compared favorably to scales of *Amphibamus* (see also Witzmann, 2007). Finally, there are gastral scales in *Edingerella* (see references in Witzmann, 2007:826), although they have not been noted or described by recent descriptions of this taxon (Steyer, 2003; Maganuco et al., 2009). All three of these codes were corrected to 207–0.

- In order to be consistent (and conservative), taxa were only coded for 207–1 when they were known from at least one articulated skeleton.
  - *Gerobatrachus, Dissorophus, Cacops*, and *Broiliellus* were newly coded for 207–1. *Metoposaurus* was uncoded from 207–1 as part of the taxonomic substitution – all material of *M. krasiejowensis* is disarticulated (Sulej, 2007). *Iberospondylus* was recoded from 207–0 to 207–1 because there is no description or figures of scalation (Laurin & Soler-Gijón, 2006). *Siderops* was recoded from 207–0 to 207–1 on the same basis (Warren & Hutchinson, 1983; Witzmann, 2007). *Peltobatrachus* was newly coded for 207–0 (Panchen, 1959; Witzmann, 2007). *Eocaecilia* was newly coded for 207–1 on the basis that postcrania of this taxon is abundant, some partially articulated, and that it was coded as lacking osteoderms (209–0). If the absence of osteoderms could be coded, then the absence of scalation should also be coded.
- Based on the naming practices of characters in which characters referring to the exact same feature share a name, character 208 would appear to refer to any form of squamation, not just gastral squamation. This would allow for coding of any taxa with scales of uncertain position or only on the dorsal surface.
  - *Nigerpeton, Tupilakosaurus, Iberospondylus, Gerobatrachus, Dissorophus, Cacops, Broiliellus, Acheloma, Phonerpeton, Lapillopsis, Chomatobatrachus, Siderops, Benthosuchus, Trematosaurus, Trematolestes, Callistomordax, Metoposaurus, Wetlugasaurus, Parotosuchus, Eryosuchus, Eocyclotosaurus,* and *Mastodonsaurus* were uncoded because there is no mention of scalation for these taxa. *Thabanchuia* was uncoded because the ossifications associated with the lower jaws were confined to this position and do not appear like other scales according to Warren (1998a:153); no other scalation was identified in this taxon. The same principle led to the uncoding of *Chenoprosopus milleri* based on Hook's (1993) description of '*Chenoprosopus lewisi*.' *Cyclotosaurus* was uncoded as part of the taxonomic substitution of *Cy. intermedius*. *Adamanterpeton, Zatrachys, Rhineceps, Sangaia,*

*Batrachosuchus*, and *Watsonisuchus* were uncoded on the basis that none of the postcrania is known.

- **210 (osteoderms, articulation):** the original 210–1 specified a double series of osteoderms with the additional caveat that the internal series was fused to the neural spines.
  - **210. Osteoderms (articulation).** Simple set of osteoderms arranged in one layer if present (0), or double set of two layers, the ventral one fused to the tip of the neural arch (1). [*added by Schoch, 2013*]

  A double series is common to both cacopine and dissorophine dissorophids, but only in the former does the internal (ventral) series of osteoderms fuse to the spines (e.g., DeMar, 1966; Schoch, 2012). However, the dissorophines *Dissorophus* and *Broiliellus* were also coded for 210–1, indicating that this additional caveat should be disregarded. Although there could be additional value in differentiating dissorophids by osteoderm-spine articulation (as in dissorophid-centric matrices), because only one cacopine is sampled, splitting of 210–1 into ‘double series, internal series not fused to spines’ and ‘double series, internal series fused to spine’ would render the character parsimony-uninformative. This approach is therefore not taken, the language of the character is adjusted, and no coding modifications were necessary.
- **210 (osteoderms, articulation); 211 (osteoderms, width):** any taxon coded for state 209–0 (osteoderms absent) cannot be coded for these characters.
  - **210. Osteoderms (articulation).** Simple set of osteoderms arranged in one layer if present (0), or double set of two layers, the ventral one fused to the tip of the neural arch (1). [*added by Schoch, 2013*]
  - **211. Osteoderms (width).** An intra-dissorophid character: Narrow median osteoderms (0), or transversely extended plates (1). [*added by Schoch, 2013*]

  Corrected codes include both taxa originally coded for 209-0 and as unknown for character 209.
  - Codes were corrected from both 210–0 and 211–0 for the following taxa that were coded for 209–0: *Proterogyrinus, Greererpeton, Cochleosaurus, Dendrerpeton, Balanerpeton, Trimerorhachis, Dvinosaurus, Isodectes, Thabanchuia, Capetus, Iberospondylus, Acanthostomatops, Micromelerpeton, Limnogyrinus, Branchierpeton, Branchiosaurus, Apateon, Micropholis, Platyrhinops, Amphibamus, Doleserpeton, Gerobatrachus, Onchiodon, Eryops, Sclerocephalus, Glanochthon, Archegosaurus, Australerpeton, Platyoposaurus, Lapillopsis, Rhineceps, Uranocentrodon, Broomistega, Lydekkerina, Siderops, Edingerella, Trematolestes, Callistomordax, Metoposaurus, Paracyclotosaurus, Eocyclotosaurus*, and *Mastodonsaurus.*
  - *Acheloma* and *Phonerpeton* were erroneously coded for 209–1 and should be 209–0; therefore, their codes were also corrected to inapplicable for 210 and 211.
  - *Chomatobatrachus, Sangaia,* and *Batrachosuchus* were originally coded for 209–0 but should be 209–?. Because no postcrania is known for these taxa; therefore, these taxa were also corrected for 210 and 211.
  - The following were coded as unknown for 209 but coded for 210–0 and 211–0 and were corrected to unknown for 210 and 211 here on the basis that the presently known material is too incomplete to confidently assess character 209: *Edops, Adamanterpeton, Nigerpeton, Chenoprosopus, Neldasaurus, Acroplous,*

*Tupilakosaurus, Zatrachys, Watsonisuchus, Benthosuchus, Trematosaurus, Wetlugasaurus, Parotosuchus,* and *Cyclotosaurus.*

- *Eryosuchus* was originally uncoded for 209 but coded for 210 and 211. Complete skeletons of this taxon are known (Gower & Sennikov, 1997:61), though they have not been described (but most postcranial characters were coded), and since there is no mention of osteoderms (as with all other capitosaurs), it can be reliably inferred that none were present. The same applies to *Eocyclotosaurus*, known from the entire skeletal region but only through disarticulated material (Rinehart & Lucas, 2016). Because the material comes from bonebeds, it is felt that this is sufficient to code ‘osteoderms absent.’ Both taxa were then corrected from state 0 to inapplicable for characters 210 and 211.
- *Triadobatrachus, Leptodactylus, Xenopus, Karaurus, Ambystoma, Hynobius, Cryptobranchus, Ichthyophis, Epicrionops,* and *Eocaecilia* were updated from unknown to inapplicable for consistency.

- **213 (skull, length):** this character, originally taken from Anderson et al. (2008), very precisely differentiates small-bodied taxa; in historical analyses of lissamphibian origins, most candidate taxa (e.g., amphibamiform dissorophoids and ‘microsaurs’) have skull lengths less than 50 mm. Only 10 of the 54 taxa coded by Anderson et al. have skull lengths exceeding 70 mm (state 1–0 there and 213–0 here). By comparison, only 15 of the 72 taxa sampled by Schoch (2013) have skull lengths less than or equal to 70 mm. Therefore, the character is asymmetric in that it delineates small-bodied taxa but fails to capture trends among medium to large taxa, which are certainly informative. Taxa as disparately sized as *Acanthostomatops* (largest specimen around 75 mm; Witzmann & Schoch, 2006) and *Mastodonsaurus* (exceeding 1,000 mm; Schoch, 1999) were aggregated into a single state despite differing by a much greater magnitude than the taxa differentiated by the other states (between 0 and 70 mm). Even if the objective of a particular study is the question of lissamphibian origins, which necessarily focuses on small-bodied taxa, proper treatment of all taxa should more comparably account for all body sizes.

  To address this, additional size bins are appended to this character here. It is not possible to define states in 20 mm increments given limitations on number of character states in standard analytical programs, so new bins are semi-arbitrarily defined and increase in increment (30, 50, 100, 200). Bins were added in this fashion until the next state would result in the character being parsimony-uninformative.

  - Four new states are appended to the ‘front’ of the existing character to maintain an ordered sequence. All taxa coded for original states 1, 2, or 3, were automatically ‘increased’ by four (i.e. a taxon coded for 213–1 is now 213–5), but codes were then checked for all taxa.
  - *Balanerpeton* was originally coded for ‘>70 mm’ but should be ’30-50 mm’ (Milner & Sequeira, 1993:334).
  - *Limnogyrinus* was originally coded for ’30-50 mm’ but should be ‘< 30 mm’ (Schoch & Witzmann, 2018:296; see also previous scale of reconstructions, e.g., Schoch & Milner, 2014:fig. 25A).
  - *Apateon* was originally coded for ’30-50 mm’ but should be ‘< 30 mm’ (Schoch, 2004; see also previous scale of reconstructions, e.g., Boy, 1987:fig. 2C; Schoch & Milner, 2014:fig. 26C).

    - *Cacops* was changed to partial uncertainty for the revised 2/3 because of uncertainty over the synonymy of '*Trematopsis seltini*' with *C. aspidephorus* specifically (Milner, 1985a).
- **225 (septomaxilla):** like palpebral ossifications, gular osteoderms, scalation, and trunk osteoderms, septomaxillae may be subject to taphonomic loss, especially in taxa in which it is an intranarial ossification that is not integrated with the dorsal surface of the skull roof (and even then, loss is not uncommon; e.g., Jeannot et al., 2006). The question becomes what standard is needed to code definitive absence (225–1), and the previous codings (entirely by Pardo et al., 2017a) are difficult to understand. Among non-stereospondyls, only *Proterogyrinus* and *Doleserpeton* were left uncoded. These taxa are known from fairly complete but semi-disarticulated and distorted skulls, but so too are taxa coded for 225–1, such as *Balanerpeton*, *Micromelerpeton*, *Limnogyrinus*, *Apateon*, and *Platyrhinops*. Incidentally, Sigurdsen & Bolt (2010:1361, fig. 2A) did report septomaxillae in the holotype of *Doleserpeton*. Codes do not appear linked to verbatim descriptions; Clack & Milner (2010:279) stated that no septomaxilla was observed in *Platyrhinops* but mentioned the presence in other 'amphibamids' and suggested that it might be found in all 'amphibamids' and merely be too difficult to observe in some specimens. Similarly, Milner & Sequeira (1993:335) noted that despite the lack of evidence for septomaxillae in *Balanerpeton*, "none has a well-preserved snout and such a small bone may have been subject to postmortem loss." Among stereospondyls, several taxa known from complete and undistorted skulls at the time of their study were also left uncoded (e.g., *Australerpeton, Chomatobatrachus*, *Edingerella*, *Benthosuchus*), while taxa known from only one or two skulls of similar preservation were coded for 225–1 (e.g., *Plagiosuchus*, *Trematolestes*, *Paracyclotosaurus*).
    - Because of this ambiguity, the character was recoded in full, with modifications to the states. The revised 225–0 refers to an ossified septomaxilla that is entirely intranarial. The revised 225–1 is maintained as referring to an unossified septomaxilla and is coded only when at least one complete and undistorted snout is known for a taxon (e.g., *Greererpeton*; Smithson, 1982). The new 225–2 ossified refers to a septomaxilla that is integrated with the dorsal surface of the skull roof and that is ornamented. The two 'present' states are left as a bracket around the 'absent' state as there is no clear fashion in which to order them to create a logically ordered character.
    - *Neldasaurus* was originally coded as lacking a septomaxilla, in agreement with the original description by Chase (1965), but this ossification was reported and figured by Schoch (2018a), and the code was accordingly updated.
    - *Capetus* was newly coded for 225–0/2; the uncertainty reflects Sequeira & Milner's (1993:661) comments that the septomaxilla probably articulated with the anterior edge of the lacrimal, but it is dislodged in the only specimen in which it is preserved, and it is unclear whether it might have had an ornamented dorsal exposure. *Metoposaurus* was changed from 225–1 to 225–0/1; a septomaxilla has never been confidently identified in this taxon, but there are disputed reports of septomaxillae in other metoposaurids (e.g., Chowdhury, 1965), and Chakravorti & Sengupta (2018) mentioned a thin plate-like ossification within the naris of *M. krasiejowensis* that they interpreted as a septomaxilla; as this has yet to be figured in this taxon, it is treated as partial uncertainty.

    - *Australerpeton*, *Chomatobatrachus*, *Edingerella*, *Benthosuchus*, *Trematosaurus*, and *Cyclotosaurus* were newly coded for 225–1. *Lydekkerina* was newly coded for 225–2.
- **257 (sphenethmoid); 324 (sphenethmoid); 328 (dorsomedial process of the sphenethmoid); 329 (dorsomedial process of the sphenethmoid); 330 (dorsal sutural surface of the sphenethmoid); 332 (length of lateral walls of sphenethmoid); 333 (floor of sphenethmoid); 341 (anterior wall of sphenethmoid); 342 (anterolateral process of sphenethmoid):** Any taxon coded for 257–1 (sphenethmoid unossified) cannot be coded for these characters.
    - *Lyrocephaliscus* was originally coded for 257–1, but Säve-Söderbergh (1936:23, 27–30, figs. 7, 9–10) described an ossified sphenethmoid in great detail (figures of which confirm that the identification is correct and not an outdated use of the name as with some other anatomical terms in that publication). It was changed for this character and then newly coded as 324–0, 332–0, 340–0/2, 341–1, and 342–0.
    - Character 324: Codes were corrected from 324–0 (sphenethmoid covered dorsally) for *Acroplous, Isodectes, Acanthostomatops, Apateon, Amphibamus, Australerpeton,* and *Laidleria*. *Lapillopsis* was uncoded from 324–0 because it was originally left uncoded for 257. *Eryops* and *Gerrothorax* were newly coded for 324–0/1.
    - Character 333: it could be argued that the 'floor' of the sphenethmoid can be ascertained from the dorsal surface of the cultriform process. However, this presumes that the sphenethmoid articulates with the process and forms a single unpaired ossification (rather than paired ossifications that flank the process but that do not override it dorsally). In taxa without an ossified sphenethmoid, this cannot be known, and cartilaginous structures are not coded in this matrix (e.g., a taxon with unossified carpals likely has cartilaginous carpals but is coded as lacking carpals). Therefore, codes were corrected from 333–0 for *Acanthostomatops, Apateon, Amphibamus, Australerpeton, Lapillopsis, Siderops, Batrachosuchus, Trematolestes, Callistomordax, Metoposaurus,* and *Rileymillerus*. Codes were corrected from 333–2 for *Acroplous, Isodectes,* and *Laidleria*
    - Codes for these seven characters were updated from unknown to inapplicable (when originally coded as unknown) for the above taxa for consistency.
- **259 (pterygoid, lateral process):** the name and character states of this character are confusing.
    - **259. Lateral process of pterygoid contribution to posttemporal fossa absent (0); present (1).** [*added by Pardo et al., 2017a*]

It is inferred here that the 'posttemporal fossa' is the same opening in the occiput that may alternatively be termed the posttemporal fenestra or the posttemporal foramen in temnospondyls (e.g., Warren, 1998a:173; Bolt & Chatterjee, 2000:674; Steyer et al., 2006:23; Eltink et al., 2016:847). However, this opening is only formed by some combination of the tabular, the postparietal, the exoccipital, and rarely the opisthotic in temnospondyls (see *Eryops* as an example with an opisthotic contribution; Sawin, 1941); the pterygoid has no role in this opening, at least not in posterior view. State 1 was coded for *Balanerpeton* (but not *Dendrerpeton*), most dissorophoids (but not for *Dissorophus* and as unknown for *Micromelerpeton*, *Limnogyrinus*, *Acheloma,* and *Phonerpeton*), eryopids, *Archegosaurus* and *Platyoposaurus* (but not sclerocephalids), *Lapillopsis*, *Peltobatrachus*, *Laidleria*, and *Metoposaurus* (but no other 'higher stereospondyls'). Most other temnospondyls were coded for 259–0, although some of the best-known taxa like *Trimerorhachis* were left uncoded. This character was previously sampled in early tetrapod matrices by Anderson et al. (2008; character 120), Huttenlocker et al. (2013; character 120), and Pardo et al. (2017b; character 85), among other analyses. In those broader samples of early tetrapods, 259–1 was restricted to some temnospondyls and as partial uncertainty in various lissamphibians; curiously, this uncertainty was not coded for any lissamphibians here (instead they were coded for 259–0).

Firstly, it is questionable whether the posttemporal fossa can even be defined or proven to exist in some of these taxa only represented by flattened or disarticulated specimens, such as *Balanerpeton* and *Gerobatrachus*. The character should not be referring to the transverse flange of the pterygoid (already captured in character 121) and its relation to the subtemporal fossa/vacuity for the adductor musculature or to the presence or absence of a palatoquadrate fissure (character 120). It also cannot refer to the opening sometimes called the 'subtympanic fossa', which occurs along the anteromedial edge of the pterygoid medial to a contact with the squamosal (e.g., *Gerrothorax*; Schoch & Witzmann, 2012), as many taxa coded for 259–1 lack this opening entirely (e.g., *Archegosaurus*; Witzmann, 2005). If the character refers to some internal aspect of the fossa rather than the contributing elements on the occiput, this too would be very hard to reconcile in crushed or distorted specimens and perhaps even in undistorted specimens for which the braincase is not well-known (practically all temnospondyls given the dearth of CT data). The frequency of taxa represented by such specimens (most Carboniferous taxa) is discordant with the relatively high coding rate for this character; only 13 temnospondyls and the karaurids were left as unknown by Pardo et al. Finally, it is not actually clear which process is the 'lateral' one; whether the quadrate ramus or the palatine ramus is more laterally oriented is variable. However, if the interpretation of 'posttemporal fossa' is correct, it would have to be the quadrate ramus, which frequently has an ascending lamina that may contact the skull roof (or form the palatoquadrate fissure when it fails to do so).

In my opinion, the character is so ambiguous as to be irreproducible. I can identify no condition shared among all taxa coded for 259–1 that is not found in 259–0, whether that is the nature of the transverse flange (e.g., rounded versus angular corner, ventrally descending versus flat), the ascending flange (e.g., contacting skull roof or not, forming oblique crest or not), or in the posttemporal fossa (e.g., which elements frame it). For this reason, the character is excluded.

- **265 (splenial exposed laterally); 267 (ventral border of Meckel's fossa); 274 (symphysis); 310 (dorsal margin of splenial); 322 (splenial teeth)**: any taxon without a splenial (264–2; exclusive to lissamphibians) cannot be coded for these characters.
    - Character 265: Codes were corrected to inapplicable from 265–1 (lateral exposure absent) for *Eocaecilia*.
    - Character 274: Codes were corrected to inapplicable from 274–1 (symphysis formed by dentary alone) for *Triadobatrachus, Leptodactylus, Xenopus, Ambystoma, Hynobius,* and *Cryptobranchus.*
    - Character 322: Codes were corrected to inapplicable from 322–0 (splenial teeth present) for *Ichthyophis* and *Epicrionops*. Even if the splenial is incorporated into

the pseudodentary, the lingual tooth row was treated as belonging to the coronoid and cannot also be treated as belonging to the splenial. Codes were corrected from 322–1 (splenial teeth absent) for *Triadobatrachus*, *Leptodactylus*, *Ambystoma*, *Hynobius*, and *Cryptobranchus*.

    - Codes for these five characters were updated from unknown to inapplicable (when originally coded as unknown) for the above taxa as well as for *Eocaecilia* for consistency. Note that for some of these characters, certain taxa were originally coded as inapplicable, but this was apparently reverted or lost in translation by Schoch et al. (2020); for example, all five extant batrachians were coded as inapplicable for character 267.

- **266 (Meckelian fossae):** this character was confusing largely because of the terminology; the openings on the lingual surface of the lower jaw are not typically referred to as ‘fossae’ in temnospondyls but more commonly as ‘fenestrae,’ ‘foramina,’ or ‘windows.’ This can then be complicated by inconsistent use between workers. Some differentiate the large opening found anteroventral to the adductor fossa that is usually framed by some combination of the postsplenial, the angular, and the prearticular in temnospondyls, from additional, smaller openings that are often entirely within the postsplenial (e.g., Sawin, 1941:429–430; Milner & Sequeira, 1993:338; Hewison, 2007:36), whereas others refer to any opening on this surface by the same name regardless of relative size or position (e.g., Langston, 1953:387). Based on how the character was originally coded (e.g., for ‘2 or more’ in taxa with only one relatively large opening but several smaller ones; Fig. 28A-B), it seems that any opening on the lower half of the lingual surface (i.e., excluding more posteriorly or dorsally situated ones near the glenoid) is counted as a Meckelian fossa in this instance, and this definition is applied here in newly coding the following taxa:

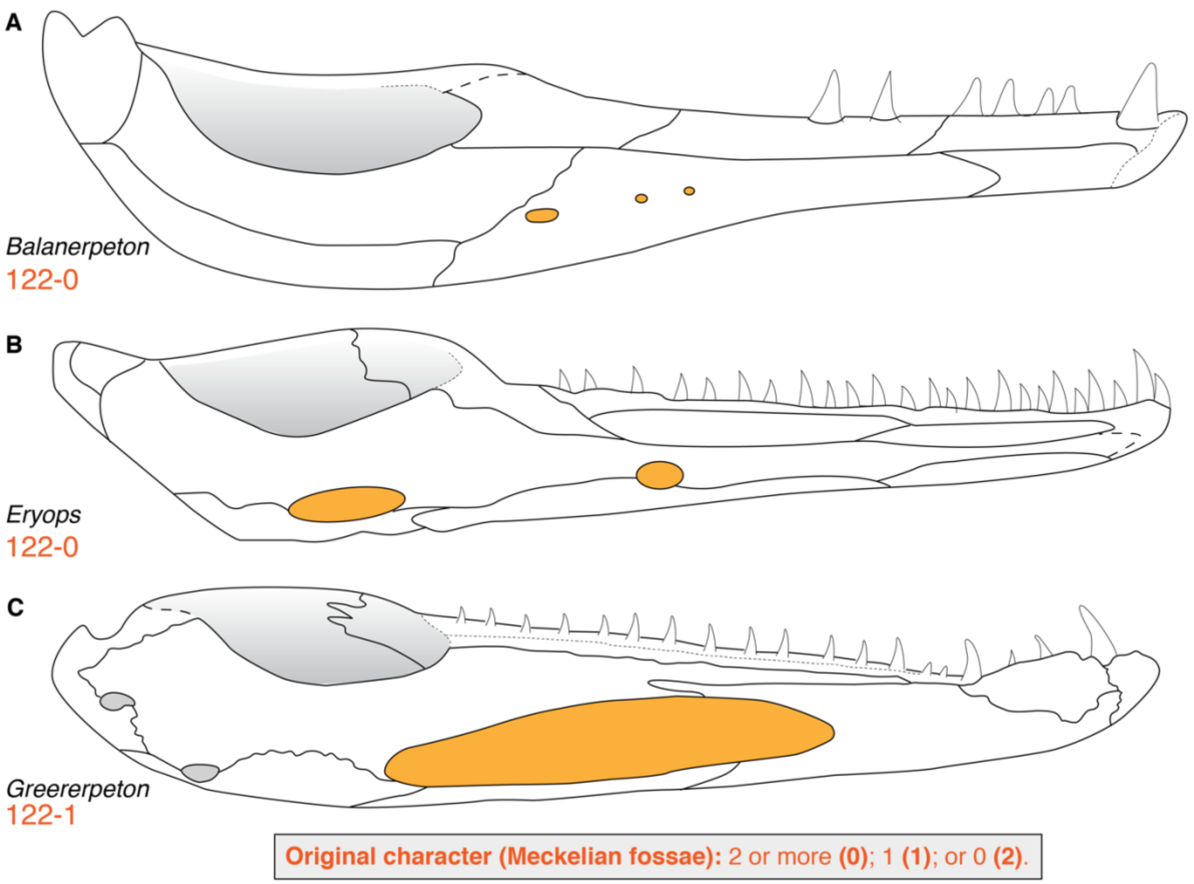

**Figure 28. Comparison of number of Meckelian fossae among temnospondyls and *Greererpeton*. A**, the 'dendrerpetid' *Balanerpeton woodi* (after Milner & Sequeira, 1993); **B**, the eryopoid *Eryops megacephalus* (after Schoch & Milner, 2014); **C**, the colosteid *Greererpeton burkemorani* (after Bolt & Lombard, 2001). Coronoid dentition is intentionally omitted for simplicity. Orange text represents the original character and codes from Schoch (2013). Silhouettes not to scale.

  - Codes for partial uncertainty of 266–0/1 were newly added for *Proterogyrinus, Chenoprosopus, Acanthostomatops, Micromelerpeton, Doleserpeton, Gerobatrachus, Acheloma, Glanochthon, Plagiosuchus, Trematosaurus, Parotosuchus*, and *Paracyclotosaurus*. *Cacops* was newly coded for 266–1/2. This reflects the observation that there is at least one large posterior opening, but the possible presence of additional small openings anteriorly is unclear due to limited exposure or poor preservation. *Zatrachys, Phonerpeton, Metoposaurus*, and *Cyclotosaurus* were newly coded for 266–0.
  - *Dendrerpeton* was corrected from 266–1 to 266–0 (Holmes et al., 1998; Arbez et al., 2022).
  - *Chomatobatrachus* was corrected from 266–0 to 266–1 (Cosgriff, 1974:61, fig. 42c).
- **267 (ventral border of Meckel's fossa):** there is some ambiguity in which fossa this character refers to because of the interpretation of character 266. Here, it is explicitly restricted to the posterior foramen found in practically all temnospondyls, typically larger than any other foramen on the lingual surface, and thus excludes any additional openings

situated more anteriorly. There is also some ambiguity in the treatment of the character in the previous analysis by Pardo et al. (2017a).

- **267. Ventral border of Meckel's fossa.** Splenial (0), angular (1). [*added by Pardo et al., 2017a*]

It appears that 'splenial' (267–0) refers to both the splenial and the postsplenial based on taxa in which it participates in the opening, but the greater issue is that the two states are not mutually exclusive. For example, in *Mastodonsaurus*, the postsplenial and the angular contribute to the ventral border, but this taxon was coded as 267–0 ('splenial'). This was treated as the same condition as in *Greererpeton*, in which the splenial, postsplenial, and angular all contribute to the ventral margin (Bolt & Lombard, 2001:figs. 4.2, 5.2; Fig. 29A) and to *Balanerpeton*, in which a much smaller opening in the same position is entirely enclosed by the postsplenial (Milner & Sequeira, 1993:fig. 5D; Fig. 29B. Therefore, this character is substituted with one from McHugh (2012) that more fully captures the different configurations. All taxa were recoded.

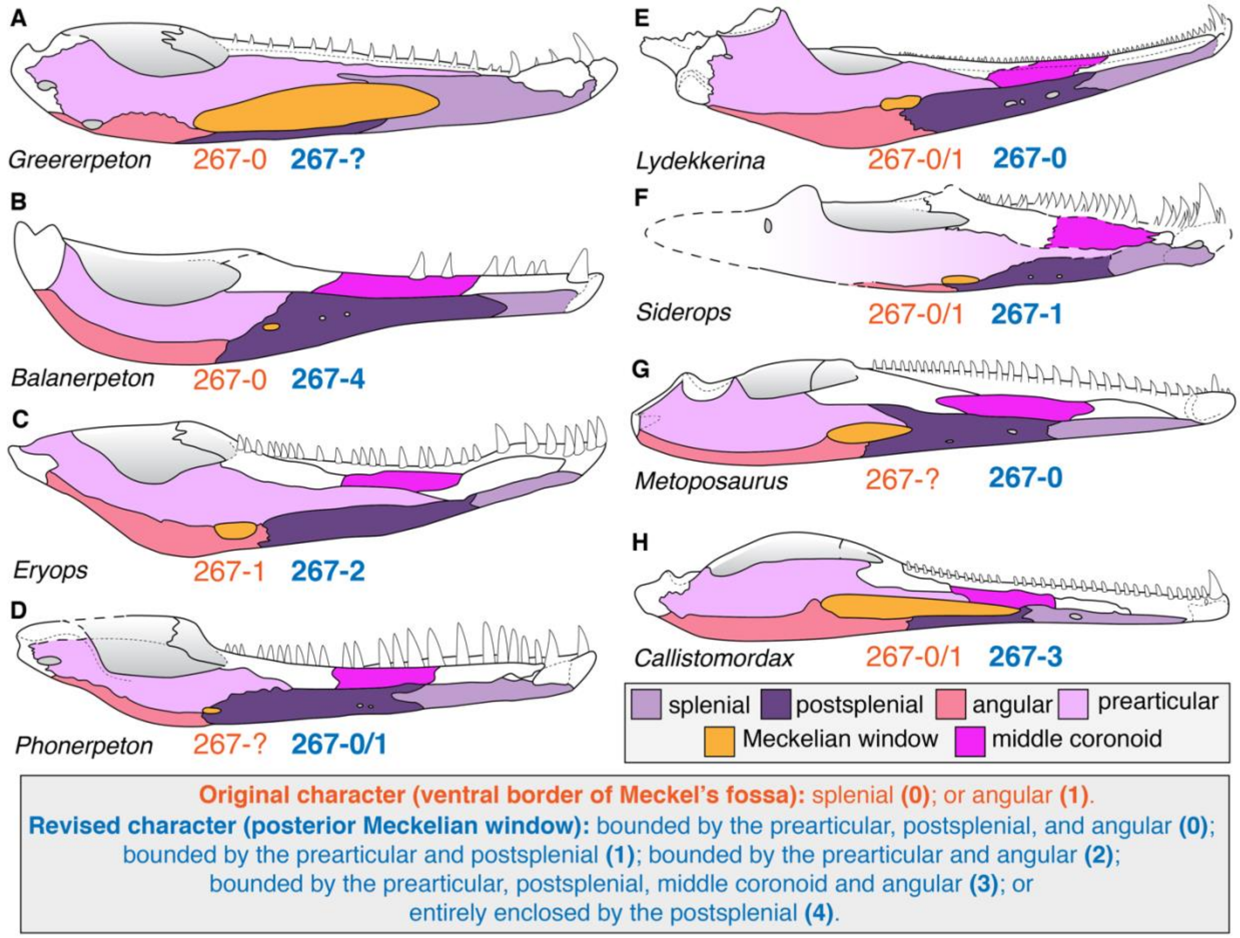

**Figure 29. Comparison of the elements framing the posterior Meckelian window in temnospondyls and *Greererpeton*.** **A**, the colosteid *Greererpeton burkemorani* (after Bolt & Lombard, 2001); **B**, the 'dendrerpetid' *Balanerpeton woodi* (after Milner & Sequeira, 1993); **C**, the eryopoid *Eryops megacephalus* (after Schoch & Milner, 2014); **D**, the trematopid *Phonerpeton pricei* (after Dilkes, 1990); **E**, the lydekkerinid *Lydekkerina huxleyi* (after Hewison, 2007); **F**, the chigutisaurid *Siderops kehli* (after Warren & Hutchinson, 1983); **G**, the metoposaurid *Metoposaurus krasiejowensis* (after Sulej, 2007); **H**, the metoposauroid *Callistomordax kugleri* (after Schoch, 2008). Coronoid dentition is intentionally omitted for simplicity. There are coronoids adjacent to the dentary in G. burkemorani, but their differentiation is unclear posteriorly, and they are not widely exposed lingually. Orange text

represents the original character and codes from Pardo et al. (2017a); blue text represent the revised character and codes. Silhouettes not to scale.

- **300 (coracoid foramina); 301 (scapulocoracoid):** state 300–2 was changed from ‘2’ to ‘2 or more’ to account for the presence of up to three foramina in *Sclerocephalus* (Boy, 1988). The only coding modification that appears linked to this revision to the state was newly coding *Sclerocephalus* for 300–2. Any taxon coded for 301–1 (only scapula) or 301–2 (scapulocoracoid absent) should be coded as inapplicable.
    - *Trimerorhachis*, *Acroplous*, *Isodectes*, *Limnogyrinus*, *Apateon*, *Micropholis*, *Glanochthon*, *Broomistega*, *Laidleria*, *Siderops*, *Edingerella*, *Trematolestes*, *Metoposaurus*, and *Paracyclotosaurus* were updated from unknown to inapplicable.
- **311 (parietal, anterior waisting):** this character refers to whether the parietal is constricted anteriorly and appears to overlap with a character added by Schoch et al. (2020; 360 therein) for the shape of the parietal.
    - **311. Parietal anterior waisting.** Absent (0), present (1). [*added by Pardo et al., 2017a*]
    - **360. Parietal.** Rectangular or quadrangular plate (0), or L-shaped (1). [*added by Schoch et al., 2020*]

  Schoch et al.’s coding intended to capture an exclusively lissamphibian feature in which the parietal is ‘L-shaped’ as a result of a marked posteromedial expansion of the orbit, which, in the absence of the postfrontals, results in a prominent incision into the parietal. However, this is the same as an ‘anteriorly waisted parietal’ (Fig. 30).

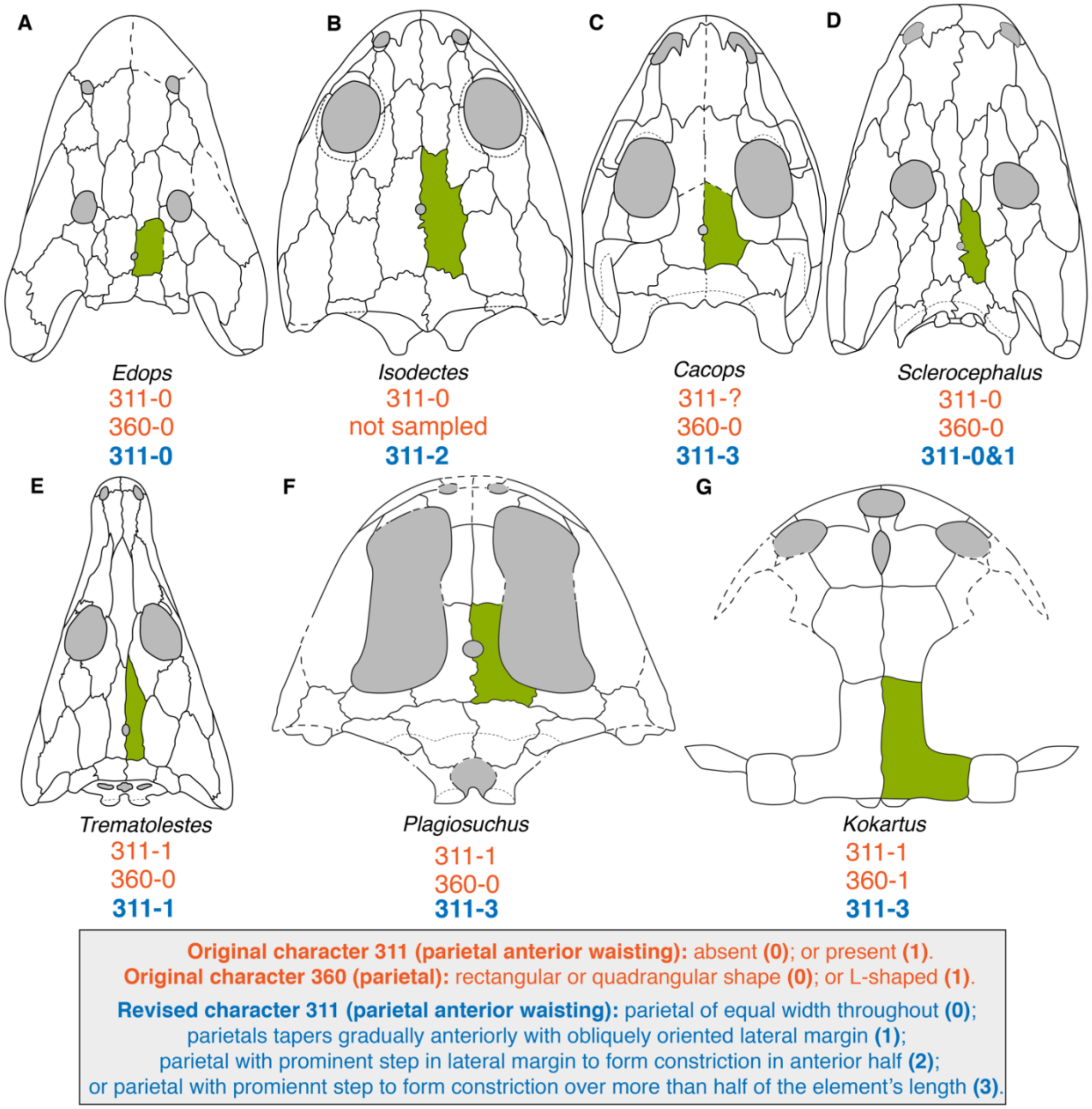

**Figure 30. Comparison of parietal morphologies among temnospondyls and lissamphibians. A**, the edopoid *Edops craigi* (after Schoch & Milner, 2014); **B**, the dvinosaur *Isodectes obtusus* (after Sequeira, 1998); **C**, the dissorophid *Cacops aspidephorus* (after Anderson et al., 2020); **D**, the stereospondylomorph *Sclerocephalus haeuseri* (after Schoch & Witzmann, 2009a); **E**, the trematosaur *Trematolestes hagdorni* (after Schoch, 2006); **F**, the plagiosaurid *Plagiosuchus pustuliferus* (after Damiani et al., 2009); **G**, the karaurid *Kokartus honorarius* (after Skutschas & Martin, 2011). Orange text represents the original characters and codes from Pardo et al. (2017a) and Schcoh et al. (2020); blue text represents the revised character and codes. Silhouettes not to scale.

The issue of redundancy is irrelevant for this study since character 360 is not sampled, but it provided an opportunity to modify character 311 to more precisely delineate different conditions since a 'waisted' parietal could be one in which the margins taper anteriorly but are not markedly incised by the postorbital, or they could appear to taper more abruptly and to be 'stepped' because of a prominent incision. State 0 remains as the condition where there is no anterior waisting (i.e. the lateral margins are parallel; Fig. 30A). State 1 was redefined as where the parietal tapers in width with a continuous lateral margin (i.e. it is not markedly incised / stepped; Fig. 30E). The new state 2 and state 3 then refer to taxa with a markedly incised / stepped lateral margin to produce 'anterior waisting.' They are differentiated by the relative length of the constricted region (< 50% of the total parietal length vs. > 50% of the total parietal length); the new state 3 captures the condition of most lissamphibians (Fig. 30G), although it also occurs in some temnospondyls (e.g., *Plagiosuchus*; Fig. 30F). All taxa were recoded.

- **316 (squamosal-frontal contact):** the original coding for this character was peculiar, as this contact is absent in all temnospondyls in which these two elements are defined, but the character was largely coded as unknown for temnospondyls except for the majority of capitosaurs and trematosaurs. These codes were updated for all taxa where the frontal is defined posteriorly, and the squamosal is defined anteriorly; subsequently, every temnospondyl other than *Rileymillerus* was coded for 316–1.
- **319 (quadrate-maxillopalatine):** this is a caecilian character that should not be coded for taxa lacking a maxillopalatine, even if they retain separate maxillae and palatines (quadrate-maxilla separation is covered by character 318). A few temnospondyls with known palates were also previously left as uncoded (e.g., *Zatrachys, Micromelerpeton, Chomatobatrachus*; Langston, 1953; Cosgriff, 1974; Boy, 1995; Schoch, 1997), but almost all temnospondyls were coded for 319–1. Codes were therefore changed to inapplicable for all non-lissamphibians, including *Chinlestegophis,* which lacks a maxillopalatine regardless of the validity of the interpreted fusion of the maxilla and lacrimal in this taxon.
    - Codes were corrected from 319–1 (quadrate and maxillopalatine do not contact laterally) for *Proterogyrinus, Greererpeton, Edops, Adamanterpeton, Cochleosaurus, Nigerpeton, Chenoprosopus, Dendrerpeton, Balanerpeton, Trimerorhachis, Neldasaurus, Acroplous, Isodectes, Capetus, Iberospondylus, Acanthostomatops, Apateon, Micropholis, Platyrhinops, Amphibamus, Doleserpeton, Gerobatrachus, Dissorophus, Acheloma, Phonerpeton, Onchiodon, Eryops, Sclerocephalus, Glanochthon, Archegosaurus, Platyoposaurus, Lapillopsis, Peltobatrachus, Rhineceps, Broomistega, Lydekkerina, Laidleria, Siderops, Batrachosuchus, Plagiosuchus, Gerrothorax, Edingerella, Benthosuchus, Trematolestes, Lyrocephaliscus, Callistomordax, Parotosuchus, Paracyclotosaurus, Mastodonsaurus, Rileymillerus, Chinlestegophis, Leptodactylus, Xenopus, Karaurus, Ambystoma, Hynobius,* and *Cryptobranchus*. This set of recodings rendered the character parsimony-uninformative.
    - Codes were updated from unknown to inapplicable for *Limnogyrinus, Cacops, Ecolsonia, Australerpeton, Chomatobatrachus, Trematosaurus, Metoposaurus, Cyclotosaurus,* and *Eocaecilia*.
    - *Sangaia* was uncoded from 319–1 because the region of the palate where the palatine normally occurs is unknown, and it has otherwise not been identified (e.g., as a dislodged element), so the presence or absence of a maxillopalatine cannot be assessed.
- **321 (tentacle):** the a tentacle is a derived condition found in gymnophionomorphs. The character was coded for 321–0 (tentacle absent) in the majority of temnospondyls but was uncoded for the overwhelming majority of stereospondyls (except for *Parotosuchus,*

*Paracyclotosaurus*, and *Mastodonsaurus*). There were instances where one member of a clade was coded but another member of no poorer quality was not (e.g., *Acanthostomatops* was coded for 321–0 but *Zatrachys* was left uncoded). In the absence of any comments on personal observation of certain taxa by Pardo et al. (2017a), there is no evidence that the minority of coded taxa were all personally examined by those workers. The possible correlate of the tentacle that Pardo et al. (2017a) identified in *Chinlestegophis* is a sulcus below the orbit in the position where the nasolacrimal duct normally passes through the lacrimal. Uncoded taxa were generally, but not always, aquatic taxa with lateral line systems, and the coding of some taxa with lateral lines (e.g., *Mastodonsaurus*) and the lack of coding for some without (e.g., *Cacops*) suggests that these cells were not left uncoded because of uncertainty about whether a groove represented a lateral line, the nasolacrimal duct, or a tentacle. The identification of a tentacular correlate in *Chinlestegophis* remains to be further explored (Pardo et al., 2017a, equivocated on the issue and coded it for partial uncertainty).
    - All taxa with sufficiently preserved suborbital regions were newly coded for 321–0 in the absence of the identification of a similar sulcus. This includes *Capetus, Zatrachys, Micromelerpeton, Limnogyrinus, Cacops, Broiliellus, Australerpeton, Platyoposaurus, Lapillopsis, Rhineceps, Uranocentrodon, Broomistega, Lydekkerina, Chomatobatrachus, Sangaia, Laidleria, Siderops, Batrachosuchus, Plagiosuchus, Gerrothorax, Edingerella, Benthosuchus, Trematosaurus, Trematolestes, Lyrocephaliscus, Callistomordax, Metoposaurus, Cyclotosaurus,* and *Rileymillerus*.
- **325 (larval stage):** the original coding for this character is difficult to interpret and therein to apply to newly added taxa. Pardo et al. (2017a) originally coded most lissamphibians and a number of European temnospondyls with well-sampled ontogenies (e.g., *Micromelerpeton*, *Sclerocephalus*) as having larval stages (325–0). Conversely, *Gerrothorax, Trematolestes, Callistomordax*, and *Mastodonsaurus* were coded as having direct development (325–1). The stereospondyls coded for 325–1 appear entirely random. At least one, *Trematolestes*, is stated to have a larval form (Schoch, 2006:38). Literature on the other three makes no mention of direct development, which would be unexpected in these obligately aquatic, paedomorphic taxa, and arguably it cannot be well-assessed in at least *Callistomordax* based on the narrow known ontogenetic range (Schoch, 2008a) or in *Mastodonsaurus* based on the absence of known small individuals (Schoch, 1999). These are also not the only sampled stereospondyls with partial ontogenetic trajectories (e.g., *Lydekkerina, Edingerella,, Trematosaurus, Metoposaurus*), some much more complete than the coded taxa (e.g., *Benthosuchus*) so it is unclear why only these four specifically were coded. The character does not prescribe anatomical features considered to be unequivocal evidence of larval stages (e.g., Boy & Sues, 2000).
    - *Trematolestes* was recoded to 325–0, and the other three stereospondyls were recoded as unknown.
    - *Limnogyrinus* was newly coded for 325–0.
- **344 (dentary, marginal dentition):** this is an intuitive character to capture the double marginal tooth row that occurs in gymnophionomorphs (344–1), but the treatment of character 271 (coronoid teeth) and character 272 (size of coronoid teeth) by Pardo et al. (2017a) means that this character is simultaneously redundant and in conflict with those characters; the teeth cannot be considered to be both coronoid teeth homologous with

those on distinct coronoids of non-gymnophionomorphs and a second row of marginal teeth distinct from all temnospondyls. By the same token, the partial lingual row of teeth in the symphysial region of *Doleserpeton* also should not be treated as a double row (it was originally coded as having a single row). Therefore, this character was changed to refer to the presence or absence of a pseudodentary, which avoids having to make any coding modifications other than coding taxa that were previously treated as unknown and which adds another character that is useful for supporting gymnophionomorph monophyly.

  - All temnospondyls other than *Chinlestegophis* were previously coded for 344–0 or were left as unknown. *Chinlestegophis* was changed to 344–0 and should not have been coded for 344–1 under the original interpretations since the coronoids remain distinct, and there is only one tooth row on the dentary. *Chenoprosopus, Neldasaurus, Zatrachys, Micromelerpeton, Limnogyrinus, Dissorophus, Cacops, Glanochthon, Peltobatrachus, Trematosaurus, Metoposaurus, Parotosuchus, Cyclotosaurus, Triadobatrachus, Leptodactylus, Xenopus, Kokartus*, and *Karaurus* were newly coded as 344–0.

The following are characters with partial dependencies that did not result in any coding changes but for which cells coded as '?' were updated to '-' for consistency.

- **175–177 (cleithrum):** any taxon coded for 348–1 (cleithrum absent) cannot be coded for these characters. Codes updated for: *Triadobatrachus, Leptodactylus, Xenopus, Karaurus, Kokartus, Ambystoma, Hynobius, Cryptobranchus, Ichthyophis, Epicrionops,* and *Eocaecilia.*
- **309 (number of distal tarsals):** any taxon coded for 206–1 (tarsals all unossified) cannot be coded for this character. Codes updated for: *Trimerorhachis, Acanthostomatops, Micromelerpeton, Apateon, Sclerocephalus, Glanochthon, Siderops, Plagiosuchus, Gerrothorax, Trematolestes, Callistomordax, Metoposaurus, Paracyclotosaurus, Mastodonsaurus,* and *Karaurus*.
- **335 (os basale):** any taxon without an os basale should be coded as inapplicable. *Proterogyrinus*, *Greererpeton*, and all sampled temnospondyls were updated from unknown.

#### Appendix 2.5. Taxon-level modifications.

##### Appendix 2.5.1. Taxon sampling.

The following section details relevant comments on the treatment of certain taxa sampled in my analysis related to discrepancies in the concepts of taxa; revised osteologies published subsequent to some or all of the previous versions of this matrix; and taxon substitutions. Taxa are listed in the same order as they appear in the matrix, which mostly follows taxonomic placement. Character numbering in this section refers to the original numbering of Schoch et al. (2020) regardless of the omission of several characters from the dataset used in our analyses (Characters 14, 47, 48, 61, 78, 81, 100, 115, 231, 232, 253, 259, 334); the new numbering scheme is detailed below in Appendix 6.

- ***Chenoprosopus milleri***: Cranial material of this taxon is well-characterized, primarily by Langston (1953), but postcranial material is lacking. Conversely, a second species,

‘*Chenopropsopus lewisi*,’ has abundant postcranial material known from the holotype (Hook, 1993). The synonymizing of *C. lewisi* with *C. milleri* by Reisz et al. (2005) is generally accepted by other workers (e.g., Schoch & Milner, 2014), and would therefore permit coding of postcranial postcranial characters *C. milleri*. However, the holotype and only specimen of ‘*C. lewisi*’ is much smaller than the largest known specimens of *C. milleri*, with a skull length around 95 mm compared to a range of about 190 to 330 mm for skulls originally referred to *C. milleri*. As a result, aspects of the postcranial skeleton of ‘*C. lewisi*’ likely reflect a degree of immaturity. To avoid creating a chimeric OTU blending morphologies from different ontogenetic stages, any characters (cranial and postcranial) that can only be coded from ‘*C. lewisi*’ were left as unknown unless a given condition can be reasonably inferred as the adult condition based on what is known of edopoid and temnospondyl ontogeny (easier to discriminate for presence/absence characters compared to qualitative or quantitative comparisons). For example, the presence of intercentra is regarded as codable because the intercentra never disappear in late stages of ontogeny, whereas the absence of an ossified pubis is not regarded as codable because this is typically one of the last elements to ossify in other temnospondyls (e.g., *Sclerocephalus*; Schoch & Witzmann, 2009a).

- ***Dendrerpeton helogenes:*** The OTU long called ‘*Dendrerpeton acadianum*’ in temnospondyl analyses that derives from material described by Godfrey et al. (1987) and Holmes et al. (1998) is *Dendrysekos helogenes* sensu Schoch & Milner (2014:25), which means that it is *Dendry. helogenes* and not *Dendre. acadianum* that has served as a common outgroup in temnospondyl analyses; the latter was considered a distinct taxon by Schoch & Milner. Here, codings are restricted to the holotype (NHMUK 4149, a partial skull roof exposed ventrally; Steen, 1934) and to the two specimens described by Godfrey et al. (1987) and Holmes et al. (1998). This did not lead to any coding changes, although it should be questioned whether occipital features can be coded from these specimens (it is unclear from the literature) or if these could only be coded from what is now a taxonomically ambiguous specimen like the occiput described by Robinson et al. (2005) on the basis of CT data (the occiput does not contain any differential characters sensu Schoch & Milner, 2014). Recently, Arbez et al. (2022) restudied the specimen described by Holmes et al. (NSM 987 GF 99.1), synonymized *Dendrysekos* with *Dendrerpeton* (the latter with seniority), and attributed this specimen to *Dendrerpeton* cf. *helogenes* pending future revision of other species of *Dendrerpeton*. Their *Dendre. helogenes* therefore comprises the same specimens as the *Dendry. helogenes* of Schoch & Milner, and this OTU is termed ‘*Dendrerpeton helogenes*’ throughout this study.
    - The following changes were made strictly based on the revision by Arbez et al.: 28 (0 → ?; orbital ossifications may be palpebral or scleral), 57 (0 → ?; the authors identified what they argued was an alary process of the jugal posterodorsal to the ectopterygoid [fig. 2B therein], but it is not clear that this was exposed on the palate, [fig. 2D therein], so the code is left as unknown to be conservative), 257 (? → 0), 310 (0 → 1), 330 (? → 1), 332 (? → 0). Note that several previously coded sphenethmoid characters (e.g., 333, sphenethmoid [floor]) were justified by this revision but should not have been coded previously given that character 257 for the presence/absence of the sphenethmoid was left uncoded.

- ***Neldasaurus wrightae*:** This taxon was described by Chase (1965). The skull was reconstructed in a differing fashion by Schoch & Milner (2014:fig. 20C) without redescription or comment, with proper revision published by Schoch (2018). The following are coding changes made entirely on the basis of the more recent literature that at least post-date Schoch (2013) if not also Pardo et al. (2017a):
    - 39 (0 → 0&1), 87 (? → 0), 164 (? → 0), 228 (3 → 0), 249 (1 → ?; this relates to uncertainty over whether denticles were removed during preparation), 251 (1 → ?; the same uncertainty as with character 249).
- ***Branchiosaurus salamandroides:*** This taxon is known from many specimens, but whether historical specimens from Nýřany that were referred to this taxon are branchiosaurids or rather non-branchiosaurid amphibamiforms or larval forms of much larger temnospondyls (e.g., Milner & Sequeira, 2003; Schoch & Milner, 2008; Werneburg, 2012; Milner, 2018) remains contentious. This raises the question of whether referred material from other localities might also not be branchiosaurid in nature, or at least not definitively referable to *B. salamandroides*. Schoch & Milner's (2014:fig. 26A) reconstruction of the skull is based on the lectotype and "other uncontroversial specimens," although it is unclear what such specimens are (e.g., is it the same as those listed by Schoch & Milner, 2008) and why they are "uncontroversial" (i.e. is it because they have been recently examined by many workers and explicitly agreed upon as being *B. salamandroides* or merely because they were originally identified as such and have never been explicitly questioned as possibly belonging to another taxon). Because the primary dissenter from the historical assignment of at least some specimens is Werneburg, his 2012 description was employed as the primary literature used to assess codes, with the reconstructions of Boy (1987, the literary source for Schoch, 2013) and Schoch & Milner (2014) accepted as a faithful restoration of the complete skull. However, their reconstruction of the portion of the palate visible through the orbit (e.g., with the characteristic 'Y-shaped' ectopterygoid of branchiosaurids) is not adopted because Werneburg (2012:12) indicated that the palate is very poorly known, with no definitive vomer or ectopterygoid known. This led to a substantial number of codes being changed to unknown for these characters (and others).
- ***Cacops aspidephorus:*** The one cacopine OTU has always been a chimera of *Cacops aspidephorus* and *Cacops morrisi* (but not *Cacops woehri*), as noted by Schoch (2013:table 1). The inferred rationale is that the cranial osteology of *Cacops aspidephorus* was essentially unknown until recently (Anderson et al., 2020), but the postcranial skeleton was entirely known (Williston, 1910; Dilkes & Brown, 2007; Dilkes, 2009, 2014), whereas the cranial osteology of *Cacops morrisi* was entirely known (Reisz et al., 2009), but the postcranial skeleton was almost entirely unknown until recently (Gee & Reisz, 2018a). For this reason, the OTU has sometimes been displayed as '*Cacops aspidephorus morrisi*' (e.g., Pardo et al., 2017a). No other taxon was coded from two or more species or as a terminal of a higher order clade, and now that *Cacops aspidephorus* can be more completely coded, I prefer to restrict the OTU to this single species, as there are differences between them that would be captured in phylogenetic codes (Anderson et al., 2020; Gee, 2021) but that apparently have not been previously based on the lack of polymorphisms. Corrections are listed in the taxon-specific section and represent changes made to codes that were mischaracterized regardless of which species is sampled (e.g.,

coding the absence of orbital rims in this OTU; 27–0). The following changes are based strictly on the restriction of the OTU to one species.

  - The following characters were recoded from one state to another: 13 (1 → 1/2), 87 (1 → 0).
  - The following characters were recoded from one state to unknown (?): 102, 104, 126.
  - The following characters were newly coded: 246 (0), 270 (0), 271 (0), 275 (1), 310 (1), 311 (1), 312 (0), 313 (0), 319 (-), 321 (0), 322 (1), 323 (0), 324 (0), 326 (1).

- ***Phonerpeton pricei:*** *Phonerpeton pricei*, as coded by Schoch (2013), was characterized following the concept of Dilkes (1990) and not that of Schoch & Milner (2014) in which some of this material was considered to belong to a separate species of *Phonerpeton*, *Phonerpeton whitei*. Dilkes' concept is followed here.
- ***Acheloma cumminsi:*** This taxon follows the concept of Gee (2020), in which material of '*Acheloma dunni*' from the Richards Spur locality is considered to be synonymous with *Acheloma cumminsi*. Schoch (2013:table 1) noted personal observation of material from the AMNH (where only material of *Acheloma cumminsi* sensu Polley & Reisz, 2011, is held), but Schoch (2013) cited only Polley & Reisz (2011), who described material of '*Acheloma dunni*' reposited at the BMRP and OMNH, as the literary source. The only difference between these two species was the purported absence of a LEP in *A. cumminsi* (e.g., Dilkes & Reisz, 1987; Polley & Reisz, 2011), which was subsequently identified in the holotype (Gee et al., 2019). This OTU was originally coded as having a LEP. It is thus unclear whether this OTU may have originally been a chimeric OTU of two species like with *Cacops*.
- ***Sclerocephalus haeuseri*****:** This taxon is complicated by the noted intraspecific variation within and between different stratigraphically separated lake populations (Schoch & Witzmann, 2009a). This variation reflects ecological differences, with some populations remaining aquatic and retaining features like a lateral line system, whereas others were terrestrial and have correspondingly robust postcrania (Schoch, 2009). Some of these differences are captured in phylogenetic characters (e.g., lateral line sulci). Schoch (2013:table 1) listed Schoch & Witzmann (2009a) as the only literary source of data for this taxon, and the latter specifically focused on characterizing the Lake Jeckenbach population (L–O 6; see p. 137, 145 therein), from which the largest sample of adults is known. I therefore interpret the existing states for this taxon to be exclusively for this population and do not expand the coding here as part of my conservative approach to minimizing coding changes.
- ***Australerpeton cosgriffi*****:** The osteology of this taxon has been substantially expanded subsequent to Schoch (2013) by Eltink & Langer (2014); Eltink et al. (2016); and Azevedo et al. (2017). However, contrary to some of the stereospondyls listed further below with similar situations regarding new literature, the newer literature for *A. cosgriffi* rarely led to a new code for a previously uncoded cell; most coding modifications, especially for postcrania, could be sourced from Dias & Schultz (2003, the literary source for Schoch, 2013), and the cranial reconstruction of Eltink et al. differs little from the original by Barberena (1998). Therefore, all coding changes are listed in the next section.
- ***Batrachosuchus browni***: The single brachyopid sampled by Schoch (2013:table 1) was listed as *Batrachosuchus browni*, but the examined material refers to specimen(s) in the

Natural History Museum London (NHMUK); there is only one specimen of *Batrachosuchus browni*, and it is reposited in the South African Museum (SAM). Conversely, *Bathignathus watsoni*, originally placed in *Batrachosuchus* until Damiani & Jeannot (2002), is known from one specimen reposited at the NHMUK, and the OTU was listed in the matrix as '*Batrachosuchus watsoni*'. The listed reference in the analysis of Schoch (2013), Watson (1956), does not resolve this issue, as both species were redescribed in that work (though *Batr. browni* in much less detail). Both historic and recent diagnoses of these taxa (e.g., Welles & Estes, 1969; Damiani & Jeannot, 2002; Schoch & Milner, 2014:98) encompass differences that are not captured by the character sample (e.g., the presence of an interfrontal in *Batr. browni*). Because *Batr. browni* was more recently described and figured (Chernin, 1977) than *Bath. watsoni* (reassigned but not redescribed by Damiani & Jeannot, and last redescribed by Watson, 1956), I opted to use *Batr. browni* (which is also represented by slightly more complete material).

- ***Siderops kehli:*** The skull of *Siderops kehli*, despite being largely complete, is poorly known with respect to the skull roof. Virtually all of the cranial roof sutures are reconstructed (Warren & Hutchinson, 1983:fig. 2), and there is only one known specimen. The poor understanding of its cranial osteology is evidenced by diagnoses that rely primarily on qualitative cranial or postcranial features and an absence of propagated reconstructions compared to less reconstructed taxa (e.g., Schoch & Milner, 2000; Warren & Marsicano, 2000). A reconstruction might be acceptable for coding if it were formed as a composite from multiple specimens. However, because there is only one specimen, the reconstruction of sutures is not really that of *S. kehli* but rather of the concept of a generalized chigutisaurid skull roof in the 1980s; what sutures are known (e.g., paired frontals) are predominantly temnospondyl plesiomorphies. This means that there is no justifiable basis for coding based on the reconstruction compared to other taxa for which a composite from multiple specimens can be made.
- ***Gerrothorax pulcherrimus, Plagiosuchus pustuliferus*:** Among the notable features of plagiosaurids is the single centrum, with intersegmental neural arches. Whether this centrum represents co-ossification of the pleurocentra and the intercentrum (Shishkin, 1987, 1989); an enlarged pleurocentrum and the loss of the intercentrum (Panchen, 1959); or an enlarged intercentrum with loss or fusion of the pleurocentra with the neural arch (Warren & Snell, 1991; Hellrung, 2003) remains unresolved (e.g., Witzmann et al., 2014; Danto et al., 2016, 2017). The first two conditions would be apomorphic among temnospondyls, whereas metoposaurids and perhaps *Laidleria gracilis* share large, dorsally closed intercentra in the absence of pleurocentra. There is further variability within the clade (e.g., Konietzko-Meier et al., 2014, suggested different development between *Gerrothorax* and *Plagiosuchus*). Original codes by Schoch (2013) for characters 163–170 reflect the hypothesis that the centra are strictly intercentra (i.e. the pleurocentra are coded as absent), but because the homology remains an open question based on the recent literature, all characters related to specifically intercentra or pleurocentra (versus a generic 'centra') should not be coded.
    - Characters 163–168 and 170 were coded as unknown (?) for both taxa.
    - Characters 278, 287 were coded as unknown (?) for *Gerrothorax*.
- ***Trematosaurus brauni:*** The osteology of *Trematosaurus brauni* was outdated prior to Schoch's (2019) revised study, so most characters added by Pardo et al. (2017a) were

probably not coded because they deemed the existing literature at the time to be insufficient to do so. Following Schoch's publication, many of these can now be coded.

  - The following characters were newly coded: 216 (0), 217 (0), 218 (0), 219 (1), 220 (0&1), 222 (0), 223 (0), 224 (0), 225 (1), 226 (0), 227 (0), 228 (0), 229 (0), 230 (0), 233 (0), 234 (0), 235 (0), 236 (0), 237 (0), 238 (0), 239 (0), 240 (0), 242 (0), 243 (0), 244 (0), 245 (0), 246 (0), 248 (0), 250 (0), 251 (1), 252 (1), 254 (0), 255 (1), 256 (0), 257 (1), 258 (1), 260 (0), 262 (0), 264 (0), 265 (0), 267 (0), 268 (0), 269 (1), 270 (0), 275 (0), 311 (0&1&2), 312 (0&1), 313 (0), 314 (0), 315 (0), 316 (0), 317 (1), 318 (0), 319 (-), 321 (0), 323 (0), 324 (-), 328 (-), 329 (-), 330 (-), 332 (-), 333 (-), 335 (-), 341 (-), 342 (-). New codes for any character that was completely replaced or revised is noted in the character-specific section (e.g., for 344 [pseudodentary]).
    - With the exception of characters 244, 255–257, 275, 324, 328–330, 332–333, and 341–342, Schoch & Milner's (2000:figs. 71, 77) reconstructions of the lingual surface of the lower jaw and dorsal, ventral, and occipital views of the skull would have been sufficient to confidently code all newly coded characters. The reconstructions of Schoch (2019) are essentially identical to the earlier reconstructions.

- ***Metoposaurus diagnosticus:*** *Metoposaurus diagnosticus* originally comprised two subspecies, *Metoposaurus diagnosticus diagnosticus* and *Metoposaurus diagnosticus krasiejowensis* (Sulej, 2002). This taxonomic framework was utilized until Brusatte et al. (2015) elevated the latter to species rank (*Metoposaurus krasiejowensis*) on the philosophical basis that subspecies are not appropriate for extinct vertebrates, a framework subsequently adopted by other workers. Schoch (2013:table 1) indicated that the osteology of *Metoposaurus diagnosticus* was based solely on a combination of older literature (Fraas, 1889, who described only *M. diagnosticus diagnosticus* in Sulej's framework) and personal observation, which are collectively prohibitive to coding by other workers (evidenced by the large amount of missing data for characters added by Pardo et al., 2017a). I changed this OTU to *Metoposaurus krasiejowensis* because it is more recently and comprehensively documented for the entire skeleton (e.g., Sulej, 2002, 2007). At present, there are no differences between these species that are captured in the sampled characters other than polymorphisms only known in *M. krasiejowensis* (likely due to its much larger sample size), and this change is made with the intent to make the coding more reproducible from the literature.
  - Characters recoded as a result of this taxonomic shift (most of which are newly coded polymorphisms for previous overgeneralized states): 20 (0 → 0&1), 39 (0 → 0&1), 40 (0 → 0&1), 44 (0 → 0&1), 45 (0 → 0&1), 58 (0 → 0&1), 141 (0 → ?), 145 (2 → 1), 146 (1 → 0&1), 150 (1 → 0), 220 (1 → 0&1).
  - Characters newly coded as a result of this taxonomic shift: 214 (0), 255 (1), 257 (1), 260 (0), 261 (0), 262 (0), 263 (1), 264 (0), 265 (0), 266 (0), 268 (0), 269 (1), 270 (1), 271 (0), 272 (1), 273 (-), 274 (0), 275 (0), 276 (1), 279 (0), 280 (0), 281 (0), 282 (0), 283 (0), 284 (1), 285 (0), 286 (0), 287 (0), 288 (1), 289 (1), 290 (2), 291 (2), 292 (2), 296 (1), 299 (0), 300 (-), 301 (1), 302 (1), 304 (1), 305 (0), 306 (-), 307 (0), 318 (0), 321 (0), 322 (1), 323 (0), 324 (-), 326 (1), 327 (0), 328 (-), 329 (-), 330 (-), 332 (-),335 (-), 341 (-), 342 (-), 343 (2), 344 (0).

- ***Eocyclotosaurus wellesi*:** This taxon was described on the basis of numerous skulls and lower jaws by Schoch (2000), but no other material of this taxon is known. However, the OTU was coded for postcranial characters representing nearly the entire postcranial skeleton, most of which would be substantiated if the OTU was changed to the more recently described *Eocyclotosaurus appetolatus* (Rinehart et al., 2015), a study in which Schoch was involved. While Schoch (2000) only indicated material of this taxon from Arizona, Schoch & Milner (2000:150–151) included material of what is now *E. appetolatus* from New Mexico with the species that became *E. wellesi* (as "*Eocyclotosaurus* nov. spec.; this is *E. wellesi* based on the type designation). This OTU is therefore changed to be *E. appetolatus* and was coded based on Rinehart et al. (2015) and Rinehart & Lucas (2016).

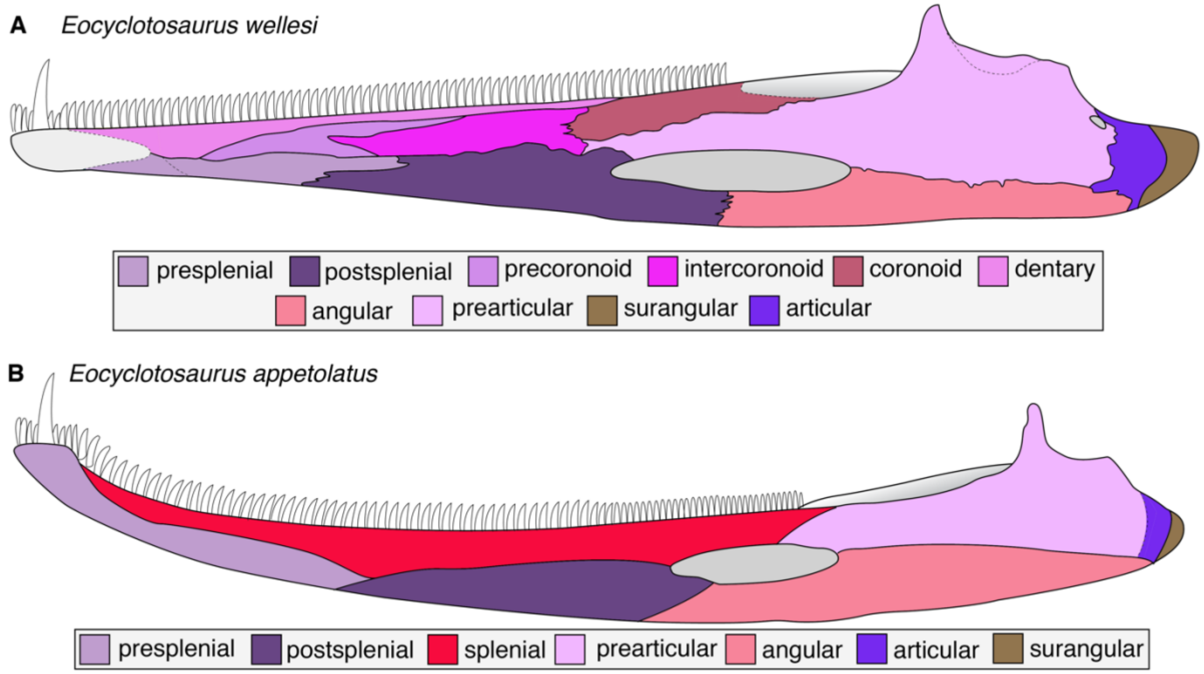

**Figure 31. Comparison of hemimandibular suture interpretations for Eocyclotosaurus. A**, *E. wellesi* (after Schoch, 2000); **B**, E. *appetolatus* (after Rinehart et al., 2015). Coronoid dentition is intentionally omitted for simplicity. Silhouettes not to scale.

One important caveat related to this substitution is the interpretation of the element along the dorsal margin on the lingual surface of the lower jaw (Fig. 31). Rinehart et al. (2015:10, fig. 6) and Rinehart & Lucas (2016:34, fig. 34) identified this as a single, elongate splenial, extending from near the symphysis to the adductor fossa. However, this does not represent a dorsolateral extension of the splenial in the sense of most other workers; this element is normally restricted to the anteroventrolingual surface and was instead identified as the 'presplenial' by Rinehart and colleagues. Because those workers also identified a postsplenial (equivalent to the element of the same name as applied by other workers), this would mean that there are three splenials in *E. appetolatus*. In no early tetrapod (e.g., Ahlberg & Clack, 1998), let alone another temnospondyl, including *E. wellesi* (Schoch, 2000:402, fig. 5), are there three splenials. Furthermore, the 'true' splenial does not extend dorsolaterally based on digital or traditional thin sections (e.g., Carroll, 1964; Chase, 1965; Gee et al., 2019; Gruntmejer et al., 2021; Arbez et al., 2022), and it never extends for most of the lingual surface anterior to the adductor fossa. Although the coronoids of *E. appetolatus* were stated to be barely preserved and without discernible sutures, it seems more likely that the very thin fragments identified as the coronoids are fractured surfaces of the ‘splenial,’ and that the ‘splenial’ is at least one coronoid (but probably three without discernible sutures). The anteroposterior extent of the ‘splenial’ is the same as that of the coronoids in most temnospondyls, and the 10 teeth borne on the ‘splenial’ just anterior to the adductor fossa and parallel to the marginal tooth row in *E. appetolatus* are in the same position as the 13 teeth borne by the posterior coronoid in *E. wellesi* (and similar conditions of other capitosaurs). Therefore, the ‘splenial’ of *E. appetolatus* is treated as a trio of coronoids without discernible sutures.

- ***Cyclotosaurus robustus***: Like *Metoposaurus diagnosticus*, this taxon was almost entirely uncoded for characters added by Pardo et al. (2017a), which reflects the fact that the osteology of this taxon has only been depicted in simple photographs (e.g., Schoch & Milner, 2000:pl. 16) and reconstructions (e.g., Schoch & Milner, 2000:101; Schoch, 2008:fig. 4D) in recent literature. However, Schoch (2013:table 1) cited Sulej & Majer (2005) as the literature source for this taxon. Sulej & Majer described a new species of *Cyclotosaurus* (*Cyclotosaurus intermedius*), which despite some contention that it is synonymous with *Cyclotosaurus robustus* (Lucas et al., 2007), is widely accepted as a distinct taxon by temnospondyl workers (e.g., Schoch, 2008; Witzmann et al., 2016; Marzola et al., 2017). Schoch (2013) also listed personal observation of material at Stuttgart (SMNS), but material of *Cyclotosaurus intermedius* is reposited at the Institute of Paleobiology in Warsaw (ZPAL). Therefore, the OTU is either chimeric (*Cyclotosaurus robustus* + *Cyclotosaurus intermedius*) and was not stated as such, or one of these sources is in error. Because recent literary documentation of *Cyclotosaurus robustus* is poor, I restricted this OTU to *Cyclotosaurus intermedius.*
    - Characters recoded from one state to another as a result of this taxonomic shift: 71 (1 → 0), 81 (1 → 0), 193 (1 → 0).
    - Characters recoded from one state to unknown as a result of this taxonomic shift: 150–153, 160–170, 174, 202, 208, 212.
    - Characters newly coded as a result of this taxonomic shift: 175 (0), 176 (0), 177 (0), 178 (0), 213 (0), 215 (0), 216 (0), 217 (0), 218 (0), 219 (1), 220 (0), 222 (0), 223 (0), 224 (1), 225 (1), 226 (0), 227 (0), 228 (0), 229 (0), 230 (0), 233 (0), 234 (0), 235 (0), 236 (0), 237 (0), 238 (0), 239 (0), 240 (0), 241 (1), 242 (0), 243 (0), 244 (0), 245 (0), 246 (0), 248 (1), 249 (0), 250 (0), 251 (0), 252 (0), 254 (2), 258 (1), 260 (0), 261 (0), 262 (0), 263 (1), 264 (0), 265 (0), 266 (0), 267 (0), 268 (0), 269 (0), 270 (0), 271 (0), 272 (1), 273 (-), 275 (0), 299 (0), 301 (0), 302 (0), 310 (0), 311 (1&2), 312 (1), 313 (0), 314 (0), 315 (0), 316 (1), 317 (0), 318 (0), 319 (-), 321 (0), 323 (0), 324 (0), 326 (1), 335 (-), 344 (0).
- ***Rileymillerus cosgriffi*:** As originally interpreted by Bolt & Chatterjee (2000), this taxon lacks a lacrimal (Fig. 32E-F, 32H-I). It was coded as such (21–1) by Pardo et al. (2017a) but was changed to 21–0 by Schoch et al. (2020; who did not newly code other characters related to the lacrimal that were left as unknown by Pardo et al.). This change is based on the interpretation of the LEP identified by Bolt & Chatterjee. I favor Pardo et al.’s identification of a LEP, but I question the validity of the original interpretation of the nasal in this taxon, as it would invoke an aberrant morphology in which the nasal extends along the lateral edge of the frontal. Crucially, this taxon is only represented by one

cranial specimen with an incomplete snout. The morphology interpreted by Bolt & Chatterjee is not observed in any other temnospondyl regardless of the presence or absence of either an LEP or the lacrimal, and there was no express rationale for the identification of the incomplete element as the nasal (rather than as the lacrimal), nor is there a distinctive aspect of temnospondyl nasals that permits confident identification. Instead, the partial 'nasal' is in the same position as the lacrimal of *Almasaurus habbazi* (Dutuit, 1976:figs. 81, 95; Fig. 32A, 32C), among some other later-diverging stereospondyls in which the lacrimal can be long yet widely separated from the orbit. Most importantly, if the 'nasal' is actually a lacrimal, it would contact the frontal, a condition not found in most other temnospondyls but that would be intriguingly shared with *A. habbazi*, which also has an anteriorly shortened prefrontal that terminates posterior to the level of the anterior orbital margin (also very rare among temnospondyls).

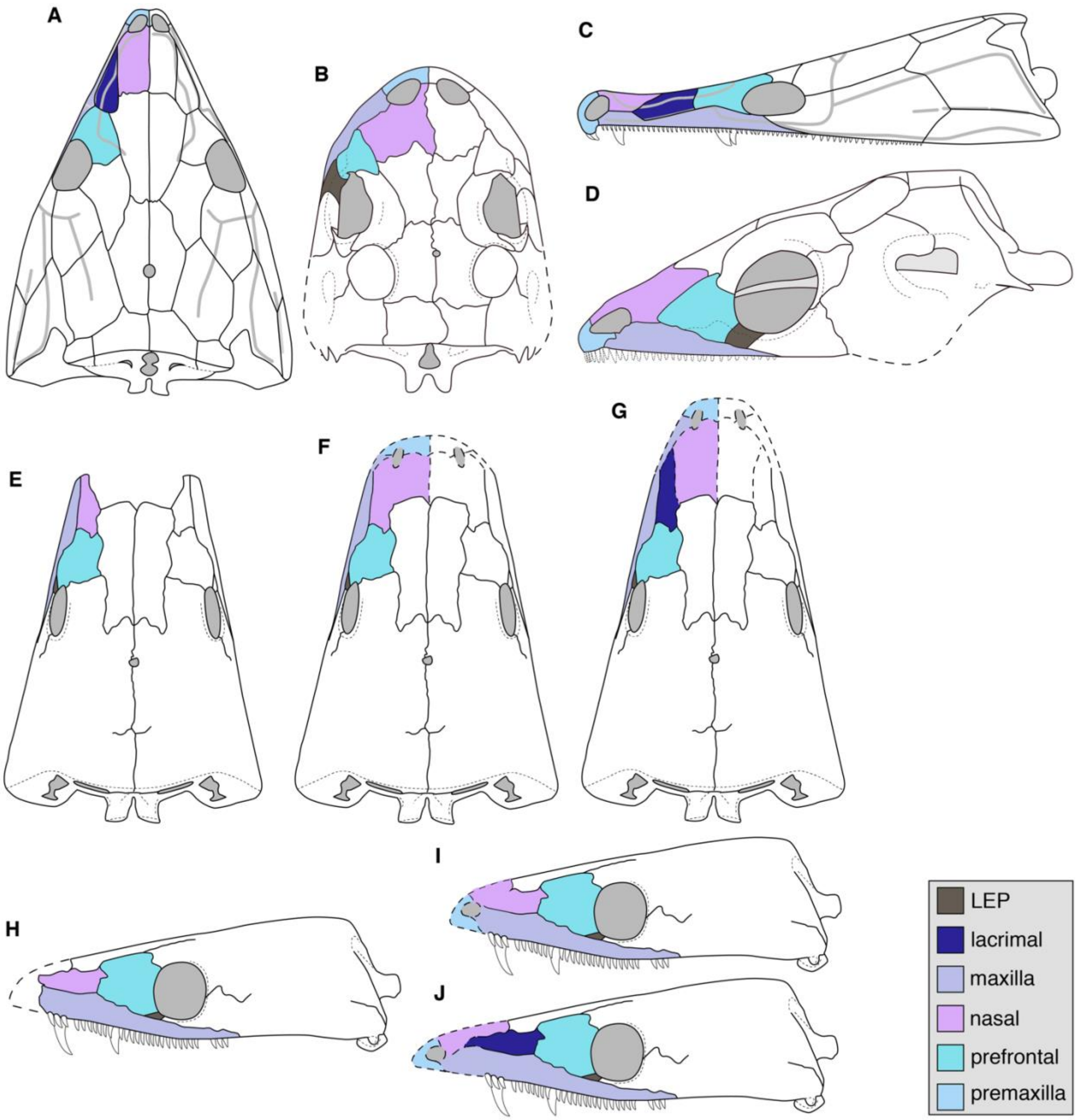

**Figure 32. Comparison of enigmatic small-bodied Late Triassic temnospondyls. A**, skull roof in dorsal view of the putative trematosaur *Almasaurus habbazi* from Morocco (after Dutuit, 1976); **B**, skull roof in dorsal view of the putative stem caecilian *Chinlestegophis jenkinsi* from Colorado (after Pardo et al., 2017); **C**, skull roof of *A. habbazi* in left lateral view (after Dutuit, 1976); **D**, skull roof of *C. jenkinsi* in left lateral view (after Pardo et al., 2017); **E**, original interpretive line drawing in dorsal view of the holotype of *Rileymillerus cosgriffi* (Bolt & Chatterjee, 2000); **F**, reconstruction of the skull of *R. cosgriffi* based on Bolt & Chatterjee's extrapolation of the missing area; **G**, reconstruction of the skull of *R. cosgriffi* based on a reinterpretation of the 'nasal' of this taxon as a lacrimal (contra Bolt & Chatterjee); **H**, original interpretive line drawing in left lateral view of the holotype of *R. cosgriffi* (Bolt & Chatterjee, 2000); **I**, reconstruction of the skull of *R. cosgriffi* based on Bolt & Chatterjee's extrapolation of the missing area (dashed line representing inferred snout margin reproduced from their line drawing); **J**, reconstruction of the skull of *R. cosgriffi* based on a reinterpretation of the 'nasal' of this taxon as a lacrimal (contra Bolt & Chatterjee). Silhouettes not to scale.

Because I do not have positive evidence that the 'nasal' is a lacrimal, I did not code previously unknown characters for the lacrimal or code the presence of a lacrimal (21–0). However, to be symmetrical in reflecting the uncertainty, I also uncoded any character for which the interpretation of the nasal would have to be correct and did not restore the original code of a lacrimal absent. Realistically, this only led to the recoding of a few characters to unknown (?); most nasal characters require the entire element to be known (e.g., 10, 16, 31). This issue will require redress from future workers to more clearly elucidate the relationships of small-bodied stereospondyls, which remain undersampled in most matrices and which may thus form spurious topological clusters due to shared features associated with small adult size (Gee et al, 2021).

  - 20 (0 → ?), 38 (1 → ?), 59 (0 → ?; this was originally coded by Pardo et al. and is an error in their interpretation that the lacrimal is absent)

- ***Hynobius japonicus:*** There is no such species within *Hynobius*, although there is a Japanese hynobiid with the same species epithet, *Onychodactylus japonicus* Houttuyn, 1782. Based on Pardo et al.'s (2017a) analysis, the OTU labeled as '*Hynobius japonicus*' was sourced from Maddin et al. (2012:2), who coded an unspecified species of *Hynobius* from personal observations of one specimen (in the earlier study, it was coded as a terminal "salamanders" OTU). I treated this OTU as a genus-level OTU for *Hynobius*.

Appendix 2.5.2. Partial uncertainty. In PAUP* and Mesquite, partial uncertainty is coded with the use of a slash (/), and this is algorithmically differentiated from polymorphism (indicated with '&'); these are expressed as different syntax as well, with polymorphism represented by parentheses and partial uncertainty by braces. TNT scripts cannot accommodate partial uncertainty, and any such codings are converted to a polymorphism encompassing the possible states (e.g., partial uncertainty of states 0 and 1 becomes a polymorphism of 0 and 1) when Mesquite exports a TNT script. Pardo et al. (2017a), who analyzed the matrix in PAUP*, included several partial uncertainties, although most of these may have been intended as polymorphisms because they are largely for binary characters (except characters 248 and 321). I did not adjust codes to differentiate between intended partial uncertainty and intended polymorphism because this has no effect on the resultant topologies, only on the length of the MPTs. Because these cells are visually distinctive using Mesquite's display option to color cells by character state, I noticed that none of these appear in the matrix of Schoch et al. (2020), in which the corresponding cells were coded as unknown. However, Schoch et al. (2020) did not note changing these codings, and it may be an error in translation between software or scripts (their analysis was conducted in TNT). None of these cells were present in Schoch (2013) and therefore originate in Pardo et al. (2017a). The following codings were restored from Pardo et al.:

- **Character 10:** restored to '0/1' from '?' for *Ichthyophis* and *Epicrionops*.
- **Character 17:** restored to '0/1' from '?' for *Hynobius*.
- **Character 106:** restored to '0/1' from '0' for *Eocaecilia*.
- **Character 160:** restored to '0/1' from '?' for *Triadobatrachus*.
- **Character 246:** restored to '0/1' from '?' for *Cacops*.
- **Character 248:** restored to '1/2' from '?' for *Xenopus*.
- **Character 253:** originally coded as '0/1' by Pardo et al. and changed here to '0&1' for *Gerrothorax* (also noted in the taxon-specific changes) .

- **Character 267:** restored to '0/1' from '?' for *Cochleosaurus*, *Lydekkerina*, *Siderops*, and *Callistomordax*.
- **Character 321:** restored to '0/1' from '?' for *Chinlestegophis*.
- **Character 324:** restored to '0/1' from '?' for *Eryops* and *Gerrothorax*.

Appendix 2.5.3. Taxon-specific changes. As with the above section, taxa are referred to only by the genus and should be assumed to be the species sampled in the matrix (for multi-species genera) unless otherwise stated (e.g., a reference to *Eryops* is specifically to *Eryops megacephalus*).

***Proterogyrinus scheelei***

- **24 (lacrimal, posterior extension):** 0 → ?. Holmes (1984:447, fig. 3) indicated that the suture between the lacrimal and the jugal is entirely unknown, and there is no comment on the anterior extent of the latter in the text. As reconstructed by Holmes, the lacrimal would extend past the antorbital region (24–1).
- **35 (infraorbital sulcus):** 0 → ?. Holmes (1984:440–441) noted that the grooves are poorly developed and only appear on the "the postfrontal, postorbital, the suborbital ramus of the jugal, and on the lateral edge of the nasal." In taxa with a lacrimal, like *Proterogyrinus*, any flexure of the infraorbital sulcus would occur on the lacrimal, which does not have any groove per Holmes' description.
- **57 (jugal, ventral process):** 0 → 1. Holmes (1984:453) stated that "the relation between the posterior end of the ectopterygoid and the jugal is not clear in any of the articulated specimens. However, in CMNH 11067 the right jugal clearly shows a modest processus alaris that would almost certainly have made contact with the pterygoid mesially and excluded the ectopterygoid from the rim of the adductor chamber, as it does in *Eogyrinus* and *Palaeoherpeton*." Although a ventral exposure of the jugal is typically thought of as a stereospondyl feature, it occurs also in *Cochleosaurus* and *Eryops* (and was originally coded as such).
- **58 (jugal, anterior extension):** 0 → ?. As with character 24.
- **59 (jugal-lacrimal):** 0 → ?. As with character 24. While a interceding LEP (or LEE) would be highly unexpected in an embolomere, it cannot be assumed that one was absent because this taxon is an embolomere (circular logic).
- **85 (transvomerine tooth row):** 0 → ?. Holmes (1984:452) stated that there is no identifiable vomer in any specimen, and the only putative fragment of a vomer confers practically no information. Associated taxon-specific changes:
  - **86 (additional vomerine fangs):** 0 → ?. As with character 85. Alternatively, this could be coded as '-' based on Holmes' inference that there were no vomerine fangs because these are often large in early tetrapods, and there was no evidence for such fangs in material of this taxon.
  - **93 (vomer):** 0 → ?. As with character 85.
  - **94 (vomer, paired anterior depressions):** 0 → ?. As with character 85.
  - **96 (vomerine ridges):** 0 → ?. As with character 85.
  - **97 (vomerine pit and fontanelle):** 0 → ?. As with character 85.
  - **98 (vomerine septum):** 0 → ?. As with character 85.
  - **126 (palatine, vomer):** 0 → ?. As with character 85.

- **120 (pterygoid, squamosal):** 0 → -. Holmes (1984:453–454, fig. 4) showed that it is the epipterygoid, not the pterygoid, that sutures to the skull roof. There is therefore neither pterygoid-squamosal contact (120–0) nor an open fissure (120–1).
- **127 (LEP):** 0 → ?. As with character 24.
- **144 (preglenoid process):** 0 → ?. This refers to the dorsal margin of the coronoid process as formed by the surangular (i.e. is it mostly flat or convex); Holmes (1984:460, figs. 14–15) stated that there is a prominent dorsal process and reconstructed a markedly convex margin well above the glenoid (144–1), but the latter morphology is only inferred.
- **145 (Meckelian window):** 0 → 1/2. Holmes (1984:461–462, fig. 15) indicated that this region of the lingual surface is not well-preserved, but it was reconstructed based on partial preservation of a "large" window.
- **151 (stapes, ventral process):** 0 → ?. Following modifications to characters 150 and 153; Holmes (1984) did not describe any stapes, fragmentary or otherwise, and Clack et al. (2016:84) noted that stapes are found in only two embolomeres: *Palaeoherpeton* and *Pholiderpeton*.
- **152 (stapes, shape):** 0 → ?. As with character 151.
- **173 (ribs, uncinate blades):** 0 → -. This character requires the presence of uncinate blades, which are absent.
- **174 (ribs, uncinate spines):** 0 → -. As with character 173.
- **196 (carpals):** 0 → 1. Holmes (1984:481, fig. 28) depicted two ossified distal carpals.
- **218 (postfrontal, shape):** ? → 1. The postfrontal is known (Holmes, 1984: figs. 3, 5, 7) and is more similar in shape to that of *Balanerpeton* (coded for state 1; Milner & Sequeira, 1993:figs. 2–3) than to that of *Greererpeton* (coded for state 0; Smithson, 1982:figs. 5–7).
- **266 (Meckelian fossa):** ? → 0/1. Holmes (1984:461–462) noted one anterior fossa near the anterior end of the prearticular. He tentatively suggested there was a second one more posteriorly, which is typical for early tetrapods, but the degree of confidence expressed is not sufficient to code it exclusively for 266–0 (two fossae).
- **271 (coronoids):** ? → 0/1. Holmes (1984) only identified two coronoids, which would be somewhat unexpected. Because the lingual exposure of the jaw is not well-known, some workers have treated the coronoid count as unknown (e.g., Ahlberg & Clack, 1998; character 4 therein) or as partial uncertainty with at least one known (e.g., Pardo et al., 2017b; character 100 therein), whereas others have coded the presence of three coronoids (e.g., Marjanović & Laurin, 2019; characters 167, 171, 175 therein). The character is left partially unknown to be conservative, but it is felt that Holmes' identification of at least two is sufficient to restrict to '0/1' (compared to '0/1/2' in Pardo et al.).
- **312 (parietal width):** ? → 0. As shown by Holmes (1984:figs. 5–6).

***Greererpeton burkemorani***

- **15 (snout, internarial distance):** 0 → 1. The nares of this taxon are not exposed dorsally in adults (this is an ontogenetic shift; Godfrey, 1989), and consequently, the interorbital distance is much narrower (see also, Smithson, 1982).
- **24 (lacrimal, posterior extent):** 0 → 1. The lacrimal forms a substantive portion of the lateral/ventral orbital margin (e.g., Smithson, 1982:figs. 5, 7, 9; Godfrey, 1989a:fig. 4).

- **45 (prefrontal-maxilla):** 0 → 1. There is a prominent contact between these elements (Romer, 1969:figs. 2–3; Smithson, 1982:44, fig. 9; Godfrey, 1989a:fig. 4).
- **60 (intertemporal):** 0 → 0&1. This is intraspecifically variable, and the intertemporal tends to be absent (60–1) in this taxon (Smithson, 1982:42).
- **62 (squamosal-tabular):** 0 → 1. In the absence of an otic notch, there is a broad contact between these elements.
- **63 (tabular, squamosal):** 0 → 1. Both elements are posterolaterally extensive such that the occipital margin is broadly concave.
- **120 (pterygoid, squamosal):** 0 → -. It is the epipterygoid, not the pterygoid, that sutures to the skull roof. There is therefore neither pterygoid-squamosal contact nor an open fissure (Smithson, 1982:42, 44, fig. 12).
- **157 (branchial denticles):** 0 → ?. This character requires branchial denticles, which are not found in this taxon.
- **196 (carpals):** 0 → 1. Godfrey (198b9:109) made brief mention of ossified carpals.
- **227 (external naris in dorsal view):** ? → 1. The naris is entirely laterally facing.
- **273 (coronoid, teeth, size):** 1 → 2. Bolt & Lombard (2001:1032) stated that all "teeth" were "small"; the largest coronoid "teeth" on the posteriormost coronoid were stated to be intermediate between those anteriorly and the marginal dentition.
- **310 (splenial-coronoids):** ? → 1. As reconstructed by Bolt & Lombard (2001:fig. 5.2), the splenial only contacts the first coronoid and is separated from the second coronoid by a slender anterior process of the prearticular.
- **332 (sphenethmoid, lateral walls, length):** ? → 0. Smithson (1982:52–54, fig. 15) described and figured a relatively long sphenethmoid.
- **340 (shape of optic foramen):** ? → 0. This is a small, circular perforation.
- **342 (sphenethmoid, anterolateral process):** ? → 0. As with character 332.

***Edops craigi***

- **31 (frontal-nasal, length):** 0 → 1. Whether measured along the midline or as the greatest length of the respective elements, the nasal is longer than the frontal (Romer & Witter, 1942:figs. 1–2; Schoch & Milner, 2014:fig. 13A).
- **168 (pleurocentrum, presence):** ? → 0. Romer & Witter (1942:952) described pleurocentra. Note that characters 169 and 170, which refer to specific attributes of the pleurocentra, were originally coded.
- **192 (humerus, entepicondylar foramen):** 0 → 1. A foramen is absent (Romer & Witter, 1942:fig. 13).
- **204 (femur, intercondylar fossa):** ? → 0. Romer & Witter's (1942:fig. 4C) reconstruction depicts a typically deep fossa.
- **214 (skull to trunk):** 1 → ?. Vertebral material of this taxon is isolated, and the length of the trunk is unknown (Romer & Witter, 1942:951–952). This code may have been accidentally entered for this taxon instead of for *Cochleosaurus bohemicus* (one line below), which has a well-characterized skeleton and which was not originally coded.

***Adamanterpeton ohioensis***

- **30 (interorbital distance):** 0 → 1. This region is much wider than the orbit (Milner & Sequeira, 1998:fig. 9).
- **57 (jugal, ventral process):** ? → 1. Milner & Sequeira (1998:269) described a ventral exposure of the jugal that separates the maxilla and the quadratojugal.

- **70 (quadrate, dorsal process):** 0 → ?. There is no description or reconstruction of the quadrate, and this general region of the skull is entirely conjectural (Milner & Sequeira, 1998:fig. 9).
- **71 (posterior skull rim):** 0 → ?. As with character 70; the posteriormost extent of the tabular is also unknown.
- **75 (quadrate and occipital condyles):** 0 → ?. As with character 70; there are no occipital condyles preserved either.
- **76 (epipterygoid):** 0 → ?. There is no description or figuring of an epipterygoid.
- **103 (vomer, anterior part):** 1 → ?. This character is complicated by the slanted anterior margin of the choana, but whether the landmark is placed at the anteromedial corner (next to the vomerine fang) or at the more anteriorly placed anterolateral corner, the reconstruction by Milner & Sequeira (1998:fig. 9) depicts a much shorter pre-choanal region (state 0). However, those same authors list (p. 265 therein) "prefang length of vomer shorter or equal to length of postfang vomer" as a differential feature from *Cochleosaurus florensiensis* (state 0/1). The best way to reconcile this is to leave it as uncoded.
- **118 (pterygoid, exoccipital):** 0 → ?. There is no clear palatal exposure of the exoccipitals documented in this taxon.
- **119 (pterygoid, basioccipital):** 0 → ?. There is no description or figuring of the basioccipital in this taxon.
- **120 (pterygoid, squamosal):** 0 → ?. This condition can only be assessed in non-compressed specimens visible in occipital view (or rendered through CT analysis).
- **121 (pterygoid, flange):** 0 → 1. There is a small flange as described and figured by Milner & Sequeira (1998:276, figs. 2, 10).
- **135 (quadrate trochlea):** 0 → ?. As with character 70.
- **145 (Meckelian window):** 0 → 1. The opening is nearly half as long as the adductor fossa and is much larger than in other taxa coded for state 0, such as *Balanerpeton*.
- **201 (ilium, orientation):** 0 → ?. There is no postcrania associated with this taxon (Milner & Sequeira, 1998:264–265; Schoch & Milner, 2014:21).
- **202 (ilium, tip):** 1 → ?. As with character 201.
- **212 (rib cage):** 0 → ?. As with character 201.
- **243 (number of maxillary teeth):** 1 → 0/1. Milner & Sequeira (1998:277) stated that there is room for "about 30." This ambiguity spans these two states.
- **245 (occipital profile):** 0 → ?. As with character 74 (posttemporal fenestra), this requires an undistorted occiput, which is unknown in this taxon.
- **271 (coronoids):** ? → 0/1/2. At least one coronoid is present.
- **310 (splenial-anterior coronoid):** 0 → ?. While at least one coronoid is clearly present, the sutures between coronoids are not resolved or reconstructed.

***Cochleosaurus bohemicus***

- **121 (pterygoid, flange):** 0 → 1. Sequeira (2004:30, fig. 7) noted a prominent posterolateral flange.
- **192 (humerus, entepicondylar foramen):** 0 → 1. The loss of the foramen is an ontogenetic feature (Sequeira, 2009:147).
- **195 (humerus, torsion):** ? → 1. Sequeira (2009:147) noted a torsion angle of only about 30 degrees.

- **205 (femur, trochanter):** ? → 0. Sequeira (2009:148) noted a small trochanter.
- **214 (skull to trunk):** ? → 1. Reconstructions of the skeleton have been published by Schoch & Milner (2014:fig. 8B).
- **278 (trunk intercentra):** ? → 0. As described by Sequeira (2009:142).
- **292 (atlas neural arch):** ? → 0. As described by Sequeira (2009:142).
- **293 (proatlantes):** ? → 0. As described by Sequeira (2009:142).
- **305 (olecranon process):** ? → 1. As described by Sequeira (2009:147).
- **312 (parietal width):** ? → 1. As with character 311.

***Nigerpeton ricqlesi***

- **29 (pineal foramen):** 0 → 1. Steyer et al. (2006:21) noted the presence of a foramen in the smaller specimen but not in the larger specimen, a typical ontogenetic transformation in edopoids.
- **96 (vomerine ridges):** 1 → 0. The only ridge mentioned by Steyer et al. (2006) borders the choana and is not the same as the more extensive anteromedially radiating ridges captured by 96–1, and Sidor (2013:fig. 3B) figured a vomer without any radiating ridges.
- **103 (vomer, anterior part):** 1 → ?. Neither the anterior nor the posterior margins of the vomer are defined in any specimen (Steyer et al., 2006; Sidor, 2013).
- **104 (vomer, extension):** 0 → ?. As with character 103.
- **139 (dentigerous palatal ossicles):** 0 → 1. As with character 87.
- **140 (postglenoid area):** 1 → ?. The most posterior portion of the mandible that is known is the end of the dentary and part of the angular; this is insufficient to assess the glenoid region, let alone the post-glenoid region or lack thereof.
- **141 (postglenoid area, types):** 0 → ?. As with character 140.
- **144 (preglenoid process):** 0 → ?. There is no indication that the surangular is preserved.
- **145 (Meckelian window):** 0 → ?. There is no description, figuring, or reconstruction of this opening, and the absence of description of the postsplenial, the prearticular, and the angular in lingual view suggests that it is not known.
- **201 (ilium, orientation):** 0 → ?. There is no pelvic material associated with this taxon (Steyer et al., 2006; Sidor, 2013; Schoch & Milner, 2014:21).
- **202 (ilium, tip):** 1 → ?. As with character 201.

***Chenoprosopus milleri***

- **29 (pineal foramen):** 1 → 0. Langston (1953:fig. 5) reconstructed the skull as lacking the pineal foramen, as is common for large edopoids, but Reisz et al. (2005:254, fig. 2) identified this foramen in a larger specimen and noted that the largest specimen used by Langston does not have a well-preserved surface where the pineal would be predicted.
- **39 (prefrontal, frontal):** 0 → 0&1. This is variable within an individual (Langston, 1953:fig. 5).
- **56 (semilunar flange):** 0 → 1. As depicted by Langston (1953:fig. 8).
- **57 (jugal, ventral process):** ? → 0. No ventral exposure has been depicted by previous workers.
- **113 (cultriform process, structure):** 0 → 1/2. Langston (1953:366) described the posterior half as having a “keel-like edge.”
- **121 (pterygoid, flange):** 0 → 1. Langston (1953:365, figs. 7–8) noted a prominent transverse flange projecting into the subtemporal vacuity.
- **178 (clavicle, ventral blade):** ? → 0. As figured by Hook (1993:fig. 2A).

- **191 (humerus, adult shaft):** 1 → ?. No humerus is reported in "*C. lewisi*," and even if one was, it could not be coded for this character, which prescribes the adult condition.
- **204 (femur):** ? → 0. As described by Hook (1993:284).
- **212 (rib cage):** 0 → ?. No ribs are reported for any specimen.
- **247 (jugular foramen):** 0 → 1. Langston (1953:369) stated that nerves IX through XI pass through the exoccipital; nerve X typically passes through the same foramen as the jugular vein.
- **250 (teeth on pterygoid):** ? → 0. There is no description, figuring, or reconstruction of teeth on the pterygoids, only denticles (Langston, 1953).
- **278 (trunk intercentra):** ? → 0. As reported by Hook (1993:282, fig. 3).
- **310 (splenial-coronoid):** 0 → ?. Langston (1953:369) was unable to discern contacts between the splenial and the postsplenial, as well as the margins of the middle coronoid, and Hook (1993:281) notes that the coronoids are entirely obscured in the holotype of "*C. lewisi*."

***Dendrerpeton helogenes***

- There are no additional changes to those made at the character level.

***Balanerpeton woodi***

- There are no additional changes to those made at the character level.

***Trimerorhachis insignis***

- **30 (interorbital distance):** 0 → 0&1. There is intraspecific variation in this feature (e.g., Milner & Schoch, 2013:fig. 2); reconstructions depict an interorbital distance that is wider than the orbit.
- **39 (prefrontal, frontal):** 1 → 0&1. This is polymorphic (e.g., Milner & Schoch, 2013:figs. 1–2).
- **244 (maxillary tooth count):** 1 → 0. Schoch & Milner (2013:106), citing Case (1935:237), noted room for up to 60 maxillary teeth.
- **281 (haemal arches):** ? → 0. Pawley (2007:876) made brief mention of these features.
- **286 (transverse processes):** ? → 0. Pawley (2007:fig. 5) illustrated these processes on the arch.
- **290 (atlas neural arch):** ? → 0. Pawley (2007:875) indicated there is no fusion between components of the atlas.
- **292 (atlas neural arch):** ? → 0/1. Pawley (2007:875) indicated the two counterparts are "not fused" to each other.
- **295 (cervical rib distal shape):** ? → 0. Pawley (2007:876, fig. 4) indicated distally expanded ribs to form a more 'spatulate' shape.
- **302 (deltopectoral crest):** ? → 1/2. Termed the 'deltoid' and 'pectoral' crests by Pawley (2007:881), these are "well-developed" and "rugose."
- **305 (olecranon process):** ? → 1. Pawley (2007:883, fig. 10) described and figured a moderately developed process.
- **307 (femur):** ? → 0. As figured by Pawley (2007:figs. 12–13). This is clearly a subjective feature with 'long' and 'short' states; the coding here is by comparison with previously coded taxa and is at least more elongate than other taxa coded for 307–0 like *Acanthostomatops*.
- **322 (splenial teeth):** 0 → 1. No previous worker has described or figured teeth on the splenial (e.g., Williston, 1913; Case, 1935; Milner & Schoch, 2013).

***Neldasaurus wrightae***

*Except for the change to character 71, all of the coding modifications that are listed here are those that are not the result of novel data / revisions by Schoch (2018) and could have been justified based strictly on the original description by Chase (1965).

- **36 (supraorbital sulcus):** ? → 1. Chase (1965:fig. 1) and Schoch (2018:fig. 1A) depicted the groove passing through the nasal and the prefrontal but not the lacrimal.
- **65 (tabular horn):** 1 → 0. The tabular forms a small, rounded posterolateral projection. This is arguably more developed than in *Trimerorhachis* (coded for 65–0) and is distinct from the condition of *Isodectes* (coded for 65–1) and tupilakosaurids (coded for 65–1) in which the posterior margin of the tabular is squared-off.
- **71 (posterior skull rim):** 0 → 1. As reconstructed by Chase (1965:fig. 3) and Schoch & Milner (2014:fig. 20C), this character was originally coded correctly, but Schoch (2018:fig. 3B) reconstructed the quadrates as being anterior or in line with the tabular horns (depending on whether this refers to the most posterior extent of each element or the general position of the entire body); either results in a correction to 71–1.
- **121 (pterygoid, flange):** 0 → 1. The lateral edge of the palatine ramus features a notable projection into the subtemporal vacuity such that this margin is not smoothly confluent (Schoch, 2018:figs. 2–3).
- **163 (intercentrum, shape):** ? → 0. Schoch (2018:7) described the intercentra as “wedge-shaped.” This is also shown and described by Chase (1965:190–192, fig. 9).
- **165 (intercentrum, ventral surface):** ? → 0. As figured by Chase (1965:fig. 9).
- **166 (intercentrum, anterior surface):** ? → 0. As figured by Chase (1965:fig. 8).
- **167 (parapophysis):** ? → 0. As described by Chase (1965:190) and Schoch (2018:7).
- **171 (ribs, length):** ? → 1. As described by Chase (1965:193–196, fig. 10) and Schoch (2018:7).
- **172 (ribs, ventral extension):** ? → 0. As described by Chase (1965:193–196, fig. 10).
- **174 (ribs, uncinate spines):** 0 → -. There are no uncinate blades (Chase, 1965:fig. 10). Character 173 was updated from unknown to inapplicable for consistency on the same basis.
- **178 (clavicle, ventral blade):** ? → 0. As described by Chase (1965:196–198, fig. 11B) and Schoch (2018:10, fig. 4H).
- **183 (interclavicle, posterior margin):** ? → 0. As described by Chase (1965:197–198).
- **186 (interclavicle, posterior stylus):** ? → 0. As with character 183.
- **188 (scapula, glenoid facet):** ? → 1. As described by Chase (1965:198).
- **189 (scapula, height):** ? → 0. As with character 188.
- **191 (humerus, adult shaft):** ? → 1. As described by Chase (1965:200).
- **192 (humerus, entepicondylar foramen):** ? → 1. No foramen was described or figured by Chase (1965:200, fig. 12).
- **193 (humerus, supinator):** ? → 0. As with character 191.
- **194 (humerus, condyles):** ? → 1. As figured by Chase (1965:fig. 12)
- **195 (humerus, torsion):** ? → 1. As with character 191.
- **196 (carpals):** ? → 0. As described by Chase (1965:200–201, fig.12).
- **197 (manual digit count):** ? → 1. As with character 196.
- **198 (ilium, shaft):** ? → 0. As described by Chase (1965:201–202, fig. 13).
- **199 (ilium, dorsal end):** ? → 0. As with character 198.

- **200 (ilium, height):** ? → 0. As with character 198.
- **203 (pubis):** ? → 0. There is no evidence for a pubis (Chase, 1965:201).
- **205 (femur, trochanter):** ? → 0. Chase (1965:202) noted the presence of a trochanter.
- **271 (coronoids):** ? → 0/1/2. At least one coronoid was identified in thin section (Chase, 1965:186), but it is technically impossible to be certain whether a coronoid identified in a section from a different position is another coronoid.
- **272 (coronoid teeth):** 0 → ?. The coronoid dentition is only identified in cross-section, with three teeth spanning the width. Taxa with coronoid "teeth" rather than denticles typically do not have them in longitudinal rows, and the distribution of dentition is also important for differentiating between 'teeth' and 'denticles.' This is best left as unknown until the coronoids can be in greater detail.
    - **273 (coronoid teeth):** 2 → ?. Following the revision to character 272.
- **296 (ribs anterior to sacrum):** ? → 0. As with character 171.
- **299 (supraglenoid foramen):** ? → 0. As noted by Chase (1965:199).
- **322 (splenial teeth):** ? → 1. Chase (1965) made no mention of splenial teeth in the partially exposed splenial, and none were identified in his thin sections.

***Dvinosaurus primus***

- **57 (jugal, ventral process):** 0 → 1. A short process posterior to the ectopterygoid and contacting the pterygoid is figured by many workers (e.g., Schoch & Milner, 2014:fig. 21B).
- **127 (LEP):** ? → 0. It is unclear why this taxon was originally uncoded for this character. Among dvinosaurs, only tupilakosaurids have a LEP, and the cranial osteology of *D. primus* is well-established.
- **137 (exoccipital condyles):** 0 → 1. Like many other dvinosaurs, the exoccipitals projects well posterior to the dorsal margin of the postparietals and tabulars (e.g., Schoch & Milner, 2014:fig. 21A).

***Acroplous vorax***

- **137 (exoccipital condyles):** 0 → 1. The condyles project far posteriorly in dorsal and lateral view (Englehorn et al., 2008:figs. 1C, 4C, 5C, 7A, 7C).
- **145 (Meckelian window):** 0 → -. There is only one opening in the lingual surface of the lower jaw, and it is unusually anteriorly situated between the dentary, anterior and middle coronoids, splenial, and postsplenial. This is not homologous with the more commonly found posterior Meckelian window that is commonly situated between the postsplenial, the angular, and the prearticular and ventral to the adductor fossa in other temnospondyls.
- **159 (caudal count):** 0 → ?. As with character 158.
- **171 (ribs, length):** ? → 1. Coldiron (1978:15) described the ribs as short (and provided corresponding measurements relative to the neural spine).
- **175 (cleithrum):** 0 → ?. Coldiron (1978:14) stated that only the descending rod is exposed.
- **176 (cleithrum):** 0 → ?. As with character 175.
- **177 (cleithrum):** 0 → ?. As with character 175.
- **191 (humerus, shaft):** ? → 0/1. Englehorn et al. (2008:300) state that this is a stout element.
- **193 (humerus, supinator):** ? → 1. As described by Coldiron (1978:15).
- **195 (humerus, torsion):** ? → 1. As with character 193.

- **198 (ilium, shaft):** ? → 1. As figured by Englehorn et al. (2008:fig. 11B).
- **205 (femur, trochanter):** ? → 0. As described and figured by Englehorn et al. (2008:300, fig. 11A).
- **214 (skull to trunk):** 1 → ?. As with character 158.
- **255 (stapes):** ? → 0. Englehorn et al. (2008:298) described a stapedial foramen.
- **312 (parietal width):** 1 → 0. As shown by Englehorn et al. (2008:fig. 7A).

***Isodectes obtusus***

- **16 (nasal, width):** 1 → 0. As shown by Sequeira (1998:fig. 9). While the nasal appears to be more equant in the holotype of "*Isodectes megalops*" (Sequeira, 1998:fig. 3B), this is a relatively smaller specimen, and the nasal is definitively longer than wide in larger specimens (e.g., holotype of "*Trimerorhachis conangulus*; Sequeira, 1998:fig. 4B). Sequeira (1998:249) notes the elongation of the nasal as an ontogenetic feature.
- **34 (sulci, skull roof):** 1 → 0. Sequeira (1998:252) noted the presence of sulci on the supratemporal (not a circumorbital element), reaffirmed by Schoch & Milner (2014:35), and also noted the possible presence of a discontinuous groove represented by deep pitting along the posterior margin.
- **62 (squamosal-tabular, dorsal):** 0 → 1. These elements contact dorsally (Sequeira, 1998:figs. 1–2, 4, 9).
- **137 (exoccipital condyles):** 0 → 1. The condyles project far posteriorly in dorsal and lateral view (Sequeira, 1998:251, figs. 2, 4, 6, 9).
- **271 (coronoids):** ? → 0/1/2. At least one coronoid was identified by Sequeira (1998:251).
- **272 (coronoid teeth):** 0 → 1. The coronoid dentition is only listed as a patch of denticles.
    - **273 (coronoid teeth):** 2 → -. Following the revision to character 272.

***Thabanchuia oomie***

- **38 (maxilla-nasal):** 0 → ?. The snout is not well-preserved, and Warren (1998b) expressly stated that the sutures between the premaxillae, maxillae, and nasal were unknown. She reconstructed the taxon as having a prefrontal that divides the maxilla and the nasal by entering the naris, whereas Schoch & Milner (2014:fig. 20I) reconstructed this as polymorphic, with a narrow maxilla-nasal contact on the left side. The conservative treatment is to leave it as unknown.
- **40 (prefrontal, anterior end):** 0 → 1. The prefrontal has a transversely oriented, squared-off end that is nearly straight except for a short portion that enters the naris. This is closer to the condition of taxa originally coded for 40–1 (e.g., *Micromelerpeton*) than to the condition of taxa originally coded for 40–0 (e.g., *Neldasaurus*).
- **49 (postorbital):** 0 → 0&1. The right side of the reconstructed skull (Warren, 1998b:figs. 1A, 5A) depicts a postorbital with a posterolateral expansion into the jugal that does expand past the lateralmost extent of the orbit (see also Schoch & Milner, 2014:fig. 20I). This condition is not found on the left side.
- **106 (basicranium, contact):** 0 → 1. Warren (1998b:149, figs. 1B, 5B) note a long interdigitating suture between the pterygoid and the parasphenoid.
- **108 (parasphenoid-exoccipitals):** 0 → ?. Dislodgement of the exoccipitals precludes a confident characterization, but as reconstructed (Warren, 1998b:fig. 5), the parasphenoid

only clearly sutures with the basioccipital posteriorly, as this element forms the entire ventral portion of the condylar surface.

- **123 (pterygoid, ectopterygoid):** 0 → ?. While this is reconstructed by Warren (1998b:fig. 5B), the pterygoid-ectopterygoid suture is entirely conjectural, and there is nothing in the text description to definitively support this coding.
- **124 (pterygoid-palatine-ectopterygoid):** 1 → ?. While this is reconstructed by Warren (1998b:fig. 5B), the palatine-ectopterygoid suture is entirely conjectural, and there is nothing in the text description to definitively support this coding.
- **125 (palatine, ectopterygoid):** 0 → ?. As with character 124.
- **127 (LEP):** ? → 1. This is a strange character to have been left uncoded, as the LEP was well-documented by Warren (1998b:147–148, figs. 1–3, 5) and has been carried forward by other workers (e.g., Schoch & Milner, 2014:fig. 20I).
- **131 (ectopterygoid, length):** 0 → ?. As with character 124.
- **137 (exoccipital condyles):** 0 → 1. The condyles project far posteriorly in dorsal and lateral view (Warren, 1998b:figs. 1A, 5A, 5D).
- **145 (Meckelian window):** 0 → ?. Warren (1998b:151–152, fig. 6) states that most of the lingual surface is obscured, made no comments on any lingual opening, and did not reconstruct one.
- **146 (symphyseal teeth):** 0 → ?. Warren (1998b:152) stated that it is not possible to determine the presence or absence of these teeth. Her reconstruction (fig. 6 therein) reflects an inferred absence, as in many non-stereospondyls, but the more conservative treatment is as unknown.
- **149 (mandibular osteoderms):** 0 → 1. The only osteoderms described by Warren (1998b:153) are "are closely associated with the inner margin of the jaw and extend posteromedially from the anterior end of the angular to terminate before the pectoral girdle," and "positioned as they are, they would afford protection to the branchial area." This appears to meet the criteria for mandibular osteoderms.
- **155 (basibranchial):** ? → 1. Per Warren (1998b:252) and affirmed by Witzmann (2013:151, fig. 4a).
- **159 (caudal count):** 0 → ?. As with the uncoding for character 158: at most 6.5 presacral positions are known. There is also no description or figuring of any caudal positions.
- **167 (parapophysis):** 0 → ?. Warren (1998b:155) noted an inability to find any parapophyses.
- **170 (pleurocentrum, ventral extension):** 0 → 2. Warren (1998b:154–155) noted that the pleurocentra and intercentra are apparently indistinguishable in size and form. While other characters for these can be coded since all centra are identical, state 170–0 (wedged between successive intercentra and not reaching ventral margin of intercentra) is not a valid interpretation for this taxon, as they do reach the ventral margin of the corresponding intercentra.
- **173 (ribs, uncinate blades):** 0 → -. There is no indication of any uncinate processes on the ribs.
- **174 (ribs, uncinate spines):** 0 → -. As with character 173.
- **182 (interclavicle, proportions):** 0 → ?. While Warren's inference of a posterior stylus being present in some form is accepted here, its posterior extent is inherently unknown.

- **185 (interclavicle, anterior stylus):** 0 → ?. As with character 182.
- **187 (interclavicle, clavicles):** 0 → 2. As described and figured by Warren (1998b:155, fig. 7).
- **192 (humerus, entepicondylar foramen):** 1 → ?. No humerus was confidently identified (Warren, 1998b:156) and other humeral characters were originally uncoded.
- **196 (carpals):** 0 → ?. While the lack of carpals may be predicted because dvinosaurs are paedomorphic, limb elements are not even confidently identified (see character 192), and there is not enough of the forelimb preserved, even if all putative limb elements are indeed as much, to be confident (i.e. no elements of the manus are preserved).
- **201 (ilium, orientation):** 0 → ?. No pelvic material is preserved.
- **202 (ilum, tip):** 1 → ?. As with character 201.
- **203 (pubis):** 1 → ?. As with character 201.
- **204 (femur):** 1 → ?. There is no hindlimb material preserved.
- **206 (tarsals):** 0 → ?. As with character 204.

***Tupilakosaurus wetlugensis***

- **16 (nasal, width):** 1 → ?. The snout region is poorly preserved such that the anterior margin is entirely unknown. In the similarly preserved *Thabanchuia*, in which the code for 16–1 is maintained, the nasal is more posteriorly truncated, allowing for a reasonable inference that it was wider than long. Conversely, it is more posteriorly extensive in *Tupilakosaurus* such that previous inferences of a premaxilla-nasal suture (e.g., Shishkin, 1961:fig. 1; Schoch & Milner, 2014:fig. 20H) produce a nearly equant element (i.e. it straddles the division between character states). For this reason, it is felt that treating it as unknown is more conservative.
- **38 (maxilla-nasal):** 0 → 1. There is a broad maxilla-nasal contact reported in all previous descriptions and figures.
- **40 (prefrontal, anterior end):** 0 → 1. Although the anterior margin is not perfectly straight, there is no tapering anterior process, and the overall margin is squared-off.
- **57 (jugal, ventral process):** 0 → 1. A short ventral exposure was figured by Shishkin (1961:fig. 1).
- **71 (posterior skull rim):** 1 → ?. As originally described and figured by Shishkin (1961), the quadrates are not preserved.
- **75 (quadrate and occipital condyles):** 2 → ?. As with character 71.
- **106 (basicranium, contact):** 0 → 1. Shishkin (1961:fig. 2) depicted a long interdigitating suture between the pterygoid and the parasphenoid.
- **124 (pterygoid-palatine-ectopterygoid):** 1 → 0. The pterygoid contacts both elements (Shishkin, 1961).
- **127 (LEP):** ? → 1. As with *Dvinosaurus* and *Thabanchuia*, t is unclear why this character was not coded. Although Shishkin's original interpretation was as the lacrimal, the original description preceded the recognition that the palatine could be laterally exposed. Schoch & Milner's comparative dvinosaur figure (fig. 20I) explicitly colored the LEP in this taxon and in *Thabanchuia* (also originally uncoded for the character) and lists a LEP as the first of eleven diagnostic features of Tupilakosauridae (p. 37 therein); Schoch (2013) also coded both taxa as lacking a lacrimal (21–1) and a LEE (132–0), so it is unclear what alternative element this could be proposed to be.
- **135 (quadrate trochlea):** 0 → ?. As with character 71.

- **137 (exoccipital condyles):** 0 → 1. Like *Thabanchuia* and many other dvinosaurs, the exoccipitals projects well posterior to the dorsal margin of the postparietals and tabulars.
- **159 (caudal count):** 0 → ?. Presacral count (character 158), the reference point for this character, was originally uncoded, so this character should not be coded.

***Acanthostomatops vorax***

- **39 (prefrontal, frontal):** 1 → 0. The prefrontal terminates well anterior to the frontal except in the most immature individuals (Boy, 1989:fig. 1; Witzmann & Schoch, 2006:figs. 2–3).
- **75 (quadrate and occipital condyles):** 1 → -. There are no ossified occipital condyles.
- **76 (epipterygoid):** 0 → -. Witzmann & Schoch (2006:373) indicated the absence of an ossified epipterygoid.
- **121 (pterygoid, flange):** 0 → 1. Boy (1989:fig. 2) and Witzmann & Schoch (2006:fig. 4) figured pterygoids with a prominent posterolateral expansion of the palatine ramus into the subtemporal fenestra.
- **145 (Meckelian window):** 0 → 1. Witzmann & Schoch (2006:374) estimated the length of the posterior Meckelian window to be one-sixth of the length of the lower jaw, which is certainly much larger than the smaller openings found in taxa like *Balanerpeton* or *Trimerorhachis* (145–0).
- **247 (Jugular foramen):** ? → -. As with character 73.
- **266 (Meckelian fossae):** ? → 0/1. As with character 145. This is expressed as partial uncertainty because the anteromedial surface is not as well-preserved (Boy, 1989:fig. 3B), so the presence of additional fossae is unclear.
- **271 (coronoids):** ? → 0/1/2. At least one coronoid was identified by Boy (1989:fig. 3B).
- **275 (jaw sculpture):** ? → 0. The angular is well-ornamented on the ventral and labial surfaces.
- **305 (olecranon process):** ? → 0. Witzmann & Schoch (2006:378) stated that an olecranon process is absent.

***Zatrachys serratus***

- **11 (prenarial region):** 0 → 1. As also seen in the larger zatracheid *Dasyceps* (not sampled here; Paton, 1975:figs. 1, 5), the prenarial region is at least as long if not longer than the frontal (contrary to *Acanthostomatops* and to most other temnospondyls) (Langston, 1953:fig. 13; Schoch, 1997:fig. 2).
- **24 (lacrimal, posterior extension):** 0 → 1. As shown by Langston (1953:fig. 13) and Schoch (1997:fig. 1), the lacrimal forms a moderate portion of the lateral orbital margin.
- **39 (prefrontal, frontal):** 1 → 0. The prefrontal terminates well anterior to the frontal (Schoch, 1997:fig. 2).
- **102 (vomer-pterygoid):** 0 → 0&1. Schoch (1997:233, fig. 3) noted a specimen in which the elements are separated, but other specimens appear to feature this contact (Langston, 1953:fig. 13) .
- **121 (pterygoid, flange):** 0 → 1. Langston (1953:fig. 13) and Schoch (1997:fig. 3) figured pterygoids with a prominent posterolateral expansion of the palatine ramus into the subtemporal fenestra.
- **149 (mandibular osteoderms):** 0 → 1. Schoch (1997:fig. 4) depicted "ventral osteoderms" on the palate of a skull, which Schoch & Rubidge (2005:510) cited as the same gular osteoderms that are found in *Micropholis*.

- **153 (stapes, curvature):** ? → 0. Langston (1953:388) stated that this bone is long and curved.
- **202 (ilium, tip):** 1 → ?. Postcrania are entirely unknown for this taxon, as reflected in other postcranial codes (Langston, 1953; Schoch & Milner, 2014).
- **214 (skull to trunk):** 0 → ?. As with character 202.
- **258 (pterygoids contact anteriorly):** ? → 1. Langston (1953:fig. 13) and Schoch (1997:fig. 3) figured wide separation.
- **260 (dentary):** ? → 0. As per Langston (1953:386–388, fig. 15).
- **261 (dentary, coronoid process):** ? → 0. As with character 260.
- **262 (surangular, normal):** ? → 0. As with character 260.
- **264 (number of splenials):** ? → 0. As with character 260.
- **266 (Meckelian fossae):** ? → 0. Langston (1953:387) identified two Meckelian foramina, one within the first third of the length of the postsplenial, the posterior Meckelian foramen bordered by the angular and the prearticular, and another "smaller opening" adjacent to the postsplenial-angular suture.
- **269 (articulation to tooth row):** ? → 0. As with character 260.
- **271 (number of coronoids):** ? → 0. As with character 260.
- **275 (jaw sculpture):** ? → 0. As with character 260.
- **312 (parietal width):** ? → 1. As shown by Langston (1953:fig. 13) and Schoch (1997:figs. 1–2).
- **314 (prefrontals):** ? → 0. As with character 312.
- **315 (postfrontals):** ? → 0. As with character 312.
- **317 (zygokrotaphy):** ? → 0. As with character 312.
- **345 (dorsolateral osteoderms fused to ribs):** 0 → ?. As with character 202.

***Capetus palustris***

- **56 (semilunar flange):** 0 → 1. The reconstruction by Sequeira & Milner (1993:fig. 9C) depicts this ventral flange.
- **57 (jugal, ventral process):** 0 → ?. The palate is only exposed in one specimen, in which the ectopterygoids are entirely unknown. Because this element would articulate with a ventral exposure of the jugal, the character is best left as unknown since most of the palate has been reconstructed based simply on the plesiomorphic condition.
- **88 (ectopterygoid, fangs):** 0 → ?. Sequeira & Milner (1993:665) stated that the ectopterygoid is not exposed in any specimen. Other associated changes:
    - **123 (pterygoid, ectopterygoid):** 0 → ?
    - **124 (pterygoid-palatine-ectopterygoid):** 0 → ?
    - **125 (palatine, ectopterygoid):** 0 → ?
    - **128 (palatine, ectopterygoid (ontogeny)):** 0 → ?
    - **129 (palatine, ectopterygoid (width)):** 1 → ?
    - **130 (palatine, ectopterygoid (continuous tooth row)):** 0 → ?
    - **131 (ectopterygoid (length)):** 1 → ?
    - **133 (ectopterygoid (Y-shaped)):** 0 → ?
- **94 (vomer, paired anterior depressions):** 0 → ?. Sequeira & Milner (1993:665), citing Broili (1908), mentioned previous identification of paired vomerine depressions but did not identify such depressions in the material available to them and urged caution in interpretation of this feature.

- **95 (anterior palatal depression):** 0 → ?. As with character 94.
- **126 (palatine, vomer):** 0 → ?. It is not clear that there is a palatine-vomer suture (Sequeira & Milner, 1993:figs. 7, 9B).
- **142 (postglenoid area, dorsal):** 0 → ?. The dorsal and lingual surfaces are not well-exposed in any specimen (Sequeira & Milner, 1993:665, figs. 7–8), and there is very little information on the mandible in general.
- **145 (Meckelian window):** 0 → ?. As with character 142.
- **169 (pleurocentrum, lateral surface):** 0 → ?. There is no mention of axial material in this taxon (Sequeira & Milner, 1993:660). Notably, character 168 (presence/absence of pleurocentra) was originally uncoded.
- **170 (pleurocentrum, ventral extension):** 0 → ?. As with character 169.
- **202 (ilium, tip):** 1 → ?. There is no pelvic material known for this taxon (Sequeira & Milner, 1993:660).
- **207 (gastral squamation):** 0 → ?. The only dermal scales are from posterior to the skull and are not specified as being from the ventral region of the skeleton (Sequeira & Milner, 1993:670).
- **212 (ribcage):** 0 → ?. There are no ribs preserved (Sequeira & Milner, 1993:660).
- **225 (septomaxilla):** ? → 0. Sequeira & Milner (1993:661) describe a dislodged septomaxilla.
- **243 (maxillary tooth count):** 1 → 0. Sequeira & Milner (1993:670) estimated 38 maxillary positions.
- **249 (denticles on pterygoid):** ? → 0. Sequeira & Milner (1993:670) described this feature.
- **250 (teeth on pterygoid):** ? → 0. As with character 249.
- **251 (denticles on vomer):** ? → 0. As with character 249.
- **254 (cultriform process contact with vomer):** 0 → ?. This contact is entirely reconstructed (Sequeira & Milner, 1993:66, fig. 9B).
- **258 (pterygoids contact anteriorly):** 1 → ?. As with character 254; this region is entirely reconstructed.
- **310 (dorsal margin of splenial only contacts first coronoid):** 0 → ?. There is no information on the first coronoid.

***Iberospondylus schultzei***

- **9 (preorbital region, length):** 1 → 0. In no specimen is the preorbital length more than twice that of the posterior skull table (Laurin & Soler-Gijón, 2001:fig. 1A; Laurin & Soler-Gijón, 2006:fig. 1A).
- **15 (snout, internarial distance):** 1 → ?. The internarial distance is definitively wider (15–1) than the interorbital distance in a small referred specimen (Laurin & Soler-Gijón, 2006:fig. 1A), but in the larger holotype (Laurin & Soler-Gijón, 2001:fig. 1A), the medial edge of the left naris slightly closer to the midline than the medial edge of the left orbit (this condition is not captured by the character). Because the left premaxilla is slightly disarticulated, this character is changed to unknown.
- **31 (frontal-nasal, length):** 0 → 1. The frontal is shorter (albeit slightly) in the referred specimens (Laurin & Soler-Gijón, 2006:285) and appears as much in the holotype (Laurin & Soler-Gijón, 2001:fig. 1A).

- **39 (prefrontal-frontal):** 0 → 1. The prefrontal extends well past the frontal in the larger of two referred specimens but not in the holotype, which is larger and in which it ends at the same level as the frontal (Laurin & Soler-Gijón, 2001:fig. 1A; Laurin & Soler-Gijón, 2006:fig. 1A).
- **46 (posterior skull table, length):** 0 → 1. Even when measured in the most liberal fashion (posterior orbital margin to posteriormost extent of the tabular), the posterior skull table is less than 0.5 times as long as it is wide.
- **70 (quadrate, dorsal process):** 0 → 1. This feature is specifically called out in previous descriptions, in which it is suggested that it may be present outside of dissorophoids (Laurin & Soler-Gijón, 2001:497; Laurin & Soler-Gijón, 2006:287).
- **105 (basipterygoid ramus):** ? → 1. As figured by Laurin & Soler-Gijón (2001: figs. 1–2) and described by and Laurin & Soler-Gijón (2006:288).
- **109 (basicranium, carotids):** ? → 0. As described by Laurin & Soler-Gijón (2006:289).
- **121 (pterygoid, flange):** 0 → 1. As figured by Laurin & Soler-Gijón (2001: figs. 1–2) and described by and Laurin & Soler-Gijón (2006:288)
- **136 (occipital condyle):** ? → 1. As described by Laurin & Soler-Gijón (2006:289).
- **162 (neural spine, height):** 0 → 1. As described and figured by Laurin & Soler-Gijón (2006:290, fig. 9).
- **175 (cleithrum):** 0 → ?. There is no pectoral girdle material.
- **176 (cleithrum):** 0 → ?. As with character 175.
- **177 (cleithrum):** 0 → ?. As with character 175.
- **178 (clavicle, ventral blade):** 0 → ?. As with character 175.
- **202 (ilum, tip):** 1 → ?. There is no pelvic girdle material.
- **243 (number of maxillary teeth):** 1 → 0. Laurin & Soler-Gijón (2006:285) noted 30 positions in the holotype.
- **274 (symphysis):** 1 → 0. As described by Laurin & Soler-Gijón (2006:290).
- **282 (haemal arches):** 1 → ?. There is no description of the contact with the intercentra, and the one figured specimen with these arches does not appear to preserve intercentra with the haemal arches (Laurin & Soler-Gijón, 2006:figs. 9–10).
- **287 (atlas-axis intercentra):** ? → 0. As described and figured by Laurin & Soler-Gijón (2006:290, figs. 6, 9).
- **292 (atlas neural arch):** ? → 0. As described by Laurin & Soler-Gijón (2006:290).
- **293 (proatlantes):** 1 → 0. Laurin & Soler-Gijón (2006:290) describe articulation facets on the atlantal neural arch for the proatlas.
- **310 (dorsal margin of splenial):** 0 → ?. The distinction between the first and second coronoids is unknown (Laurin & Soler-Gijón, 2006:fig. 8B).

***Micromelerpeton credneri***

- **39 (prefrontal-frontal):** 0 → 0&1. This is variable within a single individual (Boy, 1995:fig. 2B; Schoch & Milner, 2014:fig. 25D).
- **121 (pterygoid, flange):** 1 → 0&1. This is polymorphic (Boy, 1995:fig. 3).
- **131 (ectopterygoid length):** 0 → 0&1. This is polymorphic (Boy, 1995:fig. 3).
- **250 (teeth on pterygoid):** ? → 0. Boy (1995) did not make any mention of teeth on the pterygoid (or any other part of the palate), nor did he reconstruct this type of dentition or indicate it in figures.

- **251 (denticles on vomers):** ? → 0. Boy (1995:435, fig. 3) stated that all palatal surfaces are covered in denticles.
- **252 (denticles on palatines):** ? → 0. As with character 251.
- **253 (denticles on parasphenoid):** ? → 0. As with character 251.
- **258 (pterygoids contact anteriorly):** ? → 1. Boy (1995:fig. 3) figured wide separation.
- **271 (number of coronoids):** ? → 0/1. Boy (1995:fig. 4) identified as least two coronoids; there is a possible greatly truncated first coronoid (anterior coronoid) near the symphysis.
- **312 (parietal, width):** ? → 0. As shown by Boy (1995) among others.
- **314 (prefrontals):** ? → 0. As shown by Boy (1995) among others.
- **315 (postfrontals):** ? → 0. As with character 314.
- **317 (zygokrotaphy):** ? → 0. As with character 311.

***Limnogyrinus elegans***

- **71 (posterior skull rim):** 1 → ?. Despite reconstructing the quadrates, there is no mention of these elements in the description by Schoch & Witzmann (2018) nor any figures showing them in a given specimen.
- **75 (quadrate and occipital condyles):** 1 → ?. As with characters 71 and 73.
- **88 (ectopterygoid, fangs):** 0 → ?. Schoch & Witzmann (2018:298) stated that the dentition of the ectopterygoid is unknown.
- **91 (anterior palatal opening(s)):** 0 → ?. Schoch & Witzmann (2018:298) stated that the anterior region of the vomer is unknown in any specimen due to articulation with the lower jaws; the premaxilla-vomer suture is entirely reconstructed.
- **92 (anterior palatal opening(s)):** 0 → ?. As with character 91.
- **121 (pterygoid, flange):** 0 → 0&1. As reconstructed by Schoch & Witzmann (2018:fig. 3F), the right pterygoid has a more prominent posterolateral projection from the palatine ramus, which does not have a confluent posterior margin, in contrast to the left pterygoid in which the margin is smoothly confluent.
- **126 (palatine, vomer):** 0 → ?. Schoch & Witzmann (2018:298) stated that the suture is unclear.
- **127 (LEP):** ? → 0. As reconstructed by Schoch & Witzmann (2018:fig. 1).
- **131 (ectopterygoid, length):** 0 → 1. Schoch & Witzmann (2018:298) stated that the ectopterygoid is probably only about half the length of the palatine.
- **155 (basibranchial):** ? → 1. As reported by Schoch & Witzmann (2018:298–299, fig. 3B).
- **166 (intercentrum, anterior surface):** 0 → ?. As noted by Schoch & Witzmann (2018:300), the central elements do not ossify; note that characters 163–165 were originally uncoded.
- **167 (parapophysis):** 0 → ?. As with character 166.
- **168 (pleurocentrum, presence):** 0 → 1. As with character 166.
- **169 (pleurocentrum, lateral surface):** 0 → ?. As with character 166.
- **188 (scapula, glenoid):** 1 → -. Schoch & Witzmann (2018:300) noted that the scapulocoracoid remains unossified. In the original character sampling of Schoch (2013), this character could have been used as a proxy for an entirely unossified scapulocoracoid, but such a character exists in the expanded sample (character 301), and the absence of a

glenoid in taxa with at least a partially ossified scapula/scapulocoracoid is definitively not homologous with the absence of a glenoid in taxa without any such ossification.

- **189 (scapula, height):** 0 → -. As with character 188.
- **228 (posterior skull margin):** 0 → 1/2. As reconstructed by Schoch & Witzmann (2018:fig. 1E-F), the margin is convex in smaller specimens and either straight or barely convex in larger ones. Note that this differs from previous reconstructions (e.g., Schoch & Milner, 2014:fig. 25A).
- **237 (postparietal length):** 0 → 1. This taxon has very long and slender postparietals (Schoch & Witzmann, 2018:figs. 1–2).
- **246 (supraoccipital):** ? → 0. Schoch & Witzmann (2018:300) stated that no part of the braincase is ossified.
- **249 (denticles on pterygoid):** ? → 1. As described by Schoch & Witzmann (2018: 298), there is a loose row of teeth on the anterior part of the palatine ramus that are of comparable size to those in *Apateon* and larger than other micromelerpetids. This dentition is therefore treated as 'teeth,' not as 'denticles,' and since there are no other smaller forms of dentition on the pterygoid, this feature is coded as such.
- **250 (teeth on denticles):** ? → 1. As with character 249 (*Apateon pedestris* was originally coded as 250–1).
- **251 (denticles on vomer):** ? → 1. Similar to the pterygoid, there are patches of teeth that were not treated as denticles in character 81. This is followed here.
- **252 (denticles on palatine):** ? → 1. As with characters 249 and 251; these were described as "teeth," not as "denticles" by Schoch & Witzmann (2018:298).
- **255 (stapes):** ? → 0. As described by Schoch & Witzmann (2018:300).
- **256 (pleurosphenoid):** ? → 0. As with character 246.
- **257 (sphenethmoid):** ? → 1. As with character 246.
- **258 (pterygoids contact anteriorly):** ? → 1. As figured by Schoch & Witzmann (2018:fig. 3B).
- **260 (dentary):** ? → 0. As with character 258.
- **274 (symphysis):** ? → 0. As with character 258.
- **278 (trunk intercentra):** ? → 1. As with character 166.
- **296 (ribs anterior to sacrum):** ? → 0. As figured by Schoch & Witzmann (2018:fig. 4I-J).
- **301 (scapulocoracoid ossification):** ? → 2. As with character 188.
- **307 (femur):** ? → 0. As described and figured by Schoch & Witzmann (2018:300, fig. 5).
- **312 (parietal, width):** ? → 0. As described and figured by Werneburg (1994, 2012), Schoch & Milner (2014), and Schoch & Witzmann (2018).
- **313 (skull table):** ? → 0. As with character 312.
- **315 (prefrontals):** ? → 0. As with character 312.
- **316 (postfrontals):** ? → 0. As with character 312.
- **317 (zygokrotaphy):** ? → 0. As with character 312.
- **323 (choana, size):** ? → 0. As with character 318.

***Branchierpeton amblystomum***

- **71 (posterior skull rim):** 1 → ?. It is not clear that the quadrates are known for this taxon. Reconstructions of the skull in dorsal view do not clearly show it, and no recent worker has reconstructed it in ventral view. Werneburg (1991) neither figured nor described the quadrate in this taxon. Note that character 70 (quadrate, dorsal process) was originally left as unknown for this taxon.

- **75 (quadrate and occipital condyles):** 1 → ?. As with characters 71 and 73.
- **88 (ectopterygoid, fangs):** 0 → 1. Werneburg (1991:92) only described a long row of small teeth, and his reconstruction (fig. 9b) only shows palatine ‘fangs,’ not ectopterygoid ‘fangs.’
- **121 (pterygoid, flange):** 0 → 1. There is a prominent posterolateral projection of the palatine ramus.
- **130 (palatine, ectopterygoid, continuous tooth row):** 0 → 1. As described and reconstructed by Werneburg (1991).
- **135 (quadrate trochlea):** 0 → ?. As with character 71.
- **166 (intercentrum anterior surface):** 0 → ?. Characters 163–165 were originally uncoded (all pertain to the intercentrum), which follows Werneburg (1991:92) in making no express identification of any centra.
- **167 (parapophysis):** 0 → ?. As with character 166.
- **168 (pleurocentrum):** 0 → 1. As per Werneburg (1991). Note also that character 170 (pleurocentrum, ventral extension) was originally uncoded.
- **169 (pleurocentrum, lateral surface):** 0 → -. As with character 168.
- **173 (ribs, uncinate blades):** 0 → - There is no description or figuring of uncinate blades.
- **174 (ribs, uncinate spines):** 0 → - As with character 173.
- **188 (scapula, glenoid facet):** 1→ ?. The most recent revision of Werneburg (1991) made no mention of the scapula, let alone of the glenoid.
- **189 (scapula, height):** 0 → ?. As with character 188.
- **199 (ilium, dorsal end):** ? → 0&1. Werneburg (1991:fig. 16) depicted similarly sized ilia with differing conditions.

***Branchiosaurus salamandroides***

- **40 (prefrontal, anterior end):** 0 → 1. Schoch & Milner’s reconstruction depicts an anteriorly blunted prefrontal. The condition is at least polymorphic in Boy’s reconstruction and trends towards only state 40–1. A ‘wide, blunted’ anterior margin is also seen in Werneburg’s (2012:fig. 3) specimen illustrations.
- **50 (postorbital, postfrontal):** 0 → -. The postfrontal is the longest element of the four specified here, while the postorbital is the shortest.
- **71 (posterior skull rim):** 1 → ?. It is not clear that the quadrates are known for this taxon. Reconstructions of the skull in dorsal view do not clearly show it, and no recent worker has reconstructed it in ventral view. Werneburg (2012) neither figured nor described the quadrate in this taxon. Note that character 70 (quadrate, dorsal process) was originally uncoded for this taxon.
- **75 (quadrate and occipital condyles):** 1 → ?. As with characters 71 and 73.
- **85 (transverse tooth row, transvomerine):** 0 → ?. The vomer is not confidently identified in any specimen (Werneburg, 2012), and what little tentatively identified portion there is does not confer substantive information.
- **86 (additional vomerine fangs):** 0 → ?. As with character 85.
- **88 (ectopterygoid, fangs):** 0 → ?. The ectopterygoid was not positively identified.

- **91 (anterior palatal openings):** 0 → ?. As with character 85.
- **92 (anterior palatal openings):** 0 → ?. As with character 85.
- **93 (vomer):** 1 → ?. As with character 85.
- **94 (vomer, paired anterior depressions):** 0 → ?. As with character 85.
- **95 (anterior palatal depression):** 0 → ?. As with character 85.
- **96 (vomerine ridges):** 0 → ?. As with character 85.
- **97 (vomerine pit and fontanelle):** 0 → ?. As with character 85.
- **98 (vomerine septum):** 0 → ?. As with character 85.
- **123 (pterygoid, ectopterygoid):** 0 → ?. As with character 88.
- **124 (pterygoid-palatine-ectopterygoid):** 0 → ?. As with character 88.
- **125 (palatine, ectopterygoid):** 0 → ?. As with character 88.
- **126 (palatine, vomer):** 0 → ?. As with character 81.
- **128 (palatine, ectopterygoid, ontogeny):** 0 → ?. As with character 88.
- **129 (palatine, ectopterygoid, width):** 0 → ?. As with character 88.
- **130 (palatine, ectopterygoid, continuous tooth row):** 0 → ?. As with character 88.
- **131 (ectopterygoid, length):** 0 → ?. As with character 88.
- **132 (LEE):** 0 → ?. As with character 88.
- **133 (ectopterygoid, Y-shaped**): 0 → ?. As with character 88.
- **135 (quadrate trochlea):** 0 → ?. As with character 71.
- **166 (intercentrum anterior surface):** 0 → ?. Characters 163–165 were originally uncoded (all pertain to the intercentrum), which follows Werneburg (2012:12) that centra are not ossified in any specimen.
- **167 (parapophysis):** 0 → ?. As with character 166.
- **168 (pleurocentrum):** 0 → 1. As per Werneburg (2012). Note also that character 170 (pleurocentrum, ventral extension) was originally uncoded.
- **169 (pleurocentrum, lateral surface):** 0 → -. As with character 168.
- **173 (ribs, uncinate blades):** 0 → -. There is no description or figuring of uncinate blades.
- **174 (ribs, uncinate spines):** 0 → -. As with character 173.
- **190 (humerus, ontogeny):** 2 → 1/2. By comparison with other taxa originally coded for 190–2, it is debatable whether the humerus is always a "long rod from small stages on." Werneburg (2012:12, fig. 32 caption therein) described the element as stocky and "short and robust" and noted that length was variable based on quality of preservation. Partial uncertainty accounts for this.

***Apateon pedestris***

- **39 (prefrontal, frontal):** 1 → 0. This taxon has been repeatedly illustrated with a prefrontal well anterior to the frontal (e.g., Schoch & Milner, 2008:fig. 3D), which separates it from other species that have in-line anterior margins (e.g., *A. caudacus*).
- **121 (pterygoid, flange):** 1 → 0. There is no posterolateral process from the palatine or quadrate ramus of the pterygoid (e.g., Schoch & Milner, 2014:fig. 27G).
- **165 (intercentrum, ventral surface):** 0 → ?. Branchiosaurids almost never ossify the intercentra (e.g., Schoch & Fröbisch, 2006; Marjanović & Laurin, 2008; Fröbisch & Schoch, 2009), and it is peculiar that the two preceding characters for intercentrum shape and intercentrum width were originally left as unknown. Therefore, the character is uncoded.

    - **166 (intercentrum, anterior surface):** 0 → ?. As with character 165.
- ***Micropholis stowi***
    - **31 (frontal-nasal, length):** 0 → 0&1. This intraspecific variation is associated with the two morphs (broad- and slender-headed) that are noted by Schoch & Rubidge (2005:506).
    - **62 (squamosal-tabular, dorsal):** 1 → 0&1. This variation is noted to occur on at least the dorsal surface of the roof (which is what the character refers to) by Schoch & Rubidge (2005:507).
    - **65 (tabular, horn):** 0 → 0&1. Schoch & Rubidge (2005:507) noted the absence of a tabular horn in at least one large specimen.
    - **86 (additional vomerine fangs):** 1 → 0&1. Schoch & Rubidge (2005:507) noted this as a variable feature between morphs; 86–0 (only one fang pair) occurs in the slender-headed morph, while 86–1 (three pairs) occurs in the broad-headed morph (see also their comparative reconstructions in figure 2 therein).
    - **88 (ectopterygoid, fangs):** 1 → 0. Schoch & Rubidge (2005:507) stated that "the ectopterygoid has one fang-pair that is smaller than the palatine one," which is reflected in their reconstruction (fig. 2 therein); this might also be polymorphic (it is not reconstructed for the broad-headed morph, but they note that the ectopterygoid is not well-known from this morph).
    - **110 (parasphenoid plate):** 2 → 1&2. This is a polymorphic condition differing between morphs (Schoch & Rubidge, 2005:fig. 2). The slender-headed morph has a proportionately narrower basal plate.
    - **124 (pterygoid-palatine-ectopterygoid):** 1 → 0. Schoch & Rubidge (2005:508, fig. 2) noted substantial contact of the pterygoid with the palatine in the slender-headed morph. They indicated that the contact in the broad-headed morph was likely "faint," as in their reconstruction, which is why this is corrected to 124–0 rather than as polymorphic.
    - **149 (mandibular osteoderms):** 0 → 1. Schoch & Rubidge (2005:509–510) noted the presence of "oval, polygonal, or irregularly shaped ossicles emplaced between the basal plate of the parasphenoid and the ventral elements of the dermal pectoral girdle." They also indicate that these are "true osteoderms and not dislocated tooth plates" and term them as "osteoderms of the throat region."
    - **205 (femur, trochanter):** 1 → 0. Schoch & Rubidge (2005:513) noted "a prominent, boss-like internal trochanter" on the femur, although this is not clear from their single-profile illustration. They compared it favorably to *Eoscopus*, which has a trochanter like other non-branchiosaurid amphibamiforms (e.g., *Doleserpeton*) based on Daly's (1994:fig. 15) illustrations and description.
    - **224 (frontal-orbit):** 0 → 1. Entry of the frontal was documented by Boy (1985:fig. 2) and Schoch & Rubidge (2005:figs. 1–3).
    - **242 (number of premaxillary teeth):** ? → 0/1. Schoch & Rubidge (2005:509) stated that there is room for 8–10 premaxillary teeth.
    - **243 (number of maxillary teeth):** ? → 0. Schoch & Rubidge (2005:509) stated that there is room for 31–42 maxillary teeth (variation between morphs).
    - **271 (coronoids):** 0 → 0/1/2. Only the posterior coronoid has been positively identified (Schoch & Rubidge, 2005:509).

***Platyrhinops lyelli***

- **30 (interorbital distance):** 0 → 1. The interorbital region is unusually wide compared to other amphibamiforms, and the orbits are likewise relatively small and laterally situated as a result (Clack& Milner, 2010).
- **100 (choana, medial):** 1 → 0. The choanae form elongate ovals without any broadening anteromedially (Clack & Milner, 2009:figs. 5, 9).
- **110 (parasphenoid plate):** 2 → 1. The condition of this taxon (Clack & Milner, 2009) is much closer to taxa coded for 110–1, such as olsoniforms and zatracheids, than to amphibamids and salientians (coded for 110–2).
- **250 (teeth on pterygoid):** ? → 0. Clack & Milner (2009) did not make any mention of teeth on the pterygoid (or any other part of the palate), nor do they reconstruct it or indicate it in figures showing the ventral surface of the pterygoid.
- **251 (denticles on vomers):** ? → 0. Clack & Milner (2009:288) stated all palatal surfaces are covered in denticles.
- **252 (denticles on palatines):** ? → 0. As with character 251.
- **253 (denticles on parasphenoid):** ? → 0. As with character 251.
- **264 (splenials):** 2 → 0. Clack & Milner (2009:286) described both the splenial and the postsplenial, as with other temnospondyls.
- **271 (coronoids):** ? → 0. Clack & Milner (2010:fig. 6a) figured the lingual view of one jaw with three demarcated coronoids.

***Doleserpeton annectens***

- **143 (hamate process):** 0 → ?. The lower jaws of this taxon are very poorly known because of the frequent disarticulation of material, which is further complicated by the presence of several amphibamiforms at the type locality. Previous workers have barely described or figured any part of the lower jaw (e.g., Bolt, 1969:889; Bolt, 1977b; Sigurdsen & Bolt, 2010:1366, fig. 3C). The character is uncoded here.
- **156 (hypobranchials):** 0 → 1. Witzmann's (2013:153) review of the hyobranchial apparatus cited Sigurdsen & Bolt (2010) for evidence of ossified hypobranchials.
- **252 (denticles on palatine):** 0 → 1. This taxon has neither a pair of fangs nor any denticles; Sigurdsen & Bolt (2010:1365) stated that "a shagreen of denticles covers the vomers, the lateral shelf of the pterygoids, and a diamond-shaped field at the base of the cultriform process of the parasphenoid" without mention of the palatines.
- **266 (number of Meckelian fossae):** 0 → 0/1. Sigurdsen & Bolt (2010:1366) only described the posteriorly situated large opening; there is no reference to additional qualifying openings, but their figures are insufficient to determine whether smaller foramina might be present anteriorly within the postsplenial or splenial.
- **271 (number of coronoids):** 0 → 0/1. As with character 148; at least two coronoids have been identified.
- **346 (ectopterygoid):** This character, added by Schoch et al. (2020), is a simple presence/absence character for the ectopterygoid. Although originally coded for absent (346–1), as noted by Schoch (2019a:148), whether an ectopterygoid is lost or merely very reduced in *D. annectens* is uncertain; the small size of specimens and any possible ectopterygoid greatly complicates this. The most recent study of *D. annectens* (Sigurdsen & Bolt, 2010:1365) state that "the ectopterygoid is either absent or greatly reduced." To be conservative, this code is changed to unknown. Associated coding changes (88–133, not inclusive):
    - **88 (ectopterygoid, fangs):** 1 → ?

- **124 (pterygoid-palatine-ectopterygoid):** 1 → ?
  - **125 (palatine, ectopterygoid):** 0 → ?
  - **128 (palatine, ectopterygoid (ontogeny)):** 0 → ?
  - **129 (palatine, ectopterygoid (width)):** 1 → ?
  - **130 (palatine, ectopterygoid (continuous tooth row)):** 0 → ?
  - **131 (ectopterygoid (length)):** 1 → ?
  - **132 (LEE):** 0 → ?. Although it could be argued that there is definitively no exposure of this element on the dorsal skull roof and that the code should be maintained, the treatment would be different if the ectopterygoid was definitively absent (in which case this character should be coded as inapplicable).
  - **133 (ectopterygoid (Y-shaped)):** 0 → ?

***Amphibamus grandiceps***

- **143 (hamate process):** 0 → ?. The lower jaws of this taxon are very poorly known because of the typical articulation with the skull and the overall compression of Mazon Creek specimens. Previous workers have barely described or figured any part of the lower jaw (e.g., Carroll, 1964:232–233, fig. 22; Bolt, 1979; Milner, 1982; Daly, 1994). The character is uncoded here.
- **317 (zygokrotaphy):** ? → 0. As with character 42 regarding documentation of the skull roof configuration.

***Gerobatrachus hottoni***

- **3 (ornament, elements):** 0 → ?. The only known specimen is only exposed ventrally (Anderson et al., 2008:515).
- **4 (ornament, snout):** 0 → ?. As with character 3.
- **5 (ornament, general):** 0 → ?. As with character 3.
- **6 (ornament, intensive growth):** 0 → ?. As with character 3.
- **7 (ornament, preorbital):** 0 → ?. As with character 3.
- **24 (lacrimal, posterior extension):** 0 → ?. The posterior terminus of the lacrimal and the lateral margins of the orbit are not preserved (Anderson et al., 2008:fig. 2).
- **27 (orbit margins):** 1 → ?. As with character 3.
- **39 (prefrontal-frontal):** ? → 1. The prefrontal's anterior margin is sufficiently defined to code this character (as with character 40, which was originally coded).
- **55 (supratympanic flange):** 0 → ?. As with character 3.
- **58 (jugal, anterior extension):** 0 → ?. The lateral orbital margins are not defined (Anderson et al., 2008:fig. 2).
- **59 (jugal-lacrimal):** 1 → ?. As with character 58.
- **73 (postparietal-exoccipital):** 1 → ?. The parabasisphenoid is barely preserved and only at the basicranial articulation, let alone posteriorly (Anderson et al., 2008:fig. 2), and there was no mention of the exoccipitals.
- **75 (quadrate and occipital condyles):** 1 → ?. As with character 73.
- **86 (additional vomerine fangs):** 0 → -. There are no vomerine fangs.
- **108 (parasphenoid):** 0 → ?. As with character 73.
- **118 (pterygoid, exoccipital):** 0 → ?. As with character 73.
- **119 (pterygoid, basioccipital):** 0 → ?. As with character 73; no basioccipital is identified or described.

- **134 (palate structure):** 0 → ?. The specimen is completely flattened and cannot be observed in undistorted occipital view.
- **136 (occipital condyle):** 1 → ?. As with character 73.
- **137 (exoccipital condyles):** 0 → ?. As with character 73.
- **138 (basioccipital, length):** 1 → ?. As with characters 73 and 119.
- **142 (postglenoid area, dorsal):** 0 → ?. This refers to the dorsal surface, which is obscured due to articulation with the compressed skull.
- **188 (scapula, glenoid facet):** 0 → ?. There is no scapula preserved in the holotype. Character 189 (scapula, height) was originally left as unknown.
- **216 (supratemporal):** 1 → 0. A supratemporal was figured.
- **230 (postorbital):** ? → 0. A postorbital is present.
- **243 (number of maxillary teeth):** 0 → ?. In contrast to the premaxillary tooth count, no maxillary count or estimate was given. The tooth row appears incomplete posteriorly (markedly so), and if it were interpreted as being nearly complete, it is clearly much shorter than the premaxilla and would thus contain fewer than 21 positions. However, it is left as unknown to be conservative.
- **245 (occipital profile):** 0 → ?. As with character 134.
- **266 (Meckelian fossae):** ? → 0/1. There is at least oen small opening posteriorly between the prearticular and the angular.

***Dissorophus multicinctus***

- **75 (quadrate and occipital condyles):** 2 → 1. Schoch (2012:fig. 3D) reconstructed this taxon for 75–2, but the most recent description of this taxon by DeMar (1968:fig. 2B) depicted a condition closer to 75–1, which is what Dilkes (2020:22) advocated for in a reanalysis of a dissorophid-focused matrix.
- **271 (coronoids):** ? → 0/1/2. At least one coronoid has been identified (DeMar, 1968).

***Cacops aspidephorus***

*Note that these changes are considered to be applicable regardless of whether only *C. aspidephorus* or a composite of *C. aspidephorus* + *C. morrisi* is employed. This includes data that would have only been available to all previous authors of this matrix viz. *C. morrisi* and that has been substantiated in *C. aspidephorus* by Anderson et al. (2020).

- **27 (orbit margins):** 1 → 0. All three species of *Cacops* have prominent orbital rims (Reisz et al., 2009:fig. 2; Fröbisch & Reisz, 2012:fig. 2; Anderson et al., 2020:5, fig. 2A).
- **237 (postparietal, proportions):** 0 → 1. The foreshortened postparietal, forming a transversely elongate rectangle, is a feature of cacopines (e.g., Reisz et al., 2009; Schoch, 2012).
- **264 (splenials):** 1 → 0. A pre- and postsplenial were described by Anderson (2005).
- **270 (angular, extent in lateral view):** ? → 0. The labial side of the lower jaw of *C. morrisi* was figured by Reisz et al. (2009); the same condition was confirmed in *C. aspidephorus* by Anderson et al.(2020).
- **271 (coronoids):** ? → 0. Anderson (2005) identified three coronoids.
- **274 (symphysis):** ? → 1. As described by Anderson (2005).
- **275 (jaw sculpture):** ? → 0. As figured by Williston (1910:pl. 8) and Reisz et al. (2009:fig. 1b).
- **278 (trunk intercentra):** ? → 0. As noted by Williston (1910:261).

- **279 (trunk neural arch to centrum):** ? → 0. As noted by Williston (1910:261–262, pl. 9), Dilkes & Brown (2007:401, figs. 4–5), and Dilkes (2009:1014, fig. 1).
- **281 (haemal arches):** ? → 0. As noted by Williston (1910:265, pl. 9).
- **282 (haemal arches):** ? → 0. As with character 281.
- **283 (haemal arches):** ? → 0. As with character 281.
- **286 (transverse process):** ? → 0. As noted by Williston (1910:262, pl. 9).
- **287 (atlas-axis intercentra):** ? → 0. Williston (1910:262–263) described intercentra of both positions.
- **288 (atlas anterior centrum):** ? → 1. As with character 287.
- **289 (atlas centrum):** ? → 1. As with character 287.
- **290 (atlas neural arch):** ? → 1/2. Whether the arch is co-ossified or merely tightly sutured with the centrum is unclear in light of the preservation of most material of this taxon.
- **292 (atlas, neural arch):** ? → 0. The arch forms two separate prongs that are separated medially by an anterior expansion of the axial neural arch (Williston, 1910:263, pl. 14).
- **293 (proatlantes):** ? → 1. The completeness with which the skeleton is known confers confidence in the absence of proatlantes.
- **294 (second cervical arch):** ? → 0. As noted by Williston (1910:263).
- **299 (supraglenoid foramen):** ? → 0. As noted by Williston (1910:268).
- **301 (scapulocoracoid ossification):** ? → 0. As described by Williston (1910:268–269).
- **302 (deltopectoral crest):** ? → 1/2. As figured by Williston (1910:pl. 11).
- **303 (humerus, length):** ? → 0. As reconstructed by Williston (1910:pls. 9, 11, 17).
- **304 (radius:humerus):** ? → 0. As reconstructed by Williston (1910:pls. 14, 17).
- **305 (olecranon process):** ? → 1. As described by Williston (1910:274).
- **307 (femur):** ? → 0. As figured by Williston (1910:pl. 13).
- **312 (parietal, width):** ? → 0. As figured by Reisz et al. (2009:fig. 3)
- **313 (skull table):** ? → 0. As figured by Williston (1910) and Reisz et al. (2009), among others.
- **314 (prefrontals):** ? → 0. As with character 312.
- **315 (postfrontals):** ? → 0. As with character 312.
- **317 (zygokrotaphy):** ? → 0. As with character 313.
- **318 (quadrate-maxilla):** ? → 0. As with character 312.
- **322 (splenial, teeth):** ? → 0. Anderson (2005) did not describe any teeth on the splenial, and as this does not occur in any temnospondyl, the absence of description can be taken as evidence for an absence of the feature (he also noted coronoid dentition).
- **323 (choana, size):** ? → 0. As with character 313.
- **324 (sphenethmoid, dorsal exposure):** ? → 0. As figured by Williston (1910:255, pls. 1–2).
- **326 (dentary, anterior teeth, biscupidity):** ? → 0. As figured by Williston (1910:pl. 3).

***Broiliellus texensis***

- **312 (parietal width):** ? → 0. As figured by Williston (1914:fig. 1) and reproduced by Schoch (2012:fig. 1E).
- **313 (skull table):** ? → 0. As with character 311.
- **314 (prefrontals):** ? → 0. As with character 311.

- **315 (postfrontals):** ? → 0. As with character 311.
- **317 (zygokrotaphy):** ? → 0. As with character 311.

***Acheloma cumminsi***

*Note that these changes are considered to be applicable regardless of whether *A. cumminsi* (sensu Polley & Reisz, 2011) or ‘*A. dunni*’ (sensu Polley & Reisz, 2011) is employed.

- **24 (lacrimal, posterior extent):** 0 → 1. The lacrimal forms part of the lateral/ventral orbital margin.
- **37 (maxilla, anterior margin):** 0 → 1. Lateral swelling in this region is characteristic of this taxon due to its caniniform dentition (e.g., Williston, 1909:fig. 1; Dilkes & Reisz, 1987:figs. 2A, 3A; Polley & Reisz, 2011:fig. 1).
- **59 (jugal-lacrimal):** 1 → 0. Contrary to other dissorophoids with an LEP, the lacrimal and jugal meet because the LEP (and LEE) are separated from the orbit; this is an autapomorphic condition among all temnospondyls.
- **214 (skull to trunk):** ? → 0. Williston’s (1909:fig. 6) reconstruction is based on the holotype of ‘*Trematops milleri*’, which has an essentially complete presacral column and skull.
- **220 (lacrimal-orbit):** ? → 1. This contact is well-documented (Olson, 1941:figs. 2A, 6; Dilkes & Reisz, 1987:fig. 3; Polley & Reisz, 2011:figs. 1–5).
- **242 (premaxilla, tooth count):** 1 → 0&1. Polley & Reisz (2011:794) reported 13 premaxillary positions, whereas Dilkes & Reisz (1987) reported nine positions.
- **249 (denticles on pterygoid):** ? → 0. As reported by Polley & Reisz (2011:803).
- **251 (denticles on vomers):** ? → 0. Polley & Reisz (2011:801) stated that the vomer is covered in denticles.
- **252 (denticles on palatines):** ? → 0. Polley & Reisz (2011:fig. 1) did not comment on the presence or absence of denticles in the text but did figure a loose covering.
- **253 (denticles on parasphenoid):** ? → 0. Polley & Reisz (2011:804) stated that this is present in both the holotypes of *Acheloma cumminsi* and ‘*Acheloma dunni*’.
- **266 (Meckelian fossae):** ? → 0/1. There is at least the typical large posterior Meckelian foramen (Olson, 1941:167), but the lingual surface has rarely been described or figured. The specimen described by Gee et al. (2019) only appears to have this single foramen but was a juvenile referred to cf. *Acheloma* and is not incorporated here.
- **271 (coronoids):** ? → 0/1/2. At least one coronoid has been identified (Polley & Reisz, 2011).
- **287 (atlas-axis intercentra):** ? → 0. Williston (1909:645–646) and Polley & Reisz (2011:805) described intercentra of both positions.
- **288 (atlas anterior centrum):** ? → 1. As with character 287.
- **289 (atlas centrum):** ? → 1. As with character 287.
- **290 (atlas neural arch):** ? → 1/2. Whether the arch is co-ossified or merely tightly sutured with the centrum is unclear.
- **292 (atlas, neural arch):** ? → 0. The arch forms two separate prongs (Polley & Reisz, 2011:fig. 12C).
- **293 (proatlantes):** ? → 1. The completeness with which the skeleton is known confers confidence in the absence of proatlantes.
- **305 (olecranon process):** ? → 1. This process was documented in the most mature specimens by Olson (1941:168).

- **309 (number of distal tarsals):** ? → 1. The number of tarsals was documented by Dilkes (2014).

***Phonerpeton pricei***

- **37 (maxilla, anterior margin):** 0 → 1. Lateral swelling in this region is characteristic of this taxon due to its caniniform dentition (Dilkes, 1990:224, fig. 1A-B).
- **179 (interclavicle, length):** 0 → ?. There is no documented interclavicle for this taxon (Dilkes, 1990), and other interclavicle characters (180–187) were not originally coded.
- **220 (lacrimal-orbit):** ? → 1. This contact is well-documented (Dilkes, 1990:figs. 1, 3–4).
- **251 (denticles on vomers):** ? → 0. Dilkes (1990:231, fig. 1B) stated all palatal surfaces are covered in denticles.
- **252 (denticles on palatines):** ? → 0. As with character 251.
- **266 (Meckelian fossae):** ? → 0. As described and figured by Dilkes (1990:234:figs. 6–7).
- **271 (coronoids):** ? → 0. As with character 266.
- **305 (olecranon process):** ? → 1. This process is documented by Dilkes (1990:237).

***Ecolsonia cutlerensis***

- **31 (frontal-nasal, length):** 0 → 1. Whether measured as the median suture or as the greatest length, the nasal is longer than the frontal in the only specimen in which these elements are complete (Berman et al., 1985:fig. 5).
- **37 (maxilla, anterior margin):** 0 → 1. Because of the presence of caniniform dentition, the maxilla bows slightly anteriorly (Berman et al., 1985:fig. 5; Schoch & Milner, 2014:fig. 32E).
- **142 (postglenoid area, dorsal):** 0 → ?. The only known lower jaw is only exposed medially, posterolaterally, and ventrally (Berman et al., 1985:16–17, fig. 6).
- **146 (symphyseal teeth):** 0 → ?. As with character 142.
- **152 (stapes, shape):** ? → 1/2. The stapes is preserved in articulation, but only the distal end is exposed projecting through the otic notch (Berman et al., 1985:fig. 5); this is considered sufficient to discern that it was not “robust” or “tetrahedral” (152–0).
- **159 (caudal count):** 0 → ?. Character 158 (presacral count) was originally uncoded, and Berman et al. (1985:17) did not provide an estimate because of incompleteness of the column and the intermingling of material of at least two specimens that made definitive association impossible. They did note at least 12 caudal positions, which is still much less than the typical presacral count in trematopids (e.g., 24 in *Acheloma cumminsi*; Dilkes & Reisz, 1987:5).
- **163 (intercentrum, shape):** 0 → 1. This taxon is distinctive among trematopids for having so-called “ring intercentra” in the presacral region (Berman et al., 1985:18, fig. 9F-G). Alternatively, it could be coded as polymorphic given that the absence of a complete articulated column precludes confident determination of the last presacral.
- **164 (intercentrum, width):** 0 → 1. The notochordal canal is tiny in this taxon (Berman et al., 1985:18, fig. 9F-G).
- **165 (intercentrum, ventral surface):** 0 → 1. In ventral view, the ventral surface is practically a square (Berman et al., 1985:fig. 9F-G).
- **169 (pleurocentrum, lateral surface):** 0 → ?. As figured (Berman et al., 1985:figs. 9F-G, 10B), the pleurocentra have a much smaller lateral surface (169–1), but because there

are no figured corresponding sets of centra from the same position or the same individual, the character is best left as unknown.

- **179 (interclavicle, length):** ? → 0. A skull and corresponding interclavicle are figured by Berman et al. (1985:fig. 5).
- **196 (carpals):** 1 → ?. There is no description of carpals.
- **243 (number of maxillary teeth):** ? → 0/1. Berman et al. (1985:12) estimated room for 27 to 30 maxillary teeth.
- **310 (dorsal margin of splenial only contacts first coronoid):** ? → 1. As shown by Berman et al. (1985:fig. 6).
- **314 (prefrontals):** ? → 0. Prefrontals are present.
- **315 (postfrontals):** ? → 0. Postfrontals are present.
- **322 (splenial teeth):** ? → 1. The splenial is fully exposed lingually, and there is no description or figuring of teeth, as shown by Berman et al. (1985:fig. 6).

***Onchiodon labyrinthicus***

- **121 (pterygoid, flange):** 0 → 1. A prominent flange is found in large individuals (Boy, 1990:fig. 3B).
- **243 (maxillary tooth count):** 2 → 0. Boy (1990:296) stated that there is room for 30–34 teeth on the maxilla.

***Eryops megacephalus***

- **49 (postorbital, lateral process):** 0 → 0&1. As depicted by Sawin (1941:fig. 1), there is a slender lateral process from the postorbital that extends past the lateral margin of the orbit on at least one side of the skull. This could also be coded strictly for state 1.
- **121 (pterygoid, flange):** 0 → 1. A prominent flange is found in large individuals (Sawin, 1941:pl. 2).
- **251 (vomer, denticles):** ? → 0. Sawin (1941:415) stated that denticles cover the palate except for the parasphenoid and were not figured in his reconstruction (pl. 2 therein).
- **252 (palatine, denticles):** ? → 0. As with character 251.
- **271 (coronoids):** ? → 0. The typical suite of three coronoids has been identified (Sawin, 1941).
- **287 (atlas-axis intercentra):** ? → 0. Moulton (1974:26–27; figs. 1, 12) described these ossifications.
- **288 (atlas anterior centrum):** ? → 1. As with character 287.
- **289 (atlas centrum):** ? → 1. As with character 287.
- **290 (atlas neural arch):** ? → 0. As with character 287.
- **292 (atlas neural arch):** ? → 0&2. Moulton (1974:26–27) noted that the arches may be separate or co-ossified.
- **293 (proatlantes):** ? → 0. Moulton (1974:27) noted the presence of the proatlas.
- **305 (olecranon process):** ? → 1. A well-developed process was documented by Miner (1925:figs. 7, 21) and Pawley & Warren (2006:572–573, fig. 7.7–7.10).

***Sclerocephalus haeuseri***

- **33 (lateral line sulci, adults):** 0 → 0&1. Schoch & Witzmann (2009a:146) noted specimens from L–O 6 that have lateral line grooves but also mention a specimen from L–O 6 that was first noted by Boy (1988) to lack grooves.
- **121 (pterygoid, flange):** 0 → 1. A prominent flange comparable to the transverse flange reported in *Archegosaurus* is found in this taxon (Schoch & Witzmann, 2009a:fig. 4B).

- **156 (hypobranchials):** 0 → 1. Witzmann (2013:155, fig. 5F) identified a pair of elements as hypobranchials.
- **159 (caudal count):** 0 → 1. Schoch & Witzmann (2009a:153) only provided a caudal count for one specimen, but it is from L–O 6 and has 27 caudal positions, compared to 24–25 presacral positions.
- **251 (denticles on vomers):** 1 → 0. Schoch & Witzmann (2009a:148) stated that all palatal bones (exclusive of the parasphenoid) are covered in denticles.
- **252 (denticles on palatines):** 1 → 0. As with character 251.
- **254 (cultriform process-vomer):** 1 → 0. The process is anteriorly extensive and has a long lateral contact with the vomers that separates them for over half of their total midlength.
- **281 (haemal arches):** ? → 0. Schoch & Witzmann (2009a:153, fig. 7E) noted these features.
- **282 (haemal arches):** ? → 0. As with character 281.
- **283 (haemal arches):** ? → 0. As with character 281.
- **284 (haemal arch shape):** ? → 0. As with character 281.
- **287 (atlas-axis intercentra):** ? → 0. Schoch & Witzmann (2009a:151, fig. 7) described and figured most of this complex.
- **288 (atlas anterior centrum):** ? → 1. As with character 287.
- **290 (atlas neural arch):** ? → 0. As with character 287.
- **292 (atlas neural arch):** ? → 0/1. As with character 287; Schoch & Witzmann stated that they were "at best loosely attached."
- **294 (second cervical arch):** ? → 0. As figured by Schoch & Witzmann (2009a:fig. 7A-B).
- **295 (cervical rib distal shape):** ? → 0. As figured by Schoch & Witzmann (2009a:fig. 7C).
- **296 (ribs anterior to sacrum):** ? → 0. As with character 295.
- **298 (number of caudal rib pairs):** ? → 0. Schoch & Witzmann (2009a:153) noted the presence of six pairs.
- **299 (supraglenoid foramen):** ? → 0. As described and figured by Boy (1988:122, fig. 8).
- **301 (scapulocoracoid ossification):** ? → 0. As with character 300.
- **302 (deltopectoral crest):** ? → 2. Schoch & Witzmann (2009a:154, fig. 8A-C) depicted a well-developed thickened crest, which is better developed than in *Glanochthon* (previously coded for 302–1).
- **306 (basale commune):** ? → 0. As determined by figures of the pes (Schoch & Witzmann, 2009a:fig. 8E).
- **307 (femur):** ? → 1. As described by Schoch & Witzmann (2009a:157).
- **308 (elongate tibiale and fibulare):** ? → 0. As with character 306.

***Glanochthon latirostris***

- **37 (maxilla, anterior margin):** 0 → 1. Schoch & Witzmann (2009b:12) listed "locally differentiated teeth in the snout, with larger teeth producing lateral excursions of margin (*Intasuchus*, *Sclerocephalus*)" as a feature shared between *Glanochthon* and the aforementioned stereospondylomorphs.

- **121 (pterygoid, flange):** 0 → 1. A prominent flange comparable to the transverse flange reported in *Archegosaurus* is found in this taxon (Boy, 1993:fig. 3; Schoch & Witzmann, 2009b:fig. 4D).
- **271 (coronoids):** ? → 0/1/2. At least one coronoid was identified by Boy (1993), but the sutures are unclear.

***Archegosaurus decheni***

- **15 (internarial distance):** 1 → 0. This condition is ontogenetically variable (Witzmann, 2005:134), with state 15–1 found in juveniles (e.g., Witzmann, 2005:fig. 2a), but 15–0 occurs in adults (Witzmann, 2005:fig. 5a).
- **39 (prefrontal, frontal):** 0 → 0&2. This is variable within a single individual (e.g., Witzmann, 2005:fig. 15).
- **85 (transverse tooth row, transvomerine):** 1 → 0. Witzmann (2005:145) only noted the presence of a tooth row along the medial edge of the choana, which may be continuous with, but is not equivalent to, the transvomerine row. Witzmann (2005)'s 'type B' palate (fig. 17B therein) does have a trio of teeth near the midline, but this is not a row, and the slight anteromedial angling of the parachoanal teeth also does not form a row; even if either could be considered a 'row', this character would have to be coded as polymorphic to include 'absence.'
- **88 (ectopterygoid fangs):** 0 → 1. Witzmann (2005:146) stated that no fangs are present on the ectopterygoids.
- **121 (pterygoid, flange):** 0 → 1. A prominent flange is found in large individuals (Witzmann, 2005:146, fig. 17).

***Platyoposaurus stuckenbergi***

- **15 (snout, internarial distance):** 1 → 0. As figured by Schoch & Milner (2000, figs. 47–48).
- **39 (prefrontal-frontal):** 0 → 2. As figured by Schoch & Milner (2000, figs. 47–48).
- **104 (vomer, extension):** 0 → 1. As figured by Schoch & Milner (2000, figs. 47–48).
- **121 (pterygoid, flange):** 0 → 1. As figured by Schoch & Milner (2000, figs. 47–48).
- **123 (pterygoid, ectopterygoid):** 0 → 1. As figured by Schoch & Milner (2000, figs. 47–48).
- **312 (parietal width):** 1 → 0. As figured by Schoch & Milner (2000, figs. 47–48).

***Australerpeton cosgriffi***

- **15 (snout, internarial distance):** 1 → 0. As figured by Barberena (1998:fig. 3), Schoch & Milner (2000:fig. 48), and Eltink et al. (2016:figs. 2A, 3, 5).
- **27 (orbit margins):** 0 → 1. Eltink et al. (2016:842) states that the orbital margins are flush with the skull roof.
- **31 (frontal-nasal, length):** 0 → 1. As figured by Barberena (1998:fig. 3) and Eltink et al. (2016:figs. 2D, 5); this holds true regardless of whether the length is measured along the midline or as the greatest length of each element.
- **39 (prefrontal-frontal):** 0 → 0&1. As described and figured by Eltink et al. (2016:845, figs. 2, 5); only one specimen appears to have a more anteriorly extensive prefrontal (39–0; fig. 2f therein).
- **50 (postorbital, postfrontal):** 0 → -. As reconstructed by Barberena (1998:fig. 3) and Eltink et al. (2016:fig. 5), the postfrontal is longer than the parietal and the supratemporal, but the postorbital is only longer than the supratemporal.

- **67 (tabular, ventral crest):** 1 → 2. The original code was one feature separating this taxon from all other nominal rhinesuchids, which may partially explain why it was not recovered with these taxa in previous derivates, contrary to rhinesuchid-focused studies (Eltink et al., 2016, 2019; Marsicano et al., 2017). The occipital reconstructions by Barberena (1998:fig. 5) and Eltink et al. (2016:fig. 7) depict a very developed ventral crest that is of similar proportions to other rhinesuchids (Watson, 1962:fig. 7; Shishkin & Rubidge, 2000:fig. 4) and is much taller than in other taxa originally coded for 67–1 (e.g., *Lydekkerina*, *Chomatobatrachus*, *Sclerocephalus*; Cosgriff, 1974:fig. 37; Jeannot et al., 2006:fig. 4B; Schoch & Witzmann, 2009:fig. 4D).
- **88 (ectopterygoid, fangs):** 0 → 1. As described and figured by Barberena (1998:fig. 4) and Eltink et al. (2016:848, figs. 3C, 4C, 6).
- **102 (vomer-pterygoid):** 0 → 1. As described and figured by Barberena (1998:fig. 4) and Eltink et al. (2016:848, figs. 3C, 4C, 6).
- **106 (basicranium, contact):** 0 → 1. There is a long sutural contact spanning the anterior half of the length of the basal plate (Barberena, 1998:134, fig. 4; Eltink et al., 2016:848–849, figs. 3C, 4C, 6).
- **145 (Meckelian window):** 0 → 1/2. Although the adductor fossa is not completely defined (Eltink & Langer, 2014), the posterior Meckelian window is a very long and slender oval that is definitively larger than other taxa originally coded for 145–0.
- **159 (caudal count):** 0 → ?. Caudal remains are very fragmentary and isolated, and an estimate of the full tail length or number of caudal positions is unknown (Dias & Schultz, 2003; Eltink & Langer, 2014).
- **160 (transverse process, orientation):** 0 → 1. Dias & Schultz (2003:32, figs. 7–8) stated that the transverse processes extend laterally at a right angle.
- **174 (ribs, uncinate spines):** 0 → -. The uncinate processes were treated as uncinate blades (173–1), and there is no secondary process.
- **188 (scapula, glenoid facet):** 1 → 0. Dias & Schultz (2003:36, figs. 12–13) state that "there is a prominent dorsal articular glenoid facet in both specimens, but a complete glenoid cavity is not preserved." This indicates that the glenoid is partially ossified.
- **194 (humerus, condyles):** ? → 1. As figured by Dias & Schultz (2003:fig. 13).
- **196 (carpals):** 0 → 1. As described and figured by Dias & Schultz (2003:36, figs. 12–13).
- **199 (ilium, dorsal end):** ? → 0. As described and figured by Eltink & Langer (2014:529, fig. 4).
- **200 (ilium, height):** ? → 0. As with character 199.
- **203 (pubis):** ? → 1. As with character 199.
- **204 (femur):** ? → 1. As figured and described by Eltink & Langer (2014:529, fig. 5H).
- **205 (femur, trochanter):** ? → 0. As figured and described by Dias & Schultz (2003:37, fig. 14) and Eltink & Langer (2014:529, fig. 5).
- **242 (number of premaxillary teeth):** ? → 1. As reconstructed by Barberena (1998:fig. 4) and Eltink et al. (2016:fig. 6), there are no more than nine premaxillary teeth (the upper bound for 242–1).
- **251 (denticles on vomers):** 1 → 0. As described by Eltink et al. (2016:847) and contra Barberena (1998:132).

- **278 (trunk intercentra):** ? → 0. As described and figured by Dias & Schultz (2003:32, 34, figs. 2–5, 7–8).
- **279 (trunk neural arch to centrum):** ? → 0. As described and figured by Dias & Schultz (2003:32, 34, figs. 2–5, 7–8).
- **281 (haemal arches):** ? → 0. As described and figured by Dias & Schultz (2003:34, figs. 2–3).
- **282 (haemal arches):** ? → 1. As described and figured by Dias & Schultz (2003:34, figs. 2–3).
- **283 (haemal arches):** ? → 1. As described and figured by Dias & Schultz (2003:34, figs. 2–3).
- **286 (transverse process):** ? → 0. As described and figured by Dias & Schultz (2003:32, figs. 7–8).
- **299 (supraglenoid foramen):** ? → 0. As described and figured by Dias & Schultz (2003:36, fig. 12).
- **300 (number of coracoid foramina):** ? → 1. As with character 299.
- **301 (scapulocoracoid ossification):** ? → 0. As with character 299.
- **302 (deltopectoral crest):** ? → 1/2. As described by Dias & Schultz (2003:36).
- **305 (olecranon process):** ? → 1. As described by Dias & Schultz (2003:36).

***Lapillopsis nana***

- **21 (lacrimal)**: **? → 0.** Warren & Hutchinson (1990) originally interpreted the lacrimal as being absent in the holotype and paratype, although sutures in this region are not actually preserved by their own admonition. Their rationale was that the sutures that were identified "leave no room for a lacrimal" (p. 150 therein). Yates (1999) described additional specimens from the type locality that preserve a short lacrimal separated from both the nares and the orbit. In the holotype, this was originally misinterpreted as a posterior expansion of the medially expanded maxilla, and in the paratype, this was originally misinterpreted as a less expanded portion of the maxilla. The lateral suture of the lacrimal was identified by Warren & Hutchinson but as that of other elements.
- **22 (lacrimal, lateral suture)**: **? → 0.** Based on Yates' (1999) revised identification of the lacrimal.
- **23 (lacrimal, position)**: **? → 0.** Based on Yates' (1999) revised identification of the lacrimal.
- **24 (lacrimal, posterior extension)**: **? → 0.** Based on Yates' (1999) revised identification of the lacrimal.
- **40 (prefrontal, anterior end): 1 → 0&1**. The holotype and paratype both have prefrontals with relatively blunted ends, but the left prefrontal of QM F14497 (Yates, 1999:fig. 9B) is definitively pointed. This is also depicted in Yates' reconstruction (fig. 5 therein) but on the shaded side of the skull in which the sutures are not as stark.
- **43 (prefrontal, process): ? → 0**. This feature warrants particular mention because it is an ubiquitous feature among dissorophoids (termed the 'ventral prefrontal process' or 'VPP'). It is perplexing that this taxon was not coded for this character because it was coded for every other temnospondyl other than *Peltostega*, even though this region is not any worse preserved or illustrated in *Lapillopsis* than in other taxa. Additionally, Warren & Hutchinson (1990:155) explicitly state that this process is absent in *Lapillopsis*. This interpretation is not altered by Yates' (1999) identification of a lacrimal; the VPP may

only correlate with the presence of a lateral exposure of the palatine (LEP), which is absent in *Lapillopsis* but also common in dissorophoids.

- **45 (prefrontal-maxilla): 1 → 0**. This appears to be a coding error based on the original interpretation of Warren & Hutchinson (1990). The lacrimal identified by Yates (1999) separates these elements.
- **55 (supratympanic flange): 1 → 0.** This is another feature shared only with certain dissorophoids in the original coding of Schoch (2013). Warren & Hutchinson (1990:155) explicitly state that this flange is absent in *Lapillopsis*. The original code may be a misinterpretation of the unornamented medially directed flange from the squamosal just anterior to the quadrate. This is visible only in dorsal view and is therefore not homologous with (or equivalent to) the large unornamented flange extending dorsally within the otic notch that is often marked by striations for a presumed tympanum. The supratympanic flange (Bolt, 1974) is considered to be found only in olsoniforms and the amphibamiform *Eoscopus* (Daly, 1994) but not *Micropholis* (Schoch & Rubidge, 2005). The condition of *L. nana* is not unlike other terrestrial temnospondyls like other amphibamiforms for which state 0 was coded.
- **59 (jugal-lacrimal): ? → 0.** This appears to be another code related to Warren & Hutchinson's (1990) original interpretation of an absent lacrimal.
- **75 (quadrate and occipital condyles): 1 → ?.** Yates' (1999) reconstruction depicts the posterior face of the exoccipitals about in line with the posterior face of the quadrates. However, the former is clearly inferred as indicated by the dashed lines – complete exoccipitals are not known from any specimen, let alone in one with the quadrate preserved, and Yates stated that they were too incomplete to even determine whether one or two condyles were present. It is best left as unknown.
- **77 (dentition, marginal): 0 → 1**. Marginal dentition is unknown in the type specimens. Yates (1999:304) stated that "maxillary teeth […] are rarely well preserved but when they are present they are small, acutely conical and closely spaced." If the dentary is considered for this character, the largely complete dentary of the paratype preserves uniformly sized, evenly spaced teeth. In my opinion, this is sufficient to code this taxon for state 1, which occurs in all post-rhinesuchid stereospondyls and some amphibamiforms.
- **85 (transvomerine tooth row): ? → 0.** The only dentition on the palate are small tusks, and the region where transvomerine teeth typically occur is preserved in at least two specimens of *Lapillopsis nana* (QM F35386, QM F14502; Yates, 1999:figs. 9C, 9I).
- **91 (anterior palatal opening): ? → 1.** This appears to be an outdated code based on Warren & Hutchinson (1990), who only inferred a continuous contact between the premaxillae and vomers; Yates (1999) described specimens that clearly show this opening.
- **92 (anterior palatal opening[s]): ? → 0.** As with character 91.
- **120 (pterygoid, squamosal): 0 → 1.** Yates (1999:306) stated that "there is a complete palatoquadrate fissure because at no point does the ascending ramus contact the dorsal skull roof."
- **121 (pterygoid, flange): 0 → 1.** Previously state 1 was restricted to dissorophoids and lissamphibians, but there is clearly a strongly laterally projecting flange from the palatine ramus, and as in some dissorophoids, it angles ventrally (Yates, 1999).

- **136 (occipital condyle): 1 → ?.** Yates (1999:308) stated it was impossible to determine whether the condyle was single and bilobed or formed by two separate condyles.
- **137 (exoccipital condyles): 0 → ?**. As with character 75.
- **140 (postglenoid area): 0 → 1**. There is a distinct long process posterior to the glenoid (Yates, 1999:figs. 7B, 12).
- **142 (postglenoid area, dorsal)**: **0 → ?**. The dorsal surface of the PGA is not figured, described, or reconstructed in a sufficient fashion to determine whether it was "plain" or with an elongated groove. Yates (1999:308) states that the PGA is not preserved in the type specimens, and referred specimens can be viewed only in "lateral, medial and ventral views."
- **143 (hamate process): 0 → 1.** Yates (1999:308) describes a hamate process.
- **146 (symphyseal teeth): 0 → ?**. There is no specimen figured with the symphysis exposed in dorsal view, nor is there any description of the dentition in this region or a reconstruction of this profile.
- **150 (stapes, quadrate process): ? → 0.** The stapes figured by Yates (1999:fig. 7B) is sufficient to code this character.
- **151 (stapes, ventral process): ? → 1.** As with character 150.
- **152 (stapes, shape): ? → 1.** As with character 150
- **179 (interclavicle, length): 0 → ?.** There is no complete interclavicle for this taxon; both anterior and posterior extents are unknown (Yates, 1999).
- **182 (interclavicle, proportions): 1 → 0/1.** As with character 179; the length is clearly greater than the width, but the extent is unclear.
- **191 (humerus, adult shaft): ? → 1/2.** Yates (1999:309) remarked that is "is a remarkably elongated bone with a distinct cylindrical shaft"; it is unclear if it is sufficiently elongate to meet state 2's criterion of more than 50% of the humeral length because of the limited exposed profile.
- **193 (humerus, supinator): ? → 1.** Both Warren & Hutchinson (1990) and Yates (1999) indicate a supinator is absent.
- **202 (ilium, tip): 1 → ?.** The ilium of this taxon is unknown.
- **205 (femur, trochanter): ? → 1.** Both Warren & Hutchinson (1990) and Yates (1999) indicate that no distinctive process is formed as a trochanter.
- **212 (rib cage): 0 → ?**. The ribs are too incompletely preserved to even be described (Yates, 1999), let alone to characterize the ribcage.
- **219 (lacrimal-naris): ? → 1.** The lack of coding appears to follow Pardo et al.'s (2017) undocumented modifications to previously coded lacrimal characters (see characters 21–24).
- **220 (lacrimal-orbit): ? → 0.** As with character 219.
- **228 (posterior skull margin): 0 → 0&3.** While the type specimens have a smoothly concave margin, at least one specimen (QM F14501) appears to have an undulating margin, with short rounded projections from the postparietals (Yates, 1999:fig. 7). This is also what Yates reconstructed in his composite, even though he did not comment on the discrepancies.
- **247 (jugular foramen): 0 → 1.** Yates (1999:307) stated that the exoccipitals bear a large foramen for cranial nerves IX and X; there is no mention of a contribution from the opisthotic.

- **249 (pterygoid denticles): 0 → 1.** Yates (1999:307) stated that there were no denticles on the palatal bones. This conflicts with Warren & Hutchinson (1990), who stated denticles were present on the parasphenoid and possibly on the basipterygoid ramus of the pterygoid. Yates' interpretation is accepted since there are additional specimens with these elements preserved that he examined, and these regions appear to be covered in ornamentation instead of denticles.
- **271 (number of coronoids): 0 → 0/1**. Yates (1999) identified a posterior and a middle coronoid but did not confidently identify a third coronoid, and therefore merely assumed as much based on the stereotypical condition of temnospondyls.
- **307 (femur): ? → 0.** The femur is relatively long and in line with other taxa coded for state 0.
- **310 (splenial, dorsal margin): 0 → ?**. The suture between the first and second coronoids is unknown (see also character 271), as indicated in Yates' (1999) reconstruction in which he reconstructed the splenial as contacting both of these elements dorsally (state 0 here).
- **345 (dorsolateral osteoderms fused to ribs): 0 → ?** Yates (1999:308) notes that the only specimen with ribs has ribs that are too incomplete and obscured to allow description. Other rib characters in this matrix were not coded.

***Peltobatrachus pustulatus***

- **22 (lacrimal, lateral suture):** 0 → ?. There is no positive evidence of a lacrimal (only that a lacrimal did not enter the orbit if present; Panchen, 1959:212, figs. 1, 3); note also that character 21 (lacrimal presence/absence) was originally uncoded.
- **24 (lacrimal, posterior extension):** 0 → ?. As with character 22.
- **39 (prefrontal-frontal):** ? → 0. As illustrated by Panchen (1959:fig. 1), the prefrontals extend past the frontal-nasal suture, although their full extent is uncertain.
- **45 (prefrontal-maxilla):** 0 → ?. The lateral region of the snout is entirely unknown, including the anterior and lateral extent of the prefrontals; the maxilla is entirely reconstructed (Panchen, 1959:figs. 3–4).
- **50 (postorbital, postfrontal):** 0 → -. The right postfrontal appears to be at least as long as the right supratemporal and probably both parietals (anterior margins poorly defined), but the postorbital, as preserved, is shorter than both. B
- **59 (jugal-lacrimal):** 0 → ?. As with characters 22 and 24; no lacrimal is definitively known, and the anterolateral and lateral sutures around the orbit are entirely unknown. A lacrimal excluded from the orbit as Panchen suggested does not preclude a jugal-lacrimal contact.
- **86 (additional vomerine fangs):** 0 → ?. There is no discussion of vomerine dentition, and the vomers are very incomplete in the only specimen with preserved palatal elements (Panchen, 1959:214, fig. 1).
- **104 (vomer, extension):** 0 → ?. This character requires vomer-pterygoid contact (102–0), but character 102 was originally uncoded, and Panchen (1959:fig. 4) reconstructed the palate without this contact.
- **125 (palatine, ectopterygoid):** 0 → ?. The sutures of these elements is entirely conjectural based on other temnospondyls (Panchen, 1959:221).
- **126 (palatine, vomer):** 0 → ?. As with character 125.

- **127 (LEP):** 0 → ?. The palatine is not confidently identified in any capacity, and the region lateral/ventral to the orbit where an LEP could occur is not preserved on either side (Panchen, 1959:214, 221, fig. 1).
- **129 (palatine, ectopterygoid, width):** 0 → ?. As with character 125. There is not even a cursory identification of an ectopterygoid (Panchen, 1959:221).
- **131 (ectopterygoid, length):** 0 → ?. As with characters 125 and 129.
- **132 (LEE):** 0 → ?. As with characters 125, 127, and 129.
- **133 (ectopterygoid, Y-shaped**): 0 → ?. As with characters 125 and 129.
- **159 (caudal count):** 0 → ?. Character 158 (presacral count) was originally uncoded, and there are no more than seven articulated presacral positions or more than 12 associated presacral positions from one individual. This is more than the maximum number of associated caudal positions.
- **178 (clavicle, ventral blade):** 0 → ?. Panchen (1959:236) stated that the clavicle is not preserved.
- **193 (humerus, supinator):** 1 → 0. Panchen (1959:236, fig. 10) identified a supinator process.
- **226 (prefrontal contribution to external naris):** 0 → 0/1. The anterior terminus of the prefrontals is unclear, and no margin of the naris is preserved (Panchen, 1959:figs. 1, 3).
- **258 (pterygoids contact anteriorly):** 1 → ?. The anterior extent of the pterygoids and most of the anterior palate are unknown.
- **271 (number of coronoids):** ? → 0/1/2. At least the posteriormost coronoid is documented (Panchen, 1959:fig. f).
- **312 (parietal width):** 1 → 0. As figured by Panchen (1959:figs. 1, 3).

***Rhineceps nyasaensis***

- **50 (postorbital, postfrontal):** 0 → -. The short left postorbital confounds coding of this character. It is shorter than the parietal and the supratemporal, but the left postfrontal is at least as long if not longer than both. On the right side, the postorbital is as long as the parietal and the supratemporal, and the left postfrontal is definitively longer than either. Because two elements are specified here, the character is changed to unknown rather than to polymorphic (i.e. neither the postorbital nor the postfrontal alone is sufficient to code the character).
- **57 (jugal, ventral process):** 1 → ?. There is no figured or described ventral process per Watson (1962:fig. 6) and Marsicano et al. (2017:fig. 7B); instead, the pterygoid apparently extends a thin process laterally from the palatine ramus to separate the ectopterygoid from the subtemporal vacuity. Marsicano et al. (2017; character 73) coded an equivalent character for the ventral process in this taxon, whereas Eltink et al. (2019; character 55) left an equivalent character uncoded. The most conservative approach is to leave this as unknown.
- **88 (ectopterygoid fangs):** 0 → 0&1. Watson (1962:235, fig. 7) remarked on the asymmetry of the ectopterygoid dentition. While it is possible, as he suggested, that the lack of enlarged teeth on one ectopterygoid is a matter of replacement at the time of death, this has not been considered in the coding of other taxa that symmetrically lack enlarged ‘fangs’ or ‘tusks,’ and the possibility remains hypothetical until it can be tested (e.g., through histology).

- **159 (caudal count):** 0 → ?. There are no postcrania known for this taxon (Watson, 1962; Schoch & Milner, 2000; Marsicano et al., 2017), and character 158 (presacral count) was originally uncoded.
- **160 (transverse process, orientation):** 1 → ?. As with character 159.
- **161 (transverse process, length):** 0 → ?. As with character 159.
- **162 (neural spine, height):** 0 → ?. As with character 159.
- **163 (intercentrum, shape):** 0 → ?. As with character 159.
- **164 (intercentrum, width):** 0 → ?. As with character 159.
- **165 (intercentrum, ventral surface):** 0 → ?. As with character 159.
- **166 (intercentrum, anterior surface):** 0 → ?. As with character 159.
- **167 (parapophysis):** 0 → ?. As with character 159.
- **168 (pleurocentrum, presence):** 0 → ?. As with character 159.
- **169 (pleurocentrum, lateral surface):** 0 → ?. As with character 159.
- **170 (pleurocentrum, ventral extension):** 0 → ?. As with character 159.
- **171 (ribs, length):** 0 → ?. As with character 159.
- **172 (ribs, ventral extension):** 0 → ?. As with character 159.
- **174 (ribs, uncinate spines):** 0 → ?. As with character 159.
- **176 (cleithrum):** 0 → ?. As with character 159; note also that character 175 (cleithrum) and character 177 (cleithrum), were originally uncoded.
- **178 (clavicle, ventral blade):** 0 → ?. As with character 159.
- **179 (interclavicle, length):** 1 → ?. As with character 159.
- **181 (interclavicle, central ornamented area):** 0 → ?. As with character 159.
- **182 (interclavicle, proportions):** 1 → ?. As with character 159.
- **183 (interclavicle, posterior margin):** 0 → ?. As with character 159.
- **184 (interclavicle, anterior margin):** 1 → ?. As with character 159.
- **185 (interclavicle, anterior stylus):** 0 → ?. As with character 159.
- **186 (interclavicle, posterior stylus):** 0 → ?. As with character 159.
- **187 (interclavicle, clavicles):** 0 → ?. As with character 159.
- **202 (ilium, tip):** 1 → ?. As with character 159.
- **209 (osteoderms):** 0 → ?. As with character 159.
- **212 (rib cage):** 0 → ?. As with character 159.
- **243 (number of maxillary teeth):** 1 → 0. No tooth count has been previously given for this taxon, but there are nearly 30 illustrated positions on the right maxilla with substantial gaps between teeth unaccounted for.
- **322 (splenial teeth):** ? → 1. There is no indication of splenial teeth (Watson, 1962:237–238, fig. 8).
- **345 (dorsolateral osteoderms fused to ribs):** 0 → ?. As with character 159.

***Uranocentrodon senekalensis***

- **249 (denticles on pterygoid):** ? → 0. Broom (1930:fig. 2) depicted denticles on the pterygoid. This agrees with both earlier work (e.g., Haughton, 1925:228) and the phylogenetic coding by Marsicano et al. (2017), the most recent revision of Rhinesuchidae.
- **251 (denticles on vomer):** ? → 0. As with character 249.

- **312 (parietal width):** ? → 1. As most recently documented by Marsicano et al. (2017:fig. 4; but see also Schoch & Milner, 2000:fig. 55, for example).
- **313 (skull table):** ? → 0. As with character 311.
- **314 (prefrontals):** ? → 0. As with character 311.
- **315 (postfrontals):** ? → 0. As with character 311.
- **317 (zygokrotaphy):** ? → 0. As with character 311.

***Broomistega putterilli***

- **33 (lateral line sulci, adults):** 0 → ?. There are no sensory grooves present on the larger holotype skull, but due to incompleteness, this should not be treated as definitive absence.
- **34 (sulci, skull roof):** 0 → ?. While the grooves are not found across the skull roof in the smaller referred specimens (34–0; there is no postorbital groove), the two grooves present (the supraorbital and infraorbital sulci) are not confined to the circumorbital region (34–1).
- **102 (vomer-pterygoid):** ? → 1. As figured by Shishkin & Rubidge (2000:fig. 5B). Note also that character 104, which requires this contact (102–0), was previously coded.
- **117 (pterygoid, ventral ornament):** 0 → 1. Shishkin & Rubidge (2000:664, fig. 5A) specifically noted a field of groove-and-ridge ornamentation.
- **152 (stapes, shape):** ? → 1/2. The proximal end (Shishkin & Rubidge, 2000:fig. 5A) is sufficient to determine that it was not "robust" or "tetrahedral" (152–0).
- **255 (stapes):** ? → 1. Shishkin & Rubidge (2000:664) stated that there is no stapedial foramen.

***Lydekkerina huxleyi***

- **38 (maxilla-nasal):** 1 → 0&1. This condition is intraspecifically variable, independent of taphonomic loss of the septomaxilla, per Jeannot et al. (2006:fig. 5).
- **75 (quadrate and occipital condyles):** 1 → 0. Illustrations and reconstructions (Shishkin et al., 1996:fig. 7; Schoch & Milner, 2000:fig. 57; Jeannot et al., 2006:figs. 2, 7; Hewison 2007:figs. 30, 32) depict occipital condyles anterior to the quadrate condyles.
- **88 (ectopterygoid, fangs):** 1 → 0&1. Jeannot et al. (2006), citing Shishkin et al. (1996:1646), stated this character is polymorphic.
- **124 (pterygoid-palatine-ectopterygoid):** 0 → 0&1. Jeannot et al. (2006:829) noted that the ectopterygoid is excluded from the interpterygoid vacuity by a point contact between the pterygoid and the palatine (as with Shishkin et al., 1996:fig. 7), whereas Hewison (2007:22) noted the absence of this suture in other specimens.
- **145 (Meckelian window):** 1 → 0. The size of this opening is proportionately in line with the plesiomorphic condition observed in most non-stereospondyls (e.g., Jeannot et al., 2006:fig. 4; Hewison, 2007:fig. 34)
- **155 (basibranchials):** ? → 1. Jeannot et al. (2006:fig. 6B) and Witzmann (2013:155) noted the presence of a basibranchial.
- **196 (carpals):** ? → 1. Pawley & Warren (2005:289, fig. 5) described and figured carpals.
- **219 (lacrimal-naris contact):** 1 → 0&1. Jeannot et al. (2006:828, fig. 5) cited this as an example of non-taphonomic intraspecific variation.
- **243 (maxillary tooth count):** 1 → 0. Jeannot et al. (2006:830) were able to identify at least 29 teeth on the left maxilla of the holotype and 38 teeth on the right. Hewison (2007:12) only identified 31 teeth on the right maxilla but reconstructed several additional positions.

- **272 (coronoid teeth):** 1 → 0. Jeannot et al. (2006) and Hewison (2007:35) noted the presence of dentition of a similar size to the marginal dentition, intermingled with much smaller 'denticles.'
- **273 (coronoid teeth, size):** ? → 1. As with character 272.
- **287 (atlas-axis centra):** ? → 0. Jeannot et al. (2006:832) reported the presence of intercentra associated with these two positions.
- **293 (proatlantes):** ? → 1. Pawley & Warren (2005:282), citing Broili and Schröder (1937), considered the absence of a proatlas among the abundance of material to be genuine.
- **312 (parietal, width):** 1 → 0. Jeannot et al. (2006) and Hewison (2007) depict parietals that are wider than the frontals.

***Chomatobatrachus halei***

- **146 (symphyseal teeth):** ? → 0. Cosgriff (1974:62–63, fig. 40A) only figures symphyseal tusks and denticles on this region.
- **185 (interclavicle, anterior stylus):** 0 → ?. The only postcrania of this taxon consists of clavicles (Cosgriff, 1974:44; Schoch & Milner, 2000:74), and all other interclavicle characters were originally uncoded.
- **202 (ilium, tip):** 1 → ?. As with character 185; all other iliac characters were originally uncoded.
- **212 (rib cage):** 0 → ?. As with character 185.
- **243 (number of maxillary teeth):** ? → 0. Cosgriff (1974:50) identified 68 positions on the upper tooth row; although the premaxillary-maxillary suture is not identified, their relative lengths are typical for temnospondyls (as viewed dorsally; fig. 31 therein), and therefore it can be concluded that well more than half of the tooth row occurred on the maxilla.
- **246 (supraoccipital):** ? → 0. As described by Cosgriff (1974:52–54, fig. 37).
- **247 (jugular foramen):** ? → 1. As described by Cosgriff (1974:52, fig. 37).
- **249 (denticles on pterygoid):** ? → 0. As described by Warren et al. (2006:881, fig. 4c).
- **250 (teeth on pterygoid):** ? → 0. As described by Cosgriff (50–51) and by Warren et al. (2006:881, fig. 4c).
- **251 (denticles on vomers):** ? → 0. As with character 249.
- **252 (denticles on palatines):** ? → 0. As with character 249.
- **254 (cultriform process contact with vomer):** ? → 0. As with character 248.
- **257 (sphenethmoid):** ? → 0. As described by Cosgriff (1974:55).
- **258 (pterygoids contact anteriorly):** ? → 1. As with character 248.
- **259 (lateral process of pterygoid):** ? → 0. As with character 248.
- **263 (angular):** ? → 1. As figured by Cosgriff (1974:fig. 42).
- **270 (angular extent in lateral view):** 0 → 1. As figured by Cosgriff (1974:fig. 42b).
- **323 (choanal openings):** ? → 0. As with character 248.

***Peltostega erici***

- **11 (premaxilla, prenarial portion):** 0 → ?. The snout is not preserved in the prenarial region of the most complete specimens (e.g., Janvier, 1983:fig. 2; reproduced in Schoch & Milner, 2000:fig. 60). The only specimen with a premaxilla is a very fragmentary snout figured and described only in ventral view (Janvier, 1983:fig. 3B) and that was not incorporated into Janvier’s reconstruction (the most recent to date). No specimen has both the prenarial region and the frontal (the specifier for prenarial length comparison).

- **37 (maxilla, anterior margin):** 0 → ?. The lateral margins of the maxilla are unknown anterior to the orbit (e.g., Janvier, 1983:figs. 2, 3; reproduced in Schoch & Milner, 2000:fig. 60).
- **89 (interpterygoid vacuities):** 0 → 1. Especially in the context of lissamphibians, the condition of this taxon is more similar to all other temnospondyls than to outgroups (e.g., Cosgriff & Zawiskie, 1979:fig. 10D).
- **226 (prefrontal contributes to external naris):** 0 → ?. The prefrontal’s anterior extent and the posterior narial margin are unknown, and as the prefrontal often expands in the absence of a lacrimal (as often reconstructed for this taxon), the character is best left as unknown.
- **249 (denticles on pterygoid):** ? → 0. Cosgriff & Zawiskie (1979:17) mentioned a few scattered denticles in the holotype that might have represented a more complete shagreen.
- **251 (vomer, denticles):** ? → 1. As with Cosgriff & Zawiskie (1979); denticles are not found on the rest of the palate.
- **252 (palatine, denticles):** ? → 1. As with character 251.
- **312 (parietal, width):** ? → 0&1. This is intraspecifically variable (Janvier, 1983:fig. 2; Schoch & Milner, 2000:fig. 60).
- **313 (skull table):** ? → 0. Enough of the skull is preserved to code it like other temnospondyls.
- **314 (prefrontals):** ? → 0. Prefrontals are definitively present.
- **315 (postfrontals):** ? → 0. Postfrontals are definitively present.
- **317 (zygokrotaphy):** ? → 0. As with character 313.

***Laidleria gracilis***

- **40 (prefrontal, anterior end):** 1 → 0. The prefrontal tapers to a point and is no less “pointed” than other taxa coded for 40–0 (Warren, 1998a:fig. 4; Schoch & Milner, 2000:fig. 62; Dias-da-Silva & Marsicano, 2011:fig. 1F).
- **88 (ectopterygoid, fangs):** 1 → ?. The reconstruction (Warren, 1998a:fig. 4) does not show enlarged teeth at the anterior end of the ectopterygoid, only a continuation of the tooth row of the palatine (88–1). Warren (p.173 therein) also states it is not possible to determine whether ectopterygoid tusks were present.
- **123 (pterygoid, ectopterygoid):** 0 → ?. Sutures other than the parasphenoid-pterygoid, the parasphenoid-vomer, and the palatine-vomer sutures are entirely conjectural (Warren, 1998a:171, figs. 2B, 4B).
- **124 (pterygoid-palatine-ectopterygoid):** 1 → ?. As with character 123.
- **125 (palatine, ectopterygoid):** 0 → ?. As with character 123.
- **131 (ectopterygoid, length):** 0 → ?. As with character 123.
- **137 (exoccipital condyles):** 0 → 1. The exoccipitals have a broad exposure in dorsal view (Warren, 1998a:figs. 2A, 4A).
- **168 (pleurocentra):** 1 → ?. Warren (1998a:177) identified small fragments that might be pleurocentra and equivocated on whether the pleurocentra might be merely reduced or entirely unossified.

- **170 (pleurocentrum, ventral extension):** 0 → ?. As with character 168; it also should not have been coded under the original interpretation that pleurocentra were unossified (168–1).
- **172 (ribs, uncinate blades):** 0 → -. The ribs are simple without distinctive uncinate processes (Kitching, 1958:fig. 17).
- **173 (ribs, uncinate spines):** 0 → -. As with character 172.
- **185 (interclavicle, anterior stylus):** 0 → ?. The interclavicle is entirely unknown (Kitching, 1958; Warren, 1998a), and all other interclavicle characters were originally uncoded.
- **249 (denticles on pterygoid):** 0 → ?. Warren (1998:173) discussed the uncertainty regarding previous reports of pterygoid denticles by Cosgriff & Zawiskie (1979) but not by Kitching (1958).

***Siderops kehli***

- **11 (premaxilla, prenarial portion):** 0 → ?. The reconstruction of the premaxillary-nasal suture is entirely conjectural (Warren & Hutchinson, 1983:9–10, fig. 2).
- **12 (premaxilla, outline):** 0 → ?. As with character 11.
- **31 (frontal-nasal, length):** 0 → ?. The reconstruction of the frontal-nasal suture is entirely conjectural (Warren & Hutchinson, 1983:fig. 2).
- **32 (frontal-nasal, suture**): 0 → ?. As with character 31.
- **35 (infraorbital sulcus):** 0 → ?. The sensory groove canals are both asymmetrical and interrupted (Warren & Hutchinson, 1983:9–10, figs. 1–2); it is not clear that this groove is sufficiently posteriorly extensive to capture any flexure like that found in some other taxa. If it is considered to be sufficient, then it would be state 1 (S-shaped flexure) based on a prominent curve from the left maxilla to the left prefrontal; the right side does not preserve a groove in the same region.
- **38 (maxilla-nasal):** 1 → ?. Despite the absence of a lacrimal, this does not ensure maxilla-nasal contact, and this suture nor the anterior terminus of the prefrontal are confidently known (Warren & Hutchinson, 1983:fig. 2).
- **41 (prefrontal, lateral suture):** 0 → ?. Only a small portion of the anterolateral margin is preserved, and the region around the orbit (referenced in state 1) is unknown (Warren & Hutchinson, 1983:fig. 2).
- **44 (prefrontal-jugal):** 1 → ?. the anterior extent of the jugal is unknown, and the anterolateral and lateral orbital margins are not well-preserved (Warren & Hutchinson, 1983:fig. 2).
- **50 (postorbital, postfrontal):** 1 → ?. None of the four elements specified for relative length here (postorbital, postfrontal, supratemporal, parietal) are completely defined at both anterior and posterior ends (Warren & Hutchinson, 1983:fig. 2).
- **51 (otic notch):** 1 → 0. Warren & Hutchinson (1983:7, 9, fig. 2) expressly reconstructed an otic notch (so coding off of the reconstruction, as would have to be done for other previously coded characters, would produce this coding change) and described features of the occiput that indicate support for a notch (when ignoring the reconstruction in coding).
- **54 (supratemporal, lateral margin):** 0 → ?. The posterolateral margin is not complete on either side, and given the revised treatment in relation to character 62 (which was uncoded here), this too should be uncoded.

- **56 (semilunar flange):** 0 → ?. It is not possible to discern whether the supratemporal reached the otic notch (Warren & Hutchinson, 1983:fig. 2), so it cannot be determined that it had no ventral projection.
- **58 (jugal, anterior extension):** 0 → ?. As with character 44.
- **62 (squamosal-tabular, dorsal):** 1 → ?. This sutural relationship is entirely conjectural (Warren & Hutchinson, 1983:fig. 2).
- **71 (posterior skull rim):** 1 → ?. Although there is sufficient evidence for a tabular horn, its posterior extent is unknown, and thus this feature cannot be assessed (Warren & Hutchinson, 1983:fig. 2).
- **85 (transverse tooth row, transvomerine):** 0 → 1. Warren & Hutchinson (1983:15, fig. 9) described and figured these teeth.
- **127 (LEP):** 0 → ?. The region in which a hypothetical LEP could be found is not resolved suturally (Warren & Hutchinson, 1983:fig. 2).
- **168 (pleurocentra):** 0 → 1. Warren & Hutchinson (1983:31) stated that there were no ossified pleurocentra in association with the skeleton. Considering that practically the entire axial skeleton was recovered, the absence of pleurocentra suggests a biological absence (as in *Pelorocephalus*, as noted by Warren & Hutchinson); this may be more widespread in chigutisaurids or brachyopoids at large, but the paucity of postcranial remains for most taxa remains a hindrance.
- **170 (pleurocentrum, ventral extension):** 0 → -. As with character 168; note that character 169 (pleurocentrum, lateral surface) was originally uncoded.
- **196 (carpals):** 0 → ?. The humerus is the only element of the forelimb that is known.
- **214 (skull to trunk):** ? → 1. The presacral column and skull are nearly complete in this taxon, sufficiently to estimate this ratio.
- **222 (quadratojugal):** ? → 0. A quadratojugal is confidently identified (Warren & Hutchinson, 1983:6, 11).
- **226 (prefrontal-naris**): 1 → ?. As with character 38.
- **237 (squamosal-jugal):** 0 → ?. This sutural relationship is entirely conjectural (Warren & Hutchinson, 1983:fig. 2).
- **239 (tabular):** ? → 0. Warren & Hutchinson's (1983:7, 9) justification is considered sufficient for the inference of a tabular based on the occipital anatomy; other characters related to the tabular that were added by Pardo et al. (2017a) were coded by those workers (e.g., character 234, parietal-tabular contact).
- **268 (retroarticular process):** 0 → ?. Warren & Hutchinson (1983:21) stated that the process is lost in both lower jaws.

***Batrachosuchus browni***

*Note that corrections for characters 140–148, 174–175, 177–178, 202, 260–263, 268–272, 322, 326, and 344 (all mandibular or postcranial) are considered to be applicable regardless of whether the more restrictive concept of *Batrachosuchus browni* or *Bathignathus watsoni* is employed; neither taxon is known from more than a partial skull.

- **11 (premaxilla, prenarial portion):** 0 → ?. Chernin (1977:102) stated that "no premaxillary or maxillary sutures are discernible and only the posterior half of the right narial border is preserved as an impression." The reconstruction of the premaxilla is expressly based on other species of *Batrachosuchus* (p. 99–100 therein).
- **12 (premaxilla, outline):** 0 → ?. As with character 10.

- **19 (naris, flange):** 0 → ?. As with character 11 regarding narial preservation.
- **35 (infraorbital sulcus):** 0 → ?. The only specimen lacks the anterolateral margins where this sulcus typically occurs, and there was no discussion of the groove in the text (Chernin, 1977).
- **57 (jugal, ventral exposure):** 1 → ?. There was no comment or figuring of a ventral exposure by Chernin (1977). Neither was this reconstructed or reported in *Bathignathus watsoni* (Watson, 1956:fig. 6; Schoch & Milner, 2014:fig. 47B).
- **58 (jugal, anterior extent):** 0 → ?. The anterolateral margins of the orbits are not preserved, there is no comment on the anterior extent of the jugal, and the maxillary sutures are not identifiable. Schoch & Milner (2014:fig. 46E) reconstructed this taxon (and *Bathignathus watsoni*) as having jugals extending anterior to the orbit (58–1), but I prefer to leave this uncoded to be conservative.
- **91 (anterior palatal opening(s)):** 1 → ?. As with character 11 regarding the absence of premaxillae. The preserved anteriormost edge of the vomers is straight and without indicate of a pair of fossae or foramina in this position. The foramen situated entirely within the vomers (Chernin, 1977:fig. 5b) is not homologous with the paired anterior perforations for symphyseal fangs that are found in other temnospondyls and based on the specific inclusion of language regarding the fangs in this character, the median vomerine foramen is best regarded as being irrelevant to this character. Chernin (1977:104) also suggested that this foramen might be the same as a poorly ossified region in *Hadrokkosaurus bradyi* (Welles & Estes, 1969:fig. 26b), which is well posterior to the vomer-premaxilla suture and the anterior palatal opening.
- **92 (anterior palatal opening(s)**: 0 → ?. As with character 91.
- **140 (postglenoid area):** 1 → ?. There is no lower jaw material for this taxon (Chernin, 1977:102; Warren & Marsicano, 2000:465; Schoch & Milner, 2014:98).
- **141 (postglenoid area (types):** 0 → ?. As with character 140.
- **142 (postglenoid area (dorsal):** 0 → ?. As with character 140.
- **143 (hamate process):** 0 → ?. As with character 140.
- **144 (preglenoid process):** 0 → ?. As with character 140.
- **145 (Meckelian window):** 0 → ?. As with character 140.
- **146 (symphyseal teeth):** 0 → ?. As with character 140.
- **147 (posterior coronoid teeth):** 0 → ?. As with character 140.
- **148 (anterior, middle coronoid teeth):** 0 → ?. As with character 140.
- **174 (ribs, uncinate spines):** 0 → ?. No postcrania are known for this taxon (Chernin, 1977:102; Warren & Marsicano, 2000:465; Schoch & Milner, 2014:98), as reflected in the absence of codes for most other postcranial characters.
- **175 (cleithrum):** 0 → ?. As with character 174.
- **177 (cleithrum):** 0 → ?. As with character 174.
- **178 (clavicle, ventral blade):** 0 → ?. As with character 174.
- **202 (ilium, tip):** 1 → ?. As with character 174.
- **222 (quadratojugal):** ? → 0. The element is not exposed dorsally but is exposed in occipital view (Chernin, 1977:fig. 5C).
- **260 (dentary):** 0 → ?. As with character 140.
- **261 (dentary, coronoid process):** 0 → ?. As with character 140.
- **262 (surangular):** 0 → ?. As with character 140.

- **263 (angular):** 0 → ?. As with character 140.
- **268 (retroarticular process):** 0 → ?. As with character 140.
- **269 (articulation to tooth row):** 1 → ?. As with character 140.
- **270 (angular):** 0 → ?. As with character 140.
- **271 (number of coronoids):** 2 → ?. As with character 140. Note also that this is a particularly strange original code, as state 2 refers to only one coronoid (as in the plagiosaurid *Plagiosuchus* but contrary to all nominal brachyopoids, which have three).
- **272 (coronoid teeth):** 1 → ?. As with character 140.
- **322 (splenial teeth):** 1 → ?. As with character 140.
- **326 (anterior dentary teeth):** 1 → ?. As with character 140.
- **344 (dentary marginal dentition):** 0 → ?. As with character 140.

***Plagiosuchus pustuliferus***

- **88 (ectopterygoid fangs):** 1 → 0. Damiani et al. (2009:359) identified two large tooth sockets at the anterior end of the ectopterygoid that they interpreted as fangs; these are substantially larger than either more posterior teeth or marginal teeth.
- **130 (palatine, ectopterygoid, continuous tooth row):** 1 → ?. Damiani et al. (2009:359) stated that the palatine dentition is unknown.
- **209 (osteoderms):** 1 → 2. The osteoderms cover the entire body (Hellrung, 2003; Witzmann & Soler-Gijón, 2010).
- **219 (lacrimal-naris):** 0 → 1. The lacrimal is well-separated from the naris (Damiani et al., 2009:fig. 7).

***Sangaia lavina***

- **11 (premaxilla, prenarial portion):** 0 → ?. The premaxillae are not preserved in either the holotype of referred specimen (Dias-da-Silva et al., 2006:383, fig. 3), and the reconstruction of this element is entirely conjectural.
- **12 (premaxilla, outline):** 0 → ?. As with character 11.
- **58 (jugal, anterior extent):** 0 → 1. The jugal extends anterior to the orbit (Dias-da-Silva et al., 2006:384, figs. 3, 6).
- **89 (interpterygoid vacuities):** 1 → ?. Partial preservation of the parasphenoid and pterygoid (Dias-da-Silva et al., 2006:fig. 4) is insufficient to discern the shape of these openings.
- **91 (anterior palatal opening(s)):** 1 → ?. There is no indication that the anterior part of the palate is preserved in any specimen; there is no description or reconstruction of the vomers.
- **92 (anterior palatal opening(s)):** 0 → ?. As with character 91.
- **93 (vomer):** 1 → ?. As with character 91.
- **94 (vomer, paired anterior depressions):** 0 → ?. As with character 91.
- **95 (anterior palatal depression):** 0 → ?. As with character 91.
- **96 (vomerine ridges):** 0 → ?. As with character 91.
- **97 (vomerine pit and fontanelle):** 0 → ?. As with character 91.
- **98 (vomerine septum):** 0 → ?. As with character 91.
- **102 (vomer-pterygoid):** 1 → ?. As with character 91.
- **113 (cultriform process, structure):** 0 → ?. The cultriform process is not preserved (Dias-da-Silva et al., 2006:385, fig. 4B).

- **124 (pterygoid-palatine-ectopterygoid)**: 0 → ?. Neither the ectopterygoid nor the palatine are exposed, if they are even preserved, so while Dias-da-Silva et al. (2006:385) conjectured that the pterygoid contacted the palatine, this is not documented in any specimen, and it is unclear whether the pterygoid is fully complete anteriorly.
- **129 (palatine, ectopterygoid (width)):** 0 → ?. As with character 124.
- **130 (palatine, ectopterygoid (continuous tooth row)):** 1 → ?. As with character 124.
- **131 (ectopterygoid (length):** 0 → ?. As with character 124.
- **133 (ectopterygoid (Y-shaped):** 0 → ?. As with character 124.
- **141 (postglenoid area (types):** 0 → ?. There is no lower jaw material, let alone a preserved PGA, as indicated by the lack of coding for character 140 (postglenoid area, presence/absence).
- **143 (hamate process):** 0 → ?. As with character 141.
- **144 (preglenoid process):** 0 → ?. As with character 141.
- **145 (Meckelian window):** 0 → ?. As with character 141.
- **167 (parapophysis):** 0 → ?. No postcrania are known for this taxon (Dias-da-Silva et al., 2006), and other postcranial characters were largely uncoded.
- **178 (clavicle, ventral blade):** 0 → ?. As with character 167.
- **202 (ilium, tip)**: 1 → ?. As with character 167.
- **212 (rib cage)**: 0 → ?. As with character 167.
- **220 (lacrimal-orbit):** 1 → 0. The lacrimal is excluded by a prefrontal-jugal contact (44–1).
- **258 (pterygoids contact anteriorly):** 1 → ?. As with character 124.

***Gerrothorax pulcherrimus***

- **58 (jugal, anterior extent):** 0 → 0&1. Schoch & Witzmann (2012:fig. 1) indicated that this condition is variable within a single specimen.

***Edingerella madagascariensis***

- **9 (preorbital region, length):** 0 → 0&1. Maganuco et al. (2009:10) stated that the preorbital region in adults is between 49% to 59% of the total skull length, and the specimen that they described has a preorbital-postorbital ratio of 2.36 (their table 1). Steyer's (2003) previous description does not provide measurements, but it does clearly show that the largest specimens have a similar ratio.
- **57 (jugal, ventral exposure):** 1 → 0. There is no figuring or description of this exposure (Lehman, 1961:fig. 9; Steyer, 2003:fig. 2B; Maganuco et al., 2009:fig. 10).
- **75 (quadrate and occipital condyles):** 1 → 0. The occipital condyles terminate well before the posterior face of the quadrate condyles (e.g., Maganuco et al., 2009:figs. 9–10).
- **113 (parasphenoid, cultriform process):** 0 → 1. Maganuco et al. (2009:13, 15) described a longitudinal keel.
- **125 (palatine, ectopterygoid):** 1 → 0. As shown by Maganuco et al. (2009:fig. 10), the palatine-pterygoid contact is mediated by an anteriorly extensive pterygoid overlapping onto the palatine (which has a transverse suture with the ectopterygoid), not by a posteromedial process of the palatine (as in most other capitosaurs).
- **145 (Meckelian window):** 1 → 0. The size of this opening is proportionately in line with the plesiomorphic condition observed in most non-stereospondyls (e.g., Lehman, 1961:fig. 11) in which state 0 is coded (e.g., *Eryops*, *Sclerocephalus*).

- **202 (ilium tip):** 1 → ?. No pelvic material is known for this taxon (Steyer, 2003:545; Maganuco et al., 2009:9), and the four other iliac characters were not coded originally.
- **215 (skull, proportions):** 1 → 0. The skull is longer than it is wide.
- **237 (postparietal, length):** 1 → 0. The element is nearly a square and more anteriorly extensive than the tabular (Steyer, 2003:figs. 1–2, 6; Maganuco et al., 2009:fig. 6), as with most other stereospondyls.
- **251 (vomer, denticles):** 0 → 1. There are no denticles in this taxon per the common usage of 'denticle'; Maganuco et al. (2009:18) describe a single row of denticles that represent the transvomerine tooth row found in other stereospondyls; one of the other authors of that study (Steyer) has used the term 'denticle' for any tooth smaller than the marginal dentition in an earlier study of this taxon (Steyer, 2003).

***Watsonisuchus gungaj***

- **9 (preorbital region, length):** 0 → 1. Both reconstructions of *W. gunganj* and '*W. rewanensis*' have preorbital regions more than twice as long as the postorbital region (Warren, 1980:figs. 2, 4).
- **34 (sulci, skull roof):** 0 → ?. Warren (1980:26) stated that "lateral line grooves are poorly impressed on the skull and can be positively identified only on the left postorbital and the left and right prefrontal bones" in the holotype of '*W. rewanensis*'; these regions are circumorbital in nature (34–1). She also stated (p. 29 therein) that "lateral line grooves could not be determined except for a small area on the lateral part of each jugal" in the holotype of *W. gunganj* (the referred specimen is too incomplete to identify grooves).
- **35 (infraorbital sulcus):** 1 → ?. As with character 34; this groove was not identified.
- **50 (postorbital, postfrontal):** 0 → -. Because of the anteromedial projection of the postorbital into the jugal, the longitudinal length of this element is at least equal to, if not slightly greater than, that of the supratemporal in both taxa.
- **57 (jugal, ventral process):** 1 → ?. There is no mention of a ventral exposure of the jugal, nor is one figured for a specimen. Warren's reconstruction of *W. gunganj* (fig. 4 therein) depicts either a short gap or an interceding element between the ectopterygoid and the maxilla on the left side of the skull, but not on the right, and this might merely be a gap based on the corresponding photograph (fig. 3) therein. To be conservative, it is uncoded here.
- **85 (transverse tooth row, transvomerine):** 1 → 1&2. Warren (1980:28) described a 'V-shaped' row in '*W. rewanensis*.' The condition in *W. gunganj* is a little unclear because the vomers have shifted slightly, but Warren argued that the best restored fit favored a straight transverse row.
- **88 (ectopterygoid, fangs):** 0 → 1. Ectopterygoid 'fangs' are neither described nor figured.
- **113 (parasphenoid, cultriform process):** 0 → 0&1. Warren (1980:28, 30) described a shallow longitudinal keel in '*W. rewanensis*' but not in *W. gunganj*.
- **178 (clavicle, ventral blade):** 0 → ?. There is no postcrania known for the Rewan taxa.
- **185 (interclavicle, anterior stylus):** 1 → ?. As with character 178.
- **202 (ilium, tip):** 1 → ?. As with character 178.
- **212 (ribcage):** 0 → ?. As with character 178.

***Benthosuchus sushkini***

- **75 (quadrate and occipital condyles):** 1 → 0. The occipital condyles terminate before the posterior face of the quadrate condyles (Bystrow & Efremov, 1940:figs. 64, 66–67; Damiani, 2001:fig. 11; Schoch, 2008:fig. 5B).
- **145 (Meckelian window):** 1 → 0. The size of this opening is proportionately in line with the plesiomorphic condition observed in most non-stereospondyls (e.g., Bystrow & Efremov, 1940:fig. 26; Schoch, 2000:fig. 7B; Damiani, 2001:fig. 12B) in which state 0 is coded (e.g., *Eryops*, *Sclerocephalus*).
- **168 (pleurocentra):** 0 → 1. Bystrow & Efremov (1940) did not identify or describe any pleurocentra (contra Romer, 1947, and as with Howie, 1970). Given the large size range and large sample size, the lack of articulated skeletons is not considered prohibitive to recoding this taxon.
- **169 (pleurocentrum, lateral surface):** 0 → -. As with character 168.
- **170 (pleurocentrum, ventral extension):** 0 → -. As with character 168.
- **279 (trunk neural arch to centrum):** ? → 0. As figured by Bystrow & Efremov (1940:fig. 39).
- **286 (transverse process):** ? → 0. As with character 279.
- **299 (scapula, supraglenoid foramen):** ? → 0. As figured by Bystrow & Efremov (1940:fig. 47).
- **300 (coracoid, coracoid foramina):** ? → 0. As with character 299.
- **301 (scapulocoracoid):** ? → 0. As with character 299.
- **302 (humerus, deltopectoral crest):** ? → 2. As figured by Bystrow & Efremov (1940:fig. 51).
- **305 (olecranon process):** ? → 1. As figured by Bystrow & Efremov (1940:fig. 52).
- **307 (femur):** ? → 1. As figured by Bystrow & Efremov (1940:fig. 54).

***Trematosaurus brauni***

- **31 (frontal, nasal):** 0 → 0&1. Schoch & Milner (2000:fig. 77) and Schoch (2019:53, fig. 6) noted variability in this feature.
- **44 (prefrontal-jugal):** 1 → 0&1. This condition is intraspecifically variable (Schoch, 2019a:fig. 5).
- **58 (jugal, anterior extent):** 0 → 0&1. This condition is intraspecifically variable (Schoch, 2019a:figs. 4–5).

**137 (exoccipital condyles):** 0 → 0&1. Schoch (2019a:53, fig. 6) noted variability in this feature.

***Trematolestes hagdorni***

- **9 (preorbital region, length):** 1 → 0. The orbits are situated in the anterior half of the skull (Schoch, 2006:fig. 4A).
- **44 (prefrontal-jugal):** 0 → 0&1. This condition is variable within a referred specimen (Schoch, 2006:fig. 3A).
- **57 (jugal, ventral exposure):** 1 → 0. This exposure is not described or reconstructed by Schoch (2006:fig. 4B).
- **58 (jugal, anterior extent):** 0 → 0&1. This variability is related to character 44; a more anteriorly extensive jugal contacts the prefrontal.
- **88 (ectopterygoid fangs):** 1 → 0. Schoch (2006) made no definitive statements either way, but the reconstruction in that study (fig. 4B therein) depicted teeth at the anteriormost region of the ectopterygoid that are larger than the posteriormost teeth of the

palatine. Based on other taxa with this condition that are coded for state 0 (e.g., *Glanochthon*, *Lyrocephaliscus*), the code is changed.

- **120 (pterygoid-squamosal):** ? → 0. Despite crushing of the holotype, other taxa only represented by flattened specimens were coded (e.g., *Balanerpeton, Micromelerpeton, Apateon*) for this character, so it is felt that the balance favors the incorporation of the reconstruction of Schoch (2006:fig. 4).
- **137 (exoccipitals):** 0 → 1. Figures and reconstruction based on the holotype (Schoch, 2006:figs. 1–2, 4) indicate well-exposed condyles in dorsal view.
- **237 (postparietal, length):** 1 → 0. The element is nearly square, more anteriorly extensive than the tabular (Schoch, 2006:figs. 2–4), and in stark contrast to the condition of zatracheids or highly nested amphibamiforms in which it is an anteroposteriorly short, transversely elongate rectangle (e.g., Witzmann & Schoch, 2006:fig. 3; Clack & Milner, 2009:fig. 9a; Sigurdsen & Bolt, 2010:fig. 3A).
- **278 (trunk intercentra):** ? → 0. As described and figured by Schoch (2006:35, figs. 5–6).
- **279 (trunk neural arch to spine):** ? → 0. As described and figured by Schoch (2006:36, figs. 5–6).
- **281 (haemal arches):** ? → 0. As figured by Schoch (2006:figs. 5–6).
- **282 (haemal arch, articulation):** ? → 0. As figured by Schoch (2006:figs. 5–6).
- **283 (haemal arch, length):** ? → 0. As figured by Schoch (2006:fig. 5).
- **284 (haemal arch, shape):** ? → 0. As figured by Schoch (2006:fig. 6).
- **301 (scapulocoracoid):** ? → 1. As described by Schoch (2006:36).
    - **300 (coracoid, foramina):** ? → -.
- **302 (humerus, deltopectoral crest):** ? → 0. As described by Schoch (2006:36).
- **303 (humerus, length):** ? → 1. As figured by Schoch (2006:fig. 5).
- **304 (radius:humerus):** ? → 0. As figured by Schoch (2006:fig. 5).
- **305 (olecranon process):** ? → 0. Schoch (2006:37) stated that this is unossified.
- **307 (femur):** ? → 0. As described by Schoch (2006:37).
- **313 (skull table):** 1 → 0. By comparison with other taxa coded for 313–1, it is felt that the condition of this taxon is closer to the majority of all taxa (coded for 313–0, ‘rectangular’). In particular, it is unclear how this taxon differs from other more longirostrine temnospondyls like *Archegosaurus*, *Platyoposaurus*, or *Australerpeton* (all coded for 313–0).

***Lyrocephaliscus euri***

- **85 (transverse tooth row, transvomerine):** 2 → 1. Save-Söderbergh (1935:fig. 5) and Schoch & Milner (2000:fig. 83) depicted only two transvomerine teeth. These cannot form a ‘V-shaped’ row.
- **89 (interpterygoid vacuities):** 0 → 1. Especially in the context of lissamphibians or lepospondyls, the condition of this taxon is more similar to all other temnospondyls than to outgroups (e.g., Mazin & Janvier, 1983:fig. 3).
- **110 (parasphenoid, plate, proportions):** 0 → 1. The basal plate is more like a quadrangle with the posterior end expanded such that the greatest width is at the posterior end and is about as long as the greatest length (e.g., Schoch & Milner, 2000:fig. 83).

- **111 (parasphenoid, plate, width):** 0 → 1. As with character 110; because of the long parasphenoid-pterygoid suture, there is only a very short region posterior to the basicranial articulation, and it is much wider than the parietals.
- **161 (transverse process, length):** ? → 0. This feature can be coded based on Mazin & Janvier (1983:24–27, figs. 9–10).
- **162 (neural spine, height):** ? → 1. As with character 161.
- **163 (intercentrum, shape):** ? → 0. As with character 161.
- **164 (intercentrum, width):** ? → 0. As with character 161.
- **168 (pleurocentrum, presence):** ? → 0. As with character 161.
- **169 (pleurocentrum, lateral surface):** ? → 1. As with character 161.
- **170 (pleurocentrum, ventral extension):** ? → 0. As with character 161.
- **172 (ribs, ventral extension):** ? → 1. As with character 161.
- **173 (ribs, uncinate blades):** ? → 0. As with character 161.
- **174 (ribs, uncinate spines):** ? → 0. As with character 161.
- **178 (clavicle, ventral blade):** ? → 0. This feature can be coded based on Mazin & Janvier (1983:27, fig. 11).
- **185 (interclavicle, anterior stylus):** ? → 1. This feature can be coded based on Mazin & Janvier (1983:27, fig. 12).
- **219 (lacrimal-naris):** 0 → 1. The lacrimal is widely separated from the naris (e.g., Schoch & Milner, 2000:fig. 83).
- **242 (premaxilla, tooth count):** ? → 0. As figured by Schoch & Milner (2000:fig. 83).
- **243 (maxilla, tooth count):** ? → 0. As figured by Schoch & Milner (2000:fig. 83).
- **251 (vomer, denticles):** ? → 1. As figured by Säve-Söderbergh (1936:fig. 5) and Schoch & Milner (2000:fig. 83).

***Callistomordax kugleri***

- **30 (interorbital distance):** 0 → 1. The interorbital distance is wider than the orbit in specimens and reconstructions figured by Schoch (2008a:figs. 1–5).
- **66 (tabular, extension):** 0 → 1. Schoch (2008a:89, figs. 1–2) stated that "its [the tabular's] blunt posterolateral end never extends posteriorly."
- **75 (quadrate and occipital condyles):** 1 → 2. Figures and reconstruction by Schoch (2008a:91, figs. 1–2, 5) show the occipital condyles projecting beyond the quadrates, as in most metoposaurids, which is further substantiated by the text ("the condyles reach a level well posterior to the occipital rim of the dermal skull and behind the quadrate condyles").
- **88 (ectopterygoid, fangs):** 1 → 0. Schoch (2008a:90) stated that this element "bears a pair of small tusks, succeeded posteriorly by a row of 12 much smaller teeth." This is reflected in the corresponding illustrations and reconstructions (figs. 1–2 therein).
- **237 (postparietal, length):** 1 → 0. The element is nearly square, more anteriorly extensive than the tabular (Schoch, 2008a:figs. 1–5), and in stark contrast to the condition of zatracheids or highly nested amphibamiforms in which it is an anteroposteriorly short, transversely elongate rectangle (e.g., Witzmann & Schoch, 2006:fig. 3; Clack & Milner, 2009:fig. 9a; Sigurdsen & Bolt, 2010:fig. 3A).
- **287 (atlas-axis intercentra):** ? → 0. Schoch (2008a:92) described this complex.
- **288 (atlas anterior centrum):** ? → 1. As with character 287.
- **289 (atlas centrum):** ? → 1. As with character 287.

- **293 (proatlantes):** ? → 1. The completeness of the holotype confers support for the absence of the proatlas.

***Metoposaurus krasiejowensis***

*Note that these changes are considered to be applicable regardless of whether *M. diagnosticus* or *M. krasiejowensis* (sensu Brusatte et al., 2015) is employed.

- **66 (tabular, extension):** 0 → 1. like in the metoposauroid *Callistomordax*, the tabular horn of *Metoposaurus* points far more laterally than it does posteriorly (e.g., Fraas, 1889:taf. XII; Sulej, 2002:figs. 3–4; Sulej, 2007:figs. 1B, 5, 13; Brusatte et al., 2015:figs. 2B, 11).
- **168 (pleurocentra):** 0 → 1. Pleurocentra are widely recognized to be absent in all metoposaurids except perhaps *Dutuitosaurus ouazzoui* (e.g., Dutuit, 1972, 1976; Hunt, 1993, but see Sulej, 2007, for comments on that particular taxon).
- **214 (skull to trunk):** ? → 0. A partial skeleton figured by Fraas (1889:taf. 14–15) and by Hellrung (2003:fig. 22) would have been sufficient to code this for *M. diagnosticus*; the reconstruction of *M. krasiejowensis* by Sulej (2007) is based on articulated skeletons of *Dutuitosaurus ouazzoui* and is considered reliable for coding.
- **237 (postparietal length):** 1 → 0. The postparietals are typically longer than they are wide (e.g., Fraas, 1889:145, taf. XII; Sulej, 2002:figs. 3–4; Sulej, 2007:figs. 1B, 4–14; Brusatte et al., 2015:figs. 2B, 3, 11).
- **242 (number of premaxillary teeth):** 1 → 0. Like other highly nested stereospondyls, metoposaurids have a relatively high tooth count, with more than 15 positions on the premaxilla (e.g., Sulej, 2007:34).
- **278 (trunk intercentra):** ? → 0. The presence of intercentra was established for *M. diagnosticus* by Fraas (1889) and Warren & Snell (1991), among others, and is well-documented in isolation for *M. krasiejowensis* (Sulej, 2007).
- **293 (proatlas):** ? → 1. The proatlas has never been identified in articulated or disarticulated material.
- **313 (skull table):** ? → 0. As with other temnospondyls.
- **314 (prefrontals):** ? → 0. As with all metoposaurids.
- **315 (postfrontals):** ? → 0. As with all metoposaurids
- **317 (zygokrotaphy):** ? → 0. As with all metoposaurids

***Wetlugasaurus angustifrons***

- **31 (frontal-nasal, length):** 0 → ?. Marked intraspecific variation (e.g., Damiani, 2001:fig. 37A; Schoch & Milner, 2000:fig. 73) complicates assessment of this character. If the common midline suture is used as the reference, the frontal is longer than the nasal (31–0). If the greatest longitudinal length is used, the condition is polymorphic. Because of uncertainty about how the feature was measured, and the lack of congruency between methods, this was changed to unknown.
- **39 (prefrontal-frontal):** 0 → 0&1. As with character 39, variation in the length of the frontal produces polymorphism.

***Parotosuchus nasutus***

- **113 (cultriform process, structure):** 0 → 1. Schoch (2018:118) noted a median ventral ridge extending from the basal plate onto the cultriform process and extending for the posterior half of the process.

- **168 (pleurocentra):** 0 → ?. Schoch (2018:119) stated that postcrania are difficult to confidently identify due to disarticulation at the type locality and the presence of multiple temnospondyl taxa and that the vertebrae are entirely unknown.
- **170 (pleurocentrum, ventral extension):** 0 → ?. As with character 168; note that character 169 (pleurocentrum, lateral surface) was originally uncoded.
- **182 (interclavicle, proportions):** ? → 0. As figured by Schoch (2018:fig. 9B).
- **183 (interclavicle, posterior margin):** ? → 0. As with character 182.
- **185 (interclavicle, anterior stylus):** ? → 1. As with character 182.
- **186 (interclavicle, posterior stylus):** ? → 1. As with character 182; the more discrete offset along the lateral margin warrants coding this for state 1 rather than state 0 as in other stereospondyls.
- **202 (ilium, tip):** 1 → ?. There is no indication that the ilium is known (Schoch, 2018), and the other four iliac characters were originally uncoded.
- **266 (Meckelian fossa):** ? → 0/1. There is at least the one typical large opening ventral to the adductor fossa.

***Eryosuchus garjainovi***

- **30 (interorbital distance):** 0 → 1. Per Schoch & Milner (2000:fig. 97).
- **39 (prefrontal, frontal):** 0 → 0&1. Per Schoch & Milner (2000:fig. 97).
- **40 (prefrontal, anterior end):** 1 → 0. The prefrontal has a distinct triangular end that tapers to a point where it meets the lacrimal and the nasal.
- **110 (parasphenoid plate):** 1 → 0. The basal plate is distinctly longer than it is wide.
- **118 (pterygoid, exoccipital):** 0 → 1. Per Schoch & Milner (2000:fig. 97).

***Paracylotosaurus davidi***

- **75 (quadrate and occipital condyles):** 1 → 0. Figures by Watson (1958:fig. 1) and Damiani (2001:fig. 21) show the occipital condyles situated slightly anterior to the quadrate condyles, contrary to Watson’s description that the condyles lie “essentially in the same transverse plane” (p. 244 therein).
- **91 (anterior palatal opening):** 1 → 0. There is no indication of any perforations for the symphyseal fangs (Watson:1958:fig. 1).
- **92 (anterior palatal opening(s)):** 0 → -. As with the change to character 91.
- **168 (pleurocentra):** 0 → 1. Watson (1958:247) stated that the pleurocentra were entirely cartilaginous. While their position can be ascertained from the articulated holotype, this does not allow them to be coded for any pleurocentra character given the unossified state.
- **169 (pleurocentrum, lateral surface):** 1 → -. As with character 168.
- **170 (pleurocentrum, ventral extension):** 0 → -. As with character 168.
- **197 (manual count):** 1 → 0. Watson (1958:251) noted five manual digits. Konietzko-Meier et al. (2020:1152–1153) also commented on this with respect to the present mounting of the holotype with four fingers as perhaps a modification from the original mounting with five fingers in light of the now disproven perception that all temnospondyls have four manual digits.
- **271 (coronoids):** ? → 0. Watson (1958:245) described three coronoids.
- **272 (coronoid, teeth):** ? → 1. As with character 272.
    - **273 (coronoid, teeth, size):** ? → -.

- **287 (atlas-axis intercentra):** ? → 0. Watson (1958:245–246, fig. 8) described and figured this complex.
- **288 (atlas anterior centrum):** ? → 1. As with character 287.
- **289 (atlas centrum):** ? → 0. As with character 287.

***Eocyclotosaurus appetolatus***

- **31 (frontal-nasal, length):** 0 → 1. As shown by Rinehart et al. (2015:fig. 2), the frontal is shorter than the nasal whether length is measured along the midline suture or as the greatest longitudinal length. In the holotype of *E. wellesi*, the elements are equal in length along the midline, but the nasal is longer in its greatest length (Schoch, 2000:fig. 1). Schoch (p. 394 therein) described the nasals as slightly longer than the frontals.
- **50 (postorbital, postfrontal):** 0 → -. The parietal is longer than the postfrontal and the postorbital, whereas the supratemporal is shorter than at least the postorbital.
- **110 (parasphenoid plate):** 1 → 0. The basal plate is distinctly longer than it is wide in both *E. wellesi* (Schoch, 2000:fig. 2) and *E. appetolatus* (Rinehart et al., 2015:fig. 3B).
- **145 (Meckelian window):** 2 → 1. The opening is only about 75% of the length of the adductor chamber (Rinehart & Lucas, 2016:34, fig. 34A).
- **164 (intercentrum, width):** 0 → 1. The notochordal canal is narrow (Rinehart & Lucas, 2016:37, fig. 39).
- **168 (pleurocentrum, presence):** 0 → 1. Rinehart & Lucas (2016:35) note that no pleurocentra were identified in the Tecolotito bonebed and inferred that they were unossified given the density of elements and overall representation of other skeletal elements. Schoch (2000) did not report any postcrania for *E. wellesi*, so this is not considered to be a possible interspecific difference.
- **169 (pleurocentrum, lateral surface):** 1 → -. As with character 168.
- **170 (pleurocentrum, ventral extension):** 0 → -. As with character 168.
- **191 (humerus, adult shaft):** ? → 1. As described by Rinehart & Lucas (2016:50, fig. 57).
- **194 (humerus, condyles):** ? → 1. As with character 191.
- **198 (ilium, shaft):** ? → 0. As described by Rinehart & Lucas (2016:47, fig. 53).
- **199 (ilium, dorsal end):** ? → 0. As with character 198.
- **200 (ilium, height):** ? → 0. As with character 198.
- **201 (ilium, orientation):** ? → 0. As with character 198.
- **203 (pubis):** ? → 0. Given the extensive sampling of the Tecolotito bonebed and the identification of ilia and ischia but no pubes, it can be reasonably inferred that this element was not ossified, as in most other stereospondyls.
- **204 (femur):** ? → 1. As described by Rinehart & Lucas (2016:50–51, fig. 60).
- **205 (femur, trochanter):** ? → 1. As with character 204.

***Cyclotosaurus intermedius***

*Note that these changes are considered to be applicable regardless of whether *C. robustus* or *C. intermedius* is employed.

- **75 (quadrate and occipital condyles):** 1 → 2. The occipital condyles extend beyond the quadrates (Schoch, 2008a:fig. 5C).

***Mastodonsaurus giganteus***

- **15 (internarial distance):** 0 → 1. The narrow interorbital distance of this taxon is well-documented (Schoch, 1999:25, fig. 1)

- **141 (postglenoid area (types)):** 0 → ?. Schoch (1999:80) stated that the PGA of this taxon is highly variable and displays characteristics of both types within a single jaw (i.e. it cannot be discretized into this binary); for this reason, I code the character as unknown rather than as polymorphic.
- **196 (carpals):** 0 → 1. Schoch (1999:130, fig. 45) described and figured at least two proximal carpals, which are known from a partial articulated skeleton from Gaildorf.
- **287 (atlas-axis intercentra):** ? → 0. Schoch (1999:98–99, figs. 29–30) described and figured this complex.
- **288 (atlas anterior centrum):** ? → 1. As with character 287.
- **289 (atlas centrum):** ? → 1. As with character 287.
- **290 (atlas neural arch):** ? → 2. As with character 287.
- **291 (atlas parapophyses):** ? → 2. As with character 287.
- **292 (atlas neural arch):** ? → 1. As with character 287.
- **293 (proatlantes):** ? → 1. Schoch (1999:95) states that the proatlas is absent.

***Chinlestegophis jenkinsi***

- **21 (lacrimal):** ? → 0. Regardless of whether the lacrimal indeed fused with the maxilla or was 'lost' through some other process, there is no dorsal expression of the lacrimal or identification of a distinct lacrimal through tomography. This is considered sufficient to restore the original code of Pardo et al. (2017a), which was changed by Schoch et al. (2020). The same character is not restored for *Rileymillerus cosgriffi* because CT analysis has not been performed for this taxon and because of suspicions over identity of the 'nasal' in this taxon (see comments on this taxon). Correspondent with this change, codes for characters 22–24 were updated to '-' from '?'.
- **39 (prefrontal, frontal):** 1 → 0. The prefrontal extends well beyond the frontal (Pardo et al., 2017a:fig. 1I).
- **56 (semilunar flange):** 1 → 0. The supratemporal is widely excluded from the otic notch and has no ventral flange (Pardo et al., 2017a:fig. 1I-J).
- **58 (jugal, anterior extent):** ? → 0. The presence of an LEP truncates the jugal anteriorly (Pardo et al., 2017a:fig. 1I-J). Note that this was originally left uncoded by Pardo et al. (2017a) and was not changed as a result of Schoch et al.'s (2020) interpretation of the LEP.
- **127 (LEP):** ? → 1. This code is restored from that of Pardo et al. (2017a), following their original interpretation. This was changed to unknown by Schoch et al. (2020), who stated that it might be a lacrimal, as has previously suggested for similar exposures in some trematosaurs (Schoch, 2006, 2008), but those studies have not demonstrated that an element in the same position is either definitively a lacrimal or definitively not a LEP (e.g., via CT analyses to identify the nasolacrimal duct passage or to assess the articulation of the palate with the roof). Furthermore, the aforementioned studies did not address the possibility that other temnospondyl clades without lacrimals might have similar overplating, and if *Chinlestegophis jenkinsi* were to be left coded as unknown for this character, all other taxa without lacrimals should have been treated the same. I do not opt for this approach here and restore the original code on the basis that there was no unequivocal basis for overturning it.
- **237 (postparietal, length):** 1 → 0. The element is nearly square, more anteriorly extensive than the tabular (Pardo et al., 2017a:fig. 1I), and in stark contrast to the condition of zatracheids or highly nested amphibamiforms in which it is an

anteroposteriorly short, transversely elongate rectangle (e.g., Witzmann & Schoch, 2006:fig. 3; Clack & Milner, 2009:fig. 9a; Sigurdsen & Bolt, 2010:fig. 3A).

- **271 (coronoids):** ? → 0. This is another strange omission from the original coding by Pardo et al.; they identified and figured three coronoids.

***Rileymillerus cosgriffi***

- **57 (jugal, ventral process):** ? → 1. Bolt & Chatterjee (2000:676) mentioned a "jugal processus alaris" several times and code this feature as present in their matrix (appendix 1 therein).
- **75 (quadrate and occipital condyles):** 1 → 2. The occipital clearly extend past the level of the quadrate condyles (Bolt & Chatterjee, 2000:figs. 2.1–2.2).
- **85 (transverse tooth row, transvomerine):** 0 → 2. Bolt & Chatterjee (2000:678) described two sets of vomerine teeth (exclusive of 'fangs'): one that extends posteriorly from a position medial to the anterior fang position and that "continues near the intervomerine suture" and the other than borders the medial margin of the internal naris. The former is vaguely illustrated in their reconstruction (fig. 2.2 therein) and forms a sharp V-shaped profile, similar to the condition seen in some trematosaurs (including *Almasaurus habbazi*; Dutuit, 1976:figs. 83–84), with the same point of origination and the same orientation towards the midline suture.
- **108 (parasphenoid):** 0 → ?. Because the sutures in this region of the palate are entirely unknown (Bolt & Chatterjee, 2000:fig 2.2) and the parasphenoid is heavily damaged, this character is uncoded.
- **114 (cultriform process, outline):** 1 → 0. Based on how the character was originally coded by Schoch (2013), it refers to the condition of eryopoids where there is a prominent and abrupt lateral expansion anterior to the base of the cultriform process; it does not refer to a gradual broadening onto the base, as in this taxon.
- **118 (pterygoid, exoccipital):** 1 → ?. As with character 107.
- **120 (pterygoid, squamosal):** 0 → 1. Bolt & Chatterjee (2000:675-676) discussed whether the term 'palatoquadrate fissure' was applicable to this taxon, but since the character refers only to a 'fissure,' it is coded as state 1 since there is a wide separation between the pterygoid and the squamosal.
- **127 (LEP):** ? → 1. This code is restored from that of Pardo et al. (2017a), following the original interpretation of Bolt & Chatterjee (2000:674, fig. 2). This was reversed by Schoch et al. (2020), and it was previously argued that it is merely a short lacrimal (Schoch, 2008b:83–84) but compelling evidence to overturn the original interpretation (e.g., CT data) or to prove that exposures in similar positions in other taxa are lacrimals and not LEPs (e.g., identifying the nasolacrimal duct) have not been provided. The position is in line with where an LEP occurs in dissorophoids and dvinosaurs, which already represent two independent acquisitions of this feature. Furthermore, past authors have not addressed the possibility that other temnospondyl clades without lacrimals might have similar overplating, and if *Chinlestegophis jenkinsi* were to be left uncoded for this character, all other taxa without lacrimals (e.g., *Siderops*, *Laidleria*) should have been treated the same. I do not opt for this approach here and restore the original code on the basis that there was no unequivocal basis for overturning it.
- **145 (Meckelian window):** 0 → ?. Bolt & Chatterjee, 2000:677, fig. 4) noted that they were unable to locate any Meckelian opening and that only the posterior quarter of the lower jaw is exposed and well-preserved.

- **237 (postparietal, length):** 1 → ?. The element’s lateral margins are not discernible on the roof (Bolt & Chatterjee, 2000:675, figs. 2.1, 3), but it was clearly a relative anteroposteriorly long element, concurrent with the skull’s general proportions. Mapping the postparietal-tabular suture from the occiput onto the skull roof would indicate an element far longer than it is wide (favoring 237–0), but because it cannot be assumed that the postparietal’s lateral margin was longitudinally oriented, this character is recoded as unknown.
- **251 (vomer, denticles):** 0 → 1. Bolt & Chatterjee (2000:678) stated that no palatal denticles were identified.
- **271 (coronoids):** ? → 0/1/2. At least one coronoid was identified.

***Triadobatrachus massinoti***

- **39 (prefrontal, frontal):** 1 → 0. The prefrontal extends beyond the frontal (e.g., Rage & Roček, 1989:figs. 2, 4; Ascarrunz et al., 2016:fig. 4). Alternatively, the character could be coded as inapplicable because the frontal and parietal are co-ossified in this taxon.

***Leptodactylus mystacinus***

- There are no additional changes to those made at the character level.

***Xenopus tropicalis***

- There are no additional changes to those made at the character level.

***Karaurus sharovi***

- **238 (squamosal-jugal):** 1 → ?. There is no jugal in this taxon.

***Kokartus honorarius***

- **223 (frontals, paired):** ? → 0. As figured and described by Skutschas & Martin (2011:826:fig. 4).
- **228 (posterior skull margin):** ? → 1/3. As figured and described by Skutschas & Martin (2011:826:fig. 4). There is a very slight indentation in the posterior margin of the parietal, but it is not as ‘undulating’ as in taxa originally coded for this state, so partial uncertainty is newly coded.
- **233 (parietal-squamosal):** ? → 1. As figured and described by Skutschas & Martin (2011:827:figs. 3, 5, 9).
- **249 (pterygoid, denticles):** ? → 1. As described by Skutschas & Martin (2011:827).
- **250 (pterygoid, teeth):** ? → 0. As with character 249.
- **271 (number of coronoids):** ? → 0/1/2. Only one coronoid was described in articulation with the dentary (Skutschas & Martin, 2011:827); because no complete lower jaw is known and only three elements have been identified, this is coded as partial uncertainty.
- **272 (coronoid, teeth):** ? → 0. As with character 271.
- **273 (coronoid, teeth, size):** ? → 2. As with character 271.
- **279 (trunk neural arch to centrum):** ? → 2. As figured by Skutschas & Martin (2011:fig. 8).
- **312 (parietal width):** ? → 0. As figured by Skutschas & Martin (2011:figs. 4, 9)

***Ambystoma opacum***

- There are no additional changes to those made at the character level.

***Hynobius* spp.**

- There are no additional changes to those made at the character level.

***Cryptobrachus alleghenienisis***

- There are no additional changes to those made at the character level.

***Ichthyophis bannanicus***

- **114 (cultriform process, outline):** 1 → 0. Based on how the character was originally coded by Schoch (2013), it refers to the condition of eryopoids where there is a prominent and abrupt lateral expansion anterior to the base of the cultriform process; it does not refer to a gradual broadening onto the base, as in this taxon.
- **271 (number of coronoids):** 0 → 2. Based on the embryological data suggesting that there is only a single coronoid in gymnophionomorphs (that does not clearly represent fusion of the plesiomorphic three coronoids found in most temnospondyls), I recoded this character (e.g., Wake & Hanken, 1982; Müller et al., 2005; Müller, 2006; Theska et al., 2019). Alternatively, the character could be coded as unknown since this specific taxon has not been studied developmentally.

***Epicrionops bicolor***

- **114 (cultriform process, outline):** 1 → 0. Based on how the character was originally coded by Schoch (2013), 114–1 refers to the condition of eryopoids where there is a prominent and abrupt lateral expansion anterior to the base of the cultriform process (e.g., Sawin, 1941:pls. 2–3; Boy, 1990:fig. 3B); it does not refer to a gradual broadening onto the base, as in this taxon (e.g., Reiss, 1996:fig. 2D).
- **271 (number of coronoids):** 0 → 2. Based on the embryological data suggesting that there is only a single coronoid in gymnophionomorphs (that does not clearly represent fusion of the plesiomorphic three coronoids found in most temnospondyls), I recoded this character (e.g., Wake & Hanken, 1982; Müller et al., 2005; Müller, 2006; Theska et al., 2019). Alternatively, the character could be coded as unknown since this specific taxon has not been studied developmentally.

***Eocaecilia micropodia***

- **39 (prefrontal, frontal):** 1 → 0. The prefrontal extends beyond the frontal.
- **266 (number of Meckelian fossae):** ? → 1. Jenkins et al. (2007:319, 322, figs. 24, 27) identified one opening that they termed the "intramandibular foramen" and which they suggested was occupied by a remnant of Meckel's cartilage, as in extant gymnophionans (coded for 266–1). More anteriorly situated foramina all occur just below the lingual tooth row and likely relate to tooth replacement, not to Meckel's cartilage.
- **271 (number of coronoids):** 0 → 2. Based on the embryological data suggesting that there is only a single coronoid in gymnophionomorphs (that does not clearly represent fusion of the plesiomorphic three coronoids found in most temnospondyls), I recoded this character (e.g., Wake & Hanken, 1982; Müller et al., 2005; Müller, 2006; Theska et al., 2019). Alternatively, the character could be coded as unknown since this specific taxon cannot be studied developmentally.

### Appendix 3. Previous coding modifications.

#### Appendix 3.1. Coding modifications of Dilkes (2015).

Dilkes (2015:appendix 1) made several dozen coding changes that he documented and justified in detail. As noted in the main text of this study and previously by Marjanović & Laurin (2019:116), most of these were adopted by Pardo et al. (2017a) without attribution. However, there were 12 individual cell coding modifications and several character-level coding modifications made by Dilkes that were not propagated by Pardo et al. All of Dilkes' modifications are repeated here, though without his detailed justifications, in order to demonstrate which changes have been propagated and how I approached those changes that were not propagated by Pardo et al. Note that some changes that were propagated by Pardo et al. may have been overturned here, either based on my own perception of the data or based on modification of the character.

**Changes that were adopted, without attribution, by Pardo et al. (2017a) are listed in bold in the following sections 3.1.1. and 3.1.2**. Those changes that were not adopted here have annotations explaining whether they were implemented in this study. An asterisk (*) is used to denote a coding modification based only on Dilkes' personal observations.

##### Appendix 3.1.1. Character-level modifications of Dilkes (2015)

Character numbering refers to the original numbering of Schoch (2013), which is identical to that of Pardo et al. (2017a). There is a frameshift of –1 in the Dilkes' matrix after original character 32.

- 13: "wording changed by deletion of "separating the two alary processes" because presence of alary processes not required (e.g., *Acroplous*)."
    - This character was independently modified along with original character 14 (internarial fontanelle), and the revised version does not contain this specifier to the alary process.
- 26: "The old character concerned the location of the orbits relative to the midline of the skull as an expression of the width of the jugals. Wide jugals for laterally placed orbits and narrow jugals for medially placed orbits. However, there is no quantification of jugal height to allow determination as either wide or narrow. The new character expresses height of the jugal relative to the height of the maxilla beneath the orbit. A jugal with a height greater than that of the maxilla would be equivalent to a wide jugal in the original character."
    - This modification captures the longstanding problem of discretizing continuous data into qualitative states. Because the character was partially recoded here to ensure consistency in treatment of state 2, the quantitative specifier was not added.
- 33, 34: These characters were combined.
    - In keeping with my conservative approach of not modifying characters unless expressly necessary, this modification was not made.
- **34: "All taxa that, according to coding in character 33, lack lateral line sulci are recoded as - (inapplicable) rather than the original 0 code indicating presence of the sulci."**
    - These modifications were maintained, with the addition of the uncoding of *Micropholis*.
- 69: "The order of the states was reversed to match the coding in the data matrix."
    - As noted in Appendix _____, while most codes were indeed reversed, a few taxa were correctly coded as is, so the entire character was recoded with the defined states left in their original configuration.
- Addition of nine new characters (212–220 in Dilkes, 2015)
    - These characters all refer to the carpals and tarsals, which are rarely preserved in temnospondyls. They were added by Dilkes based on the focus of his study on the

carpus and tarsus but are considered here to introduce a high degree of uncertainty due to a high proportion of missing data and were therefore excluded.

##### Appendix 3.1.2. Taxon-level modifications of Dilkes (2015)

Thirty-six of the 48 modifications (75%) listed below were adopted by Pardo et al. (2017a), without comment or attribution. Of those 36, eight were strictly made by Dilkes based on his personal observations (all of *Dissorophus multicinctus*).

*Proterogyrinus scheelei*

- 195: 0 → 1
    - This change was independently made here on the same basis of partial preservation of the carpus.

*Greererpeton burkemorani*

- 195: 0 → 1
    - This change was independently made here on the same basis of partial preservation of the carpus.

*Balanerpeton woodi*

- 28: 0 → 0&1
    - Dilkes argued that this should be coded as polymorphic following Sigurdsen & Bolt (2010), who coded it as such because the palpebral ossifications are not as developed as in dissorophoids. However, a polymorphism is not the correct approach here (partial uncertainty would be better); either splitting the character states further or lumping all palpebral ossifications together is more appropriate. The latter is adopted here.

*Nigerpeton ricqlesi*

- **33: 1 → 0**

*Thabanchuia oomie*

- 34: 0 → -
    - This taxon was not propagated by Pardo et al. (2017a), and this coding modification was made independently here.

*Tupilakosaurus wetlugensis*

- 34: 0 → -
    - This taxon was not propagated by Pardo et al. (2017a), and this coding modification was made independently here.

*Trimerorhachis insignis*

- **35: 0 → 1**

*Neldasaurus wrightae*

- **35: 0 → -**

*Isodectes obtusum*

- **35: 0 → -**

*Acroplous vorax*

- **10: 1 → 0**
- **13: 0 → 1**
- **34: 0 → -**
- **35: 0 → -**

*Dvinosaurus primus*

- **35: 0 → -**

*Iberospondylus schultzei*

- **33: 1 → 0**
- **35: 0 → -**

*Doleserpeton annectens*

- **146: ? → 0**
- **147: ? → 0**
- **157: ? → 1**

*Gerobatrachus hottoni*

- **33: 1 → ?**
- **80: 1 → 0**
- 81: ? → 0
    - The presence of pedicelly in *Gerobatrachus* is contentious (e.g., Marjanović & Laurin, 2008; Sigurdsen & Green, 2011). In the absence of any new data beyond the brief original description, the code is left as unknown here (as with Marjanović & Laurin).
- **195: 1 → ?**

*Cacops* spp.

- **31: 0 → 1**
- **146: ? → 0**
- **147: ? → 0**

*Broiliellus texensis*

- *80: ? → 0.
    - This change was adopted here based on Dilkes' personal observation.

*Dissorophus multicinctus*

- ***13: 0 → 1**
- ***28: ? → 1**
- ***29: 1 → 0**
- ***74: 2 → 0**
- ***80: ? → 0**
- ***87: ? → 1**
- ***146: ? → 0**
- ***147: ? → 0**
- *157: 2 → 0
    - Based on Dilkes' comments, the revised code should have been 157–1 (as it is in Pardo et al., 2017a, and subsequent derivates). The code of Pardo et al. was adopted here.
- **158: 1 → 0**
- *183: 0 → ?
    - This change was adopted based on Dilkes' personal observation (it is considered sufficient to overturn the original code).

*Acheloma cumminsi*

- **13: 0 → 1**

*Phonerpeton pricei*

- **146: ? → 0**

- **147: ? → 0**
- **161: ? → 0**

*Ecolsonia cutlerensis*

- **146: ? → 0**
- **147: ? → 0**
- **195: 1 → ?**

*Archegosaurus decheni*

- 195: 0 → 1.
  - Dilkes' treatment is accepted here based on his justification.
- 205: 1 → 0
  - Dilkes' treatment is accepted here based on his justification.

*Siderops kehli*

- 1 → ?.
  - This change was independently made here on the same basis of incomplete preservation.

#### Appendix 3.2. Coding modifications of Pardo et al. (2017a)

As previously noted, Pardo et al. (2017a) did not indicate that they made any coding modifications and stated that this matrix was derived from Schoch (2013) without attribution of many changes made by Dilkes (2015) that were adopted. Because some of these changes were based strictly on Dilkes' personal observations of specific specimens, it seems unlikely that some or all of these changes were made independently.

Additionally, while in the process of identifying unsubstantiated codes and comparing them to previous matrices, I inadvertently identified additional coding modifications that were made by Pardo et al. to previous codes (these were also not documented). As a result, some of the coding modifications made here to cells originally coded by Schoch (2013) actually represent 'reversions' rather than 'corrections.' Because Pardo et al. did not note any changes to previously coded cells, the omission of documentation of any single coding change cannot be attributed to accidental omission. The failure to document any of these coding changes is problematic in how it misrepresents the dataset as a derivation of an existing one, and coding changes were not evenly distributed across characters or across taxa (e.g., a relatively large number of codes were changed for *Lapillopsis*). Subsequently, I compared the original codes of Schoch (2013) with the equivalent codes of Pardo et al. (2017a) and detail additional undocumented changes here.

Of the following changes, 66 were algorithmically significant (i.e. not changing inapplicable to unknown). There were no additional changes beyond those noted by Dilkes (2015) for the following taxa: *Proterogyrinus, Edops, Cochleosaurus, Chenoprosopus, Dendrerpeton, Capetus, Iberospondylus, Acanthostomatops, Micromelerpeton, Limnogyrinus, Apateon, Micropholis, Dissorophus, Broiliellus, Acheloma, Phonerpeton, Onchiodon, Eryops, Sclerocephalus, Glanochthon, Platyoposaurus, Australerpeton, Rhineceps, Uranocentrodon, Lydekkerina, Chomatobatrachus, Edingerella, Benthosuchus, Trematosaurus, Trematolestes, Lyrocephaliscus, Callistomordax, Metoposaurus, Parotosuchus, Paracyclotosaurus, Cyclotosaurus,* and *Mastodonsaurus*. This encompasses all sampled micromelerpetids, branchiosaurids, eryopoids, sclerocephalids, lydekkerinids, trematosaurs, and capitosaurs. The most extensively recoded taxa were *Broomistega* and *Lapillopsis*; the former was likely based on the high-resolution CT scan of a complete skeleton that was published but not fully described by Fernandez et al. (2014) based on all new codes being for postcranial characters (Shishkin & Rubidge, 2000, only very briefly described and figured a few postcranial elements). The rationale for extensive modifications to *Lapillopsis*, many of which were reverting original codes of Schoch (2013) that are substantiated by the most recent study (Yates, 1999), is uncertain.

*Greererpeton burkemorani*
- 52: - → ?

*Adamanterpeton ohioensis*
- 30: 1 → 0

*Nigerpeton ricqlesi*
- 139: 1 → 0
- 140: ? → 1

*Balanerpeton woodi*
- 28: 0 → 1.
    - This was a cell where Dilkes (2015) modified the code (to a polymorphism in this instance), but this change was not propagated by Pardo et al. (2017a), who instead independently modified it.
- 112: 0 → 2

*Neldasaurus wrightae*
- 52: - → 0

*Acroplous vorax*
- 52: - → ?
- 112: 0 → 2

*Isodectes obtusus*
- 52: - → ?

*Zatrachys serratus*
- 112: 0 → 2

*Platyrhinops lyelli*
- 26: 0 → 2
- 112: 0 → 2

*Amphibamus grandiceps*
- 26: 0 → 2
- 188: 0 → ?

*Doleserpeton annectens*
- 26: 0 → 2

*Gerobatrachus hottoni*
- 26: 0 → 2
- 112: 0 → 2
- 212: 0 → ?

*Cacops morrisi* + *C. aspidephorus*
- 31: 0 → 1
- 147: ? → 0
- 148: ? → 0

*Ecolsonia cutlerensis*

- 34: 0 → ?
- 147: ? → 0
- 148: ? → 0

*Archegosaurus decheni*

- 187: 0 → ?

*Lapillopsis nana*

- 15: 0 → 1
- 19: 0 → ?
- 20: 0 → 1
- 21: 0 → ?
- 22: 0 → ?
- 23: 0 → ?
- 24: 0 → ?
- 35: 0 → ?
- 36: 0 → ?
- 42: 0 → 1
- 43: 0 → ?
- 45: 0 → 1
- 52: 2 → 0
- 55: 0 → 1
- 58: 0 → 1
- 59: 0 → ?
- 186: ? → 1
- 187: ? → 0
- 188: ? → 0
- 189: ? → 0

*Peltobatrachus pustulatus*

- 163: - → ?
- 169: - → ?
- 212: 0 → ?

*Broomistega putterilli*

- 191: ? → 1
- 192: ? → 1
- 193: ? → 0
- 194: ? → 1
- 195: ? → 0
- 196: ? → 0
- 197: ? → 1
- 198: ? → 0
- 199: ? → 0
- 200: ? → 0
- 201: ? → 0
- 203: ? → 0
- 204: ? → 1

- 205: ? → 1
- 206: ? → 1
- 207: ? → 0

*Sangaia lavina*

- 52: - → ?
- 91: ? → 1
- 92: ? → 0
- 145: ? → 0

*Peltostega erici*

- 29: 0 → 1
- 75: 1 → ?

*Laidleria gracilis*

- 23: - → ?
- 52: - → ?
- 59: - → ?

*Siderops kehli*

- 23: - → ?
- 52: - → ?
- 59: - → ?

*Batrachosuchus/Bathignathus*

- 23: - → ?
- 52: - → ?
- 59: - → ?

*Plagiosuchus pustuliferus*

- 17: 0 → 1
- 52: - → ?

*Gerrothorax pulcherrimus*

- 52: - → ?
- 169: - → ?

### Appendix 4. Literature sources for temnospondyl geographic occurrences.

**Table 2. Reference literature used to map the global geographic distribution of temnospondyl body fossils in Figure 2A of the main text.** The listed references are not intended to represent a comprehensive sample, either with respect to the temporal range of publications on material from a given region or the total taxonomic diversity known from said region. Especially for the richest collecting regions (e.g., United States, Germany), the cited literature are not fully representative of the breadth of taxa or of previous work. They are only meant to provide the necessary data for reproducing Figure 2A of the main text (i.e. literature providing clear and convincing documentation of temnospondyls from a given region) and are therefore skewed towards more recent literature, as this is more likely to have stronger documentation (e.g., high-quality photographs) and less likely to have become outdated by the most recent work.

| Country | References |
|---|---|
| Algeria | Lehman (1971); Jalil (1999); Nedjari et al. (2010); Dahoumane et al. (2016) |

| | |
|---|---|
| Antarctica | Barrett et al. (1968); Colbert & Cosgriff (1974); Cosgriff & Hammer (1984); Sidor et al. (2007, 2008, 2014); Gee & Sidor (2021a, 2021b); Gee et al. (2021) |
| Argentina | Rusconi (1948, 1950, 1951, 1953; 1955); Bonaparte (1963); Marsicano (1990, 1994, 1996, 1999, 2005) |
| Australia | Watson (1958); Cosgriff (1974); Warren & Hutchinson (1983); Warren & Black (1985); Warren et al. (1997, 2001, 2006); Warren & Marsicano (1998); Damiani (1999, 2000); Yates (1999, 2000); Nield et al. (2006) |
| Austria | Stur (1873); Sander & Meyer (1991) |
| Brazil | Dias & Barberena (2001); Dias-da-Silva et al. (2006, 2009, 2011, 2012); Dias-da-Silva & Ilha (2009); Ramos & Vega (2011); Dias-da-Silva & Dias (2013); Strapasson et al. (2014); Cisneros et al. (2015); Pacheco et al. (2017); Dias et al. (2020) |
| Canada | Steen (1934); Carroll (1967); Rieppel (1980); Milner (1982a, 1996); Godfrey et al. (1987); Hunt (1993); Holmes et al. (1995, 1998); Sues & Olsen (2015); Sues et al. (2021); Arbez et al. (2022) |
| China | von Huene (1958); Young (1963, 1966, 1973); Dong (1985); Li & Cheng (1999); Maisch et al. (2004); Liu & Wang (2005); Maisch & Matzke (2005); Skutschas et al. (2009); Liu (2016, 2018) |
| Czech Republic | Steen (1938); Sequeira & Milner (1993); Werneburg (1994, 2012); Milner & Sequeira (2003); Sequeira (2004, 2009); Milner et al. (2007); Klembara & Steyer (2012); Milner (2018); Schoch & Witzmann (2018); Werneburg et al. (2020) |
| Ethiopia | Warren et al. (1998) |
| France | Dutuit & Heyler (1994); Heyler (1994); Steyer et al. (1997, 2000a, 2000b); Werneburg & Steyer (1999); Werneburg et al. (2007); Werneburg (2019) |
| Germany | Boy (1972, 1994); Werneburg (1991); Schoch & Werneburg (1999); Schoch (1999, 2006, 2008a, 2009, 2011); Lucas & Schoch (2002); Milner & Schoch (2004); Schoch & Witzmann (2006; 2009a, 2009b); Witzmann (2006); Schoch & Seegis (2016); Werneburg et al. (2019) |
| Greenland | Säve-Söderbergh (1935); Nielsen (1954, 1967); Bjerring (1997, 1999); Jenkins et al. (2008); Marzola et al. (2017, 2018) |
| India | von Huene & Sahni (1958); Tripathi (1961a, 1961b, 1969); Chowdhury (1965, 1970); Chatterjee & Chowdhury (1974); Sengupta (1995, 2002, 2003); Werneburg & Schneider (1996); Mukherjee & Sengupta (1998); Yates & Sengupta (2002) |
| Ireland | Milner (1980); Sequeira (1996) |
| Israel | Mazin (1986); Zanon (1991) |
| Italy | Koken (1913); Ronchi & Tintori (1997) |
| Japan | Nakajima & Schoch (2011) |
| Jordan | Schoch (2011) |
| Kazakhstan | Lozovsky & Shishkin (1974); Shishkin (1986) |
| Kyrgyzstan | Nesov (1990); Averianov et al. (2008) |
| Lesotho | Dutuit & Ginsburg (1982); Steyer & Damiani (2005) |

| | |
|---|---|
| Luxembourg | Milner et al. (1996) |
| Madagascar | Swinton (1956); Lehman (1961, 1966, 1974); Janvier (1992); Hewison (1996); Steyer (2002, 2003); Maganuco & Pasini (2009); Maganuco et al. (2009, 2014); Fortuny et al. (2017, 2018) |
| Malawi | Watson (1962) |
| Mongolia | Nesov (1988); Shishkin (1991); Shishkin et al. (2000) |
| Morocco | Dutuit (1972a, 1972b, 1976); Jalil (1999); Steyer & Jalil (2009); Khaldoune et al. (2017); Buffa et al. (2019); Werneburg et al. (2019) |
| Namibia | Keyser (1973); Warren et al. (2001) |
| Niger | Sidor et al. (2005); Damiani et al. (2006); Steyer et al. (2006); Sidor (2013) |
| Norway | Woodward (1904); Wiman (1910, 1913, 1914, 1916, 1917); Säve-Söderbergh (1936, 1937); Nilsson (1942, 1943, 1946); Janvier (1983); Mazin & Janvier (1983); Kear et al. (2016); Slodownik et al. (2021) |
| Pakistan | Maisch (2020) |
| Poland | Maryanska & Shishkin (1996); Sulej (2002, 2007); Sulej & Majer (2005); Shishkin & Sulej (2009); Sulej & Niedźwiedzki (2013) |
| Portugal | Witzmann & Gassner (2008); Brusatte et al. (2015) |
| Russia | Amalitzky (1924); Efremov (1933); Konzhukova (1955); Ochev (1966, 1972); Shishkin (1973, 1986, 1996); Gubin (1980, 1991, 1993); Shishkin et al. (2000); Werneburg (2009); Novikov (2007, 2010, 2012a, 2012b); Golubev & Bulanov (2018) |
| Slovenia | Lucas et al. (2008) |
| Spain | Gaete et al. (1993a, 1993b, 1994); Laurin & Soler-Gijón (2001, 2006); Fortuny et al. (2011); de Miguel Chaves et al. (2015) |
| South Africa | Kitching (1978); Shishkin & Welman (1994); Warren & Damiani (1999); Damiani et al. (2000, 2001); Hancox et al. (2000); Morales & Shishkin (2002); Damiani & Hancox (2003); Damiani & Kitching (2003); Damiani & Rubidge (2003); Damiani (2004) |
| Sweden | Hellrung (2003) |
| Switzerland | Leonardi (1952); Scheyer et al. (2022) |
| Tanzania | Haughton (1932); Panchen (1959); Howie (1970); Steyer et al. (2021) |
| Thailand | Ingavat & Janvier (1981); Suteethorn et al (1988); Buffetaut et al. (1994a, 1994b); Nonsrirach et al. (2021) |
| Tunisia | Soussi et al. (2017) |
| Turkey | Fortuny et al. (2015) |
| United Kingdom | Watson (1921); Paton (1975); Milner (1980); Milner et al. (1990); Milner & Sequeira (1993) |
| USA | Williston (1911); Branson & Mehl (1929); Gregory (1950); Langston (1953); Carroll (1964); Olson (1965, 1970); Milner (1982b); Hook & Baird (1986, 1994); Long & Murry (1995); Sequeira (1998); Milner & Schoch (2013); Gee et al. (2019b); Sues et al. (2021) |
| Uruguay | Marsicano et al. (2000); Piñeiro et al. (2007a, 2007b, 2007c, 2012) |

| | |
|---|---|
| Zambia | Chernin (1974); Chernin & Cosgriff (1975); Chernin (1977); Steyer et al. (2021) |

**Table 3. Reference literature used to map the North American (lower 48 American states and Canadian Maritimes) geographic distribution of temnospondyl body fossils in Figure 2B of the main text.** The listed references are not intended to represent a comprehensive sample, either with respect to the temporal range of publications on material from a given region or the total taxonomic diversity known from said region. Especially for the richest collecting regions (e.g., Texas), the cited literature are not fully representative of the breadth of taxa or of previous work. They are only meant to provide the necessary data for reproducing Figure 2B of the main text (i.e. literature providing clear and convincing documentation of temnospondyls from a given region) and are therefore skewed towards more recent literature, as this is more likely to have stronger documentation (e.g., high-quality photographs) and less likely to have become outdated by the most recent work. The asterisk (*) for the single source for occurrences in Nevada is a personal communication to the authors of Martz et al. (2017) and thus should be treated more conservatively.

| State / Province | References |
|---|---|
| Arizona | Brown (1933); Welles (1965, 1967, 1993); Welles & Estes (1969); Long & Murry (1995); Schoch (2000); Ruta & Bolt (2008) |
| Arkansas | Lane (1932) |
| Colorado | Lewis & Vaughn (1965); Pardo et al. (2017a); Gee et al. (2020) |
| Illinois | Gregory (1950); Milner (1982); Godfrey (2003); Mann & Gee (2019) |
| Kansas | Hotton (1959); Eaton (1973); Milner (1985b); Foreman (1990); Daly (1994); Englehorn et al. (2008); Martin & Merriam (2011); May et al. (2011) |
| Nebraska | Huttenlocker et al. (2005, 2007); Englehorn et al. (2008); May et al. (2011) |
| New Mexico | Langston (1953); Vaughn (1963, 1969); Berman (1973); Berman et al. (1985, 1987); Hunt et al. (1996); Schoch (1997); Lucas et al. (2010); Werneburg et al. (2010, 2021); Spielmann & Lucas (2012); Holmes et al. (2013); Rinehart et al. (2015) |
| New Jersey | Baird (1986) |
| Nevada | Martz et al. (2017)* |
| North Carolina | Leidy (1856); Colbert & Imbrie (1956); Heckert et al. (2012) |
| Nova Scotia | Steen (1934); Carroll (1967); Rieppel (1980); Milner (1982a, 1996); Godfrey et al. (1987); Hunt (1993); Holmes et al. (1995, 1998); Sues & Olsen (2015); Sues et al. (2021); Arbez et al. (2022) |
| Ohio | Olson (1970b); Vaughn (1971); Berman & Berman (1975); Hook & Baird (1984, 1986, 1994); Milner & Sequeira (1998, 2011); Clack & Milner (2010); Schoch & Milner (2021); Schoch et al. (2021) |
| Oklahoma | Bolt (1969, 1974a, 1977, 1980); Olson (1965, 1967, 1970a, 1972, 1991); Daly (1973); Sullivan et al. (2000); Ruta & Bolt (2006); Fröbisch & Reisz (2008, 2012); Anderson & Bolt (2013); Gee & Reisz (2018a, 2018b); Gee et al. (2019b) |
| Pennsylvania | Lucas & Sullivan (1996); Berman et al. (2010); Werneburg & Berman (2012); Sues & Schoch (2013); Gee & Jasinski (2021) |
| Prince Edward Island | Brink et al. (2013) |
| Texas | Williston (1910, 1911, 1914); Case (1922, 1931, 1932, 1935); Romer & Witter (1942); Sawin (1945); Romer (1947); Wilson (1948); |

| | |
|---|---|
| | Carroll (1964); Chase (1965); Hook (1993); Sequeira (1998); Bolt & Chatterjee (2000); Anderson et al. (2008); Milner & Schoch (2013) |
| Utah | Vaughn (1962, 1964); Welles (1967); Huttenlocker et al. (2018) |
| West Virginia | Murphy (1971); Lund (1975, 1976); Urban & Berman (2007); Werneburg & Berman (2012) |
| Wyoming | Branson (1905); Branson & Mehl (1929); Gee et al. (2018); Kufner & Gee (2021) |

**Table 4. Reference literature used to map the Russian geographic distribution of temnospondyl body fossils in Figure 2C of the main text.** The listed references are not intended to represent a comprehensive sample, either with respect to the temporal range of publications on material from a given region or the total taxonomic diversity known from said region. Especially for the richest collecting regions (e.g., Orenburg Oblast), the cited literature are not fully representative of the breadth of taxa or of previous work. They are only meant to provide the necessary data for reproducing Figure 2C of the main text (i.e. literature providing clear and convincing documentation of temnospondyls from a given region) and are therefore skewed towards more recent literature, as this is more likely to have stronger documentation (e.g., high-quality photographs) and less likely to have become outdated by the most recent work.

| **Federal subject** | **References** |
|---|---|
| Arkhangelsk Oblast | Amalitzky (1921, 1924); Bystrow (1938); Efremov & Viushkov (1955); Shishkin (1960a; 1966, 1967, 1973); Novikov (1990); Novikov & Orlov (1992); Novikov & Shishkin (1992) |
| Astrakhan Oblast | Sushkin (1927); Efremov (1932); Shishkin (1960b); Novikov (2007) |
| Ivanovo Oblast | Hartmann-Weinburg & Kuz'min (1936); Efremov (1940); Efremov & Viushkov (1955); |
| Kirov Oblast | Trautschold (1884); Efremov (1933); Shishkin (1966, 1967, 1980); Gubin (1987); Golubev (1995) |
| Komi Republic | Konzhukova (1956); Gubin (1983, 1984, 1986, 1987; 1993); Novikov (1986, 1990); Schoch & Milner (2000); Werneburg et al. (2020) |
| Kostroma Oblast | Getmanov (1989) |
| Krasnoyarsk Krai | Shishkin (1998); Werneburg (2009) |
| Mari El Republic | Golubev & Bulanov (2018) |
| Nizhny Novgorod Oblast | Efremov & Viushkov (1955); Shishkin (1961, 1968) |
| Novosibirsk Oblast | Schoch & Milner (2000) |
| Orenburg Oblast | Twelvetrees (1880); Konzhukova (1965); Vjushkov (1955); Ochev (1958, 1966); Shishkin (1966, 1967; 1986, 1987, 1994); Gubin (1980, 1984, 1987, 1989, 1991, 1993); Sennikov (1981); Getmanov (1982; 1989); Novikov (2012b, 2018); Sennikov & Novikov (2018) |
| Perm Krai | Zalesskiy (1947); Gubin (1980, 1986, 1991) |
| Primorsky Krai | Efremov & Viushkov (1955); Shishkin & Lozovsky (1979) |
| Republic of Bashkortostan | Eichwald (1848); Efremov (1937); Hartmann-Weinberg (1939); Efremov & Viushkov (1955); Konzhukova (1955); Ochev (1972); Gubin (1980, 1981); Shishkin (1986) |
| Republic of Tatarstan | Efremov & Viushkov (1955); Konzhukova (1955); Gubin (1991) |
| Sakha Republic | Shishkin & Vavilov (1985) |
| Samara Oblast | Getmanov (1989); Novikov (2012a, 2016) |

| | |
|---|---|
| Udmurt Republic | Golubev (1995) |
| Vladimir Oblast | Shishkin (1968); Gubin (2004) |
| Volgograd Oblast | Shishkin et al. (2000); Novikov (2010) |
| Vologda Oblast | Efremov (1929); Efremov & Viushkov (1955); Morkovin (2020) |
| Yaroslavl Oblast | Hartmann-Weinburg & Kuz'min (1936); Ivahknenko (1972); Getmanov (1986); Schoch & Milner (2000) |

**Table 5. Reference literature used to map the Brazilian geographic distribution of temnospondyl body fossils in Figure 2D of the main text.** The listed references are not intended to represent a comprehensive sample, either with respect to the temporal range of publications on material from a given region or the total taxonomic diversity known from said region. They are only meant to provide the necessary data for reproducing Figure 2D of the main text (i.e. literature providing clear and convincing documentation of temnospondyls from a given region) and are therefore skewed towards more recent literature, as this is more likely to have stronger documentation (e.g., high-quality photographs) and less likely to have become outdated by the most recent work.

| State | References |
|---|---|
| Maranhão | Price (1948); Cox & Hutchinson (1991); Cisneros et al. (2015); Marsicano et al. (2021) |
| Paraná | Barberena & Daemon (1974); Barberena (1980, 1998); Ramos & Vega (2011); Eltink et al. (2014, 2016); Azevedo et al. (2017); Dias et al. (2020) |
| Piauí | Cisneros et al. (2015); Marsicano et al. (2021) |
| Santa Catarina | Strapasson et al. (2014) |
| São Paulo | Chahud & Petri (2010) |
| Rio Grande do Sul | Lavina & Barberena (1985); Dias & Barberena (2001); Malabarba et al. (2003); Dias-da-Silva et al. (2005, 2006, 2009, 2011, 2012); Dias-da-Silva & Schultz (2008); Da-Rosa et al. (2009); Dias-da-Silva & Ilha (2009); Dias-da-Silva & Milner (2010); Eltink et al. (2017); Pacheco et al. (2017) |

**Table 6. Reference literature used to map the Chinese geographic distribution of temnospondyl body fossils in Figure 2E of the main text.** The listed references are not intended to represent a comprehensive sample, either with respect to the temporal range of publications on material from a given region or the total taxonomic diversity known from said region. They are only meant to provide the necessary data for reproducing Figure 2E of the main text (i.e. literature providing clear and convincing documentation of temnospondyls from a given region) and are therefore skewed towards more recent literature, as this is more likely to have stronger documentation (e.g., high-quality photographs) and less likely to have become outdated by the most recent work. The asterisk (*) for the single source for occurrences in Guizhou is a personal communication to the authors of Schoch & Milner (2014) and thus should be treated more conservatively.

| Province | References |
|---|---|
| Gansu | Li & Cheng (1999); Liu (2018) |
| Guizhou | Schoch & Milner (2014)* |
| Henan | Yang (1979) |
| Hubei | Liu & Wang (2005); Liu (2016) |
| Shanxi | von Huene (1958); Young (1963) |
| Shaanxi | Cheng (1980, 1981); Wu et al. (2022) |
| Sichuan | Dong (1985) |

| | |
|---|---|
| Xinjiang | Young (1966, 1973, 1978); Cheng (1980); Maisch et al. (2004); Maisch & Matzke (2005); Skutschas et al. (2009) |
| Yunnan | Sun (1962) |

**Table 7. Reference literature used to map the Australian geographic distribution of temnospondyl body fossils in Figure 2F of the main text.** The listed references are not intended to represent a comprehensive sample, either with respect to the temporal range of publications on material from a given region or the total taxonomic diversity known from said region. Especially for the richest collecting regions (e.g., Queensland), the cited literature are not fully representative of the breadth of taxa or of previous work. They are only meant to provide the necessary data for reproducing Figure 2F of the main text (i.e. literature providing clear and convincing documentation of temnospondyls from a given region) and are therefore skewed towards more recent literature, as this is more likely to have stronger documentation (e.g., high-quality photographs) and less likely to have become outdated by the most recent work.

| State | References |
|---|---|
| Queensland | Warren (1980; 1981; 1985); Warren & Hutchinson (1983, 1987, 1990); Warren & Black (1985); Damiani & Warren (1996); Yates (2000); Nield et al. (2006); Warren et al. (2006, 2009) |
| New South Wales | Watson (1958); Cosgriff (1972); Damiani & Warren (1997); Marsicano & Warren (1998); Warren & Marsicano (1998); Damiani (1999); Hart et al. (2021) |
| South Australia | Pledge (2013) |
| Tasmania | Cosgriff (1974); Cosgriff & DeFauw (1987); Warren & Marsicano (1998); Yates (1999); Akerman & Rozefelds (2011); Warren et al. (2011) |
| Victoria | Warren et al. (1997) |
| Western Australia | Cosgriff (1965, 1969); Cosgriff & Garbutt (1972); Damiani (2000) |

### Appendix 5. References.

Ahlberg, P. E., and J. A. Clack. 1998. Lower jaws, lower tetrapods–a review based on the Devonian genus *Acanthostega*. Earth and Environmental Science Transactions of The Royal Society of Edinburgh 89:11–46.

Akerman, K., and A. Rozefelds. 2011. Message in a bottle: a tale of two Triassic temnospondyl (labyrinthodont) femora from Tasmania. Papers and Proceedings of the Royal Society of Tasmania 145:5–7.

Amalitzky, V. P. 1921. [Dvinosauridae. North Dvina excavations of Prof. V. P. Amalitzky.] – Akademiya Nauk, Petrograd. 16 pp. [Russian]

Amalitzky, V. P. 1924. On the Dvinosauridae, a family of labyrinthodonts from the Permian of North Russia. Annals and Magazine of Natural History 9:50–64.

Anderson, J. S. 2005. On the skull of *Cacops aspidephorus* Williston (Tetrapoda; Temnospondyli; Dissorophidae) from the Lower Permian of Texas. Part 1: lower jaw anatomy. New Mexico Museum of Natural History & Science Bulletin 30:15.

Anderson, J. S. 2008. Focal review: the origin (s) of modern amphibians. Evolutionary Biology, 35(4):231–247.

Anderson, J. S., and J. R. Bolt. 2013. New information on amphibamids (Tetrapoda, Temnospondyli) from Richards Spur (Fort Sill), Oklahoma. Journal of Vertebrate Paleontology 33:553–567.

Anderson, J. S., D. Scott, and R. R. Reisz. 2020. The anatomy of the dermatocranium and mandible of *Cacops aspidephorus* Williston, 1910 (Temnospondyli: Dissorophidae), from the Lower Permian of Texas. Journal of Vertebrate Paleontology 40:e1776720.

Anderson, J. S., R. R. Reisz, D. Scott, N. B. Fröbisch, and S. S. Sumida. 2008. A stem batrachian from the Early Permian of Texas and the origin of frogs and salamanders. Nature 453:515–518.

Arbez, T., A. Dahoumane, and J.-S. Steyer. 2017. Exceptional endocranium and middle ear of *Stanocephalosaurus* (Temnospondyli: Capitosauria) from the Triassic of Algeria revealed by micro-CT scan, with new functional interpretations of the hearing system. Zoological Journal of the Linnean Society 180:910–929.

Arbez, T., J. B. Atkins, and H. C. Maddin. 2022. Cranial anatomy and systematics of *Dendrerpeton* cf. *helogenes* (Tetrapoda, Temnospondyli) from the Pennsylvanian of Joggins, revisited through micro-CT scanning. Papers in Palaeontology 8:e1421.

Ascarrunz, E., J.-C. Rage, P. Legreneur, and M. Laurin. 2016. *Triadobatrachus massinoti*, the earliest known lissamphibian (Vertebrata: Tetrapoda) re-examined by μCT scan, and the evolution of trunk length in batrachians. Contributions to Zoology 85:201–234.

Averianov, A. O., T. Martin, P. P. Skutschas, A. S. Rezvyi, and A. A. Bakirov. 2008. Amphibians from the Middle Jurassic Balabansai Svita in the Fergana Depression, Kyrgyzstan (Central Asia). Palaeontology 51:471–485.

Azevedo, K. L., C. S. Vega, and M. B. Soares. 2017. A new specimen of *Australerpeton cosgriffi* Barberena, 1998 (Stereospondyli: Rhinesuchidae) from the Middle/Upper Permian Rio do Rasto Formation, Paraná Basin, Brazil. Revista Brasileira de Paleontologia 20:333–344.

Baird, D. 1986. Some Upper Triassic reptiles, footprints and an amphibian from New Jersey. The Mosasaur 3:125–135.

Barberena, M. C. 1998. *Australerpeton cosgriffi* n.g., n. sp., a Late Permian Rhinesuchoid amphibian from Brazil. Anais da Academia Brasileira de Ciências 70:125–137.

Barberena, M. C., and R. F. Daemon. 1974. A primeira ocorrência de Amphibia (Labirintodontia) na Formação Rio do Rasto: implicações geocronológicas e estratigráficas. Anais do Congresso Brasileiro de Geologia 2:251–261.

Barberena, M. C., N. R. Correia, and J. Aumond. 1980. Contribuição à estratigrafia e bioestratigrafia do Grupo Passa Dois na Serra do Cadeado (Nordeste do Paraná, Brasil). Revista Brasileira de Geociências 10:208–275.

Barrett, P. J., R. J. Baillie, and E. H. Colbert. 1968. Triassic amphibian from Antarctica. Science 161:460–462.

Berman, D. S. 1973. A trimerorhachid amphibian from the Upper Pennsylvanian of New Mexico. Journal of Paleontology 47:932–945.

Berman, D. S., A. C. Henrici, D. K. Brezinski, and A. D. Kollar. 2010. A new trematopid amphibian (Temnospondyli: Dissorophoidea) from the Upper Pennsylvanian of western Pennsylvania: earliest record of terrestrial vertebrates responding to a warmer, drier climate. Annals of Carnegie Museum 78:289–318.

Berman, D. S., A. C. Henrici, T. Martens, S. S. Sumida, and J. S. Anderson. 2011. *Rotaryus gothae*, a new trematopid (Temnospondyli: Dissorophoidea) from the Lower Permian of Central Germany. Annals of Carnegie Museum 80:49–65.

Berman, D. S., and S. L. Berman. 1975. *Broiliellus hektotopos* sp.nov. (Temnospondyli: Amphibia), Washington Formation, Dunkard Group, Ohio; pp 69–78 in J. A. Barlow (ed.), The Age of the Dunkard, Proceedings of the First I. C. White Memorial Symposium. West Virginia Geological & Economic Survey, Morgantown, West Virginia.

Berman, D. S., R. R. Reisz, and D. A. Eberth. 1985. *Ecolsonia cutlerensis*, an Early Permian dissorophid amphibian from the Cutler Formation of north-central New Mexico. New Mexico Bureau of Mines & Mineral Resources Circular 191:1–31.

Berman, D. S., R. R. Reisz, and D. A. Eberth. 1987. A new genus and species of trematopid amphibian from the Late Pennsylvanian of north-central New Mexico. Journal of Vertebrate Paleontology 7:252–269.

Bolt, J. R. 1969. Lissamphibian origins: possible protolissamphibian from the Lower Permian of Oklahoma. Science 166:888–891.

Bolt, J. R. 1974a. A trematopsid skull from the Lower Permian, and analysis of some characters of the dissorophoid (Amphibia: Labyrinthodontia) otic notch. Fieldiana: Geology 30:67–79.

Bolt, J. R. 1974b. Evolution and functional interpretation of some suture patterns in Paleozoic labyrinthodont amphibians and other lower tetrapods. Journal of Paleontology 48:434–458.

Bolt, J. R. 1977a. *Cacops* (Amphibia: Labyrinthodontia) from the Fort Sill locality, Lower Permian of Oklahoma. Fieldiana: Geology 37:61–73.

Bolt, J. R. 1977b. Dissorophoid relationships and ontogeny, and the origin of the Lissamphibia. Journal of Paleontology 51:235–249.

Bolt, J. R. 1979. *Amphibamus grandiceps* as a juvenile dissorophid: evidence and implications; pp. 529–563 in M. H. Nitecki (ed.), Mazon Creek Fossils. Academic Press, New York, New York.

Bolt, J. R. 1991. Lissamphibian origins; pp. 194–222 in H.-P. Schultze and L. Trueb (eds.), Origins of the Higher Groups of Tetrapods: Controversy and Consensus. Cornell University Press, Ithaca, New York.

Bolt, J. R., and R. E. Lombard. 2001. The mandible of the primitive tetrapod *Greererpeton*, and the early evolution of the tetrapod lower jaw. Journal of Paleontology 75:1016–1042.

Bolt, J. R., and S. Chatterjee. 2000. A new temnospondyl amphibian from the Late Triassic of Texas. Journal of Paleontology 74:670–683.

Boy, J. 1987. Studien über die Branchiosauridae (Amphibia: Temnospondyli; Ober-Karbon-Unter-Perm). 2. Systematische Übersicht. Neues Jahrbuch für Geologie und Paläontologie. Abhandlungen 174:75–104.

Boy, J. A. 1972. Die Branchiosaurier (Amphibia) des saarpfälzischen Rotliegenden (Perm, SW-Deutschland. Abhandlungen des hessischen Landesamt fçr Bodenforschung 65:6–137.

Boy, J. A. 1985. Über *Micropholis*, den letzten Überlebenden der Dissorophoidea (Amphibia, Temnospondyli; Unter-Trias). Neues Jahrbuch für Geologie und Paläontologie-Monatshefte 1985:29–45.

Boy, J. A. 1988. Über einige Vertreter der Eryopoidea (Amphibia: Temnospondyli) aus dem europäischen Rotliegend (? höchstes Karbon—Perm) 1. *Sclerocephalus*. Paläontologische Zeitschrift 62:107–132.

Boy, J. A. 1989. Über einige Vertreter der Eryopoidea (Amphibia: Temnospondyli) aus dem europäischen Rotliegend (? höchstes Oberkarbon—Perm) 2. *Acanthostomatops*. Paläontologische Zeitschrift 63:133–151.

Boy, J. A. 1990. Über einige Vertreter der Eryopoidea (Amphibia: Temnospondyli) aus dem europäischen Rotliegend (? höchstes Karbon-Perm) 3. *Onchiodon*. Paläontologische Zeitschrift 64:287–312.

Boy, J. A. 1993. Über einige vertreter der Eryopoidea (Amphibia: Temnospondyli) aus dem europäischen Rotliegend (? höchstes Karbon-Perm) 4. *Cheliderpeton latirostre*. Paläontologische Zeitschrift 67:123–143.

Boy, J. A. 1995. Über die Micromelerpetontidae (Amphibia: Temnospondyli). 1. Morphologie und Paläoökologie des *Micromelerpeton credneri* (Unter-Perm; SW-Deutschland). Paläontologische Zeitschrift 69:429–457.

Boy, J. A., and H.-D. Sues. 2000 Branchiosaurs: larvae, metamorphosis, and heterochrony in temnospondyls and seymouriamorphs; pp. 1150–1197 in H. Heatwole and R. Carroll (eds.), Amphibian Biology, Volume 4: Palaeontology. Surrey Beatty, Chipping Norton.

Branson, E. B. 1905. Structure and relationships of American Labyrinthodontidae. Journal of Geology 13:568–610.

Branson, E. B., and M. G. Mehl. 1929. Triassic amphibians from the Rocky Mountain region. University of Missouri Studies 4:154–253.

Brazeau, M. D. 2011. Problematic character coding methods in morphology and their effects. Biological Journal of the Linnean Society 104:489–498.

Broili, F., and H. C. Schröder. 1937. Beobachtungen an wirbeltieren der Karrooformation. XXV. Über *Micropholis* Huxley. SSVI. Über *Lydekkerina* Broom. Sitzungsberichte der Bayerischen Akademie der Wissenschaften, Mathematisch-Naturwissenschaftliche Abteilung 1937:19–57.

Broom, R. 1930. Notes on some labyrinthodonts in the Transvaal Museum. Annals of the Transvaal Museum 14:1–10.

Brown, B. 1933. A new genus of Stegocephalia from the Triassic of Arizona. American Museum Novitates 640:1–4.

Brusatte, S. L., R. J. Butler, O. Mateus, and J.-S. Steyer. 2015. A new species of *Metoposaurus* from the Late Triassic of Portugal and comments on the systematics and biogeography of metoposaurid temnospondyls. Journal of Vertebrate Paleontology 35:e912988.

Buffa, V., N.-E. Jalil, and J.-S. Steyer. 2019. Redescription of *Arganasaurus* (*Metoposaurus*) *azerouali* (Dutuit) comb. nov. from the Upper Triassic of the Argana Basin (Morocco), and the first phylogenetic analysis of the Metoposauridae (Amphibia, Temnospondyli). Papers in Palaeontology 5:699–717.

Buffetaut, E., H. Tong, and V. Suteethorn. 1994a. First post-Triassic labyrinthodont amphibian in South East Asia: a temnospondyl intercentrum from the Jurassic of Thailand. Neues Jahrbuch für Geologie und Paläontologie-Monatshefte 1994:385–390.

Buffetaut, E., L. Raksaskulwong, V. Suteethorn, and H. Tong. 1994b. First post-Triassic temnospondyl amphibians from the Shan-Thai block: intercentra from the Jurassic of peninsular Thailand. Geological Magazine 131:837–839.

Bystrow, A. P. 1938. *Dvinosaurus* als neotenische Form der Stegocephalen. Acta Zoologica, 19:209–295.
Bystrow, A. P., and J. A. Efremov. 1940. [*Benthosuchus sushkini* Efr., a labyrinthodont from the Eotriasic of Sharzhenga River.] Trudy Paleozoologicheskogo Instituta Akademia Nauk SSSR 10:1–152. [Russian]
Carroll, R. L. 1964. Early evolution of the dissorophid amphibians. Bulletin of the Museum of Comparative Zoology 131:161–250.
Carroll, R. L. 1967. Labyrinthodonts from the Joggins formation. Journal of Paleontology 41:111–142.
Carroll, R. L. 2007. The Palaeozoic ancestry of salamanders, frogs and caecilians. Zoological Journal of the Linnean Society 150:1–140.
Carroll, R. L., and R. Holmes. 1980. The skull and jaw musculature as guides to the ancestry of salamanders. Zoological Journal of the Linnean Society 68:1–40.
Case, E. C. 1922. New reptiles and stegocephalians from the Upper Triassic of western Texas. Carnegie Institute of Washington 321:7–84.
Case, E. C. 1931. Description of a new species of *Buettneria*, with a discussion of the brain case. Contributions from the Museum of Paleontology, University of Michigan 3:187–206.
Case, E. C. 1932. A collection of stegocephalians from Scurry County, Texas. Contributions from the Museum of Paleontology, University of Michigan 4:1–56.
Case, E. C. 1935. Description of a collection of associated skeletons of *Trimerorhachis*. Contributions from the Museum of Paleontology, University of Michigan 4:227–244.
Chahud, A., and S. Petri. 2010. Anfíbio e Palaeonisciformes da Porção Basal do Membro Taquaral, Formação Irati (Permiano), Estado de São Paulo, Brasil. Geologia USP. Série Científica 10:29–37.
Chakravorti, S., and D. P. Sengupta. 2019. Taxonomy, morphometry and morphospace of cranial bones of *Panthasaurus* gen. nov. *maleriensis* from the Late Triassic of India. Journal of Iberian Geology 45:317–340.
Chase, J. N. 1965. *Neldasaurus wrightae*, a new rhachitomous labyrinthodont from the Texas Lower Permian. Bulletin of the Museum of Comparative Zoology 133:155–230.
Chatterjee, S., and T. Roy-Chowdhury. 1974. Triassic Gondwana vertebrates from India. Indian Journal of Earth Science 1:96–112.
Cheng, Z. 1980. VII. [Vertebrate fossils]; pp. 115–170 in [The Mesozoic Stratigraphy and Paleontology of Shaanxi-Gansu-Ningxia Basin, Volume 2]. Publishing House of Geology: Beijing. [Chinese]
Cheng, Z. 1981. Permo-Triassic continental deposits and vertebrate faunas of China; pp. 65–70 in M. M. Creswell, and P. Vella (eds.), Gondwana Five. Balkema, Rotterdam.
Chernin, S. 1974. Capitosaurid amphibians from the Upper Luangwa Valley, Zambia. Palaeontologia Africana 17:29–55.
Chernin, S. 1977. A new brachyopid, *Batrachosuchus concordi* sp. nov. from the upper Luangwa Valley, Zambia with a redescription of *Batrachosuchus browni* Broom, 1903. Palaeontologia Africana 20:87–109.
Chernin, S., and J. W. Cosgriff. 1975. Further consideration of the capitosaurids from the Upper Luangwa Valley, Zambia. Palaeontologia Africana 18:143–148.
Chowdhury, T. R. 1965. A new metoposaurid amphibian from the Upper Triassic Maleri Formation of Central India. Philosophical Transactions of the Royal Society of London. Series B, Biological Sciences 250:1–52.

Cisneros, J. C., C. Marsicano, K. D. Angielczyk, R. M. Smith, M. Richter, J. Fröbisch, C. F. Kammerer, and R. W. Sadleir. 2015. New Permian fauna from tropical Gondwana. Nature Communications 6:1–8.

Clack, J. A., and A. R. Milner. 2009 (for 2010). Morphology and systematics of the Pennsylvanian amphibian *Platyrhinops lyelli* (Amphibia: Temnospondyli). Earth and Environmental Science Transactions of the Royal Society of Edinburgh 100:275–295.

Clack, J. A., and A. R. Milner. 2015. Basal Tetrapoda; pp. 1–92 in H.-D. Sues (ed.), Encyclopedia of Paleoherpetology, Part 3A1. Verlag Dr. Friedrich Pfeil, Munich.

Clack, J. A., and J. S. Anderson. Early tetrapods: experimenting with form and function; pp. 71–106 in J. A. Clack, R. R. Fay, and A. N. Popper (eds.), Evolution of the Vertebrate Ear. Springer, Cham, Switzerland.

Colbert, E. H., and J. Imbrie. 1956. Triassic metoposaurid amphibians. Bulletin of the American Museum of Natural History 110:399–452.

Colbert, E. H., and J. W. Cosgriff. 1974. Labyrinthodont amphibians from Antarctica. American Museum Novitates 2552:1–30.

Coldiron, R. W. 1978. *Acroplous vorax* Hotton (Amphibia, Saurerpetontidae) restudied in light of new material. American Museum Novitates 2662:1–27.

Cope, E. D. 1889. The Batrachia of North America. United States National Museum Bulletin 34:1–525.

Cosgriff, J. W. 1965. A new genus of Temnospondyli from the Triassic of Western Australia. Journal of the Royal Society of Western Australia 48:65–90.

Cosgriff, J. W. 1969. *Blinasaurus*, a brachyopid genus from Western Australia and New South Wales. Journal of the Royal Society of Western Australia 52:65–88.

Cosgriff, J. W. 1972. *Parotosaurus wadei*, a new capitosaurid from the Gosford Formation of New South Wales. Journal of Paleontology 46:545–555.

Cosgriff, J. W. 1974. Lower Triassic Temnospondyli of Tasmania. Special Papers of the Geological Society of America 149:1–134.

Cosgriff, J. W., and J. M. Zawiskie. 1979. A new species of the Rhytidosteidae from the *Lystrosaurus* Zone and a review of the Rhytidosteoidea. Palaeontologia Africana 22:1–27.

Cosgriff, J. W., and N. K. Garbutt. 1972. *Erythrobatrachus noonkanbahensis*, a trematosaurid species from the Blina Shale. Journal of the Royal Society of Western Australia 55:5–18.

Cosgriff, J. W., and S. L. Defauw. 1987. A capitosaurid labyrinthodont from the Early Scythian of Tasmania. Alcheringa 11:21–41.

Cosgriff, J. W., and W. R. Hammer. 1984. New material of labyrinthodont amphibians from the Lower Triassic Fremouw Formation of Antarctica. Journal of Vertebrate Paleontology 4:47–56.

Cox, C. B., and P. Hutchinson. 1991. Fishes and amphibians from the Late Permian Pedra de Fogo Formation of northern Brazil. Palaeontology 34:561–573.

Credner, H. 1881. Die Stegocephalen aus dem Rotliegenden des Plauen'schen Grundes bei Dresden. Zweiter Theil. Zeitschrift der Deutschen Geologischen Gesellschaft 33:574–603.

Da-Rosa, A. A. S., G. Piñeiro, S. Dias-Da-Silva, J. C. Cisneros, F. F. Feltrin, and L. W. Neto. 2009. Bica São Tomé, um novo sítio fossilífero para o Triássico Inferior do sul do Brasil. Revista Brasileira de Paleontologia 12:67–76.

Dahoumane, A., A. Nedjari, R. Aït-Ouali, P. Taquet, R. Vacant, and J.-S. Steyer. 2016. A new mastodonsauroid temnospondyl from the Triassic of Algeria: implications for the biostratigraphy and palaeoenvironments of the Zarzaïtine Series, northern Sahara. Comptes Rendus Palevol 15:918–926.

Daly, E. 1973. A Lower Permian vertebrate fauna from southern Oklahoma. Journal of Paleontology 47:562–589.

Daly, E. 1994. The Amphibamidae (Amphibia: Temnospondyli), with a description of a new genus from the Upper Pennsylvanian of Kansas. University of Kansas Miscellaneous Publications 85:1–59.

Damiani R. J., and A. M. Jeannot. 2002. A brachyopid temnospondyl from the lower Cynognathus Assemblage Zone in the northern Karoo Basin, South Africa. Palaeontologia Africana 38:57–69.

Damiani R. J., and J. W. Kitching. 2003. A new brachyopid temnospondyl from the *Cynognathus* Assemblage Zone, Upper Beaufort Group, South Africa. Journal of Vertebrate Paleontology 23:67–78.

Damiani, R. 2004. Cranial anatomy and relationships of *Microposaurus casei*, a temnospondyl from the Middle Triassic of South Africa. Journal of Vertebrate Paleontology 24:533–541.

Damiani, R. J. 1999. Giant temnospondyl amphibians from the Early to Middle Triassic Narrabeen Group of the Sydney Basin, New South Wales, Australia. Alcheringa 23:87–109.

Damiani, R. J. 2000. Early Triassic mastodonsaurids (Temnospondyli, Stereospondyli) from Western Australia, with remarks on mastodonsauroid palaeobiogeography. Alcheringa 24:299–305.

Damiani, R. J. 2001. A systematic revision and phylogenetic analysis of Triassic mastodonsauroids (Temnospondyli, Stereospondyli). Zoological Journal of the Linnean Society 133:379–499.

Damiani, R. J. 2004. Temnospondyls from the Beaufort Group (Karoo Basin) of South Africa and their biostratigraphy. Gondwana Research 7:165–173.

Damiani, R. J., and A. Warren. 1996. A new look at members of the Superfamily Brachyopoidea (Amphibia, Temnospondyli) from the Early Triassic of Queensland and a preliminary analysis of brachyopoid relationships. Alcheringa 20:277–300.

Damiani, R. J., and A. Warren. 1997. Re-interpretation of *Parotosuchus wadei* Cosgriff, a capitosaurid from the Triassic Narrabeen Group at Gosford, New South Wales, with comments on its growth stage. Alcheringa 21:281–289.

Damiani, R. J., and B. S. Rubidge. 2003. A review of the South African temnospondyl amphibian record. Palaeontologia africana 39:21–36.

Damiani, R. J., and P. J. Hancox. 2003. New mastodonsaurid temnospondyls from the *Cynognathus* assemblage zone (upper Beaufort Group; Karoo Basin) of South Africa. Journal of Vertebrate Paleontology 23:54–66.

Damiani, R. J., J. Neveling, and P. J. Hancox. 2001. First record of a mastodonsaurid (Temnospondyli, Stereospondyli) from the Early Triassic Lystrosaurus Assemblage Zone (Karoo Basin) of South Africa. Neues Jahrbuch für Geologie und Paläontologie-Abhandlungen 221:133–144.

Damiani, R., C. A. Sidor, J.-S. Steyer, R. M., Smith, H. C. Larsson, A. Maga, and O. Ide. 2006. The vertebrate fauna of the Upper Permian of Niger. V. The primitive temnospondyl *Saharastega moradiensis*. Journal of Vertebrate Paleontology 26:559–572.

Damiani, R., J. Neveling, J. Hancox, and B. Rubidge. 2000. First trematosaurid temnospondyl from the Lystrosaurus Assemblage Zone of South Africa and its biostratigraphic implications. Geological Magazine 137:659–665.

Damiani, R., R. R. Schoch, H. Hellrung, R. Werneburg, and S. Gastou. 2009. The plagiosaurid temnospondyl *Plagiosuchus pustuliferus* (Amphibia: Temnospondyli) from the Middle Triassic of Germany: anatomy and functional morphology of the skull. Zoological Journal of the Linnean Society 155:348–373.

Danto, M., F. Witzmann, and N. B. Fröbisch. 2016. Vertebral development in Paleozoic and Mesozoic tetrapods revealed by paleohistological data. PLOS ONE 11:e0152586.

Danto, M., F. Witzmann, F., S. E. Pierce, and N. B. Fröbisch. 2017. Intercentrum versus pleurocentrum growth in early tetrapods: A paleohistological approach. Journal of Morphology 278:1262–1283.

Daza, J. D., E. L. Stanley, A. Bolet, A. M., Bauer, J. S. Arias, A. Čerňanský, J. J. Bevitt, P. Wagner, and S. E. Evans. 2020. Enigmatic amphibians in mid-Cretaceous amber were chameleon-like ballistic feeders. Science 370:.687–691.

de Miguel Chaves, C., A. Pérez-García, A. Cobos, R. Royo-Torres, F. Ortega, and > Alcalá. 2015. A diverse Late Triassic tetrapod fauna from Manzanera (Teruel, Spain). Geobios 48:479–490.

DeMar, R. 1968. The Permian labyrinthodont amphibian *Dissorophus multicinctus*, and adaptations and phylogeny of the family Dissorophidae. Journal of Paleontology 42:1210–1242.

DeMar, R. E. 1966. The functional and phylogenetic significance of the armor of dissorophid amphibians. Fieldiana: Geology 16:55–88.

Dias Dias-da-Silva, S., C. Marsicano, and C. L. Schultz. 2005. Early Triassic temnospondyl skull fragments from southern South America (Paraná Basin, Brazil). Revista Brasileira de Paleontologia 8:165–172.

Dias-da-Silva, S. 1998. Na Formação Sanga Do Cabral (Eotriássico Da Bacia Do Paraná). Acta Geologica Leopoldensia 21:101–108.

Dias-Da-Silva, S., and A. L. R. Ilha. 2009. On the presence of a pustulated temnospondyl in the Lower Triassic of southern Brazil. Acta Palaeontologica Polonica 54:609–614.

Dias-Da-Silva, S., and A. R. Milner. 2010. The pustulated temnospondyl revisited—a plagiosternine plagiosaurid from the Lower Triassic of Brazil. Acta Palaeontologica Polonica 55:561–563.

Dias-da-Silva, S., and C. Marsicano. 2011. Phylogenetic reappraisal of Rhytidosteidae (Stereospondyli: Trematosauria), temnospondyl amphibians from the Permian and Triassic. Journal of Systematic Palaeontology 9:305–325.

Dias-da-Silva, S., and E. V. Dias. 2013. A comprehensive survey of Triassic stereospondyls from southern Brazil with comments on their overall significance. Bulletin of the New Mexico Museum of Natural History and Science 61:1–93.

Dias-da-Silva, S., and R. Hewison. 2013. Phylogenetic analysis and palaeobiogeography of the Pangaean Lower Triassic Lydekkerinidae (Temnospondyli, Stereospondyli). 73rd Meeting of the Society of Vertebrate Paleontology Program and Abstract Book, 116.

Dias-da-Silva, S., C. Marsicano, and C. L. Schultz. 2006. Rhytidosteid temnospondyls in Gondwana: a new taxon from the Lower Triassic of Brazil. Palaeontology 49:381–390.

Dias-da-Silva, S., D. P. Sengupta, S. F. Cabreira, and L. R. Da Silva. 2012. The presence of *Compsocerops* (Brachyopoidea: Chigutisauridae)(Late Triassic) in southern Brazil with comments on chigutisaurid palaeobiogeography. Palaeontology 55:163–172.

Dias-da-Silva, S., E. V. Dias, and C. L. Schultz. 2009. First record of stereospondyls (Tetrapoda, Temnospondyli) in the Upper Triassic of southern Brazil. Gondwana Research 15:131–136.

Dias-da-Silva, S., S. F. Cabreira, and L. R. Da Silva. 2011. Occurrence of giant stereospondyl remains in the Santa Maria Formation (Middle/Upper Triassic of southern Brazil). Alcheringa 35:11–19.

Dias, E. V., and C. L. Schultz., 2003. The first Paleozoic temnospondyl postcranial skeleton from South America. Revista Brasileira de Paleontologia 6:29–42.

Dias, E. V., and M. C. Barberena. 2001. A temnospondyl amphibian from the Rio do Rasto Formation, Upper Permian of southern Brazil. Anais da Academia Brasileira de Ciências 73:135–143.

Dias, E. V., S. Dias-da-Silva, and C. L. Schultz. 2020. A new short-snouted rhinesuchid from the Permian of southern Brazil. Revista Brasileira de Paleontologia 23:98–122.

Dilkes, D. 2015. Carpus and tarsus of Temnospondyli. Vertebrate Anatomy Morphology Palaeontology 1:51–87.

Dilkes, D. W. 1990. A new trematopsid amphibian (Temnospondyli: Dissorophoidea) from the Lower Permian of Texas. Journal of Vertebrate Paleontology 10:222–243.

Dilkes, D. W. 1993. Biology and evolution of the nasal region in trematopid amphibians. Palaeontology 36:839–853.

Dilkes, D. W. 2009. Comparison and biomechanical interpretations of the vertebrae and osteoderms of *Cacops aspidephorus* and *Dissorophus multicinctus* (Temnospondyli, Dissorophidae). Journal of Vertebrate Paleontology 29:1013–1021.

Dilkes, D. W. 2020. Revision of the Early Permian dissorophid '*Dissorophus*' *angustus* (Temnospondyli: Dissorophoidea). Journal of Vertebrate Paleontology 40:e1801704.

Dilkes, D. W., and R. Reisz. 1987. *Trematops milleri* Williston, 1909, identified as a junior synonym of *Acheloma cumminsi* Cope, 1882: with a revision of the genus. American Museum Novitates 2902:1–12.

Dilkes, D., and L. E. Brown. 2007. Biomechanics of the vertebrae and associated osteoderms of the Early Permian amphibian *Cacops aspidephorus*. Journal of Zoology 271:396–407.

Dong, Z. 1985. [A Middle Jurassic labyrinthodont (*Sinobrachyops placenticephalus* gen. et sp. nov.) from Dashanpu, Zigong, Sichuan Province]. Vertebrata PalAsiatica 23:301–306. [Chinese]

Dutuit, J-M. 1972b. Un nouveau genre de stégocéphale du Trias supérieur marocain :*Almasaurus habbazi*. Bulletin du Muséum National D'Histoire Naturelle, Série C, Sciences de la Terre 72:73–81.

Dutuit, J.-M. 1972a. Découverte de pleurocentres dans les vertébrés de stégocephales métoposaurides. Comptes rendus hehdomadaires des Seances de l'Academie des Sciences, Serie II 274:536–537.

Dutuit, J.-M. 1976. Introduction à l'étude paléontologique du Trias continental Marocain. Descriptions des primaires ste- gocephales receillis dans le couloir d'Argana (Atlas Occiden- tal). Memoires du Museum National d'Histoire Naturelle 36:1–253.

Dutuit, J.-M., and L. Ginsburg. 1982. Présence d'un Stégocéphale géant dans le Trias du Lesotho (Afrique du Sud). Comptes Rendus de l'Académie des Sciences de Paris 294:233–236.

Dutuit, J.M. and Heyler, D., 1994. Rachitomes, lépospondyles et reptiles du Stéphanien (Carbonifère supérieur) du Bassin de Montceau-les-Mines (Massif Central-France) pp. 249–266 in C. Poplin and D. Heyler (eds.), Quand le Massif Central était sous l'Equateur. Un ecosystème carbonifère à Montceau-les-Mines. Comité des Travaux Historiques et Scientifiques, Paris.

Eaton, T. H. 1973. A Pennsylvanian dissorophid amphibian from Kansas. Occasional Papers, Museum of Natural History, University of Kansas 14:1–8.

Efremov, I. A. 1929. *Benthosaurus sushkini*, ein neuer Labyrinthodont der permotriassischen Ablagerungen der Sharschenga Flusses Bulletin de l'Académie des Sciences de l'URSS, VII. Série. Classe des Sciences naturelles 8:757–770.

Efremov, I. A. 1937. [On the Permo-Triassic labyrinthodonts from USSR. IV. A note on the lost forms *Zygosaurus* Eichwald and *Chalcosaurus* von Meyer.] Trudy Paleontologichesko Instituta Akademiya Nauk SSSR 8:17–27. [Russian]

Efremov, I. A. 1940. [Preliminary description of new forms of Permian and Triassic fauna of tetrapods from the USSR.] Trudy Paleontologichesko Instituta Akademiya Nauk SSSR 10:1–156. [Russian]

Efremov, I. A., and B. P. Vjushkov. 1955. [Catalogue of localities of Permian and Triassic terrestrial vertebrates in the territories of the USSR.] Trudy Paleontologicheskogo Instituta Akademiya Nauk SSSR 46:1–185. [Russian]

Efremov, J. A. 1932. On the Permo-Triassic labyrinthodonts from U.S.S.R. I. The labyrinthodonts of the Campylian beds of mountain Great Bogdo. II. On the morphology of *Dvinosaurus*. Travaux de l'Institut paléozoologique de l'Académie des sciences de l'U.R.S.S. 1:57–67.

Efremov, J. A. 1933. Über die Labyrinthodonten der U.d.S.S.R. II. Permische Labyrinthodonen des frühreren Gouvernements Wjatka. 1. Neubeschreibung des Labyrinthodonten *Platyops stuckenbergi*, Trautschold aus den oberpermischen Ablagerungen des Flusses Kitjak, eines Nebenflsuses der Wjatka. Trudy Palaeontologii Akademii Nauk SSSR 2:117–165.

Eichwald, E. 1848. Ueber die Saurier des kupferführenden Zechsteins Russlands. Bulletin de la Société des Naturalistes de Moscou 21:136–204.

Eltink, E. and M. C. Langer. 2014. A new specimen of the temnospondyl *Australerpeton cosgriffi* from the Late Permian of Brazil (Rio do Rasto Formation, Paraná Basin): comparative anatomy and phylogenetic relationships. Journal of Vertebrate Paleontology 34:524–538.∂

Eltink, E., Á. A. S. Da-Rosa, and S. Dias-da-Silva. 2017. A capitosauroid from the Lower Triassic of South America (Sanga do Cabral Supersequence: Paraná Basin), its phylogenetic relationships and biostratigraphic implications. Historical Biology 29:863–874.

Eltink, E., E. V. Dias, S. Dias-da-Silva, C. L. Schultz, and M. C. Langer. 2016. The cranial morphology of the temnospondyl *Australerpeton cosgriffi* (Tetrapoda: Stereospondyli) from the Middle-Late Permian of Paraná Basin and the phylogenetic relationships of Rhinesuchidae. Zoological Journal of the Linnean Society 176:835–860.

Eltink, E., R. R. Schoch, and M. C. Langer. 2019. Interrelationships, palaeobiogeography and early evolution of Stereospondylomorpha (Tetrapoda: Temnospondyli). Journal of Iberian Geology 45:251–267.

Englehorn, J., B. J. Small, and A. Huttenlocker. 2008. A redescription of *Acroplous vorax* (Temnospondyli: Dvinosauria) based on new specimens from the Early Permian of Nebraska and Kansas, USA. Journal of Vertebrate Paleontology 28:291–305.

Fernandez, V., F. Abdala, K. J. Carlson, D. C., Cook, B. S. Rubidge, A. Yates, and P. Tafforeau. 2013. Synchrotron reveals Early Triassic odd couple: injured amphibian and aestivating therapsid share burrow. PLOS ONE 8:e64978.

Foreman, B. C. 1990. A revision of the cranial morphology of the Lower Permian temnospondyl amphibian *Acroplous vorax* Hotton. Journal of Vertebrate Paleontology 10:390–397.

Fortuny, J., À. Galobart, and C. De Santisteban. 2011. A new capitosaur from the Middle Triassic of Spain and the relationships within the Capitosauria. Acta Palaeontologica Polonica 56:553–566.

Fortuny, J., J.-S. Steyer, and I. Hoşgör. 2015. First occurrence of temnospondyls from the Permian and Triassic of Turkey: Paleoenvironmental and paleobiogeographic implications. Comptes Rendus Palevol 14:281–289.

Fortuny, J., S. Gastou, F. Escuillié, L. Ranivoharimanana, and J.-S. Steyer. 2018. A new extreme longirostrine temnospondyl from the Triassic of Madagascar: phylogenetic and palaeobiogeographical implications for trematosaurids. Journal of Systematic Palaeontology 16:675–688.

Fortuny, J., T. Arbez, E. Mujal, and J.-S. Steyer. 2019. Reappraisal of '*Metoposaurus hoffmani*' Dutuit, 1978, and description of new temnospondyl specimens from the Middle–Late Triassic of Madagascar (Morondava Basin). Journal of Vertebrate Paleontology 39:e1576701.

Fraas, E. 1889. Die Labyrinthodonten der Schwäbischen Trias. Palaeontographica 36:1–158.

Fröbisch, N. B., and R. R. Reisz. 2008. A new Lower Permian amphibamid (Dissorophoidea, Temnospondyli) from the fissure fill deposits near Richards Spur, Oklahoma. Journal of Vertebrate Paleontology 28:1015–1030.

Fröbisch, N. B., and R. R. Reisz. 2012. A new species of dissorophid (*Cacops woehri*) from the Lower Permian Dolese quarry, near Richards Spur, Oklahoma. Journal of Vertebrate Paleontology 32:35–44.

Fröbisch, N. B., C. Bickelmann, and F. Witzmann. 2014. Early evolution of limb regeneration in tetrapods: evidence from a 300-million-year-old amphibian. Proceedings of the Royal Society B: Biological Sciences 281:20141550.

Fröbisch, N.B., and R. R. Reisz. 2012. A new species of dissorophid (*Cacops woehri*) from the Lower Permian Dolese quarry, near Richards Spur, Oklahoma. Journal of Vertebrate Paleontology 32:35–44.

Fröbisch, N.B., and R. R. Schoch. 2009b. The largest specimen of *Apateon* and the life history pathway of neoteny in the Paleozoic temnospondyl family Branchiosauridae. Fossil Record 12:83–90.

Gaete, R., A. Galobart, and X. Ros. 1993a. Primera noticia del hallazgo de un *Parotosaurus* (Capitosauridae, Amphibia) en el Triásico inferior del Macizo del Montseny. Treballs del Museu de Geologia de Barcelona 3:61–66.

Gaete, R., A. Galobart, and X. Ros. 1993b. El laberintodont de la Calma: Un amfibi de 230 milions d'anys trobat al Montseny. Lauro 5:27–40.

Gaete, R., A. Galobart, J. Palomar, and M. Marzo. 1994. Hallazgo de *Parotosuchus* sp.(Amphibia, Temnospondyli) en las facies Buntsandstein del Pla de La Calma (Cordilleras Costeras Catalanas): resultados preliminares. Geogaceta 16:61–63.

Gee, B. M. 2020a. Ecology, ontogeny, and taxonomy of the diverse early Permian dissorophoid assemblage from Richards Spur, Oklahoma. Ph.D. dissertation, University of Toronto, Toronto, Ontario; 444 pp.

Gee, B. M. 2020b. Size matters: the effects of ontogenetic disparity on the phylogeny of Trematopidae (Amphibia: Temnospondyli). Zoological Journal of the Linnean Society 190:79–113.

Gee, B. M. 2021. Returning to the roots: resolution, reproducibility, and robusticity in the phylogenetic inference of Dissorophidae (Amphibia: Temnospondyli). PeerJ 9:e12423.

Gee, B. M., and C. A. Sidor. 2021. Cold capitosaurs and polar plagiosaurs: new temnospondyl records from the upper Fremouw Formation (Middle Triassic) of Antarctica. Journal of Vertebrate Paleontology 41:e1998086.

Gee, B. M., and C. A. Sidor. 2021. First record of the amphibamiform *Micropholis stowi* from the lower Fremouw Formation (Lower Triassic) of Antarctica. Journal of Vertebrate Paleontology 41:e1904251.

Gee, B. M., and R. R. Reisz. 2018a. Cranial and postcranial anatomy of *Cacops morrisi*, a eucacopine dissorophid from the early Permian of Oklahoma. Journal of Vertebrate Paleontology 38:e1433186.

Gee, B. M., and R. R. Reisz. 2018b. Postcrania of large dissorophid temnospondyls from Richards Spur, Oklahoma. Fossil Record 21:79–91.

Gee, B. M., and R. R. Reisz. 2020. A redescription of the late Carboniferous trematopid *Actiobates peabodyi* from Garnett, Kansas. The Anatomical Record 303:2821–2838.

Gee, B. M., and S. E. Jasinski. 2021. Description of the metoposaurid *Anaschisma browni* from the New Oxford Formation of Pennsylvania. *Journal* of Paleontology 95:1061–1078.

Gee, B. M., D S. Berman, A. C. Henrici, J. D. Pardo, and A. K. Huttenlocker. 2020. New information on the dissorophid Conjunctio (Temnospondyli) based on a specimen from the Cutler Formation of Colorado, USA. Journal of Vertebrate Paleontology 40:e1877152.

Gee, B. M., D. S. Berman, A. C. Henrici, J. D. Pardo, and A. K. Huttenlocker. 2020. New information on the dissorophid *Conjunctio* (Temnospondyli) based on a specimen from the Cutler Formation of Colorado, USA. Journal of Vertebrate Paleontology 40:e1877152.

Gee, B. M., J. J. Bevitt, and R. R. Reisz. 2019a. A juvenile specimen of the trematopid Acheloma from Richards Spur, Oklahoma and challenges of trematopid ontogeny. Frontiers in Earth Science 7:38.

Gee, B. M., J. J. Bevitt, and R. R. Reisz. 2019b. Dissorophid diversity at the early Permian cave system near Richards Spur, Oklahoma, USA. Palaeontologia Electronica 22:1–32.

Gee, B. M., P. J. Makovicky, and C. A. Sidor. 2022. Upside down: '*Cryobatrachus*' and the lydekkerinid record from Antarctica. Journal of Paleontology 96:658–683.

Gee, B. M., W. G. Parker, and A. D. Marsh. 2018 (for 2020). Redescription of *Anaschisma* (Temnospondyli: Metoposauridae) from the Late Triassic of Wyoming and the phylogeny of the Metoposauridae. Journal of Systematic Palaeontology 18:233–258.

Getmanov, S. N. 1982. ‘A labyrinthodont from the Lower Triassic of the Obshchiy Syrt region.] Paleontologicheskii Zhurnal 1982:103–108. [Russian]; Paleontological Journal 16:102–106. [English]

Getmanov, S. N. 1986. [On the individual variation in the skull of benthosuchids as related to the development of the trematosaur organizational plan.] Paleontologicheskii Zhurnal 1986:81–91. [Russian]; Paleontological Journal 20:76–85. [English]

Getmanov, S. N. 1989. [Triassic amphibians of the East European platform (family Benthosuchidae Efremov).] Trudy Paleontologicheskogo Instituta Akademiya Nauk SSSR 236:1–102. [Russian]

Glienke, S. 2013. A taxonomic revision of *Batropetes* (Amphibia, Microsauria) from the Rotliegend (basal Permian) of Germany. Neues Jahrbuch für Geologie und Paläontologie-Abhandlungen 269:73–96.

Glienke, S. 2015. Two new species of the genus *Batropetes* (Tetrapoda, Lepospondyli) from the Central European Rotliegend (basal Permian) in Germany. Journal of Vertebrate Paleontology 35:e918041.

Godfrey, S. J. 1989a. Ontogenetic changes in the skull of the Carboniferous tetrapod *Greererpeton burkemorani* Romer, 1969. Philosophical Transactions of the Royal Society of London. B, Biological Sciences 323:135–153.

Godfrey, S. J. 1989b. The postcranial skeletal anatomy of the Carboniferous tetrapod *Greererpeton burkemorani* Romer, 1969. Philosophical Transactions of the Royal Society of London. B, Biological Sciences 323:75–133.

Godfrey, S. J. 2003. A diminutive temnospondyl amphibian from the Pennsylvanian of Illinois. Canadian Journal of Earth Sciences 40:507–514.

Godfrey, S. J., A. R. Fiorillo, and R. L. Carroll. 1987. A newly discovered skull of the temnospondyl amphibian *Dendrerpeton acadianum* Owen. Canadian Journal of Earth Sciences 24:796–805.

Golubev, V. K. 1995. [New species of *Melosaurus* (Amphibia, Labyrinthodontia) from Kazanian deposits of the Kama River drainage area.] Paleontologicheskii Zhurnal 1995:86–97. [Russian]; Paleontological Journal 29:107–119. [English]

Golubev, V. K., and V. V. Bulanov. 2018. Amphibians of the Permian Sundyr tetrapod assemblage of Eastern Europe. Paleontological Journal 52:639–652.

Goodman, C. M., G. F. Jongsma, J. E. Hill, E. L. Stanley, Q. M. Tuckett, D. C. Blackburn, and C. M. Romagosa. 2021. A case of mistaken identity: Genetic and anatomical evidence reveals the cryptic invasion of *Xenopus tropicalis* in central Florida. Journal of Herpetology 55:62–69.

Gower, D. J., and A. G. Sennikov. 1997. *Sarmatosuchus* and the early history of the Archosauria. Journal of Vertebrate Paleontology 17:60–73.

Gregory, J. T. 1950. Tetrapods of the Pennsylvanian nodules from Mazon Creek, Illinois. American Journal of Science 248:833–873.

Gruntmejer, K., A. Bodzioch, and D. Konietzko-Meier. 2021. Mandible histology in *Metoposaurus krasiejowensis* (Temnospondyli, Stereospondyli) from the Upper Triassic of Poland. PeerJ 9:e12218.

Gubin, Y. M. 1980. [New Permian dissorophids of the Ural forelands.] Paleontologicheskii Zhurnal 14:88–96. [Russian]; Paleontological Journal 15:108–110. [English]

Gubin, Y. M. 1983. [The first eryopids from the Permian of the Eastern European platform.] Paleontologicheskii Zhurnal 1983:110–115. [Russian]

Gubin, Y. M. 1984. [The systematic position of the intasuchids.] Paleontologicheskii Zhurnal 1984:118–120. [Russian]; Paleontological Journal 18:115–118. [English]

Gubin, Y. M. 1986. [New data on the archegosauroids of the East European platform.] Paleontologicheskii Zhurnal 1986:75–80. [Russian]; Paleontological Journal 20:70–75. [English]

Gubin, Y. M. 1987. [On the systematic position and age of some labyrinthodonts from the Upper Permian deposits of the region west of the Urals.] Paleontologicheskii Zhurnal 1987:94–99. [Russian]; Paleontological Journal 21:104–110. [English]

Gubin, Y. M. 1989. [The systematic position of the labyrinthodonts from the Malaya Kinel' locality (Orenburg Region).] Paleontologicheskii Zhurnal 1989:116–120. [Russian]; Paleontological Journal 23:115–119. [English]

Gubin, Y. M. 1991. [Permian archegosauroid amphibians of the USSR]. Trudy Paleontologicheskogo Instituta Akademiya Nauk SSSR 249:1–138. [Russian]

Gubin, Y. M. 1993. [New data on lower tetrapods from the Upper Permian of the Northern Urals and Obshchiy Syrt regions.] Paleontologicheskii Zhurnal 1993:97–105. [Russian]; Paleontological Journal 27:120–131. [English]

Gubin, Y. M. 2004. [A new dvinosaur (Amphibia, Temnospondyli) from the Upper Tatarian of the Middle Volga Region.] Paleontologicheskii Zhurnal 2004:72–80. [Russian]

Gubin. Y. M. 1981. [A new platyoposaurid from Bashkiria.] Paleontologicheskii Zhurnal 1981:141–143. [Russian]

Hancox, P. J., R. J. Damiani, and B. S. Rubidge. 2000. First occurrence of *Paracyclotosaurus* (Temnospondyli, Capitosauridae) in the Karoo Basin of South Africa and its biostratigraphic significance. South African Journal of Science 96:135–137.

Hara, S., and K. Nishikawa. 2021 (for 2022). Osteological characteristics of the Setouchi salamander *Hynobius setouchi* (Urodela, Hynobiidae). The Anatomical Record 305:1316–1342.

Hart, L. J., M. R. McCurry, M. Frese, T. J. Peachey, and J. Brocks. 2021. The first tetrapod remains from the Upper Jurassic Talbragar Fossil Fish Bed. Alcheringa 45:423–428.

Hartmann-Weinberg, A. 1939. *Melosaurus uralensis* H. v. Meyer an Upper Permian Archegosaurid. Problems of Palaeontology 5:20–31.

Hartmann-Weinberg, A. P., and F. M. Kuz'min. 1936. Lower Triassic stegosephalians of the USSR. *Lyrocephalus acutirostris* nov. sp. Problems of Palaeontology 1:75–84.

Haughton, S. H. 1925. Investigations in the South African Fossil Reptilia and Amphibia (Part 13). Annals of the South African Museum 22:227–261.

Haughton, S. H. 1932. On a collection of Karroo vertebrates from Tanganyika Territory. Quarterly Journal of the Geological Society of London 88:634-671.

Heckert, A. B., J. S. Mitchell, V. P. Schneider, and P. E. Olsen. 2012. Diverse new microvertebrate assemblage from the Upper Triassic Cumnock Formation, Sanford Subbasin, North Carolina, USA. Journal of Paleontology 86:368–390.

Hellrung, H., 2003. *Gerrothorax pustuloglomeratus*, ein Temnospondyle (Amphibia) mit knöcherner Branchialkammer aus dem Unteren Keuper von Kupferzell (Süddeutschland). Stuttgarter Beiträge zur Naturkunde Serie B 330:1–130.

Henderson, A. C. 2002. Studies on Interactions Among Xenopus Taxa Using Comparative Osteology and Other Methods: An Evolutionary Perspective. Ph.D. dissertation, University of Bristol, Bristol, 323 pp.

Hewison, R. H. 1996. The skull of *Deltacephalus whitei*, a lydekkerinid temnospondyl amphibian from the Lower Triassic of Madagascar. Palaeontology 39:305–322.

Hewison, R. H. 2007. The skull and mandible of the stereospondyl *Lydekkerina huxleyi*, (Tetrapoda: Temnospondyli) from the Lower Triassic of South Africa, and a reappraisal of the family Lydekkerinidae, its origin, taxonomic relationships and phylogenetic importance. Journal of Temnospondyl Palaeontology 1:1–80.

Heyler, D. 1994. Les branchiosaures stéphaniens et permiens de Montceau-les-Mines et des autres bassins du Massif Central; pp. 227–247 in C. Poplin and D. Heyler (eds.), Quand le Massif Central était sous l'Equateur. Un ecosystème carbonifère à Montceau-les-Mines. Comité des Travaux Historiques et Scientifiques, Paris.

Holmes, R. 1984. The Carboniferous amphibian *Proterogyrinus scheelei* Romer, and the early evolution of tetrapods. Philosophical Transactions of the Royal Society of London. B, Biological Sciences 306:431–524.

Holmes, R. B., and R. L. Carroll. 1977. A temnospondyl amphibian from the Mississippian of Scotland. Bulletin of the Museum of Comparative Zoology 147:489–511.

Holmes, R. B., R. L. Carroll, and R. R. Reisz. 1998. The first articulated skeleton of *Dendrerpeton acadianum* (Temnospondyli, Dendrerpetontidae) from the Lower Pennsylvanian locality of Joggins, Nova Scotia, and a review of its relationships. Journal of Vertebrate Paleontology 18:64–79.

Holmes, R., D. S Berman, and J. S. Anderson. 2013. A new dissorophid (Temnospondyli, Dissorophoidea) from the Early Permian of New Mexico (United States). Comptes Rendus Palevol 12:419–435.

Holmes, R., S. Godfrey, and D. Baird. 1995. Tetrapod remains from the late Mississippian Pomquet Formation near Grand Étang, Nova Scotia. Canadian Journal of Earth Sciences 32:913–921.

Hook, R. W. 1993. *Chenoprosopus lewisi*, a new cochleosaurid amphibian (Amphibia: Temnospondyli) from the Permo-Carboniferous of north-central Texas. Annals of Carnegie Museum 62:273–291.

Hook, R. W. and D. Baird. 1984. *Ichthycanthus platypus* Cope, 1877, reidentified as the dissorophoid amphibian *Amphibamus lyelli*. Journal of Paleontology 58:697–702.

Hook, R. W., and D. Baird. 1986. The diamond coal mine of Linton, Ohio, and its Pennsylvanian-age vertebrates. Journal of Vertebrate Paleontology 6:174–190.

Hook, R. W., and D. Baird. 1994, A new fish and tetrapod assemblage from the Allegheny Group (late Westphalian, upper Carboniferous) of eastern Ohio, U.S.A.: Pollichia-Buch 29:143–154.

Hotton, N. 1959. *Acroplous vorax*, a new and unusual labyrinthodont amphibian from the Kansas Permian. Journal of Paleontology 33:161–178.

Houttuyn, M. 1782. Het onderscheid der salamanderen van de haagdissen in’t algemeneen, en van de gekkoos in’t byzonder. Venhandelingen Uitgegeven door het Zeeuwsch Genootschap der Wetenschappen te Vlissingen 9:304-336.

Howie, A. A. 1970. A new capitosaurid labyrinthodont from East Africa. Palaeontology 13:210–253.

Hunt, A. P. 1993. Revision of the Metoposauridae (Amphibia: Temnospobdyli) and description of a new genus from Western North America. Museum of Northern Arizona Bulletin 59:67–97.

Hunt, A. P. 1993. Revision of the Metoposauridae (Amphibia: Temnospobdyli) and description of a new genus from Western North America. Museum of Northern Arizona Bulletin 59:67–97.

Hunt, A. P., S. G. Lucas, and D. S. Berman. 1996. A new amphibamid (Amphibia: Temnospondyli) from the Late Pennsylvanian (middle Stephanian) of central New Mexico, USA. Paläontologische Zeitschrift 70:555–565.

Huttenlocker, A. K., A. Henrici, W. J. Nelson, S. Elrick, D. S. Berman, T. Schlotterbeck, and S. S. Sumida. 2018. A multitaxic bonebed near the Carboniferous–Permian boundary (Halgaito Formation, Cutler Group) in Valley of the Gods, Utah, USA: vertebrate paleontology and taphonomy. Palaeogeography, Palaeoclimatology, Palaeoecology 499:72–92.

Huttenlocker, A. K., J. D. Pardo, and B. J. Small. 2005. An earliest Permian nonmarine vertebrate assemblage from the Eskridge Formation, Nebraska. New Mexico Museum of Natural History and Science Bulletin 30:133–143.

Huttenlocker, A. K., J. D. Pardo, B. J. Small, and J. S. Anderson. 2013. Cranial morphology of recumbirostrans (Lepospondyli) from the Permian of Kansas and Nebraska, and early morphological evolution inferred by micro-computed tomography. Journal of Vertebrate Paleontology 33:540–552.

Ingavat, R., and P. Janvier. 1981. *Cyclotosaurus* cf. *posthumus* Fraas (Capitosauridae, Stereospondyli) from the Huai Hin Lat Formation (Upper Triassic), Northeastern Thailand: with a note on capitosaurid biogeography. Geobios 14:711–725.

Ivahknenko, M. F. 1972. [A new benthosuchid from the Lower Triassic of the Upper Volga region.]. Paleontologicheskii Zhurnal 1972:93–99. [Russian]; Paleontological Journal 6:532–536. [English]

Jalil, N.-E. 1999. Continental Permian and Triassic vertebrate localities from Algeria and Morocco and their stratigraphical correlations. Journal of African Earth Sciences 29:219–226.

Janvier, P. 1983. Le toit cranien de *Peltostega* cf. *erici* WIMAN, un Rhytidosteidae (Temnospondyli) du Scythien du Spitsberg. Bulletin du Museum National d’Historie Naturelle Section C: Sciences de la Terre 5:25–32.

Janvier, P. 1992. Les écailles des Trématosaures (Tetrapoda, Temnospondyli): nouvelles données sur les Trématosaures du Trias inférieur de Madagascar. Bulletin du Muséum national d'histoire naturelle. Section C, Sciences de la terre, paléontologie, géologie, minéralogie 14:3–13.

Jeannot, A. M., R. Damiani, and B. S. Rubidge. 2006. Cranial anatomy of the Early Triassic stereospondyl *Lydekkerina huxleyi* (Tetrapoda: Temnospondyli) and the taxonomy of South African lydekkerinids. Journal of Vertebrate Paleontology 26:822–838.

Jenkins Jr., F. A., N. H. Shubin, S. M. Gatesy, and A. Warren. 2008. *Gerrothorax pulcherrimus* from the Upper Triassic Fleming Fjord Formation of East Greenland and a reassessment of head lifting in temnospondyl feeding. Journal of Vertebrate Paleontology 28:935–950.

Jenkins, F. A., D. M. Walsh, and R. L. Carroll. 2007. Anatomy of *Eocaecilia micropodia*, a limbed caecilian of the Early Jurassic. Bulletin of the Museum of Comparative Zoology 158:285–365.

Jupp, R., and A. A. Warren. 1986. The mandibles of the Triassic temnospondyl amphibians. Alcheringa 10:99–124.

Kear, B.P., Poropat, S.F. and Bazzi, M., 2016. Late Triassic capitosaurian remains from Svalbard and the palaeobiogeographical context of Scandinavian Arctic temnospondyls. *Geological Society, London, Special Publications*, *434*(1), pp.113-126.

Keyser, A. W. 1973. A new Triassic vertebrate fauna from south west Africa. Palaeontologia Africana 16:1–15.

Khaldoune, F., N.-E. Jalil, D. Germain, and J.-S. Steyer. 2016. Les vertébrés du Permien et du Trias du Maroc (bassin d'Argana, Haut-Atlas occidental): une fenêtre ouverte sur l'évolution autour de la grande crise finipaléozoïque. Mémoires de la Société Géologique de France 180:103–167.

Kitching, J. W, 1957. A new small stereospondylous labyrinthodont from the Triassic beds of South Africa. Palaeontologia africana 5:67–82.

Klembara, J., and J.-S. Steyer. 2012. A new species of *Sclerocephalus* (Temnospondyli: Stereospondylomorpha) from the early Permian of the Boskovice Basin (Czech Republic). Journal of Paleontology 86:302–310.

Kligman, B., M. Stocker, A. Marsh, S. Nesbitt, and W. Parker. 2021. New Late Triassic stem-caecilian from southwestern North America strengthens evidence for lissamphibian monophyly, and illuminates the anatomical, functional, and geographic origins of living caecilians. Society of Vertebrate Paleontology Annual Meeting Program and Abstracts, 161.

Koken, E. 1913. Beiträge zur Kenntnis der Schichten von Heiligkreuz (Abteital, Südtirol). Abhandlungen der Kaiserlich-Königlichen Geologischen Reichsanstalt 16:1–43.

Konietzko-Meier, D., and K. Wawro. 2007. Mandibular dentition in the Late Triassic temnospondyl amphibian *Metoposaurus*. Acta Palaeontologica Polonica 52:213–215.

Konietzko-Meier, D., M. Danto, and K. Gądek. 2014. The microstructural variability of the intercentra among temnospondyl amphibians. Biological Journal of the Linnean Society 112:747–764.

Konietzko-Meier, D., E. M. Teschner, A. Bodzioch, and P. M. Sander. 2020. Pentadactyl manus of the *Metoposaurus krasiejowensis* from the Late Triassic of Poland, the first record of pentadactyly among Temnospondyli. Journal of Anatomy 237:1151–1161.

Konzhukova, E. D. 1955. [Permian and Triassic labyrinthodonts of the Volga and Urals region]. Trudy Paleontologicheskogo Instituta Akademiya Nauk SSSR 49:5–88. [Russian]

Konzhukova, E. D. 1956. [The Intan Lower Permian fauna of the northern Ural region]. Trudy Paleontologicheskogo Instituta Akademiya Nauk SSSR 62:5–50. [Russian]

Konzhukova, E. D. 1965. [A new *Parotosaurus* from the Triassic of Cisuralia]. Trudy Paleontologicheskii Zhurnal 1965:97–104. [Russian]

Kufner, A. M., and B. M. Gee. 2021. Reevaluation of the holotypes of *Koskinonodon princeps* Branson and Mehl, 1929, and *Borborophagus wyomingensis* Branson and Mehl, 1929 (Temnospondyli, Metoposauridae). Journal of Vertebrate Paleontology 41:e1922067.

Lane, H. H. 1932. A new stegocephalian from the Pennsylvanian of Arkansas. Kansas University Science Bulletin 20:313–317.

Langston, W. 1953. Permian amphibians from New Mexico. University of California Publications in Geological Sciences 29:349–416.

Laurin, M., and R. Soler-Gijón. 2001. The oldest stegocephalian from the Iberian Peninsula: evidence that temnospondyls were euryhaline. Comptes Rendus de l'Académie des Sciences-Series III-Sciences de la Vie 324:495–501.

Laurin, M., and R. Soler-Gijón. 2006. The oldest known stegocephalian (Sarcopterygii: Temnospondyli) from Spain. Journal of Vertebrate Paleontology 26:284–299.

Lavina, E. L., and M. C. Barberena. 1985, Anfíbios ritidosteídeos e lidekkerinídeos da Formação Sanga do Cabral (Triássico Inferior do Rio Grande do Sul). Implicações bioestratigráficas e geocronológicas. Iheringia 10:19–27.

Lehman, J. P. 1961. Les Stégocéphales de Madagascar. Annales de Paléontologie, 47:111–154.

Lehman, J. P. 1961. Les stégocephales du Trias de Madagascar. Annales de Paléontologie 47:111–154.

Lehman, J. P. 1966. Nouveaux Stégocéphales de Madagascar. Annales de Paléontologie, 52:117–139.

Lehman, J. P. 1971. Nouveaux vertébrés fossiles du trias de la série de Zarzaitine. Annales de Paléontologie 53:71–93.

Lehman, J. P. 1974. A propos des Trématosaures de Madagascar. Comptes Rendus sommaires de la Société Géologique de France, 1974:166–167.

Lehman, J.-P. 1966. Nouveaux stégocéphales de Madagascar. Annales de Paléontologie 52:117–139.

Leidy, J. 1856. Notice of extinct vertebrated animals discovered by Prof. E. Emmons. Proceedings of the Academy of Natural Sciences, Philadelphia 8:255-256.

Leonardi, P. 1952. Un osso nasale di *Metoposaurus* nei tufi a pachicardie dell'Alpe di Siusi (Dolomiti Occidentali). Revista Italiana *di* Paleontologia 58:145-147.

Lewis, G. E., and P. P. Vaughn. 1965. Early Permian Vertebrates from the Cutler Formation of the Placerville Area Colorado. U.S.G.S. Contributions to Paleontology: Geological Survey Professional Paper 503-C:C1–C46.

Li, J., and Z. Cheng. 1999. New anthracosaur and temnospondyl amphibians from Gansu, China—the fifth report on Late Permian Dashankou Lower Tetrapod Fauna. Vertebrata Palasiatica, 37:242–247.

Liu, J. 2016. *Yuanansuchus maopingchangensis* sp. nov., the second capitosauroid temnospondyl from the Middle Triassic Badong Formation of Yuanan, Hubei, China. PeerJ 4:e1903.

Liu, J. 2018. Osteology of the large dissorophid temnospondyl *Anakamacops petrolicus* from the Guadalupian Dashankou Fauna of China. Journal of Vertebrate Paleontology 38:e1513407.

Liu, J., and Y. Wang. 2005. The first complete mastodonsauroid skull from the Triassic of China: *Yuanansuchus laticeps* gen. et sp. nov. Journal of Vertebrate Paleontology 25:725–728.

Long, R. A., and P. A. Murry. 1995. Late Triassic (Carnian and Norian) tetrapods from the southwestern United States. New Mexico Museum of Natural History and Science Bulletin 4:1–254.

Lozovsky, V.R. and Shishkin, M.A. 1974. [First discovery of a labyrinthodont in the Lower Triassic of Mangyshlak]. Doklady Akademii Nauk SSSR 214:169–172. [in Russian]

Lucas, S. G., and A. P. Hunt. 1993. A review of Triassic labyrinthodont amphibians from China. Geobios 26:121–128.

Lucas, S. G., and R. M. Sullivan. 1996, Fossils provide a Pennsylvania standard for part of Late Triassic time. Pennsylvania Geology 27:8–14.

Lucas, S. G., J. A. Spielmann, and A. P. Hunt. 2007. Biochronological significance of Late Triassic tetrapods from Krasiejów, Poland. New Mexico Musuem of Natural History and Science Bulletin 41:248–258.

Lucas, S. G., T. Kolar-Jurkovšek, and B. Jurkovšek. 2008. First record of a fossil amphibian in Slovenia (Lower Triassic, Olenekian). Rivista Italiana di Paleontologia e Stratigrafia 114:323–326.

Lucas, S.G., L. F. Rinehart, K. Krainer, J. A. Spielmann, and A. B. Heckert. 2010. Taphonomy of the Lamy amphibian quarry: a Late Triassic bonebed in New Mexico, USA. Palaeogeography, Palaeoclimatology, Palaeoecology 298:388–398.

Lund, R. 1975. Vertebrate fossil zonation and correlation of the Dunkard Basin; pp. 171–182 in J. A. Barlow (ed.), The Age of the Dunkard, Proceedings of the First I. C. White Memorial Symposium. West Virginia Geological & Economic Survey, Morgantown, West Virginia.

Lund, R. 1976. General geology and vertebrate biostratigraphy of the Dunkard Basin; pp 225–239 in H. Falke (ed.), The Continental Permian in Central, West and South Europe. D. Reidel, Dordrecht.

Maddin, H. C., and J. S. Anderson. 2012. Evolution of the amphibian ear with implications for lissamphibian phylogeny: insight gained from the caecilian inner ear. Fieldiana: Life and Earth Sciences 2012:59–76.

Maddin, H. C., F. A. Jenkins Jr., and J. S. Anderson. 2012. The braincase of *Eocaecilia micropodia* (Lissamphibia, Gymnophiona) and the origin of caecilians. PLOS ONE 7:e50743.

Maddin, H. C., N. B. Fröbisch, D. C. Evans, and A. R. Milner. 2013. Reappraisal of the Early Permian amphibamid *Tersomius texensis* and some referred material. Comptes Rendus Palevol 12:447-461.

Maddison, W. P. 1993. Missing data versus missing characters in phylogenetic analysis. Systematic Biology 42:576–581.

Maganuco, S., and G. Pasini. 2009. A new specimen of trematosaurian temnospondyl from the Lower Triassic of NW Madagascar, with remarks on palatal anatomy and taxonomic affinities. Atti Societ italiana Museo Civico di Storia Naturale Milano 150:91–112.

Maganuco, S., G. Pasini, and M. Auditore. 2014. A revision of the short-faced stereospondyls *Mahavisaurus dentatus* and *Lyrosaurus australis* from the Lower Triassic of Madagascar: cranial anatomy, ontogenetic remarks, palaeoecology and rhytidosteid phylogeny. Memoire della Società Italiana di Scienze Naturali e del Museo Civico di Storia Naturale di Milano 39:1–64.

Maganuco, S., J.-S. Steyer, G. Pasini, M. Boulay, S. Lorrain, A. Bénéteau, and M. Auditore. 2009. An exquisite specimen of *Edingerella madagascariensis* (Temnospondyli) from the Lower Triassic of Madagascar; cranial anatomy, phylogeny, and restorations. Memoire della Società Italiana di Scienze Naturali e del Museo Civico di Storia Naturale di Milano 36:1–72.

Maisch, M. W. 2020. *Aphaneramma kokeni* (von Huene, 1920), a lonchorhynchine trematosaurid (Amphibia: Temnospondyli) from the Lower Triassic of Pakistan. Neues Jahrbuch für Geologie und Paläontologie-Abhandlu*ngen* 295:211–241.

Maisch, M.W. and Matzke, A.T., 2005. Temnospondyl amphibians from the Jurassic of the southern Junggar Basin (NW China). *Paläontologische Zeitschrift*, *79*(2), pp.285-301.

Maisch, M.W., Matzke, A.T. and Sun, G., 2004. A relict trematosauroid (Amphibia: Temnospondyli) from the Middle Jurassic of the Junggar Basin (NW China). *Naturwissenschaften*, *91*(12), pp.589-593.

Malabarba, M. C., F. Abdala, F. E. Weiss, and P. A. Perez. 2003. New data on the late Permian vertebrate fauna of Posto Queimado, Rio do Rasto Formation, southern Brazil. Revista Brasileira de Paleontologia 6:49–54.

Mann, A., and B. M. Gee. 2019. Lissamphibian-like toepads in an exceptionally preserved amphibamiform from Mazon Creek. Journal of Vertebrate Paleontology 39:e1727490.

Marjanović, D., and Laurin, M. 2019. Phylogeny of Paleozoic limbed vertebrates reassessed through revision and expansion of the largest published relevant data matrix. PeerJ 6:e5565.

Marjanović, D., and M. Laurin. 2008. A reevaluation of the evidence supporting an unorthodox hypothesis on the origin of extant amphibians. Contributions to Zoology 77:149–199.

Marsicano, C. A. 1990. Redescripción del craneo del holotipo de *Pelorocephalus mendozensis* Cabrera, 1944 (Amphibia, Temnospondyli) del Triasico del Cerro Bayo, Provincia de Mendoza. Ameghiniana 27:39–44.

Marsicano, C. A. 1994. Postcranial skeleton of a brachyopoid (Amphibia, Temnospondyli) from the Triassic of Mendoza (Argentina). Alcheringa 17:185–197.

Marsicano, C. A. 1996. *Otuminisaurus limensis* Rusconi, 1948: a problematic temnospondyl amphibian from the Triassic of Argentina. Journal of Vertebrate Paleontology 16:785–786.

Marsicano, C. A. 1999. Chigutisaurid amphibians from the Upper Triassic of Argentina and their phylogenetic relationships. Palaeontology 42:545–565.

Marsicano, C. A. 2005. A new temnospondyl record from the Upper Triassic of Argentina. Ameghiniana 42:501–504.

Marsicano, C. A., and A. Warren. 1998. The first Palaezoic rhytidosteid: *Trucheosaurus major* (Woodward, 1909) from the late Permian of Australia, and a reassessment of the Rhytidosteidae (Amphibia, Temnospondyli). Bulletin of the Natural History Museum, Geology Series 54:147–154.

Marsicano, C. A., D. Perea, and M. Ubilla. 2000. A new temnospondyl amphibian from the Lower Triassic of South America. Alcheringa 24:119–123.

Marsicano, C. A., E. Latimer, B. Rubidge, and R. M. Smith. 2017. The Rhinesuchidae and early history of the Stereospondyli (Amphibia: Temnospondyli) at the end of the Palaeozoic. Zoological Journal of the Linnean Society 181:357–384.

Marsicano, C., K. D. Angielczyk, J. C. Cisneros, M. Richter, C. F. Kammerer, J. Fröbisch, and R. M. Smith. 2021. Brazilian Permian dvinosaurs (Amphibia, Temnospondyli): revised description and phylogeny. Journal of Vertebrate Paleontology 41:e1893181.

Martin, L. D., and D. Merriam 2011. A labyrinthodont amphibian from the margin of the Pennsylvanian epicontinental sea in Kansas. Transactions of the Kansas Academy of Science 114:3–4.

Martz, J. W., B. Mueller, S. J. Nesbitt, M. R. Stocker, W. G. Parker, M. Atanassov, N. Fraser, J. Weinbaum, and J. R. Lehane. 2012. A taxonomic and biostratigraphic re-evaluation of the Post Quarry vertebrate assemblage from the Cooper Canyon Formation (Dockum Group, Upper Triassic) of southern Garza County, western Texas. Earth and Environmental Science Transactions of the Royal Society of Edinburgh 103:339–364.

Martz, J., J. Kirkland, A. Milner, W. Parker, and V. Santucci. 2017. Upper Triassic lithostratigraphy, depositional systems, and vertebrate paleontology across southern Utah. Geology of the Intermountain West 4:99–180.

Maryanska, T., and M. A. Shishkin. 1996. New cyclotosaurid (Amphibia: Temnospondyli) from the Middle Triassic of Poland and some problems of interrelationships of capitosauroids. Prace Muzeum Ziemi 43:53–83.

Marzola, M., O. Mateus, N. H. Shubin, and L. B. Clemmensen. 2017. *Cyclotosaurus naraserluki*, sp. nov., a new Late Triassic cyclotosaurid (Amphibia, Temnospondyli) from the Fleming Fjord Formation of the Jameson Land Basin (East Greenland). Journal of Vertebrate Paleontology 37:e1303501.

May, W., A. K. Huttenlocker, J. D. Pardo, J. Benca, and B. J. Small. 2011. New Upper Pennsylvanian armored dissorophid records (Temnospondyli, Dissorophoidea) from the US midcontinent and the stratigraphic distributions of dissorophids. Journal of Vertebrate Paleontology 31:907–912.

Mazin, J.-M. 1986. *Negevodus ramonensis* n. g. n. sp., un nouveau placodonte du Trias moyen du Negev (Israel). Comptes Rendus de I'Academic des Sciences, Paris , Serie II 302:927–929.

Mazin, J.-M., and P. Janvier. 1983. L'anatomie de *Lyrocephaliscus euri* (WIMAN), trématosaure du Trias Inférieur du Spitsberg: arrière-crâne, squelette axial et ceinture scapulaire. Palaeovertebrata 13:13–31.

McHugh, J. B. 2012. Temnospondyl ontogeny and phylogeny, a window into terrestrial ecosystems during the Permian-Triassic mass extinction. Ph.D. dissertation, The University of Iowa, Iowa City, 198 pp.

Milner, A. 1996. A revision of the temnospondyl amphibians from the Upper Carboniferous of Joggins, Nova Scotia. Special Papers in Palaeontology 52:81–103.

Milner, A. R. 1980. The temnospondyl amphibian *Dendrerpeton* from the Upper Carboniferous of Ireland. Palaeontology 23:125-141.

Milner, A. R. 1982a. A small temnospondyl amphibian from the Lower Pennsylvanian of Nova Scotia. Journal of Paleontology 56:1302–1305.

Milner, A. R. 1982b. Small temnospondyl amphibians from the Middle Pennsylvanian of Illinois. Palaeontology 25:635–664.

Milner, A. R. 1985a. On the identity of *Trematopsis seltini* (Amphibia: Temnospondyli) from the Lower Permian of Texas. Neues Jahrbuch für Geologie und Paläontologie-Monatshefte 1985:357–367.

Milner, A. R. 1985b. On the identity of the amphibian *Hesperoherpeton garnettense* from the Upper Pennsylvanian of Kansas. Palaeontology 28:767–776.

Milner, A. R. 2003. *Longiscitula houghae* DeMar, 1966 (Amphibia: Temnospondyli), a junior synonym of *Dissorophus multicinctus* Cope, 1895. Journal of Vertebrate Paleontology 23:941–944.

Milner, A. R. 2018. Two primitive trematopid amphibians (Temnospondyli, Dissorophoidea) from the Upper Carboniferous of the Czech Republic. Earth and Environmental Science Transactions of the Royal Society of Edinburgh 109:201–223.

Milner, A. R., and R. R. Schoch. 2004. The latest metoposaurid amphibians from Europe. Neues Jahrbuch für Geologie und Paläontologie Abhandlungen 232:231–252.

Milner, A. R., and R. R. Schoch. 2013. *Trimerorhachis* (Amphibia: Temnospondyli) from the Lower Permian of Texas and New Mexico: cranial osteology, taxonomy and biostratigraphy. Neues Jahrbuch für Geologie und Paläontologie, Abhandlungen 270:91–128.

Milner, A. R., and S. E. K. Sequeira. 1993. The temnospondyl amphibians from the Viséan of east Kirkton, West Lothian, Scotland. Earth and Environmental Science Transactions of the Royal Society of Edinburgh 84:331–361.

Milner, A. R., and S. E. K. Sequeira. 1998. A cochleosaurid temnospondyl amphibian from the Middle Pennsylvanian of Linton, Ohio, USA. Zoological Journal of the Linnean Society 122:261–290.

Milner, A. R., and S. E. K. Sequeira. 2003a. On a small *Cochleosaurus* described as a large *Limnogyrinus* [Amphibia, Temnospondyli] from the Upper Carboniferous of the Czech Republic. Acta Palaeontologica Polonica 48:143–147.

Milner, A. R., and S. E. K. Sequeira. 2011. The amphibian *Erpetosaurus radiatus* (Temnospondyli, Dvinosauria) from the Middle Pennsylvanian of Linton, Ohio: morphology and systematic position. Special Papers in Palaeontology 86:57–73.

Milner, A. R., B. G. Gardiner, N. C. Fraser, and M. A. Taylor. 1990. Vertebrates from the Middle Triassic Otter Sandstone Formation of Devon. Palaeontology 33:873–892.

Milner, A. R., C. Duffin, and D. Delsate. 1996. Plagiosaurid and capitosaurid amphibian material from the Late Triassic of Medernach, Grand-Duchy of Luxembourg: preliminary note. Bulletin de la Socièté belge de Gèologie 104:42–53.

Milner, A. R., J. Klembara, and O. Dostál. 2007. A zatrachydid temnospondyl from the Lower Permian of the Boskovice Furrow in Moravia (Czech Republic). Journal of Vertebrate Paleontology 27:711–715.

Milner, A.R., and S. E. K. Sequeira. 2003b. Revision of the amphibian genus *Limnerpeton* [Temnospondyli] from the Upper Carboniferous of the Czech Republic. Acta Palaeontologica Polonica 48:123–141.

Miner, R. W. 1925. The pectoral limb of *Eryops* and other primitive tetrapods. Bulletin of the American Museum of Natural History 51:145–312.

Moodie, R. L. 1916. The Coal Measures Amphibia of North America. Publications of the Carnegie Institution of Washington 238:1–222.

Morales, M., and M. A. Shishkin. 2002. A re-assessment of *Parotosuchus africanus* (Broom), a capitosauroid temnospondyl amphibian from the Triassic of South Africa. Journal of Vertebrate Paleontology 22:1–11.

Morkovin, B. I. 2020. A new benthosuchid (Amphibia: Temnospondyli) from the Lower Triassic of the Moscow Syncline. Paleontological Journal 54:401–409.

Moulton, J. M. 1974. A description of the vertebral column of *Eryops* based on the notes and drawings of A. S.  Romer. Breviora 428:1–10.

Müller, H. 2006. Ontogeny of the skull, lower jaw, and hyobranchial skeleton of *Hypogeophis rostratus* (Amphibia: Gymnophiona: Caeciliidae) revisited. Journal of Morphology 267:968–986.

Müller, H., O. V. Oommen, and P. Bartsch. 2005. Skeletal development of the direct-developing caecilian *Gegeneophis ramaswamii* (Amphibia: Gymnophiona: Caeciliidae). Zoomorphology 124:171–188.

Murphy, J. L. 1971. Eryopsid remains from the Conemaugh Group, Braxton County, West Virginia. Southeastern Geology 13:265–273.

Nakajima, Y., and R. R. Schoch. 2011. The first temnospondyl amphibian from Japan. Journal of Vertebrate Paleontology 31:1154–1157.

Nedjari, A., R. Aït Ouali, L. Bitam, J.-S. Steyer, P. Taquet, R. Vacant, W. Bouzidi, and B. Kedadra. 2010. Découverte d'un nouveau gisement de stégocéphales d'une conservation exceptionnelle dans le Trias d'in Amenas (bassin d'Illizi, Algérie). Bulletin du Service Géologique National 21:1–18.

Nesov L. A., P. V. Fedorov, D. O. Potanov, and L. S. Golovnyeva. 1996. [The structure of the skulls of caudate amphibians collected from the Jurassic of Kirgizstan and the Cretaceous of Uzbekistan]. Vestnik Sankt-Petersburgiskogo Universiteta, Seriya 7, Geologiya, Geografiya 1:3–11. [Russian]

Nesov, L. A. 1988. Late Mesozoic amphibians and lizards of Soviet Middle Asia. Acta Zoologica Cracoviensia 31:475-486.

Nesov, L. A. 1990. [A Late Jurassic labyrinthodont (Amphibia, Labyrinthodontia) among other relict groups of vertebrates of northern Fergana]. Paleontologicheskii Zhurnal 1990:82–90. [Russian]

Nield, C. M., R. Damiani, and A. Warren. 2006. A short-snouted trematosauroid (Tetrapoda, Temnospondyli) from the Early Triassic of Australia: the oldest known trematosaurine. Alcheringa 30:263–271.

Nielsen, E. 1967. New observations on the skull-roof of the holotype of *Tupilakosaurus heilmani* Nielsen. Zoological Journal of the Linnean Society 47:225–229.

Nilsson, T. 1942. *Sassenisaurus*, a new genus of Eotriassic stegocephalian from Spitsbergen. Bulletin of the Geological Institution of the University of Upsala 30:91–102.

Nilsson, T. 1943. On the Morphology of the Lower Jaw of Stegocephalia with Special Reference to Eotriassic Stegocephalians from Spitsbergen. Descriptive Part I. Kungliga Svenska Vetenskapsakademiens Handlingar, 20.

Nilsson, T. 1946. A new find of *Gerrothorax rhaeticus*, a plagiosaurid from the Rhaetic of Scania. Lunds Universitets Årsskrift, N.F. 42:1–42.

Nilsson, T. 1946. On the Genus *Peltostega* Wiman and the Classification of the Triassic Stegocephalians. Kungliga Svenska Vetenskapsakademiens Handlingar, 23.

Nonsrirach, T., S. Manitkoon, and K. Lauprasert. 2021. First occurrence of brachyopid temnospondyls in Southeast Asia and review of the Mesozoic amphibians from Thailand. Fossil Record 24:33–47.

Nonsrirach, T., S. Manitkoon, and K. Lauprasert. 2021. First occurrence of brachyopid temnospondyls in Southeast Asia and review of the Mesozoic amphibians from Thailand. Fossil Record 24:33–47.

Novikov, I. V. 1986. [A new species of *Parotosuchus* (Amphibia: Labyrinthodontia) from the Triassic deposits of the Vychegda river basin. Paleontologicheskii Zhurnal 1986:129–131. [Russian]; Paleontological Journal 20:126–128. [English]

Novikov, I. V. 1990. [New early Triassic labyrinthodonts from Central Timan.] Paleontologicheskii Zhurnal 1990:87–100. [Russian]; Paleontological Journal 24:48–60. [English]

Novikov, I. V. 2007. New data on trematosauroid labyrinthodonts of eastern Europe: 1. Genus *Inflectosaurus* Shishkin, 1960. Paleontological Journal 41:167–174.

Novikov, I. V. 2010. New data on trematosauroid labyrinthodonts of Eastern Europe: 2. *Trematosaurus galae* sp. nov.: Cranial morphology. Paleontological Journal 44:457–467.

Novikov, I. V. 2012a. New data on trematosauroid labyrinthodonts of Eastern Europe: 3. *Qantas samarensis* gen. et sp. nov. Paleontological Journal 46:177–186.

Novikov, I. V. 2012b. New data on trematosauroid labyrinthodonts of Eastern Europe: 4. Genus *Benthosuchus* EFREMOV, 1937. Paleontological Journal 46:400–411.

Novikov, I. V. 2016. New temnospondyl amphibians from the basal Triassic of the Obshchii Syrt Highland, Eastern Europe. Paleontological Journal 50:297–310.

Novikov, I. V. 2018. New data on Early Triassic lonchorhynchids (Amphibia, Temnospondyli) of Eastern Europe. Paleontological Journal 52:1428–1431.

Novikov, I. V., and M. A. Shishkin. 1992. [New Middle Triassic labyrinthodonts from the Pechora Urals.] Paleontologicheskii Zhurnal 1992:71-80. [Russian]; Paleontological Journal 92–102. [English]

Novikov, I.V., and A. N. Orlov. 1992. [New early Triassic labyrinthodonts from Kolguyev Island.] Paleontologicheskii Zhurnal 1992:133–136. [Russian]; Paleontological Journal 26:80–84. [English]

Ochev, V. G. 1958. [New data on the fauna of Triassic vertebrates from the Orenburg Transural area.] Doklady Akademiya Nauk SSSR 122:485–488. [Russian]

Ochev, V. G. 1966. [Systematics and Phylogeny of Capitosauroid Labyrinthodonts.] Izdatel'stvo Saratovskogo Universiteta, Saratov, Russia, 184 pp. [Russian]

Ochev, V. G. 1972. [Capitosaurodi Labyrinthodonts from the South-Eastern European Part of the USSR.] Izdatel'stvo Saratovskogo Universiteta, Saratov, Russia, 269 pp. [Russian]

Olson, E. C. 1941. The family Trematopsidae. The Journal of Geology 49:149–176.

Olson, E. C. 1955. Fauna of the Vale and Choza: *Trimerorhachis*: including a revision of pre-Vale species. Fieldiana: Geology 10:225–274.

Olson, E. C. 1965. New Permian vertebrates from the Chickasha Formation in Oklahoma. Oklahoma Geological Survey Circular 70:1–70.

Olson, E. C. 1967. Early Permian vertebrates of Oklahoma. Oklahoma Geological Survey Circular 74:1–111.

Olson, E. C. 1970a. New and little known genera and species of vertebrates from the Lower Permian of Oklahoma. Fieldiana: Geology 18:358–434.

Olson, E. C. 1970b. *Trematops stonei* sp. nov. (Temnospondyli: Amphibia) from the Washington Formation, Dunkard Group, Ohio. Kirtlandia 8:1–12.

Olson, E. C. 1972. *Fayella chickashaensis*, the Dissorophoidea and the Permian terrestrial radiations. Journal of Paleontology 46:104–114.

Olson, E. C. 1991. An eryopid (Amphibia: Labyrinthodontia) from the Fort Sill fissures, Lower Permian, Oklahoma. Journal of Vertebrate Paleontology 11:130–132.

Pacheco, C. P., E. Eltink, R. T. Müller, and S. Dias-da-Silva. 2017. A new Permian temnospondyl with Russian affinities from South America, the new family Konzhukoviidae, and the phylogenetic status of Archegosauroidea. Journal of Systematic Palaeontology 15:241–256.

Panchen, A. L. 1959. A new armoured amphibian from the Upper Permian of East Africa. Philosophical Transactions of the Royal Society of London. Series B, Biological Sciences 242(691):207–281.

Pardo, J. D., and J. S. Anderson. 2016. Cranial morphology of the Carboniferous-Permian tetrapod *Brachydectes newberryi* (Lepospondyli, Lysorophia): new data from µCT. PLOS ONE 11:e0161823.

Pardo, J. D., B. J. Small, and A. K. Huttenlocker. 2017a. Stem caecilian from the Triassic of Colorado sheds light on the origins of Lissamphibia. PNAS:E5389–E5395.

Pardo, J. D., M. Szostakiwskyj, P. E. Ahlberg, and J. S. Anderson. 2017b. Hidden morphological diversity among early tetrapods. Nature, 546:642–645.

Parker, W. K. 1887. XV. On the structure and development of the skull in the urodelous amphibia.—Part I. Philosophical Transactions of the Royal Society of London 167:529–597.

Paton, R. L. 1975. A Lower Permian temnospondylous amphibian from the English Midlands. Palaeontology 18:831–845.

Pawley, K. 2007. The postcranial skeleton of *Trimerorhachis insignis* Cope, 1878 (Temnospondyli: Trimerorhachidae): a plesiomorphic temnospondyl from the Lower Permian of North America. Journal of Paleontology 81:873–894.

Pawley, K., 2006. The postcranial skeleton of temnospondyls (Tetrapoda: Temnospondyli). Ph.D. dissertation, La Trobe University, Melbourne, Victoria, 442 pp.

Pawley, K., and A. Warren. 2005. A terrestrial stereospondyl from the Lower Triassic of South Africa: the postcranial skeleton of *Lydekkerina huxleyi* (Amphibia: Temnospondyli). Palaeontology 48:281–298.

Pawley, K., and A. Warren. 2006. The appendicular skeleton of *Eryops megacephalus* COPE, 1877 (Temnospondyli: Eryopoidea) from the Lower Permian of North America. Journal of Paleontology 80:561–580.

Piñeiro, G., A. Ramos, and C. Marsicano, C. 2012. A rhinesuchid-like temnospondyl from the Permo-Triassic of Uruguay. Comptes Rendus Palevol, 11:65–78.

Piñeiro, G., C. A. Marsicano, and R. Damiani. 2007a. Mandibles of mastodonsaurid temnospondyls from the Upper Permian–Lower Triassic of Uruguay. Acta Palaeontologica Polonica, 52:695–703.

Piñeiro, G., C. A. Marsicano, C. Goso, and E. Morosi. 2007b. Temnospondyl diversity of the Permian-Triassic Colonia Orozco local fauna (Buena Vista Formation) of Uruguay. Revista Brasileira de Paleontologia 10:169–180.

Piñeiro, G., C. Marsicano, and N. Lorenzo. 2007c. A new temnospondyl from the Permo-Triassic Buena Vista Formation of Uruguay. Palaeontology 50:627–640.

Pledge, N. S. 2013. First South Australian labyrinthodont: a possible chigutisaurid stereospondyl amphibian from the Late Triassic at Leigh Creek. Transactions of the Royal Society of South Australia 137:127–134.

Polley, B. P., and R. R. Reisz. 2011. A new Lower Permian trematopid (Temnospondyli: Dissorophoidea) from Richards Spur, Oklahoma. Zoological Journal of the Linnean Society 161:789–815.

Ponssa, M. L. 2008. Cladistic analysis and osteological descriptions of the frog species in the *Leptodactylus fuscus* species group (Anura, Leptodactylidae). Journal of Zoological Systematics and Evolutionary Research 46:249–266.

Price, L. I. 1948. Um anfíbio labirintodonte da Formação Pedra de Fogo, estado do Maranhão. Boletim do Departamento Nacional de Produção Mineral, Divisão de Geologia e Mineralogia 124:–33.

Rage, J.-C., and Z. Roček. 1989. Redescription of *Triadobatrachus massinoti* (Piveteau, 1936) an anuran amphibian from the early Triassic. Palaeontographica A 206:1–16.

Ramos, B. D., and C. S. Vega. 2011. Temnospondyl remains from the late Permian Rio do Rasto Formation (Paraná Basin) of Brazil. Revista Brasileira de Paleontologia 14:67–74.

Reese, A. M. 1906. Anatomy of *Cryptobranchus allegheniensis*. The American Naturalist 40:287–326.

Reisz, R. R., D. S. Berman, and A. C. Henrici. 2005. A new skull of the cochleosaurid amphibian *Chenoprosopus* (Amphibia: Temnospondyli) from the Early Permian of New Mexico. New Mexico Museum of Natural History and Science Bulletin 30:253–255.

Reisz, R. R., R. R. Schoch, and J. S. Anderson. 2009. The armoured dissorophid *Cacops* from the Early Permian of Oklahoma and the exploitation of the terrestrial realm by amphibians. Naturwissenschaften 96:789–796.

Rieppel, O. 1980. The edopoid amphibian *Cochleosaurus* from the middle Pennsylvanian of Nova Scotia. Palaeontology 23:143–149.

Rinehart, L. F., and S. G. Lucas. 2016. *Eocyclotosaurus appetolatus, a* Middle Triassic amphibian: osteology, life history, and paleobiology. New Mexico Museum of Natural History & Science Bulletin 70:1–118.

Rinehart, L. F., S. G. Lucas, and R. R. Schoch. 2015. *Eocyclotosaurus appetolatus*, a new cyclotosaurid amphibian from the Middle Triassic (Perovkan) Moenkopi Formation of New Mexico, USA. Journal of Vertebrate Paleontology 35:e929140.

Robinson, J., P. E. Ahlberg, and G. Koentges. 2005. The braincase and middle ear region of *Dendrerpeton acadianum* (Tetrapoda: Temnospondyli). Zoological Journal of the Linnean Society 143:577–597.

Romer, A. S. 1947. Review of the Labyrinthodontia. Bulletin of the Museum of Comparative Zoology 99:1–367.

Romer, A. S. 1969. A temnospondylous labyrinthodont from the Lower Carboniferous. Kirtlandia 6:1–20.

Romer, A. S., and R. V. Witter. 1942. *Edops*, a primitive rhachitomous amphibian from the Texas red beds. The Journal of Geology 50:925–960.

Ronchi, A., and A. Tintori. 1997. First amphibian find in early Permian from Sardinia (Italy). Rivista Italiana di Paleontologia e Stratigrafia 103:29–38.

Ronquist, F., M. Teslenko, P. Van Der Mark, D. L. Ayres, A. Darling, S. Höhna, B. Larget, L. Liu, M. A. Suchard, and J. P. Huelsenbeck. 2012. MrBayes 3.2: efficient Bayesian phylogenetic inference and model choice across a large model space. Systematic Biology 61:539–542.

Rusconi, C. 1948. Nuevos laberintodontes del triásico de Mendoza.Revista del Museo de Historia Natural de Mendoza 2:225–229.

Rusconi, C. 1949. Otra especie de laberintodonte del triásico de Mendoza.Revista del Museo de Historia Natural deMendoza 3:93–94.

Rusconi, C. 1951. Laberintodontes triásicos y pérmicos de Mendoza.Revista del Museo de Historia Natural de Mendoza 5:33–158.

Rusconi, C. 1953. Nuevo laberintodonte de Mendoza. Nota previa. Boletín Paleontológico de Buenos Aires 26:1.

Rusconi, C. 1955. Acerca de los grandes laberintodontes de Cacheuta.Revista del Museo de Historia Natural de Mendoza 8:85–101.

Ruta, M., and J. R. Bolt. 2006. A reassessment of the temnospondyl amphibian *Perryella olsoni* from the Lower Permian of Oklahoma. Earth and Environmental Science Transactions of The Royal Society of Edinburgh 97:113–165.

Ruta, M., and J. R. Bolt. 2008. The brachyopoid *Hadrokkosaurus bradyi* from the early Middle Triassic of Arizona, and a phylogenetic analysis of lower jaw characters in temnospondyl amphibians. Acta Palaeontologica Polonica 53:579–592.

Sanchez, S., G. De Ploeg, G. Clément, and P. E. Ahlberg. 2010. A new tool for determining degrees of mineralization in fossil amphibian skeletons: the example of the Late Palaeozoic branchiosaurid *Apateon* from the Autun Basin, France. Comptes Rendus Palevol 9:311–317.

Sander, M. P., and C. Meyer. 1991. A labyrinthodont jaw fragment from the marine Triassic of the Alps. Neues Jahrbuch für Geologie und Paläontologie-Monatshefte 1991:222–232.

Säve-Söderbergh, G. 1935. On the dermal bones of the head in labyrinthodont stegocephalians and primitive reptilia with special reference to Eotriassic stegocephalians from East Greenland. Meddelelser om Grønland 98:1–211.

Säve-Söderbergh, G. 1935. On the dermal bones of the head in labyrinthodont stegocephalians and primitive Reptilia. Meddelelser om Grønland 98:1–211.

Säve-Söderbergh, G. 1936. On the morphology of Triassic stegocephalians from Spitsbergen, and the interpretation of the endocranium in the Labyrinthodontia. Kungliga Svenska Vetenskapsakademiens Handlingar 16:1–181.

Säve-Söderbergh, G. 1936. On the morphology of Triassic stegocephalians from Spitsbergen, and the interpretation of the endocranium in the Labyrinthodontia. Kungliga Svenska Vetenskaps Akademiens Handlingar 16:1–181.

Säve-Söderbergh, G. 1937. On the dermal skulls of *Lyrocephalus*, *Aphaneramma*, and *Benthosaurus*, labyrinthodonts from the Triassic of Spitsbergen and N. Russia. Bulletin of the Geological Institution of the University of Uppsala 27:189–208.

Scheyer, T. M., U. Oberli, N. Klein, and H. Furrer. 2022. A large osteoderm-bearing rib from the Upper Triassic Kössen Formation (Norian/Rhaetian) of eastern Switzerland. Swiss Journal of Palaeontology 141:1–9.

Schoch, R. R. 1997. Cranial anatomy of the Permian temnospondyl amphibian *Zatrachys serratus* Cope 1878, and the phylogenetic position of the Zatrachydidae. Neues Jahrbuch für Geologie und Paläontologie-Abhandlungen 206:223–248.

Schoch, R. R. 1999. Comparative osteology of *Mastodonsaurus giganteus* (Jaeger, 1828) from the Middle Triassic (Lettenkeuper: Longobardian) of Germany (Baden-Württemberg, Bayern, Thüringen*)*. Stuttgarter Beiträge zur Naturkunde Serie B 278:1–175.

Schoch, R. R. 2000. The status and osteology of two new cyclotosaurid amphibians from the Upper Moenkopi Formation of Arizona (Amphibia: Temnospondyli; Middle Triassic). Neues Jahrbuch für Geologie und Paläontologie-Abhandlungen 216:387–411.

Schoch, R. R. 2003. Early larval ontogeny of the Permo-Carboniferous temnospondyl *Sclerocephalus*. Palaeontology 46:1055–1072.

Schoch, R. R. 2006. A complete trematosaurid amphibian from the Middle Triassic of Germany. Journal of Vertebrate Paleontology 26:29–43.

Schoch, R. R. 2008a. A new stereospondyl from the German Middle Triassic, and the origin of the Metoposauridae. Zoological Journal of the Linnean Society 152:79–113.

Schoch, R. R. 2008b. The Capitosauria (Amphibia): characters, phylogeny, and stratigraphy. Palaeodiversity 1:189–226.

Schoch, R. R. 2009. Life-cycle evolution as response to diverse lake habitats in Paleozoic amphibians. Evolution 63:2738–2749.

Schoch, R. R. 2011. A trematosauroid temnospondyl from the Middle Triassic of Jordan. Fossil Record 14:119–127.

Schoch, R. R. 2012. Character distribution and phylogeny of the dissorophid temnospondyls. Fossil Record 15:121–137.

Schoch, R. R. 2013. The evolution of major temnospondyl clades: an inclusive phylogenetic analysis. Journal of Systematic Palaeontology 11:673–705.

Schoch, R. R. 2017 (for 2019). The stapes of *Edops craigi* and ear evolution in the lissamphibian stem group. Acta Zoologica 100:126–134.

Schoch, R. R. 2018a. Osteology of the temnospondyl *Neldasaurus* and the evolution of basal dvinosaurians. Neues Jahrbuch für Geologie und Paläontologie-Abhandlungen 287:1–16.

Schoch, R. R. 2018b (for 2019b). The putative lissamphibian stem-group: phylogeny and evolution of the dissorophoid temnospondyls. Journal of Paleontology 93:137–156.

Schoch, R. R. 2018c. The temnospondyl *Parotosuchus nasutus* (v. Meyer, 1858) from the Early Triassic Middle Buntsandstein of Germany. Palaeodiversity 11:107–126.

Schoch, R. R. 2019a. Osteology of the temnospondyl *Trematosaurus brauni* Burmeister, 1849 from the Middle Buntsandstein of Bernburg, Germany. Palaeodiversity 12:41–63.

Schoch, R. R. 2021a. Osteology of the Permian temnospondyl amphibian *Glanochthon lellbachae* and its relationships. Fossil Record 24:49–64.

Schoch, R. R. 2021b. The life cycle in late Paleozoic eryopid temnospondyls: developmental variation, plasticity and phylogeny. Fossil Record 24:295–319.

Schoch, R. R., A. C. Henrici, and R. W. Hook. 2021. A new dissorophoid temnospondyl from the Allegheny Group (late Carboniferous) of five points, Mahoning County, Ohio (USA). Journal of Paleontology 95:638–651.

Schoch, R. R., and A. R. Milner. 2000. Stereospondyli; pp. 1–203 in P. Wellnhofer (ed.), Encyclopedia of Paleoherpetology, Part 3B. Verlag Dr. Friedrich Pfeil, Munich.

Schoch, R. R., and A. R. Milner. 2008. The intrarelationships and evolutionary history of the temnospondyl family Branchiosauridae. Journal of Systematic Palaeontology 6:409–431.

Schoch, R. R., and A. R. Milner. 2014. Temnospondyli I; pp. 1–150 in H.-D. Sues (ed.), Encyclopedia of Paleoherpetology, Part 3A2. Verlag Dr. Friedrich Pfeil, Munich.

Schoch, R. R., and A. R. Milner. 2021. Morphology and relationships of the temnospondyl *Macrerpeton huxleyi* from the Pennsylvanian of Linton, Ohio (USA). Neues Jahrbuch für Geologie und Paläontologie-Abhandlungen 299:77–98.

Schoch, R. R., and B. S. Rubidge. 2005. The amphibamid *Micropholis* from the *Lystrosaurus* Assemblage Zone of South Africa. Journal of Vertebrate Paleontology 25:502–522.

Schoch, R. R., and F. Witzmann. 2009a. Osteology and relationships of the temnospondyl genus *Sclerocephalus*. Zoological Journal of the Linnean Society 157:135–168.

Schoch, R. R., and F. Witzmann. 2009b. The temnospondyl *Glanochthon* from the Lower Permian Meisenheim Formation of Germany. Special Papers in Palaeontology 81:121–136.

Schoch, R. R., and F. Witzmann. 2012. Cranial morphology of the plagiosaurid *Gerrothorax pulcherrimus* as an extreme example of evolutionary stasis. Lethaia 45:371–385.

Schoch, R. R., and F. Witzmann. 2018. Morphology of the Late Carboniferous temnospondyl *Limnogyrinus elegans*, and the evolutionary history of the Micromelerpetidae. Neues Jahrbuch für Geologie und Paläontologie-Abhandlungen 299:293–310.

Schoch, R. R., and N. Fröbisch. 2006. Metamorphosis and neoteny: alternative pathways in an extinct amphibian clade. Evolution 60:1467–1475.

Schoch, R. R., and S. Voigt. 2019. A dvinosaurian temnospondyl from the Carboniferous-Permian boundary of Germany sheds light on dvinosaurian phylogeny and distribution. Journal of Vertebrate Paleontology 39:e1577874.

Schoch, R. R., R. Werneburg, R., and S. Voigt, S. 2020b. A Triassic stem-salamander from Kyrgyzstan and the origin of salamanders. Proceedings of the National Academy of Sciences 117:11584–11588.

Sengupta, D. P. 1995. Chigutisaurid temnospondyls from the Late Triassic of India and a review of the Family Chigutisauridae. Palaeontology 38:313–339.

Sengupta, D. P. 2002. Indian metoposaurid amphibians revised. Paleontological Research 6:41–65.

Sengupta, D. P. 2003. Triassic temnospondyls of the Pranhita–Godavari basin, India. Journal of Asian Earth Sciences 21:655–662.

Sennikov, A. G. 1981. [A new wetlugasaur from the Samara River Basin.] Paleontologicheskii Zhurnal 1981:143–148. [Russian]; Paleontological Journal 15:111–116. [English]

Sennikov, A. G., and I. V. Novikov. 2018. On possible trophic adaptations of some Rhytidosteidae (Amphibia, Temnospondyli). Paleontological Journal 52:1412–1418.

Sequeira, S. E. K. 1996. A cochleosaurid amphibian from the Upper Carboniferous of Ireland. Special Papers in Palaeontology 52:65-80.

Sequeira, S. E. K. 1998. The cranial morphology and taxonomy of the saurerpetontid *Isodectes obtusus* comb. nov.(Amphibia: Temnospondyli) from the Lower Permian of Texas. Zoological Journal of the Linnean Society 122:237–259.

Sequeira, S. E. K. 2004 (for 2003). The skull of *Cochleosaurus bohemicus* Frič, a temnospondyl from the Czech Republic (Upper Carboniferous) and cochleosaurid interrelationships. Earth and Environmental Science Transactions of The Royal Society of Edinburgh 94:21–43.

Sequeira, S. E. K. 2009. The postcranium of *Cochleosaurus bohemicus* Frič, a primitive Upper Carboniferous temnospondyl from the Czech Republic. Special Papers in Palaeontology 81:137–153.

Sequeira, S. E. K., and A. R. Milner. 1993. The temnospondyl amphibian *Capetus* from the Upper Carboniferous of the Czech Republic. Palaeontology 36:657–680.

Sereno, P. C. 2007. Logical basis for morphological characters in phylogenetics. Cladistics 23:565–587.

Shishkin, M. A. 1960a. [Yarengiidae, a new family of Triassic labyrinthodonts.] Paleontologicheskii Zhurnal 1960:97–106. [Russian]

Shishkin, M. A. 1960b. [*Inflectosaurus amplus*, a new Triassic trematosaurid.] Paleontologicheskii Zhurnal 1960:130–148. [Russian]

Shishkin, M. A. 1961. [Recent data on *Tupilakosaurus*]. Doklady Akademiya Nauk SSSR 136:938–941. [Russian]

Shishkin, M. A. 1966. [A brachyopid labyrinthodont from the Triassic of the Russian Platform.] Paleontologicheskii Zhurnal 1966:93–108. [Russian]

Shishkin, M. A. 1967. [Plagiosauria in the Triassic of the USSR.] Paleontologicheskii Zhurnal 1967:92–99. [Russian]

Shishkin, M. A. 1968. On the cranial arterial system of the labyrinthodonts. Acta Zoologica 49:1–22.

Shishkin, M. A. 1973. [The morphology of the early Amphibia and some problems of the lower tetrapod evolution]. Trudy Paleontologicheskogo Instituta Akademiya Nauk SSSR 137:1–260. [Russian]

Shishkin, M. A. 1980. [The Luzocephalidae, a new Triassic labyrinthodont family.] Paleontologicheskii Zhurnal 1980:104–119. [Russian]; Paleontological Journal 14:88–101. [English]

Shishkin, M. A. 1986. [New data on plagiosaurs from the Triassic of the USSR]. Byulleten Moskovskogo Obschchestva Ispytatelei Prirody, Otdel Geologischeskii 61:97–102. [Russian]

Shishkin, M. A. 1987. [The evolution of early amphibians (Plagiosauroidea)]. Trudy Paleontologicheskogo Instituta Akademia Nauk SSSR 225:1–143. [Russian]

Shishkin, M. A. 1989. The axial skeleton of early amphibians and the origin of resegmentation in tetrapod vertebrae. Fortschritte der Zoologie 35:180–195.

Shishkin, M. A. 1991. [A labyrinthodont from the Late Jurassic of Mongolia]. Paleontologicheskii Zhurnal 1991:81–95. [Russian].

Shishkin, M. A. 1994. [A Gondwanan rhytidosteid (Amphibia, Temnospondyli) from the Lower Triassic of southern Cisuralia. Paleontologicheskii Zhurnal 1994:97–110. [Russian]; Paleontological Journal 28:127–143. [English]

Shishkin, M. A. 1998. [*Tungussogyrinus* – a relict neotenic dissorophoid (Amphibia, Temnospondyli) from the Permo- Triassic of Siberia.] Paleontologicheskii Zhurnal 1998: 85–96. [Russian]

Shishkin, M. A. 2000. Mesozoic amphibians from Mongolia and the Central Asiatic republics; pp. 297–308 in M. J. Benton, M. A. Shishkin, D. M. Unwin, and E. N. Kurochkin (eds)., The Age of Dinosaurs in Russia and Mongolia. Cambridge University Press: Cambridge.

Shishkin, M. A., and B. S. Rubidge. 2000. A relict rhinesuchid (Amphibia: Temnospondyli) from the Lower Triassic of South Africa. Palaeontology 43:653–670.

Shishkin, M. A., and J. Welman. 1994. A new find of *Trematosuchus* (Amphibia, Temnospondyli) from the *Cynognathus* zone of South Africa. Palaeontologia africana 31:39–49.

Shishkin, M. A., and M. N. Vavilov. 1985. [A rhytidosteid (Amphibia, Labyrinthodontia) find from the Triassic of the USSR. Doklady Akadamie Nauk SSSR 282:971–975. [Russian]; Doklady Akadamie Nauk SSSR (Earth Sciences) 282:233–236. [English]

Shishkin, M. A., and T. Sulej. 2009. The Early Triassic temnospondyls of the Czatkowice 1 tetrapod assemblage. Palaeontologia Polonica 65:31–77.

Shishkin, M. A., and V. Lozovsky. 1979. [A labyrinthodont from the Triassic of southern Primov'ye.] Doklady Akadamie Nauk SSSR 246:201–205. [Russian]

Shishkin, M. A., B. S. Rubidge, and J. W. Kitching. 1996. A new lydekkerinid (Amphibia, Temnospondyli) from the lower Triassic of South Africa: implications for evolution of the early capitosauroid cranial pattern. Philosophical Transactions of the Royal Society of London. Series B: Biological Sciences 351:1635–1659.

Sidor, C. A. 2013. The vertebrate fauna of the Upper Permian of Niger–VIII. *Nigerpeton ricqlesi* (Temnospondyli: Cochleosauridae) and tetrapod biogeographic provinces. Comptes Rendus Palevol 12:463–472.

Sidor, C. A., F. R. O'Keefe, R. Damiani, J.-S. Steyer, R. M. Smith, H. C. Larsson, P. C. Sereno, O. Ide, and A. Maga. 2005. Permian tetrapods from the Sahara show climate-controlled endemism in Pangaea. Nature 434:886–889.

Sidor, C. A., J.-S. Steyer, and R. Damiani. 2007. *Parotosuchus* (Temnospondyli: Mastodonsauridae) from the Triassic of Antarctica. Journal of Vertebrate Paleontology 27:232–235.

Sidor, C. A., J.-S. Steyer, and W. R. Hammer. 2014. A new capitosauroid temnospondyl from the Middle Triassic upper Fremouw Formation of Antarctica. Journal of Vertebrate Paleontology 34:539–548.

Sidor, C. A., R. Damiani, and W. R. Hammer. 2008. A new Triassic temnospondyl from Antarctica and a review of Fremouw Formation biostratigraphy. Journal of Vertebrate Paleontology 28:656–663.

Sigurdsen, T., and D. M. Green. 2011. The origin of modern amphibians: a re-evaluation. Zoological Journal of the Linnean Society 162:457–469.

Sigurdsen, T., and J. R. Bolt. 2010. The Lower Permian amphibamid *Doleserpeton* (Temnospondyli: Dissorophoidea), the interrelationships of amphibamids, and the origin of modern amphibians. Journal of Vertebrate Paleontology 30:1360-1377.

Simões, T. R., M. W. Caldwell, A. Palci, and R. L. Nydam. 2017. Giant taxon-character matrices: quality of character constructions remains critical regardless of size. Cladistics 33:198–219.

Skutschas, P. P., T. Martin, and S. Ge. 2009. The co-occurrence of non-lissamphibian temnospondyls and salamanders in the Late Jurassic of the southern Junggar Basin (Xinjiang Autonomous Region, NW China). Proceedings of the Zoological Institute RAS 313:143–148.

Skutschas, P., and T. Martin. 2011. Cranial anatomy of the stem salamander *Kokartus honorarius* (Amphibia: Caudata) from the Middle Jurassic of Kyrgyzstan. Zoological Journal of the Linnean Society 161:816–838.

Slodownik, M.A., Mörs, T. and Kear, B.P., 2021. Reassessment of the Early Triassic trematosaurid temnospondyl Tertrema acuta from the Arctic island of Spitsbergen. *Journal of Vertebrate Paleontology*, *41*(1), p.e1900209.

Smithson, T. R. 1982. The cranial morphology of *Greererpeton burkemorani* Romer (Amphibia: Temnospondyli). Zoological Journal of the Linnean Society 76:29–90.

Soussi, M., G. Niedźwiedzki, M. Tałanda, D. Dróżdż, T. Sulej, K. Boukhalfa, J. Mermer, and B. Błażejowski. 2017. Middle Triassic (Anisian-Ladinian) Tejra red beds and Late Triassic (Carnian) carbonate sedimentary records of southern Tunisia, Saharan Platform: Biostratigraphy, sedimentology and implication on regional stratigraphic correlations. Marine and Petroleum Geology 79:222–256.

Spielmann, J. A., and S. G. Lucas. 2012. Tetrapod fauna of the Upper Triassic Redonda Formation east-central New Mexico: the characteristic assemblage of the Apachean land-vertebrate faunachron. New Mexico Museum of Natural History and Science Bulletin 55:1–119.

Steen, M. C. 1934. The amphibian fauna from the South Joggins. Nova Scotia. Proceedings of the Zoological Society of London 104:465–504.

Steen, M. C. 1938. On the Fossil Amphibia from the Gas Coal of Nýřany and other Deposits in Czechoslovakia. Proceedings of the Zoological Society of London 108:205–283.

Steyer, J.-S. 2002. The first articulated trematosaur 'amphibian' from the Lower Triassic of Madagascar: implications for the phylogeny of the group. Palaeontology 45:771–793.

Steyer, J.-S. 2003. A revision of the early Triassic "capitosaurs" (Stegocephali, Stereospondyli) from Madagascar, with remarks on their comparative ontogeny. Journal of Vertebrate Paleontology 23:544–555.

Steyer, J.-S., and N.-E. Jalil. 2009. First evidence of a temnospondyl in the Late Permian of the Argana Basin, Morocco. Special Papers in Palaeontology 81:155–160.

Steyer, J.-S., and R. Damiani. 2005. A giant brachyopoid temnospondyl from the Upper Triassic or Lower Jurassic of Lesotho. Bulletin de la Societe géologique de France 176:243–248.

Steyer, J.-S., B. R. Peecook, T. Arbez, S. J. Nesbitt, S. Tolan, M. R. Stocker, R. M. Smith, K. D. Angielczyk, and C. A. Sidor. 2021. New data on the Triassic temnospondyls from the Karoo rift basins of Tanzania and Zambia. Geodiversitas 43:365–376.

Steyer, J.-S., D. Heyler, and L. Guillot. 2000. Nouvelles données sur les stégocéphales du bassin de Souvigny (Permien inférieur de l'Allier, Massif central). Revue Scientifique du Bourbonnais 98:80-88.

Steyer, J.-S., F. Escuillie, J. M. Pouillon, J. Broutin, P. Debriette, P. Freytet, G. Gand, C. Poplin, C., J.-C. Rage, J. Rival, and J. W. Schneider. 2000. New data on the flora and fauna from the? uppermost Carboniferous-Lower Permian of Buxieres-les-Mines, Bourbon l'Archambault Basin (Allier, France); a preliminary report. Bulletin de la Société Géologique de France 171:239–249.

Steyer, J.-S., G. Gand, and J. M. Pouillon 1997. Les amphibiens du Paléozoïque et du Trias français: historique et inventaire. Bulletin de la Société d'Histoire naturelle d'Autun 162:23–40.

Steyer, J.-S., R. Damiani, C. A. Sidor, F. R. O'Keefe, H. C. Larsson, A. Maga, and O. Ide. 2006. The vertebrate fauna of the Upper Permian of Niger. IV. *Nigerpeton ricqlesi* (Temnospondyli: Cochleosauridae), and the edopoid colonization of Gondwana. Journal of Vertebrate Paleontology 26:18–28.

Stokely, P. S., and P. A. Holle. 1954. Appendicular skeleton of the Ambystomidae. Herpetologica 10:57–61.

Strapasson, A., F. L. Pinheiro, and M. B. Soares. 2014. On a new stereospondylomorph temnospondyl from the Middle—Late Permian of Southern Brazil. Acta Palaeontologica Polonica 60:843–855.

Strong, E. E., and D. Lipscomb. 1999. Character coding and inapplicable data. Cladistics 15:363–371.

Stur, D. 1873. *Mastodonsaurus giganteus* JÄGER im Lunzersandstein der Grube Prinzbach bei Kirchberg an der Pielach, in der nordöstlichen Kalkalpen. Verhandlungen der Kaiserlich-Königlichen Geologischen Reichsanstalt 1873:18–19.

Sues, H.-D., and P. Olsen. 2015. Stratigraphic and temporal context and faunal diversity of Permian-Jurassic continental tetrapod assemblages from the Fundy rift basin, eastern Canada. Atlantic Geology 51:139–205.

Sues, H.-D., and R. R. Schoch. 2013. Anatomy and phylogenetic relationships of *Calamops paludosus* (Temnospondyli, Stereospondyli) from the Triassic of the Newark Basin, Pennsylvania. Journal of Vertebrate Paleontology 33:1061–1070.

Sues, H.-D., P. E. Olsen, T. J. Fedak, and R. R. Schoch. 2021. Diverse assemblage of Middle Triassic continental tetrapods from the Newark Supergroup of Nova Scotia (Canada). Journal of Vertebrate Paleontology 41:e2023168.

Sulej, T. 2002. Species discrimination of the Late Triassic temnospondyl amphibian *Metoposaurus diagnosticus*. Acta Palaeontologica Polonica 47:535–546.

Sulej, T. 2007. Osteology, variability, and evolution of *Metoposaurus*, a temnospondyl from the Late Triassic of Poland. Palaeontologia Polonica 64:29–139.

Sulej, T., and D. Majer. 2005. The temnospondyl amphibian *Cyclotosaurus* from the Upper Triassic of Poland. Palaeontology 48:157–170.

Sulej, T., and G. Niedźwiedzki. 2013. A new large capitosaurid temnospondyl amphibian from the Early Triassic of Poland. Acta Palaeontologica Polonica 58:65–75.

Sullivan, C., R. R. Reisz, and W. J. May. 2000. Large dissorophoid skeletal elements from the Lower Permian Richards Spur fissures, Oklahoma, and their paleoecological implications. Journal of Vertebrate Paleontology 20:456–461.

Sumida, S. S., D. S. Berman, and T. Martens. 1998. A new trematopid amphibian from the Lower Permian of central Germany. Palaeontology 41:605–630.

Sun, A. 1962. Discovyer of neorhachitomous vertebrae from Lufeng, Yunnan. Vertebrata PalAsiatic 6:109–110.

Sushkin, P. P. 1927. On the modifications of the mandibular and hyoid arches and their relations to the braincase in the early Tetrapoda. Paläontologische Zeitschrift 8:263–321.

Suteethorn, V., P. Janvier, and M. Morales. 1988. Evidence for a plagiosauroid amphibian in the Upper Triassic Huai Hin Lat Formation. Journal of Southeast Asian Earth Sciences 2:185–187.

Swinton, W. E. 1956. A neorachitome amphibian from Madagascar. Annals and Magazine of Natural History 97:60–64.

Szostakiwskyj, M., J. D. Pardo, and J. S. Anderson. 2015. Micro-CT study of *Rhynchonkos stovalli* (Lepospondyli, Recumbirostra), with description of two new genera. PLOS ONE 10:e0127307.

Theska, T., M. Wilkinson, D. J. Gower, and H. Müller. 2019. Musculoskeletal development of the Central African caecilian *Idiocranium russeli* (Amphibia: Gymnophiona: Indotyphlidae) and its bearing on the re-evolution of larvae in caecilian amphibians. Zoomorphology 138:137–158.

Tihen, J. A. 1958. Comments on the osteology and phylogeny of ambystomid salamanders. Bulletin of the Florida State Museum 3:1–50.

Trautschold, H. 1884. *Die Reste Permischer Reptilien des Paläontologischen Kabinets der Universität Kasan*, 38Moscow: l'Université impériale.

Tripathi, C. 1961a. Fossil labyrinthodonts from the Panchet Series of the Raniganj coalfield. Indian Minerals 15:198–199.

Tripathi, C. 1961b. *Rhinesuchus wadiai* sp. nov.: a new labyrinthodont from Vindhya Pradesh. Records of the Geological Survey of India 89:399–405.

Tripathi, C. 1969. Fossil labyrinthodonts from the Panchet Series of the Indian Gondwanas; Palaeontologia Indica 38:1–45.

Urban, M., and D. S. Berman. 2007. First occurrence of the Late Paleozoic amphibian *Zatrachys serratus* (Temnospondyli, Zatrachydidae) in the eastern United States. Annals of Carnegie Museum 76:157–164.

Van Hoepen, E. C. N. 1915. Stegocephalia of Senekal, OFS. Annals of the Transvaal Museum 5:125–149.

Vassilieva, A. B., J. S. Lai, S. F. Yang, Y. H. Chang, and N. A. Poyarkov Jr. 2015. Development of the bony skeleton in the Taiwan salamander, *Hynobius formosanus* Maki, 1922 (Caudata: Hynobiidae): heterochronies and reductions. Vertebrate Zoology 65:117–130.

Vaughn, P. P. 1962. Vertebrates from the Halgaito Tongue of the Cutler Formation, Permian of San Juan County, Utah. Journal of Paleontology 36:529–539.

Vaughn, P. P. 1963. The age and locality of the late Paleozoic vertebrates from El Cobre Canyon, Rio Arriba County, New Mexico. Journal of Paleontology 37:283–286.

Vaughn, P. P. 1964. Vertebrates from the organ rock shale of the Cutler Group, Permian of Monument Valley and vicinity, Utah and Arizona. Journal of Paleontology 38:567–583.

Vaughn, P. P. 1969. Further evidence of close relationship of the trematopsid and dissorophid labyrinthodont amphibians with a description of a new genus and new species. Bulletin of the Southern California Academy of Sciences 68:121–130.

Vaughn, P. P. 1971. A *Platyhystrix*-like amphibian with fused vertebrae, from the Upper Pennsylvanian of Ohio. Journal of Paleontology 45:464–469.

Vjushkov, B. P. 1955. [Concerning the fauna of the Upper Permian terrestrial vertebrates from the Malaia Kinel' River.] Trudy Paleozoologicheskogo Instituta Akademia Nauk SSSR 49:176–189. [Russian]

von Huene, F. 1922. Beiträge zur Kenntnis der Organisation einiger Stegocephalen der schwäbischen Trias. Acta Zoologica 3:395–400.

von Huene, F. 1958. The first Chinese labyrinthodonts. Vertebrata PalAsiatica 2:101–102.

von Huene, F., and M. R. Sahni. 1958. On *Indobrachyops panchetensis*, gen. et sp. nov. from the Upper Panchets (Lower Trias) of the Raniganj Coalfield. Palaeontological Society of India Monographs 2:1–17.

Wake, M. H., and J. Hanken. 1982. Development of the skull of *Dermophis mexicanus* (Amphibia: Gymnophiona), with comments on skull kinesis and amphibian relationships. Journal of Morphology 173:203–223.

Warren, A. 1981. A horned member of the labyrinthodont super-family Brachyopoidea from the Early Triassic of Queensland. Alcheringa 5:273–288.

Warren, A. 1985. Triassic Australian plagiosauroid. Journal of Paleontology 59:236–241.

Warren, A. 1998a. *Laidleria* uncovered: a redescription of *Laidleria gracilis* Kitching (1957), a temnospondyl from the *Cynognathus* Zone of South Africa. Zoological Journal of the Linnean Society 122:167-185.

Warren, A. 1998b. Karoo tupilakosaurid: a relict from Gondwana. Earth and Environmental Science Transactions of The Royal Society of Edinburgh 89:145–160.

Warren, A. A. 1980. *Parotosuchus* from the early Triassic of Queensland and Western Australia. Alcheringa 4:25–36.

Warren, A. A., and L. Davey. 1992. Folded teeth in temnospondyls—a preliminary study. Alcheringa 16:107–132.

Warren, A. A., and M. N. Hutchinson. 1983. The last labyrinthodont? A new brachyopoid (Amphibia, Temnospondyli) from the early Jurassic Evergreen Formation of Queensland, Australia. Philosophical Transactions of the Royal Society of London. B, Biological Sciences 303:1–62.

Warren, A. A., and M. N. Hutchinson. 1987. The skeleton of a new hornless rhytidosteid (Amphibia, Temnospondyli). Alcheringa 11:291–302.

Warren, A. A., and M. N. Hutchinson. 1988. The Madagascan capitosaurs. Bulletin du Muséum National d'Histoire Naturelle 10C:23–30.

Warren, A. A., and M. N. Hutchinson. 1990. *Lapillopsis*, a new genus of temnospondyl amphibians from the Early Triassic of Queensland. Alcheringa 14:149–158.

Warren, A. A., and R. Damiani. 1999. Stereospondyl amphibians from the Elliot Formation of South Africa. Palaeontologia africana 35:45–54.

Warren, A. A., B. S. Rubidge, I. G. Stanistreet, H. Stollhofen, A. Wanke, E. M. Latimer, C. A. Marsicano, and R. J. Damiani. 2001. Oldest known stereospondylous amphibian from the Early Permian of Namibia. Journal of Vertebrate Paleontology 21:34–39.

Warren, A. A., R. Damiani, and A. M. Yates. 2006. The South African stereospondyl *Lydekkerina huxleyi* (Tetrapoda, Temnospondyli) from the Lower Triassic of Australia. Geological Magazine 143:877–886.

Warren, A. A., R. J. Damiani, and A. M. Yates. 2001. Palaeobiogeography of Australian fossil amphibians. Historical Biology 15:171–179.

Warren, A., A. C. Rozefelds, and S. Bull. 2011. Tupilakosaur-like vertebrae in *Bothriceps australis*, an Australian brachyopid stereospondyl. Journal of Vertebrate Paleontology 31:738–753.

Warren, A., A. M. Yates, R. J. Damiani, M. B. Goodwin, C. B. Wood, and C. R. Schaff. 1998. The first temnospondyl amphibian (Stereospondyli: Capitosauroidea) from Ethiopia. Neues Jahrbuch für Geologie und Paläontologie-Monatshefte 1998:694–704.

Warren, A., and C. A. Marsicano. 2000. A phylogeny of the Brachyopoidea (Temnospondyli, Stereospondyli). Journal of Vertebrate Paleontology 20:462–483.

Warren, A., and C. Marsicano. 1998. Revision of the Brachyopidae (Temnospondyli) from the Triassic of the Sydney, Carnarvon and Tasmania basins, Australia. Alcheringa 22:32–342.

Warren, A., and N. Snell. 1991. The postcranial skeleton of Mesozoic temnospondyl amphibians: a review. Alcheringa 15:43–64.

Warren, A., and T. Black. 1985. A new rhytidosteid (Amphibia, Labyrinthodontia) from the Early Triassic Arcadia Formation of Queensland, Australia, and the relationships of Triassic temnospondyls. Journal of Vertebrate Paleontology 5:303–327.

Warren, A., R. Damiani, and D. P. Sengupta. 2009. Unique stereospondyl mandibles from the early Triassic Panchet formation of India and the Arcadia formation of Australia. Special Papers in Palaeontology 81:161–173.

Warren, A., T. H. Rich, and P. Vickers-Rich. 1997. The last last labyrinthodonts. Palaeontographica Abteilung A 247:1–24.

Watson, D. M. S. 1921. On *Eugyrinus wildi* (ASW), a branchiosaur from the Lancashire Coal-measures. Geological Magazine 58:70–74.

Watson, D. M. S. 1956. The brachyopid labyrinthodonts. Bulletin of the British Museum (Natural History), Geology 2:317–391.

Watson, D. M. S. 1958. A new labyrinthodont (*Paracyclotosaurus*) from the Upper Trias of New South Wales. Bulletin of the British Museum of Natural History, London (Geology) 3:233–263.

Watson, D. M. S. 1962. The evolution of the labyrinthodonts. Philosophical Transactions of the Royal Society of London. Series B, Biological Sciences 245:219–265.

Welles, S. P. 1965. A revision of the labyrinthodont family Capitosauridae and a description of *Parotosaurus peabodyi*, n. sp. from the Wupatki member of the Moenkopi Formation of northern Arizona. University of California Publications in Geological Sciences 54:1–148.

Welles, S. P. 1967. Arizona's giant amphibians. Pacific Discovery 20:10–15.

Welles, S. P. 1993, A review of the lonchorhychine trematosaurs (Labyrinthodontia), and a description of a new genus and species from the lower Moenkopi Formation of Arizona. Paleobios 14:1–24.

Welles, S. P., and R. Estes. 1969. *Hadrokkosaurus bradyi* from the Upper Moenkopi Formation of Arizona, with a review of the brachyopid labyrinthodonts. University of California Publications in Geological Sciences 84:1–56.

Werneburg, R. 1991. Die Branchiosaurier aus dem Unterrotliegend des Döhlener Beckens bei Dresden. Veröffentlichungen des Naturhistorishen Museums Schleusingen 6:75–99.

Werneburg, R. 1993. *Onchiodon* (Eryopidae, Amphibia) aus dem Rotliegend des Innersudetischen Beckens (Böhmen). Paläontologische Zeitschrift 67:343–355.

Werneburg, R. 2009. The Permotriassic branchiosaurid T*ungussogyrinus* EFREMOV, 1939 (Temnospondyli, Dissorophoidea) from Siberia restudied. Fossil Record 12:105–120.

Werneburg, R. 2012. Dissorophoide Amphibien aus dem Westphalian D (Ober-Karbon) von Nýrany in Böhmen (Tschechische Republik)–der Schlüssel zum Verständnis der frühen 'Branchiosaurie.' Semana 27:3–50.

Werneburg, R. 2019. Dissorophoid amphibians from the Carboniferous-Permian boundary of France. Semana 34:11–51.

Werneburg, R., and D. S. Berman. 2012. Revision of the aquatic eryopid temnospondyl *Glaukerpeton avinoffi* Romer, 1952, from the Upper Pennsylvanian of North America. Annals of Carnegie Museum 81:33–60.

Werneburg, R., and J. Schneider. 1996. The Permian temnospondyl amphibians of India. Special Papers in Paleontology 52:105–128.

Werneburg, R., and J.-S. Steyer. 1999. Redescription of the holotype of *Actinodon frossardi* (Amphibia, Temnospondyli) from the Lower Permian of the Autun basin (France). Geobios 32:599–607.

Werneburg, R., F. Witzmann, J. W. Schneider, and R. Rößler. 2022. A new basal zatracheid temnospondyl from the early Permian Chemnitz Fossil Lagerstätte, central-east Germany. PalZ, 24 pp (ahead of print). DOI: 10.1007/s12542-022-00624-8

Werneburg, R., J. W. Schneider, S. Voigt, and A. Belahmira. 2019. First African record of micromelerpetid amphibians (Temnospondyli, Dissorophoidea). Journal of African Earth Sciences 159:103573.

Werneburg, R., J.-S. Steyer, G. Sommer, G. Gand, J. W. Schneider, and M. Vianey-Liaud. 2007. The earliest tupilakosaurid amphibian with diplospondylous vertebrae from the Late Permian of southern France. Journal of Vertebrate Paleontology 27:26–30.

Werneburg, R., S. G. Lucas, J. W. Schneider, and L. F. Rinehart. 2010. First Pennsylvanian *Eryops* (Temnospondyli) and its Permian record from New Mexico. New Mexico Museum of Natural History and Science Bulletin 49:129–135.

Werneburg, R., S. Štamberg, and J.-S. Steyer. 2020. A new stereospondylomorph, *Korkonterpeton kalnense* gen. et sp. nov., from lower Permian of the Czech Krkonoše Piedmont Basin and a redescription of *Intasuchus silvicola* from the lower Permian of Russia (Temnospondyli, Amphibia). Fossil Imprint 76:217–242.

Werneburg, R., Schneider, J.W., and S. G. Lucas. 2021. The new dvinosaurian *Bermanerpeton kinneyi* (Temnospondyli), with "branchiosaurid" characters, from the Late Pennsylvanian Kinney Brick Quarry in New Mexico. New Mexico Museum of Natural History and Science Bulletin 84:433–451.

Williston, S. W. 1909. New or little-known Permian vertebrates: *Trematops*, new genus. The Journal of Geology 17:636–658.

Williston, S. W. 1910. *Cacops*, *Desmospondylus*; new genera of Permian vertebrates. Bulletin of the Geological Society of America 21:249–284.

Williston, S. W. 1911. American Permian vertebrates. University of Chicago Press, Chicago, 130 pp.

Williston, S. W. 1913. The primitive structure of the mandible in Amphibians and Reptiles. The Journal of Geology 21:625–627.

Williston, S. W. 1914. *Broiliellus*, a new genus of amphibians from the Permian of Texas. *The* Journal of Geology 22:49–56.

Wilson, J. A. 1948. A small amphibian from the Triassic of Howard County, Texas. Journal of Paleontology 22:359–361.

Wiman, C. 1910. Ein paar Labyrinthodontenreste aus der Trias Spitzbergens. Bulletin of the Geological Institution of the University of Uppsala 9:33–41.

Wiman, C. 1913. Über das Hinterhaupt der Labyrinthodonten. Bulletin of the Geological Institution of the University of Upsala 12:1–8.

Wiman, C. 1914. Über die Stegocephalen aus der Trias Spitzbergens. Bulletin of the Geological Institution of the University of Uppsala 13:1–34.

Wiman, C. 1916. Neue Stegocephalenfunde aus dem Posidonomyaschiefer Spitzbergens. Bulletin of the Geological Institution of the University of Uppsala 13:13–30.

Wiman, C. 1917. Über die Stegocephalen *Tertrema* und *Lonchorhynchus*. Bulletin of the Geological Institution of the University of Uppsala 14:15–29.

Witzmann, F. 2005. Cranial morphology and ontogeny of the Permo-Carboniferous temnospondyl *Archegosaurus decheni* Goldfuss, 1847 from the Saar–Nahe Basin, Germany. Earth and Environmental Science Transactions of The Royal Society of Edinburgh 96:131–162.

Witzmann, F. 2007. The evolution of the scalation pattern in temnospondyl amphibians. Zoological Journal of the Linnean Society 150:815–834.

Witzmann, F. 2013. Phylogenetic patterns of character evolution in the hyobranchial apparatus of early tetrapods. Earth and Environmental Science Transactions of the Royal Society of Edinburgh 104:145–167.

Witzmann, F., and R. R. Schoch. 2005 (for 2006a). Skeletal development of the temnospondyl *Acanthostomatops vorax* from the Lower Permian Döhlen Basin of Saxony. Earth and Environmental Science Transactions of the Royal Society of Edinburgh 96:365–385.

Witzmann, F., and R. R. Schoch. 2006b. The postcranium of *Archegosaurus decheni*, and a phylogenetic analysis of temnospondyl postcrania. Palaeontology 49:1211–1235.

Witzmann, F., and R. Soler-Gijón. 2010. The bone histology of osteoderms in temnospondyl amphibians and in the chroniosuchian *Bystrowiella*. Acta Zoologica 91:96–114.

Witzmann, F., and T. Gassner. 2008. Metoposaurid and mastodonsaurid stereospondyls from the Triassic–Jurassic boundary of Portugal. Alcheringa 32:37–51.

Witzmann, F., B. M. Rothschild, O. Hampe, G. Sobral, Y. M. Gubin, and P. Asbach. 2014. Congenital malformations of the vertebral column in ancient amphibians. Anatomia, Histologia, Embryologia 43:90–102.

Witzmann, F., R. R.Schoch, A. Hilger, and N. Kardjilov. 2012. Braincase, palatoquadrate and ear region of the plagiosaurid *Gerrothorax pulcherrimus* from the Middle Triassic of Germany. Palaeontology 55:31–50.

Witzmann, F., S. Sachs, and C. J. Nyhuis. 2016. A new species of *Cyclotosaurus* (Stereospondyli, Capitosauria) from the Late Triassic of Bielefeld, NW Germany, and the intrarelationships of the genus. Fossil Record 19:83–100.

Wu, R., L. Tu, and F-L. Han. 2022. A Temnospondyl tooth from the Middle Triassic of the Ordos Basin, Shaanxi Province. Vertebrata PalAsiatica 60:54–58.

Xiong, J. L., Q. Chen, X. M. Zeng, E. M. Zhao, and L. Y. Qing. 2007. Karyotypic, morphological, and molecular evidence for *Hynobius yunanicus* as a synonym of *Pachyhynobius shangchengensis* (Urodela: Hynobiidae). Journal of Herpetology 41:664–671.

Yang, Z. 1979. [A late Permian fauna from Jiyuan, Henan]. Vertebrata PalAsiatica 17:99–113. [Chinese]

Yates, A. M. 1999. The Lapillopsidae: a new family of small temnospondyls from the Early Triassic of Australia. Journal of Vertebrate Paleontology 19:302–320.

Yates, A. M. 2000. A new tiny rhytidosteid (Temnospondyli: Stereospondyli) from the Early Triassic of Australia and the possibility of hidden temnospondyl diversity. Journal of Vertebrate Paleontology 20:484–489.

Yates, A. M., and A. Warren. 2000. The phylogeny of the 'higher' temnospondyls (Vertebrata: Choanata) and its implications for the monophyly and origins of the Stereospondyli. Zoological Journal of the Linnean Society 128:77–121.

Yates, A. M., and D. P. Sengupta. 2002. A lapillopsid temnospondyl from the Early Triassic of India. Alcheringa 26:201–208.

Young, C.-C. 1963. Note on the labyrinthodonts of the Sinokanemeyeria fauna in Shansi. Vertebrata PalAsiatica 7:331–341.

Young, C.-C. 1966. On the first capitosauroid from Sinkiang. *Vertebrate PalAsiatica*, 19: 41–46.

Young, C.-C. 1966. On the first discovery of capitosaurid from Sinkiang. Vertebrata PalAsiatica 10:58–63.

Young, C.-C. 1973. [A labyrinthodont from the Lystrosaurus Beds of Jimusuo'er County, Xinjiang]. Vertebrata PalAsiatica 11:50–51. [Chinese]

Young, C.-C. 1978. [A late Triassic vertebrate fauna from Fukang, Sinkiang]. Memoirs of the Institute of Vertebrate Paleontology and Paleoanthropology, Academia Sinica 13:14–147. [Chinese]

Zalesskiy, I. M. 1947. [First find of a branchiosaur in the Urals]. Priroda 1947:79–80. [Russian]

Zanon, R. T. 1991. *Negevodus ramonensis* Mazin, 1986, reinterpreted as a temnospondyl, not a placodont. Journal of Vertebrate Paleontology 11:515–518.

#### Appendix 6. Character list.

Numbers in parentheses represent the corresponding numbering from Schoch (2013), Pardo et al. (2017a), and Schoch et al. (2020). The following characters were omitted from the analysis: 14, 47, 48, 61, 78, 81, 100, 115, 253, 259, 334; these omitted characters are listed in full at the end of this section. All modified characters have an annotation indicating in brief what change was made; the reader is referred to the character-specific sections for more details.

1. (1) **Skull (outline)**. Parabolic (0), or triangular with pointed tip (1).

2. (2) **Skull (height)**. Skull of variable height, but occiput always deeper than snout (0), or extremely flattened to equal height throughout (1).

3. (3) **Ornament (elements)**. Reticulate ridges of various sizes (0), isolated pustules (1), dense covering of minute pits (2), or sparsely distributed pits or absent (3). [*unordered*]
    a. State 2 was modified for the condition of *Eocaecilia* and a new state 3 was added to further differentiate this from other lissamphibians with greatly reduced or absent ornamentation.

4. (4) **Ornament (snout)**. Polygons or short grooves (0), mostly radial, elongated grooves (1), or numerous circular to subcircular pits (2). [*unordered*]
    a. State 2 was added to capture the condition of gymnophionomorphs.

5. (5) **Ornament (general)**. Shallow ridges of variable height (0), or high ridges throughout (1).

6. (6) **Ornament (intensive growth)**. Elongated ridges (‘zones of intensive growth’) confined to snout only (0), or prepineal growth zone established on anterior parietal and postorbital (1).

7. (7) **Ornament (preorbital)**. Zone of subdued ornament on the medial skull bones adjacent to the midline suture absent (0), or present (1).

8. (8) **Spines**. Absent (0), or present along lateral flank of quadratojugal (1).

9. (9) **Preorbital region (length)**. Less than twice the length of posterior skull table (0), more than twice the length (1); twice as long (2). [*unordered*]
    a. State 2 was previously incorrectly defined as a pre- and postorbital region of equal length, which would be captured by state 0; state 2 was redefined to the only available option.

10. (10) **Premaxilla (posterior margin)**. Transversely straight (0), angled posterolaterally (1), forming a posterior hook-like indentation ‘(alary process’ sensu Ruta & Bolt (2006) (2), or forming a posterior plate-like indentation (3). [*unordered*]
    a. The character was redefined to differentiate among different conditions more clearly. It was also partially combined with character 350 of Schoch et al. (2020)

to capture a more uncommon morphology of the alary process in which it forms a broad, plate-like indentation in some taxa.

11. (11) **Premaxilla (prenarial portion)**. Short (0), or very expanded, equaling the length of the frontal (1).

12. (12) **Premaxilla (outline)**. Parabolically rounded (0), or box-like and anteriorly blunt (1).

13. (13) **Premaxilla (foramen)**. Premaxilla and nasal completely sutured (0), medially bearing an opening within the premaxillae (1), medially bearing an opening between the premaxillae and the nasals without prominent division of the nasals (2), or medially bearing an opening between the premaxillae and the nasals that prominently divides the nasals (3). [*unordered*]
    a. This character was combined with original character 14 (premaxillary fontanelle), the latter of which was then omitted.

14. (15) **Snout (internarial distance)**. Narrower than interorbital distance (0), or wider (1).

15. (16) **Nasal (width)**. Longer than wide (0), or as wide as long (1).

16. (17) **Naris (position)**. At lateral margin of snout, opening laterally (0), or set well medially, opening anteriorly (1).

17. (18) **Naris (extension)**. Naris uniform, with oval shaped margin (0), or posteriorly expanded with distinct anterior and posterior regions giving ‘key-hole’ shape (1).

18. (19) **Naris (flange)**. Ventral surface of skull roof medial to naris is smooth (0), or bearing a complicated bar-like structure (narial flange) (1).
    a. The character was modified to remove a mention of the lacrimal for state 0, allowing taxa without lacrimals to be coded.

19. (20) **Nasal (lateral margin)**. Straight, longitudinal (0), stepped, with lateral excursion (1), or straight, oblique (2). [*unordered*]
    a. The character was modified to remove a mention of the lacrimal for state 1, allowing taxa without lacrimals to be coded, and added a state 2 to differentiate the condition of some dvinosaurs and lissamphibians in which a straight suture is oriented mostly laterally, rather than anteriorly.

20. (21) **Lacrimal**. Present (0), or absent (1).

21. (22) **Lacrimal (lateral suture)**. Parallels medial suture (0), or posterolaterally expanded to give broader preorbital region (1).

22. (23) **Lacrimal (anterior extent)**. Extending anterior to orbit (0), or confined to lateral orbital margin (1).

23. (24) **Lacrimal (posterior extent)**. Restricted to the antorbital region (0), or extending back lateral to orbit (1).

24. (25) **Orbit and naris**. Well separated (0), or separated only by narrow gap (1).
    a. The character was modified to remove the lacrimal as a specifier in state 0.

25. (26) **Orbit location**. More medial than lateral, framed by wide jugals laterally (0), more lateral than medial, framed by slender jugals laterally (1), or central, framed by both narrow jugals and narrow frontals (2). [*unordered*]
    a. State 2 was not defined by Pardo et al. (2017a) but was inferred by comparison of taxa coded for this state and by personal communication with J. Pardo.

26. (27) **Orbit margins**. Raised well above skull plane (0), or flush with roof (1).

27. (28) **Palpebral ossifications**. Ossifications in orbit restricted to sclerotic ring (0), or numerous additional ossicles (1).

28. (29) **Pineal foramen (adults)**. Present (0), or absent (1).

29. (30) **Interorbital distance**. Narrower than or equaling orbital width (0), or wider (1).

30. (31) **Frontal, nasal (length)**. Frontal as long or longer than nasal (0), or shorter (1).

31. (32) **Frontal-nasal (suture)**. Level with or anterior to the orbital margin (0), or posterior (1).

32. (33) **Lateral line sulci (adults)**. Present (0), absent (1).

33. (34) **Sulci (skull roof)**. Throughout skull roof if present (0), or confined to circumorbital region (1).

34. (35) **Infraorbital sulcus**. With simple curve (0), or with pronounced S-shaped flexure (1).
    a. Modified to remove the lacrimal as a specifier in state 0.

35. (36) **Supraorbital sulcus**. Passing entirely through nasal (0), passing through nasal and prefrontal (1), or passing through nasal, prefrontal, and lacrimal (2). [*unordered*]
    a. State 2 was added to further differentiate taxa.

36. (37) **Maxilla (anterior margin)**. Straight (0), or laterally convex due to enlarged teeth (1).

37. (38) **Maxilla-nasal**. Separated (0), or in contact (1).

38. (39) **Prefrontal, frontal (anterior extent)**. Prefrontal extends beyond the anterior level of the frontal (0), ending at one level with frontal (1), or ending posterior to the anterior level of the frontal (2). [*ordered*]

a. State 2 was added to capture the rare condition in which the prefrontals terminate posterior to the anterior level of the frontals.

39. (40) **Prefrontal (anterior end).** Pointed (0), or wide and blunt (1).

40. (41) **Prefrontal (lateral suture with jugal)**. Suture posterolaterally oriented, either straight or continuously anterolaterally convex (0), longitudinally straight or continuously laterally convex (1), anteromedially oriented, either straight or continuously posterolaterally convex (2), stepped with distinct inflection point near orbit (3). [*unordered*]
    a. This character was completely revised to more clearly define various conditions, to restrict it to the prefrontal-jugal suture, and in an attempt to capture at least the most distinctive condition of some rhinesuchids (*Uranocentrodon*).

41. (42) **Prefrontal-postfrontal**. In contact (0), or separated (1).

42. (43) **Prefrontal (process)**. Prefrontal-lacrimal suture simple (0), or prefrontal underlying lacrimal laterally to reach the palatine (1).

43. (44) **Prefrontal-jugal**. Separated (0), or in contact (1).
    a. Modified to remove the lacrimal as a specifier in state 0.

44. (45) **Prefrontal-maxilla**. Separated (0), or in contact (1).
    a. Modified to remove the lacrimal as a specifier in state 0.

45. (46) **Posterior skull table (length)**. More than 0.6 times the width (0), or less than that (1).

46. (231) **Postorbital (posterior margin)**. Posterior margin approximately straight or squared-off (0), or margin with prominent triangular terminus (1).

47. (49) **Postorbital (lateral process)**. Not wider than orbit (0), or with substantial lateral process projecting into jugal (1).

48. (50) **Postorbital, postfrontal (length)**. Shorter than supratemporal and parietal (0), or as long or longer (1).

49. (51) **Otic notch**. Semicircular embayment between squamosal and posterior skull table (0), or straight transverse posterior skull margin without embayment between cheek and table (1).

50. (52) **Otic notch (position)**. Lateral, expanding along entire cheek to form continuous unornamented area up to quadrate (0), or slit-like (1), or small and rounded, confined to dorsomedial part of squamosal (2). [*unordered*]

51. (53) **Supratemporal (proportions)**. Longer than wide (0), or quadrangular (1).

52. (54) **Supratemporal (lateral margin)**. With straight sagittal lateral margin (0), or

posterolaterally constricted (1).

   a. A state 2 added by Pardo et al. (2017a; 'small and oval in shape') was a problematic compound character that overlapped with character 53 (specifically state 0) and that was ill-suited to be placed with this character. It was omitted. The reference to 'by an expanded otic notch' for state 1 was also removed to allow taxa without otic notches to be coded.

53. (55) **Supratympanic flange**. Squamosal continuously ornamented around margin of otic notch (0), or squamosal having dorsally exposed and ornamented area (supratympanic flange) stepping abruptly into steeply aligned, poorly ornamented portion (1).

54. (56) **Semilunar flange**. Supratemporal without ventral projection into otic notch (0), or supratemporal forming marked ventral flange participating in medial bordering of otic notch (1).

55. (57) **Jugal (ventral process)**. No ventral outgrowth (0), or insula jugalis framing subtemporal window (1).

56. (58) **Jugal (anterior extent)**. Ending at or posterior to level of anterior orbit margin (0), or extending anteriorly (1).

57. (59) **Jugal-lacrimal**. In contact (0), or separated (1).

58. (60) **Intertemporal**. Present (0), or absent (1).

59. (232) **Parietal-postorbital**. Separated (0), or in contact (1).

60. (62) **Squamosal-tabular (dorsal)**. Separated (0), or in contact (1).

61. (63) **Tabular, squamosal**. Forming either squamosal embayment or straight posterior margin (0), or projecting posteriorly, with tabular extended posterolaterally (1).

62. (64) **Squamosal (falciform crest)**. Posterior rim of squamosal straight (0), or with convex projection (falciform crest) (1).

63. (65) **Tabular horn**. Present in some form (0), or entirely absent (1).

64. (66) **Tabular horn (extension)**. Pointing posteriorly if present (0), or aligned laterally (1), or sutured with squamosal posterior to otic notch (cyclotosaur condition) (2). [*unordered*]

65. (67) **Tabular (ventral crest)**. Absent (0), or present and shallow (1), or forming a deep ridge that almost doubles the thickness of the tabular in occipital view (2). [*ordered*]

66. (68) **Quadratojugal (medial process)**. Absent (0), or present (1).

67. (69) **Quadratojugal-maxilla**. In contact (0), or separated (1).

68. (70) **Quadrate (dorsal process)**. Absent (0), or present (1).

69. (71) **Posterior skull rim**. Quadrate trochlea posterior to posteriormost extent of tabular (0), or at one level or anterior (1).
    a. Modified to change the reference point from the tabular horn (not found in all taxa) to the posteriormost extent of the tabular (the tabular horn in taxa with this projection and codeable in taxa without this projection).

70. (72) **Occipital flange**. Descending flange of occipital portion of postparietals forming a bulge (0), or long smooth blades as long as the dermal portion of the postparietal (1).

71. (73) **Postparietal-exoccipital**. Both elements ossified but separated (0), both elements ossified and in contact (1), or exoccipitals unossified (2). [*unordered*]
    a. State 2 was added here as I felt that this was the most appropriate character to add a state for the absence of ossified exoccipitals, a phylogenetically informative feature.

72. (74) **Posttemporal fenestra**. Large opening, much wider than the foramen magnum when the maximum width of each is compared (0), moderately sized, subequal in width to the foramen magnum (1), or reduced to a tiny foramen (2). [*ordered*]
    a. The previous two states did not capture a condition in which the openings were of subequal width; this was added as the new state 1 to create an ordered character. The specifier of ‘maximum width’ is also added to clarify how these openings are being compared.

73. (75) **Quadrate and occipital condyles**. Quadrate condyles posterior to occipital ones (0), or at same level (1), or well anterior (2). [*ordered*]

74. (76) **Epipterygoid**. Simple rod-like ascending process (0), or complicated and robust element with up to six processes (1).

75. (77) **Dentition (marginal, size)**. Heterogenous, varying sizes and distances (0), or homogeneous, small teeth, equidistant (1).

76. (79) **Dentition (marginal, tooth bases)**. Round or oval (0), or forming transversely broadened ovals (1).

77. (80) **Palatal tusks (cross-section)**. Round or oval (0), or laterally compressed and keeled at least on one side (1).

78. (82) **Dentition (marginal, pedicely)**. All teeth formed consisting of a single mineralized conus (0), or at least some teeth with two separate mineralization centers (1).

79. (83) **Dentition (marginal, bicuspidity)**. All teeth with single tip (0), or at least some with bicuspid crowns (1).

80. (84) **Dentition (marginal, labyrinthodonty)**. Teeth with labyrinth infolding of dentine and enamel at base (0), or never labyrinthodont (1).

81. (85) **Transverse tooth row (transvomerine)**. Absent (0), present and transverse in single row (1), V-shaped in single row (2), transverse patch of teeth (3). [*unordered*]

82. (86) **Additional vomerine fangs**. Vomer with a single pair of fangs at the medial margin of choana (0), or with additional fangs/fang pairs posteromedially (1).

83. (87) **Parasphenoid (shagreen)**. Denticle patches restricted to basal plate (0), denticles entirely absent (1); denticles expanded onto cultriform process to at least the mid-length of the process (2). [*unordered*]
    a. This character was retained over the redundant character 253 added by Pardo et al. (2017a) and was combined with characters 115 and 352 to capture the new state 2 in which the denticles extend onto the cultriform process. 'Absent' is retained as state 1 because the three characters do not form a clear morphocline.

84. (88) **Ectopterygoid (fangs)**. Present (0), or absent (1).

85. (89) **Interpterygoid vacuities (shape)**. Slender and slit-like (0), or rounded, oval (1).

86. (90) **Interpterygoid vacuities, pterygoid**. Vacuities bordered by moderately concave pterygoid (0), or laterally greatly extended at mid-level pushing pterygoid to the margin (1).

87. (91) **Anterior palatal opening(s)**. Vomer and premaxilla with continuous suture (0), or perforated to accommodate symphyseal fangs (1).

88. (92) **Anterior palatal opening(s) (paired)**. Unpaired if present (0), or paired (1).

89. (93) **Vomer (shape)**. Narrow and small (0), or large plate, widely separating choanae (1).

90. (94) **Vomer (paired anterior depressions)**. Absent (0), or present (1).

91. (95) **Anterior palatal depression**. Posterior rim round if present (0), or straight transverse (1).

92. (96) **Vomer (ridges)**. Absent (0), or present, radiating from vomerine tusks anteriorly (1).

93. (97) **Vomer (pit and fontanelle)**. Absent (0), or pit present (posterior to mandible) (1), or fenestra within such pit (2). [*unordered*]

94. (98) **Vomer (septum)**. Absent (0), or present (1).

95. (99) **Choana (orientation)**. Long axis oriented more transversely than it is sagittally (0), or long axis more sagittally oriented (1).
    a. This character was completely replaced by a new character to capture the orientation of the long axis of the choana.

96. (101) **Choana (shape)**. Oval and symmetrical (0), semi-circular and symmetrical (1), asymmetrical, expanded orally (2), or asymmetrical, expanded aborally (3). [*unordered*]
    a. This character was combined with original characters 99 and 100 to differentiate shape of the choana within a single character.

97. (102) **Vomer-pterygoid**. In contact (0), or separated (1).

98. (103) **Vomer (posteromedial process)**. No process extending posterior to the level of the anterior margin of the interpterygoid vacuities (0), process present and short (1), or process present and long (2). [*ordered*]

99. (104) **Vomer-palatine (suture)**. Anteriormost contact with the palatine is anterior to the level of the posterior choanal margin (0), or contact is at the same level or more posterior (1).
    a. This character was completely revised to more clearly differentiate a condition similar to, if not equivalent with, the original character.

100. (105) **Basipterygoid ramus**. Pterygoid with short, posteromedially curved basipterygoid process, constricting palatal vacuities posterolaterally (0), or with transversely extended process producing posteriorly wider vacuities (1).

101. (106) **Basicranium (contact)**. Joint between basal plate and pterygoid (0), or sutural contact (1).

102. (107) **Basicranium (suture)**. Suture (if present) much shorter than basal plate, reaching at best 40% its length (0), suture between 40% and 75% the length (1), or suture almost as long as basal plate (2). [*ordered*]
    a. The original two states did not capture the intermediate condition in which the suture was neither very short nor very long; this is added in the new state 1, with the original state 1 now becoming state 2 in order to create an ordered character.

103. (108) **Parasphenoid-exoccipital**. Parasphenoid suturing with exoccipitals (0), or underplating exoccipitals (1).

104. (109) **Basicranium (carotids)**. Internal carotids entered basicranium ventrally near base of cultriform process (0), or at posterolateral corner of bone (1).

105. (110) **Parasphenoid (plate, proportions)**. Basal plate sagittally rectangular (0), or quadrangular to slightly wider than long (1), or much abbreviated, transversely rectangular (2). [*ordered*]

106. (111) **Parasphenoid (plate, width)**. Basal plate substantially narrower than parietals at level posterior to basicranial joint (0), or as wide or wider (1).

107. (112) **Parasphenoid (cultriform process, width)**. Base not wider than rest, clearly set off from basal plate (0), or merging continuously into plate (1). [*unordered*]

108. (113) **Parasphenoid (cultriform process, structure)**. Ventrally flat (0), or with ridge emplaced on broader base (1), or knife-edged and keel-shaped (2). [*unordered*]

109. (114) **Parasphenoid (cultriform process, outline)**. Of similar width throughout (0), or posteriorly expanding abruptly to about twice the width (1).

110. (116) **Parasphenoid (plate, lateral notches)**. Posterolateral margin continuous (0), or with notches in the lateral margin of the basal plate (1).
    a. Because of the ambiguity related to the so-called 'lateral wings' of some taxa, this character was modified to refer to a correlated feature, incisions in the lateral margin of the basal plate, that is more clearly identifiable without confusion over terminological consistency. State 0 is also modified from 'straight' to 'continuous' to better differentiate states, as a margin could be curved and therefore neither 'straight' nor 'with notches.'

111. (117) **Pterygoid (ventral ornament)**. Palatine ramus of pterygoid smooth (0), or ornamented with reticulate ridges (1).

112. (118) **Pterygoid-exoccipital**. Separated (0), or sutured lateral to parasphenoid (1).

113. (119) **Pterygoid-basioccipital**. Separated (0), or sutured lateral to parasphenoid (1).

114. (120) **Pterygoid-squamosal**. Entirely sutured (0), or with open fissure (1).

115. (121) **Pterygoid (flange)**. Palatine ramus of pterygoid merging continuously into basipterygoid ramus (0), or broadening abruptly to form transverse flange (1).

116. (122) **Pterygoid (quadrate ramus, width)**. Ramus distinct from palatine ramus and narrower at base (0), distinct and broader at base (1), or confluent with palatine ramus (2). [*unordered*]
    a. This character was restricted to the quadrate ramus, and the width measured at the base is specified as the means of assessing relative width with the palatine ramus. As some taxa have an essentially indistinct base of these processes (e.g., *Dvinosaurus*), a new state 2 is added for the condition in which these processes merge together to be relatively indistinct except distally.

117. (123) **Palatine ramus**. Formed exclusively by pterygoid (0), or with posteromedial projection of ectopterygoid.

118. (124) **Pterygoid-palatine-ectopterygoid**. Pterygoid contacting both ectopterygoid and palatine (0), or pterygoid only in contact with ectopterygoid (1).

119. (125) **Palatine, ectopterygoid (suture)**. With straight suture (0), or palatine with posterior process along medial edge of ectopterygoid (1).

120. (126) **Palatine, vomer (suture)**. Suture aligned posterolaterally (0), or with medial wing framing the interpterygoid vacuity anteriorly (1).

121. (127) **Palatine (laterally exposed palatine, LEP)**. Absent (0), or present (1).

122. (128) **Palatine, ectopterygoid (ontogeny)**. Maintain their width (0), or become proportionally wider (1).

123. (129) **Palatine, ectopterygoid (width)**. Much wider than maxilla (0), or as narrow (1).

124. (130) **Palatine, ectopterygoid (continuous tooth row)**. Absent (0), or present (1).

125. (131) **Ectopterygoid (length)**. As long or longer than palatine (0), or markedly shorter (1).

126. (132) **Ectopterygoid (laterally exposed ectopterygoid, LEE)**. Absent (0), or present (1).

127. (133) **Ectopterygoid (Y-shaped)**. Ectopterygoid with continuous maxillary suture (0), or Y-shaped, with posterior half separated from maxilla by a gap (1).

128. (134) **Palate structure**. In occipital view, pterygoids either sloping continuously ventrolaterally or flat horizontal (0), or vertically curved ventrally at right angle with basicranium (1).

129. (135) **Quadrate trochlea**. Medial bulge only slightly larger than lateral one (0), or at least two times longer and twice as wide (1).

130. (136) **Occipital condyle**. Trilobed (0), or bilobed (1).
    a. Modified to remove the basioccipital as a specifier in both states.

131. (137) **Exoccipital condyles**. Exoccipitals not exposed in dorsal view (0), exoccipitals partially exposed, with occipital condyles visible in dorsal view (1), or almost the complete element visible in dorsal view (2). [*ordered*]
    a. This character treated taxa with no dorsal exposure the same as those with only the occipital condyles exposed, so a new state 0 was added to the 'front' of the character for the former taxa.

132. (138) **Basioccipital (length)**. Forming a long element posterior to parasphenoid plate (0), foreshortened to a narrow posterior rim (1), or unossified (2). [*ordered*]

a. State 2 was added for ‘basioccipital unossified,’ as in many paedomorphic taxa; I felt that this was the best character to append this state, a phylogenetically relevant feature, as it allows for an ordered character to be reasonably constructed.

133. (139) **Dentigerous palatal ossicles**. Absent (0), or present within interpterygoid vacuities (1).

134. (140) **Postglenoid area**. Absent or present as very faint outgrowth (0), or longer than glenoid facet (1).

135. (141) **Postglenoid area (type)**. Type 1 (0), or type 2 (1).

136. (142) **Postglenoid area (dorsal)**. Plain (0), or with elongated groove (1).

137. (143) **Hamate process**. Absent (0), or present but lower than postglenoid portion is long (1), or as high as postglenoid process is long (2). [*ordered*]

138. (144) **Preglenoid process**. Labial side of surangular with straight dorsal margin anterior to glenoid (0), or forming convex dorsal projection well above the level of the glenoid articulation (1).

139. (145) **Meckelian window**. Small round or oval opening (0), or elongate window shorter than the adductor fossa (1), or as long or longer than adductor fossa (2). [*ordered*]

140. (146) **Symphyseal teeth**. No accessory teeth posterior to symphyseal tusks (0), or a transverse row of such teeth (1), or a row aligned parallel to the labial tooth row (2). [*unordered*]
    a. A new state 2 was added to characterize taxa with a symphyseal row that is aligned parallel to the marginal tooth row.

141. (147) **Coronoids (denticles):** Present on all coronoids (0), present on posteriormost coronoid but not anteriormost coronoid (1), or absent on all coronoids (2). [*unordered*]
    a. This character was modified in tandem with characters 148 and 272 to capture the three types of dentition that may occur on the coronoids (denticles, teeth of subequal size to the marginal teeth, and tusks/fangs).
    b. In the reanalysis of Schoch (2013), the original character is restored and then modified:
    **Posterior coronoid dentition**. Denticles only (0), teeth only (1), denticles and teeth (2), or edentulous (3). [*unordered*]

142. (148) **Coronoids (fangs):** Absent (0), or present (1).
    a. This character was modified in tandem with characters 147 and 272 to capture the three types of dentition that may occur on the coronoids (denticles, teeth of subequal size to the marginal teeth, and tusks/fangs).
    b. In the reanalysis of Schoch (2013), the original character is restored and restricted to only to the middle coronoid in order to avoid a compound character.

**Middle coronoid dentition**. Denticles only (0), teeth only (1), denticles and teeth (2), or edentulous (3). [*unordered*]

143. (149) **Mandibular osteoderms**. Throat region naked (0), or covered with a mosaic of ventral osteoderms between mandible and dermal pectoral girdle (1).

144. (150) **Stapes (quadrate process)**. Absent (0), or present (1).

145. (151) **Stapes (ventral process)**. Absent (0), or present, giving the proximal region two heads (1).

146. (152) **Stapes (shape)**. Robust, tetrahedral bone (0), or rod-like element with elongated stylus (1), or blade-like (2). [*unordered*]
    a. The specifier of the quadrate process was removed from state 0.

147. (153) **Stapes (curvature)**. Stapes with pronounced dorsodistal curvature (0), or without such curvature (1).
    a. This character was restricted to curvature, with references to length omitted.

148. (154) **Ceratobranchials**. Bony elements absent (0), present in adults (1).

149. (155) **Basibranchial**. Bony element absent (0), present in adults (1).

150. (156) **Hypobranchials**. Bony elements absent (0), present in adults (1).

151. (157) **Branchial denticles**. Conical and attached to small ossicles in groups of 2–10 (0), or free and with brush-like end (1).

152. (158) **Presacral count**. More than 28 vertebrae (0), 23–28 (1), 19–22 (2), or less than 19 (3). [*ordered*]
    a. Schoch et al. (2020) added an unspecified state 3 that was inferred to be 'less than 19 positions' based on the taxa for which it was coded. Regardless of this addition, the previous states did not encompass all possible presacral counts (specifically 22, 26, 27, and 28), so states 1 and 2 were redefined to encompass these.

153. (159) **Caudal count**. Similar to presacral count or higher (0), or much lower (1).

154. (160) **Transverse process (orientation)**. Short, directed posteriorly (0), or distally extended with diapophysis pointing laterally (1).

155. (161) **Transverse process (length)**. Shorter than dorsal spine is high (0), or markedly longer (1).

156. (162) **Neural spine (height)**. As high as distance between zygapophyses (0), or higher (1).

157. (163) **Intercentrum (shape)**. Presacral intercentra form wedges (0); or dorsally closed discs (1); or dorsally closed and elongated cylinders (2). [*ordered*]

158. (164) **Intercentrum (width)**. Chordal canal wider than intercentrum is high (0), narrower (1).

159. (165) **Intercentrum (ventral surface)**. Ventral surface shorter than wide in ventral view, giving transversely rectangular outline (0), or as long as wide, quadrangular (1).

160. (166) **Intercentrum (anterior surface)**. Always concave (0), or convex at least in some presacral centra (1).

161. (167) **Parapophysis**. Segmental (0), or intersegmental (1).

162. (168) **Pleurocentrum.** Ossified (0), unossified (1).

163. (169) **Pleurocentrum (lateral surface)**. As large as that of intercentrum (0), or smaller (1).

164. (170) **Pleurocentrum (ventral extension)**. Wedged between successive intercentra and not reaching ventral margin of intercentra (0), pleurocentra ventrally expanded to near each other (1), or ventrally fused to form a single cylindrical element (2). [*ordered*]

165. (171) **Ribs (curvature)**. Curved distoventrally (0), or without distal curvature (1).
    a. This character was restricted to curvature, with references to length omitted.

166. (172) **Ribs (ventral extension)**. Rib heads (tuberculum and capitulum) confluent (0), or clearly set off and widely separated in mid-trunk region (1).

167. (173) **Ribs (uncinate blades)**. Small and spine-like (0), extensive and blade-like (1), or uncinate processes absent (2). [*unordered*]
    a. A new state 2 was added for taxa without any sort of uncinate process or projection; it is appended to the 'end' of the original character as there is no *a priori* reason to order the three states.

168. (174) **Ribs (uncinate spines)**. Short (0), or longest uncinate spine elongated, as long as shaft (1).

169. (175) **Cleithrum (dorsal head)**. With broadened dorsal head region (0), or a simple rod (1).

170. (176) **Cleithrum (dorsal head, anterior rim)**. Head with slightly convex or straight anterior rim (0), or with pronounced anterior projection (1).

171. (177) **Cleithrum (dorsal head, dorsal extent)**. Dorsal head region confined to anterior rim of scapula (0), or posteriorly extended to cover dorsal rim of scapula (1).

172. (178) **Clavicle (ventral blade)**. Wide, triangular, overlapping interclavicle broadly (0), or slender, with minor overlap (0).

173. (179) **Interclavicle (length)**. Shorter than posterior skull (0), or substantially longer (1).

174. (180) **Interclavicle (ontogeny)**. Without major proportional change in ontogeny (0), or decreasing proportionally relative to skull length (1).

175. (181) **Interclavicle (central ornamented area)**. Rhomboidal (0), or pentagonal and posteriorly widest (1).

176. (182) **Interclavicle (proportions)**. As long as wide (0), 1.3 times as long as wide (1), or more than twice as long as wide (2). [*ordered*]

177. (183) **Interclavicle (posterior margin)**. With posterior process (0), or transversely straight (1).

178. (184) **Interclavicle (anterior margin)**. Serrated (0), or smooth (1).

179. (185) **Interclavicle (anterior stylus)**. Anterior portion of interclavicle variably shaped but not longer than posterior one, as measured by the center of ornamentation (0), or substantially longer than posterior one (1).

180. (186) **Interclavicle (posterior stylus)**. Posterior end rounded or blunt (0), or with elongated stylus or parasternal process (1).

181. (187) **Interclavicle, clavicles**. Clavicles broadly separated by interclavicle ventrally (0), or leaving only narrow strip of interclavicle in between (1), or in contact and excluding anterior part of interclavicle from ventral exposure (2). [*ordered*]

182. (188) **Scapula (glenoid facet)**. Ossified (0), or unossified in adults (1).

183. (189) **Scapula (height)**. Maximally two times longer (higher) than wide (0), or dorsally extended and fully ossified, three times higher than wide (1).

184. (190) **Humerus (ontogeny)**. Short cylinder in larvae, elongates at slow rate (0), with substantial shaft in larvae (1), or very long rod from small stages on (2). [*unordered*]

185. (191) **Humerus (adult shaft)**. No shaft proper, humerus blade-like in cross-section (0), with short shaft oval in cross-section (1), or shaft elongate and slender, comprising more than 50% of the humerus with rounded cross-section (2). [*ordered*]

186. (192) **Humerus (entepicondylar foramen)**. Present (0), or absent (1).

187. (193) **Humerus (supinator)**. Present (0), or absent (1).

188. (194) **Humerus (condyles)**. Distal end at least partially ossified with identifiable condyles (0), unfinished without condyles (1), or with fully established condyles (2). [*unordered*]

189. (195) **Humerus (torsion)**. Strong, 70-90° (0), or weak, well below 60° (1).

190. (196) **Carpals**. All unossified (0), or at least some ossified (1).

191. (197) **Manual digit count**. Five (0), or four (1).

192. (198) **Ilium (shaft)**. Shaft of variable length but laterally flattened (0), or very long and slender (1).

193. (199) **Ilium (dorsal end)**. Tip of dorsal end continuous (0), or much broadened (1).

194. (200) **Ilium (height)**. Shaft more than twice the length of the base (0), or shorter (1).

195. (201) **Ilium (orientation)**. Main axis of shaft inclined posterodorsally (0), vertically (1), or anteriorly (2). [*unordered*]

196. (202) **Ilium (tip)**. Dorsal end two-headed (0), or single-headed (1).

197. (203) **Pubis**. Unossified (0), or ossified (1).

198. (204) **Femur (intercondylar fossa)**. Intercondylar fossa forming deep and elongated trough (0), or reduced to short depression (1).

199. (205) **Femur (trochanter)**. Internal trochanter present as discrete process (0), reduced to a shallow crest (1), or greatly enlarged to form a massive projection (2). [*unordered*]

200. (206) **Tarsals**. At least some ossified (0), or all unossified (1).

201. (207) **Gastral squamation**. Ossified ventral dermal scales (0), or ventral dermal scales absent (1).

202. (208) **Squamation**. Dermal scales oval to spindle-shaped (0), or broad, oval with ‘microsaur-type’ ornament (1).

203. (209) **Osteoderms**. Absent (0), or present as single sagittal row (1), or more numerous to form a carapace (2). [*unordered*]

204. (210) **Osteoderms (articulations).** Simple set of osteoderms arranged in one layer if present (0), or double set of layers (1).

205. (211) **Osteoderms (width)**. An intra-dissorophid character: Narrow median osteoderms (0), or transversely extended plates (1).

(212) **Rib cage**. Trunk narrower than or as wide as skull (0), or trunk substantially wider than lateral margin of cheeks at about mid-level (1).

206. (213) **Skull (length)**. >450 mm (0) 251-450 mm (1), 151-250 mm (2), 101-150 mm (3), 71-100 mm (4), 51-70 mm (5), 30-50 mm (6), <30 mm (7). [*ordered*]
    a. This character was expanded to capture larger body sizes (previously all taxa with skull lengths greater than 70 mm were coded for the same state); this reflects the large body size of many stereospondyls. The ranges were also modified to avoid overlap at a specific measurement (e.g., original states of '30-50' and '50-70' would overlap at 50 mm).

207. (214) **Skull to trunk**. >=0.45 (0), 0.30-0.44 (1), 0.20-0.29 (2), < 0.20 (3). [*ordered*]

208. (215) **Skull (proportions)**. Longer than wide (0), wider than long (1).

209. (216) **Supratemporal**. Present (0), absent (1).

210. (217) **Supratemporal (occipital exposure)**. Absent (0), present (1).

211. (218) **Postfrontal (shape)**. Broadly quadrangular (0), falciform (1).

212. (219) **Lacrimal-naris**. In contact (0), separated (1).

213. (220) **Lacrimal-orbit**. Separated (0), in contact (1).

214. (221) **Lacrimal (orbital processes)**. Only ventral present (0), dorsal and ventral present (1), neither present (2). [*unordered*]

215. (222) **Quadratojugal**. Present (0), absent (1).

216. (223) **Frontals (paired)**. Paired along midline (0), fused (1).

217. (224) **Frontal-orbit**. Separated (0), in contact (1).

218. (225) **Septomaxilla**. Ossified and entirely intranarial (0), unossified (1), or ossified and integrated with skull roof (2). [*unordered*]
    a. A new state was added to differentiate taxa with intranarial septomaxilla and taxa with septomaxilla partially integrated with the skull roof. The two states expressing an ossified septomaxilla do not clearly form a morphocline with the state for an unossified septomaxilla, which is why the new state 2 is appended onto the end rather than placed immediately before or after state 0.

219. (226) **Prefrontal-naris**. Distant from naris (0), near (1), in contact (2). [*ordered*]

220. (227) **External naris in dorsal view**. Exposed (0), not exposed (1).

221. (228) **Posterior skull marg**in. Concave (0), straight (1), convex (2), undulating (3). [*unordered*]

222. (229) **Large otic notch approaching orbit**. Absent (0), intermediate (1), close (2). [*ordered*]

223. (230) **Postorbital**. Present (0), absent (1).

224. (233) **Parietal-squamosal**. Separated (0), in contact (1).

225. (234) **Parietal-tabular**. Separated (0), in contact (1).

226. (235) **Postparietals**. Paired (0), fused (1), absent (2). [*unordered*]

227. (236) **Postparietal-squamosal**. Separated (0), in contact (1).

228. (237) **Postparietal (proportions)**. Large, quadrangular (0), abbreviated anteroposteriorly, Transversely elongate rectangle (1).

229. (238) **Squamosal-jugal**. In contact (0), separated (1).

230. (239) **Tabular**. Present (0), absent (1).

231. (240) **Maxilla-external naris**. In contact (0), separated (1).

232. (241) **Maxilla (contribution to ventral naris)**. Absent (0), present (1).

233. (242) **Dentition (premaxilla, tooth count)**. >=10 (0), 5-9 (1), <5 (2). [*ordered*]

234. (243) **Dentition (maxilla, tooth count)**. >=30 (0), 20-29 (1), <20 (2). [*ordered*]

235. (244) **Dentition (marginal teeth, lateral compression)**. Absent (0), present (1).

236. (245) **Skull (occipital profile)**. Low and wide (0), high and wide (1), high and narrow (2). [*unordered*]

237. (246) **Supraoccipital**. Absent (0), present (1).

238. (247) **Jugular foramen**. Between opisthotic and exoccipital (0), through exoccipital (1).

239. (248) **Palatine (anterior process)**. Process along aboral edge of choana longer than process along oral edge (0), processes of equal length (1), process along oral edge longer (2), or palatine absent (3). [*unordered*]
    a. This character was completely redefined to remove ambiguity with the differentiation of states. Based on this, there are three states comparing the relative length of the two processes of the palatine that can frame the choana

orally ('inner' edge) and aborally ('outer' edge); these terms are preferred to 'medial' and 'lateral' following the same rationale as modifications to characters 99 and 101. Compared to character 87 where a new form of 'present' was merely appended to the end of the character, all of the 'present' states are grouped together, as some workers may feel that they can be ordered with the 'absent' state (or if the 'absent' state is removed).

240. (249) **Dentition (pterygoid, denticles)**. Present (0), absent (1).

241. (250) **Dentition (pterygoid, teeth)**. Absent (0), present (1).

242. (251) **Dentition (vomer, denticles)**. Present (0), absent (1).

243. (252) **Dentition (palatine, denticles)**. Present (0), absent (1).

244. (254) **Cultriform process (anterior extent)**. Extending anterior to the anterior level of the interpterygoid vacuities (0), terminating at such level (1), or terminating posterior to such level (2). [*ordered*]
    a. This character became partially redundant with the modified character 103, so this one was modified to refer to the anterior extent of the cultriform process.

245. (255) **Stapes**. Perforated columella (0), imperforate columella (1), no columella (2). [*unordered*]

246. (256) **Pleurosphenoid**. Unossified (0), ossified (1).

247. (257) **Sphenethmoid**. Ossified (0), unossified (1).

248. (258) **Pterygoids (contact)**. In contact (0), separated (1).

249. (260) **Dentary (length)**. Long (0), short (1).

250. (261) **Dentary (coronoid process)**. Absent (0), present (1).

251. (262) **Surangular**. Normal (0), reduced (1), absent (2). [*ordered*]

252. (263) **Angular (depth)**. Narrow (0), deep (1).

253. (264) **Splenials**. Two (0), one (1), zero (2). [*ordered*]

254. (265) **Splenial (lateral exposure)**. Present (0), absent (1).

255. (266) **Meckelian fossae**. Two or more foramina found along the ventrolingual surface (0), one (1), zero (2). [*ordered*]

a. It is clarified that a 'Meckelian fossa' is any opening along the ventrolingual surface of the lower jaw (based on how taxa were originally coded) and therefore excludes any posterodorsal foramina.

256. (267) **Posterior Meckelian window**. (0) Bounded by the prearticular, postsplenial, and angular; (1) bounded by the prearticular and postsplenial; (2) bounded by the prearticular and angular; (3) bounded by the prearticular, postsplenial, middle coronoid, and angular; (4) entirely enclosed by the postsplenial. Adapted from (Warren and Marsicano 2000; McHugh, 2012). [*unordered*]
    a. This character was completely replaced and redefined to explicitly refer to what is typically the largest of the Meckelian fossae ventral to the adductor fossa.

257. (268) **Retroarticular process**. Straight (0), hooked (1).

258. (269) **Jaw articulation, tooth row**. Jaw articulation above tooth row (0), at same level (1), below (2). [*ordered*]

259. (270) **Angular (extent in lateral view)**. Posterior tooth row (0), middle of tooth row (1).

260. (271) **Coronoids**. Three (0), two (1), one (2), zero (3). [*ordered*]

261. (272) **Coronoid (teeth)**. Present on all coronoids (0), present only on posteriormost coronoid (1), or absent on all coronoids (2). [*unordered*]
    a. This character was modified to further differentiate temnospondyls with three coronoids and teeth only on the posterior one from those with three coronoids and teeth on all three.

262. (273) **Coronoid (teeth, size)**. Larger than marginal (0), equal to marginal (1), smaller than marginal (2). [*ordered*]

263. (274) **Symphysis**. Dentary and splenial (0), dentary alone (1).

264. (275) **Jaw sculpture**. Present (0), absent (1).

265. (276) **Ossified hyoids**. Present (0), absent (1).

266. (277) **Parahyoid**. Absent (0), present (1).

267. (278) **Trunk intercentra**. Present (0), absent (1).

268. (279) **Trunk neural arch to centrum**. Loosely articulated (0), sutured (1), fused (2). [*ordered*]

269. (280) **Base of neural spine**. Equal to or wider than haemal (0), smaller than haemal (1).

270. (281) **Haemal arches**. Present (0), absent (1).

271. (282) **Haemal arch (articulation)**. Loosely articulated to centrum (0), fused to mid-length of centrum (1).

272. (283) **Haemal arch (length)**. Longer than or equal to neurals (0), shorter than neurals (1).

273. (284) **Haemal arch (shape)**. Non-parallel (triangular) (0), parallel (rectangular) (1).

274. (285) **Spinal nerve foramina**. Absent (0), present (1).

275. (286) **Transverse process**. On arch pedicle (0), on centrum (1).

276. (287) **Atlas-axis intercentra**. Present (0), absent (1).

277. (288) **Atlas (anterior surface of centrum)**. Same size as posterior (0), laterally expanded (1).

278. (289) **Atlas (centrum)**. Multipartite (0), single notochordal (1), single odontoid (2). [*unordered*]

279. (290) **Atlas (neural arch, articulation)**. Loosely articulated (0), sutured (1), fused (2). [*ordered*]

280. (291) **Atlas (parapophyses)**. On centrum (0), on transverse process (1), absent (2). [*unordered*]

281. (292) **Atlas (neural arch)**. Paired (0), sutured at midline (1), fused at midline (2). [*ordered*]

282. (293) **Proatlantes**. Present (0), absent (1).

283. (294) **Second cervical arch**. Expanded to more posterior (0), equal to more posterior (1), shorter than more posterior (2). [*ordered*]

284. (295) **Cervical rib (distal shape)**. Spatulate (0), pointed (1).

285. (296) **Presacral ribs (length)**. Short (0), long (1).

286. (297) **Sacral parapophysis**. On centrum (0), on transverse process (1).

287. (298) **Caudal ribs (count)**. Five or more (0), four (1), three (2), two or fewer (3). [*ordered*]

288. (299) **Scapula (supraglenoid foramen)**. Present (0), absent (1).

289. (300) **Coracoid (number of coracoid foramina)**. zero (0), one (1), two or more (2). [*ordered*]
    a. State 2 was modified to add ‘or more’ based on *Sclerocephalus*.

290. (301) **Scapulocoracoid**. Both (0), scapula only (1), absent (2). [*ordered*]

291. (302) **Humerus (deltopectoral crest)**. Weak (0), intermediate (1), prominent (2). [*ordered*]

292. (303) **Humerus (length)**. Long (>4 trunk centra) (0), short (1).

293. (304) **Radius:humerus**. >=0.7 (0), 0.5-0.69 (1), <0.5 (2). [*ordered*]

294. (305) **Ulna (olecranon process)**. Unossified (0), ossified (1).

295. (306) **Basale commune**. Absent (0), present (1).

296. (307) **Femur (length)**. Long (0), short (1).

297. (308) **Elongate tibiale and fibulare**. Absent (0), present (1).

298. (309) **Distal tarsals (count)**. Six (0), five or fewer (1).

299. (310) **Splenial-coronoids.** Splenial contacts multiple coronoids dorsally (0), splenial contacts only first coronoid dorsally (1).

300. (311) **Parietal (anterior waisting)**. Parietal of equal width throughout (0), parietal tapers gradually anteriorly with obliquely oriented lateral margin (1), parietal with prominent step in lateral margin to form constriction in anterior half (2), parietal with prominent step to form constriction over more than half of the element’s length (3). [*ordered*]
    a. This character was combined with character 360 of Schoch et al. (2020), which refers to whether the parietal is ‘L-shaped.’ Because an ‘L-shaped’ parietal is inherently waisted, the character was modified to differentiate between taxa with only a relatively small anterior waisting (the new state 2) and those with a much greater waisting extending past the mid-length of the parietal (the new state 3). Another state (the new state 1) was also added to differentiate taxa with longitudinally straight margins (no tapering in width) from those that have anteromedially straight margins (tapering but no waisting).

301. (312) **Parietal (width relative to frontal)**. Greater (0), less than or equal to (1).

302. (313) **Skull table**. Rectangular (0), T-shaped (1).

303. (314) **Prefrontals**. Present (0), absent (1).

304. (315) **Postfrontals**. Present (0), absent (1).

305. (316) **Squamosal-frontal**. In contact (0), separated (1).

306. (317) **Zygokrotaphy**. Stegokrotaphy including roofing bones like tabulars and supratemporals (0), zygokrotaphy with muscles approaching skull midline (1), stegokrotaphy in the absence of roofing bones such as tabulars and supratemporals (2), zygokrotaphy, but muscle does not extend through temporal fossa (3). [*unordered*]

307. (318) **Quadrate-maxilla (separation)**. Only separated by pterygoid (0), separated by small pterygoid and pterygoid process of quadrate (1), separated by pterygoid process of quadrate only (pterygoid absent) (2). [*unordered*]

308. (319) **Quadrate-maxillopalatine**. In contact (0), separated (1).

309. (320) **Ceratobranchials**. 3 and 4 fused enclosing larynx (0), not fused (1).

310. (321) **Tentacle**. Absent (0), present adjacent to orbit (1), between orbit and naris (2), beneath naris (3). [*unordered*]

311. (322) **Splenial (teeth)**. Present (0), absent (1).

312. (323) **Choana (size)**. Small (0), large (1).

313. (324) **Sphenethmoid (dorsal exposure)**. Covered dorsally (0), exposed dorsally (1).

314. (325) **Larval stage**. Present (0), absent, direct development (1).

315. (326) **Dentary (anterior teeth, bicuspidity)**. Bicuspid (0), monocuspid (1).

316. (327) **Nasal septum ossification**. Not ossified (0), 20-35% of the total skull length (1), less than 20% of the total skull length (2). [*unordered*]

317. (328) **Sphenethmoid (dorsomedial process, length)**. Absent (0), 11% or more of the total skull length (1), 10% or less of the total skull length (2). [*unordered*]

318. (329) **Sphenethmoid (dorsomedial process, thickness)**. Thin and rod-like (0), thick and broad (1).

319. (330) **Sphenethmoid (dorsal sutural surface)**. Narrow or weakly tapered (0), broad and triangular and strongly tapered (1).

320. (331) **Sola nasi**. Absent (0), present and connected to the nasal septum ventrally (1), present, rod-like or tubular (2). [*unordered*]

321. (332) **Sphenethmoid (lateral walls, length)**. 14% or less than the total skull length (0), 15% or greater of the total skull length (1).

322. (333) **Sphenethmoid (floor)**. Deeply incised and U-shaped in outline (0), shallowly incised with broad coverage of os basale (1), deeply incised and U-shaped, but with midline extension (2). [*unordered*]

323. (335) **Os basale (dorsal exposure)**. Moderate exposure at the dorsal midline (0), little to no exposure near the dorsal midline (1), very wide exposure near the dorsal midline (2). [*unordered*]

324. (336) **Fenestra vestibuli**. Large, subcircular (0), anteroposteriorly elongate, oval (1), very small, subcircular (2). [*unordered*]

325. (337) **Canal for palatal ramus of the facial nerve in antotic region**. Absent (0), present (1).

326. (338) **Stapes (footplate)**. Fills the fenestra vestibule (0), much smaller than the fenestra vestibule (1).

327. (339) **Stapes (columellar process)**. No ridge present (0), ridge on distal columellar process (1).

328. (340) **Optic foramen (shape)**. Large, subcircular opening (0), narrow and slit-like opening (1), oblique anterior margin in the sphenethmoid (2), oblique posterior margin in the os basale (3). [*unordered*]

329. (341) **Sphenethmoid (anterior wall )**. Unossified (0), ossified, one pair of foramina (1), ossified, two pairs of foramina (dorsal and ventral) (2). [*ordered*]

330. (342) **Sphenethmoid (anterolateral process)**. Absent (0), present (1).

331. (343) **Parasphenoid (cultriform process)**. Narrow, tapering anteriorly throughout its length (0), parallel-sided (1); narrowest at mid-length and then flaring anteriorly or continuously flaring anteriorly from the base (2). [*unordered*]

332. (344) **Pseudodentary**. Absent (0), present (1).
    a. The original character (dentary marginal dentition, tooth rows) was completely replaced as it is not possible to treat the second tooth row of gymnophionomorphs as both a double row on the dentary and as coronoid teeth (271–1); this row was treated as coronoid teeth previously and here since the embryological data indicates the lingual row is formed on the coronoid before fusing with the dentary; this treatment also links temnospondyls with a row of teeth on the coronoid(s) to gymnophionomorphs. The substitution of this character provides another supporting character for gymnophionomorph monophyly.

333. (345) **Dorsolateral osteoderms fused to ribs**. Absent (0), present (1).

#### Appendix 6.1. Omitted characters

- (14) **Premaxillary fontanelle**. Premaxillae with or without small opening (0), or encircling large fenestra extending posteriorly between the nasals (1).
    - Merged with character 13.
- (47) **Postorbital (shape).** Long triangular, wedged deeply between squamosal and supratemporal (0), or short (1).
    - Replaced with character 231.
- (48) **Postorbital (end)**. Posterior end acutely triangular (0), or with finger-like projection (1).
    - Merged with the new character 47 (former 231).
- (61) **Intertemporal and postorbital.** Postorbital lateral to postfrontal (0), or expanding medially to replace intertemporal, contacting parietal (1).
    - Replaced with character 232.
- (78) **Dentition (upper jaw).** Conical to slightly curved inwards (0), or caniniform (1).
    - Removed due to ambiguity / possible redundancy with character 38.
- (81) **Dentition (vomer)**. Tooth patches present at least in small specimens (0), or dentition entirely restricted to vomerine fangs (1).
    - Redundant with character 251 (only omitted for reanalysis of Pardo et al., 2017).
- (100) **Choana (medial)**. Medial margin straight or gently convex (0), or anteromedially expanded giving choana a reniform outline (1).
    - Merged with character 101.
- (115) **Cultriform process (dentition)**. Main shaft edentulous except for base (0), or with elongate tooth patch (1).
    - Merged with character 87.
- (253) **Denticles on parasphenoid.** Present (0), absent (1).
    - Redundant with character 87.
- (259) **Pterygoid (lateral process).** Does not contribute to posttemporal fossa (0), contributes to posttemporal fossa (1).
    - Coded feature / philosophy unclear.
- (334) **Parasphenoid (lateral margins)**. Absence of lateral constriction posterior to basicranial articulations (0), weak constriction posterior to basicranial articulations present (1), strong constriction posterior to basicranial articulations, reaching to the carotid foramina (2).
    - Coded feature / philosophy unclear and seemingly inconsistent; character was originally parsimony-uninformative.
