## Supplemental figures for "The disadvantage of derivation: conserved systematic flaws in primary data have repeatedly biased the phylogenetic inference of Temnospondyli (Tetrapoda, Amphibia)"

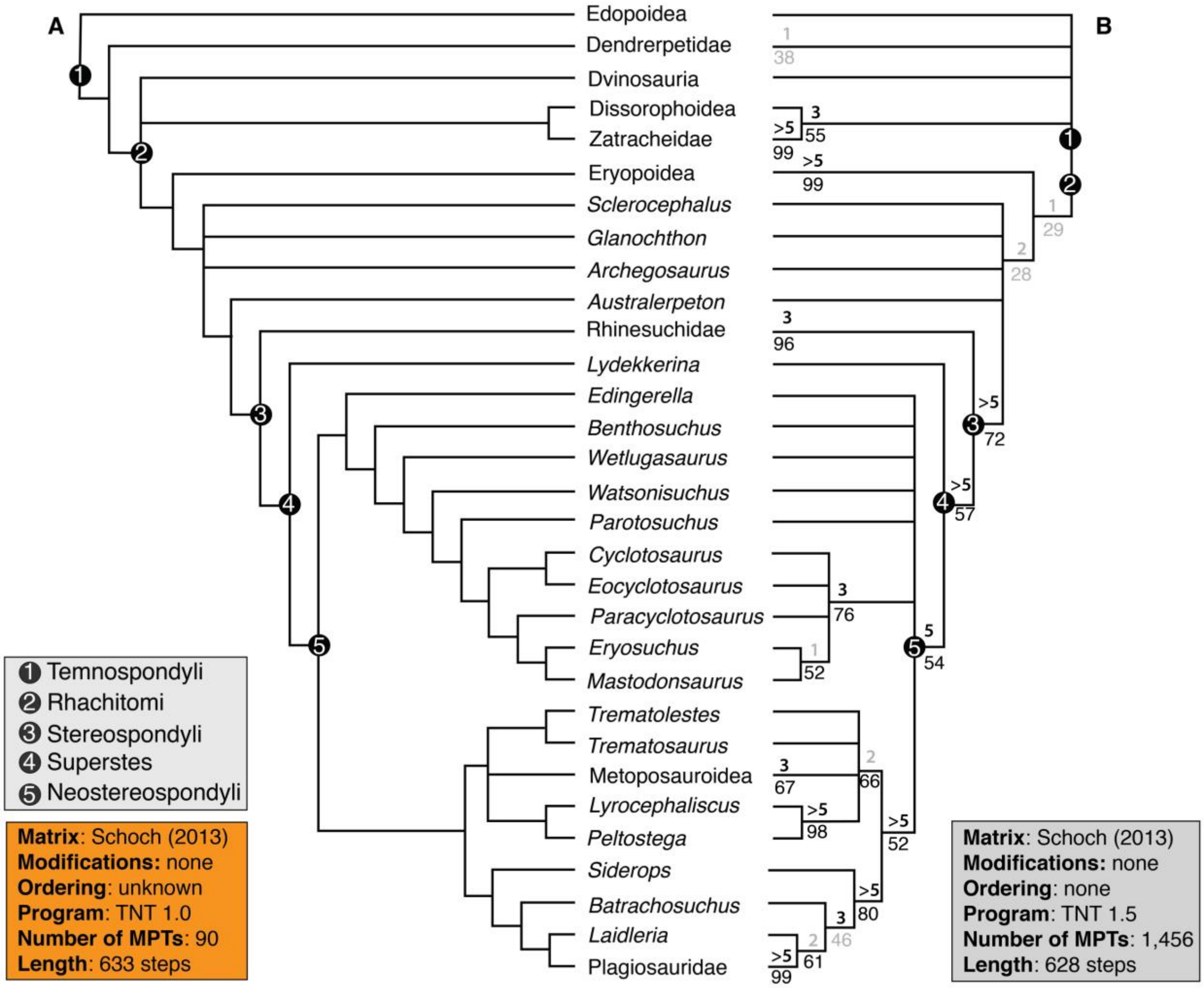


**Supplemental Figure 1. Comparison of the strict consensus topology from Schoch's (2013) analysis using TNT 1.0 (90 MPTs, length = 433 steps) with the strict consensus topology recovered from reanalysis of Schoch's original matrix using TNT 1.5 (1,456 MPTs, length = 428 steps) with all multistate characters unordered and the restricted taxon sample**. **A**, original topology as described but not figured by Schoch (2013:682); **B**, newly recovered topology from the reanalysis. The latter consensus is the same regardless of whether character 67 is ordered or unordered. Values above lines represent Bremer decay index; values below lines represent bootstrap support. Values in gray represent those below the thresholds for strong support (bootstrap ≥ 50%; Bremer index ≥ 3). Support values were not reported for this analysis by Schoch (2013) and are therefore absent from part A. Collapsed nodes representing only two species (Dendrerpetidae [*Balanerpeton*, *Dendrerpeton*], Eryopoidea [*Eryops*, *Onchiodon*], Metoposauroidea [*Callistomordax*, *Metoposaurus*], Plagiosauridae [*Gerrothorax*, *Plagiosuchus*], Rhinesuchidae [*Rhineceps*, *Uranocentrodon*], Zatracheidae [*Acanthostomatops*, *Zatrachys*]) have their support values listed at the tips and are condensed for visual clarity. Relationships of more speciose nodes (Dissorophoidea, Dvinosauria, Edopoidea) are presented in Supplemental Figure 2.

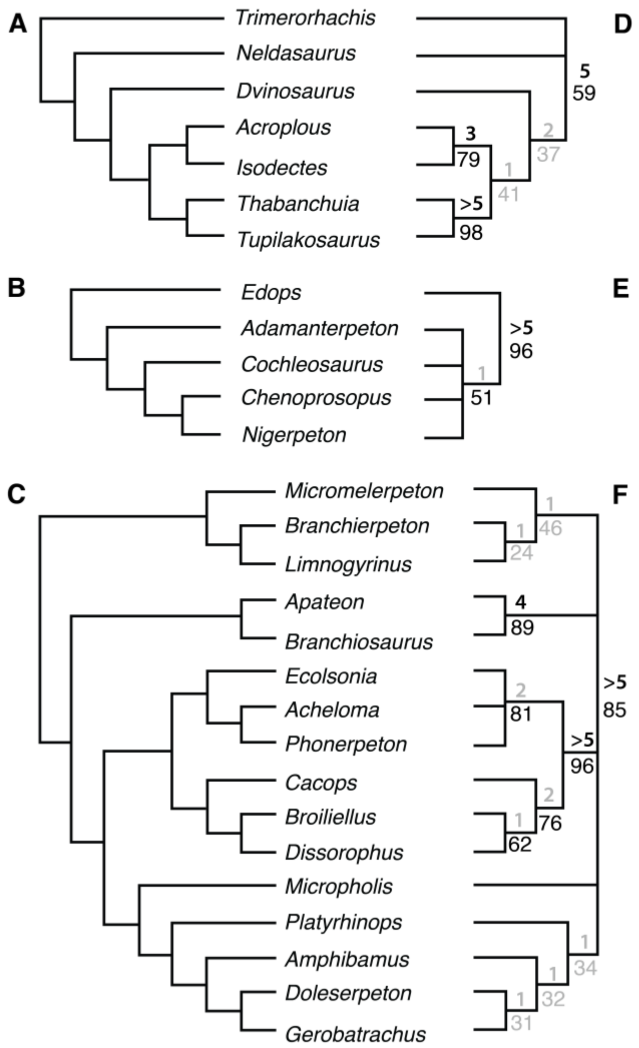


**Supplemental Figure 2. Specific relationships of Dissorophoidea, Dvinosauria, and Edopoidea recovered from reanalysis of the original matrix of Schoch (2013) in TNT 1.0. A**, original topology of Dvinosauria as described but not figured by Schoch (2013:682); **B**, original topology of Edopoidea; **C**, original topology of Dissorophoidea; **D**, newly recovered topology of Dvinosauria; **E**, newly recovered topology of Edopoidea; **F**, newly recovered topology of Dissorophoidea. The consensus of Schoch (2013) is the same regardless of whether character 67 is ordered or unordered. Values in gray represent those below the thresholds for strong support (bootstrap ≥ 50%; Bremer index ≥ 3).

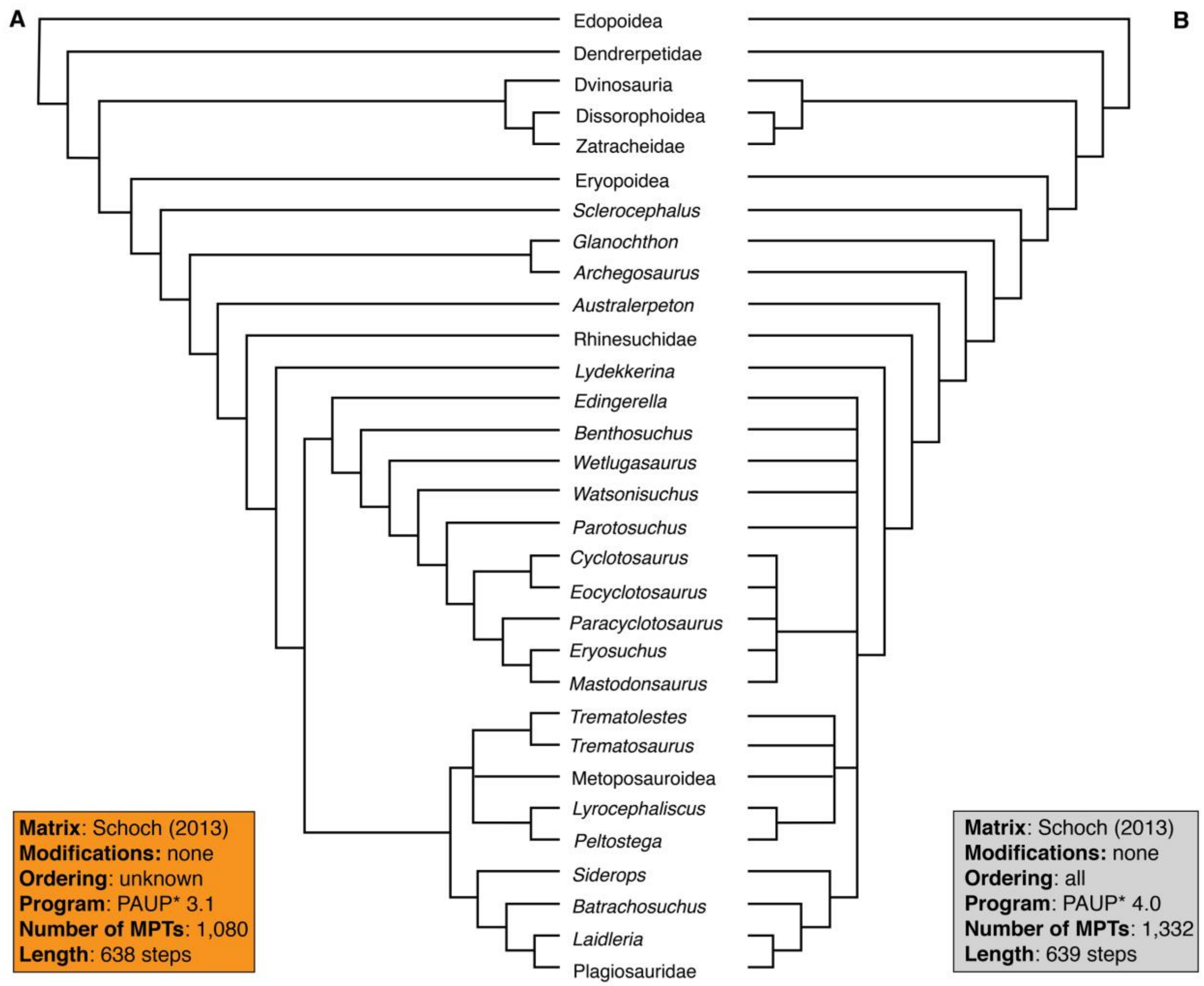


**Supplemental Figure 3. Comparison of the strict consensus topology from Schoch's (2013) analysis using PAUP* 3.1 (1,080 MPTs, length = 638 steps) with the strict consensus topology recovered from reanalysis of Schoch's original matrix using PAUP* 4.0a169 (1,332 MPTs, length = 639 steps) with all multistate characters ordered and the restricted taxon sample**. **A**, original topology as described but not figured by Schoch (2013:682); **B**, newly recovered topology from the reanalysis. Support metrics were not calculated once topological differences were identified. Collapsed nodes representing only two species (Dendrerpetidae [*Balanerpeton*, *Dendrerpeton*], Eryopoidea [*Eryops*, *Onchiodon*], Metoposauroidea [*Callistomordax*, *Metoposaurus*], Plagiosauridae [*Gerrothorax*, *Plagiosuchus*], Rhinesuchidae [*Rhineceps*, *Uranocentrodon*], Zatracheidae [*Acanthostomatops*, *Zatrachys*]) have their support values listed at the tips and are condensed for visual clarity. Relationships of more speciose nodes (Dissorophoidea, Dvinosauria, Edopoidea) are presented in Supplemental Figure 4.

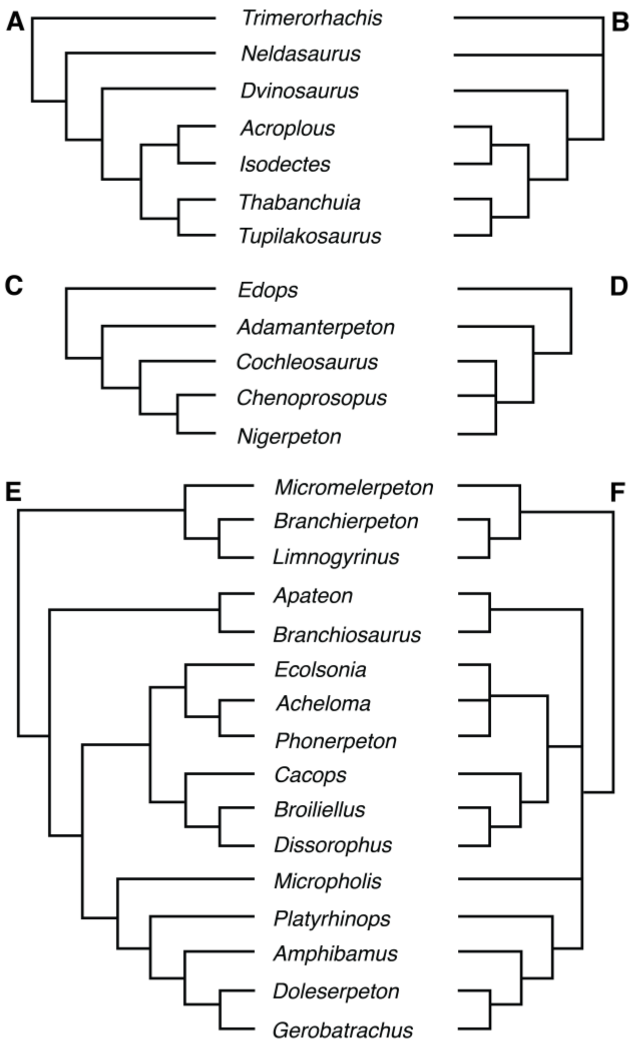


**Supplemental Figure 4. Specific relationships of Dissorophoidea, Dvinosauria, and Edopoidea recovered from reanalysis of the original matrix of Schoch (2013) in PAUP* 4.0a169. A**, original topology as described but not figured by Schoch (2013:682); **B**, original topology of Edopoidea; **C**, original topology of Dissorophoidea; **D**, newly recovered topology of Dvinosauria; **E**, newly recovered topology of Edopoidea; **F**, newly recovered topology of Dissorophoidea. The consensus of Schoch (2013) is the same regardless of whether character 67 is ordered or unordered. Values in gray represent those below the thresholds for strong support (bootstrap ≥ 50%; Bremer index ≥ 3).

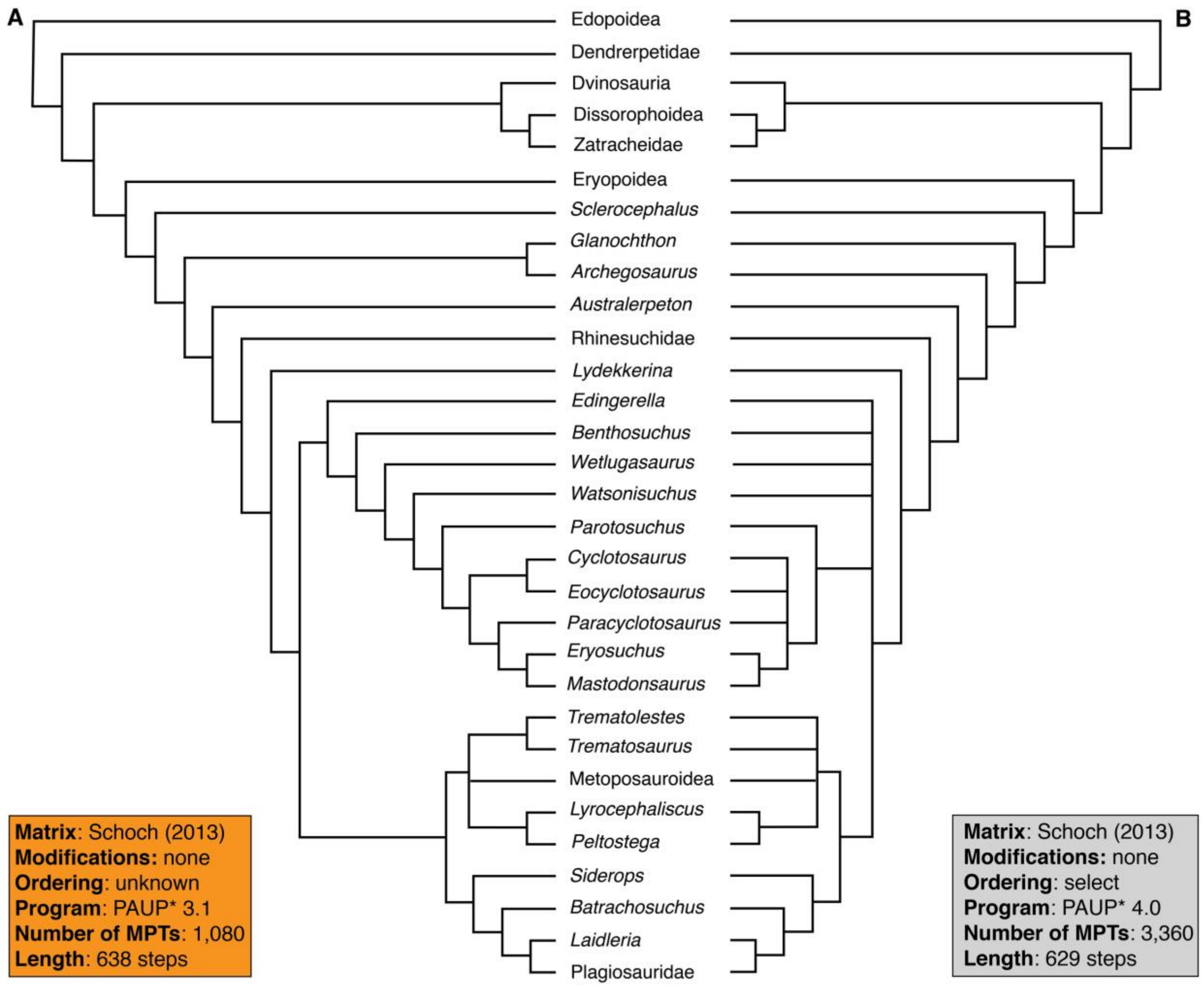


**Supplemental Figure 5. Comparison of the strict consensus topology from Schoch's (2013) analysis using PAUP* 3.1 (1,080 MPTs, length = 638 steps) with the strict consensus topology recovered from reanalysis of Schoch's original matrix using PAUP* 4.0a169 (3,360 MPTs, length = 629 steps) with select multistate characters ordered (67, 75, 110, 143, 145, 158, 163, 170, 182, 187, and 191) and the restricted taxon sample**. **A**, original topology as described but not figured by Schoch (2013:682); **B**, newly recovered topology from the reanalysis. The specific relationships of Dissorophoidea, Dvinosauria, and Edopoidea are as with Supplemental Figure 4 and are not repeated here. Support metrics were not calculated once topological differences were identified. Collapsed nodes representing only two species (Dendrerpetidae [*Balanerpeton*, *Dendrerpeton*], Eryopoidea [*Eryops*, *Onchiodon*], Metoposauroidea [*Callistomordax*, *Metoposaurus*], Plagiosauridae [*Gerrothorax*, *Plagiosuchus*], Rhinesuchidae [*Rhineceps*, *Uranocentrodon*], Zatracheidae [*Acanthostomatops*, *Zatrachys*]) have their support values listed at the tips and are condensed for visual clarity. Relationships of more speciose nodes (Dissorophoidea, Dvinosauria, Edopoidea) are presented in Supplemental Figure 6.

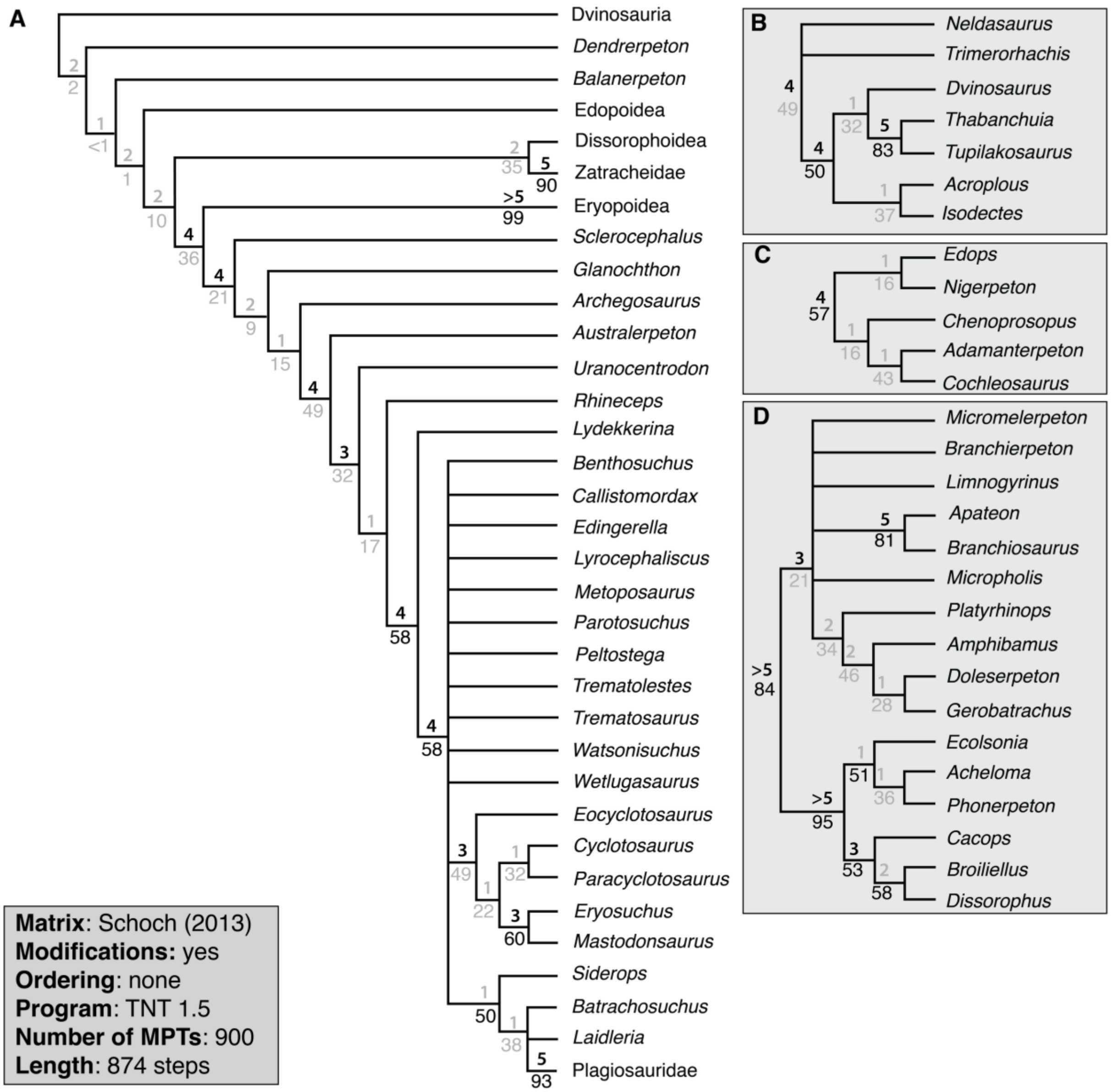


**Supplemental Figure 6. Strict consensus topology recovered from analysis of the modified Schoch (2013) matrix using TNT, with the restricted taxon sample and all multistate characters unordered.** **A**, topology of Temnospondyli, with select clades collapsed for visual clarity; **B**, relationships of Dvinosauria; **C**, relationships of Edopoidea; **D**, relationships of Dissorophoidea. Values above lines represent Bremer decay index; values below lines represent bootstrap support. Values in gray represent those below the thresholds for strong support (bootstrap ≥ 50%; Bremer index ≥ 3). Collapsed nodes representing only two species (Dendrerpetidae [*Balanerpeton*, *Dendrerpeton*], Eryopoidea [*Eryops*, *Onchiodon*], Metoposauroidea [*Callistomordax*, *Metoposaurus*], Plagiosauridae [*Gerrothorax*, *Plagiosuchus*], Rhinesuchidae [*Rhineceps*, *Uranocentrodon*], Zatracheidae [*Acanthostomatops*, *Zatrachys*]) have their support values listed at the tips and are condensed for visual clarity.

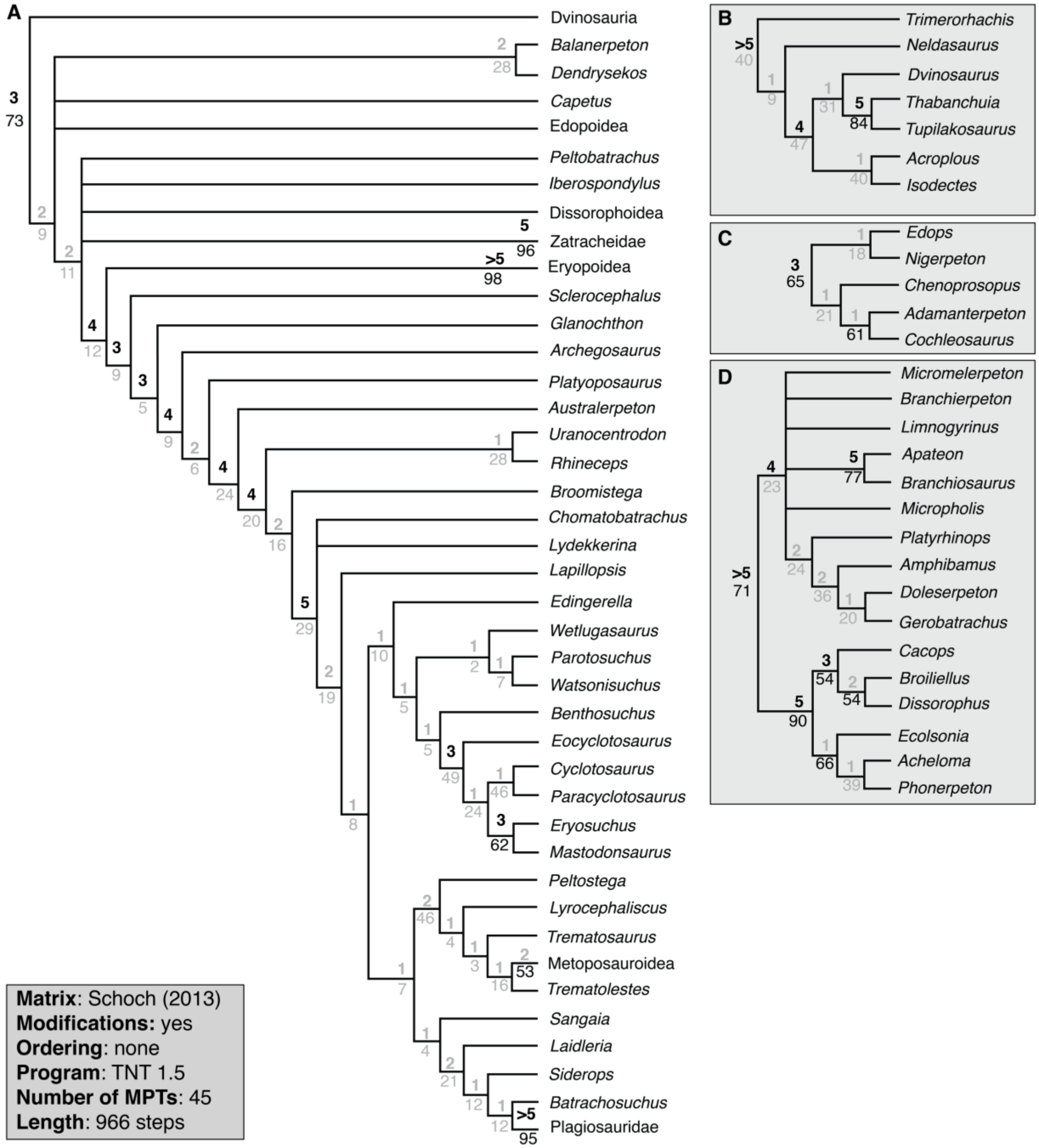


**Supplemental Figure 7. Strict consensus topology recovered from analysis of the modified Schoch (2013) matrix using TNT, with the full taxon sample and all multistate characters unordered.** **A**, topology of Temnospondyli, with select clades collapsed for visual clarity; **B**, relationships of Dvinosauria; **C**, relationships of Edopoidea; **D**, relationships of Dissorophoidea. Values above lines represent Bremer decay index; values below lines represent bootstrap support. Values in gray represent those below the thresholds for strong support (bootstrap ≥ 50%; Bremer index ≥ 3). Collapsed nodes representing only two species (Dendrerpetidae [*Balanerpeton*, *Dendrerpeton*], Eryopoidea [*Eryops*, *Onchiodon*], Metoposauroidea [*Callistomordax*, *Metoposaurus*], Plagiosauridae [*Gerrothorax*, *Plagiosuchus*], Rhinesuchidae

[*Rhineceps*, *Uranocentrodon*], Zatracheidae [*Acanthostomatops*, *Zatrachys*]) have their support values listed at the tips and are condensed for visual clarity.

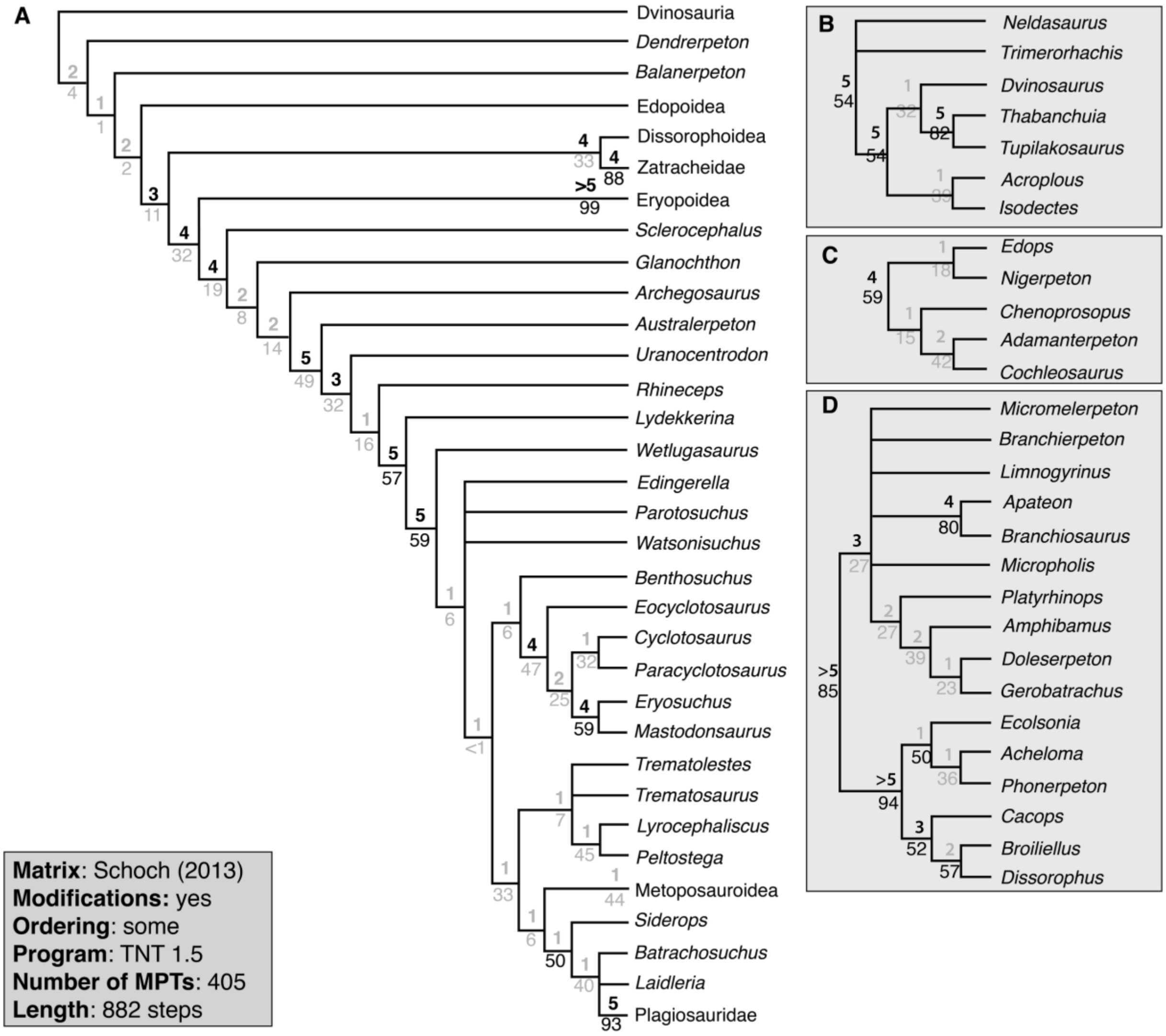


**Supplemental Figure 8. Strict consensus topology recovered from analysis of the modified Schoch (2013) matrix using TNT, with the restricted taxon sample and certain multistate characters ordered.** **A**, topology of Temnospondyli, with select clades collapsed for visual clarity; **B**, relationships of Dvinosauria; **C**, relationships of Edopoidea; **D**, relationships of Dissorophoidea. Values above lines represent Bremer decay index; values below lines represent bootstrap support. Values in gray represent those below the thresholds for strong support (bootstrap ≥ 50%; Bremer index ≥ 3). Collapsed nodes representing only two species (Dendrerpetidae [*Balanerpeton*, *Dendrerpeton*], Eryopoidea [*Eryops*, *Onchiodon*], Metoposauroidea [*Callistomordax*, *Metoposaurus*], Plagiosauridae [*Gerrothorax*, *Plagiosuchus*], Rhinesuchidae [*Rhineceps*, *Uranocentrodon*], Zatracheidae [*Acanthostomatops*, *Zatrachys*]) have their support values listed at the tips and are condensed for visual clarity.

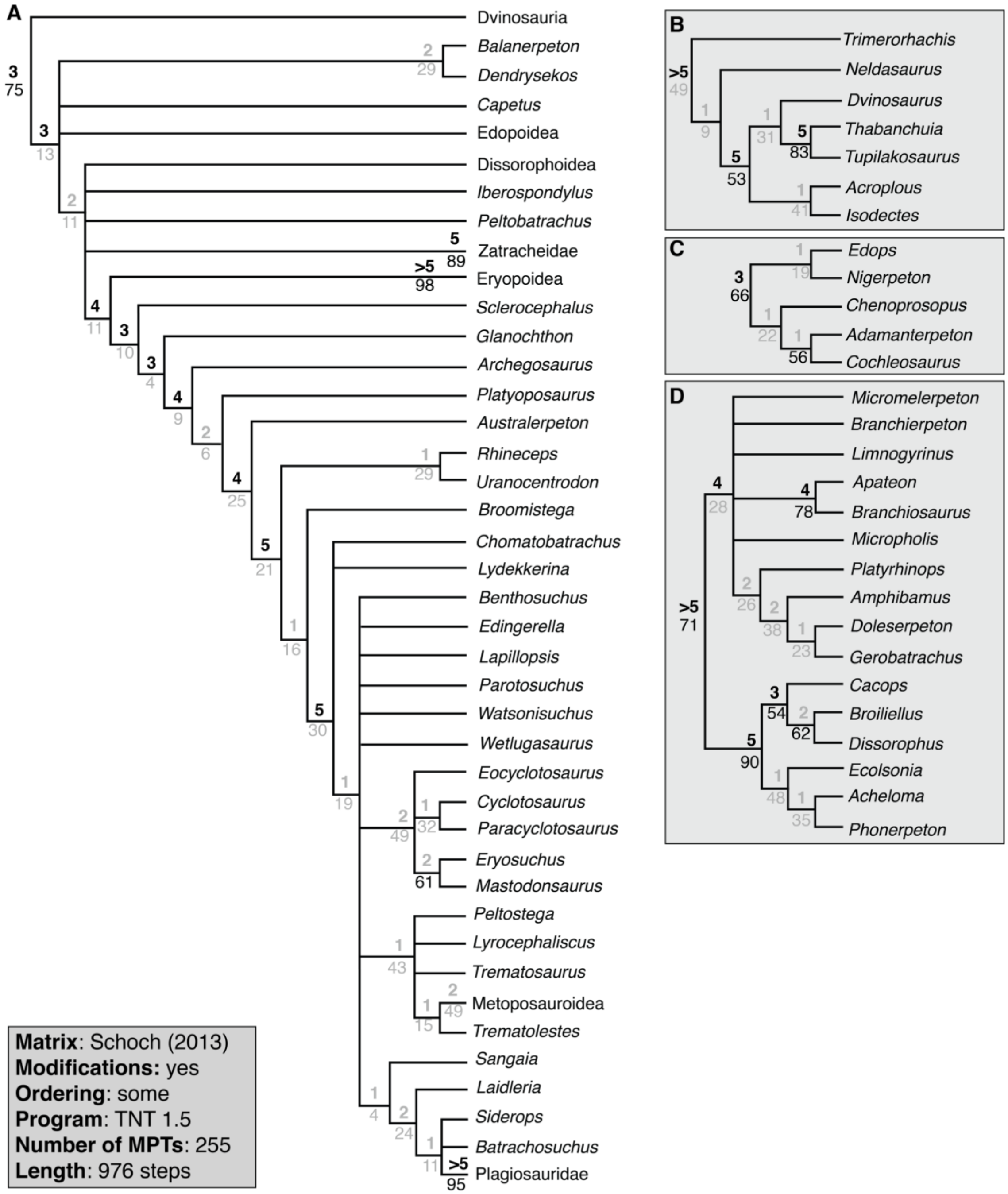


**Supplemental Figure 9. Strict consensus topology recovered from analysis of the modified Schoch (2013) matrix using TNT, with the full taxon sample and certain multistate characters ordered.** **A**, topology of Temnospondyli, with select clades collapsed for visual clarity; **B**, relationships of Dvinosauria; **C**, relationships of Edopoidea; **D**, relationships of Dissorophoidea. Values above lines represent Bremer decay index; values below lines represent bootstrap support. Values in gray represent those below the thresholds for strong support (bootstrap ≥ 50%; Bremer index ≥ 3). Collapsed nodes representing only two species (Dendrerpetidae [*Balanerpeton*, *Dendrerpeton*], Eryopoidea [*Eryops*, *Onchiodon*], Metoposauroidea [*Callistomordax*, *Metoposaurus*], Plagiosauridae [*Gerrothorax*, *Plagiosuchus*],

Rhinesuchidae [*Rhineceps*, *Uranocentrodon*], Zatracheidae [*Acanthostomatops*, *Zatrachys*]) have their support values listed at the tips and are condensed for visual clarity.
